# Partnership with both fungi and bacteria can protect *Odontotermes obesus* fungus gardens against fungal invaders

**DOI:** 10.1101/2023.08.25.554860

**Authors:** Renuka Agarwal, Manisha Gupta, Ruchira Sen, Nimisha E.S., Rhitoban Raychoudhury

## Abstract

Fungus-growing termites, like *Odontotermes obesus*, cultivate *Termitomyces* as their sole food source on fungus combs which are continuously maintained with foraged plant materials. This necessary augmentation also increases the threat of introducing pathogenic fungi capable of displacing *Termitomyces*. The magnitude of this threat and how termites prevent pathogens remain largely unknown. This study identifies this pathogenic load by establishing the pan-mycobiota of *O. obesus* from the fungus comb and termite castes. Furthermore, to maximize the identification of such pathogenic fungi, the mycobiota of the decaying stages of the unattended fungus comb were also assessed. The simultaneous assessment of the microbiota and the mycobiota of these stages identified possible interactions between the fungal and bacterial members of this community. Based on these, we propose a possible interaction among the crop fungus *Termitomyces*, the weedy fungus *Pseudoxylaria* and some bacterial mutualists. These possibilities were then tested with *in vitro* interaction assays which suggest that *Termitomyces*, *Pseudoxylaria* and bacterial mutualists all possess anti-fungal capabilities. We propose a multifactorial interaction model of these microbes, under the care of the termites, to explain how their interactions can maintain a predominantly *Termitomyces* monoculture.

## Introduction

Termites primarily utilize plant-derived lignocellulosic materials as sources of nutrition. Lignocellulose is a recalcitrant material that only some Basidiomycotan fungi can digest completely^1^. Termites have evolved symbiotic associations with different groups of organisms (bacteria, protists and fungi) to achieve lignocellulose digestion^2^. In all lower and some higher termites, it is achieved directly through nutritional symbiosis with gut-dwelling microbial symbionts^3^. However, the termites of the subfamily Macrotermitinae have evolved an indirect way of utilizing the lignocellulose-degrading capabilities of the Basidiomycotan fungus, *Termitomyces*. This fungus subsists on plant materials, brought into the termite mound by workers, and thrives on a spongy lignocellulosic structure called the fungus comb^4,5^. *Termitomyces* utilize the lignocellulose to grow^6^, whereas termites use the growing fungal nodules as food^7^. This association, which first evolved in tropical Africa around 35 MYA^5,8,9^, has become so successful that fungus cultivation in termites is considered a canonical example of the evolution of agriculture in animal societies^10^. As this crop fungus has become the sole source of nutrition for these termites, it has led to the evolution of many unique behavioral phenotypes associated with their successful cultivation^11^.

The behavioral phenotypes include but are not limited to (1) the construction of the specialized fungus-growing beds (combs) and the earthen mounds^10,12^ to protect them; (2) the use of semi-digested plant material^13^, feces and soil^14,15^ to build the combs and in the process seeding the combs with the inoculum of *Termitomyces*^13^, (3) regulation of temperature^16^, relative humidity^16,17^ and CO2 levels^17^ and (4) maintenance of fungus gardens to ensure ideal growing conditions for *Termitomyces*.

However, these ideal growing conditions for *Termitomyces* can also be conducive to the growth of other non-desirable fungi, similar to the invasion of weeds in human agricultural fields. Additionally, the continuous and unidirectional flow of dead and decaying plant materials into these mounds increases the threat of introducing pathogenic and parasitic fungi being introduced is also high. These decaying plant materials can harbor a diverse array of saprophytic communities of microfungi, yeasts, and bacteria^18,19^. Indeed, several such weedy fungi have been isolated from different fungus-growing termites^15,20–22^. Many of these can outcompete *Termitomyces* and result in a severe reduction of available nutrients for the termites. The termite-fungus symbiosis, therefore, should be under strong selection pressure to evolve mechanisms that can prevent the growth of such weedy fungi. This selection pressure can act on the termites directly, enabling them to constantly seek out weedy fungi for elimination through digestion^23^. However, recent reports^20^ indicate that such a strategy of selective de-weeding does not reduce pathogenic fungal load as these spores survive the passage through the termite gut. The near absence of any other active fungi in healthy fungus combs, which are dominated by *Termitomyces*^24^, points to mechanisms that actually prevent these fungi from proliferating within the combs. This could be due to the direct or indirect use of the resident microbial community of the fungus comb by the termites in preventing the growth of any weedy fungi^25–28^. Some indirect evidence exists for this hypothesis where the microbiota of many different fungus-growing insects shows compositional similarity^29^. This is a remarkable example of convergent evolution as most of these symbiotic microbial communities are dominated by the bacteria *Pseudomonas*^30–32^ which have also been shown to inhibit weedy fungi^33^. This can be evidence for selection^25^ to include bacterial mutualists within the resident microbiota, which can hinder the growth of weedy fungi. Accordingly, several in vitro studies have found many such resident bacteria acting as secondary mutualists capable of preventing the most dominant weedy fungi of these fungus gardens, *Pseudoxylaria*^34–36^. Several such bacterial secondary symbionts have since been identified, including *Bacillus*^26,36^, *Streptomyces*^37,38^, *Burkholderia*^33^ and *Pseudomonas*^33^ which can act as anti-fungal agents. Another evidence for this hypothesis comes from the ability of *Termitomyces* themselves to hamper the growth of some fungal contaminants when cultured on plates that previously grew *Termitomyces*^24,39^. However, what remains unknown is how and to what extent can *Termitomyces* act against these non-specific fungi.

The current research on how fungus-growing termites raise a weed-free crop has primarily been restricted to mechanisms by which *Pseudoxylaria* is controlled. However, these studies lack a comprehensive assessment of the threat from other weedy fungi. Many sequencing approaches have revealed variations in the bacterial communities from these fungus combs^33,40,41^, but since there are no reports of a core-mycobiota from any fungus-growing termites, it is difficult to know the roles of other fungi in this symbiosis and the relationship between the microbiota and the mycobiota. In this study, we try to answer these questions by using the widespread fungus-growing termite from India, *Odontotermes obesus*. First, to assess the threat from the fungal contaminants, we identify the pan-mycobiota of this symbiosis by amplifying the internal transcribed spacer (ITS) gene fragment and generating amplicons on the Nanopore platform of the fungal comb and termite castes. We augment this estimation by including samples from the decaying stages of the fungus comb after removal of the termites to identify which weedy fungi remain suppressed in an active comb but become visible with the progressive decay. Second, to identify any interactions between the myco-and microbiota of these combs, we characterize the community-wide changes in the abundance of microbes from these decaying combs by using nanopore platform and confirmed the results using qPCR for three specific microbes proven to be important in this association^33^. Third, as these temporal changes of the decaying community of microbes highlight the possibility of a few important interactions, we empirically validate these by isolating different fungi from the colonies and testing the nature of their interactions. Fourth, we test whether putative bacterial mutualists, identified in a previous study using these same termite mounds^33^, also show anti-fungal properties against these weedy fungi. Finally, we propose a model to explain the role of these interactions in the disease-free growth of *Termitomyces* in the fungus gardens of *O*. *obesus*.

## Results

### O. obesus colonies harbor a vast array of fungal contaminants

The pan-mycobiota of *O. obesus* established from the combs (fresh and decaying), alates, workers and nymphs yielded over 3.9 × 10^5^ high-quality Nanopore reads (Table 1), with the highest number of reads from female alates (∼1.06 × 10^5^ reads) and the lowest from the 120 hrs old decaying comb (∼4.0 ×10^3^ reads). The rarefaction curves (Supplementary Fig. S7 a) showed an exhaustive sampling of major workers and female alates, indicating sufficient coverage. The number of identified fungal OTU’s ranged from 197 (in major workers) to 51 (in minor workers), with 418 unique fungal genera across *O. obesus*. Figure 1 indicates the presence of the fungal OTU’s having a relative abundance of greater than 0.05% in any one of the eleven samples. The termite-free incubation of the comb yielded 236 different fungal contaminants other than *Pseudoxylaria*. These include 81 fungal contaminants alone from the 24 hrs old comb, which reduced to 69 after 120 hrs of incubation. The highest fungal diversity was found from 72 hrs old comb which showed 110 different fungal contaminants. This indicates that many of these fungal contaminants are actively suppressed in the combs for *Termitomyces* to flourish. The detailed taxonomic classification and abundances of these OTU’s are tabulated in supplementary table S8. Community similarities, using UniFrac distances in a weighted PCoA plot, indicated greater similarity among the mycobiota of the decaying combs and between both alates and major workers (Supplementary Fig. S7 c).

**Fig. 1:**
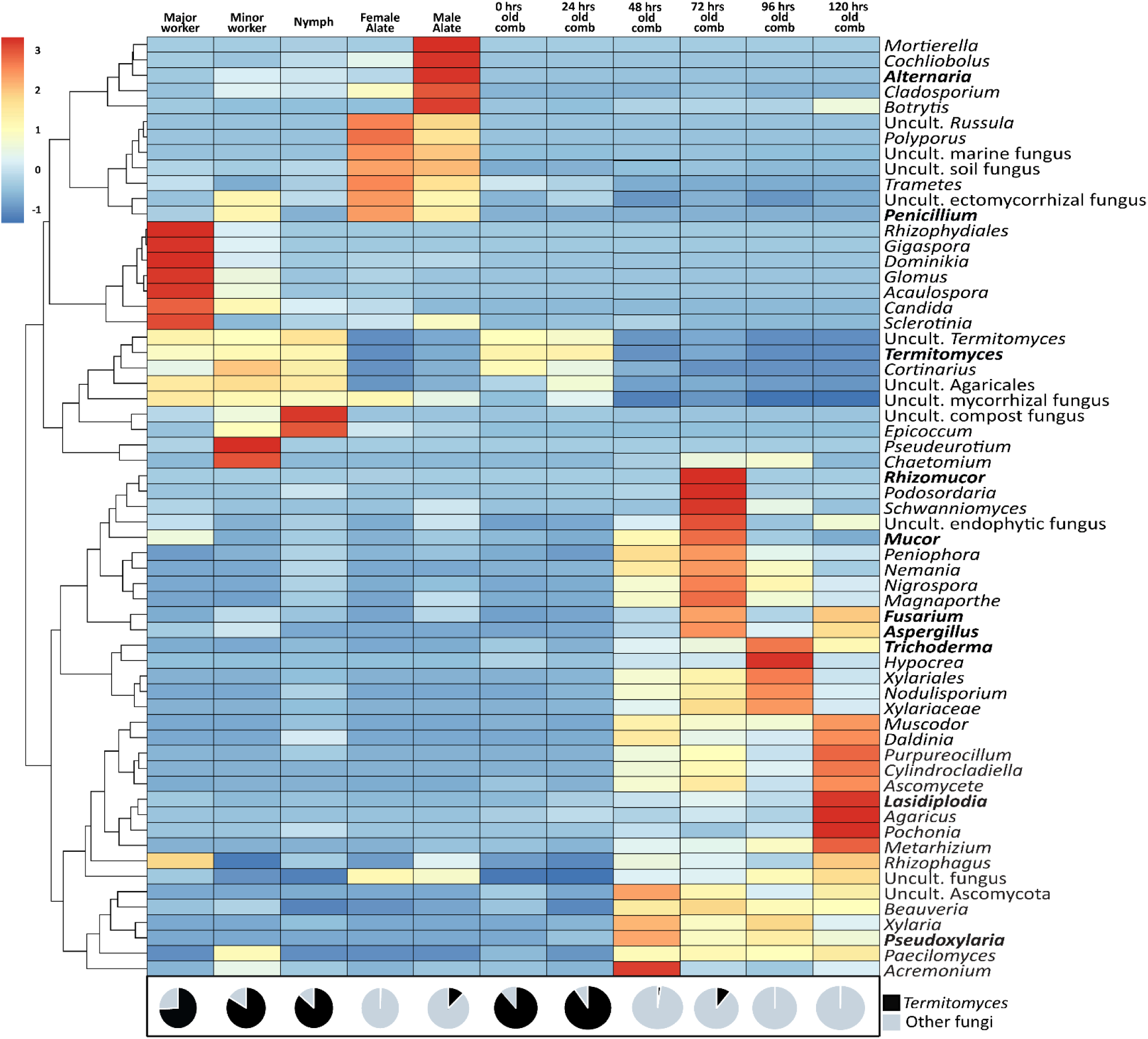
Mycobiota of *O. obesus*. Fungal OTU’s that accounted for > 0.05% abundance in any of the six samples are shown in the heatmap. The numbers below each pie chart indicates the percentage of *Termitomyces* OTU’s obtained. The genera mentioned in blue were also obtained in culture.

**Table 1:** Number of reads obtained to identify the mycobiota of *O*. *obesus*. The table also shows different diversity indices for all the samples. The details of the taxonomic distribution, as well as abundance, for all the identified reads are given in table S8.

| Sample source | Total reads (initial) | Total reads (post-cleanup) | Reads identified | Fungal genera identified | Chao1 | Shannon | Simpson | InvSimpson |
| --- | --- | --- | --- | --- | --- | --- | --- | --- |
| Major Worker | 0.92 x 10<sup>5</sup> | 0.76 x 10<sup>5</sup> | 0.76 x 10<sup>5</sup> | 197 | 170.0 | 0.32 | 0.89 | 1.09 |
| Minor Worker | 0.04 x 10<sup>5</sup> | 0.04 x 10<sup>5</sup> | 0.04 x 10<sup>5</sup> | 51 | 40.7 | 0.71 | 0.30 | 1.43 |
| Nymph | 0.22 x 10<sup>5</sup> | 0.19 x 10<sup>5</sup> | 0.17 x 10<sup>5</sup> | 87 | 110.4 | 0.50 | 0.21 | 1.27 |
| Female Alate | 1.15 x 10<sup>5</sup> | 1.06 x 10<sup>5</sup> | 1.06 x 10<sup>5</sup> | 185 | 284.3 | 0.85 | 0.52 | 2.09 |
| Male Alate | 0.19 x 10<sup>5</sup> | 0.14 x 10<sup>5</sup> | 0.12 x 10<sup>5</sup> | 98 | 126.0 | 1.28 | 0.64 | 2.82 |
| Decaying comb (0 hrs) | 0.58x10<sup>5</sup> | 0.54x10<sup>5</sup> | 0.54x10<sup>5</sup> | 111 | 199.3 | 0.53 | 0.20 | 1.25 |
| Decaying comb (24 hrs) | 0.62x10<sup>5</sup> | 0.60x10<sup>5</sup> | 0.58x10<sup>5</sup> | 83 | 116 | 0.47 | 0.17 | 1.20 |
| Decaying comb (48 hrs) | 0.16x10<sup>5</sup> | 0.18x10<sup>5</sup> | 0.14x10<sup>5</sup> | 103 | 157.4 | 1.67 | 0.75 | 4.04 |
| Decaying comb (72 hrs) | 0.16x10<sup>5</sup> | 0.17x10<sup>5</sup> | 0.14x10<sup>5</sup> | 110 | 152.5 | 1.97 | 0.80 | 5.24 |
| Decaying comb (96 hrs) | 0.17x10<sup>5</sup> | 0.18x10<sup>5</sup> | 0.15x10<sup>5</sup> | 99 | 184 | 1.54 | 0.70 | 3.43 |
| Decaying comb (120 hrs) | 0.04x10<sup>5</sup> | 0.04x10<sup>5</sup> | 0.04x10<sup>5</sup> | 71 | 126.1 | 1.47 | 0.63 | 2.73 |
|  |  |  |  | 418 unique fungal genera |  |  |  |  |

### The dynamics of the micro-and mycobiome in the decaying fungus comb

To determine how the micro-and mycobiota of fungus combs change after the removal of termites, we incubated fresh fungus combs for 120 hrs and enumerated their temporal changes every 24 hrs. Over 1.59 x 10^5^ fungal OTU’s were identified across 120 hrs of incubation. The highest number of reads were obtained from the 24 hrs old comb (0.58 × 10^5^) and the least from 120 hrs old comb (0.04 × 10^5^). The mycobiota survey confirmed (Fig. 1, Supplementary Tables S5 and S8) previous observations of a fresh comb being dominated by *Termitomyces*^24,42^ as 89% of the OTU’s were identified as *Termitomyces*. However, with each passing day, these decaying combs revealed the proliferation of various new fungi. The relative abundance of *Termitomyces* was highest in the fresh fungus combs (90.7%), which drastically decreased after 24 hrs (2.1%) and was almost undetectable by 72 hrs (Fig. 2 and Supplementary Fig. S11). This was accompanied by a corresponding increase in *Pseudoxylaria* OTU’s (Fig. 2 and Supplementary Fig. S11) which increased from 2.6 % by 24 hrs and reached the maximum in 48 hrs (26.1%) but then also experienced a reduction by 120 hrs (12.2%). The temporal changes in the abundances of these two fungi were also accompanied by a steady increase of the non-specific fungal OTU which jumped from ∼10% in 24 hrs old comb to 99.9% by 96 hrs of incubation. Thus, the increase in abundance of these contaminant fungi seems to be negatively correlated with *Termitomyces* and *Pseudoxylaria* (Supplementary Fig. S11).

**Fig. 2:**
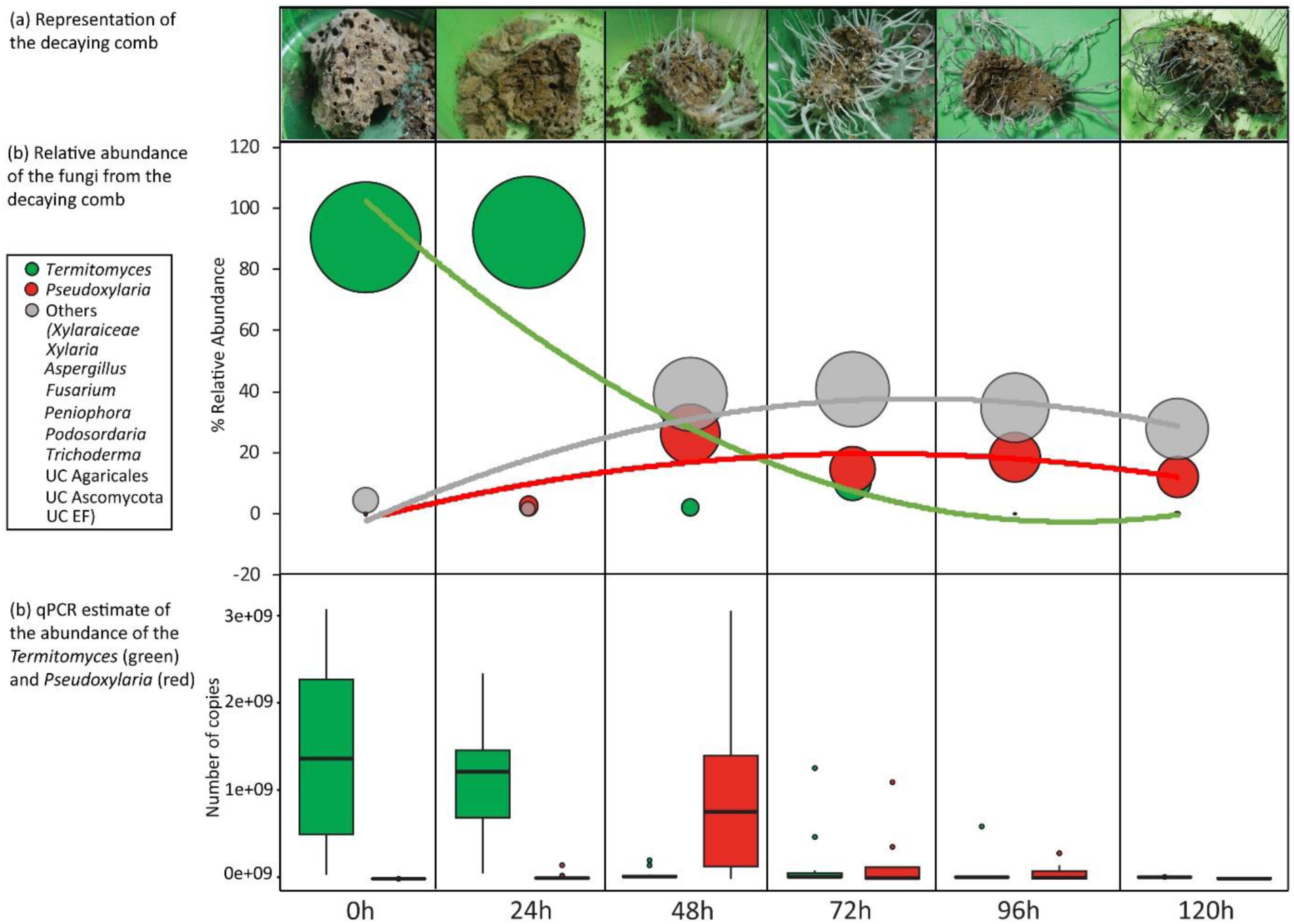
Change in the physical appearance of the decaying fungus comb and the relative abundances and copy numbers of *Termitomyces* and *Pseudoxylaria* in decaying fungus comb as determined by Nanopore Sequencing (bubble plot) and qPCR (box plot). Fungal OTU’s that had relative abundance of atleast >0.3% in any of the six samples are shown in the bubble plot.

### qPCR estimation confirmed the assessment through nanopore platform

Since, qPCR is a more sensitive estimator of actual abundance^43^, we confirmed the OTU data with qPCR estimation of copy number variations of the cultivar *Termitomyces* and the most prominent weed *Pseudoxylaria* using the same DNA source used to generate the OTU’s. The 95% confidence intervals of these copy numbers were used to assess the changes (Supplementary Table S6) and indicate a similar pattern of dynamics between the OTU and qPCR data for both these fungi (Fig. 2 and Supplementary Fig. S11). As expected, the OTU data revealed a lower threshold of the number of reads in comparison to qPCR estimates.

The corresponding microbiota of this decay was estimated from over 1.16 × 10^5^ high-quality Nanopore reads (Supplementary Table S7) across these same six samples, with the highest number of identified reads from 48 hrs old comb (∼0.36 × 10^5^ reads) and the lowest from the fresh comb (∼0.12 × 10^5^ reads). The rarefaction curves (Supplementary Fig. S8 a) indicated relatively exhaustive coverage of bacterial species diversity except for 48 hrs old comb. A weighted PCoA analysis of the Unifrac distances among these samples showed a limited clustering (Supplementary Fig. S8 c), indicating differential temporal abundances of some of these bacterial genera (Supplementary Fig. S9). Significantly, the magnitude of the changes in the bacterial community was far less severe than the mycobiome. This indicates relatively more resilient bacterial communities than fungal ones in the combs. Subsequently, to determine the correlation between the potential bacterial mutualists that have inhibitory effects against *Pseudoxylaria*^33^, the abundances of *Pseudoxylaria* and the bacterial mutualists were analyzed by comparing their number of OTU’s. This comparison also revealed a negative association (Supplementary Figs. S10 a, b). This shows the possibility that mutualistic bacteria may play a defensive role against the weedy fungus which results in healthy combs having low incidence of *Pseudoxylaria*.

To test any interaction between the bacteria and the fungi, we selected *Pseudomonas*, a secondary mutualist identified from these same mounds^33^. The number of copies of *Pseudomonas* remained largely unchanged till 48 hrs, then increased after 72 hrs and came down after 120 hrs (Supplementary Fig. S10 b). One-way Anova conducted with *Termitomyces*, *Pseudoxylaria* and *Pseudomonas* showed significant variation between the groups (p= 0.00081, F= 7.512, df= 2). Post hoc Tukey tests revealed insignificant pairwise site differences in the copy numbers of *Termitomyces* and *Pseudoxylaria* (p= 0.102) and *Pseudoxylaria* and *Pseudomonas* (p= 0.364).

Thus, these complex patterns of the abundances of the three major microbes, *Termitomyces*, *Pseudoxylaria* and the bacterial mutualists, indicate that they may have a synergistic role in preventing non-specific fungi.

### Members of both the micro-and mycobiota can stop non-specific fungi

The temporal dynamics of the micro-and mycobiota, detailed above, highlights a crucial role for both *Termitomyces* and *Pseudoxylaria*. It appears that the abundances of these two fungi are negatively correlated with the presence of other contaminant fungi (Supplementary Fig. S11). Therefore, we tested whether these two fungi can indeed inhibit the growth of some of these non-specific fungal strains in culture. We began by looking at the capability of *Termitomyces* against *Pseudoxylaria* and found that *Pseudoxylaria* easily grew over *Termitomyces* (Supplementary Fig. S12), indicating that the crop fungus cannot prevent the most prominent weedy fungus. However, as figure 3 indicates, *Termitomyces* can inhibit fourteen other fungi in direct interaction assays (Supplementary Fig. S13). *Termitomyces* showed maximum inhibition (more than 50%) against *Alternaria*-MN913751, *Curvularia*-MN913753, *Rhizomucor*-MN913756, *Syncephalastrum*-MN913763 and *Ustilago*-MN913772 (Fig. 3, Supplementary Fig. S13 and Table 2) and limited inhibition (less than 50%) against nine other fungi, *Aspergillus*-MN913749, *Aspergillus*-MN913750, *Diaporthe*-MN913762*, Fusarium*-MN913754, *Lasiodiplodia*-MN913758, *Paraconiothyrium*-MN913759, *Penicillium*-MN913771, *Phialotubus*-MN913760 and *Phoma*-MN913761 (Fig. 3, Supplementary Fig. S13 and Table 2). However, the interaction with *Rhizomucor*-MN913756 and *Diaporthe*-MN913762, was not conclusive as the patterns of growth were too ambiguous to clearly differentiate these two fungi from *Termitomyces*. The two fungi, *Mucor*-MN913757 and *Trichoderma*-MN913767 overgrew *Termitomyces*, showing no sign of inhibition. However, these results indicate that *Termitomyces* possess the capability of preventing many fungal contaminants to a variable degree.

**Fig. 3:**
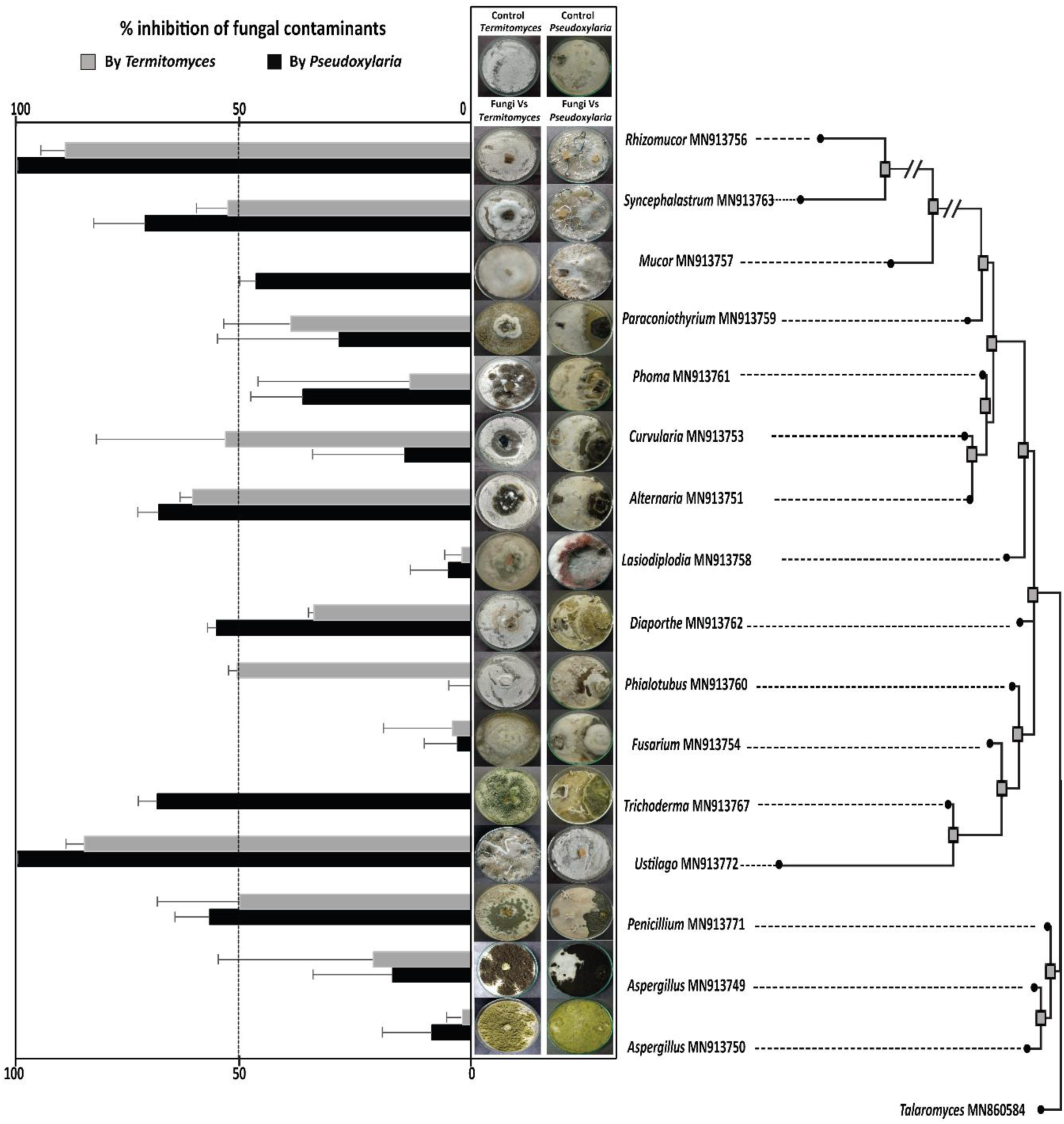
A comparison of the % inhibition (left panel) of fungal contaminants by *Termitomyces* and *Pseudoxylaria* obtained from *O. obesus* colonies. The right panel shows representative pictures of *Termitomyces* and *Pseudoxylaria* interactions with the fungal contaminants with the top panel showing the growths in control plates. The phylogenetic analysis of ITS gene was run on MEGAX with the K2+g substitution model using *Talaromyces* as the outgroup. represents > 75 bootstrap value.

**Table 2:**
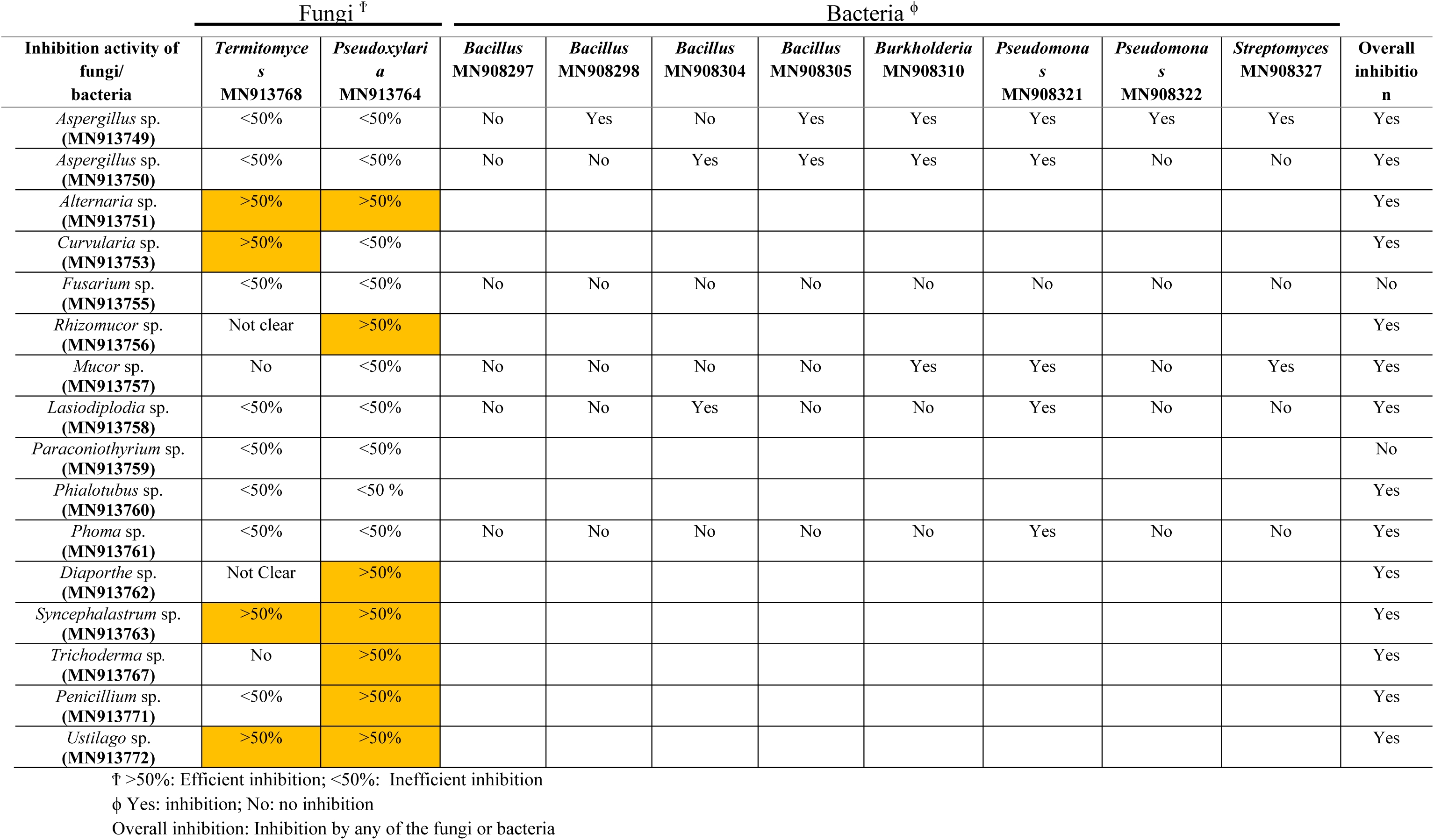
Inhibition efficiency of *Termitomyces*, *Pseudoxylaria* and bacterial mutualists on fungal contaminants.

*Pseudoxylaria* showed variable degrees of inhibition against all of tested contaminant fungi (Fig. 3 and Supplementary Fig. S14). The maximum degree of inhibition (more than 50%) was shown against *Alternaria-*MN913751, *Diaporthe*-MN913762, *Penicillium*-MN913771, *Rhizomucor-* MN913756, *Syncephalastrum*-MN913763, *Trichoderma*-MN913767, and *Ustilago*-MN913772 (Fig. 3 and Supplementary Fig. S14) and limited inhibition (less than 50%) was shown against *Aspergillus*-MN91375049, *Aspergillus-*MN913750, *Curvularia*-MN913753, *Fusarium*-MN913754, *Lasiodiplodia*-MN913758, *Mucor*-MN913757, *Paraconiothyrium*-MN913759, *Phialotubus*-MN913760 and *Phoma*-MN913761 (Fig. 3 and Supplementary Fig. S14). Interaction of *Pseudoxylaria* against *Alternaria-*MN913751 and *Syncephalastrum*-MN913763 showed vertical growth of both these interacting fungi. However, the two fungi could be easily identified and their zones of growth evaluated (Fig. 3 and Supplementary Fig. S14).

These interaction assays reveal that both *Termitomyces* and *Pseudoxylaria* mirror each other in their inhibitory effects (Table 2; Supplementary Fig. S15). As figure 3 indicates, certain contaminant fungi are inhibited by both. The two exceptions are *Mucor*-MN913757 and *Phialotubus*-MN913760 where the major inhibition is shown only by *Pseudoxylaria* and *Termitomyces*, respectively. However, *Aspergillus*-MN913749, *Aspergillus*-MN913750, *Fusarium*-MN913754, *Lasiodiplodia*-MN913758, *Mucor*-MN913757 and *Phoma*-MN913761 were neither inhibited by *Termitomyces* nor by *Pseudoxylaria* but to a degree less than 50% and hence these fungi were then assayed with the bacterial mutualists.

The eight bacterial mutualists, from four different genera^33^, showed inhibitory effects against all but one (*Fusarium*-MN913755) of these fungi (Fig. 4, Table 2). As figure 4 indicates, out of the eight bacterial cultures, six showed a significant capability to prevent the growth of these fungi (Table 2). These interaction assays suggest that termites possess multifaceted capability of preventing the growth of contaminant fungi through *Termitomyces*, *Pseudoxylaria* and bacterial mutualists.

**Fig. 4:**
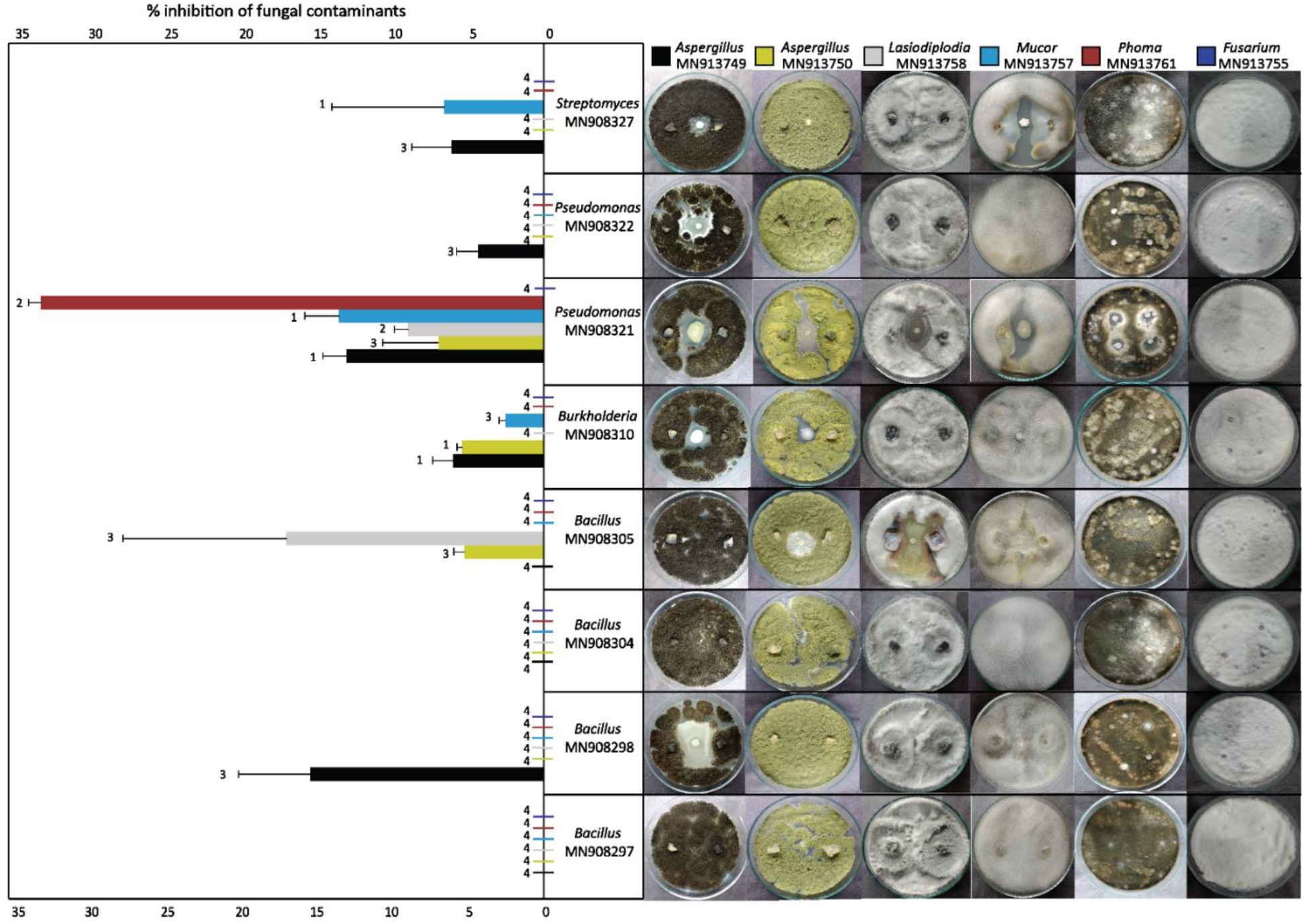
% inhibition (left panel) of the fungal contaminants by the 8 bacterial mutualists (right panel) obtained from *O. obesus* colonies. Whisker over horizontal bars represent standard deviation and number indicates the type of inhibition shown by the bacterial strains (1= clear zone of inhibition, 2= reduced growth near bacteria, 3= contact inhibition, 4= negligible inhibition).

## Discussion

The fungus gardens of *O. obesus* appear to be a monoculture, but the pan-mycobiota of *O. obesus*-*Termitomyces* symbiosis indicates that the threat from many potential contaminants remains suppressed in a fresh comb. A total of 418 different fungal strains were successfully annotated from the nearly 2.69 × 10^5^ OTU’s generated from the Nanopore platform (Table 1), with 236 unique genera alone from the fungus comb. This remains a conservative estimate as rarefaction curves of the fungal OTU’s from many samples (Supplementary Fig. S7 a) did not reach saturation. These results are contrary to Bos et al^20^ and Otani et al^24^, which reported significantly fewer non-specific fungi with relatively low abundances of non-*Termitomyces* genera from fungus combs which ranged from <0.001% (454 sequencing datasets) to 0.07% relative abundances (MiSeq sequencing datasets) of non-specific fungi^20,24^. Only one of the four samples from Bos et al^20^ identified 75% of the reads as *Termitomyces* and 24% as *Xylaria*. These discrepancies could be the result of methodological differences or could be characteristic of *O. obesus* colonies. Our initial assumption of workers bringing in potentially contaminating fungi was also supported as the major workers revealed the presence of 197 different fungal OTU’s. The abundance of such contaminants in the alates, with only 0.6-12% relative abundance of *Termitomyces* (Fig. 1), also suggests vertical transmission of these contaminants. How these fungi are suppressed by alates during nest founding and establishing *Termitomyces* monocultures merits future studies^44^. Our results suggest that the ideal growing conditions for *Termitomyces* can also harbor a vast array of contaminant and weedy fungi and identifies the magnitude of the ‘fungal threat’ that this symbiosis needs to overcome. Consequently, these results also highlight the efficiency of termites suppressing the growth of non-cultivar fungi for the successful proliferation of *Termitomyces*.

Termites are reported to select the best candidates among the different strains of *Termitomyces* available for cultivation and propagation^8,45^. Natural selection should favour both choosing (termites) and chosen (cultivar strain) partner if the cultivar is adept at preventing the proliferation of weedy fungi. As figure 3 indicates, *Termitomyces* can indeed inhibit several non-specific fungi obtained from these same mounds, but crucially, not the most prominent weed, *Pseudoxylaria* (Supplementary Fig. S12). However, as Supplementary figure S11 illustrates, the presence of both *Termitomyces* and *Pseudoxylaria* is correlated with the contaminating fungi being in check for at least the first 48 hrs. This can be crucial evidence for the synergistic effects of both *Termitomyces* and *Pseudoxylaria* in preventing other non-specific fungi. However, this necessitates the demonstration of the anti-fungal effects of *Pseudoxylaria*^46^. Therefore, we tested *Pseudoxylaria* for anti-fungal properties and as figure 3 indicates, *Pseudoxylaria* shows some inhibitory effects against almost all the fungi tested. Moreover, the eight bacterial mutualists previously identified^33^ also show similar growth inhibitory capabilities. Our results point to the presence of microbes, both bacterial and fungal, with fungicidal capabilities present in this symbiosis. Given the magnitude of the ‘fungal threat’ to this monoculture, this seems unsurprising. However, what remains unknown is what role, if any, selection has played in bringing about the composition of this community and how precisely these microbes bring about a successful monoculture.

Our results confirm that the presence of termites is indispensable for a functioning monoculture, as their removal initiates the proliferation of contaminant fungi. This indicates that the termites need constant weeding to keep any such non-specific fungi at bay. Previous studies^23^ reported that *O*. *obesus* workers bury any visible *Pseudoxylaria* when given portions of cultured hyphae, indicating that they can recognize this as a contaminant to be removed, possibly through olfactory cues^47^. However, the qPCR data indicated that *Pseudoxylaria* is present in appreciable numbers even in a fresh comb (Supplementary Fig. S11) and is obviously prevented from proliferating. This can be indicative of a dynamic equilibrium where the growth of *Pseudoxylaria* is tolerated by the termites to some extent, perhaps to prevent other non-specific fungi, but is also simultaneously inhibited from taking over the comb. This can be achieved by constant weeding out of *Pseudoxylaria* outbreaks within a functioning comb. However, how termites perform such a selective de-weeding remains unknown. As figure 2 indicates, *Pseudoxylaria* starts proliferating after 24 hrs and the termites probably have less than 48 hrs to remove it as by that time, *Termitomyces* experiences a drastic reduction (Supplementary Fig. S4). This decline progresses steadily with the corresponding increase of non-specific fungi. Thus, if termites weed out *Pseudoxylaria* by 48 hrs, then the comb can still perhaps be used to grow *Termitomyces,* but beyond 72 hrs, *Termitomyces* is all but wiped out by the proliferation of both *Pseudoxylaria* and other contaminating fungi (Fig. 2 and Supplementary Fig. S11). *Pseudoxylaria*, with its faster growth and constitutive fungicidal activity (Fig. 3, Supplementary Fig. S12, and Table 2), can be a more efficient inhibitor of non-specific fungi than *Termitomyces*. This hints towards its potential beneficial role in this symbiosis, but such a contention remains controversial. *Pseudoxylaria* has been considered to be an opportunistic fungus that can take over crop monoculture when conditions are suitable for it^48^. As termites cannot feed on any other fungus^49^, the invasion of *Pseudoxylaria* is indicative of the collapse of this symbiosis^50^. However, if the anti-fungal capabilities of *Pseudoxylaria* and *Termitomyces*, as seen in culture assays (Fig. 3), are a reliable indicator of their capabilities within a fungus comb, then it is clear that *Termitomyces* is a less efficient inhibitor of non-specific fungi than *Pseudoxylaria*. This inefficiency can be a selection pressure for the accommodation of *Pseudoxylaria* into this symbiosis. First, the presence of *Pseudoxylaria* in the core-mycobiota of *O. obesus* (Fig. 1 and Supplementary Table S8) argues against its role as an incidental infection^48^. Moreover, its presence in the alates (Fig. 1) also hints at successful transmission across generations. The extensive taxonomic review of this group^51,52^ puts *Pseudoxylaria* to be exclusively associated with Macrotermitinae termites. Therefore, *Pseudoxylaria* is either an active member of this symbiosis or is an extremely specialized weed or parasite. Second, Fricke et al^53^ reported that the genome architecture of *Pseudoxylaria* is consistent with a symbiotic lifestyle as they reveal significantly reduced genome sizes with a loss of lignin metabolizing capabilities. Therefore, this could be indicative of a mutualistic role for *Pseudoxylaria* involved in preventing other fungi while simultaneously being dependent on *Termitomyces* for nutrition through lignocellulose digestion. Third, Visser et al^48^ reported that both *Pseudoxylaria* and *Termitomyces* can use the same Carbon sources to grow, making them strong competitors with the potential of one eliminating the other within a fungus comb. However, as Supplementary figure S12 show, *Termitomyces*, which has inhibitory effects against other fungi, cannot prevent *Pseudoxylaria* growth and is easily overtaken by it. This inability of *Termitomyces* to prevent *Pseudoxylaria* is contrary to the expected outcome of the evolution of some mutual inhibitory capabilities, especially if they are competing for the same resources. However, there are several fungal (Fig. 3) as well as bacterial strains (Fig. 4)^33^ that can do the same and yet a functioning comb is still found to be dominated by *Termitomyces* (Fig. 2). Fourth, a mutualistic role of *Pseudoxylaria* predicts its presence in appreciable amounts even in a fresh comb. As figure 2 indicates, the OTU data barely supports this as only 51 reads identified as *Pseudoxylaria* from the fresh fungus comb. However, qPCR estimates, which are a better estimator of abundance, indicate the presence of 6.0×10^5^ copies (95% confidence intervals; Supplementary Table S6). When taken together, these two estimates prove the presence of *Pseudoxylaria* in fresh combs and are contrary to previous estimates^24^. The control of *Pseudoxylaria* can perhaps be enforced by secondary mutualistic bacteria in these colonies, as several such bacterial strains have been identified from these same mounds^33^. This suggests a complex interdependent relationship between *Termitomyces*, *Pseudoxylaria*, and the mutualistic bacteria. However, how such precise control of these various microbes is achieved remains to be investigated. The broad outlines of these arguments are detailed in figure 5 as a hypothetical model. This model posits that *Termitomyces* monocultures face a constant threat from incoming microbes, possibly brought by foragers into the mound (Fig. 5). Since termites construct an ideal growth chamber to grow *Termitomyces* monocultures, the presence of these fungal contaminants can be detrimental as they can easily parasitize the same resources. However, these fungal contaminants are prevented by *Termitomyces* (Fig. 3 and Supplementary Fig. S13), *Pseudoxylaria* (Fig. 3 and Supplementary Fig. S14), as well as by some bacterial mutualists (Fig. 4 and Supplementary Figs. S16-S23) from proliferating. The second prediction of this model assumes the presence of bacterial mutualists in the fresh comb, possibly for the prevention of both *Pseudoxylaria* and other contaminating fungi. Supplementary table S9 indicates the presence of 991 OTU’s from the four genera of bacterial mutualists in fresh comb samples^33^. As a representative of these mutualists, we undertook qPCR estimation of *Pseudomonas* and found 3.0×10^5^ copies (95% confidence intervals; Supplementary Table S6) in the same fresh comb samples. Therefore, the presence of these bacteria indicates that the control of *Pseudoxylaria* can perhaps be enforced by secondary mutualistic bacteria in these colonies with additional control by termites through weeding^23^. This model, however, awaits empirical validation, and future studies with more mounds and different species of termites are needed to establish whether this model is validated or the role of *Pseudoxylaria* remains that of a weed and/or parasite.

**Fig. 5:**
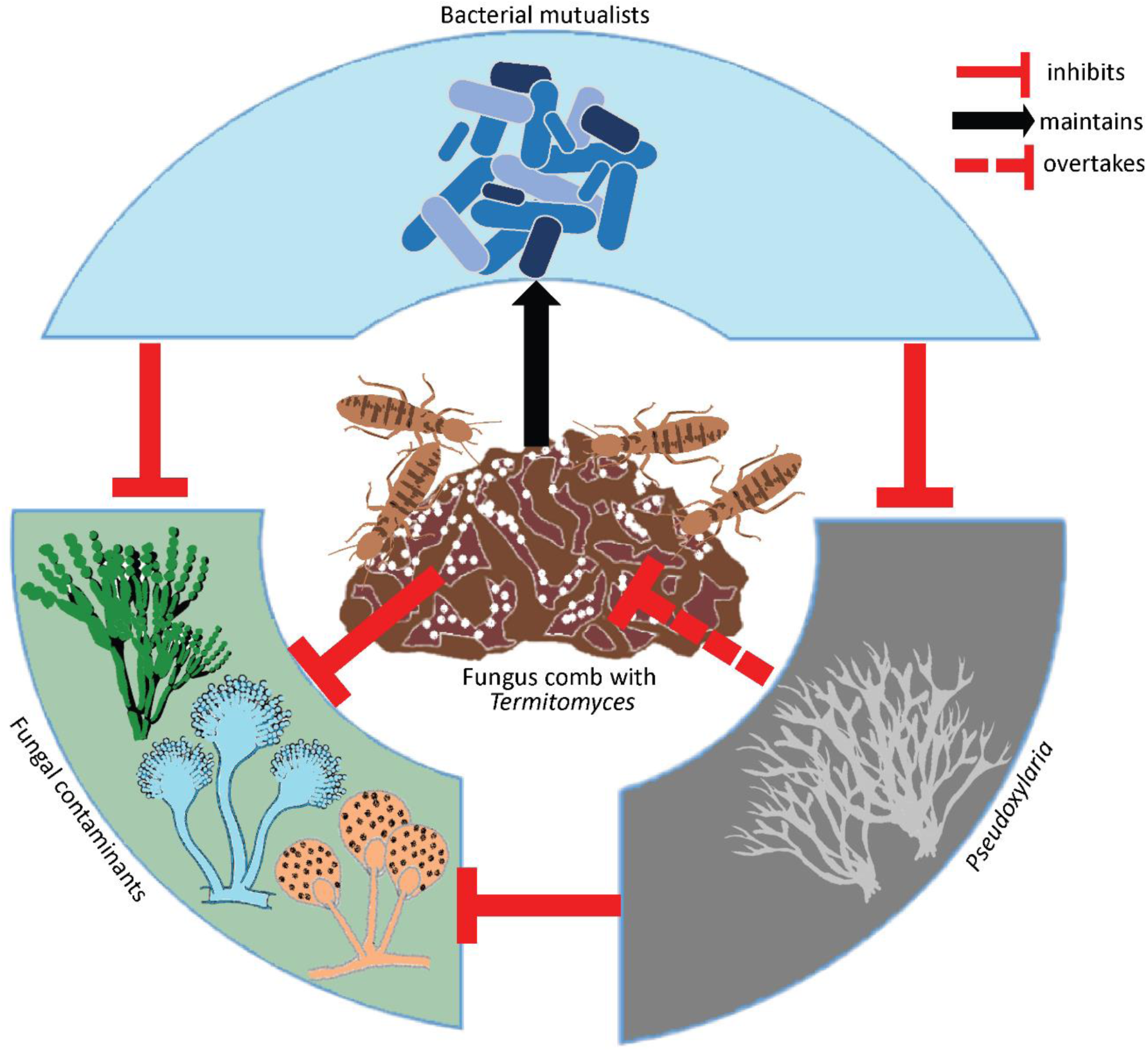
Verbal model for understanding interactions among microbes, fungi and termites in fungus-growing termites.

## Methods

### Collection of termites, fungus combs and their DNA extraction

Different castes (major workers, minor workers, male and female alates and nymphs) and fungus combs of *O. obesus* were collected from two different mounds within the IISER Mohali campus These are the same mounds from which the core-microbiota was obtained in a previous study^33^. (Supplementary Figs. S1 and S2). Male and female alates were collected from these mounds during swarming, which began after the first rains in June 2018, while other castes (major workers, minor workers and nymphs) were collected along with the fungus combs (Supplementary Fig. S3). The details of the collection and DNA extraction are detailed in supplementary note 1 and 2.

Fresh fungus combs were collected across three months (August-October, 2018) from the same two mounds as mentioned above and were set up to decay for 120 hrs. The collected combs were immediately removed of their resident termites, kept in sterile plastic containers, then crushed with autoclaved pestles, divided into roughly 0.5 g portions, and incubated at 30 °C inside sterilized glass containers which were then screwed shut to prevent any possible contamination (Supplementary Fig. S4). For each incubation assay, three such containers were set up from each mound with 12 such incubation assays in all. DNA was extracted from these combs consecutively for six days, i.e., 0 hrs to 120 hrs. For 0 hrs of incubation, DNA was extracted from each portion separately within 3 hours of the collection. DNA samples were amplified in replicates and then the PCR products from the same day were pooled in equimolar concentrations (20-25 ng µl^-1^) into a single sample. Six such pooled PCR products, one from each day (0 hrs – 120 hrs), were prepared for microbiota and mycobiota analysis on the Nanopore platform (Supplementary Table S3).

### Isolation and identification of fungal strains

*Termitomyces* was directly isolated from fresh nodules growing on the fungus comb, successively washed with 50% ethanol and sterilized water, inoculated on plates containing Potato Dextrose Agar (PDA) with Yeast Malt Extract (PYME)^22^ and incubated at 30 °C. *Pseudoxylaria* hyphae were obtained from a 48-72 hrs old post-collection fungus comb and directly cultured on Potato Dextrose Agar (PDA) plates. To isolate other fungi, comb fragments and termite castes were washed with sterilized water, homogenized in 500 µl of sterile 1X PBS buffer (pH -7.4) and a dilution series (10x-10^-6^x) of these homogenates were plated on PDA plates containing 250 μg/mL chloramphenicol (Himedia), and incubated at 30 °C for 4-7 days. Fungal cultures obtained were initially screened visually for unique growth characteristics and identified by sequencing the Internal Transcribed Spacer (ITS) gene fragment, which was amplified with the primer set ITS4*/*ITS5^54^ (Supplementary Note 3). Any fungal cultures with more than one Single Nucleotide Polymorphism (SNP) were considered unique and used in the study (Supplementary Table S1). The sequences of these fungi have been submitted to GenBank (accession number MN913749 – MN913772). Since multiple strains within some fungal genera were obtained, the nomenclature of these strains was modified to read as ‘Genus-NCBI Accession Number’ (Supplementary Table S1).

### Running the Nanopore platform and obtaining Fungal and Bacterial Operational taxonomic Units (OTU’s)

The mycobiota was obtained by amplifying the ITS fragment with the primer set ITS5/ITS4^54^, while the V3-V4 region of the 16S rRNA gene was amplified to characterize the microbiota using the primers 341F/806R^55^. DNA from different castes and six different comb samples of varying degrees of incubation were used to obtain the pan-mycobiota (Supplementary Tables S2-S3). These six comb samples were also amplified to characterize their microbiota (Supplementary Table S3).

The sequencing library was run on FLO-MIN106 flowcell for 48 hrs using MinKNOW software with the protocol *NC_48Hr_sequencing_FLO-MIN106_SQK-LSK108_plus_Basecaller* (Supplementary Note 4). All the cleaned sequences were deposited to NCBI Sequence Read Archive (SRA). The five mycobiota were deposited under the BioProject PRJNA665655 (SAMN16262892 - SAMN16262897), the microbiota of decaying comb samples were deposited under BioProject PRJNA758817 (SAMN21036816 – SAMN21036821) and the mycobiota of decaying comb as PRJNA758827 (SAMN21037110 – SAMN21037115), respectively.

### Taxonomic identification of OTU’s

The fungal OTU’s were identified by comparing them against the fungus repository from the NCBI FTP site with an additional modification. Initial annotation revealed several ‘*Pseudoxylaria*-like’ sequences, which were not identified till the genus level. Therefore, to refine the annotation we used the previous study by Hsieh et al^52^ to phylogenetically characterize these sequences from NCBI to see whether these belong to *Pseudoxylaria* or not. This was done by amplifying the α-actin gene using the primer set ACT-512F/ACT-783R^52^ and comparing it with the other members of the Xylariaceae family. The α-actin sequences (accession number OQ503187 and OQ503188) from the *Pseudoxylaria* isolated from the two mounds under study were initially aligned with Clustal Omega ^56^ and manually edited in Bioedit v 7.0.5.3^57^. Phylogenetic analysis of these sequences was performed in MrBayes v 3.2.7^58^. After confirming that the two sequences belonged to the *Pseudoxylaria* clade, we aligned them with sequences from NCBI annotated as *Xylaria*, Xylariaceae, and Xylariales and performed another phylogenetic analysis. Sequences that formed a monophyletic clade with these two previously identified *Pseudoxylaria* sequences were renamed as *Pseudoxylaria* (Supplementary Figs. S5-S6 and Supplementary Table S8) and updated in the reference database used for annotation. Annotation was done using LAST v 973, with the following parameters: match score of 1, gap opening penalty of 1, and gap extension penalty of 1^59^. The identified reads were sorted from the phylum to the genus level and their relative abundances were calculated. To estimate the sequencing depth, rarefaction curves were generated using the Vegan package v 2.5-4 in R v 3.6.2. The estimated community richness (Chao1) and diversity indices (Evenness, Shannon, Simpson, and Inv Simpson) were also calculated in R v 3.6.2. To test how representative these communities are across the different fungus-growing termites, a comparison was done with the data obtained in this study with previously published data from Bos et al^20^ and Otani et al^24^.

The bacterial sequences generated from the six comb samples (0 hrs-120 hrs) were identified by comparing them against the customized microbial repository made by combining 16S rRNA gene fragments from NCBI FTP site and the DictDb v 3.0 database^60^. The OTU’s for the potential four bacterial mutualists were filtered, combined and compared with the absolute abundances of *Termitomyces* and *Pseudoxylaria*.

### Quantitative PCR for Termitomyces, Pseudoxylaria and Pseudomonas in fungus combs

The decaying comb samples (0-120 hrs) were also used to quantify *Termitomyces, Pseudoxylaria* and *Pseudomonas* copy numbers. The specificity of the designed primers was checked with Primer-BLAST (last performed on July 2021). The sequence of these primers and their respective annealing temperatures are given in supplementary table S4.

qPCR was performed in a CFX96^TM^ Real-Time System (Bio-Rad) with SYBR Green I assay. The amplification reaction was prepared to a final volume of 10 µl containing 2.7 µl of sterilized distilled water, 5 µl of *i*Taq Universal SYBR^®^ Green Supermix (BIO-RAD), 0.05 µl (10 µM) of each primer, 0.2 µl of Bovine Serum Albumin (BSA) and 20 ng of template DNA. The qPCR amplification began with an initial incubation for 3 minutes at 95 °C, followed by 40 cycles of 95°C for 10 s and a 30 s incubation for primer annealing (Supplementary Table S4). All the reactions were set up in triplicates and a mixture of DNA from other non-target fungi was used as the negative control, while autoclaved distilled water was used as the non-template control. DNA of pure cultures of *Termitomyces*, *Pseudoxylaria* and *Pseudomonas* were amplified using their specific primer sets (Supplementary Table S4) in a serial dilution of 10^-1^ x to 10^-5^ to obtain the standard curve. Target copy numbers for each reaction were calculated from the standard curves^61^ using the formula:

Number of copies/ µl = ((amount used in ng) * (6.022 * 1023 molecules/mole)) / (length of amplicon) * (650 g/mole)

The specificity of the reactions was assessed by the analysis of the melting curve. The copy numbers obtained were then used to generate the number of amplicons in the target samples with the following formula^62^:

X_օsample_ = E_AMPsample_ [b_abs_ ^*log^ (E_AMPsample_ E_AMPabs_)-Cq_sample_]

X_օsample_ = number of copies

E_AMPsample_= Exponential amplification of the sample EAMPabs= Exponential amplification of the standards

C_qsample_= Quantification cycle for the sample

b_abs_= Standard curve of the intercept

The statistical significance of these copy number variations was done with a two-tailed t-test at p< 0.05.

### In vitro anti-fungal assays with Pseudoxylaria and Termitomyces

*In vitro* interaction experiments were done to identify whether *Termitomyces* and *Pseudoxylaria* can prevent the growth of the fungal contaminants (Supplementary Note 5). The magnitude of inhibition by test fungi was evaluated by using the following formula from Royse & Ries^63^:

% Inhibition = ((Total area of test fungus in control in mm^2^) – (Total area of test fungus in the interaction in mm^2^)) * 100 / Total area of test fungus in control in mm^2^

For easy comprehension, the interactions were categorized into the following three categories: more than 50% inhibition, less than 50% inhibition and no inhibition. To explore the effect of *Termitomyces*, the spread method was used ^33^ where a liquid suspension of *Termitomyces* nodules/mycelia was prepared in 1X PBS and spread on PYME ^22^ plates (Supplementary Fig. S13). These plates were first incubated for 36 h at 30°C before placing a plug of the test fungus at the center. For interaction with *Pseudoxylaria*, a dual plug assay was used (Supplementary Fig. S14) in 90 mm diameter Petri dishes containing PDA and 250 µg/mL chloramphenicol. Plates were inoculated separately with a 1 cm^2^ disc of actively growing *Pseudoxylaria*, 10 mm away from the edge of the plates. 1 cm^2^ disc of the test fungus was placed separately in the same plate but 60 mm away from the *Pseudoxylaria* plug (Supplementary Fig. S14). The control growth plates were set up for the test fungi and *Pseudoxylaria*. These were then incubated in the dark at 30 °C in Heratherm^TM^ Compact Microbiological Incubator (Thermo Fisher Scientific). All the interaction plates (with both *Pseudoxylaria* and *Termitomyces*) were evaluated for growth inhibition by taking photographs using the Panasonic Lumix G2 camera. Photographs from the 7^th^ day were taken as the final data and were analyzed in Adobe Photoshop CS6 by measuring the area of test fungi growth in their respective interactions (both test and control plates) in pixels and converting it into millimeters (100 pixels = 1 mm). All interaction assays were repeated with sample sizes ranging from 3-5. Representative figures for all the interaction assays are given in supplementary figures S13-S14.

## *In vitro* anti-fungal assays with mutualistic bacteria

The fungal contaminants which were inhibited to less than 50% by both the *Termitomyces* and *Pseudoxylaria* were further selected for the bacterial-fungal assays. Paper disc diffusion assays were performed to identify whether the previously identified eight bacterial mutualists from the same mounds^33^ prevented the growth of these fungal contaminants. The interactions were categorized into four broad categories of clear zone of inhibition, reduced growth of fungus near the bacteria, contact inhibition, and negligible inhibition^33^. The magnitude of inhibition of the fungal growth by these bacteria was evaluated using the formula mentioned above from Royse & Ries^63^. The cultures of *Phoma* and *Fusarium* grow much slower than the other fungi tested. To avoid any discrepancy in the results due to differences in growth, the spread method was used for these two fungi and four bacterial discs were placed compared to a single disc. The culture of *Paraconiothyrium*-MN913759 was lost during this study and therefore, bacterial interactions against this fungus could not be done.

## Data availability

Data Accessibility: The mycobiome dataset generated is available as supplementary information file to this article. ITS gene fragments obtained have been submitted to GenBank (accession number MN913749–MN913772) and Nanopore sequencing generated have been submitted to NCBI Sequence Read Archive (SRA) under the BioProjects PRJNA665655 (SAMN16262892 - SAMN16262897), PRJNA758817 (SAMN21036816 – SAMN21036821) and PRJNA758827 (SAMN21037110 – SAMN21037115).

## Acknowledgments

We thank Abin Antony and Raunak Dhar for their help with the fieldwork, Dr. Shashi Bhushan Pandit and Paras Verma for their technical help with the data analysis softwares. We acknowledge all the EVOGEN lab members for their valuable comments on the manuscript.

## Author Contributions

RR and RA devised the study. RA performed field collections, sequencing, microbial culturing, data collection and analysis. NES helped in the fieldwork, culturing of fungi and pilot experiments. Nanopore sequencing was done by RA and MG and its analysis was done by RA. The statistical analysis of interaction experiments was done by RA and RS. RA, RS and RR wrote the paper.

