## Supplementary_file for "Partnership with both fungi and bacteria can protect *Odontotermes obesus* fungus gardens against fungal invaders"

### Table of Contents

| | Page No. |
|---|---|
| Supplementary Note 1: Sample Collection/Mounds used | S3 |
| Supplementary Note 2: DNA extraction and PCR amplification | S4 |
| Supplementary Note 3: Identification of isolated fungal strains | S5 |
| Supplementary Note 4: Nanopore Sequencing | S6-S7 |
| • Sample preparation and PCR amplification | S6 |
| • Library prepartion | S6 |
| • Data analysis workflow | S7 |
| Supplementary Note 5: In vitro anti-fungal assays with *Pseudoxylaria* and *Termitomyces* | S8 |
| Fig. S1: Location of *O. obesus* mounds | S9 |
| Fig. S2: Different castes of *O. obesus* | S10 |
| Fig. S3: Reproductives of *O. obesus* | S11 |
| Fig. S4: Setting up the fungus combs to decay | S12 |
| Fig. S5: A Phylogenetic tree of *Pseudoxylaria* using α-actin gene | S13 |
| Fig. S6: A Phylogenetic tree of *Pseudoxylaria* using ITS gene | S14 |
| Fig. S7: Pan-mycobiota analysis | S15 |
| Fig. S8: Microbiota of the decaying combs | S16 |
| Fig S9: Relative abudance of bacterial genera in degrading fungus comb | S17 |
| Fig. S10: Change in abundances of *Termitomyces*, *Pseudoxylaria* and bacterial mutualists in decomposing fungus comb. | S18 |
| Fig. S11: Change in abundances of *Termitomyces*, *Pseudoxylaria* and other fungi in decomposing fungus comb**.** | S19 |
| Fig. S12: Interaction assay between *Pseudoxylaria* and *Termitomyces* | S20 |
| Fig. S13: *Termitomyces* Vs Fungal contaminants | S21-22 |
| Fig. S14: *Pseudoxylaria* Vs Fungal contaminants | S23-24 |
| Fig. S15: Re-using media from the test experimental plates | S25 |
| Fig. S16: *Bacillus*- MN908297 Vs Fungal contaminants | S26 |
| Fig. S17: *Bacillus*- MN908298 Vs Fungal contaminants | S27 |
| Fig. S18: *Bacillus*- MN908304 Vs Fungal contaminants | S28 |
| Fig. S19: *Bacillus*- MN908305 Vs Fungal contaminants | S29 |
| Fig. S20: *Burkholderia*- MN908310 Vs Fungal contaminants | S30 |
| Fig. S21: *Pseudomonas*- MN908321 Vs Fungal contaminants | S30 |
| Fig. S22: *Pseudomonas*- MN908322 Vs Fungal contaminants | S31 |
| Fig. S23: *Streptomyces*- MN908327 Vs Fungal contaminants | S32 |
| Table S1: Details of the fungal strains isolated for the culture-dependent assay. | S33 |
| Table S2: The detailed information of termite castes used to study the mycobiota. | S34 |
| Table S3: The detailed information of the degrading comb samples used to study the micro- and mycobiota. | S35 |
| Table S4: List of primers used in this study | S36 |
| Table S5: The mycobiota of the degrading fungus comb of *O. obesus*. | S37 |
| Table S6: No. of copies of *Termitomyces, Pseudoxylaria and Pseudomonas* in different degrading stages of fungus comb using qPCR estimation | S38-S39 |
| Table S7: The microbiota of the degrading fungus comb of *O. obesus*. | S40 |
| Table S8: Relative abundances of fungal OTUs in different castes and degrading fungus comb obtained from Nanopore sequencing. | S41-S63 |
| Table S9 :Relative abundances of the bacterial communities across degrading fungus comb obtained from Nanopore sequencing | S64-118 |
| References | S119 |

**Supplementary Note 1: Sample Collection/Mounds used:**

Different castes of *O. obesus* were collected from two different mounds of the IISER Mohali campus. Fresh fungus combs were collected across three months (August-October, 2018) from two mounds to set up the incubation to decay for 120 hrs. The collected combs were immediately kept in sterile plastic containers and were then crushed with autoclaved pestles, divided into roughly 0.5 g portions, and incubated at 30 °C inside sterilized glass containers. For each day, three such containers were set up from each mound. Major workers, minor workers and nymphs collected from the combs were kept in separate sterile vials. Regular visits were made to these mounds, in anticipation of emerging alates, after the first monsoon shower in June 2018. Male and female alates were collected directly from the mounds and kept individually in sterile vials. DNA was extracted from individual termite and comb samples using the CTAB (Cetyl Trimethyl Ammonium Bromide) method.

**Supplementary Note 2: DNA extraction**

1 ml of CTAB buffer containing 100 mM Tris (pH 8), 2% CTAB, 1.4 M NaCl, 20 mM EDTA, 1% β-mercaptoethanol and 0.3 g $mL^{-1}$ proteinase K was used for DNA extraction from termite castes and fungus comb samples. The samples were crushed in the buffer and then incubated at 60˚C for 30 minutes. These were then subjected to Phenol: Chloroform: Isoamyl alcohol (PCI: 25:24:1) precipitation and the purified DNA pellet obtained was dissolved in 1X TE (pH 8) buffer. Extracted DNA from fungus combs often have extensive humic acid contamination which can hinder downstream PCR reactions. To remove this humic acid, the precipitation step with Phenol: Chloroform: Isoamyl alcohol was repeated until the final pellet obtained was white in color. DNA samples were used for PCR only if the 260/280 nm ratio were within the range of 1.3-1.9, which were obtained through a NanoDrop Spectrophotometer 2000 (Thermo-Fisher Scientific). The 260/280 nm ratio was variable for the DNA extracted from the comb samples due to the natural presence of humic acid[1]. DNA from the combs was extracted consecutively for six days, i.e., 0 hrs to 120 hrs. For 0 hrs, DNA was extracted from each portion separately within 3 hours of the collection.

**Supplementary Note 3: Identification of isolated fungal strains**

DNA from morphologically distinct fungi were extracted with CTAB buffer and amplified with the primer set ITS4/ITS5[2]. The amplification reaction was prepared to a final volume of 20 µl containing: 14.5 µl sterile distilled water, 0.4 µl dNTPs (10 µM), 2 µl 17.5 mM buffer, 0.5 µl of each primer (10 µM), 2 µl template and 0.1 µl *Taq* Polymerase (Himedia). PCR was performed under following conditions: an initial denaturation step at 95°C for 3 minutes, followed by 37 cycles of denaturation (95°C, 45 seconds), annealing (56°C, 45 seconds), extension (72°C, 1 minute) with a final extension at 72°C for 10 minutes.

The PCR products were cleaned with Exonuclease I and Shrimp alkaline phosphatase and then sequenced with BigDye® Terminator v.3.1 cycle sequencing kit for both strands. The chromatograms obtained were cleaned with Sequencher v.5.2.4 (GeneCodes Corp.) and the sequences were manually edited with Bioedit v.7.0.5.3[3]. Taxonomic identification of these sequences was achieved through BLAST with the parameters of >90% sequence identity and/or the first hit. The sequences were aligned with other similar homologues, obtained from NCBI, using the ClustalW in Bioedit. The Maximum likelihood phylogenetic tree of aligned sequences was constructed using MEGAX v.10.1.7[4] with 1000 bootstrap replicates. Suitable substitution models for various phylogenetic analyses were also obtained through MEGAX[4]. All the fungi selected for use in these experiments had unique sequence profiles in their Internal Transcribed Spacer (ITS) gene.

**Supplementary Note 4: Nanopore Sequencing**

- **Sample preparation and PCR amplification**

Two separate sets of nanopore runs were done. The first was to identify the pan-mycobiota and the second was to identify the microbiota of the degrading combs. To obtain a comprehensive assessment of fungal diversity, three separate DNA extractions per mounds were pooled in equimolar concentrations. This was done for all the termite castes, except alates, where three individual DNA extractions from males and females were pooled (Table S2 and S3). Five different DNA samples (major worker, minor worker, nymph, male alate and female alate) were thus prepared for mycobiota estimation and the other six comb samples (0 hrs, 24 hrs, 48 hrs, 72 hrs, 96 hrs and 120 hrs) were prepared for both micro- and mycobiota estimation on the Nanopore platform.

To identify the mycobiota present within the termite castes and fungus comb, the ITS fragment was amplified using the primers ITS5/ITS4[2] whereas to characterize the microbiota in a degrading fungus comb, *16S* rRNA[5] gene regions was amplified. The amplification reaction was prepared to a final volume of 20 µl containing: 14.5 µl sterile distilled water, 0.4 µl dNTPs (10 µM), 2 µl 17.5 mM buffer, 0.5 µl of each primer (10 µM), 2 µl template and 0.1 µl *Taq* Polymerase (Himedia). PCR conditions involved an initial denaturation step at 95°C for 3 minutes, followed by 33 cycles of denaturation (95°C, 45 seconds), annealing (56°C, 45 seconds), extension (72°C, 1 minute) with a final extension at 72°C for 10 minutes. PCR products were cleaned using Wizard® SV Gel and PCR clean-Up System (Promega). Purified PCR products were quantified using NanoDrop 2000 (Thermo Scientific) and Qubit 3.0 Fluorometer (Life Technologies).

- **Library prepartion**

An initial library preparation was done using the SQK-LSK108 Ligation Sequencing Kit 1D-vR9 (Oxford Nanopore Technologies) by Ultra II End-prep enzyme mix. The resultant mixture was purified using 1X AMPure XP beads and eluted in nuclease-free water. These were individually ligated with barcode adaptors and purified using 1X AMPure XP beads and eluted in nuclease-free water. These different samples were then individually barcoded with unique barcodes using the EXP-PBC001 PCR barcoding kit I (https://community.nanoporetech.com/protocols).

Barcoded samples were then purified using Wizard® SV Gel and the PCR clean-Up System (Promega) and quantified using Qubit 3.0 Fluorometer (Life Technologies). All the twelve barcoded samples were pooled in equal amount (20-25 ng/µl) before being loaded on the Nanopore flowcell. 5 µl of λ phage DNA was added to the library to serve as a control. Samples were sequenced on a MinION platform (MinION Mk1B). The sequencing library was run on FLO-MIN106 flowcell for 48 hours using MinKNOW software with the protocol *NC_48Hr_sequencing_FLO-MIN106_SQK-LSK108_plus_Basecaller*.

- **Data analysis workflow**

After the completion of the run, sequences were first separated according to their barcodes by using the program ont-albacore (Oxford Nanopore Technologies Ltd.) These were then converted to FASTA and FASTQ files using *Poretools* (v 0.5.1) for downstream analysis[6]. High-quality reads (average read quality score ≥ 10) were selected using the program *Nanofilt*[7]. The obtained high-quality sequences were then further processed where the barcodes and adaptors were removed using Porechop (https://github.com/rrwick/Porechop). The sequencing exhibited the average accuracy of about 89% which was estimated by aligning control λ phage DNA with existing NCBI λ sequences (NC_001416.1) using LAST v. 973. The fungal OTU's were identified by comparing them against the fungus repository from NCBI FTP site and the bacterial sequences generated from the six comb samples (0 hrs- 120 hrs) were identified by comparing them against the customized microbial repository made by combining 16S rRNA gene fragments from NCBI FTP site and the DictDb v 3.0 database[8] . Annotation was done using LAST v 973, with the following parameters: match score of 1, gap opening penalty of 1, and gap extension penalty of 1[9]. The identified reads were sorted from the phylum to the genus level and their relative abundances were calculated.

**Supplementary Note 5: *In vitro* anti-fungal assays with *Pseudoxylaria* and *Termitomyces***

*In vitro* interaction experiments were done to identify whether *Termitomyces* and *Pseudoxylaria* can prevent the growth of the fungal contaminants. A simple explanation for the inhibitory capabilities seen for both *Pseudoxylaria* and *Termitomyces* could be the depletion of nutrients in the PDA plates. However, control experiments using previously used media to grow these fungi indicate the presence of sufficient nutrients in these plates for adequate fungal growth (Fig. S15).

### Fig. S1: Location of *O. obesus* mounds

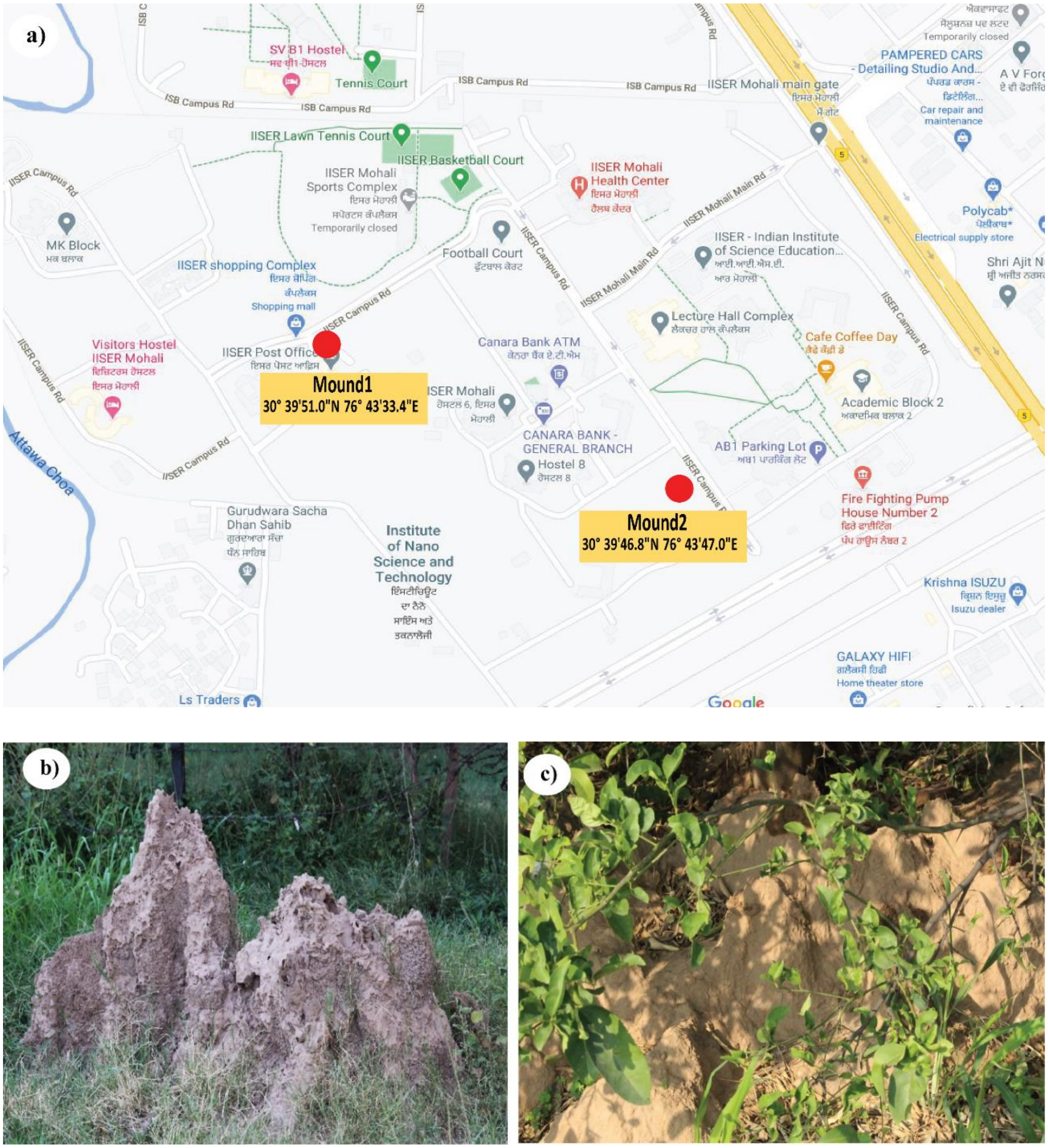

Fig. S1: a) Geographical location of *Odontotermes obesus* colonies in IISER Mohali campus. b) & c) Two mounds of *O. obesus* used in the study.

**Fig. S2: Different castes of *O. obesus***

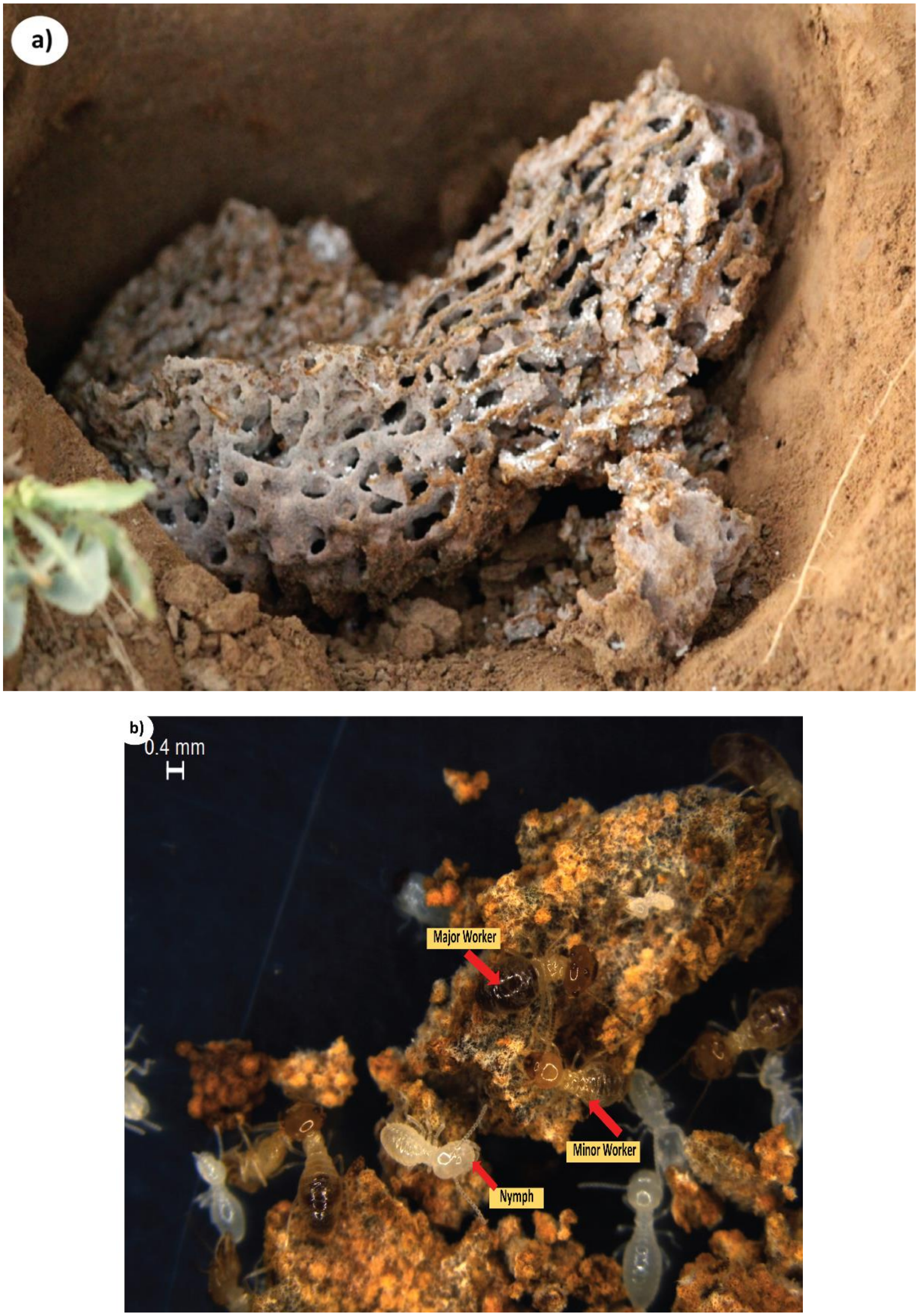

Fig. S2: a) Detailed view of fungus comb of *O. obesus* b) Different castes of termites present over the fungus comb.

**Fig. S3: Reproductives of *O. obesus***

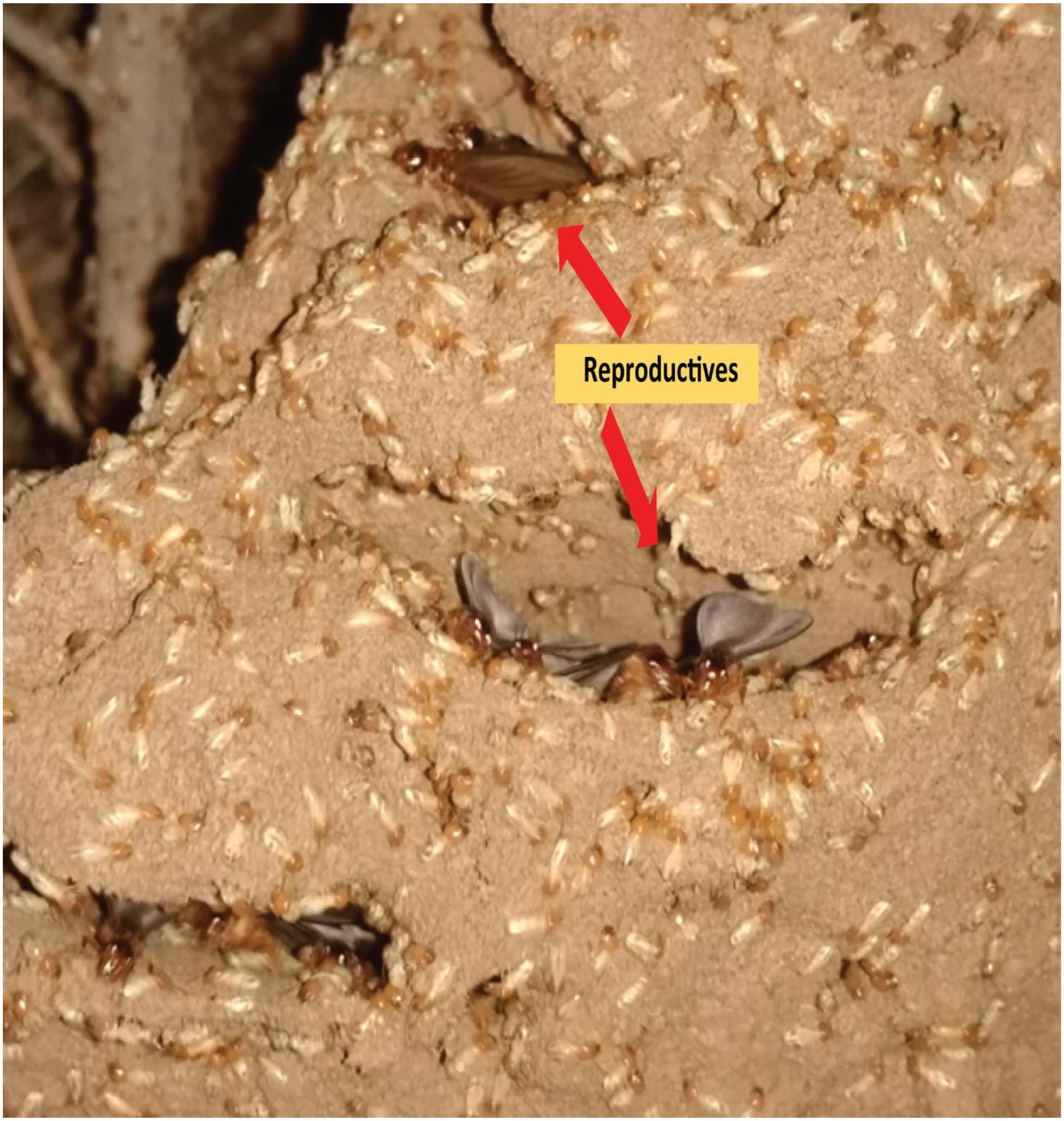

Fig. S3: Reproductives emerging out from the gaps in mound during monsoon season from one of the mounds in this study.

### Fig. S4: Setting up the fungus combs to decay

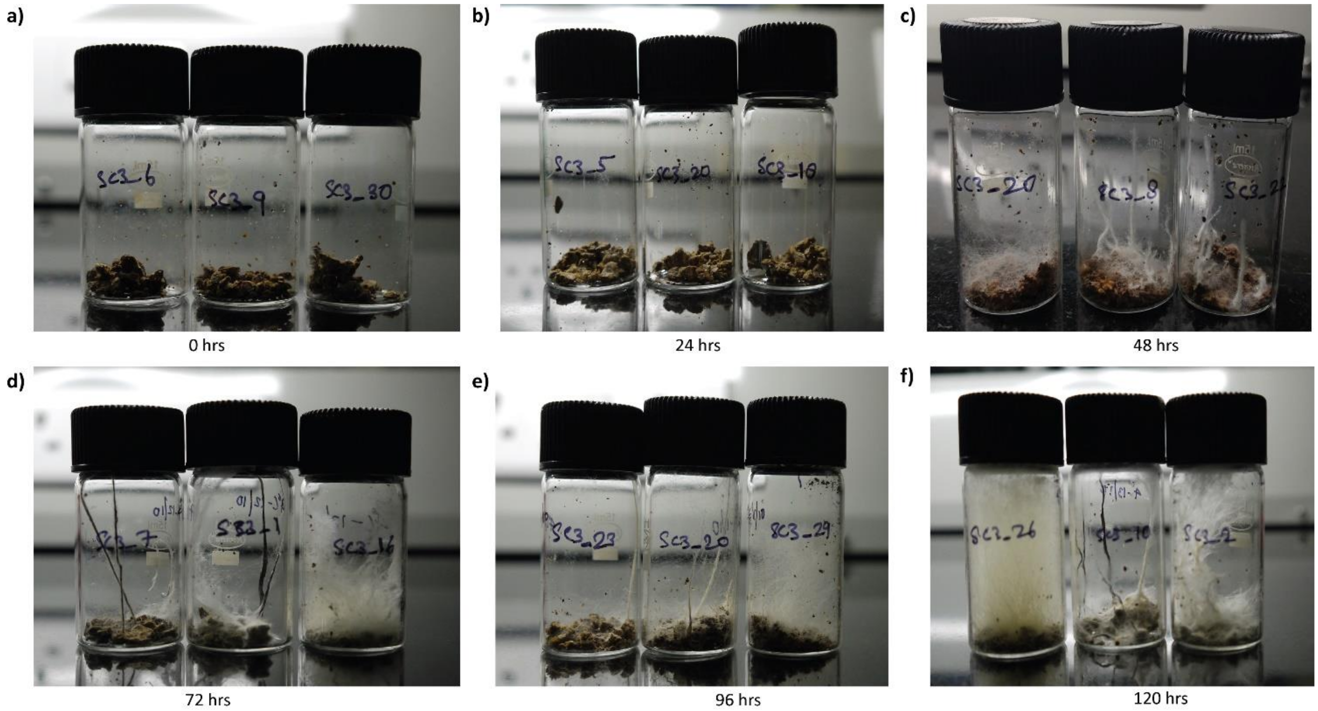

Fig. S4: Crushed fungus combs inside sterilized glass containers. Fresh fungus combs were collected across three months (August-October, 2018) from two different mounds and were crushed with autoclaved pestles, divided into roughly 0.5 g portions. For each day, three such containers were set up from each mound which were further used for DNA extraction and sequencing.

**Fig. S5: A Phylogenetic tree of *Pseudoxylaria* using α-actin gene**

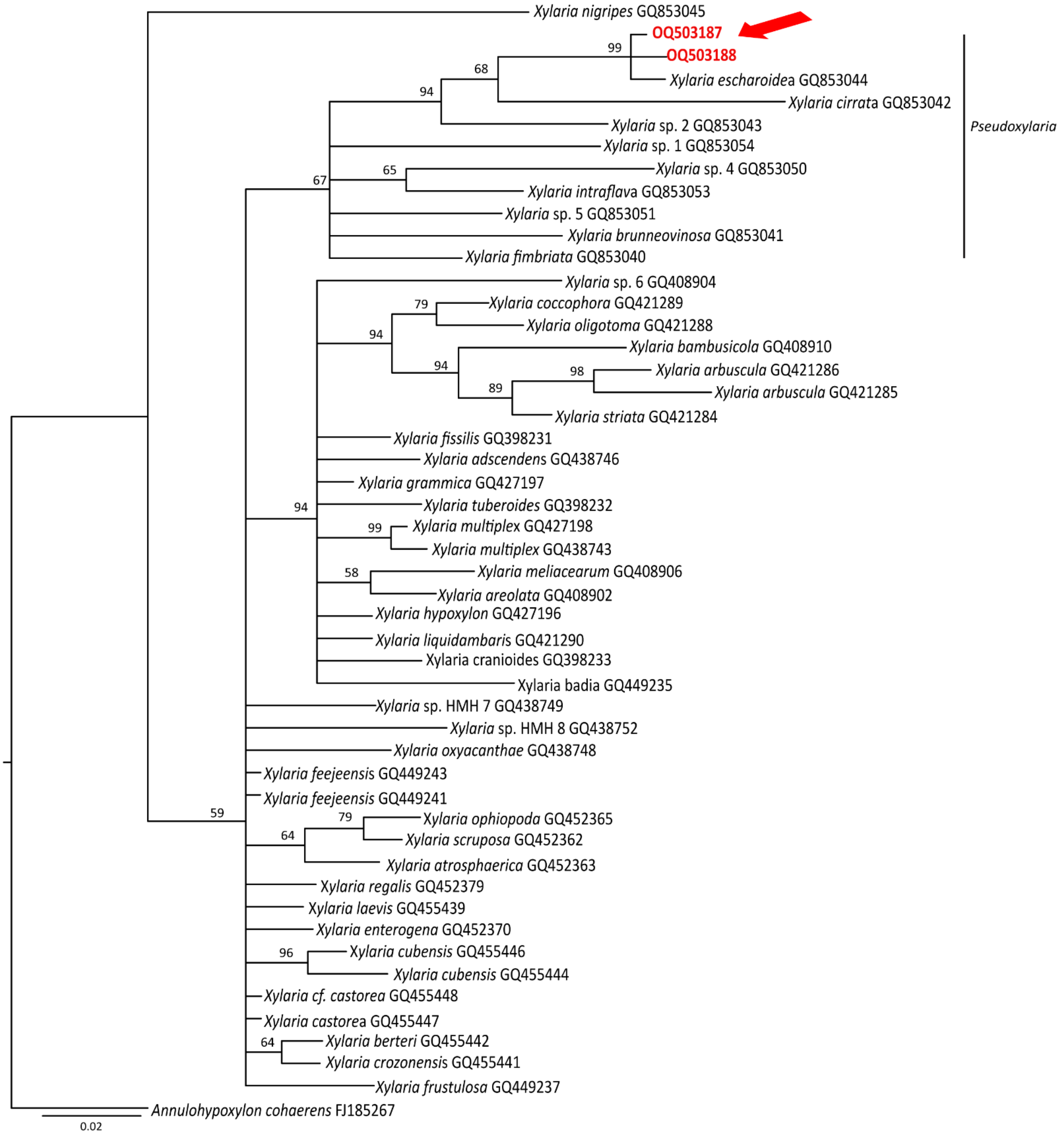

Fig. S5: A phylogenetic tree of *Pseudoxylaria* generated by using Bayesian analysis from the α-actin gene. *Annulohypoxylon cohaerens* was used as the outgroup taxa. Taxa in bold red and also indicated with a red arrow were obtained from this study and other all taxa were obtained from Hsieh et al.[10].

**Fig. S6: A Phylogenetic tree of *Pseudoxylaria* using ITS gene**

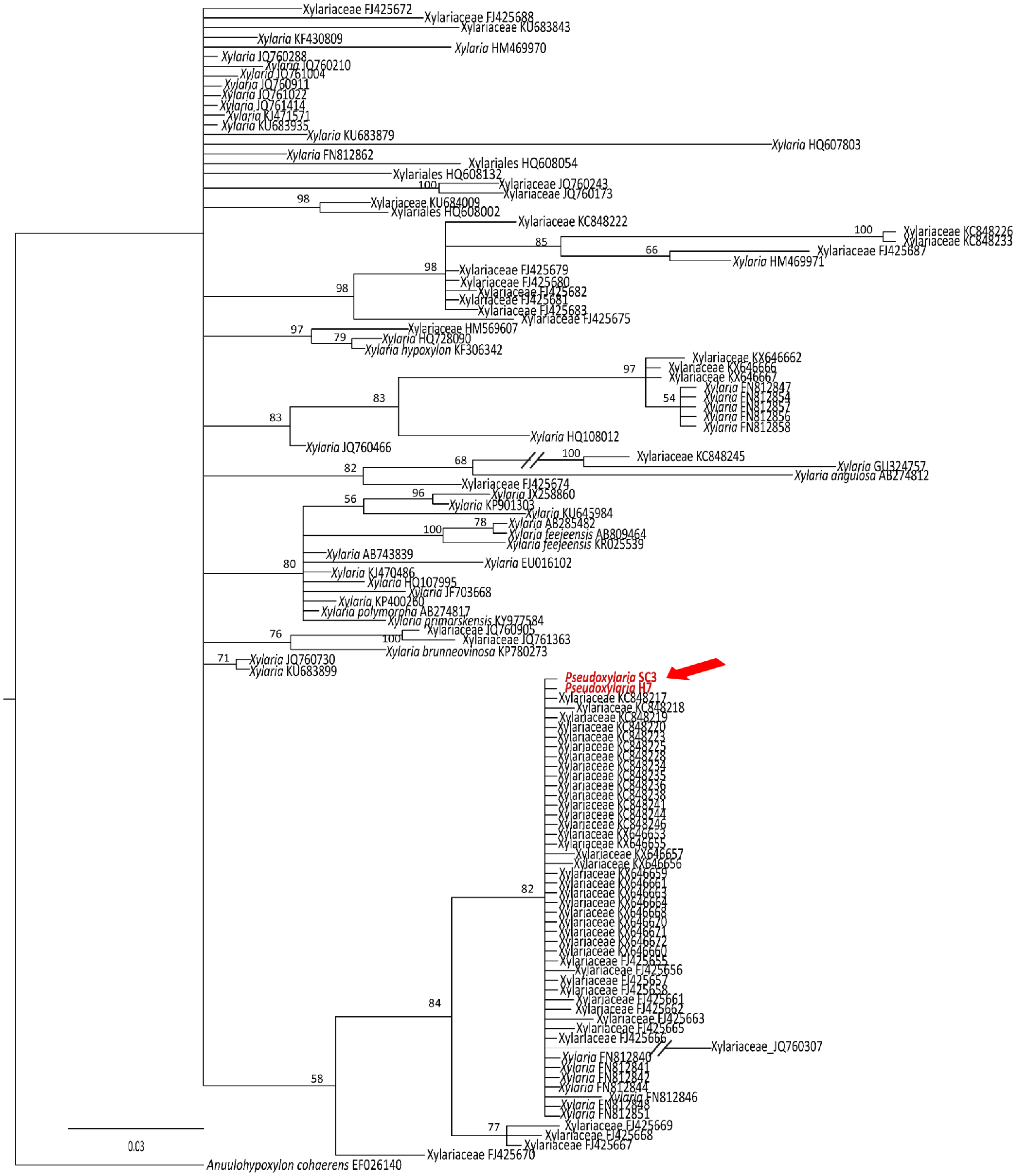

Fig. S6: A phylogenetic tree of *Pseudoxylaria* generated by using Bayesian analysis from the ITS gene. *Annulohypoxylon cohaerens* was used as the outgroup taxa. Taxa in bold red and also indicated in red arrow were obtained in culture and other all taxa were obtained from the Nanopore sequencing of this study.

### Fig. S7: Pan-mycobiota analysis

a)

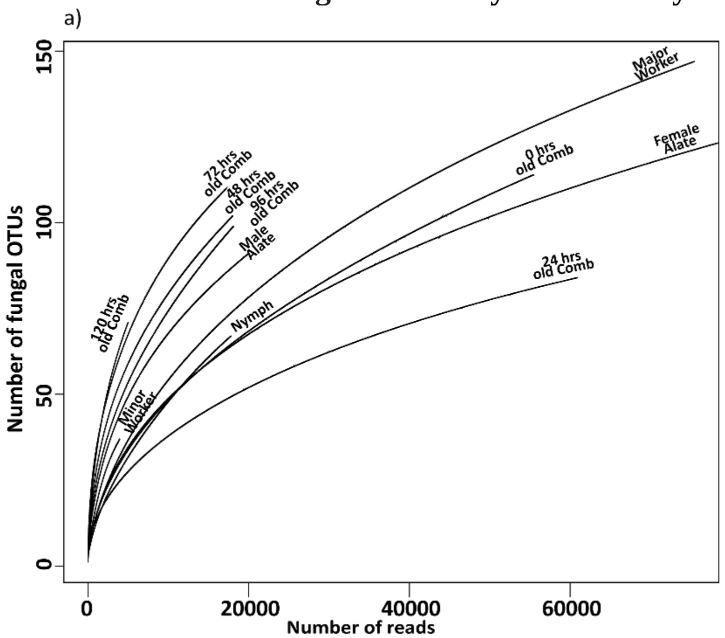

b)

| Sample source | Total reads |
|---|---|
| Major Worker | $0.92 \times 10^5$ |
| Minor Worker | $0.04 \times 10^5$ |
| Nymph | $0.22 \times 10^5$ |
| Female Alate | $1.15 \times 10^5$ |
| Male Alate | $0.19 \times 10^5$ |
| Decaying comb (0 hrs) | $0.58\times10^5$ |
| Decaying comb (24 hrs) | $0.62\times10^5$ |
| Decaying comb (48 hrs) | $0.16\times10^5$ |
| Decaying comb (72 hrs) | $0.16\times10^5$ |
| Decaying comb (96 hrs) | $0.17\times10^5$ |
| Decaying comb (120 hrs) | $0.04\times10^5$ |

c)

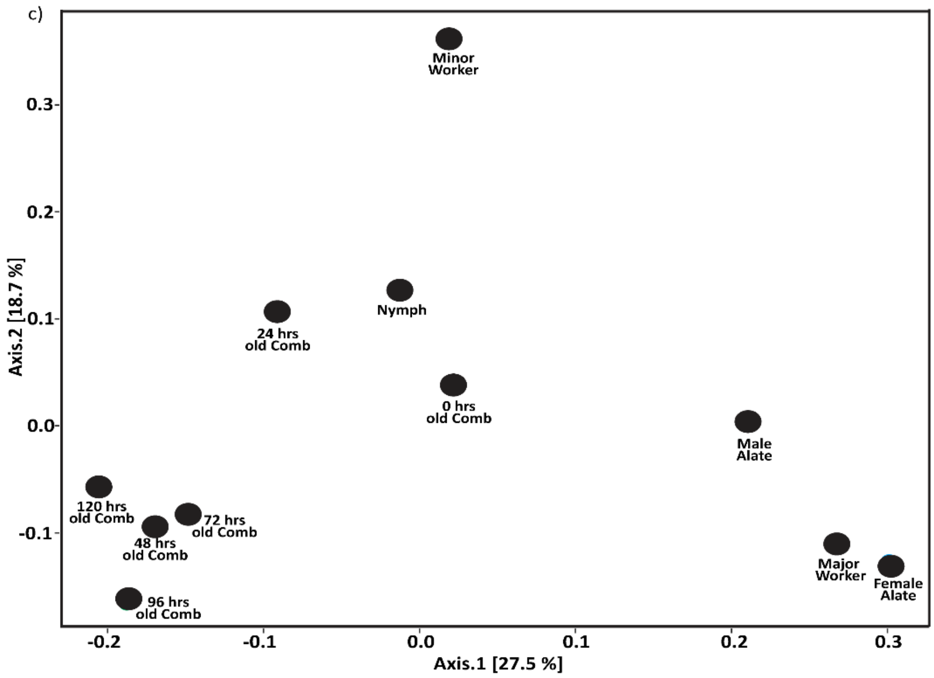

Fig. S7: The pan-mycobiota analysis of degrading fungus comb a) Rarefaction analysis of fungal OTUs from six different sources of *O. obesus* colony, b) Number of reads obtained to identify the mycobiota of *O. obesus* from Nanopore Sequencing c) PCoA similarity analysis of six samples of this study using weighted Unifrac distance.

### Fig. S8: Microbiota of the decaying combs

a)

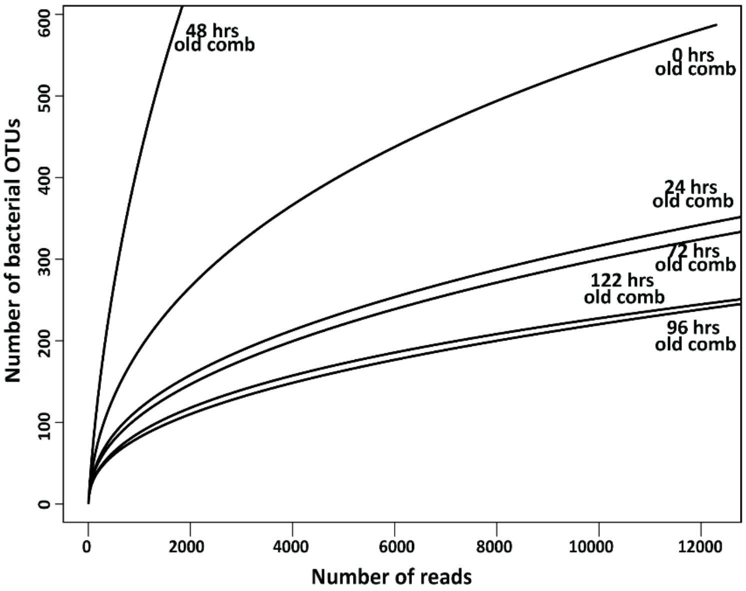

b)

| Sample source | Total reads |
|---|---|
| **Decaying comb (0 hrs)** | $0.14x10^{5}$ |
| **Decaying comb (24 hrs)** | $0.94x10^{5}$ |
| **Decaying comb (48 hrs)** | $0.38x10^{5}$ |
| **Decaying comb (72 hrs)** | $1.54x10^{5}$ |
| **Decaying comb (96 hrs)** | $0.63x10^{5}$ |
| **Decaying comb (120 hrs)** | $1.33x10^{5}$ |

c)

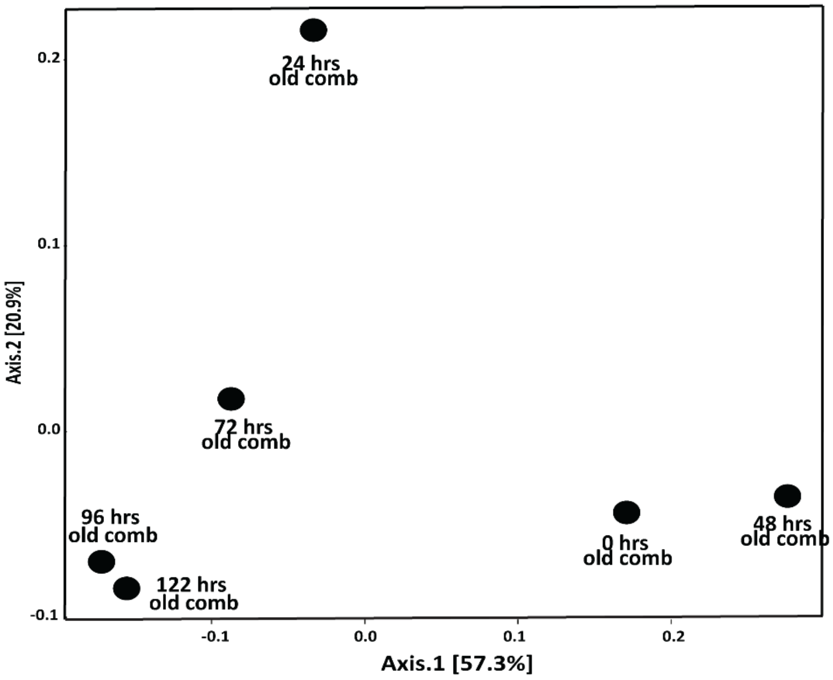

Fig. S8: Microbiota analysis of degrading fungus comb. a) Rarefaction analysis of the six samples, b) Number of reads obtained to identify the microbiota of *O. obesus* from Nanopore sequencing c) Weighted Unifrac PCoA analysis.

**Fig. S9: Relative abudances of the bacterial genera in the decaying combs**

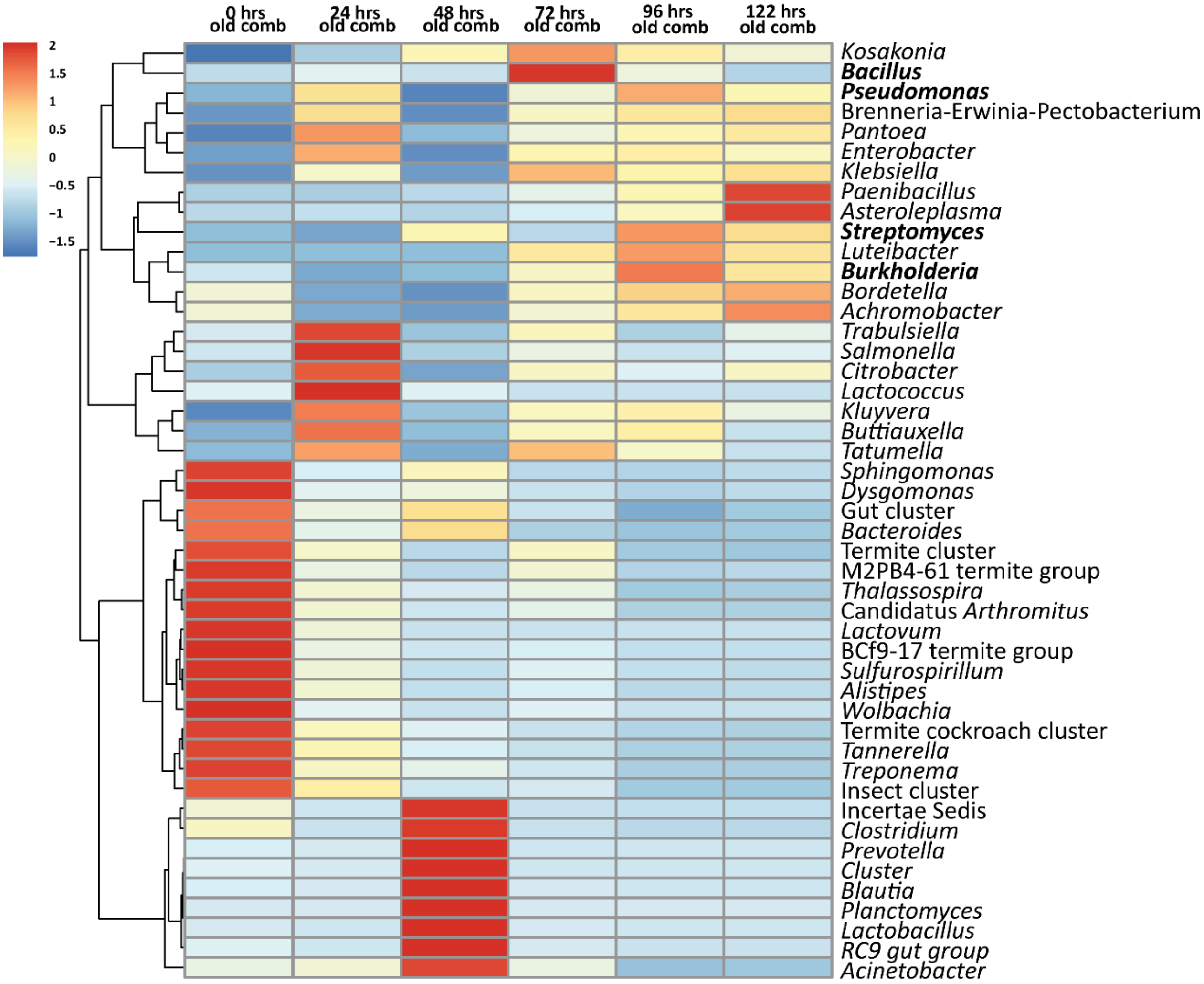

Fig S9: Heatmap of the bacterial genera with relative abundance of at least 0.05% in any of the samples.

**Fig. S10: Change in abundances of *Termitomyces*, *Pseudoxylaria* and bacterial mutualists in decomposing fungus comb.**

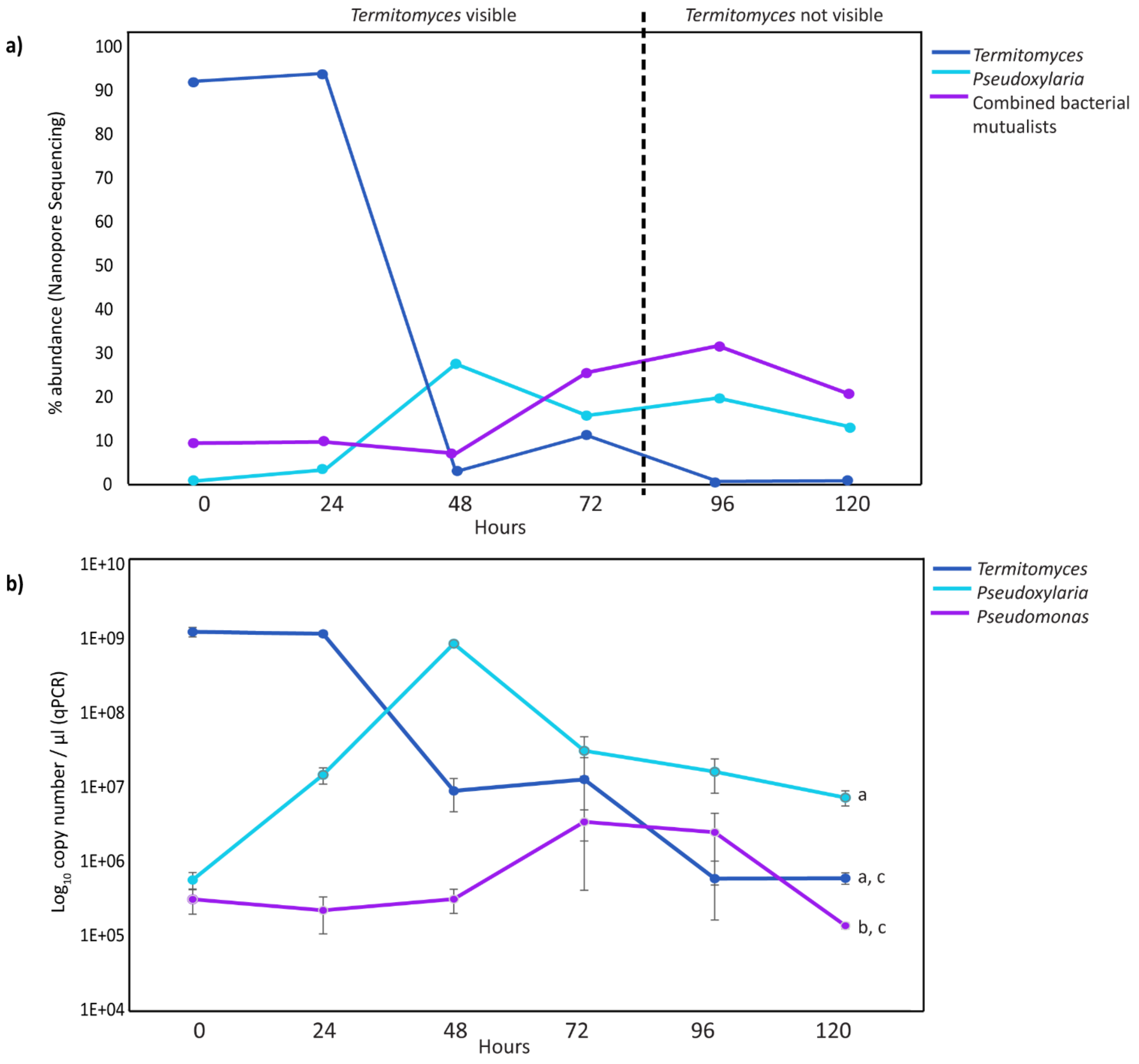

Fig. S10: a) Change in absolute abundances of *Termitomyces*, *Pseudoxylaria* and bacterial mutualists in different decomposing stages of fungus comb. b) Change in copy numbers of *Termitomyces*, *Pseudoxylaria* and *Pseudomonas* as determined by qPCR analysis. One- way Anova showed significant variation between the groups (p= 0.00081, F= 7.512, df= 2). Post hoc Tukey tests revealed insignificant pairwise site differences in the copy numbers of *Termitomyces* vs *Pseuodxylaria* (p= 0.102) and *Pseudoxylaria* and *Pseudomonas* (p= 0.364).

**Fig. S11: Change in abundances of *Termitomyces*, *Pseudoxylaria* and other fungi in decomposing fungus comb.**

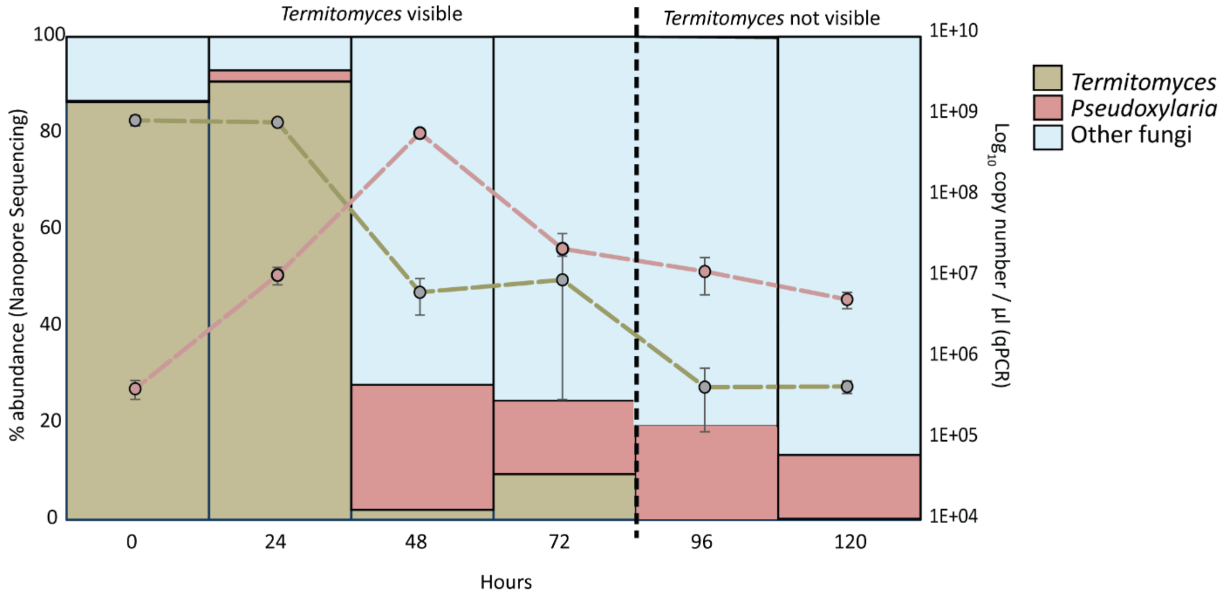

Fig. S11: Change in abundance (in percentages) of *Termitomyces*, *Pseudoxylaria* and other fungi, and $\log_{10}$ copy numbers of *Termitomyces* and *Pseudoxylaria* in different stages of the decay of the fungus comb as determined by Nanopore Sequencing (bar graph) and qPCR (line graph).

**Fig. S12: Interaction assay between *Pseudoxylaria* and *Termitomyces***

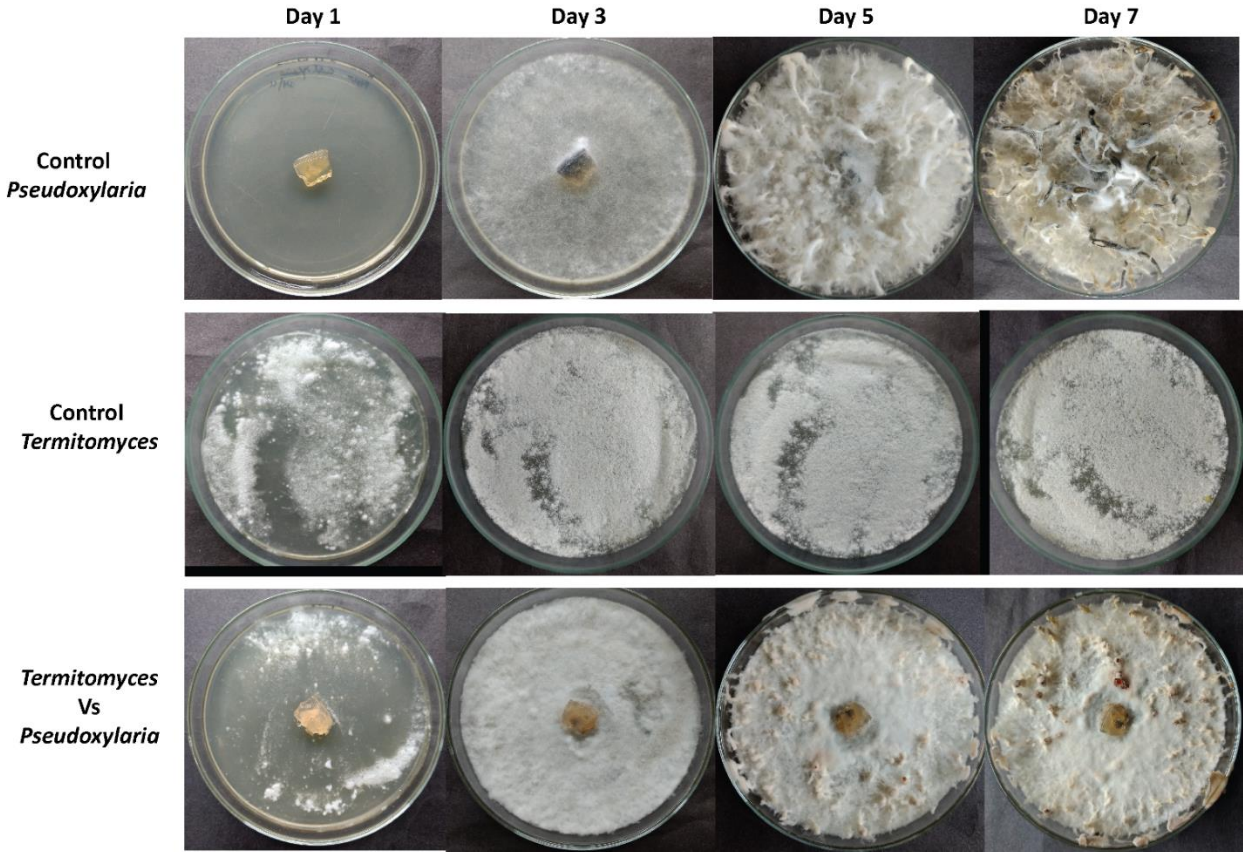

Fig. S12: Interaction assay between *Pseudoxylaria* and *Termitomyces*. Interaction is represented with its control and test plate. *Pseudoxylaria* easily grow over *Termitomyces* indicating that the crop fungus cannot prevent the most prominent weedy fungus. Also with its faster growth rate, *Pseudoxylaria* can be a more efficient inhibitor of non-specific fungi than *Termitomyces*.

**Fig. S13: *Termitomyces* Vs Fungal contaminants**

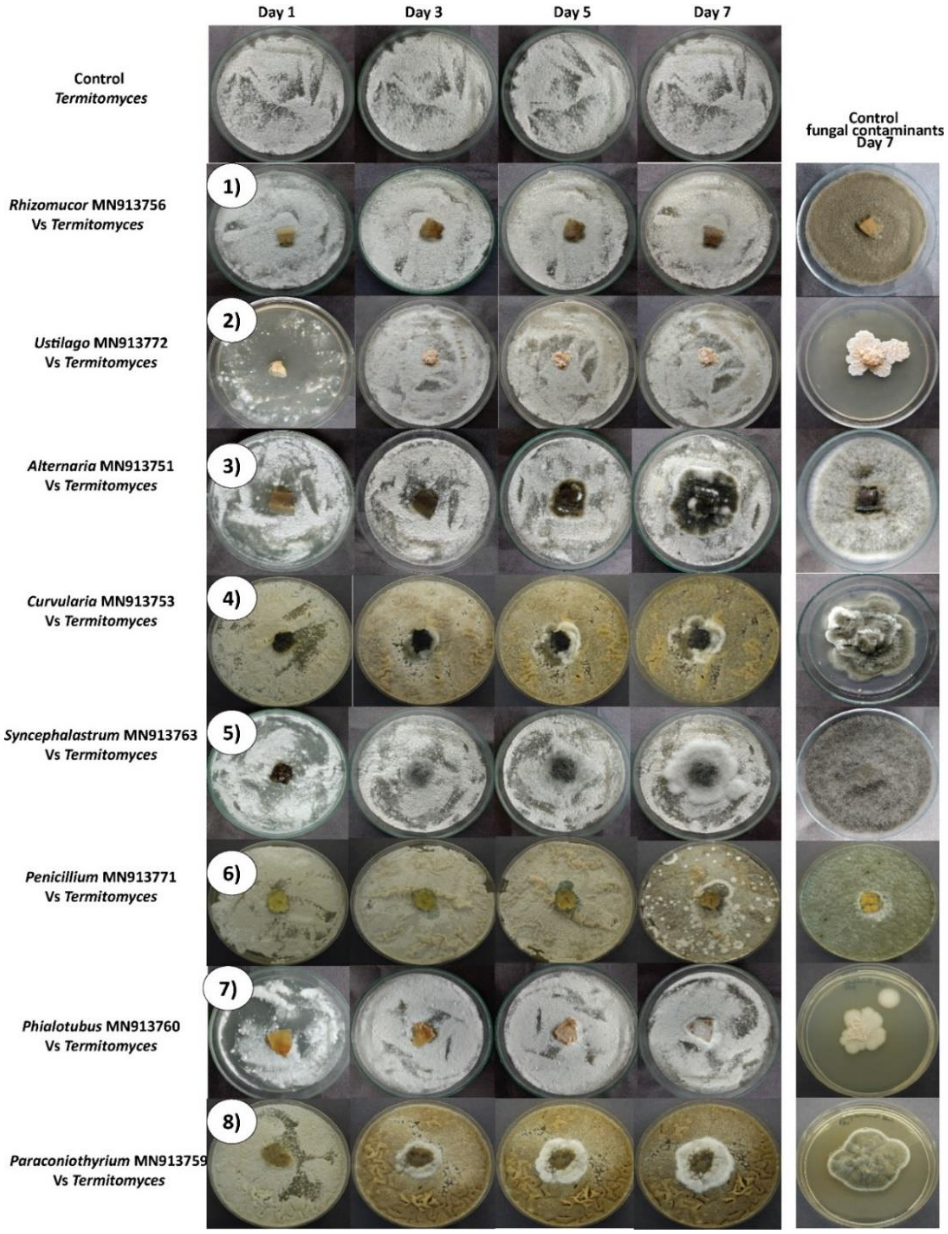

Fig. S13: Interaction assay of *Termitomyces* against fungal contaminants. All the interactions are categorized into three types: **1)-8) more than 50% inhibition 8) - 14) less than 50 % inhibition** and **15)-16) negigible inhibition** by the *Termitomyces*. Each interaction is represented with its fungal and *Termitomyces* controls and a representative test plate. *Termitomyces* showed maximum inhibition (more than 50%) against *Alternaria*-MN913751, *Curvularia*-MN913753, *Rhizomucor*-MN913756, *Syncephalastrum*-MN913763 and *Ustilago*-MN913772. The other nine fungi, *Aspergillus*-MN913749, *Aspergillus*-MN913750, *Diaporthe*-MN913762, *Fusarium*-MN913754, *Lasiodiplodia*-MN913758, *Paraconiothyrium*-MN913759, *Penicillium*-MN913771, *Phialotubus*- MN913760 and *Phoma*-MN913761 showed less than 50% inhibition. However, *Mucor*-MN913757 and *Trichoderma*-MN913767 overgrew *Termitomyces*, showing no sign of inhibition. All the photos are from day 7 of the beginning of the experiment.

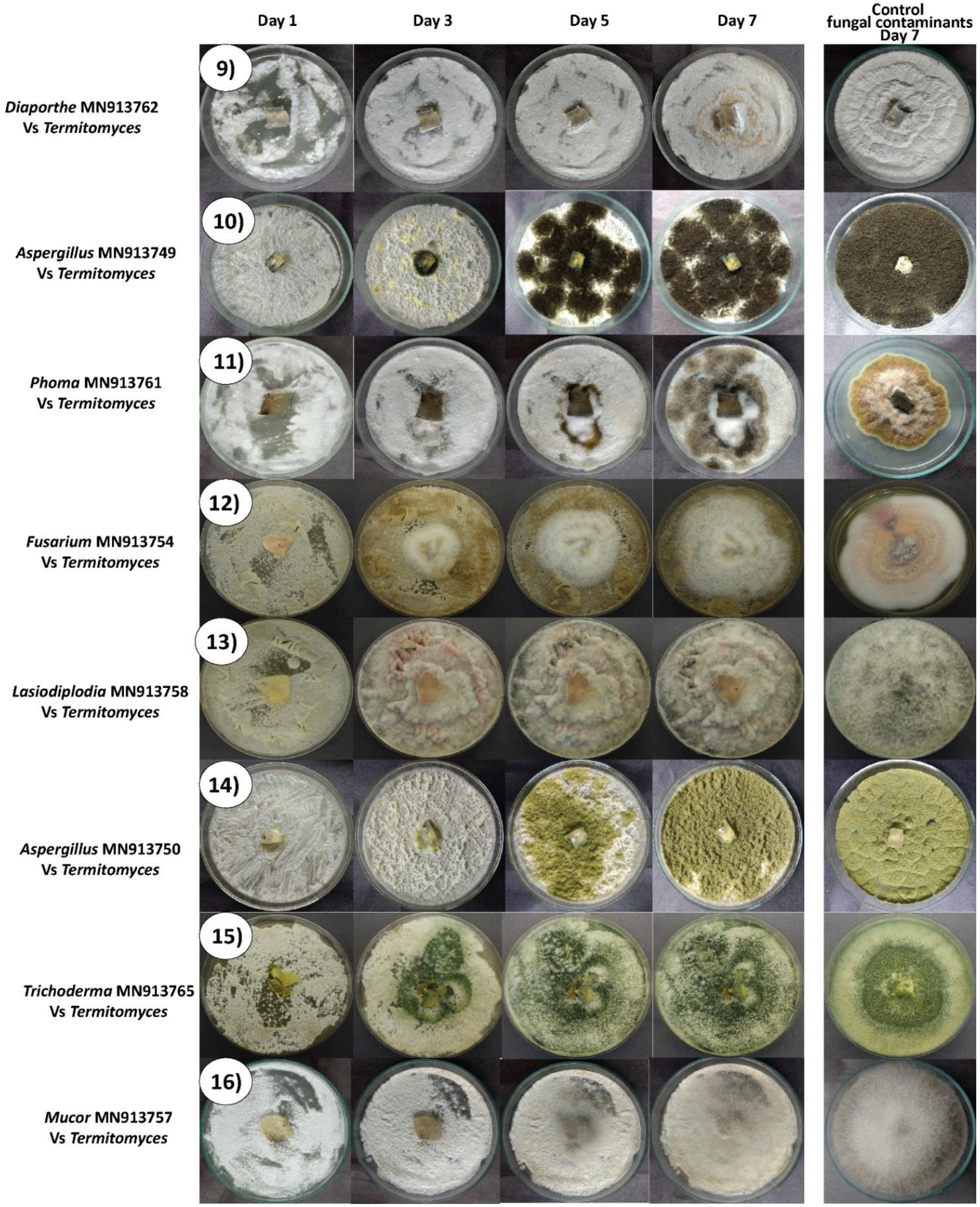

Fig. S13 continued

**Fig. S14: *Pseudoxylaria* Vs Fungal contaminants**

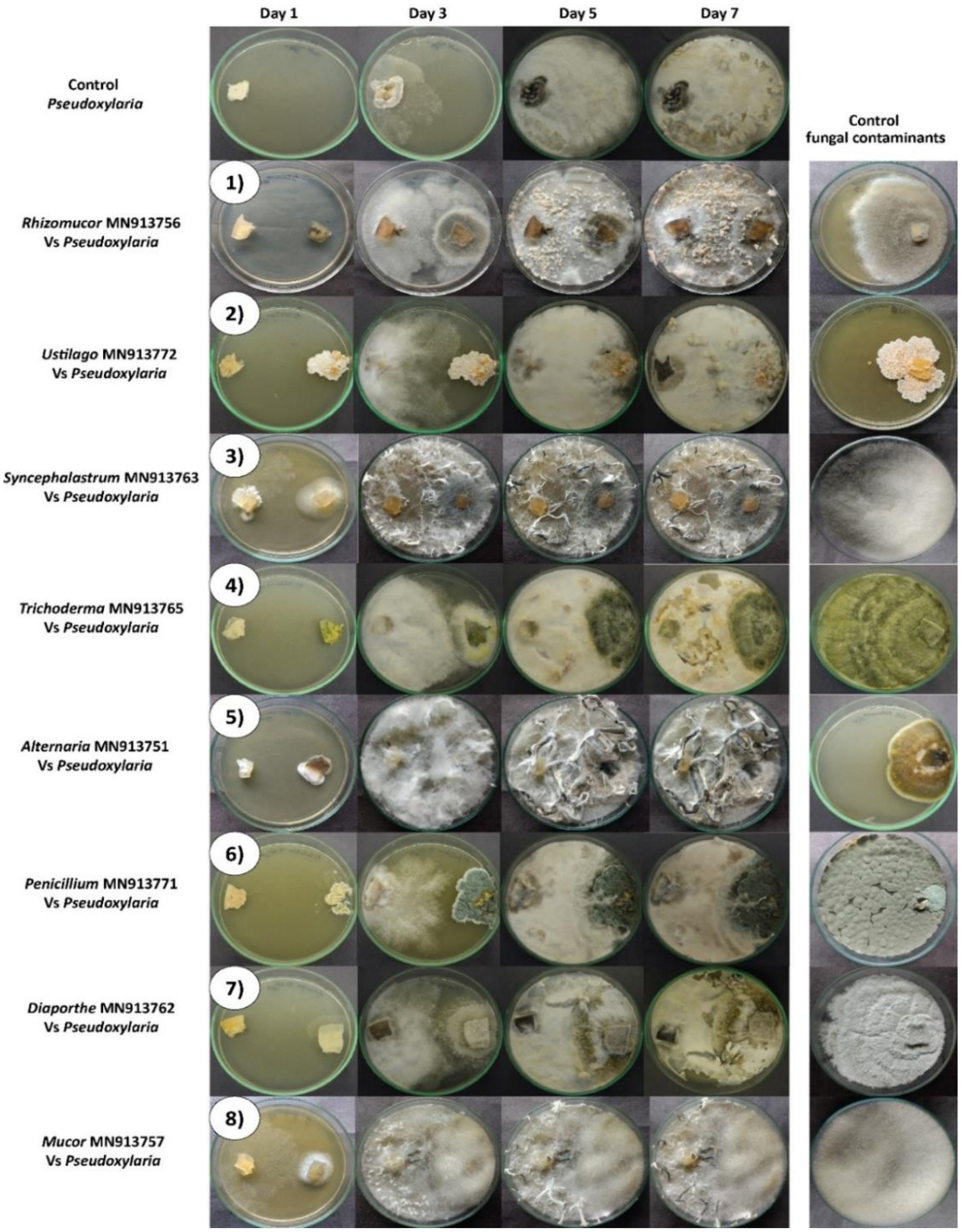

Fig. S14: Interaction assays showing the prevention of the fungal contaminants by *Pseudoxylaria*. All the interactions are categorized into two types: **1)-7) more than 50% inhibition, 8) - 16) less than 50% inhibition**. Each interaction is represented with its fungal and *Pseudoxylaria* control and one representative test plate. More than 50%of inhibition was shown against *Alternaria*-MN913751, *Diaporthe*-MN913762, *Penicillium*-MN913771, *Rhizomucor*-MN913756, *Syncephalastrum*-MN913763, *Trichoderma*-MN913767, and *Ustilago*-MN913772. The other nine fungi (*Aspergillus*-MN91375049, *Aspergillus*-MN913750, *Curvularia*-MN913753, *Fusarium*-MN913754, *Lasiodiplodia*-MN913758, *Mucor*-MN913757, *Paraconiothyrium*-MN913759, *Phialotubus*-MN913760 and *Phoma*-MN913761) were also inhibited by *Pseudoxylaria* but to a lesser degree (less than 50%). All the photos are from day 7 of the beginning of the experiment.

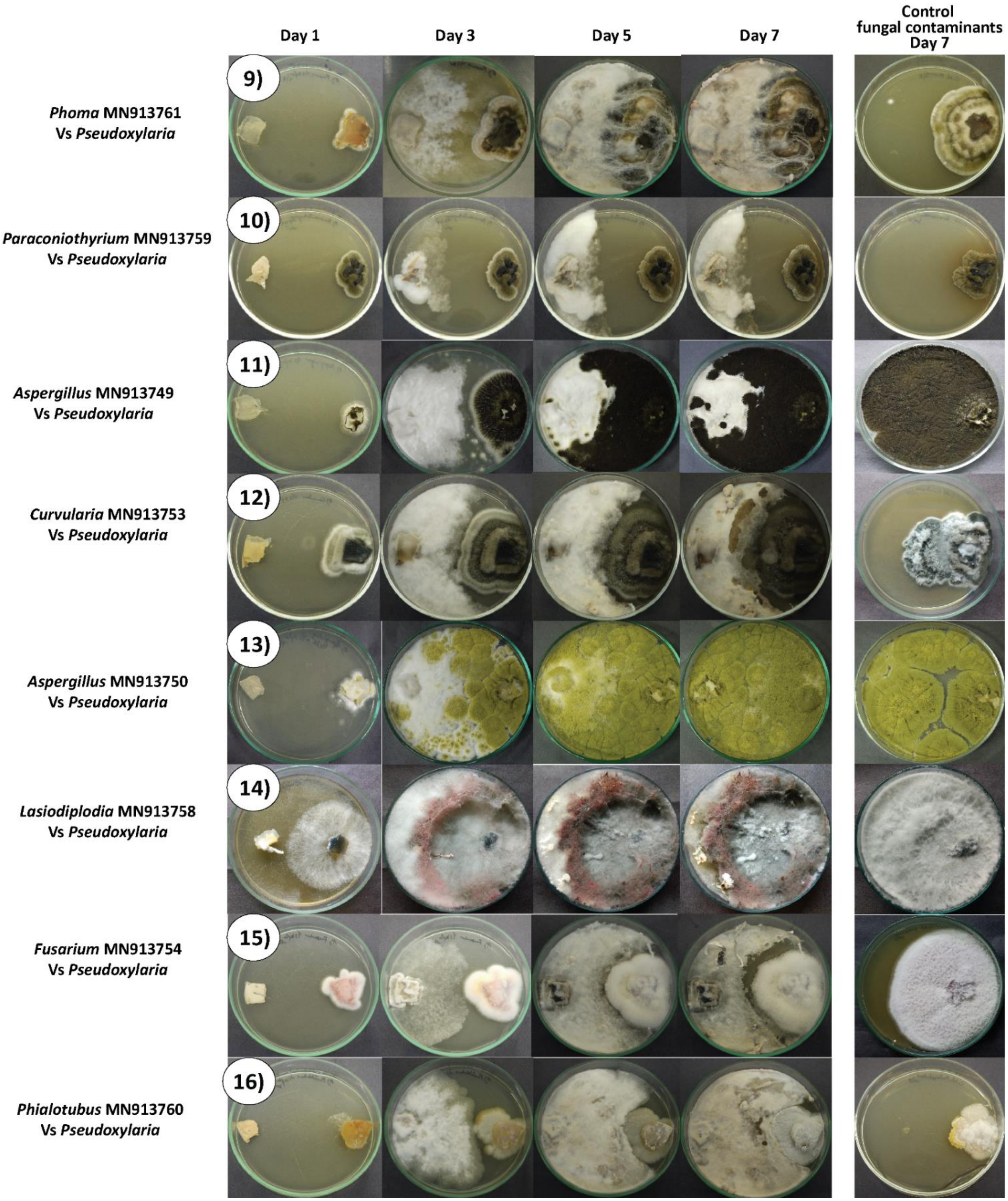

Fig. S14 continued.

**Fig. S15: Re-using media from the test experimental plates**

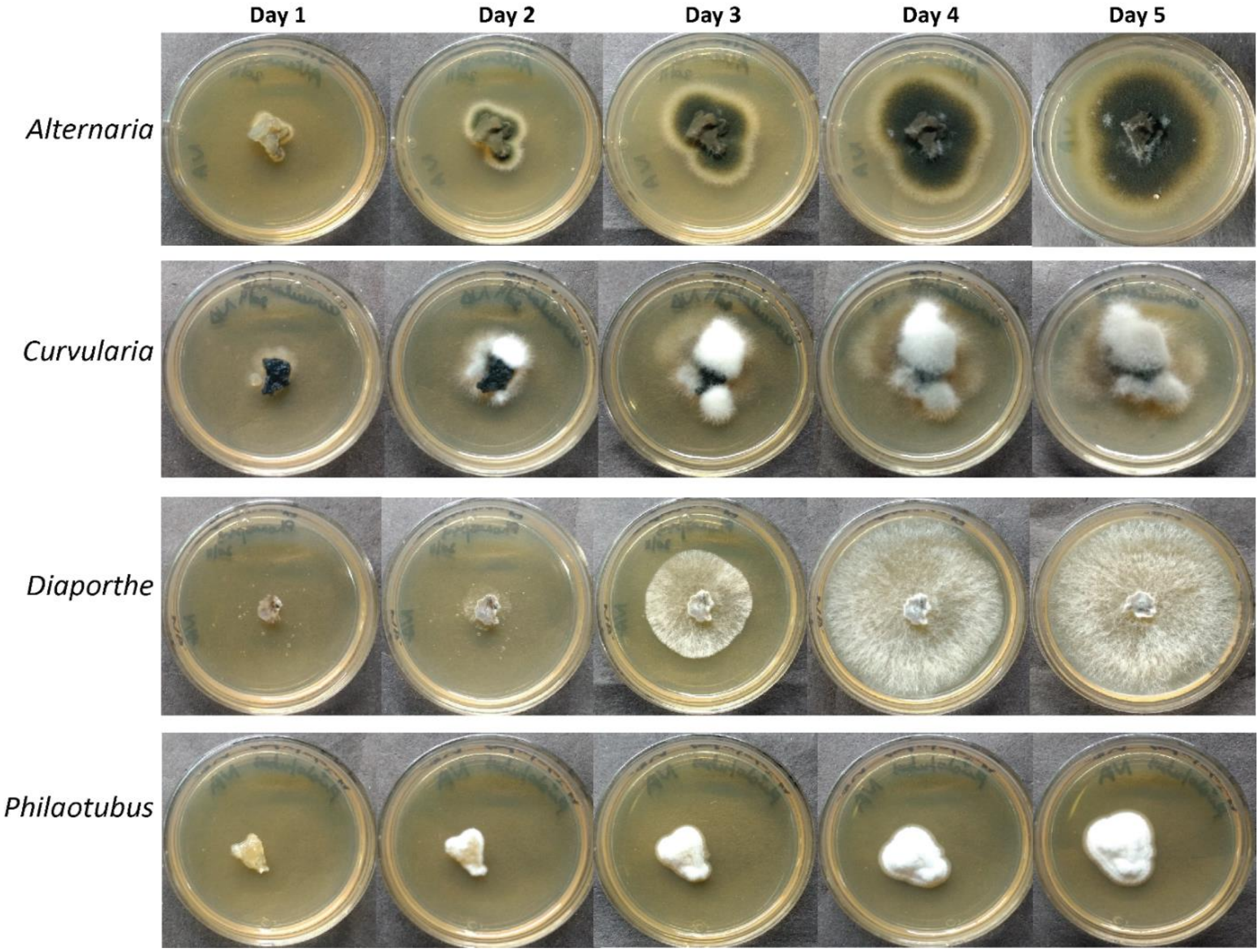

Fig. S15: Representative pictures of few fungi grown on previously used media from the interaction assay (test plates) that were re-plated. A simple explanation for the inhibitory capabilities seen for both *Pseudoxylaria* and *Termitomyces* could be the depletion of nutrients in the PDA plates. However, control experiments using previously used media to grow these fungi indicate the presence of sufficient nutrients in these plates for adequate fungal growth

**Fig. S16: *Bacillus*- MN908297 Vs Fungal contaminants**

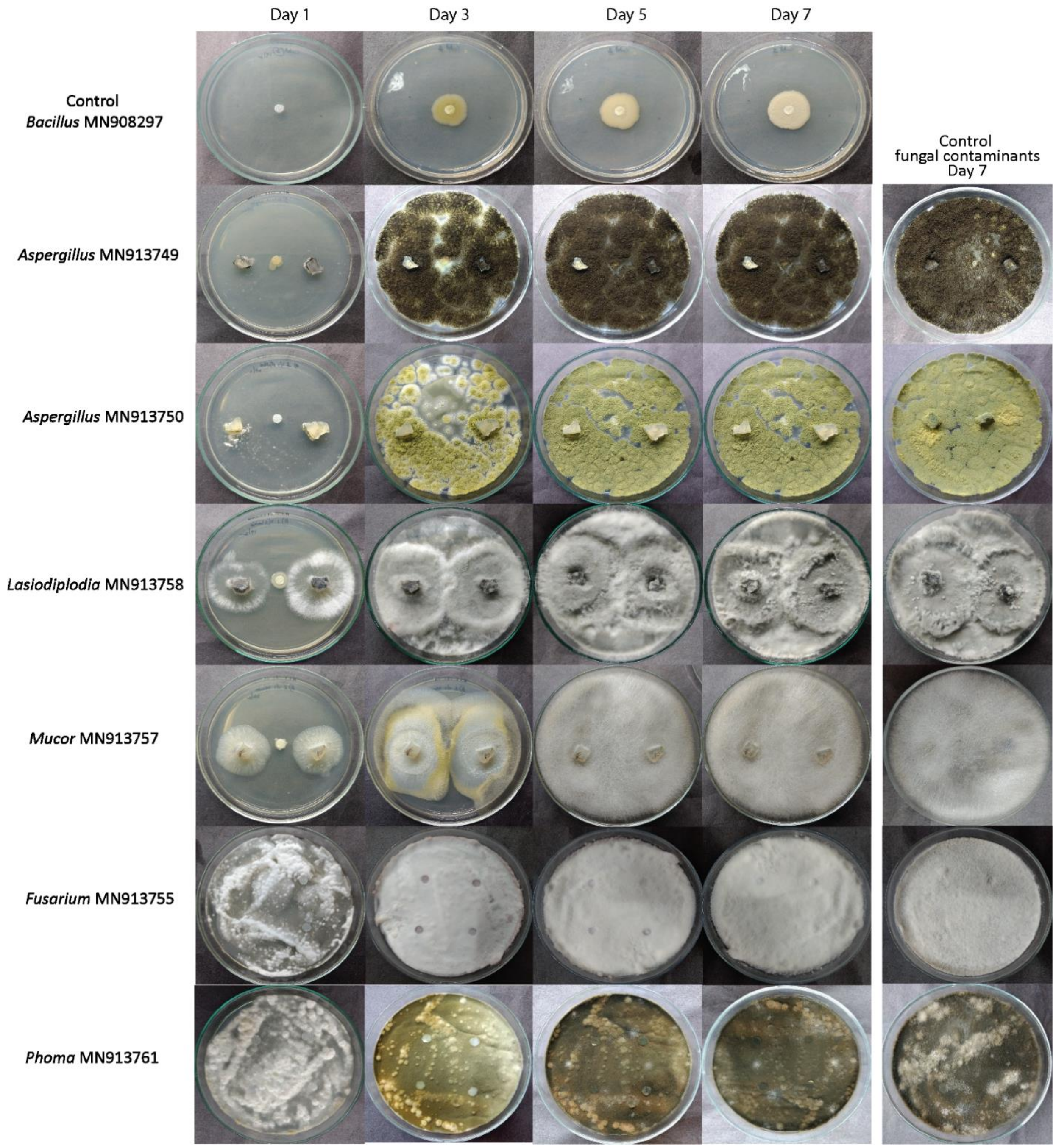

Fig. S16: Assay for the prevention of the growth of fungal contaminants by *Bacillus*- MN908297.

**Fig. S17: *Bacillus*- MN908298 Vs Fungal contaminants**

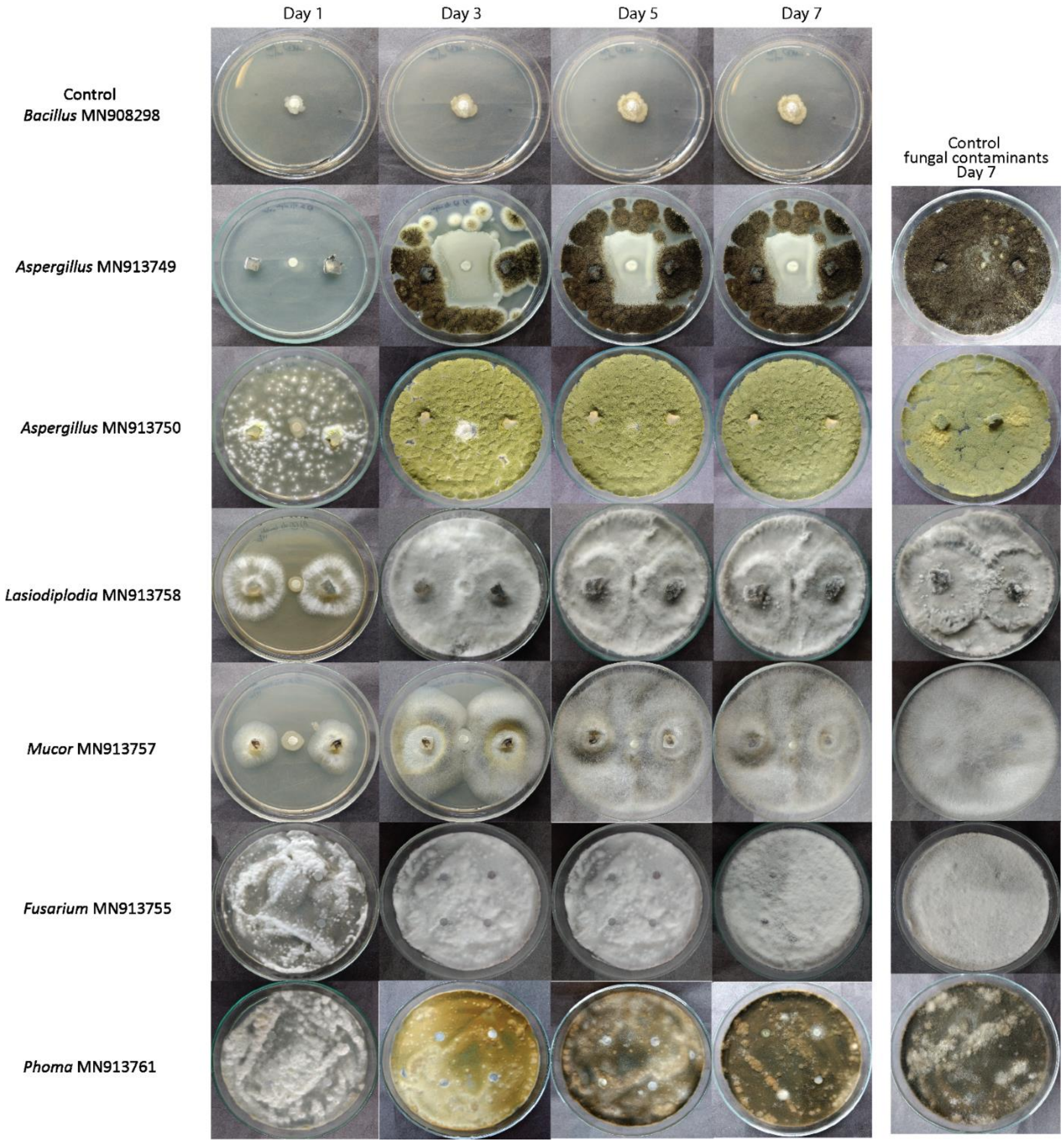

Fig. S17: Assay for the prevention of the growth of fungal contaminants by *Bacillus*- MN908298.

**Fig. S18: *Bacillus*- MN908304 Vs Fungal contaminants**

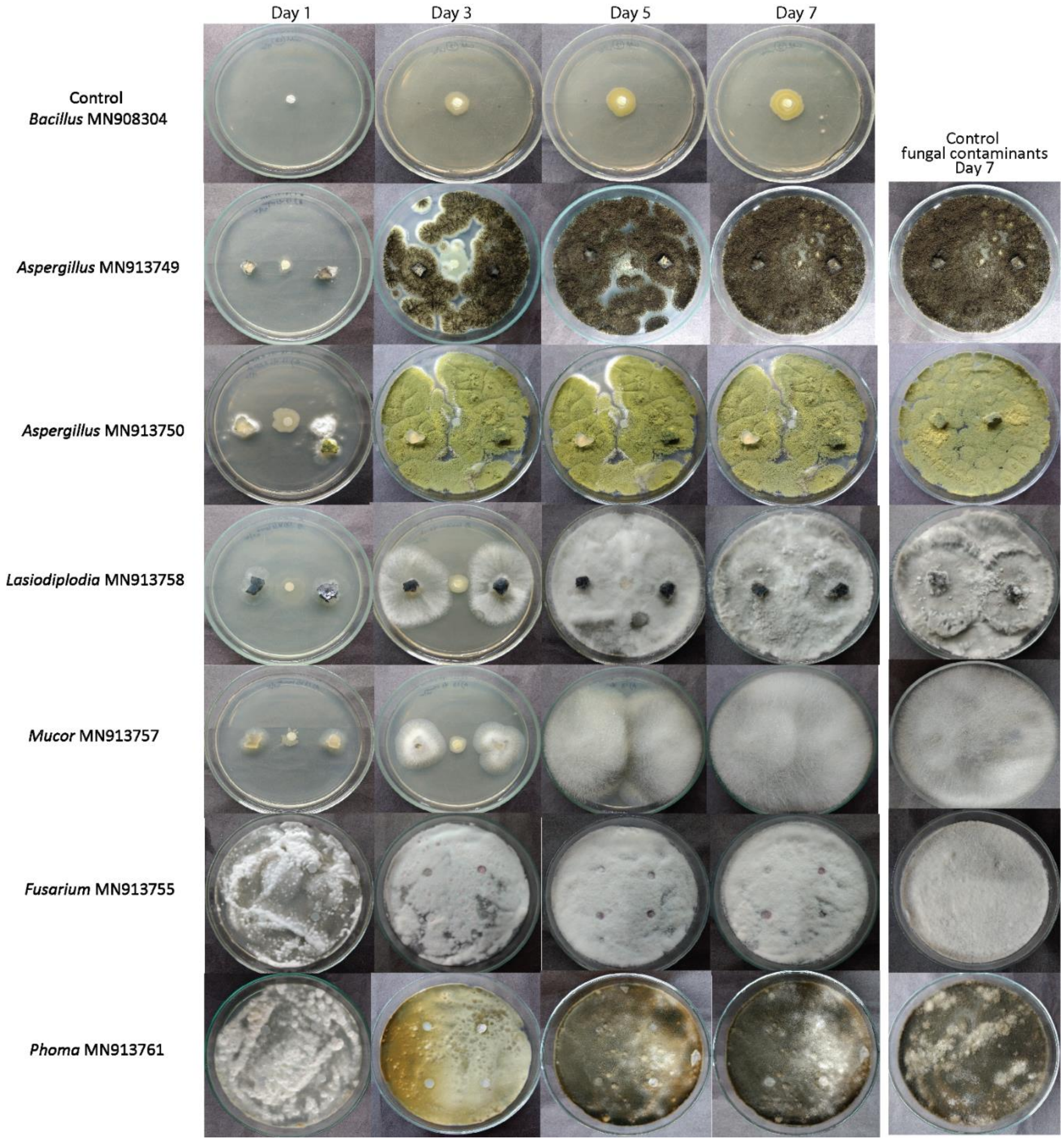

Fig. S18: Assay for the prevention of the growth of fungal contaminants by *Bacillus*- MN908304.

**Fig. S19: *Bacillus*- MN908305 Vs Fungal contaminants**

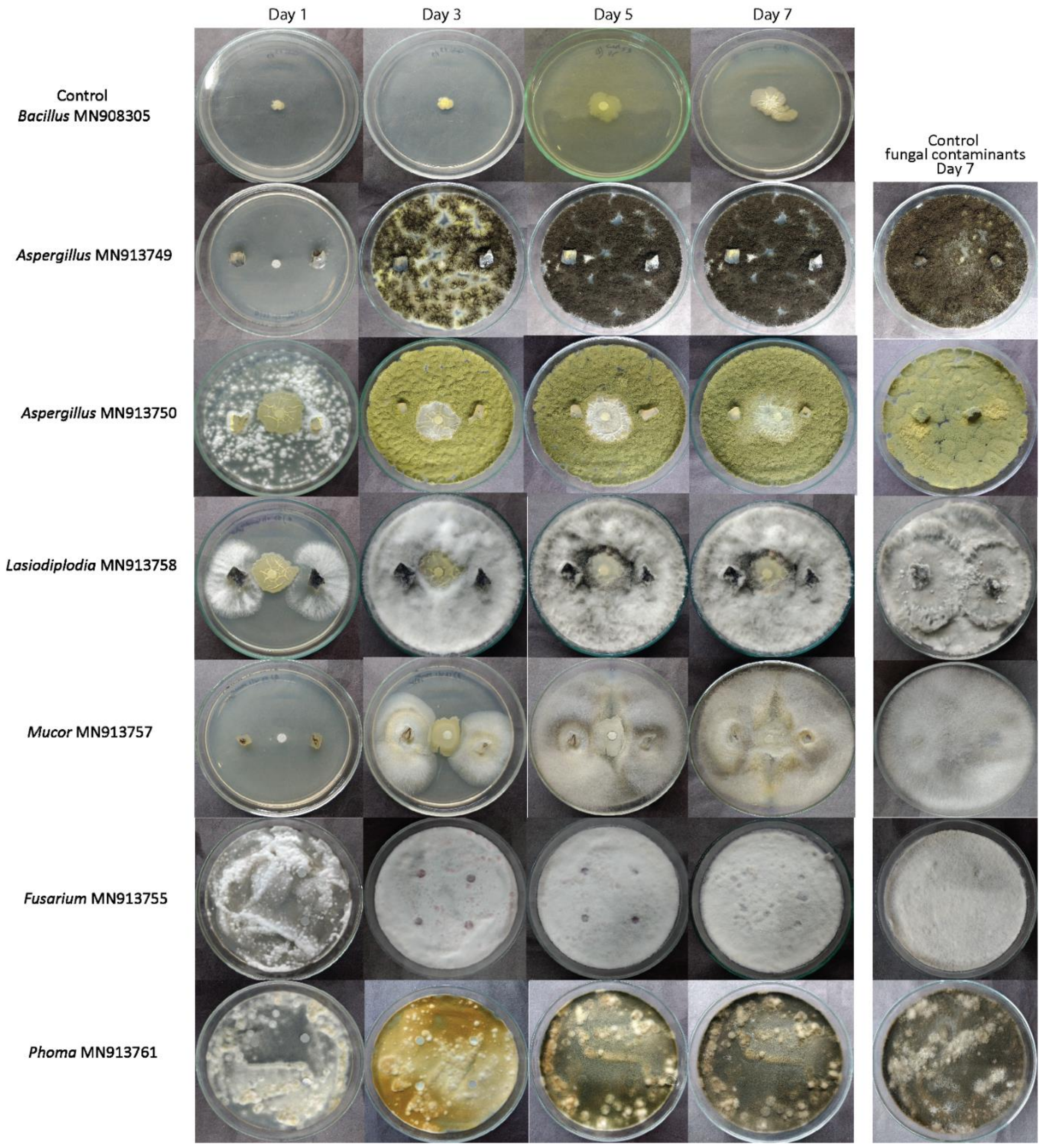

Fig. S19: Assay for the prevention of the growth of fungal contaminants by *Bacillus*- MN908305.

**Fig. S20: *Burkholderia*- MN908310 Vs Fungal contaminants**

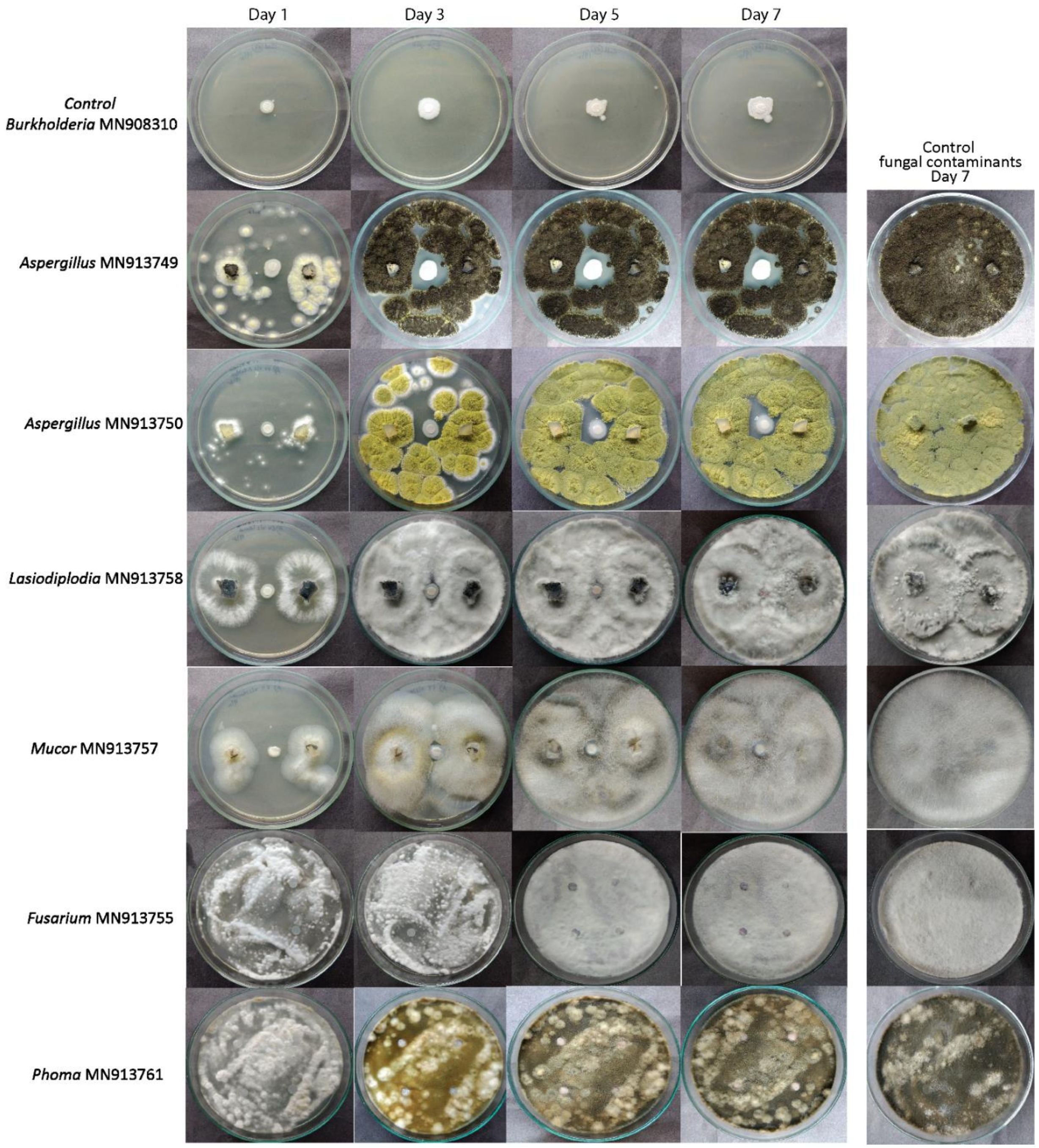

Fig. S20: Assay for the prevention of the growth of fungal contaminants by *Burkholderia*- MN908310.

**Fig. S21: *Pseudomonas*- MN908321 Vs Fungal contaminants**

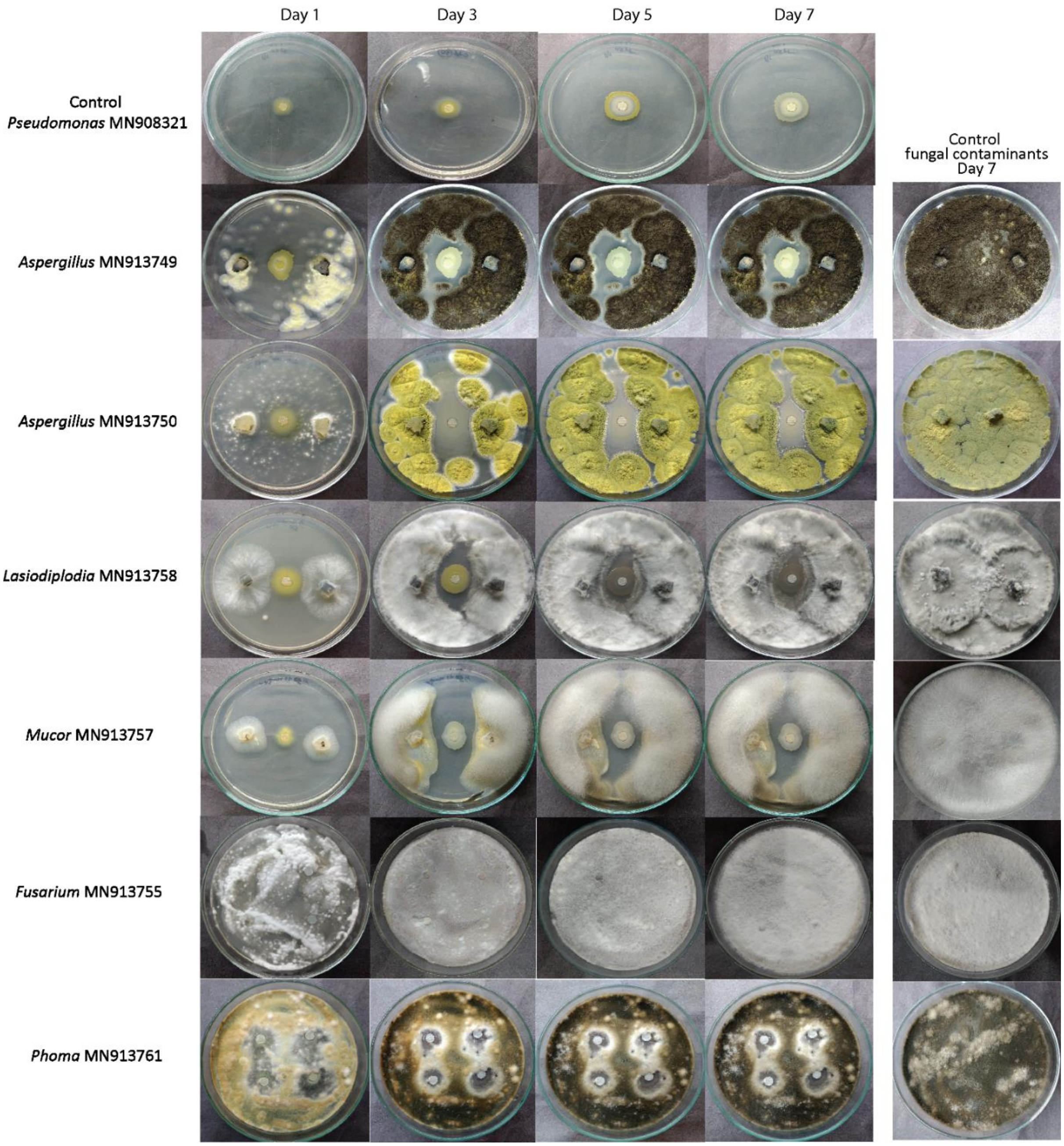

Fig. S21: Assay for the prevention of the growth of fungal contaminants by *Pseudomonas*- MN908321.

**Fig. S22: *Pseudomonas*- MN908322 Vs Fungal contaminants**

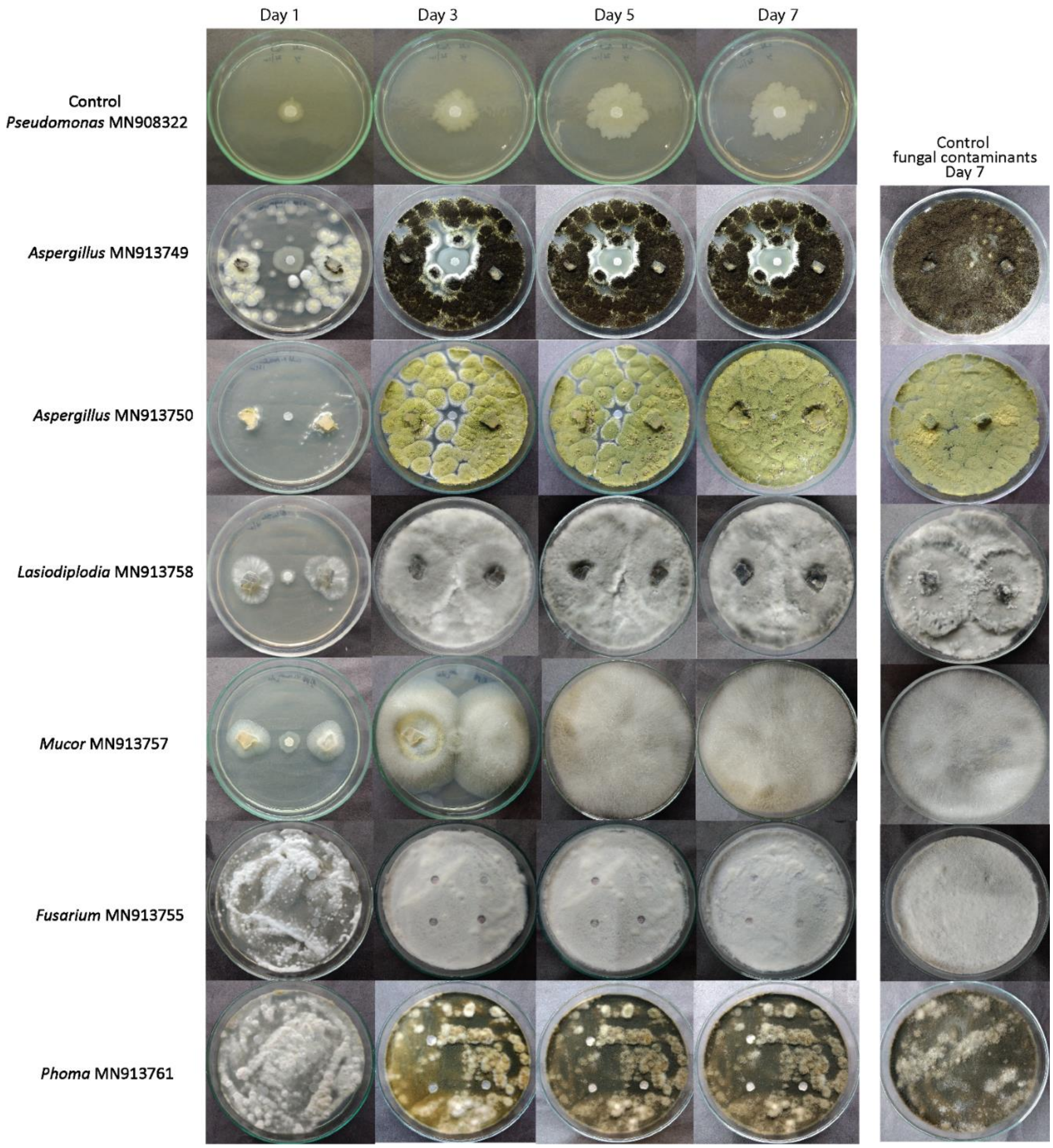

Fig. S22: Assay for the prevention of the growth of fungal contaminants by *Pseudomonas*- MN908322.

**Fig. S23: *Streptomyces*-MN908327 Vs Fungal contaminants**

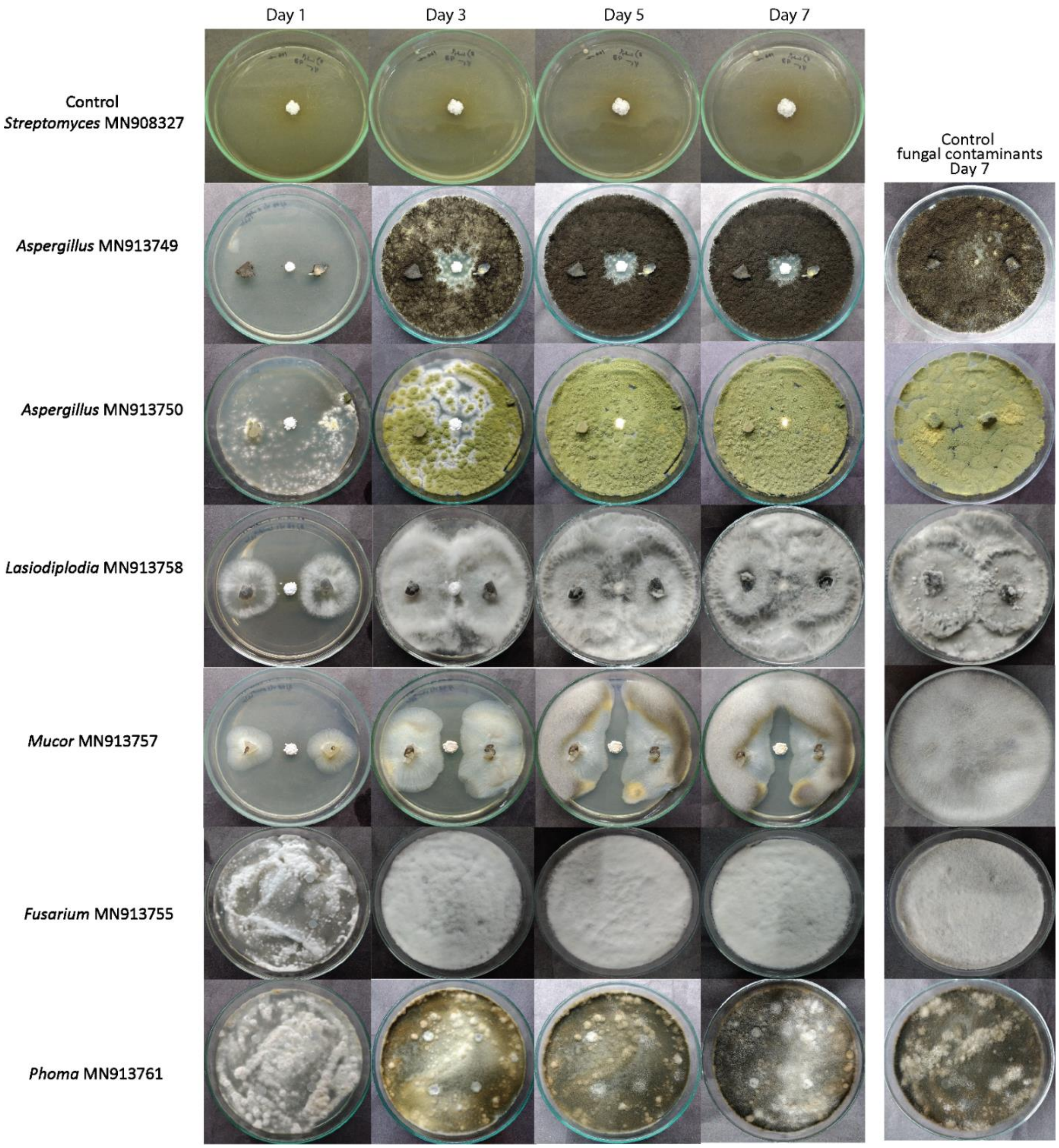

Fig. S23: Assay for the prevention of the growth of fungal contaminants by *Streptomyces*-MN908327.

**Table S1: Details of the fungal strains isolated for the culture-dependent assay.**

| Isolated cultures (Genus-Accession no.) | Isolated from | Nearest neighbor in GenBank | % identity |
|---|---|---|---|
| *Aspergillus* sp. **(MN913749)** | Alate, Comb | KX928746 | 99.66 |
| *Aspergillus* sp. **(MN913750)** | Alate | KU866665 | 100 |
| *Alternaria* sp. **(MN913751)** | Alate | KT192402 | 100 |
| *Curvularia* sp. **(MN913753)** | Alate | MN053866 | 100 |
| *Fusarium* sp. **(MN913755)** | Comb | MW308555 | 100 |
| *Rhizomucor* sp. **(MN913756)** | Alate, Comb | MT254752 | 99.83 |
| *Mucor* sp. **(MN913757)** | Alate, Comb | MN853752 | 99.49 |
| *Lasiodiplodia* sp. **(MN913758)** | Comb | MT302844 | 100 |
| *Paraconiothyrium* sp. **(MN913759)** | Nymph | LT796893 | 100 |
| *Phialotubus* sp. **(MN913760)** | Comb | JN831360 | 100 |
| *Phoma* sp. **(MN913761)** | Alate | MT448809 | 99.42 |
| *Diaporthe* sp. **(MN913762)** | Alate | KU663500 | 99.81 |
| *Syncephalastrum* sp. **(MN913763)** | Alate | MK968099 | 98.95 |
| *Pseudoxylaria* sp. **(MN913764)** | Comb | MK048540 | 100 |
| *Trichoderma* sp. **(MN913767)** | Comb | MT530036 | 100 |
| *Termitomyces* sp. **(MN913768)** | Comb | KR154979 | 100 |
| *Penicillium* sp. **(MN913771)** | Comb | MT588795 | 99.82 |
| *Ustilago* sp. **(MN913772)** | Alate | AY740168 | 99.74 |

Tabe S1: The fungal strains isolated in this study with the *Odontotermes obesus* colony part they were isolated from are listed. Molecular identification of each fungal strain is based on BLASTn results of ITS gene sequences. The list has the accession numbers of the top BLASTn hits for each of their ITS sequences and the percentage identity to these BLAST hits.

**Table S2: The detailed information of termite castes used to study the mycobiota.**

| Samples ID | Biological replicates | Mound 1 | Mound 2 | Total replicates |
|---|---|---|---|---|
| Major worker | MaW1 | + | + | 6 |
| | MaW2 | + | + | |
| | MaW3 | + | + | |
| Minor worker | MiW1 | + | + | 6 |
| | MiW2 | + | + | |
| | MiW3 | + | + | |
| Nymph | Ny1 | + | + | 6 |
| | Ny2 | + | + | |
| | Ny3 | + | + | |
| Male Alate | MA1 | ++ | | 6 |
| | MA2 | ++ | | |
| | MA3 | ++ | | |
| Female Alate | FM1 | ++ | | 6 |
| | FM2 | ++ | | |
| | FM3 | ++ | | |

++ : Two technical replicates

**Table S3: The detailed information of the degrading comb samples used to study the micro- and mycobiota.**

| Microbiota survey | | | | | | | |
|---|---|---|---|---|---|---|---|
| Comb Samples | Mound 1 | | | Mound 2 | | | Total Replicates/day |
| | August | September | October | August | September | October | |
| O hrs | + | + | + | + | + | + | 6 |
| 24 hrs | + | + | + | + | + | + | 6 |
| 48 hrs | + | + | + | + | + | + | 6 |
| 72 hrs | + | + | + | + | + | + | 6 |
| 96 hrs | + | + | + | + | + | + | 6 |
| 120 hrs | + | + | + | + | + | + | 6 |
| **Mycobiota survey** | | | | | | | |
| Comb Samples | Mound 1 | | | Mound 2 | | | Total Replicates/day |
| | August | September | October | August | September | October | |
| O hrs | + | + | + | + | + | + | 6 |
| 24 hrs | + | + | + | + | + | + | 6 |
| 48 hrs | + | + | + | + | + | + | 6 |
| 72 hrs | + | + | + | + | + | + | 6 |
| 96 hrs | + | + | + | + | + | + | 6 |
| 120 hrs | + | + | + | + | + | + | 6 |

**Table S4: List of primers used in this study**

| Primer name | Function | Samples amplified | Target | Sequence (5′→3′) | Annealing Temperature (°C) |
|---|---|---|---|---|---|
| Termito_F1 | Forward | Pure culture | *Termitomyces* | CATCGTTTTCAACCACCTGTGC | 60 |
| Termito_R1 | Reverse | | | GCCTATCCAAGCTCACAAAAGC | |
| Xyla_F1 | Forward | Pure culture | *Pseudoxylaria* | GGCCCTGAAAACTCTGTTTCG | 56 |
| Xyla_R1 | Reverse | | | GCCCAGCTGCAGACGAGAATA | |
| Pseudo_F1 | Forward | Pure culture | *Pseudomonas* | GAGTAATGCCTAGGAATCTGC | 56 |
| Pseudo_R1 | Reverse | | | CAGTATCAGTCCAGGTGGTCG | |
| qTermito_F | Forward | Fungus comb | *Termitomyces* | TCAACCACCTGTGCACCTTTTG | 51 |
| qTermito_R | Reverse | | | GCATAGACCGGAAATGCAG | |
| qXyla_F | Forward | Fungus comb | *Pseudoxylaria* | TCGTTGCTTAGCGTTGGGAGC | 61 |
| qXyla_R | Reverse | | | CTAGAGCGTGAAACCGACTC | |
| qPseudo_F1 | Forward | Fungus comb | *Pseudomonas* | TCAACCTGGGAACTGCATCCA | 60 |
| Pse686R[11] | Reverse | | | ACACAGGAAATTCCACCACCC | |

**Table S5: The mycobiota degrading fungus comb of *O. obesus*. The table also shows different diversity indices for all the samples.**

| Sample source | Total read (initial) | Total reads (post-cleanup) | Reads identified | Fungal genera identified | Chao1 | Shannon | Simpson | InvSimspon |
|---|---|---|---|---|---|---|---|---|
| **Decaying comb (0 hrs)** | $0.58x10^5$ | $0.54x10^5$ | $0.54x10^5$ | 111 | 199.3 | 0.53 | 0.20 | 1.25 |
| **Decaying comb (24 hrs)** | $0.62x10^5$ | $0.60x10^5$ | $0.58x10^5$ | 83 | 116 | 0.47 | 0.17 | 1.20 |
| **Decaying comb (48 hrs)** | $0.16x10^5$ | $0.18x10^5$ | $0.14x10^5$ | 103 | 157.4 | 1.67 | 0.75 | 4.04 |
| **Decaying comb (72 hrs)** | $0.16x10^5$ | $0.17x10^5$ | $0.14x10^5$ | 110 | 152.5 | 1.97 | 0.80 | 5.24 |
| **Decaying comb (96 hrs)** | $0.17x10^5$ | $0.18x10^5$ | $0.15x10^5$ | 99 | 184 | 1.54 | 0.70 | 3.43 |
| **Decaying comb (120 hrs)** | $0.04x10^5$ | $0.04x10^5$ | $0.04x10^5$ | 71 | 126.1 | 1.47 | 0.63 | 2.73 |
| | | | | 236 unique fungal genera | | | | |

**Table S6: Number of copies of *Termitomyces, Pseudoxylaria and Pseudomonas* in different degrading stages of fungus comb using qPCR estimation. The highlighted cells are number of copies within 95% Confidence intervals.**

| | | No. of copies/ ul | | |
|---|---|---|---|---|
| | Samples | *Termitomyces* | *Pseudoxylaria* | *Pseudomonas* |
| 0 hrs | Comb_1 | 1360107542 | 89981 | 175388 |
| | Comb_2 | 1527079637 | 261070 | 120571 |
| | Comb_3 | 3010663121 | 1662542 | 109453 |
| | Comb_4 | 29961793 | 480150 | 16482 |
| | Comb_5 | 1379035572 | 550258 | 12115 |
| | Comb_6 | 66598339 | 965224 | 454345 |
| | Comb_7 | 3076097533 | 10680 | 5262 |
| | Comb_8 | 573685238 | 106523 | 868021 |
| | Comb_9 | 703381182 | 200563 | 334247 |
| | Comb_10 | 408063859 | 198522 | 1016288 |
| | Comb_11 | 3019443861 | N/A | 6140510 |
| 24 hrs | Comb_1 | 1439414705 | 9648600 | 6059579 |
| | Comb_2 | 533834327 | 2797119 | 64542 |
| | Comb_3 | 2337609707 | 3912787 | 210228 |
| | Comb_4 | 682213164 | 5980566 | 113264 |
| | Comb_5 | 1456291250 | 8215710 | 165481 |
| | Comb_6 | 44139049 | 7859628 | 1527 |
| | Comb_7 | 984861560 | 2525442 | 3716 |
| | Comb_8 | 1210550436 | 12659094 | 13237 |
| | Comb_9 | 991863875 | 13882737 | 232467 |
| | Comb_10 | 1516168056 | 8318347 | 218101 |
| | Comb_11 | 1224549561 | 38050364 | 69040 |
| | Comb_12 | 1605096212 | 27805362 | 1331129 |
| 48 hrs | Comb_1 | 2775237 | 933660096 | 69281 |
| | Comb_2 | 695691 | 752072080 | 76725 |
| | Comb_3 | 9064272 | 462970 | 58965 |
| | Comb_4 | 8440967 | 180809717 | 73434 |
| | Comb_5 | 195786877 | 23597478 | 12983 |
| | Comb_6 | 13193507 | 3003351792 | 43232 |
| | Comb_7 | 3787981 | 1824323516 | 697610 |
| | Comb_8 | 137221322 | 892522016 | 268384 |
| | Comb_9 | 11569396 | 185587389 | 986642 |
| | Comb_10 | 30363382 | 2047154667 | 264786 |
| | Comb_11 | 5539178 | 99217932 | 110009 |
| | Comb_12 | 3968439 | N/A | 742690 |

| | | | | |
|---|---|---|---|---|
| 72 hrs | Comb_1 | 735628 | 17242698 | 68775 |
| | Comb_2 | 1252119357 | 1083222891 | 445587 |
| | Comb_3 | 249477 | 7752681 | 149178 |
| | Comb_4 | 1504145 | 201581 | 56405 |
| | Comb_5 | 1874901 | 445655 | 6115877 |
| | Comb_6 | 75413343 | 2824057 | 11582345 |
| | Comb_7 | 5363423 | 4306131 | 27078 |
| | Comb_8 | 462079025 | 8967365 | 14130 |
| | Comb_9 | 37516583 | 358315851 | 3659116 |
| | Comb_10 | 4398251 | 2942430 | 467746 |
| | Comb_11 | 551510 | 130507319 | 135188 |
| | Comb_12 | 276314 | 135926460 | 11997329 |
| 96 hrs | Comb_1 | 28614 | 3940039 | 50105 |
| | Comb_2 | 85917 | 54880292 | 3873646 |
| | Comb_3 | 2703112 | 116695 | 53624 |
| | Comb_4 | 1430558 | 121515648 | 183775 |
| | Comb_5 | 583616597 | 945957 | 57043 |
| | Comb_6 | 1996 | 965994 | 17790 |
| | Comb_7 | 197580 | 152511231 | 24126 |
| | Comb_8 | 158590 | 13736078 | 8492 |
| | Comb_9 | 346307 | 31014431 | 88413 |
| | Comb_10 | 1200402 | 5246721 | 22053078 |
| | Comb_11 | 137577 | 287839668 | 675092 |
| | Comb_12 | 182311 | | 173260 |
| 120 hrs | Comb_1 | 24967 | 3555896 | 113598 |
| | Comb_2 | 75815 | 9409891 | 355757 |
| | Comb_3 | 650499 | 559521 | 117656 |
| | Comb_4 | 90228 | 15718962 | 32026 |
| | Comb_5 | 632982 | 658760 | 85201 |
| | Comb_6 | 1440370 | 5423933 | 1139999 |
| | Comb_7 | 294558 | 10737477 | 139542 |
| | Comb_8 | 2440949 | 12765611 | 61357038 |
| | Comb_9 | 194647 | 1416038 | 164270 |
| | Comb_10 | 582163 | 1182903 | 175860 |
| | Comb_11 | 942411 | 26059452 | 153213 |
| | Comb_12 | 488913 | N/A | 151121 |

**Table S6 continued**

**Table S7: The microbiota of the degrading fungus comb of *O. obesus*. The table also shows different diversity indices for all the samples.**

| Sample source | Total reads (initial) | Total reads (post-cleanup) | Reads identified | Bacterial genera identified | Chao1 | Shannon | Simpson | InvSimspon |
|---|---|---|---|---|---|---|---|---|
| **Decaying comb (0 hrs)** | $0.14x10^5$ | $0.12x10^5$ | $0.12x10^5$ | 587 | 587 | 4.4 | 0.9 | 32.0 |
| **Decaying comb (24 hrs)** | $0.94x10^5$ | $0.88x10^5$ | $0.16x10^5$ | 390 | 390 | 3.7 | 0.9 | 19.9 |
| **Decaying comb (48 hrs)** | $0.38x10^5$ | $0.50x10^5$ | $0.36x10^5$ | 2014 | 2014 | 6.0 | 0.9 | 137.4 |
| **Decaying comb (72 hrs)** | $1.54x10^5$ | $1.45x10^5$ | $0.16x10^5$ | 374 | 374 | 3.7 | 0.9 | 22.7 |
| **Decaying comb (96 hrs)** | $0.63x10^5$ | $0.59x10^5$ | $0.20x10^5$ | 298 | 298 | 3.3 | 0.9 | 13.9 |
| **Decaying comb (120 hrs)** | $1.33x10^5$ | $1.25x10^5$ | $0.16x10^5$ | 278 | 278 | 3.4 | 0.9 | 15.4 |
| | | | | 2, 182 unique bacterial genera | | | | |

**Table S8: Relative abundances of fungal OTUs in different castes and degrading fungus comb obtained from Nanopore sequencing.**

| Phylum | **Class** | **Order** | **Family** | **Genera** | **Major Worker** | **Minor Worker** | **Nymph** | **Female Alate** | **Male Alate** | **0 hrs** | **24 hrs** | **48 hrs** | **72 hrs** | **96 hrs** | **120 hrs** |
|---|---|---|---|---|---|---|---|---|---|---|---|---|---|---|---|
| Basidiomycota | Agaricomycetes | Polyporales | Podoscyphaceae | Abortiporus_biennis | 0.000 | 0.000 | 0.000 | 0.001 | 0.000 | 0.000 | 0.000 | 0.000 | 0.000 | 0.000 | 0.000 |
| Basidiomycota | Exobasidiomycetes | Exobasidiales | Cryptobasidiaceae | Acaromyces_sp. | 0.001 | 0.000 | 0.000 | 0.000 | 0.000 | 0.000 | 0.000 | 0.000 | 0.000 | 0.000 | 0.000 |
| Mucoromycota | Glomeromycetes | Diversisporales | Acaulosporaceae | Acaulospora_cavernata | 0.001 | 0.000 | 0.000 | 0.000 | 0.000 | 0.000 | 0.000 | 0.000 | 0.000 | 0.000 | 0.000 |
| Mucoromycota | Glomeromycetes | Diversisporales | Acaulosporaceae | Acaulospora_colossica | 0.005 | 0.000 | 0.000 | 0.000 | 0.000 | 0.000 | 0.000 | 0.000 | 0.000 | 0.000 | 0.000 |
| Mucoromycota | Glomeromycetes | Diversisporales | Acaulosporaceae | Acaulospora_excavata | 0.005 | 0.025 | 0.000 | 0.000 | 0.000 | 0.000 | 0.000 | 0.000 | 0.000 | 0.000 | 0.000 |
| Mucoromycota | Glomeromycetes | Diversisporales | Acaulosporaceae | Acaulospora_nivalis | 0.210 | 0.050 | 0.000 | 0.001 | 0.000 | 0.000 | 0.000 | 0.000 | 0.000 | 0.000 | 0.000 |
| Mucoromycota | Glomeromycetes | Diversisporales | Acaulosporaceae | Acaulospora_scrobiculata | 0.030 | 0.000 | 0.000 | 0.000 | 0.000 | 0.000 | 0.000 | 0.000 | 0.000 | 0.000 | 0.000 |
| Mucoromycota | Glomeromycetes | Diversisporales | Acaulosporaceae | Acaulospora_sp. | 0.004 | 0.000 | 0.000 | 0.000 | 0.000 | 0.000 | 0.000 | 0.000 | 0.000 | 0.000 | 0.000 |
| Mucoromycota | Glomeromycetes | Diversisporales | Acaulosporaceae | Acaulospora_spinosissima | 0.014 | 0.000 | 0.000 | 0.000 | 0.000 | 0.000 | 0.000 | 0.000 | 0.000 | 0.000 | 0.000 |
| Ascomycota | Sordariomycetes | Hypocreales | Unclassified | Acremonium_exuviarum | 0.000 | 0.000 | 0.000 | 0.000 | 0.000 | 0 | 0 | 0 | 0 | 0 | 0.02 |
| Ascomycota | Sordariomycetes | Hypocreales | Unclassified | Acremonium_sclerotigenum | 0.000 | 0.000 | 0.000 | 0.000 | 0.005 | 0.000 | 0.000 | 0.000 | 0.000 | 0.000 | 0.000 |
| Ascomycota | Sordariomycetes | Hypocreales | Unclassified | Acremonium_sp. | 0.000 | 0.025 | 0.006 | 0.000 | 0.000 | 0.0071 | 0 | 0.0943 | 0.0116 | 0.0055 | 0 |
| Ascomycota | Dothideomycetes | Pleosporales | Morosphaeriaceae | Acrocalymma_cycadis | 0.000 | 0.000 | 0.000 | 0.000 | 0.000 | 0.0072 | 0 | 0 | 0 | 0 | 0 |
| Ascomycota | Sordariomycetes | Sordariales | Chaetomiaceae | Acrophialophora_fusispora | 0.000 | 0.000 | 0.000 | 0.000 | 0.000 | 0 | 0.0016 | 0 | 0 | 0.0055 | 0 |
| Ascomycota | Sordariomycetes | Sordariales | Chaetomiaceae | Acrophialophora_levis | 0.000 | 0.000 | 0.000 | 0.000 | 0.000 | 0 | 0 | 0 | 0 | 0.0055 | 0 |
| Basidiomycota | Agaricomycetes | Agaricales | Agaricaceae | Agaricus_sp. | 0.001 | 0.000 | 0.000 | 0.000 | 0.000 | 0.0071 | 0.0082 | 0.0055 | 0.0116 | 0.0055 | 0.08 |
| Ascomycota | Eurotiomycetes | Onygenales | Ajellomycetaceae | Ajellomyces_capsulatus | 0.001 | 0.000 | 0.000 | 0.000 | 0.000 | 0.000 | 0.000 | 0.000 | 0.000 | 0.000 | 0.000 |
| Ascomycota | Sordariomycetes | Hypocreales | Stachybotryaceae | Albifimbria_verrucaria | 0.000 | 0.000 | 0.000 | 0.000 | 0.010 | 0 | 0.0016 | 0.0055 | 0.0058 | 0 | 0 |
| Ascomycota | Saccharomycetes | Saccharomycetales | Alloascoideaceae | Alloascoidea_hylecoeti | 0.001 | 0.000 | 0.000 | 0.000 | 0.000 | 0.000 | 0.000 | 0.000 | 0.000 | 0.000 | 0.000 |
| Ascomycota | Dothiodeomycetes | Pleosporales | Pleosporaceae | Alternaria_alternata | 0.009 | 0.050 | 0.051 | 0.036 | 0.370 | 0.0035 | 0 | 0 | 0 | 0 | 0 |
| Ascomycota | Dothiodeomycetes | Pleosporales | Pleosporaceae | Alternaria_brassicae | 0.000 | 0.000 | 0.000 | 0.001 | 0.005 | 0.000 | 0.000 | 0.000 | 0.000 | 0.000 | 0.000 |
| Ascomycota | Dothiodeomycetes | Pleosporales | Pleosporaceae | Alternaria_infectoria | 0.000 | 0.000 | 0.006 | 0.000 | 0.000 | 0.000 | 0.000 | 0.000 | 0.000 | 0.000 | 0.000 |
| Ascomycota | Dothiodeomycetes | Pleosporales | Pleosporaceae | Alternaria_sp. | 0.004 | 0.050 | 0.028 | 0.017 | 0.165 | 0.0143 | 0 | 0 | 0 | 0 | 0 |
| Ascomycota | Dothiodeomycetes | Pleosporales | Pleosporaceae | Alternaria_tenuissima | 0.001 | 0.000 | 0.000 | 0.000 | 0.010 | 0.0035 | 0 | 0 | 0 | 0 | 0 |
| Basidiomycota | Agaricomycetes | Agaricales | Amanitaceae | Amanita_fulva | 0.004 | 0.000 | 0.000 | 0.001 | 0.000 | 0.0035 | 0.0082 | 0 | 0 | 0 | 0 |
| Basidiomycota | Agaricomycetes | Agaricales | Amanitaceae | Amanita_melleiceps | 0.001 | 0.000 | 0.000 | 0.000 | 0.000 | 0 | 0.0016 | 0 | 0 | 0 | 0 |
| Basidiomycota | Agaricomycetes | Agaricales | Amanitaceae | Amanita_muscaria | 0.000 | 0.000 | 0.000 | 0.001 | 0.000 | 0.000 | 0.000 | 0.000 | 0.000 | 0.000 | 0.000 |

**Table S8 continued.**

| | | | | | | | | | | | | | | | |
|---|---|---|---|---|---|---|---|---|---|---|---|---|---|---|---|
| Basidiomycota | Agaricomycetes | Agaricales | Amanitaceae | Amanita_pantherina | 0.000 | 0.000 | 0.000 | 0.000 | 0.000 | 0 | 0.0049 | 0 | 0 | 0 | 0 |
| Basidiomycota | Agaricomycetes | Agaricales | Amanitaceae | Amanita_phalloides | 0.000 | 0.000 | 0.006 | 0.000 | 0.000 | 0.000 | 0.000 | 0.000 | 0.000 | 0.000 | 0.000 |
| Basidiomycota | Agaricomycetes | Agaricales | Amanitaceae | Amanita_pseudoporphyria | 0.001 | 0.000 | 0.000 | 0.000 | 0.000 | 0.0072 | 0 | 0 | 0 | 0 | 0 |
| Basidiomycota | Agaricomycetes | Agaricales | Amanitaceae | Amanita_velosa | 0.000 | 0.000 | 0.000 | 0.000 | 0.000 | 0 | 0.0016 | 0 | 0 | 0 | 0 |
| Basidiomycota | Agaricomycetes | Agaricales | Amanitaceae | Amanita_verna | 0.000 | 0.000 | 0.000 | 0.000 | 0.000 | 0 | 0.0016 | 0 | 0 | 0 | 0 |
| Ascomycota | Eurotiomycetes | Onygenales | Onygenaceae | Amauroascus_albicans | 0.000 | 0.000 | 0.000 | 0.000 | 0.000 | 0.000 | 0.000 | 0.000 | 0.000 | 0.000 | 0.000 |
| Mucoromycota | Glomeromycetes | Archaeosporales | Ambisporaceae | Ambispora_sp. | 0.000 | 0.000 | 0.000 | 0.000 | 0.000 | 0.000 | 0.000 | 0.000 | 0.000 | 0.000 | 0.000 |
| Basidiomycota | Agaricomycetes | Atheliales | Atheliaceae | Amphinema_byssoides | 0.000 | 0.000 | 0.006 | 0.000 | 0.000 | 0 | 0.0016 | 0 | 0 | 0 | 0 |
| Ascomycota | Sordariomycetes | Xylariales | Hypoxylaceae | Annulohypoxylon_sp. | 0.000 | 0.000 | 0.000 | 0.000 | 0.000 | 0 | 0 | 0.0055 | 0 | 0 | 0 |
| Ascomycota | Sordariomycetes | Xylariales | Hypoxylaceae | Annulohypoxylon_stygium | 0.000 | 0.000 | 0.000 | 0.000 | 0.000 | 0 | 0 | 0 | 0 | 0.0055 | 0 |
| Ascomycota | Sordariomycetes | Xylariales | Hypoxylaceae | Annulohypoxylon_truncatum | 0.000 | 0.000 | 0.000 | 0.000 | 0.000 | 0 | 0.0016 | 0 | 0 | 0 | 0 |
| Ascomycota | Sordariomycetes | Xylariomycetidae | Xylariales | Apiosporaceae_sp. | 0.000 | 0.000 | 0.000 | 0.000 | 0.000 | 0 | 0 | 0 | 0 | 0 | 0.04 |
| Ascomycota | Sordariomycetes | Xylariales | Apiosporaceae | Arthrinium_phaeospermum | 0.000 | 0.000 | 0.000 | 0.000 | 0.000 | 0 | 0.0016 | 0 | 0 | 0.0055 | 0.02 |
| Ascomycota | Sordariomycetes | Xylariales | Apiosporaceae | Arthrinium_sp. | 0.000 | 0.000 | 0.000 | 0.000 | 0.000 | 0 | 0 | 0.0111 | 0.0174 | 0.0055 | 0 |
| Ascomycota | Orbiliomycetes | Orbiliales | Orbiliaceae | Arthrobotrys_oligospora | 0.001 | 0.000 | 0.000 | 0.001 | 0.005 | 0.000 | 0.000 | 0.000 | 0.000 | 0.000 | 0.000 |
| Ascomycota | Eurotiomycetes | Onygenales | Arthrodermataceae | Arthroderma_redellii | 0.000 | 0.000 | 0.000 | 0.020 | 0.025 | 0.000 | 0.000 | 0.000 | 0.000 | 0.000 | 0.000 |
| Ascomycota | Dothiodeomycetes | Eremomycetales | Eremomycetaceae | Arthrographis_curvata | 0.000 | 0.000 | 0.000 | 0.001 | 0.000 | 0.000 | 0.000 | 0.000 | 0.000 | 0.000 | 0.000 |
| Ascomycota | Dothiodeomycetes | Eremomycetales | Eremomycetaceae | Arthrographis_kalrae | 0.000 | 0.000 | 0.000 | 0.000 | 0.005 | 0.000 | 0.000 | 0.000 | 0.000 | 0.000 | 0.000 |
| Ascomycota | Sordariomycetes | Xylariales | Xylariaceaea | Arthroxylaria_elegans | 0.000 | 0.000 | 0.006 | 0.000 | 0.000 | 0.000 | 0.000 | 0.000 | 0.000 | 0.000 | 0.000 |
| Ascomycota | Saccharomycetes | Saccharomycetales | Dipodascaceae | Arxula_adeninivorans | 0.000 | 0.000 | 0.000 | 0.000 | 0.000 | 0 | 0 | 0 | 0 | 0.0055 | 0 |
| Ascomycota | Sordariomycetes | Hypocreales | Clavicipitaceae | Aschersonia_sp. | 0.000 | 0.000 | 0.000 | 0.000 | 0.000 | 0 | 0.0033 | 0.0388 | 0.0232 | 0.0221 | 0 |
| Ascomycota | Dothiodeomycetes | Pleosporales | Didymellaceae | Ascochyta_pinodes | 0.000 | 0.000 | 0.000 | 0.000 | 0.000 | 0.000 | 0.000 | 0.000 | 0.000 | 0.000 | 0.000 |
| Ascomycota | Leotiomycetes | Helotiales | Helotiaceae | Ascocoryne_sarcoides | 0.004 | 0.000 | 0.000 | 0.000 | 0.000 | 0 | 0 | 0 | 0 | 0.0055 | 0 |
| Unclassified | Unclassified | Unclassified | Unclassified | Ascomycota_sp. | 0.000 | 0.000 | 0.000 | 0.000 | 0.000 | 0.1976 | 0.0164 | 0.4714 | 0.714 | 0.2318 | 1.0404 |
| Ascomycota | Sordariomycetes | Xylariales | Xylariaceaea | Ascotricha_sp. | 0.000 | 0.000 | 0.000 | 0.000 | 0.000 | 0 | 0.0016 | 0.0055 | 0.0058 | 0.0055 | 0 |
| Ascomycota | Eurotiomycetes | Eurotiales | Aspergillaceae | Aspergillus_aculeatinus | 0.000 | 0.000 | 0.000 | 0.000 | 0.000 | 0 | 0 | 0.0055 | 0.0813 | 0.0166 | 0.08 |
| Ascomycota | Eurotiomycetes | Eurotiales | Aspergillaceae | Aspergillus_clavatus | 0.001 | 0.000 | 0.000 | 0.002 | 0.005 | 0 | 0 | 0 | 0 | 0.0055 | 0 |
| Ascomycota | Eurotiomycetes | Eurotiales | Aspergillaceae | Aspergillus_flavus | 0.000 | 0.025 | 0.000 | 0.000 | 0.000 | 0 | 0 | 0 | 0.0058 | 0 | 0 |
| Ascomycota | Eurotiomycetes | Eurotiales | Aspergillaceae | Aspergillus_fumigatus | 0.003 | 0.000 | 0.000 | 0.000 | 0.000 | 0.0035 | 0 | 0.0055 | 0.0116 | 0.011 | 0 |
| Ascomycota | Eurotiomycetes | Eurotiales | Aspergillaceae | Aspergillus_japonicus | 0.000 | 0.000 | 0.000 | 0.003 | 0.000 | 0 | 0.0016 | 0.0555 | 0.0755 | 0.0166 | 0.06 |
| Ascomycota | Eurotiomycetes | Eurotiales | Aspergillaceae | Aspergillus_niger | 0.032 | 0.025 | 0.000 | 0.002 | 0.000 | 0 | 0 | 0.0166 | 0.18 | 0.0386 | 0.1 |
| Ascomycota | Eurotiomycetes | Eurotiales | Aspergillaceae | Aspergillus_nomius | 0.000 | 0.000 | 0.000 | 0.000 | 0.000 | 0 | 0 | 0 | 0.0058 | 0 | 0 |

**Table S8 continued.**

| | | | | | | | | | | | | | | | |
|---|---|---|---|---|---|---|---|---|---|---|---|---|---|---|---|
| Ascomycota | Eurotiomycetes | Eurotiales | Aspergillaceae | Aspergillus_oryzae | 0.001 | 0.050 | 0.000 | 0.000 | 0.000 | 0 | 0 | 0 | 0 | 0.011 | 0 |
| Ascomycota | Eurotiomycetes | Eurotiales | Aspergillaceae | Aspergillus_puniceus | 0.000 | 0.000 | 0.000 | 0.000 | 0.000 | 0 | 0 | 0 | 0 | 0.0055 | 0 |
| Ascomycota | Eurotiomycetes | Eurotiales | Aspergillaceae | Aspergillus_ruber | 0.001 | 0.000 | 0.000 | 0.000 | 0.005 | 0.000 | 0.000 | 0.000 | 0.000 | 0.000 | 0.000 |
| Ascomycota | Eurotiomycetes | Eurotiales | Aspergillaceae | Aspergillus_sp. | 0.003 | 0.000 | 0.000 | 0.000 | 0.000 | 0 | 0 | 0 | 0 | 0.011 | 0 |
| Ascomycota | Eurotiomycetes | Eurotiales | Aspergillaceae | Aspergillus_stellifer | 0.000 | 0.000 | 0.000 | 0.000 | 0.000 | 0 | 0 | 0 | 0.0058 | 0 | 0 |
| Ascomycota | Eurotiomycetes | Eurotiales | Aspergillaceae | Aspergillus_tamarii | 0.000 | 0.025 | 0.000 | 0.000 | 0.000 | 0.000 | 0.000 | 0.000 | 0.000 | 0.000 | 0.000 |
| Ascomycota | Eurotiomycetes | Eurotiales | Aspergillaceae | Aspergillus_tubingensis | 0.025 | 0.000 | 0.006 | 0.000 | 0.000 | 0 | 0.0033 | 0.0111 | 0.0987 | 0.0276 | 0.12 |
| Ascomycota | Eurotiomycetes | Eurotiales | Aspergillaceae | Aspergillus_ustus | 0.000 | 0.000 | 0.000 | 0.000 | 0.000 | 0 | 0 | 0 | 0.0058 | 0.0055 | 0 |
| Ascomycota | Dothideomycetes | Dothideales | Saccotheciaceae | Aureobasidium_iranianum | 0.001 | 0.000 | 0.000 | 0.000 | 0.000 | 0.000 | 0.000 | 0.000 | 0.000 | 0.000 | 0.000 |
| Ascomycota | Dothideomycetes | Dothideales | Saccotheciaceae | Aureobasidium_pullulans | 0.001 | 0.000 | 0.000 | 0.000 | 0.005 | 0.000 | 0.000 | 0.000 | 0.000 | 0.000 | 0.000 |
| Ascomycota | Dothideomycetes | Dothideales | Saccotheciaceae | Aureobasidium_sp. | 0.003 | 0.000 | 0.000 | 0.000 | 0.000 | 0.000 | 0.000 | 0.000 | 0.000 | 0.000 | 0.000 |
| Basidiomycota | Agaricomycetes | Auriculariales | Auriculariaceae | Auricularia_auricula-judae | 0.001 | 0.000 | 0.000 | 0.002 | 0.000 | 0.000 | 0.000 | 0.000 | 0.000 | 0.000 | 0.000 |
| Ascomycota | Sordariomycetes | Xylariales | Sporocadaceae | Bartalinia_sp. | 0.004 | 0.000 | 0.000 | 0.000 | 0.000 | 0.000 | 0.000 | 0.000 | 0.000 | 0.000 | 0.000 |
| Unclassified | Unclassified | Unclassified | Unclassified | Basidiomycete_sp. | 0.001 | 0.000 | 0.000 | 0.000 | 0.000 | 0 | 0.0033 | 0.0111 | 0.0116 | 0.0055 | 0.04 |
| Basidiomycota | Unclassified | Unclassified | Unclassified | Basidiomycota_sp. | 0.001 | 0.000 | 0.011 | 0.000 | 0.000 | 0.000 | 0.000 | 0.000 | 0.000 | 0.000 | 0.000 |
| Chytridiomycota | Chytridiomycetes | Rhizophydiales | Unclassified | Batrachochytrium_dendrobatidis | 0.003 | 0.000 | 0.000 | 0.000 | 0.000 | 0.000 | 0.000 | 0.000 | 0.000 | 0.000 | 0.000 |
| Ascomycota | Sordariomycetes | Hypocreales | Cordycipitaceae | Beauveria_bassiana | 0.018 | 0.025 | 0.000 | 0.005 | 0.010 | 0.0354 | 0.0016 | 0.0721 | 0.0755 | 0.0607 | 0.06 |
| Ascomycota | Sordariomycetes | Hypocreales | Cordycipitaceae | Beauveria_sp. | 0.000 | 0.000 | 0.000 | 0.000 | 0.000 | 0 | 0 | 0 | 0.0058 | 0 | 0 |
| Ascomycota | Sordariomycetes | Xylariales | Beltraniaceae | Beltrania_pseudorhombica | 0.001 | 0.000 | 0.000 | 0.000 | 0.000 | 0.000 | 0.000 | 0.000 | 0.000 | 0.000 | 0.000 |
| Ascomycota | Sordariomycetes | Xylariales | Beltraniaceae | Beltraniella_endiandrae | 0.000 | 0.000 | 0.000 | 0.000 | 0.000 | 0 | 0 | 0 | 0 | 0.011 | 0 |
| Ascomycota | Leotiomycetes | Erysiphales | Erysiphaceae | Blumeria_graminis | 0.000 | 0.000 | 0.000 | 0.000 | 0.000 | 0.000 | 0.000 | 0.000 | 0.000 | 0.000 | 0.000 |
| Ascomycota | Leotiomycetes | Helotiales | Dermataceae | Blumeriella_kerriae | 0.001 | 0.000 | 0.000 | 0.000 | 0.000 | 0.000 | 0.000 | 0.000 | 0.000 | 0.000 | 0.000 |
| Basidiomycota | Agaricomycetes | Boletales | Boletaceae | Boletus_pinophilus | 0.000 | 0.000 | 0.000 | 0.016 | 0.005 | 0.000 | 0.000 | 0.000 | 0.000 | 0.000 | 0.000 |
| Ascomycota | Dothideomycetes | Botryosphaeriales | Botryosphaeriaceae | Botryosphaeria_dothidea | 0.000 | 0.050 | 0.000 | 0.000 | 0.000 | 0.000 | 0.000 | 0.000 | 0.000 | 0.000 | 0.000 |
| Ascomycota | Dothideomycetes | Botryosphaeriales | Botryosphaeriaceae | Botryosphaeria_sp. | 0.000 | 0.000 | 0.000 | 0.000 | 0.005 | 0 | 0 | 0 | 0.0058 | 0 | 0 |
| Ascomycota | Leotiomycetes | Helotiales | Sclerotiniaceae | Botrytis_cinerea | 0.003 | 0.000 | 0.000 | 0.000 | 0.065 | 0.0035 | 0 | 0.0055 | 0 | 0.0055 | 0.02 |
| Ascomycota | Leotiomycetes | Helotiales | Sclerotiniaceae | Botrytis_sp. | 0.000 | 0.000 | 0.000 | 0.000 | 0.000 | 0 | 0 | 0 | 0.0058 | 0 | 0 |
| Ascomycota | Eurotiomycetes | Eurotiales | Thermoascaceae | Byssochlamys_spectabilis | 0.000 | 0.000 | 0.000 | 0.000 | 0.000 | 0.0035 | 0 | 0 | 0 | 0 | 0 |
| Unclassified | Unclassified | Unclassified | Unclassified | Calluna_vulgaris | 0.000 | 0.000 | 0.000 | 0.000 | 0.000 | 0 | 0 | 0 | 0.0058 | 0 | 0 |
| Ascomycota | Sordariomycetes | Hypocreales | Nectriaceae | Calonectria_canadiana | 0.000 | 0.000 | 0.000 | 0.000 | 0.000 | 0 | 0 | 0.0055 | 0.0116 | 0 | 0 |
| Ascomycota | Sordariomycetes | Hypocreales | Nectriaceae | Calonectria_pseudonaviculata | 0.000 | 0.000 | 0.000 | 0.000 | 0.000 | 0 | 0 | 0 | 0.0058 | 0 | 0 |
| Ascomycota | Sordariomycetes | Hypocreales | Nectriaceae | Calonectria_pseudoreteaudii | 0.000 | 0.000 | 0.006 | 0.000 | 0.000 | 0.0035 | 0.0016 | 0.0277 | 0 | 0 | 0 |

**Table S8 continued.**

| | | | | | | | | | | | | | | | |
|---|---|---|---|---|---|---|---|---|---|---|---|---|---|---|---|
| Ascomycota | Saccharomycetes | Saccharomycetales | Unclassfied | Candida_anutae | 0.000 | 0.000 | 0.000 | 0.003 | 0.000 | 0.000 | 0.000 | 0.000 | 0.000 | 0.000 | 0.000 |
| Ascomycota | Saccharomycetes | Saccharomycetales | Unclassfied | Candida_parapsilosis | 0.000 | 0.000 | 0.000 | 0.000 | 0.000 | 0.000 | 0.000 | 0.000 | 0.000 | 0.000 | 0.000 |
| Ascomycota | Saccharomycetes | Saccharomycetales | Unclassfied | Candida_sp. | 0.050 | 0.000 | 0.006 | 0.003 | 0.000 | 0.000 | 0.000 | 0.000 | 0.000 | 0.000 | 0.000 |
| Ascomycota | Saccharomycetes | Saccharomycetales | Unclassfied | Candida_tropicalis | 0.001 | 0.000 | 0.006 | 0.000 | 0.000 | 0.000 | 0.000 | 0.000 | 0.000 | 0.000 | 0.000 |
| Ascomycota | Saccharomycetes | Saccharomycetales | Unclassfied | Candida_vespimorsuum | 0.000 | 0.025 | 0.000 | 0.000 | 0.000 | 0.000 | 0.000 | 0.000 | 0.000 | 0.000 | 0.000 |
| Ascomycota | Sordariomycetes | Xylariales | Unclassified | Castanediella_acaciae | 0.001 | 0.000 | 0.000 | 0.000 | 0.000 | 0.000 | 0.000 | 0.000 | 0.000 | 0.000 | 0.000 |
| Ascomycota | Dothideomycetes | Unclassified | Gloniaceae | Cenococcum_geophilum | 0.000 | 0.000 | 0.000 | 0.001 | 0.010 | 0.000 | 0.000 | 0.000 | 0.000 | 0.000 | 0.000 |
| Ascomycota | Sordariomycetes | Microascales | Unclassified | Cephalotrichiella_penicillata | 0.000 | 0.000 | 0.000 | 0.000 | 0.000 | 0 | 0 | 0 | 0 | 0.0055 | 0 |
| Basidiomycota | Agaricomycetes | Polyporales | Irpicaceae | Ceriporia_lacerata | 0.001 | 0.000 | 0.000 | 0.000 | 0.000 | 0.000 | 0.000 | 0.000 | 0.000 | 0.000 | 0.000 |
| Basidiomycota | Agaricomycetes | Polyporales | Meruliaceae | Ceriporiopsis_rivulosa | 0.001 | 0.000 | 0.000 | 0.000 | 0.000 | 0.000 | 0.000 | 0.000 | 0.000 | 0.000 | 0.000 |
| Ascomycota | Sordariomycetes | Sordariales | Chaetomiaceae | Chaetomium_globosum | 0.000 | 0.000 | 0.000 | 0.000 | 0.000 | 0 | 0 | 0.0055 | 0.0058 | 0.011 | 0 |
| Ascomycota | Sordariomycetes | Sordariales | Chaetomiaceae | Chaetomium_sp. | 0.000 | 0.076 | 0.000 | 0.000 | 0.000 | 0 | 0 | 0 | 0.0174 | 0.0166 | 0 |
| Ascomycota | Sordariomycetes | Chaetosphaeriales | Chaetosphaeriaceae | Chloridium_virescens_var._chlamydosporum | 0.000 | 0.000 | 0.000 | 0.000 | 0.000 | 0 | 0 | 0.0055 | 0.0116 | 0 | 0 |
| Basidiomycota | Agaricomycetes | Agaricales | Schizophyllaceae | Chondrostereum_sp. | 0.000 | 0.000 | 0.000 | 0.000 | 0.000 | 0.0035 | 0 | 0 | 0 | 0 | 0 |
| Ascomycota | Eurotiomycetes | Onygenales | Unclassified | Chrysosporium_leigongshanense | 0.000 | 0.000 | 0.000 | 0.000 | 0.000 | 0.0035 | 0 | 0 | 0 | 0 | 0 |
| Ascomycota | Eurotiomycetes | Onygenales | Unclassified | Chrysosporium_sp. | 0.000 | 0.000 | 0.000 | 0.000 | 0.000 | 0.000 | 0.000 | 0.000 | 0.000 | 0.000 | 0.000 |
| Ascomycota | Lecanoromycetes | Lecanorales | Cladoniaceae | Cladonia_macilenta | 0.001 | 0.000 | 0.000 | 0.000 | 0.000 | 0.000 | 0.000 | 0.000 | 0.000 | 0.000 | 0.000 |
| Ascomycota | Lecanoromycetes | Lecanorales | Cladoniaceae | Cladonia_metacorallifera | 0.009 | 0.000 | 0.000 | 0.000 | 0.000 | 0.000 | 0.000 | 0.000 | 0.000 | 0.000 | 0.000 |
| Ascomycota | Eurotiomycetes | Chaetothyriales | Herpotrichiellaceae | Cladophialophora_chaetospira | 0.000 | 0.000 | 0.000 | 0.016 | 0.010 | 0.000 | 0.000 | 0.000 | 0.000 | 0.000 | 0.000 |
| Ascomycota | Eurotiomycetes | Onygenales | Unclassified | Cladosporium_cladosporioides | 0.018 | 0.177 | 0.158 | 0.325 | 0.781 | 0 | 0 | 0 | 0.0174 | 0 | 0 |
| Ascomycota | Dothideomycetes | Cladosporiales | Cladosporiaceae | Cladosporium_halotolerans | 0.000 | 0.000 | 0.000 | 0.000 | 0.000 | 0.000 | 0.000 | 0.000 | 0.000 | 0.000 | 0.000 |
| Ascomycota | Dothideomycetes | Cladosporiales | Cladosporiaceae | Cladosporium_herbarum | 0.000 | 0.025 | 0.000 | 0.000 | 0.000 | 0.000 | 0.000 | 0.000 | 0.000 | 0.000 | 0.000 |
| Ascomycota | Dothideomycetes | Cladosporiales | Cladosporiaceae | Cladosporium_oxysporum | 0.000 | 0.000 | 0.000 | 0.000 | 0.000 | 0.000 | 0.000 | 0.000 | 0.000 | 0.000 | 0.000 |
| Ascomycota | Dothideomycetes | Cladosporiales | Cladosporiaceae | Cladosporium_sp. | 0.009 | 0.000 | 0.000 | 0.018 | 0.055 | 0 | 0 | 0.0055 | 0.0174 | 0 | 0 |
| Ascomycota | Dothideomycetes | Cladosporiales | Cladosporiaceae | Cladosporium_sphaerospermum | 0.000 | 0.000 | 0.000 | 0.002 | 0.000 | 0.000 | 0.000 | 0.000 | 0.000 | 0.000 | 0.000 |
| Ascomycota | Dothideomycetes | Cladosporiales | Cladosporiaceae | Cladosporium_tenuissimum | 0.000 | 0.000 | 0.000 | 0.000 | 0.000 | 0.000 | 0.000 | 0.000 | 0.000 | 0.000 | 0.000 |
| Ascomycota | Sordariomycetes | Phyllachorales | Unclassified | Clathrosporium_retortum | 0.000 | 0.000 | 0.000 | 0.000 | 0.000 | 0 | 0 | 0.0055 | 0 | 0 | 0 |
| Ascomycota | Sordariomycetes | Hypocreales | Clavicipitaceae | Claviceps_purpurea | 0.000 | 0.000 | 0.000 | 0.000 | 0.000 | 0 | 0 | 0.0111 | 0.0116 | 0 | 0.02 |
| Ascomycota | Sordariomycetes | Hypocreales | Unclassified | Clavicipitaceae_sp. | 0.000 | 0.000 | 0.000 | 0.000 | 0.000 | 0.0035 | 0 | 0 | 0 | 0.0055 | 0.02 |
| Ascomycota | Sordariomycetes | Hypocreales | Bionectriaceae | Clonostachys_rosea | 0.000 | 0.000 | 0.000 | 0.000 | 0.000 | 0 | 0 | 0 | 0 | 0 | 0.02 |
| Ascomycota | Sordariomycetes | Hypocreales | Bionectriaceae | Clonostachys_sp. | 0.000 | 0.000 | 0.000 | 0.000 | 0.000 | 0 | 0 | 0.0055 | 0 | 0 | 0 |
| Ascomycota | Dothideomycetes | Pleosporales | Pleosporaceae | Cochliobolus_sp. | 0.001 | 0.000 | 0.006 | 0.013 | 0.060 | 0.000 | 0.000 | 0.000 | 0.000 | 0.000 | 0.000 |

**Table S8 continued.**

| | | | | | | | | | | | | | | | |
|---|---|---|---|---|---|---|---|---|---|---|---|---|---|---|---|
| Basidiomycota | Pucciniomycetes | Pucciniales | Coleosporiaceae | Coleosporium_tussilaginis | 0.000 | 0.000 | 0.000 | 0.001 | 0.000 | 0.000 | 0.000 | 0.000 | 0.000 | 0.000 | 0.000 |
| Ascomycota | Sordariomycetes | Glomerellales | Glomerellaceae | Colletotrichum_axonopodi | 0.000 | 0.000 | 0.000 | 0.000 | 0.000 | 0 | 0 | 0 | 0 | 0.0055 | 0 |
| Ascomycota | Sordariomycetes | Glomerellales | Glomerellaceae | Colletotrichum_capsici | 0.000 | 0.000 | 0.000 | 0.000 | 0.000 | 0 | 0 | 0 | 0 | 0.0055 | 0 |
| Ascomycota | Sordariomycetes | Glomerellales | Glomerellaceae | Colletotrichum_orbiculare | 0.000 | 0.000 | 0.000 | 0.000 | 0.000 | 0 | 0 | 0 | 0.0058 | 0 | 0 |
| Basidiomycota | Agaricomycetes | Hymenochaetales | Hymenochaetaceae | Coniferiporia_sulphurascens | 0.001 | 0.000 | 0.000 | 0.000 | 0.000 | 0.000 | 0.000 | 0.000 | 0.000 | 0.000 | 0.000 |
| Ascomycota | Sordariomycetes | Coniochaetales | Coniochaetaceae | Coniochaeta_ligniaria | 0.004 | 0.000 | 0.000 | 0.000 | 0.000 | 0.000 | 0.000 | 0.000 | 0.000 | 0.000 | 0.000 |
| Basidiomycota | Agaricomycetes | Agaricales | Psathyrellaceae | Coprinellus_disseminatus | 0.000 | 0.050 | 0.000 | 0.000 | 0.000 | 0.000 | 0.000 | 0.000 | 0.000 | 0.000 | 0.000 |
| Basidiomycota | Agaricomycetes | Agaricales | Psathyrellaceae | Coprinellus_micaceus | 0.000 | 0.000 | 0.000 | 0.000 | 0.000 | 0.0037 | 0.0016 | 0 | 0 | 0 | 0 |
| Basidiomycota | Agaricomycetes | Agaricales | Psathyrellaceae | Coprinopsis_sp. | 0.000 | 0.000 | 0.000 | 0.000 | 0.000 | 0.000 | 0.000 | 0.000 | 0.000 | 0.000 | 0.000 |
| Basidiomycota | Agaricomycetes | Agaricales | Agaricaceae | Coprinus_comatus | 0.000 | 0.000 | 0.006 | 0.000 | 0.000 | 0.000 | 0.000 | 0.000 | 0.000 | 0.000 | 0.000 |
| Chytridiomycota | Chyrtidiomycetes | Rhizophydiales | Coralloidiomyces | Coralloidiomyces_digitatus | 0.007 | 0.000 | 0.000 | 0.001 | 0.000 | 0.000 | 0.000 | 0.000 | 0.000 | 0.000 | 0.000 |
| Ascomycota | Sordariomycetes | Hypocreales | Cordycipitaceae | Cordyceps_bassiana | 0.000 | 0.000 | 0.000 | 0.000 | 0.000 | 0 | 0 | 0 | 0.0058 | 0 | 0 |
| Ascomycota | Sordariomycetes | Hypocreales | Cordycipitaceae | Cordyceps_militaris | 0.021 | 0.000 | 0.000 | 0.000 | 0.000 | 0.0176 | 0.0016 | 0.0333 | 0.0058 | 0.011 | 0.04 |
| Basidiomycota | Agaricomycetes | Corticiales | Unclassified | Corticiales_sp. | 0.000 | 0.000 | 0.006 | 0.027 | 0.045 | 0.000 | 0.000 | 0.000 | 0.000 | 0.000 | 0.000 |
| Basidiomycota | Agaricomycetes | Agaricales | Cortinariaceae | Cortinarius_collybianus | 0.000 | 0.000 | 0.000 | 0.000 | 0.000 | 0.007 | 0.000 | 0.000 | 0.000 | 0.000 | 0.000 |
| Basidiomycota | Agaricomycetes | Agaricales | Cortinariaceae | Cortinarius_magellanicus | 0.001 | 0.000 | 0.000 | 0.000 | 0.000 | 0 | 0.0016 | 0 | 0 | 0 | 0 |
| Basidiomycota | Agaricomycetes | Agaricales | Cortinariaceae | Cortinarius_turgidus | 0.003 | 0.000 | 0.000 | 0.000 | 0.000 | 0.007 | 0.000 | 0.000 | 0.000 | 0.000 | 0.000 |
| Ascomycota | Sordariomycetes | Hypocreales | Nectriaceae | Cosmospora_flavoviridis | 0.020 | 0.050 | 0.039 | 0.000 | 0.005 | 0.0614 | 0.023 | 0.0055 | 0 | 0 | 0 |
| Ascomycota | Sordariomycetes | Diaporthales | Cryphonectriaceae | Cryphonectria_parasitica | 0.000 | 0.000 | 0.000 | 0.000 | 0.000 | 0.0035 | 0 | 0.0055 | 0 | 0 | 0 |
| Ascomycota | Sordariomycetes | Diaporthales | Cryphonectriaceae | Cryphonectria_parasitica | 0.001 | 0.000 | 0.000 | 0.000 | 0.000 | 0 | 0 | 0 | 0 | 0.0055 | 0 |
| Basidiomycota | Tremellomycetes | Tremellales | Cryptococcaceae | Cryptococcus | 0.000 | 0.000 | 0.000 | 0.002 | 0.005 | 0.000 | 0.000 | 0.000 | 0.000 | 0.000 | 0.000 |
| Basidiomycota | Tremellomycetes | Tremellales | Cryptococcaceae | Cryptococcus_cf._cellulolyticus | 0.003 | 0.000 | 0.000 | 0.000 | 0.000 | 0.000 | 0.000 | 0.000 | 0.000 | 0.000 | 0.000 |
| Basidiomycota | Tremellomycetes | Tremellales | Cryptococcaceae | Cryptococcus_magnus | 0.001 | 0.000 | 0.000 | 0.000 | 0.005 | 0.000 | 0.000 | 0.000 | 0.000 | 0.000 | 0.000 |
| Ascomycota | Tremellomycetes | Tremellales | Cryptococcaceae | Cryptococcus_neoformans | 0.000 | 0.000 | 0.000 | 0.000 | 0.000 | 0 | 0 | 0 | 0.0058 | 0 | 0 |
| Basidiomycota | Tremellomycetes | Tremellales | Cryptococcaceae | Cryptococcus_neoformans_var._grubii | 0.020 | 0.000 | 0.000 | 0.002 | 0.005 | 0.004 | 0.000 | 0.000 | 0.000 | 0.000 | 0.000 |
| Basidiomycota | Tremellomycetes | Tremellales | Cryptococcaceae | Cryptococcus_sp. | 0.001 | 0.000 | 0.000 | 0.000 | 0.000 | 0.000 | 0.000 | 0.000 | 0.000 | 0.000 | 0.000 |
| Ascomycota | Sordariomycetes | Xylariales | Diatrypaceae | Cryptosphaeria_pullmanensis | 0.000 | 0.000 | 0.000 | 0.000 | 0.000 | 0 | 0 | 0.0055 | 0 | 0 | 0 |
| Mucoromycota | Mucoromycetes | Mucorales | Cunninghamellaceae | Cunninghamella_sp. | 0.001 | 0.000 | 0.000 | 0.000 | 0.000 | 0.000 | 0.000 | 0.000 | 0.000 | 0.000 | 0.000 |
| Ascomycota | Dothideomycetes | Pleosporales | Pleosporaceae | Curvularia_hawaiiensis | 0.000 | 0.000 | 0.000 | 0.005 | 0.015 | 0.000 | 0.000 | 0.000 | 0.000 | 0.000 | 0.000 |
| Ascomycota | Dothideomycetes | Pleosporales | Pleosporaceae | Curvularia_hominis | 0.000 | 0.000 | 0.006 | 0.000 | 0.000 | 0.000 | 0.000 | 0.000 | 0.000 | 0.000 | 0.000 |
| Ascomycota | Dothideomycetes | Pleosporales | Pleosporaceae | Curvularia_lunata | 0.000 | 0.000 | 0.006 | 0.000 | 0.010 | 0.000 | 0.000 | 0.000 | 0.000 | 0.000 | 0.000 |
| Ascomycota | Dothideomycetes | Pleosporales | Pleosporaceae | Curvularia_sp. | 0.000 | 0.000 | 0.000 | 0.000 | 0.005 | 0.000 | 0.000 | 0.000 | 0.000 | 0.000 | 0.000 |

**Table S8 continued.**

| | | | | | | | | | | | | | | | |
|---|---|---|---|---|---|---|---|---|---|---|---|---|---|---|---|
| Ascomycota | Dothideomycetes | Pleosporales | Pleosporaceae | Curvularia_tsudae | 0.000 | 0.000 | 0.000 | 0.000 | 0.010 | 0.000 | 0.000 | 0.000 | 0.000 | 0.000 | 0.000 |
| Ascomycota | Sordariomycetes | Hypocreales | Nectriaceae | Cylindrocladiella_sp._E6809b | 0.000 | 0.000 | 0.000 | 0.000 | 0.000 | 0 | 0 | 0 | 0.0058 | 0 | 0 |
| Ascomycota | Sordariomycetes | Hypocreales | Nectriaceae | Cylindrocladiella_sp._ZH60 | 0.000 | 0.000 | 0.000 | 0.000 | 0.000 | 0 | 0 | 0.0388 | 0.0464 | 0.0276 | 0.1 |
| Chytridiomycota | Neocallimastigomycetes | Neocallimastigales | Neocallimastigaceae | Cyllamyces_sp. | 0.001 | 0.000 | 0.000 | 0.000 | 0.000 | 0.000 | 0.000 | 0.000 | 0.000 | 0.000 | 0.000 |
| Basidiomycota | Agaricomycetes | Agaricales | Agaricaceae | Cystoderma_carpaticum | 0.001 | 0.000 | 0.006 | 0.000 | 0.005 | 0.0072 | 0.0049 | 0 | 0 | 0 | 0 |
| Ascomycota | Sordariomycetes | Diaporthales | Valsaceae | Cytospora_sp. | 0.005 | 0.000 | 0.000 | 0.000 | 0.000 | 0.000 | 0.000 | 0.000 | 0.000 | 0.000 | 0.000 |
| Ascomycota | Sordariomycetes | Hypocreales | Nectriaceae | Dactylonectria_vitis | 0.000 | 0.000 | 0.000 | 0.000 | 0.000 | 0 | 0 | 0 | 0.0174 | 0 | 0 |
| Ascomycota | Sordariomycetes | Xylariales | Hypoxylaceae | Daldinia_eschscholzii | 0.000 | 0.000 | 0.023 | 0.000 | 0.000 | 0 | 0.0016 | 0.0555 | 0.029 | 0.0221 | 0.08 |
| Ascomycota | Sordariomycetes | Hypocreales | Nectriaceae | Dematiocladium_celtidicola | 0.000 | 0.000 | 0.000 | 0.000 | 0.000 | 0 | 0 | 0.0055 | 0 | 0 | 0 |
| Ascomycota | Sordariomycetes | Diaporthales | Diaporthaceae | Diaporthe_longicolla | 0.000 | 0.000 | 0.000 | 0.000 | 0.000 | 0 | 0 | 0 | 0.0058 | 0.0055 | 0 |
| Ascomycota | Sordariomycetes | Diaporthales | Diaporthaceae | Diaporthe_melonis | 0.000 | 0.000 | 0.000 | 0.003 | 0.010 | 0.000 | 0.000 | 0.000 | 0.000 | 0.000 | 0.000 |
| Ascomycota | Sordariomycetes | Diaporthales | Diaporthaceae | Diaporthe_sp. | 0.000 | 0.000 | 0.000 | 0.000 | 0.000 | 0.000 | 0.000 | 0.000 | 0.000 | 0.000 | 0.000 |
| Ascomycota | Dothideomycetes | Pleosporales | Didymellaceae | Didymella_sp. | 0.000 | 0.000 | 0.000 | 0.001 | 0.000 | 0.000 | 0.000 | 0.000 | 0.000 | 0.000 | 0.000 |
| Ascomycota | Dothideomycetes | Pleosporales | Didymellaceae | Didymellaceae_sp. | 0.001 | 0.000 | 0.000 | 0.000 | 0.000 | 0.000 | 0.000 | 0.000 | 0.000 | 0.000 | 0.000 |
| Ascomycota | Leotiomycetes | Helotiales | Helotiaceae | Dimorphospora_foliicola | 0.000 | 0.000 | 0.000 | 0.000 | 0.000 | 0.000 | 0.000 | 0.000 | 0.000 | 0.000 | 0.000 |
| Ascomycota | Sordariomycetes | Chaetosphaeriales | Chaetosphaeriaceae | Dinemasporium_trichophoricola | 0.000 | 0.000 | 0.000 | 0.000 | 0.000 | 0.0035 | 0 | 0 | 0 | 0 | 0 |
| Mucoromycota | Glomeromycetes | Diversisporales | Diversisporaceae | Diversispora_sp. | 0.004 | 0.000 | 0.000 | 0.000 | 0.000 | 0.000 | 0.000 | 0.000 | 0.000 | 0.000 | 0.000 |
| Mucoromycota | Glomeromycetes | Glomerales | Glomeraraceae | Dominikia_bernensis | 0.529 | 0.076 | 0.006 | 0.025 | 0.035 | 0.000 | 0.000 | 0.000 | 0.000 | 0.000 | 0.000 |
| Ascomycota | Dothideomycetes | Unclassified | Unclassified | Dothideomycetes_sp. | 0.000 | 0.025 | 0.006 | 0.000 | 0.000 | 0.000 | 0.000 | 0.000 | 0.000 | 0.000 | 0.000 |
| Ascomycota | Dothideomycetes | Pleosporales | Pleosporaceae | Edenia_gomezpompae | 0.001 | 0.025 | 0.000 | 0.000 | 0.000 | 0.000 | 0.000 | 0.000 | 0.000 | 0.000 | 0.000 |
| Ascomycota | Dothideomycetes | Myriangiales | Elsinoaceae | Elsinoe_genipae-americanae | 0.001 | 0.000 | 0.000 | 0.000 | 0.000 | 0.000 | 0.000 | 0.000 | 0.000 | 0.000 | 0.000 |
| Ascomycota | Dothideomycetes | Myriangiales | Elsinoaceae | Elsinoe_othonnae | 0.000 | 0.000 | 0.000 | 0.001 | 0.000 | 0.000 | 0.000 | 0.000 | 0.000 | 0.000 | 0.000 |
| Ascomycota | Dothideomycetes | Myriangiales | Elsinoaceae | Elsinoe_punicae | 0.001 | 0.000 | 0.000 | 0.000 | 0.000 | 0.000 | 0.000 | 0.000 | 0.000 | 0.000 | 0.000 |
| Ascomycota | Eurotiomycetes | Eurotiales | Aspergillaceae | Emericella_rugulosa | 0.000 | 0.000 | 0.000 | 0.000 | 0.000 | 0 | 0 | 0 | 0 | 0.0055 | 0 |
| Ascomycota | Eurotiomycetes | Verrucariales | Verrucariaceae | Endocarpon_pusillum | 0.001 | 0.000 | 0.000 | 0.000 | 0.000 | 0 | 0.0033 | 0 | 0.0058 | 0 | 0 |
| Ascomycota | Unclassified | Unclassified | Unclassified | Endophytic_ascomycete | 0.003 | 0.000 | 0.000 | 0.000 | 0.000 | 0.000 | 0.000 | 0.000 | 0.000 | 0.000 | 0.000 |
| Ascomycota | Agaricomycetes | Agaricales | Entolomataceae | Entoloma_pseudoconferendum | 0.001 | 0.025 | 0.000 | 0.000 | 0.000 | 0.0037 | 0.0016 | 0 | 0 | 0 | 0 |
| Unclassified | Unclassified | Unclassified | Unclassified | Environmental_fungal | 0.001 | 0.000 | 0.000 | 0.000 | 0.000 | 0.000 | 0.000 | 0.000 | 0.000 | 0.000 | 0.000 |
| Unclassified | Unclassified | Unclassified | Unclassified | Epacris_microphylla | 0.000 | 0.000 | 0.000 | 0.000 | 0.000 | 0 | 0 | 0.0055 | 0 | 0 | 0 |
| Ascomycota | Dothideomycetes | Pleosporales | Didymellaceae | Epicoccum_nigrum | 0.001 | 0.025 | 0.056 | 0.009 | 0.005 | 0.000 | 0.000 | 0.000 | 0.000 | 0.000 | 0.000 |
| Ascomycota | Dothideomycetes | Pleosporales | Didymellaceae | Epicoccum_sp. | 0.000 | 0.000 | 0.000 | 0.000 | 0.000 | 0.000 | 0.000 | 0.000 | 0.000 | 0.000 | 0.000 |
| Ascomycota | Saccharomycetes | Saccharomycetales | Saccharomycetaceae | Eremothecium_gossypii | 0.000 | 0.000 | 0.000 | 0.000 | 0.000 | 0 | 0 | 0 | 0.0058 | 0 | 0 |

**Table S8 continued.**

| | | | | | | | | | | | | | | | |
|---|---|---|---|---|---|---|---|---|---|---|---|---|---|---|---|
| Ascomycota | Unclassified | Unclassified | Unclassified | Ericoid_mycorrhizal | 0.001 | 0.000 | 0.000 | 0.000 | 0.000 | 0.000 | 0.000 | 0.000 | 0.000 | 0.000 | 0.000 |
| Ascomycota | Sordariomycetes | Hypocreales | Hypocreaceae | Escovopsioides_nivea | 0.000 | 0.000 | 0.006 | 0.000 | 0.000 | 0 | 0 | 0.0111 | 0.0058 | 0 | 0.06 |
| Ascomycota | Sordariomycetes | Hypocreales | Hypocreaceae | Escovopsis_microspora | 0.000 | 0.000 | 0.000 | 0.000 | 0.000 | 0 | 0 | 0.0055 | 0.0058 | 0 | 0 |
| Ascomycota | Sordariomycetes | Hypocreales | Hypocreaceae | Escovopsis_sp. | 0.000 | 0.000 | 0.000 | 0.000 | 0.000 | 0 | 0 | 0.0111 | 0.0174 | 0 | 0.02 |
| Ascomycota | Sordariomycetes | Hypocreales | Unclassified | Eucasphaeria_capensis | 0.000 | 0.000 | 0.000 | 0.000 | 0.000 | 0 | 0 | 0 | 0 | 0.0055 | 0.04 |
| Ascomycota | Sordariomycetes | Hypocreales | Unclassified | Eucasphaeria_cf._capensis | 0.001 | 0.000 | 0.000 | 0.000 | 0.000 | 0.000 | 0.000 | 0.000 | 0.000 | 0.000 | 0.000 |
| Ascomycota | Dothideomycetes | Pleosporales | Leptosphaeriaceae | Eudarluca_caricis | 0.000 | 0.000 | 0.006 | 0.000 | 0.000 | 0.0072 | 0 | 0 | 0 | 0 | 0 |
| Ascomycota | Leotiomycetes | Erysiphales | Erysiphaceae | Euoidium_longipes | 0.000 | 0.000 | 0.000 | 0.000 | 0.000 | 0.000 | 0.000 | 0.000 | 0.000 | 0.000 | 0.000 |
| Ascomycota | Eurotiomycetes | Eurotiales | Aspergillaceae | Eurotium_amstelodami | 0.000 | 0.000 | 0.000 | 0.000 | 0.000 | 0 | 0 | 0 | 0 | 0.011 | 0 |
| Ascomycota | Sordariomycetes | Xylariales | Diatrypaceae | Eutypa_lata | 0.034 | 0.000 | 0.000 | 0.000 | 0.000 | 0 | 0 | 0.0166 | 0 | 0.0166 | 0 |
| Ascomycota | Sordariomycetes | Xylariales | Diatrypaceae | Eutypella_sp. | 0.000 | 0.000 | 0.000 | 0.000 | 0.000 | 0.000 | 0.000 | 0.000 | 0.000 | 0.000 | 0.000 |
| Ascomycota | Eurotiomycetes | Chaetothyriales | Herpotrichiellaceae | Exophiala_sp. | 0.000 | 0.000 | 0.000 | 0.000 | 0.005 | 0.000 | 0.000 | 0.000 | 0.000 | 0.000 | 0.000 |
| Basidiomycota | Agaricomycetes | Polyporales | Unclassified | Fibroporia_radiculosa | 0.003 | 0.000 | 0.006 | 0.000 | 0.000 | 0.000 | 0.000 | 0.000 | 0.000 | 0.000 | 0.000 |
| Basidiomycota | Agaricomycetes | Atheliales | Atheliaceae | Fibularhizoctonia_sp. | 0.001 | 0.000 | 0.000 | 0.000 | 0.000 | 0.0035 | 0.0016 | 0 | 0 | 0 | 0 |
| Basidiomycota | Agaricomycetes | Agaricales | Strophoriaceae | Flammula_sp. | 0.003 | 0.000 | 0.000 | 0.000 | 0.000 | 0 | 0.0016 | 0 | 0 | 0 | 0 |
| Basidiomycota | Agaricomycetes | Agaricales | Physalacriaceae | Flammulina_velutipes | 0.003 | 0.000 | 0.011 | 0.000 | 0.000 | 0 | 0.0049 | 0 | 0 | 0 | 0 |
| Basidiomycota | Agaricomycetes | Polyporales | Irpicaceae | Flavodon_flavus | 0.001 | 0.000 | 0.006 | 0.000 | 0.000 | 0.000 | 0.000 | 0.000 | 0.000 | 0.000 | 0.000 |
| Basidiomycota | Agaricomycetes | Polyporales | Formitopsidaceae | Fomitopsis_palustris | 0.001 | 0.000 | 0.000 | 0.000 | 0.000 | 0.000 | 0.000 | 0.000 | 0.000 | 0.000 | 0.000 |
| Ascomycota | Leotiomycetes | Helotiales | Discinellaceae | Fontanospora_sp. | 0.000 | 0.000 | 0.000 | 0.000 | 0.000 | 0.0035 | 0 | 0 | 0 | 0 | 0 |
| Ascomycota | Sordariomycetes | Hypocreales | Nectriaceae | Fusarium_cf._incarnatum | 0.000 | 0.000 | 0.000 | 0.000 | 0.000 | 0 | 0 | 0 | 0.0116 | 0 | 0 |
| Ascomycota | Sordariomycetes | Hypocreales | Nectriaceae | Fusarium_equiseti | 0.000 | 0.000 | 0.017 | 0.001 | 0.005 | 0 | 0 | 0.0055 | 0.0697 | 0 | 0 |
| Ascomycota | Sordariomycetes | Hypocreales | Nectriaceae | Fusarium_graminearum | 0.004 | 0.000 | 0.000 | 0.000 | 0.000 | 0.000 | 0.000 | 0.000 | 0.000 | 0.000 | 0.000 |
| Ascomycota | Sordariomycetes | Hypocreales | Nectriaceae | Fusarium_incarnatum | 0.000 | 0.000 | 0.000 | 0.000 | 0.000 | 0 | 0 | 0.0166 | 0.0058 | 0 | 0 |
| Ascomycota | Sordariomycetes | Hypocreales | Nectriaceae | Fusarium_oxysporum | 0.003 | 0.000 | 0.000 | 0.000 | 0.000 | 0 | 0 | 0.0222 | 0.0522 | 0.0166 | 0.12 |
| Ascomycota | Sordariomycetes | Hypocreales | Nectriaceae | Fusarium_proliferatum | 0.000 | 0.000 | 0.006 | 0.000 | 0.000 | 0.000 | 0.000 | 0.000 | 0.000 | 0.000 | 0.000 |
| Ascomycota | Sordariomycetes | Hypocreales | Nectriaceae | Fusarium_solani | 0.000 | 0.000 | 0.000 | 0.000 | 0.000 | 0 | 0 | 0 | 0.0232 | 0.0442 | 0.06 |
| Ascomycota | Sordariomycetes | Hypocreales | Nectriaceae | Fusarium_sp. | 0.007 | 0.076 | 0.017 | 0.001 | 0.065 | 0 | 0.0016 | 0.0222 | 0.1567 | 0.0055 | 0.1 |
| Ascomycota | Sordariomycetes | Hypocreales | Nectriaceae | Fusarium_tricinctum | 0.000 | 0.000 | 0.000 | 0.000 | 0.000 | 0 | 0.0016 | 0 | 0 | 0 | 0 |
| Ascomycota | Sordariomycetes | Magnaporthales | Magnaporthaceae | Gaeumannomyces_graminis_var._graminis | 0.000 | 0.000 | 0.000 | 0.000 | 0.000 | 0.000 | 0.000 | 0.000 | 0.000 | 0.000 | 0.000 |
| Ascomycota | Sordariomycetes | Magnaporthales | Magnaporthaceae | Gaeumannomyces_incrustans | 0.000 | 0.000 | 0.000 | 0.000 | 0.000 | 0 | 0 | 0 | 0 | 0.0055 | 0 |
| Basidiomycota | Agaricomycetes | Polyporales | Polyporaceae | Ganoderma_adspersum | 0.000 | 0.000 | 0.000 | 0.000 | 0.000 | 0.0035 | 0.0016 | 0 | 0 | 0 | 0 |
| Basidiomycota | Agaricomycetes | Polyporales | Polyporaceae | Ganoderma_applanatum | 0.000 | 0.000 | 0.000 | 0.000 | 0.000 | 0.000 | 0.000 | 0.000 | 0.000 | 0.000 | 0.000 |

**Table S8 continued.**

| | | | | | | | | | | | | | | | |
|---|---|---|---|---|---|---|---|---|---|---|---|---|---|---|---|
| Basidiomycota | Agaricomycetes | Polyporales | Polyporaceae | Ganoderma_lingzhi | 0.000 | 0.000 | 0.000 | 0.000 | 0.000 | 0.000 | 0.000 | 0.000 | 0.000 | 0.000 | 0.000 |
| Basidiomycota | Agaricomycetes | Polyporales | Polyporaceae | Ganoderma_lucidum | 0.001 | 0.000 | 0.006 | 0.000 | 0.000 | 0 | 0.0016 | 0 | 0 | 0 | 0 |
| Ascomycota | Leotiomycetes | Unclassified | Unclassified | Gelatinomyces_siamensis | 0.000 | 0.000 | 0.000 | 0.000 | 0.005 | 0.000 | 0.000 | 0.000 | 0.000 | 0.000 | 0.000 |
| Ascomycota | Sordariomycetes | Xylariales | Xylariaceae | Geniculisynnema_termiticola | 0.000 | 0.000 | 0.000 | 0.000 | 0.000 | 0 | 0 | 0.1553 | 0.1858 | 0.0055 | 0.02 |
| Ascomycota | Leotiomycetes | Unclassified | Pseudeurotiaceae | Geomyces_auratus | 0.000 | 0.000 | 0.000 | 0.000 | 0.000 | 0 | 0 | 0 | 0 | 0 | 0.06 |
| Ascomycota | Leotiomycetes | Unclassified | Pseudeurotiaceae | Geomyces_destructans | 0.000 | 0.000 | 0.000 | 0.000 | 0.005 | 0 | 0 | 0 | 0.0116 | 0 | 0 |
| Ascomycota | Sordariomycetes | Hypocreales | Nectriaceae | Gibberella_moniliformis | 0.000 | 0.000 | 0.006 | 0.000 | 0.000 | 0 | 0 | 0.0055 | 0 | 0 | 0 |
| Mucoromycota | Glomeromycetes | Diversisporales | Gigasporaceae | Gigaspora_margarita | 0.407 | 0.076 | 0.000 | 0.002 | 0.000 | 0.000 | 0.000 | 0.000 | 0.000 | 0.000 | 0.000 |
| Mucoromycota | Glomeromycetes | Diversisporales | Gigasporaceae | Gigaspora_rosea | 0.038 | 0.000 | 0.000 | 0.000 | 0.000 | 0.000 | 0.000 | 0.000 | 0.000 | 0.000 | 0.000 |
| Ascomycota | Leotiomycetes | Helotiales | Helotiaceae | Glarea_lozoyensis | 0.000 | 0.000 | 0.000 | 0.022 | 0.020 | 0 | 0 | 0 | 0 | 0.0055 | 0 |
| Ascomycota | Sordariomycetes | Hypocreales | Bionectriaceae | Gliomastix_polychroma | 0.000 | 0.000 | 0.000 | 0.000 | 0.000 | 0.000 | 0.000 | 0.000 | 0.000 | 0.000 | 0.000 |
| Mucoromycota | Unclassified | Unclassified | Unclassified | Glomeromycota_sp. | 0.017 | 0.000 | 0.000 | 0.000 | 0.000 | 0.000 | 0.000 | 0.000 | 0.000 | 0.000 | 0.000 |
| Mucoromycota | Glomeromycetes | Glomerales | Glomeraceae | Glomus_compressum | 0.062 | 0.000 | 0.000 | 0.000 | 0.000 | 0.000 | 0.000 | 0.000 | 0.000 | 0.000 | 0.000 |
| Mucoromycota | Glomeromycetes | Glomerales | Glomeraceae | Glomus_eburneum | 0.000 | 0.000 | 0.000 | 0.000 | 0.000 | 0.000 | 0.000 | 0.000 | 0.000 | 0.000 | 0.000 |
| Mucoromycota | Glomeromycetes | Glomerales | Glomeraceae | Glomus_indicum | 0.003 | 0.000 | 0.000 | 0.000 | 0.000 | 0.000 | 0.000 | 0.000 | 0.000 | 0.000 | 0.000 |
| Mucoromycota | Glomeromycetes | Glomerales | Glomeraceae | Glomus_intraradices | 0.007 | 0.000 | 0.000 | 0.003 | 0.000 | 0.000 | 0.000 | 0.000 | 0.000 | 0.000 | 0.000 |
| Mucoromycota | Glomeromycetes | Glomerales | Glomeraceae | Glomus_mosseae | 0.003 | 0.000 | 0.000 | 0.000 | 0.000 | 0.000 | 0.000 | 0.000 | 0.000 | 0.000 | 0.000 |
| Mucoromycota | Glomeromycetes | Glomerales | Glomeraceae | Glomus_proliferum | 0.018 | 0.025 | 0.000 | 0.002 | 0.000 | 0.000 | 0.000 | 0.000 | 0.000 | 0.000 | 0.000 |
| Mucoromycota | Glomeromycetes | Glomerales | Glomeraceae | Glomus_sp. | 0.001 | 0.000 | 0.000 | 0.000 | 0.000 | 0.000 | 0.000 | 0.000 | 0.000 | 0.000 | 0.000 |
| Basidiomycota | Agaricomycetes | Boletales | Gomphidiaceae | Gomphidius_roseus | 0.000 | 0.000 | 0.000 | 0.000 | 0.000 | 0.000 | 0.000 | 0.000 | 0.000 | 0.000 | 0.000 |
| Mucoromycota | Mucoromycetes | Mucorales | Cunninghamellaceae | Gongronella_butleri | 0.004 | 0.000 | 0.000 | 0.002 | 0.000 | 0.000 | 0.000 | 0.000 | 0.000 | 0.000 | 0.000 |
| Ascomycota | Sordariomycetes | Microascales | Graphiaceae | Graphium_jumulu | 0.013 | 0.000 | 0.000 | 0.000 | 0.000 | 0 | 0 | 0 | 0 | 0 | 0.06 |
| Ascomycota | Sordariomycetes | Ophiostomatales | Ophiostomataceae | Grosmannia_clavigera | 0.000 | 0.000 | 0.000 | 0.000 | 0.000 | 0.0035 | 0 | 0 | 0.0058 | 0 | 0 |
| Ascomycota | Lecanaromycetes | Teloschisatales | Teloschistaceae | Gyalolechia_flavorubescens | 0.046 | 0.025 | 0.000 | 0.000 | 0.000 | 0 | 0 | 0 | 0.0058 | 0 | 0 |
| Basidiomycota | Agaricomycetes | Agaricales | Cortinariaceae | Gymnopilus_suberis | 0.001 | 0.000 | 0.000 | 0.000 | 0.005 | 0.0035 | 0 | 0 | 0 | 0 | 0 |
| Ascomycota | Eurotiomycetes | Onygenales | Unclassified | Harorepupu_aotearoa | 0.000 | 0.000 | 0.000 | 0.005 | 0.000 | 0.000 | 0.000 | 0.000 | 0.000 | 0.000 | 0.000 |
| Zoopagomycota | Harpellomycetes | Harpellales | Harpellaceae | Harpellomyces_sp. | 0.001 | 0.000 | 0.000 | 0.000 | 0.000 | 0.000 | 0.000 | 0.000 | 0.000 | 0.000 | 0.000 |
| Basidiomycota | Agaricomycetes | Agaricales | Cortinariaceae | Hebeloma_hiemale | 0.000 | 0.000 | 0.000 | 0.000 | 0.000 | 0.0035 | 0 | 0 | 0 | 0 | 0 |
| Basidiomycota | Tremellomycetes | Filobasidiales | Filobasidiaceaea | Heterocephalacria_bachmannii | 0.000 | 0.000 | 0.000 | 0.000 | 0.000 | 0 | 0 | 0 | 0.0058 | 0 | 0 |
| Ascomycota | Sordariomycetes | Hypocreales | Ophiocordycipitaceae | Hirsutella_citriformis | 0.000 | 0.000 | 0.000 | 0.000 | 0.000 | 0 | 0 | 0 | 0.0058 | 0.0055 | 0 |
| Ascomycota | Sordariomycetes | Hypocreales | Ophiocordycipitaceae | Hirsutella_minnesotensis | 0.000 | 0.000 | 0.000 | 0.000 | 0.000 | 0.0143 | 0 | 0.0277 | 0.0232 | 0.0166 | 0.02 |
| Ascomycota | Sordariomycetes | Hypocreales | Ophiocordycipitaceae | Hirsutella_sp. | 0.000 | 0.000 | 0.000 | 0.000 | 0.000 | 0.000 | 0.000 | 0.000 | 0.000 | 0.000 | 0.000 |

**Table S8 continued.**

| | | | | | | | | | | | | | | | |
|---|---|---|---|---|---|---|---|---|---|---|---|---|---|---|---|
| Ascomycota | Dothideomycetes | Dothideales | Dothioraceae | Hormonema_dematioides | 0.001 | 0.000 | 0.000 | 0.000 | 0.000 | 0.000 | 0.000 | 0.000 | 0.000 | 0.000 | 0.000 |
| Ascomycota | Eurotiomycetes | Chaetothyriales | Unclassified | Hyalocladosporiella_tectonae | 0.000 | 0.000 | 0.000 | 0.000 | 0.000 | 0.0035 | 0 | 0 | 0 | 0 | 0 |
| Ascomycota | Pezizomycetes | Pezizales | Discinaceae | Hydnotrya_bailii | 0.000 | 0.000 | 0.000 | 0.000 | 0.000 | 0.000 | 0.000 | 0.000 | 0.000 | 0.000 | 0.000 |
| Ascomycota | Pezizomycetes | Pezizales | Discinaceae | Hydnotrya_tulasnei | 0.000 | 0.000 | 0.000 | 0.001 | 0.005 | 0.000 | 0.000 | 0.000 | 0.000 | 0.000 | 0.000 |
| Basidiomycota | Agaricomycetes | Boletales | Hygrophoropsidaceae | Hygrophoropsis_sp. | 0.003 | 0.000 | 0.000 | 0.017 | 0.000 | 0 | 0.0016 | 0 | 0 | 0.0055 | 0 |
| Ascomycota | Sordariomycetes | Hypocreales | Hypocreaceae | Hypocrea_koningii | 0.000 | 0.000 | 0.000 | 0.000 | 0.000 | 0 | 0 | 0 | 0 | 0.0055 | 0 |
| Ascomycota | Sordariomycetes | Hypocreales | Hypocreaceae | Hypocrea_lixii | 0.000 | 0.000 | 0.000 | 0.000 | 0.000 | 0.0176 | 0.0016 | 0.0222 | 0.0174 | 0.1325 | 0.02 |
| Ascomycota | Sordariomycetes | Hypocreales | Hypocreaceae | Hypocrea_lutea | 0.000 | 0.000 | 0.000 | 0.000 | 0.000 | 0 | 0 | 0 | 0.0058 | 0.0055 | 0 |
| Ascomycota | Sordariomycetes | Hypocreales | Hypocreaceae | Hypocrea_virens | 0.000 | 0.000 | 0.000 | 0.000 | 0.000 | 0.0035 | 0 | 0 | 0 | 0.0055 | 0 |
| Ascomycota | Sordariomycetes | Unclassified | Unclassified | Hypocreales_sp. | 0.000 | 0.000 | 0.000 | 0.000 | 0.000 | 0.0035 | 0 | 0.0111 | 0.0058 | 0.011 | 0.02 |
| Ascomycota | Sordariomycetes | Hypocreales | Hypocreaceae | Hypocreopsis_amplectens | 0.000 | 0.000 | 0.000 | 0.000 | 0.000 | 0 | 0 | 0.0055 | 0.0058 | 0 | 0 |
| Ascomycota | Sordariomycetes | Hypocreales | Hypocreaceae | Hypocreopsis_lichenoides | 0.000 | 0.000 | 0.000 | 0.000 | 0.000 | 0 | 0.0016 | 0 | 0 | 0 | 0 |
| Ascomycota | Sordariomycetes | Hypocreales | Hypocreaceae | Hypocreopsis_rhododendri | 0.000 | 0.000 | 0.000 | 0.000 | 0.000 | 0 | 0 | 0 | 0.0058 | 0 | 0 |
| Ascomycota | Sordariomycetes | Xylariales | Hypoxylaceae | Hypoxylon_sp. | 0.000 | 0.000 | 0.000 | 0.000 | 0.000 | 0 | 0 | 0.0055 | 0.0058 | 0.0055 | 0 |
| Ascomycota | Sordariomycetes | Hypocreales | Nectriaceae | Ilyonectria_macrodidyma | 0.000 | 0.000 | 0.000 | 0.000 | 0.000 | 0 | 0 | 0 | 0 | 0.011 | 0.02 |
| Ascomycota | Sordariomycetes | Hypocreales | Nectriaceae | Ilyonectria_radicicola | 0.000 | 0.000 | 0.000 | 0.000 | 0.000 | 0 | 0 | 0.0055 | 0.0058 | 0 | 0 |
| Basidiomycota | Agaricomycetes | Agaricales | Inocybaceae | Inocybe_lacera | 0.001 | 0.000 | 0.000 | 0.000 | 0.000 | 0.000 | 0.000 | 0.000 | 0.000 | 0.000 | 0.000 |
| Ascomycota | Sordariomycetes | Hypocreales | Cordycipitaceae | Isaria_farinosa | 0.000 | 0.000 | 0.000 | 0.000 | 0.000 | 0 | 0 | 0.0111 | 0.0058 | 0.0055 | 0.02 |
| Ascomycota | Sordariomycetes | Hypocreales | Cordycipitaceae | Isaria_poprawskii | 0.000 | 0.000 | 0.000 | 0.000 | 0.000 | 0 | 0 | 0 | 0 | 0 | 0.02 |
| Ascomycota | Sordariomycetes | Hypocreales | Cordycipitaceae | Isaria_tenuipes | 0.000 | 0.000 | 0.000 | 0.001 | 0.000 | 0.000 | 0.000 | 0.000 | 0.000 | 0.000 | 0.000 |
| Ascomycota | Saccharomycetes | Saccharomycetales | Pichiaceae | Issatchenkia_orientalis | 0.000 | 0.000 | 0.000 | 0.002 | 0.000 | 0.000 | 0.000 | 0.000 | 0.000 | 0.000 | 0.000 |
| Basidiomycota | Exobasidiomycetes | Microstromatales | Unclassified | Jaminaea_angkoriensis | 0.001 | 0.000 | 0.000 | 0.000 | 0.000 | 0.000 | 0.000 | 0.000 | 0.000 | 0.000 | 0.000 |
| Ascomycota | Saccharomycetes | Saccharomycetales | Kluyveromyces | Kluyveromyces_dobzhanskii | 0.001 | 0.000 | 0.000 | 0.000 | 0.000 | 0.000 | 0.000 | 0.000 | 0.000 | 0.000 | 0.000 |
| Ascomycota | Sordariomycetes | Xylariales | Xylariaceae | Kretzschmaria_deusta | 0.003 | 0.000 | 0.000 | 0.000 | 0.000 | 0.000 | 0.000 | 0.000 | 0.000 | 0.000 | 0.000 |
| Basidiomycota | Agaricomycetes | Agaricales | Tricholomataceae | Laccaria_bicolor | 0.009 | 0.000 | 0.000 | 0.000 | 0.000 | 0.0072 | 0.0016 | 0 | 0 | 0 | 0 |
| Basidiomycota | Agaricomycetes | Agaricales | Tricholomataceae | Laccaria_fibrillosa | 0.000 | 0.000 | 0.000 | 0.000 | 0.000 | 0 | 0 | 0 | 0.0058 | 0 | 0 |
| Basidiomycota | Agaricomycetes | Agaricales | Tricholomataceae | Laccaria_masoniae | 0.007 | 0.000 | 0.006 | 0.000 | 0.000 | 0.004 | 0.000 | 0.000 | 0.000 | 0.000 | 0.000 |
| Ascomycota | Saccharomycetes | Saccharomycetales | Saccharomycetaceae | Lachancea_nothofagi | 0.003 | 0.000 | 0.000 | 0.001 | 0.010 | 0.000 | 0.000 | 0.000 | 0.000 | 0.000 | 0.000 |
| Ascomycota | Saccharomycetes | Saccharomycetales | Saccharomycetaceae | Lachancea_sp. | 0.003 | 0.000 | 0.000 | 0.000 | 0.000 | 0.000 | 0.000 | 0.000 | 0.000 | 0.000 | 0.000 |
| Ascomycota | Dothideomycetes | Botryosphaeriales | Botryosphaeriaceae | Lasiodiplodia_crassispora | 0.001 | 0.000 | 0.006 | 0.000 | 0.000 | 0 | 0 | 0.0111 | 0.0058 | 0 | 0.02 |
| Ascomycota | Dothideomycetes | Botryosphaeriales | Botryosphaeriaceae | Lasiodiplodia_pseudotheobromae | 0.003 | 0.000 | 0.000 | 0.000 | 0.000 | 0 | 0.0049 | 0.0166 | 0.029 | 0.011 | 0.1601 |
| Ascomycota | Dothideomycetes | Botryosphaeriales | Botryosphaeriaceae | Lasiodiplodia_theobromae | 0.007 | 0.000 | 0.000 | 0.000 | 0.000 | 0 | 0.0115 | 0.0055 | 0.0174 | 0.0166 | 0.04 |

**Table S8 continued.**

| | | | | | | | | | | | | | | | |
|---|---|---|---|---|---|---|---|---|---|---|---|---|---|---|---|
| Ascomycota | Sordariomycetes | Hypocreales | Cordycipitaceae | Lecanicillium_psalliotae | 0.000 | 0.000 | 0.017 | 0.000 | 0.000 | 0.000 | 0.000 | 0.000 | 0.000 | 0.000 | 0.000 |
| Ascomycota | Sordariomycetes | Hypocreales | Cordycipitaceae | Lecanicillium_sp._569 | 0.000 | 0.000 | 0.000 | 0.000 | 0.000 | 0 | 0 | 0.0055 | 0.0174 | 0 | 0 |
| Basidiomycota | Agaricomycetes | Agaricales | Omphalotaceae | Lentinula_edodes | 0.003 | 0.000 | 0.000 | 0.000 | 0.000 | 0.000 | 0.000 | 0.000 | 0.000 | 0.000 | 0.000 |
| Basidiomycota | Agaricomycetes | Gloeophyllales | Gloeophyllaceae | Lentinus_cyathiformis | 0.000 | 0.000 | 0.000 | 0.000 | 0.005 | 0.0037 | 0 | 0 | 0 | 0 | 0.02 |
| Basidiomycota | Agaricomycetes | Gloeophyllales | Gloeophyllaceae | Lentinus_sajor-caju | 0.001 | 0.000 | 0.000 | 0.000 | 0.000 | 0.000 | 0.000 | 0.000 | 0.000 | 0.000 | 0.000 |
| Ascomycota | Unclassified | Unclassified | Unclassified | Leotiomycetes_sp. | 0.000 | 0.000 | 0.000 | 0.000 | 0.000 | 0 | 0 | 0.0055 | 0.0058 | 0 | 0 |
| Ascomycota | Dothideomycetes | Mytilinidiales | Argynnaceae | Lepidopterella_palustris | 0.003 | 0.000 | 0.000 | 0.000 | 0.000 | 0.000 | 0.000 | 0.000 | 0.000 | 0.000 | 0.000 |
| Basidiomycota | Agaricomycetes | Agaricales | Unclassified | Lepiotaceae_sp. | 0.000 | 0.000 | 0.000 | 0.000 | 0.000 | 0.0035 | 0.0016 | 0 | 0 | 0.0055 | 0 |
| Ascomycota | Sordariomycetes | Hypocreales | Cordycipitaceae | Leptobacillium_leptobactrum | 0.000 | 0.000 | 0.000 | 0.000 | 0.000 | 0 | 0 | 0.0055 | 0 | 0 | 0 |
| Ascomycota | Dothideomycetes | Pleosporales | Leptosphaeriaceae | Leptosphaeria_maculans | 0.000 | 0.000 | 0.000 | 0.000 | 0.000 | 0.004 | 0.000 | 0.000 | 0.000 | 0.000 | 0.000 |
| Ascomycota | Dothideomycetes | Pleosporales | Leptosphaeriaceae | Leptosphaeria_sp. | 0.000 | 0.000 | 0.000 | 0.001 | 0.000 | 0.0035 | 0 | 0 | 0 | 0 | 0 |
| Ascomycota | Dothideomycetes | Pleosporales | Astrosphaeriellaceae | Leptosphaerulina_chartarum | 0.000 | 0.000 | 0.000 | 0.001 | 0.000 | 0.000 | 0.000 | 0.000 | 0.000 | 0.000 | 0.000 |
| Ascomycota | Dothideomycetes | Pleosporales | Didymellaceae | Leptosphaerulina_trifolii | 0.000 | 0.000 | 0.000 | 0.000 | 0.000 | 0.000 | 0.000 | 0.000 | 0.000 | 0.000 | 0.000 |
| Basidiomycota | Agaricomycetes | Agaricales | Agaricaceae | Leucoagaricus_gongylophorus | 0.000 | 0.000 | 0.006 | 0.000 | 0.000 | 0.004 | 0.000 | 0.000 | 0.000 | 0.000 | 0.000 |
| Basidiomycota | Agaricomycetes | Agaricales | Agaricaceae | Leucoagaricus_sp. | 0.000 | 0.000 | 0.006 | 0.000 | 0.000 | 0.0035 | 0.0033 | 0 | 0 | 0.0055 | 0 |
| Basidiomycota | Agaricomycetes | Agaricales | Agaricaceae | Leucocoprinus_sp. | 0.009 | 0.000 | 0.006 | 0.005 | 0.005 | 0.004 | 0.000 | 0.000 | 0.000 | 0.000 | 0.000 |
| Chytridiomycota | Chytridiomycetes | Lobulomycetales | Lobulomyces | Lobulomyces_sp. | 0.001 | 0.000 | 0.000 | 0.000 | 0.000 | 0.000 | 0.000 | 0.000 | 0.000 | 0.000 | 0.000 |
| Ascomycota | Sordariomycetes | Microascales | Microascaceae | Lomentospora_prolificans | 0.001 | 0.000 | 0.000 | 0.000 | 0.000 | 0 | 0.0033 | 0 | 0 | 0 | 0 |
| Basidiomycota | Agaricomycetes | Agaricales | Lyophyllaceae | Lyophyllum_semitale | 0.000 | 0.000 | 0.006 | 0.000 | 0.000 | 0.000 | 0.000 | 0.000 | 0.000 | 0.000 | 0.000 |
| Ascomycota | Sordariomycetes | Magnaporthales | Pyriculariaceae | Magnaporthe_grisea | 0.003 | 0.000 | 0.017 | 0.000 | 0.015 | 0.0144 | 0.0033 | 0.0333 | 0.0697 | 0.0221 | 0.04 |
| Ascomycota | Sordariomycetes | Magnaporthales | Pyriculariaceae | Magnaporthe_oryzae | 0.000 | 0.000 | 0.006 | 0.001 | 0.020 | 0 | 0.0033 | 0.0388 | 0.0871 | 0.0442 | 0 |
| Ascomycota | Sordariomycetes | Magnaporthales | Pyriculariaceae | Magnaporthe_poae | 0.000 | 0.000 | 0.000 | 0.000 | 0.000 | 0.000 | 0.000 | 0.000 | 0.000 | 0.000 | 0.000 |
| Basidiomycota | Agaricomycetes | Agaricales | Marasmiaceae | Marasmius_rotula | 0.000 | 0.000 | 0.006 | 0.000 | 0.000 | 0.000 | 0.000 | 0.000 | 0.000 | 0.000 | 0.000 |
| Ascomycota | Leotiomycetes | Helotiales | Dermataceae | Marssonina_brunnea | 0.004 | 0.000 | 0.006 | 0.011 | 0.025 | 0 | 0 | 0 | 0.0058 | 0 | 0 |
| Ascomycota | Pucciniomycetes | Pucciniales | Melamsporaceae | Melampsora_larici-populina | 0.007 | 0.000 | 0.000 | 0.000 | 0.000 | 0 | 0 | 0 | 0 | 0 | 0.02 |
| Basidiomycota | Ustilaginomycetes | Ustilaginales | Ustilaginaceae | Melanopsichium_pennsylvanicum | 0.003 | 0.000 | 0.006 | 0.000 | 0.000 | 0.000 | 0.000 | 0.000 | 0.000 | 0.000 | 0.000 |
| Basidiomycota | Sordariomycetes | Hypocreales | Clavicipitaceae | Metarhizium_anisopliae | 0.000 | 0.000 | 0.000 | 0.000 | 0.005 | 0.0107 | 0.0016 | 0.0166 | 0.029 | 0.0386 | 0.1 |
| Basidiomycota | Sordariomycetes | Hypocreales | Clavicipitaceae | Metarhizium_robertsii | 0.000 | 0.000 | 0.000 | 0.000 | 0.000 | 0 | 0 | 0.0111 | 0 | 0.0055 | 0 |
| Ascomycota | Microbotryomycetes | Microbotryales | Microbotryaceae | Microbotryum_heliospermae | 0.000 | 0.000 | 0.000 | 0.000 | 0.000 | 0.000 | 0.000 | 0.000 | 0.000 | 0.000 | 0.000 |
| Ascomycota | Microbotryomycetes | Microbotryales | Microbotryaceae | Microbotryum_lagerheimii | 0.000 | 0.000 | 0.000 | 0.000 | 0.005 | 0.000 | 0.000 | 0.000 | 0.000 | 0.000 | 0.000 |
| Ascomycota | Microbotryomycetes | Microbotryales | Microbotryaceae | Microbotryum_lychnidis-dioicae | 0.000 | 0.000 | 0.000 | 0.000 | 0.000 | 0.000 | 0.000 | 0.000 | 0.000 | 0.000 | 0.000 |
| Ascomycota | Dothideomycetes | Botryosphaeriales | Botryosphaeriaceae | Microdiplodia_sp. | 0.000 | 0.000 | 0.000 | 0.000 | 0.000 | 0 | 0 | 0 | 0 | 0.0055 | 0 |

**Table S8 continued.**

| | | | | | | | | | | | | | | | |
|---|---|---|---|---|---|---|---|---|---|---|---|---|---|---|---|
| Ascomycota | Sordariomycetes | Xylariales | Microdochiaceae | Microdochium_bolleyi | 0.000 | 0.000 | 0.000 | 0.000 | 0.000 | 0 | 0 | 0 | 0.0058 | 0 | 0 |
| Ascomycota | Sordariomycetes | Xylariales | Microdochiaceae | Microdochium_nivale | 0.000 | 0.000 | 0.000 | 0.000 | 0.000 | 0 | 0 | 0 | 0.0058 | 0 | 0 |
| Ascomycota | Dothideomycetes | Pleosporales | Unclassified | Microsphaeropsis_arundinis | 0.003 | 0.000 | 0.000 | 0.000 | 0.000 | 0 | 0 | 0.0055 | 0 | 0 | 0 |
| Ascomycota | Agaricomycetes | Cantharellales | Unclassified | Minimedusa_polyspora | 0.000 | 0.000 | 0.000 | 0.002 | 0.005 | 0.000 | 0.000 | 0.000 | 0.000 | 0.000 | 0.000 |
| Ascomycota | Eurotiomycetes | Eurotiales | Aspergillaceae | Monascus_pallens | 0.000 | 0.000 | 0.000 | 0.000 | 0.000 | 0.000 | 0.000 | 0.000 | 0.000 | 0.000 | 0.000 |
| Basidiomycota | Sordariomycetes | Hypocreales | Niessliaceae | Monocillium_gamsii | 0.000 | 0.000 | 0.000 | 0.000 | 0.000 | 0 | 0 | 0 | 0 | 0 | 0.02 |
| Basidiomycota | Sordariomycetes | Hypocreales | Niessliaceae | Monocillium_ligusticum | 0.000 | 0.000 | 0.000 | 0.000 | 0.000 | 0 | 0 | 0 | 0 | 0 | 0.02 |
| Basidiomycota | Dothideomycetes | Pleosporales | Unclassified | Montagnulaceae_sp. | 0.000 | 0.000 | 0.000 | 0.000 | 0.000 | 0 | 0 | 0 | 0.0058 | 0.0221 | 0 |
| Ascomycota | Pezizomycetes | Pezizales | Morchellaceae | Morchella_crassipes | 0.001 | 0.000 | 0.000 | 0.000 | 0.000 | 0.000 | 0.000 | 0.000 | 0.000 | 0.000 | 0.000 |
| Ascomycota | Mortierellomycetes | Mortierellales | Mortierellaceae | Mortierella_alpina | 0.001 | 0.000 | 0.000 | 0.000 | 0.000 | 0.000 | 0.000 | 0.000 | 0.000 | 0.000 | 0.000 |
| Ascomycota | Mortierellomycetes | Mortierellales | Mortierellaceae | Mortierella_gamsii | 0.000 | 0.000 | 0.000 | 0.000 | 0.000 | 0.000 | 0.000 | 0.000 | 0.000 | 0.000 | 0.000 |
| Ascomycota | Mortierellomycetes | Mortierellales | Mortierellaceae | Mortierella_hyalina | 0.000 | 0.000 | 0.000 | 0.001 | 0.000 | 0.000 | 0.000 | 0.000 | 0.000 | 0.000 | 0.000 |
| Ascomycota | Mortierellomycetes | Mortierellales | Mortierellaceae | Mortierella_sp. | 0.000 | 0.000 | 0.011 | 0.000 | 2.073 | 0.004 | 0.000 | 0.000 | 0.000 | 0.000 | 0.000 |
| Basidiomycota | Mortierellomycetes | Unclassified | Unclassified | Mortierellomycotina_sp. | 0.001 | 0.000 | 0.000 | 0.000 | 0.000 | 0.000 | 0.000 | 0.000 | 0.000 | 0.000 | 0.000 |
| Ascomycota | Mucoromycetes | Mucorales | Mucoraceae | Mucor_ambiguus | 0.000 | 0.000 | 0.006 | 0.000 | 0.005 | 0.0035 | 0 | 0 | 0 | 0 | 0 |
| Ascomycota | Mucoromycetes | Mucorales | Mucoraceae | Mucor_circinelloides_f._circinelloides | 0.001 | 0.000 | 0.000 | 0.000 | 0.000 | 0.000 | 0.000 | 0.000 | 0.000 | 0.000 | 0.000 |
| Ascomycota | Mucoromycetes | Mucorales | Mucoraceae | Mucor_fragilis | 0.004 | 0.000 | 0.000 | 0.000 | 0.000 | 0.000 | 0.000 | 0.000 | 0.000 | 0.000 | 0.000 |
| Ascomycota | Mucoromycetes | Mucorales | Mucoraceae | Mucor_hiemalis | 0.001 | 0.000 | 0.000 | 0.000 | 0.000 | 0.000 | 0.000 | 0.000 | 0.000 | 0.000 | 0.000 |
| Ascomycota | Mucoromycetes | Mucorales | Mucoraceae | Mucor_sp. | 0.013 | 0.000 | 0.000 | 0.000 | 0.000 | 0 | 0 | 0.0277 | 0.0522 | 0.0055 | 0 |
| Mucoromycota | Mucoromycetes | Mucorales | Unclassified | Mucorales_sp. | 0.000 | 0.000 | 0.000 | 0.000 | 0.000 | 0.000 | 0.000 | 0.000 | 0.000 | 0.000 | 0.000 |
| Mucoromycota | Sordariomycetes | Xylariales | Unclassified | Muscodor_albus | 0.000 | 0.000 | 0.000 | 0.000 | 0.000 | 0.0035 | 0 | 0.0166 | 0.0174 | 0.0055 | 0 |
| Mucoromycota | Sordariomycetes | Xylariales | Unclassified | Muscodor_sp. | 0.000 | 0.000 | 0.006 | 0.000 | 0.000 | 0 | 0 | 0.0222 | 0.0116 | 0.0221 | 0.06 |
| Mucoromycota | Sordariomycetes | Sordariales | Chaetomiaceae | Myceliophthora_thermophila | 0.000 | 0.000 | 0.000 | 0.000 | 0.000 | 0 | 0 | 0 | 0 | 0.0055 | 0 |
| Mucoromycota | Agaricomycetes | Agaricales | Mycenaceae | Mycena_galopus | 0.001 | 0.000 | 0.000 | 0.000 | 0.000 | 0.000 | 0.000 | 0.000 | 0.000 | 0.000 | 0.000 |
| Mucoromycota | Agaricomycetes | Agaricales | Omphalotaceae | Mycetinis_scorodonius | 0.000 | 0.000 | 0.000 | 0.000 | 0.000 | 0.0037 | 0.0016 | 0 | 0 | 0 | 0 |
| Mucoromycota | Sordariomycetes | Hypocreales | Hypocreaceae | Mycogone_perniciosa | 0.000 | 0.000 | 0.000 | 0.000 | 0.000 | 0 | 0 | 0.0055 | 0 | 0.011 | 0 |
| Mucoromycota | Dothideomycetes | Mycosphaerellales | Mycosphaerellaceae | Mycosphaerella_sp. | 0.001 | 0.000 | 0.000 | 0.000 | 0.000 | 0.000 | 0.000 | 0.000 | 0.000 | 0.000 | 0.000 |
| Mucoromycota | Dothiodeomycetes | Capnodiales | Mycosphaerellaceae | Mycovellosiella_phaseoli | 0.000 | 0.000 | 0.000 | 0.000 | 0.000 | 0 | 0 | 0 | 0 | 0.0055 | 0 |
| Mucoromycota | Agaricomycetes | Geastrales | Geastraceae | Myriostoma_calongei | 0.001 | 0.000 | 0.000 | 0.000 | 0.000 | 0.000 | 0.000 | 0.000 | 0.000 | 0.000 | 0.000 |
| Mucoromycota | Dothideomycetes | Valsariales | Valsariaceae | Myrmaecium_fulvopruinatum | 0.000 | 0.050 | 0.000 | 0.000 | 0.000 | 0.000 | 0.000 | 0.000 | 0.000 | 0.000 | 0.000 |
| Ascomycota | Dothideomycetes | Valsariales | Valsariaceae | Myrothecium_gramineum | 0.000 | 0.000 | 0.000 | 0.002 | 0.040 | 0.000 | 0.000 | 0.000 | 0.000 | 0.000 | 0.000 |
| Ascomycota | Dothideomycetes | Valsariales | Valsariaceae | Myrothecium_roridum | 0.000 | 0.000 | 0.000 | 0.000 | 0.000 | 0.0035 | 0 | 0 | 0.0058 | 0 | 0 |

**Table S8 continued.**

| | | | | | | | | | | | | | | | |
|---|---|---|---|---|---|---|---|---|---|---|---|---|---|---|---|
| Ascomycota | Dothideomycetes | Valsariales | Valsariaceae | Myrothecium_sp. | 0.000 | 0.000 | 0.000 | 0.000 | 0.005 | 0 | 0 | 0.0055 | 0 | 0 | 0 |
| Basidiomycota | Sordariomycetes | Hypocreales | Nectriaceae | Nectria_cinnabarina | 0.000 | 0.000 | 0.000 | 0.000 | 0.000 | 0.0035 | 0 | 0 | 0 | 0 | 0 |
| Basidiomycota | Sordariomycetes | Hypocreales | Nectriaceae | Nectria_pseudopeziza | 0.000 | 0.000 | 0.000 | 0.000 | 0.000 | 0 | 0.0016 | 0 | 0 | 0.0055 | 0 |
| Ascomycota | Sordariomycetes | Hypocreales | Nectriaceae | Nectria_sp. | 0.000 | 0.000 | 0.000 | 0.000 | 0.000 | 0 | 0 | 0.0055 | 0.0058 | 0 | 0 |
| Ascomycota | Sordariomycetes | Xylariales | Xylariaceae | Nemania_diffusa | 0.000 | 0.000 | 0.000 | 0.000 | 0.000 | 0 | 0 | 0 | 0.0058 | 0 | 0 |
| Ascomycota | Sordariomycetes | Xylariales | Xylariaceae | Nemania_serpens | 0.000 | 0.000 | 0.011 | 0.000 | 0.000 | 0.0071 | 0.0016 | 0.0943 | 0.1277 | 0.0662 | 0.02 |
| Basidiomycota | Sordariomycetes | Xylariales | Xylariaceae | Nemania_sp. | 0.000 | 0.000 | 0.017 | 0.000 | 0.000 | 0 | 0.0049 | 0.0166 | 0.0232 | 0.0166 | 0 |
| Ascomycota | Sordariomycetes | Hypocreales | Nectriaceae | Neonectria_punicea | 0.000 | 0.000 | 0.000 | 0.000 | 0.000 | 0 | 0.0016 | 0 | 0.0116 | 0.0055 | 0 |
| Ascomycota | Sordariomycetes | Hypocreales | Nectriaceae | Neonectria_radicicola | 0.000 | 0.000 | 0.000 | 0.000 | 0.000 | 0 | 0 | 0.0111 | 0.0058 | 0 | 0.02 |
| Ascomycota | Sordariomycetes | Hypocreales | Nectriaceae | Neonectria_sp. | 0.000 | 0.000 | 0.000 | 0.000 | 0.000 | 0 | 0 | 0 | 0.0058 | 0 | 0 |
| Ascomycota | Eurotiomycetes | Eurotiales | Aspergillaceae | Neosartorya_fischeri | 0.001 | 0.050 | 0.000 | 0.000 | 0.000 | 0 | 0 | 0 | 0.0058 | 0 | 0 |
| Ascomycota | Eurotiomycetes | Eurotiales | Aspergillaceae | Neosartorya_udagawae | 0.001 | 0.000 | 0.000 | 0.001 | 0.005 | 0 | 0 | 0 | 0.0116 | 0 | 0.02 |
| Ascomycota | Sordariomycetes | Hypocreales | Clavicipitaceae | Neotyphodium_occultans | 0.000 | 0.000 | 0.000 | 0.000 | 0.000 | 0 | 0 | 0 | 0 | 0 | 0.02 |
| Ascomycota | Sordariomycetes | Sordariales | Sordariaceae | Neurospora_crassa | 0.005 | 0.000 | 0.000 | 0.000 | 0.005 | 0.000 | 0.000 | 0.000 | 0.000 | 0.000 | 0.000 |
| Ascomycota | Sordariomycetes | Sordariales | Sordariaceae | Neurospora_tetrasperma | 0.000 | 0.000 | 0.011 | 0.000 | 0.000 | 0.0035 | 0.0016 | 0.0111 | 0.029 | 0.0386 | 0.02 |
| Ascomycota | Sordariomycetes | Trichosphaeriales | Trichosphaeriaceae | Nigrospora_oryzae | 0.000 | 0.000 | 0.006 | 0.000 | 0.000 | 0 | 0 | 0.0166 | 0.0406 | 0.0055 | 0 |
| Ascomycota | Sordariomycetes | Trichosphaeriales | Trichosphaeriaceae | Nigrospora_sp. | 0.000 | 0.000 | 0.006 | 0.000 | 0.005 | 0 | 0.0016 | 0.0111 | 0.029 | 0.0276 | 0.02 |
| Ascomycota | Sordariomycetes | Trichosphaeriales | Trichosphaeriaceae | Nigrospora_sphaerica | 0.000 | 0.000 | 0.000 | 0.001 | 0.000 | 0 | 0 | 0.0055 | 0.0058 | 0.011 | 0 |
| Ascomycota | Sordariomycetes | Xylariales | Xylariaceae | Nodulisporium_sp. | 0.000 | 0.000 | 0.011 | 0.000 | 0.000 | 0 | 0.0033 | 0.0333 | 0.0464 | 0.0718 | 0.02 |
| Ascomycota | Dothideomycetes | Venturiales | Sympoventuriaceae | Ochroconis_humicola | 0.000 | 0.000 | 0.000 | 0.000 | 0.000 | 0.000 | 0.000 | 0.000 | 0.000 | 0.000 | 0.000 |
| Ascomycota | Leotiomycetes | Unclassified | Myxotrichaceae | Oidiodendron_maius | 0.000 | 0.000 | 0.000 | 0.000 | 0.000 | 0.0037 | 0 | 0 | 0 | 0.0055 | 0 |
| Ascomycota | Leotiomycetes | Unclassified | Myxotrichaceae | Oidiodendron_sp. | 0.000 | 0.000 | 0.000 | 0.000 | 0.000 | 0 | 0 | 0 | 0 | 0 | 0.02 |
| Ascomycota | Sordariomycetes | Hypocreales | Ophiocordycipitaceae | Ophiocordyceps_sinensis | 0.000 | 0.000 | 0.000 | 0.001 | 0.015 | 0.000 | 0.000 | 0.000 | 0.000 | 0.000 | 0.000 |
| Ascomycota | Sordariomycetes | Hypocreales | Ophiocordycipitaceae | Ophiocordyceps_sp. | 0.000 | 0.000 | 0.000 | 0.000 | 0.000 | 0 | 0 | 0.0055 | 0.0116 | 0 | 0.02 |
| Ascomycota | Sordariomycetes | Diaporthales | Gnomoniaceae | Ophiognomonia_clavigignenti-juglandacearum | 0.000 | 0.000 | 0.000 | 0.000 | 0.000 | 0 | 0 | 0.0055 | 0 | 0 | 0 |
| Ascomycota | Sordariomycetes | Ophiostomatales | Ophiostomataceae | Ophiostoma_eucalyptigena | 0.000 | 0.000 | 0.000 | 0.000 | 0.000 | 0.0035 | 0 | 0 | 0 | 0 | 0 |
| Ascomycota | Sordariomycetes | Ophiostomatales | Ophiostomataceae | Ophiostoma_novo-ulmi | 0.000 | 0.000 | 0.000 | 0.000 | 0.000 | 0 | 0 | 0.0111 | 0.0116 | 0.0055 | 0.02 |
| Ascomycota | Sordariomycetes | Ophiostomatales | Ophiostomataceae | Ophiostoma_pseudotsugae | 0.000 | 0.000 | 0.000 | 0.001 | 0.000 | 0.000 | 0.000 | 0.000 | 0.000 | 0.000 | 0.000 |
| Ascomycota | Sordariomycetes | Ophiostomatales | Ophiostomataceae | Ophiostoma_setosum | 0.000 | 0.000 | 0.000 | 0.000 | 0.000 | 0.0035 | 0 | 0 | 0 | 0 | 0 |
| Ascomycota | Neocallimastigomycetes | Neocallimastigales | Neocallimastigaceae | Orpinomyces_sp. | 0.007 | 0.000 | 0.000 | 0.000 | 0.000 | 0.000 | 0.000 | 0.000 | 0.000 | 0.000 | 0.000 |
| Ascomycota | Eurotiomycetes | Eurotiales | Thermoascaceae | Paecilomyces_carneus | 0.000 | 0.000 | 0.000 | 0.000 | 0.000 | 0.0035 | 0 | 0 | 0 | 0 | 0 |
| Ascomycota | Eurotiomycetes | Eurotiales | Thermoascaceae | Paecilomyces_cateniannulatus | 0.000 | 0.000 | 0.000 | 0.000 | 0.000 | 0.0035 | 0 | 0 | 0.0232 | 0.011 | 0.02 |

**Table S8 continued.**

| | | | | | | | | | | | | | | | |
|---|---|---|---|---|---|---|---|---|---|---|---|---|---|---|---|
| Ascomycota | Eurotiomycetes | Eurotiales | Thermoascaceae | Paecilomyces_lilacinus | 0.000 | 0.000 | 0.000 | 0.000 | 0.000 | 0 | 0 | 0 | 0 | 0.0055 | 0 |
| Ascomycota | Eurotiomycetes | Eurotiales | Thermoascaceae | Paecilomyces_penicillatus | 0.000 | 0.000 | 0.000 | 0.000 | 0.000 | 0.0141 | 0 | 0.0499 | 0.029 | 0.0221 | 0.02 |
| Ascomycota | Eurotiomycetes | Eurotiales | Thermoascaceae | Paecilomyces_sp. | 0.000 | 0.025 | 0.000 | 0.000 | 0.000 | 0 | 0.0016 | 0 | 0 | 0.011 | 0.02 |
| Ascomycota | Eurotiomycetes | Eurotiales | Thermoascaceae | Paecilomyces_variotii | 0.000 | 0.025 | 0.000 | 0.000 | 0.000 | 0.000 | 0.000 | 0.000 | 0.000 | 0.000 | 0.000 |
| Ascomycota | Eurotiomycetes | Eurotiales | Thermoascaceae | Paenibacillus_sp. | 0.000 | 0.000 | 0.000 | 0.000 | 0.000 | 0.0035 | 0 | 0 | 0 | 0 | 0.02 |
| Ascomycota | Glomeromycetes | Archaeosporales | Archaeosporaceae | Palaeospora_spainii | 0.000 | 0.000 | 0.000 | 0.000 | 0.000 | 0 | 0 | 0 | 0 | 0.0055 | 0 |
| Ascomycota | Dothideomycetes | Pleosporales | Massarinaceae | Paraconiothyrium_sporulosum | 0.000 | 0.000 | 0.000 | 0.000 | 0.000 | 0.0035 | 0 | 0 | 0 | 0 | 0 |
| Chytridiomycota | Sordariomycetes | Hypocreales | Nectriaceae | Penicillifer_martinii | 0.000 | 0.000 | 0.000 | 0.000 | 0.000 | 0 | 0 | 0 | 0 | 0.0055 | 0 |
| Ascomycota | Eurotiomycetes | Eurotiales | Aspergillaceae | Penicillium_brocae | 0.001 | 0.000 | 0.000 | 0.000 | 0.000 | 0.000 | 0.000 | 0.000 | 0.000 | 0.000 | 0.000 |
| Ascomycota | Eurotiomycetes | Eurotiales | Aspergillaceae | Penicillium_capsulatum | 0.001 | 0.025 | 0.000 | 0.051 | 0.050 | 0.000 | 0.000 | 0.000 | 0.000 | 0.000 | 0.000 |
| Ascomycota | Eurotiomycetes | Eurotiales | Aspergillaceae | Penicillium_citreonigrum | 0.000 | 0.000 | 0.000 | 0.006 | 0.000 | 0.000 | 0.000 | 0.000 | 0.000 | 0.000 | 0.000 |
| Ascomycota | Eurotiomycetes | Eurotiales | Aspergillaceae | Penicillium_commune | 0.000 | 0.000 | 0.000 | 0.001 | 0.000 | 0.000 | 0.000 | 0.000 | 0.000 | 0.000 | 0.000 |
| Ascomycota | Eurotiomycetes | Eurotiales | Aspergillaceae | Penicillium_crustosum | 0.000 | 0.025 | 0.000 | 0.000 | 0.000 | 0.000 | 0.000 | 0.000 | 0.000 | 0.000 | 0.000 |
| Ascomycota | Eurotiomycetes | Eurotiales | Aspergillaceae | Penicillium_digitatum | 0.001 | 0.000 | 0.000 | 0.000 | 0.000 | 0.000 | 0.000 | 0.000 | 0.000 | 0.000 | 0.000 |
| Ascomycota | Eurotiomycetes | Eurotiales | Aspergillaceae | Penicillium_janthinellum | 0.000 | 0.000 | 0.000 | 0.000 | 0.000 | 0.000 | 0.000 | 0.000 | 0.000 | 0.000 | 0.000 |
| Mucoromycota | Eurotiomycetes | Eurotiales | Aspergillaceae | Penicillium_oxalicum | 0.000 | 0.000 | 0.000 | 0.001 | 0.000 | 0.000 | 0.000 | 0.000 | 0.000 | 0.000 | 0.000 |
| Ascomycota | Eurotiomycetes | Eurotiales | Aspergillaceae | Penicillium_polonicum | 0.000 | 0.000 | 0.000 | 0.000 | 0.000 | 0.000 | 0.000 | 0.000 | 0.000 | 0.000 | 0.000 |
| Ascomycota | Eurotiomycetes | Eurotiales | Aspergillaceae | Penicillium_sclerotiorum | 0.001 | 0.000 | 0.000 | 0.000 | 0.000 | 0.004 | 0.000 | 0.000 | 0.000 | 0.000 | 0.000 |
| Ascomycota | Eurotiomycetes | Eurotiales | Aspergillaceae | Penicillium_simplicissimum | 0.000 | 0.000 | 0.000 | 0.000 | 0.000 | 0.000 | 0.000 | 0.000 | 0.000 | 0.000 | 0.000 |
| Ascomycota | Eurotiomycetes | Eurotiales | Aspergillaceae | Penicillium_singorense | 0.000 | 0.000 | 0.000 | 0.006 | 0.000 | 0.000 | 0.000 | 0.000 | 0.000 | 0.000 | 0.000 |
| Ascomycota | Eurotiomycetes | Eurotiales | Aspergillaceae | Penicillium_solitum | 0.001 | 0.000 | 0.000 | 0.002 | 0.000 | 0.000 | 0.000 | 0.000 | 0.000 | 0.000 | 0.000 |
| Ascomycota | Eurotiomycetes | Eurotiales | Aspergillaceae | Penicillium_sp. | 0.003 | 0.000 | 0.000 | 0.011 | 0.005 | 0.000 | 0.000 | 0.000 | 0.000 | 0.000 | 0.000 |
| Ascomycota | Eurotiomycetes | Eurotiales | Aspergillaceae | Penicillium_subrubescens | 0.000 | 0.000 | 0.000 | 0.001 | 0.000 | 0.000 | 0.000 | 0.000 | 0.000 | 0.000 | 0.000 |
| Ascomycota | Eurotiomycetes | Eurotiales | Aspergillaceae | Penicillium_waksmanii | 0.000 | 0.000 | 0.000 | 0.000 | 0.000 | 0.000 | 0.000 | 0.000 | 0.000 | 0.000 | 0.000 |
| Ascomycota | Agaricomycetes | Russulales | Peniophoraceae | Peniophora_sp. | 0.013 | 0.050 | 0.118 | 0.055 | 0.080 | 0.0853 | 0.0082 | 0.4326 | 0.5399 | 0.2153 | 0.1601 |
| Ascomycota | Agaricomycetes | Polyporales | Polyporaceae | Perenniporia_tephropora | 0.000 | 0.000 | 0.000 | 0.000 | 0.000 | 0 | 0.0016 | 0 | 0 | 0 | 0 |
| Ascomycota | Dothideomycetes | Pleosporales | Periconiaceae | Periconia_sp. | 0.001 | 0.000 | 0.000 | 0.001 | 0.000 | 0.000 | 0.000 | 0.000 | 0.000 | 0.000 | 0.000 |
| Ascomycota | Sordariomycetes | Xylariales | Sporocadaceae | Pestalotiopsis_fici | 0.001 | 0.000 | 0.000 | 0.003 | 0.000 | 0 | 0 | 0.0055 | 0 | 0 | 0 |
| Ascomycota | Sordariomycetes | Xylariales | Sporocadaceae | Pestalotiopsis_maculiformans | 0.000 | 0.000 | 0.000 | 0.000 | 0.000 | 0 | 0 | 0.0055 | 0 | 0.0055 | 0 |
| Ascomycota | Sordariomycetes | Xylariales | Sporocadaceae | Pestalotiopsis_neglecta | 0.000 | 0.000 | 0.000 | 0.000 | 0.000 | 0 | 0 | 0 | 0.0058 | 0 | 0 |
| Ascomycota | Sordariomycetes | Xylariales | Sporocadaceae | Pestalotiopsis_sp. | 0.001 | 0.000 | 0.006 | 0.000 | 0.000 | 0 | 0 | 0.0055 | 0.0232 | 0.0055 | 0.02 |
| Ascomycota | Dothideomycetes | Pleosporales | Didymellaceae | Peyronellaea_glomerata | 0.000 | 0.000 | 0.000 | 0.000 | 0.005 | 0.000 | 0.000 | 0.000 | 0.000 | 0.000 | 0.000 |

**Table S8 continued.**

| | | | | | | | | | | | | | | | |
|---|---|---|---|---|---|---|---|---|---|---|---|---|---|---|---|
| Ascomycota | Pezizomycetes | Unclassified | Unclassified | Pezizomycetes_sp. | 0.000 | 0.000 | 0.000 | 0.000 | 0.000 | 0.000 | 0.000 | 0.000 | 0.000 | 0.000 | 0.000 |
| Ascomycota | Sordariomycetes | Togniniales | Togniniaceae | Phaeoacremonium_parasiticum | 0.000 | 0.000 | 0.000 | 0.000 | 0.005 | 0.000 | 0.000 | 0.000 | 0.000 | 0.000 | 0.000 |
| Basidiomycota | Eurotiomycetes | Chaetothyriales | Trichomericeae | Phaeococcomyces_chersonesos | 0.000 | 0.000 | 0.000 | 0.000 | 0.005 | 0.000 | 0.000 | 0.000 | 0.000 | 0.000 | 0.000 |
| Basidiomycota | Dothideomycetes | Mycosphaerellales | Mycosphaerellaceae | Phaeophleospora_parsoniae | 0.000 | 0.000 | 0.000 | 0.000 | 0.000 | 0.000 | 0.000 | 0.000 | 0.000 | 0.000 | 0.000 |
| Ascomycota | Pucciniomycetes | Pucciniales | Phakosporaceae | Phakopsora_muscadiniae | 0.001 | 0.000 | 0.000 | 0.000 | 0.000 | 0.000 | 0.000 | 0.000 | 0.000 | 0.000 | 0.000 |
| Ascomycota | Agaricomycetes | Polyporales | Phanerochaetaceae | Phanerochaete_sordida | 0.000 | 0.000 | 0.000 | 0.003 | 0.010 | 0.004 | 0.000 | 0.000 | 0.000 | 0.000 | 0.000 |
| Ascomycota | Agaricomycetes | Polyporales | Phanerochaetaceae | Phanerochaete_sp. | 0.000 | 0.000 | 0.000 | 0.000 | 0.000 | 0.004 | 0.000 | 0.000 | 0.000 | 0.000 | 0.000 |
| Ascomycota | Agaricomycetes | Hymenochaetales | Hymenochaetaceae | Phellinus_noxius | 0.000 | 0.000 | 0.000 | 0.000 | 0.000 | 0.000 | 0.000 | 0.000 | 0.000 | 0.000 | 0.000 |
| Ascomycota | Sordariomycetes | Sordariales | Cephalothecaceae | Phialemonium_inflatum | 0.000 | 0.000 | 0.000 | 0.003 | 0.000 | 0.000 | 0.000 | 0.000 | 0.000 | 0.000 | 0.000 |
| Ascomycota | Sordariomycetes | Sordariales | Cephalothecaceae | Phialemonium_sp. | 0.000 | 0.000 | 0.000 | 0.000 | 0.000 | 0 | 0 | 0 | 0.0058 | 0 | 0 |
| Ascomycota | Leotiomycetes | Unclassified | Unclassified | Phialophora_attae | 0.001 | 0.000 | 0.000 | 0.000 | 0.000 | 0.000 | 0.000 | 0.000 | 0.000 | 0.000 | 0.000 |
| Ascomycota | Agaricomycetes | Polyporales | Meruliaceae | Phlebia_sp. | 0.000 | 0.000 | 0.000 | 0.000 | 0.000 | 0.000 | 0.000 | 0.000 | 0.000 | 0.000 | 0.000 |
| Ascomycota | Agaricomycetes | Polyporales | Meruliaceae | Phlebia_subserialis | 0.000 | 0.000 | 0.000 | 0.001 | 0.000 | 0.000 | 0.000 | 0.000 | 0.000 | 0.000 | 0.000 |
| Ascomycota | Agaricomycetes | Boletales | Boletinellaceae | Phlebopus_portentosus | 0.000 | 0.000 | 0.000 | 0.005 | 0.005 | 0.000 | 0.000 | 0.000 | 0.000 | 0.000 | 0.000 |
| Basidiomycota | Agaricomycetes | Agaricales | Strophariaceae | Pholiota_adiposa | 0.001 | 0.000 | 0.000 | 0.000 | 0.005 | 0.000 | 0.000 | 0.000 | 0.000 | 0.000 | 0.000 |
| Basidiomycota | Agaricomycetes | Agaricales | Strophariaceae | Pholiota_carbonaria | 0.000 | 0.000 | 0.000 | 0.000 | 0.000 | 0.004 | 0.000 | 0.000 | 0.000 | 0.000 | 0.000 |
| Ascomycota | Agaricomycetes | Agaricales | Strophariaceae | Pholiota_cf._lenta | 0.003 | 0.000 | 0.000 | 0.000 | 0.000 | 0.0178 | 0.0115 | 0 | 0 | 0 | 0 |
| Basidiomycota | Agaricomycetes | Agaricales | Strophariaceae | Pholiota_cf._spumosa | 0.000 | 0.000 | 0.000 | 0.000 | 0.005 | 0.0037 | 0.0033 | 0 | 0 | 0 | 0 |
| Ascomycota | Agaricomycetes | Agaricales | Strophariaceae | Pholiota_olivaceophylla | 0.001 | 0.000 | 0.006 | 0.000 | 0.000 | 0.004 | 0.000 | 0.000 | 0.000 | 0.000 | 0.000 |
| Ascomycota | Agaricomycetes | Agaricales | Strophariaceae | Pholiota_sp. | 0.004 | 0.000 | 0.000 | 0.000 | 0.000 | 0.0144 | 0.0164 | 0 | 0.0058 | 0 | 0 |
| Ascomycota | Dothideomycetes | Pleosporales | Didymellaceae | Phoma_herbarum | 0.000 | 0.000 | 0.000 | 0.001 | 0.000 | 0.000 | 0.000 | 0.000 | 0.000 | 0.000 | 0.000 |
| Basidiomycota | Dothideomycetes | Pleosporales | Didymellaceae | Phoma_medicaginis | 0.001 | 0.000 | 0.000 | 0.000 | 0.000 | 0.000 | 0.000 | 0.000 | 0.000 | 0.000 | 0.000 |
| Basidiomycota | Dothideomycetes | Pleosporales | Didymellaceae | Phoma_sp. | 0.000 | 0.000 | 0.006 | 0.004 | 0.025 | 0.0178 | 0 | 0 | 0 | 0 | 0 |
| Basidiomycota | Sordariomycetes | Diaporthales | Valsaceae | Phomopsis_vaccinii | 0.001 | 0.000 | 0.000 | 0.000 | 0.000 | 0.000 | 0.000 | 0.000 | 0.000 | 0.000 | 0.000 |
| Basidiomycota | Pucciniomycetes | Pucciniales | Phragmidiaceae | Phragmidium_tuberculatum | 0.000 | 0.000 | 0.000 | 0.000 | 0.000 | 0.000 | 0.000 | 0.000 | 0.000 | 0.000 | 0.000 |
| Basidiomycota | Sordariomycetes | Xylariales | Xylariaceae | Phylacia_bomba | 0.001 | 0.000 | 0.000 | 0.000 | 0.000 | 0 | 0 | 0 | 0.0058 | 0 | 0 |
| Basidiomycota | Sordariomycetes | Phyllachorales | Phyllachoraceae | Phyllachoraceae_sp. | 0.000 | 0.000 | 0.000 | 0.000 | 0.000 | 0 | 0 | 0.0055 | 0 | 0 | 0 |
| Basidiomycota | Agaricomycetes | Polyporales | Formitopsidaceae | Piptoporus_betulinus | 0.003 | 0.000 | 0.011 | 0.000 | 0.000 | 0.0107 | 0.0066 | 0 | 0 | 0.0055 | 0 |
| Basidiomycota | Agaricomycetes | Boletales | Pisolithaceae | Pisolithus_albus | 0.001 | 0.000 | 0.000 | 0.000 | 0.000 | 0.000 | 0.000 | 0.000 | 0.000 | 0.000 | 0.000 |
| Basidiomycota | Agaricomycetes | Boletales | Pisolithaceae | Pisolithus_tinctorius | 0.000 | 0.000 | 0.000 | 0.000 | 0.000 | 0.004 | 0.000 | 0.000 | 0.000 | 0.000 | 0.000 |
| Ascomycota | Dothideomycetes | Pleosporales | Astrosphaerellaceae | Pithomyces_chartarum | 0.000 | 0.000 | 0.000 | 0.000 | 0.000 | 0.000 | 0.000 | 0.000 | 0.000 | 0.000 | 0.000 |
| Ascomycota | Sordariomycetes | Glomerellales | Plectosphaerellaceae | Plectosphaerella | 0.000 | 0.025 | 0.000 | 0.000 | 0.000 | 0.000 | 0.000 | 0.000 | 0.000 | 0.000 | 0.000 |

**Table S8 continued.**

| | | | | | | | | | | | | | | | |
|---|---|---|---|---|---|---|---|---|---|---|---|---|---|---|---|
| Ascomycota | Dothideomycetes | Pleosporales | Pleosporaceae | Pleosporaceae_sp. | 0.000 | 0.000 | 0.000 | 0.000 | 0.005 | 0.000 | 0.000 | 0.000 | 0.000 | 0.000 | 0.000 |
| Ascomycota | Dothideomycetes | Pleosporales | Unclassified | Pleosporales_sp. | 0.003 | 0.025 | 0.000 | 0.001 | 0.000 | 0.000 | 0.000 | 0.000 | 0.000 | 0.000 | 0.000 |
| Basidiomycota | Agaricomycetes | Agaricales | Pleurotaceae | Pleurotus_cornucopiae | 0.001 | 0.000 | 0.000 | 0.000 | 0.000 | 0.000 | 0.000 | 0.000 | 0.000 | 0.000 | 0.000 |
| Ascomycota | Agaricomycetes | Agaricales | Pleurotaceae | Pleurotus_ostreatus | 0.000 | 0.000 | 0.000 | 0.000 | 0.005 | 0 | 0 | 0.0111 | 0 | 0 | 0 |
| Ascomycota | Sordariomycetes | Hypocreales | Clavicipitaceae | Pochonia_chlamydosporia | 0.000 | 0.000 | 0.006 | 0.000 | 0.000 | 0 | 0 | 0.0055 | 0 | 0.0055 | 0.06 |
| Basidiomycota | Sordariomycetes | Xylariales | Xylariaceae | Podosordaria_tulasnei | 0.000 | 0.000 | 0.158 | 0.000 | 0.000 | 0 | 0.0098 | 0.0776 | 1.3293 | 0.0552 | 0.08 |
| Basidiomycota | Sordariomycetes | Sordariales | Chaetomiaceae | Podospora_anserina | 0.000 | 0.000 | 0.000 | 0.000 | 0.000 | 0 | 0 | 0.0055 | 0 | 0 | 0 |
| Basidiomycota | Sordariomycetes | Xylariales | Unclassified | Polyancora_globosa | 0.004 | 0.000 | 0.000 | 0.000 | 0.000 | 0.000 | 0.000 | 0.000 | 0.000 | 0.000 | 0.000 |
| Ascomycota | Agaricomycetes | Polyporales | Unclassified | Polycephalomyces_prolificus | 0.000 | 0.000 | 0.000 | 0.000 | 0.000 | 0 | 0.0016 | 0.0277 | 0.0058 | 0.0055 | 0 |
| Ascomycota | Agaricomycetes | Polyporales | Unclassified | Polyporales_sp. | 0.000 | 0.000 | 0.000 | 0.000 | 0.000 | 0.000 | 0.000 | 0.000 | 0.000 | 0.000 | 0.000 |
| Ascomycota | Agaricomycetes | Polyporales | Polyporaceae | Polyporus_umbellatus | 0.001 | 0.000 | 0.000 | 0.205 | 0.135 | 0.000 | 0.000 | 0.000 | 0.000 | 0.000 | 0.000 |
| Ascomycota | Agaricomycetes | Polyporales | Dacryobolaceae | Postia_placenta | 0.000 | 0.000 | 0.000 | 0.000 | 0.000 | 0.000 | 0.000 | 0.000 | 0.000 | 0.000 | 0.000 |
| Basidiomycota | Dothideomycetes | Pleosporales | Sporormiaceae | Preussia_australis | 0.000 | 0.000 | 0.000 | 0.000 | 0.000 | 0.0035 | 0 | 0 | 0 | 0 | 0 |
| Basidiomycota | Dothideomycetes | Pleosporales | Sporormiaceae | Preussia_sp. | 0.000 | 0.000 | 0.000 | 0.006 | 0.000 | 0.000 | 0.000 | 0.000 | 0.000 | 0.000 | 0.000 |
| Ascomycota | Laboulbeniomycetes | Laboulbeniales | Laboulbeniaceae | Prolixandromyces_triandrus | 0.001 | 0.000 | 0.000 | 0.000 | 0.000 | 0.0035 | 0 | 0 | 0 | 0 | 0 |
| Ascomycota | Agaricomycetes | Agaricales | Psathyrellaceae | Psathyrella_candolleana | 0.000 | 0.000 | 0.000 | 0.000 | 0.000 | 0.000 | 0.000 | 0.000 | 0.000 | 0.000 | 0.000 |
| Ascomycota | Leotiomycetes | Unclassified | Pseudeurotiaceae | Pseudeurotium_bakeri | 0.001 | 0.050 | 0.000 | 0.000 | 0.000 | 0.000 | 0.000 | 0.000 | 0.000 | 0.000 | 0.000 |
| Ascomycota | Sordariomycetes | Hypocreales | Unclassified | Pseudoacremonium_sacchari | 0.000 | 0.000 | 0.000 | 0.000 | 0.000 | 0 | 0 | 0 | 0.0058 | 0 | 0 |
| Basidiomycota | Leotiomycetes | Unclassified | Pseudeurotiaceae | Pseudogymnoascus_destructans | 0.000 | 0.000 | 0.006 | 0.000 | 0.000 | 0 | 0 | 0 | 0 | 0 | 0.02 |
| Basidiomycota | Leotiomycetes | Unclassified | Pseudeurotiaceae | Pseudogymnoascus_pannorum | 0.000 | 0.000 | 0.000 | 0.000 | 0.000 | 0 | 0 | 0 | 0 | 0 | 0.02 |
| Basidiomycota | Leotiomycetes | Unclassified | Pseudeurotiaceae | Pseudogymnoascus_sp. | 0.000 | 0.000 | 0.000 | 0.000 | 0.000 | 0 | 0 | 0 | 0 | 0 | 0.02 |
| Basidiomycota | Sordariomycetes | Xylariales | Xylariaceae | Pseudoxylaria | 0.064 | 0.025 | 0.918 | 0.081 | 0.140 | 0.2205 | 2.6027 | 26.159 | 14.721 | 18.591 | 12.245 |
| Ascomycota | Ustilaginomycetes | Ustilaginales | Ustilaginaceae | Pseudozyma_hubeiensis | 0.001 | 0.000 | 0.000 | 0.000 | 0.000 | 0.000 | 0.000 | 0.000 | 0.000 | 0.000 | 0.000 |
| Ascomycota | Pucciniomycetes | Pucciniales | Pucciniaceae | Puccinia_caricina | 0.001 | 0.000 | 0.000 | 0.000 | 0.000 | 0.000 | 0.000 | 0.000 | 0.000 | 0.000 | 0.000 |
| Ascomycota | Pucciniomycetes | Pucciniales | Pucciniaceae | Puccinia_kusanoi | 0.001 | 0.000 | 0.000 | 0.006 | 0.000 | 0.000 | 0.000 | 0.000 | 0.000 | 0.000 | 0.000 |
| Basidiomycota | Pucciniomycetes | Pucciniales | Pucciniaceae | Puccinia_triticina | 0.000 | 0.000 | 0.000 | 0.006 | 0.000 | 0.000 | 0.000 | 0.000 | 0.000 | 0.000 | 0.000 |
| Ascomycota | Sordariomycetes | Hypocreales | Ophiocordycipitaceae | Purpureocillium_lilacinum | 0.001 | 0.000 | 0.006 | 0.000 | 0.000 | 0 | 0 | 0.0222 | 0.029 | 0.011 | 0.06 |
| Ascomycota | Unclassified | Unclassified | Unclassified | Pycnobasidium_sp. | 0.000 | 0.000 | 0.000 | 0.000 | 0.000 | 0.004 | 0.000 | 0.000 | 0.000 | 0.000 | 0.000 |
| Ascomycota | Exobasidiomycetes | Microstromatales | Quambalariaceae | Quambalaria_cyanescens | 0.024 | 0.000 | 0.000 | 0.000 | 0.000 | 0.000 | 0.000 | 0.000 | 0.000 | 0.000 | 0.000 |
| Ascomycota | Dothideomycetes | Mycosphaerellales | Mycosphaerellaceae | Ramularia_grevilleana | 0.000 | 0.000 | 0.000 | 0.000 | 0.000 | 0 | 0 | 0 | 0 | 0.0055 | 0 |
| Ascomycota | Dothideomycetes | Mycosphaerellales | Mycosphaerellaceae | Ramularia_pratensis | 0.000 | 0.000 | 0.000 | 0.000 | 0.000 | 0 | 0 | 0 | 0.0058 | 0 | 0 |
| Ascomycota | Eurotiomycetes | Eurotiales | Trichocomaceae | Rasamsonia_eburnea | 0.000 | 0.000 | 0.000 | 0.000 | 0.000 | 0.000 | 0.000 | 0.000 | 0.000 | 0.000 | 0.000 |

**Table S8 continued.**

| | | | | | | | | | | | | | | | |
|---|---|---|---|---|---|---|---|---|---|---|---|---|---|---|---|
| Basidiomycota | Agaricomycetes | Hymenochaetales | Unclassified | Resinicium_mutabile | 0.000 | 0.000 | 0.000 | 0.000 | 0.000 | 0.0035 | 0 | 0 | 0 | 0 | 0 |
| Basidiomycota | Agaricomycetes | Cantharellales | Ceratobasidiaceae | Rhizoctonia_fragariae | 0.001 | 0.000 | 0.006 | 0.000 | 0.000 | 0.0035 | 0.0016 | 0 | 0 | 0 | 0 |
| Basidiomycota | Agaricomycetes | Cantharellales | Ceratobasidiaceae | Rhizoctonia_solani | 0.007 | 0.000 | 0.000 | 0.000 | 0.000 | 0 | 0.0016 | 0 | 0 | 0 | 0 |
| Basidiomycota | Agaricomycetes | Cantharellales | Ceratobasidiaceae | Rhizoctonia_sp. | 0.005 | 0.000 | 0.000 | 0.000 | 0.000 | 0.022 | 0.0033 | 0 | 0 | 0 | 0 |
| Ascomycota | Mucoromycetes | Mucorales | Mucoraceae | Rhizomucor_variabilis | 0.000 | 0.000 | 0.000 | 0.000 | 0.000 | 0 | 0 | 0 | 0.2322 | 0 | 0 |
| Ascomycota | Glomeromycetes | Glomerales | Glomeraceae | Rhizophagus_irregularis | 0.148 | 0.000 | 0.045 | 0.017 | 0.075 | 0.0319 | 0.0066 | 0.0887 | 0.0755 | 0.0497 | 0.1601 |
| Ascomycota | Glomeromycetes | Glomerales | Glomeraceae | Rhizophagus_proliferus | 0.003 | 0.000 | 0.000 | 0.000 | 0.000 | 0.000 | 0.000 | 0.000 | 0.000 | 0.000 | 0.000 |
| Ascomycota | Chytridiomycetes | Rhizophydiales | Unclassified | Rhizophydiales_sp. | 0.164 | 0.025 | 0.000 | 0.000 | 0.000 | 0.000 | 0.000 | 0.000 | 0.000 | 0.000 | 0.000 |
| Ascomycota | Agaricomycetes | Boletales | Rhizopogonaceae | Rhizopogon_roseolus | 0.001 | 0.000 | 0.006 | 0.049 | 0.030 | 0.0035 | 0.0016 | 0 | 0 | 0 | 0 |
| Ascomycota | Mucoromycetes | Mucorales | Rhizopodaceae | Rhizopus_microsporus | 0.003 | 0.000 | 0.006 | 0.000 | 0.000 | 0.000 | 0.000 | 0.000 | 0.000 | 0.000 | 0.000 |
| Basidiomycota | Mucoromycetes | Mucorales | Rhizopodaceae | Rhizopus_oryzae | 0.005 | 0.000 | 0.000 | 0.001 | 0.000 | 0 | 0 | 0.0111 | 0 | 0 | 0 |
| Basidiomycota | Dothideomycetes | Pleosporales | Morosphaeriaceae | Rhizopycnis_vagum | 0.000 | 0.000 | 0.000 | 0.000 | 0.000 | 0.0071 | 0 | 0 | 0 | 0 | 0 |
| Ascomycota | Leotiomycetes | Helotiales | Hyaloscyphceae | Rhizoscyphus_ericae | 0.004 | 0.000 | 0.000 | 0.000 | 0.000 | 0 | 0 | 0 | 0 | 0.0055 | 0 |
| Ascomycota | Agaricomycetes | Agaricales | Tricholomatceae | Rhodocollybia_butyracea | 0.000 | 0.000 | 0.011 | 0.000 | 0.000 | 0.000 | 0.000 | 0.000 | 0.000 | 0.000 | 0.000 |
| Ascomycota | Microbotryomycetes | Sporidiobolales | Sporidiobolaceae | Rhodosporidiobolus_odoratus | 0.004 | 0.000 | 0.000 | 0.000 | 0.000 | 0.000 | 0.000 | 0.000 | 0.000 | 0.000 | 0.000 |
| Basidiomycota | Microbotryomycetes | Sporidiobolales | Sporidiobolaceae | Rhodotorula_sp. | 0.005 | 0.000 | 0.000 | 0.000 | 0.000 | 0.000 | 0.000 | 0.000 | 0.000 | 0.000 | 0.000 |
| Basidiomycota | Sordariomycetes | Xylariales | Sporocadaceae | Robillarda_sessilis | 0.004 | 0.000 | 0.000 | 0.000 | 0.000 | 0 | 0 | 0.0111 | 0 | 0.0055 | 0 |
| Basidiomycota | Sordariomycetes | Xylariales | Xylariaceae | Rosellinia_necatrix | 0.000 | 0.000 | 0.000 | 0.000 | 0.000 | 0 | 0 | 0.0111 | 0.0348 | 0.011 | 0 |
| Basidiomycota | Agaricomycetes | Boletales | Boletaceae | Rossbeevera_yunnanensis | 0.000 | 0.000 | 0.000 | 0.000 | 0.000 | 0.000 | 0.000 | 0.000 | 0.000 | 0.000 | 0.000 |
| Basidiomycota | Agaricomycetes | Russulales | Russulaceae | Russula_bicolor | 0.000 | 0.000 | 0.000 | 0.000 | 0.005 | 0.0035 | 0 | 0 | 0 | 0 | 0 |
| Basidiomycota | Agaricomycetes | Russulales | Russulaceae | Russula_decolorans | 0.000 | 0.000 | 0.000 | 0.001 | 0.000 | 0.000 | 0.000 | 0.000 | 0.000 | 0.000 | 0.000 |
| Basidiomycota | Agaricomycetes | Russulales | Russulaceae | Russula_rhodopoda | 0.000 | 0.000 | 0.000 | 0.000 | 0.000 | 0.000 | 0.000 | 0.000 | 0.000 | 0.000 | 0.000 |
| Basidiomycota | Agaricomycetes | Russulales | Russulaceae | Russula_sp. | 0.000 | 0.000 | 0.000 | 0.001 | 0.000 | 0.000 | 0.000 | 0.000 | 0.000 | 0.000 | 0.000 |
| Mucoromycota | Agaricomycetes | Russulales | Russulaceae | Russula_vinosa | 0.000 | 0.000 | 0.000 | 0.001 | 0.000 | 0.000 | 0.000 | 0.000 | 0.000 | 0.000 | 0.000 |
| Mucoromycota | Leotiomycetes | Helotiales | Rutstroemiaceae | Rutstroemia_sp. | 0.003 | 0.000 | 0.006 | 0.000 | 0.000 | 0.000 | 0.000 | 0.000 | 0.000 | 0.000 | 0.000 |
| Mucoromycota | Dothideomycetes | Pleosporales | Massarinaceae | Saccharicola_bicolor | 0.000 | 0.000 | 0.000 | 0.000 | 0.000 | 0.0425 | 0 | 0 | 0 | 0 | 0 |
| Mucoromycota | Saccharomycetes | Saccharomycetales | Saccharomycetaceae | Saccharomyces_cerevisiae | 0.003 | 0.000 | 0.000 | 0.000 | 0.000 | 0.000 | 0.000 | 0.000 | 0.000 | 0.000 | 0.000 |
| Mucoromycota | Mucoromycetes | Mucorales | Saksenaeaceae | Saksenaea_erythrospora | 0.000 | 0.000 | 0.000 | 0.001 | 0.000 | 0.000 | 0.000 | 0.000 | 0.000 | 0.000 | 0.000 |
| Mucoromycota | Mucoromycetes | Mucorales | Saksenaeaceae | Saksenaea_vasiformis | 0.000 | 0.000 | 0.000 | 0.000 | 0.000 | 0.000 | 0.000 | 0.000 | 0.000 | 0.000 | 0.000 |
| Mucoromycota | Eurotiomycetes | Chaetothyriales | Sarcinomyces | Sarcinomyces_sp. | 0.001 | 0.000 | 0.000 | 0.000 | 0.000 | 0.000 | 0.000 | 0.000 | 0.000 | 0.000 | 0.000 |
| Mucoromycota | Agaricomycetes | Thelephorales | Thelephoraceae | Sarcodon_lundellii | 0.000 | 0.000 | 0.000 | 0.000 | 0.015 | 0.000 | 0.000 | 0.000 | 0.000 | 0.000 | 0.000 |
| Chytridiomycota | Agaricomycetes | Thelephorales | Thelephoraceae | Sarcodon_squamosus | 0.000 | 0.000 | 0.000 | 0.000 | 0.000 | 0.0035 | 0 | 0 | 0 | 0 | 0 |

**Table S8 continued.**

| | | | | | | | | | | | | | | | |
|---|---|---|---|---|---|---|---|---|---|---|---|---|---|---|---|
| Basidiomycota | Sordariomycetes | Hypocreales | Sarocladiaceae | Sarocladium_bacillisporum | 0.000 | 0.000 | 0.000 | 0.000 | 0.000 | 0.000 | 0.000 | 0.000 | 0.000 | 0.000 | 0.000 |
| Basidiomycota | Sordariomycetes | Hypocreales | Sarocladiaceae | Sarocladium_implicatum | 0.000 | 0.000 | 0.000 | 0.000 | 0.000 | 0 | 0 | 0 | 0 | 0.0055 | 0 |
| Mucoromycota | Sordariomycetes | Hypocreales | Sarocladiaceae | Sarocladium_strictum | 0.000 | 0.000 | 0.000 | 0.000 | 0.005 | 0 | 0 | 0 | 0.0058 | 0 | 0 |
| Mucoromycota | Sordariomycetes | Hypocreales | Sarocladiaceae | Sarocladium_terricola | 0.000 | 0.000 | 0.000 | 0.000 | 0.020 | 0.000 | 0.000 | 0.000 | 0.000 | 0.000 | 0.000 |
| Mucoromycota | Saccharomycetes | Saccharomycetales | Pichiaceae | Saturnispora_gosingensis | 0.004 | 0.000 | 0.000 | 0.000 | 0.000 | 0.000 | 0.000 | 0.000 | 0.000 | 0.000 | 0.000 |
| Ascomycota | Sordariomycetes | Microascales | Microascaceae | Scedosporium_apiospermum | 0.005 | 0.000 | 0.000 | 0.000 | 0.000 | 0.0035 | 0 | 0 | 0 | 0 | 0 |
| Ascomycota | Agaricomycetes | Agaricales | Schizophyllaceae | Schizophyllum_commune | 0.000 | 0.000 | 0.000 | 0.002 | 0.000 | 0.000 | 0.000 | 0.000 | 0.000 | 0.000 | 0.000 |
| Ascomycota | Agaricomycetes | Agaricales | Schizophyllaceae | Schizophyllum_radiatum | 0.001 | 0.000 | 0.000 | 0.001 | 0.000 | 0.000 | 0.000 | 0.000 | 0.000 | 0.000 | 0.000 |
| Basidiomycota | Saccharomycetes | Saccharomycetales | Schwanniomyces | Schwanniomyces_polymorphus_var._polymorphus | 0.005 | 0.000 | 0.000 | 0.000 | 0.010 | 0 | 0 | 0 | 0.0639 | 0.0166 | 0 |
| Basidiomycota | Agaricomycetes | Boletales | Sclerodermataceae | Scleroderma_suthepense | 0.001 | 0.000 | 0.000 | 0.000 | 0.000 | 0.000 | 0.000 | 0.000 | 0.000 | 0.000 | 0.000 |
| Basidiomycota | Dothideomycetes | Pleosporales | Phaeosphaeriaceae | Sclerostagonospora_sp. | 0.000 | 0.000 | 0.000 | 0.001 | 0.000 | 0.000 | 0.000 | 0.000 | 0.000 | 0.000 | 0.000 |
| Ascomycota | Leotiomycetes | Helotiales | Sclerotiniaceae | Sclerotinia_borealis | 0.001 | 0.000 | 0.000 | 0.010 | 0.005 | 0 | 0.0016 | 0 | 0 | 0 | 0 |
| Ascomycota | Leotiomycetes | Helotiales | Sclerotiniaceae | Sclerotinia_sclerotiorum | 0.062 | 0.000 | 0.006 | 0.000 | 0.020 | 0 | 0 | 0.0055 | 0 | 0 | 0 |
| Basidiomycota | Dothideomycetes | Venturiales | Sympoventuriaceae | Scolecobasidium_sp. | 0.001 | 0.000 | 0.000 | 0.000 | 0.000 | 0 | 0 | 0 | 0.0058 | 0 | 0 |
| Basidiomycota | Sordariomycetes | Microascales | Microascaceae | Scopulariopsis_brevicaulis | 0.001 | 0.000 | 0.000 | 0.000 | 0.005 | 0.0035 | 0 | 0.0055 | 0 | 0.011 | 0 |
| Basidiomycota | Sordariomycetes | Microascales | Microascaceae | Scopulariopsis_sp. | 0.000 | 0.000 | 0.000 | 0.000 | 0.000 | 0 | 0 | 0 | 0 | 0.0055 | 0 |
| Basidiomycota | Sordariomycetes | Sordariales | Chaetomiaceae | Scytalidium_lignicola | 0.000 | 0.000 | 0.000 | 0.000 | 0.000 | 0 | 0 | 0 | 0 | 0 | 0.02 |
| Basidiomycota | Sordariomycetes | Sordariales | Chaetomiaceae | Scytalidium_thermophilum | 0.000 | 0.000 | 0.000 | 0.003 | 0.010 | 0.000 | 0.000 | 0.000 | 0.000 | 0.000 | 0.000 |
| Basidiomycota | Agaricomycetes | Sebacinales | Serendipitaceae | Sebacina_vermifera | 0.004 | 0.000 | 0.000 | 0.000 | 0.000 | 0.000 | 0.000 | 0.000 | 0.000 | 0.000 | 0.000 |
| Basidiomycota | Glomeromycetes | Glomerales | Glomeraceae | Septoglomus_sp. | 0.000 | 0.000 | 0.000 | 0.000 | 0.005 | 0.000 | 0.000 | 0.000 | 0.000 | 0.000 | 0.000 |
| Ascomycota | Agaricomycetes | Boletales | Serpulaceae | Serpula_lacrymans | 0.000 | 0.000 | 0.000 | 0.000 | 0.000 | 0.004 | 0.000 | 0.000 | 0.000 | 0.000 | 0.000 |
| Ascomycota | Sordariomycetes | Hypocreales | Cordycipitaceae | Simplicillium_lamellicola | 0.000 | 0.000 | 0.000 | 0.000 | 0.000 | 0 | 0 | 0.0055 | 0 | 0.0055 | 0 |
| Ascomycota | Lecanoromycetes | Pertusariales | Icmadophilaceae | Siphula_ceratites | 0.001 | 0.000 | 0.000 | 0.000 | 0.000 | 0.000 | 0.000 | 0.000 | 0.000 | 0.000 | 0.000 |
| Ascomycota | Unclassified | Unclassified | Unclassified | Slimacomyces_isiola | 0.001 | 0.000 | 0.006 | 0.000 | 0.000 | 0.000 | 0.000 | 0.000 | 0.000 | 0.000 | 0.000 |
| Ascomycota | Agaricomycetes | Boletales | Boletaceae | Solioccasus_polychromus | 0.000 | 0.000 | 0.000 | 0.001 | 0.000 | 0.000 | 0.000 | 0.000 | 0.000 | 0.000 | 0.000 |
| Mucoromycota | Sordariomycetes | Unclassified | Unclassified | Sordariomycetes_sp. | 0.004 | 0.000 | 0.000 | 0.000 | 0.000 | 0.0035 | 0 | 0.0166 | 0.0232 | 0.0221 | 0 |
| Mucoromycota | Agaricomycetes | Geastrales | Sphaerobolaceae | Sphaerobolus_sp. | 0.001 | 0.000 | 0.000 | 0.000 | 0.005 | 0.000 | 0.000 | 0.000 | 0.000 | 0.000 | 0.000 |
| Ascomycota | Ustilaginomycetes | Ustilaginales | Ustilaginaceae | Sporisorium_sorghi | 0.001 | 0.000 | 0.000 | 0.000 | 0.025 | 0.000 | 0.000 | 0.000 | 0.000 | 0.000 | 0.000 |
| Basidiomycota | Microbotryomycetes | Sporidiobolales | Sporidiobolaceae | Sporobolomyces_sp. | 0.003 | 0.000 | 0.000 | 0.000 | 0.000 | 0.000 | 0.000 | 0.000 | 0.000 | 0.000 | 0.000 |
| Basidiomycota | Sordariomycetes | Ophiostomatales | Ophiostomataceae | Sporothrix_inflata | 0.000 | 0.000 | 0.000 | 0.000 | 0.000 | 0.0746 | 0 | 0 | 0 | 0 | 0 |
| Ascomycota | Sordariomycetes | Ophiostomatales | Ophiostomataceae | Sporothrix_phasma | 0.003 | 0.000 | 0.000 | 0.000 | 0.000 | 0.004 | 0.000 | 0.000 | 0.000 | 0.000 | 0.000 |
| Ascomycota | Sordariomycetes | Ophiostomatales | Ophiostomataceae | Sporothrix_schenckii | 0.001 | 0.000 | 0.000 | 0.000 | 0.000 | 0.000 | 0.000 | 0.000 | 0.000 | 0.000 | 0.000 |

**Table S8 continued.**

| | | | | | | | | | | | | | | | |
|---|---|---|---|---|---|---|---|---|---|---|---|---|---|---|---|
| Ascomycota | Sordariomycetes | Ophiostomatales | Ophiostomataceae | Sporothrix_sp. | 0.000 | 0.000 | 0.000 | 0.000 | 0.000 | 0 | 0 | 0 | 0.0058 | 0 | 0 |
| Ascomycota | Sordariomycetes | Hypocreales | Stachybotryaceae | Stachybotrys_chartarum | 0.000 | 0.000 | 0.000 | 0.000 | 0.005 | 0.000 | 0.000 | 0.000 | 0.000 | 0.000 | 0.000 |
| Ascomycota | Dothideomycetes | Pleosporales | Massarinaceae | Stagonospora_trichophoricola | 0.000 | 0.000 | 0.000 | 0.000 | 0.000 | 0.0462 | 0 | 0 | 0 | 0.0055 | 0 |
| Ascomycota | Sordariomycetes | Hypocreales | Unclassified | Stilbella_fimetaria | 0.000 | 0.000 | 0.000 | 0.000 | 0.000 | 0 | 0 | 0.0111 | 0.0058 | 0.0055 | 0 |
| Ascomycota | Sordariomycetes | Xylariales | Beltraniaceae | Subsessila_turbinata | 0.001 | 0.000 | 0.000 | 0.000 | 0.000 | 0.000 | 0.000 | 0.000 | 0.000 | 0.000 | 0.000 |
| Basidiomycota | Agaricomycetes | Boletales | Suillaceae | Suillus_bovinus | 0.001 | 0.000 | 0.000 | 0.000 | 0.000 | 0.000 | 0.000 | 0.000 | 0.000 | 0.000 | 0.000 |
| Basidiomycota | Exobasidiomycetes | Microstromatales | Unclassified | Sympodiomycopsis_kandeliae | 0.003 | 0.000 | 0.000 | 0.000 | 0.000 | 0.000 | 0.000 | 0.000 | 0.000 | 0.000 | 0.000 |
| Ascomycota | Eurotiomycetes | Eurotiales | Trichocomaceae | Talaromyces_diversus | 0.000 | 0.000 | 0.000 | 0.018 | 0.020 | 0 | 0.0016 | 0 | 0 | 0 | 0 |
| Basidiomycota | Eurotiomycetes | Eurotiales | Trichocomaceae | Talaromyces_minioluteus | 0.001 | 0.000 | 0.000 | 0.000 | 0.000 | 0.000 | 0.000 | 0.000 | 0.000 | 0.000 | 0.000 |
| Ascomycota | Eurotiomycetes | Eurotiales | Trichocomaceae | Talaromyces_purpureogenus | 0.013 | 0.000 | 0.000 | 0.000 | 0.000 | 0.000 | 0.000 | 0.000 | 0.000 | 0.000 | 0.000 |
| Ascomycota | Eurotiomycetes | Eurotiales | Trichocomaceae | Talaromyces_sp. | 0.000 | 0.000 | 0.000 | 0.000 | 0.000 | 0 | 0 | 0 | 0 | 0.0055 | 0 |
| Ascomycota | Eurotiomycetes | Eurotiales | Trichocomaceae | Talaromyces_verruculosus | 0.000 | 0.000 | 0.000 | 0.000 | 0.000 | 0 | 0.0016 | 0 | 0 | 0 | 0 |
| Ascomycota | Taphrinomycetes | Taphrinales | Taphrinaceae | Taphrina_deformans | 0.000 | 0.000 | 0.000 | 0.020 | 0.015 | 0.000 | 0.000 | 0.000 | 0.000 | 0.000 | 0.000 |
| Ascomycota | Agaricomycetes | Agaricales | Lyophyllaceae | Termitomyces_DKA-2007 | 0.110 | 0.076 | 0.130 | 0.002 | 0.015 | 0.1956 | 0.0591 | 0.0055 | 0 | 0 | 0 |
| Ascomycota | Agaricomycetes | Agaricales | Lyophyllaceae | Termitomyces_sp. | 2.085 | 1.186 | 2.051 | 0.012 | 0.666 | 4.0678 | 1.2964 | 0.0555 | 0.1451 | 0 | 0 |
| Ascomycota | Agaricomycetes | Agaricales | Lyophyllaceae | Termitomyces_sp._Group1 | 0.046 | 0.000 | 0.023 | 0.000 | 0.000 | 0.0994 | 0.0181 | 0 | 0.0058 | 0 | 0 |
| Ascomycota | Agaricomycetes | Agaricales | Lyophyllaceae | Termitomyces_sp._Group2 | 0.041 | 0.151 | 0.079 | 0.000 | 0.000 | 0.1521 | 0.0673 | 0 | 0 | 0 | 0 |
| Basidiomycota | Agaricomycetes | Agaricales | Lyophyllaceae | Termitomyces_sp._Group3 | 70.342 | 81.272 | 82.530 | 0.571 | 11.344 | 171.02 | 88.39 | 2.1018 | 10.013 | 0.0442 | 0.1601 |
| Mucoromycota | Agaricomycetes | Agaricales | Lyophyllaceae | Termitomyces_sp._Group4 | 0.130 | 0.000 | 0.068 | 0.000 | 0.035 | 0.239 | 0.041 | 0 | 0.0058 | 0 | 0 |
| Basidiomycota | Agaricomycetes | Agaricales | Lyophyllaceae | Termitomyces_sp._Group5 | 0.277 | 0.101 | 0.169 | 0.000 | 0.075 | 0.5278 | 0.0804 | 0.0055 | 0.0232 | 0.0055 | 0 |
| Ascomycota | Agaricomycetes | Agaricales | Lyophyllaceae | Termitomyces_sp._Group6 | 0.001 | 0.000 | 0.006 | 0.000 | 0.000 | 0 | 0.0033 | 0 | 0 | 0 | 0 |
| Ascomycota | Agaricomycetes | Agaricales | Lyophyllaceae | Termitomyces_sp._Group7 | 0.013 | 0.000 | 0.023 | 0.000 | 0.015 | 0.0292 | 0.0049 | 0 | 0 | 0 | 0 |
| Ascomycota | Agaricomycetes | Agaricales | Lyophyllaceae | Termitomyces_sp._Group8 | 1.156 | 0.883 | 1.234 | 0.003 | 0.135 | 1.9334 | 0.7565 | 0.0277 | 0.0639 | 0 | 0 |
| Basidiomycota | Agaricomycetes | Agaricales | Lyophyllaceae | Termitomyces_sp._Juja-2 | 0.001 | 0.000 | 0.000 | 0.000 | 0.000 | 0.000 | 0.000 | 0.000 | 0.000 | 0.000 | 0.000 |
| Ascomycota | Agaricomycetes | Agaricales | Lyophyllaceae | Termitomyces_sp._Juja-5 | 0.001 | 0.000 | 0.006 | 0.000 | 0.000 | 0.000 | 0.000 | 0.000 | 0.000 | 0.000 | 0.000 |
| Basidiomycota | Agaricomycetes | Agaricales | Lyophyllaceae | Termitomyces_sp._Juja-6 | 0.012 | 0.000 | 0.034 | 0.000 | 0.000 | 0.0111 | 0.0016 | 0 | 0 | 0 | 0 |
| Basidiomycota | Agaricomycetes | Agaricales | Lyophyllaceae | Termitomyces_sp._Juja-7 | 0.000 | 0.000 | 0.000 | 0.000 | 0.000 | 0.0035 | 0 | 0 | 0.0058 | 0 | 0 |
| Basidiomycota | Agaricomycetes | Agaricales | Lyophyllaceae | Termitomyces_sp._Juja-8 | 0.000 | 0.000 | 0.000 | 0.000 | 0.000 | 0.0071 | 0.0082 | 0 | 0 | 0 | 0 |
| Ascomycota | Agaricomycetes | Agaricales | Lyophyllaceae | Termitomyces_striatus | 0.000 | 0.000 | 0.000 | 0.000 | 0.000 | 0.004 | 0.000 | 0.000 | 0.000 | 0.000 | 0.000 |
| Ascomycota | Agaricomycetes | Agaricales | Lyophyllaceae | Termitomyces_symbiont_of_Allodontotermes_sp. | 0.004 | 0.000 | 0.000 | 0.000 | 0.000 | 0.0035 | 0.0033 | 0 | 0 | 0 | 0 |
| Ascomycota | Agaricomycetes | Agaricales | Lyophyllaceae | Termitomyces_symbiont_of_Macrotermes_bellicosus | 0.001 | 0.000 | 0.000 | 0.000 | 0.000 | 0.007 | 0.000 | 0.000 | 0.000 | 0.000 | 0.000 |
| Ascomycota | Agaricomycetes | Agaricales | Lyophyllaceae | Termitomyces_symbiont_of_Macrotermes_muelleri | 0.001 | 0.000 | 0.000 | 0.000 | 0.000 | 0.000 | 0.000 | 0.000 | 0.000 | 0.000 | 0.000 |

**Table S8 continued.**

| | | | | | | | | | | | | | | | |
|---|---|---|---|---|---|---|---|---|---|---|---|---|---|---|---|
| Ascomycota | Agaricomycetes | Agaricales | Lyophyllaceae | Termitomyces_symbiont_of_Microtermes_sp. | 0.000 | 0.000 | 0.000 | 0.000 | 0.000 | 0.0035 | 0 | 0 | 0 | 0 | 0 |
| Ascomycota | Dothideomycetes | Pleosporales | Tetraplosphaeriaceae | Tetraplosphaeria_tetraploa | 0.000 | 0.000 | 0.000 | 0.000 | 0.000 | 0.000 | 0.000 | 0.000 | 0.000 | 0.000 | 0.000 |
| Ascomycota | Pucciniomycetes | Pucciniales | Pucciniastraceae | Thekopsora_areolata | 0.000 | 0.000 | 0.000 | 0.000 | 0.000 | 0.000 | 0.000 | 0.000 | 0.000 | 0.000 | 0.000 |
| Ascomycota | Agaricomycetes | Thelephorales | Thelephoraceae | Thelephoraceae_sp. | 0.001 | 0.000 | 0.000 | 0.000 | 0.000 | 0.000 | 0.000 | 0.000 | 0.000 | 0.000 | 0.000 |
| Basidiomycota | Sordariomycetes | Microascales | Ceratocystidaceae | Thielaviopsis_basicola | 0.000 | 0.000 | 0.000 | 0.000 | 0.000 | 0 | 0 | 0 | 0 | 0 | 0.02 |
| Basidiomycota | Exobasidiomycetes | Tilletiales | Tilletiaceae | Tilletia_controversa | 0.000 | 0.000 | 0.006 | 0.000 | 0.000 | 0.0035 | 0 | 0 | 0 | 0 | 0 |
| Ascomycota | Exobasidiomycetes | Tilletiales | Tilletiaceae | Tilletia_ehrhartae | 0.000 | 0.000 | 0.000 | 0.007 | 0.000 | 0.000 | 0.000 | 0.000 | 0.000 | 0.000 | 0.000 |
| Ascomycota | Sordariomycetes | Hypocreales | Ophiocordycipitaceae | Tolypocladium_album | 0.000 | 0.000 | 0.000 | 0.000 | 0.000 | 0.0035 | 0 | 0.0166 | 0.0116 | 0 | 0 |
| Ascomycota | Sordariomycetes | Hypocreales | Ophiocordycipitaceae | Tolypocladium_cylindrosporum | 0.000 | 0.000 | 0.000 | 0.000 | 0.000 | 0 | 0 | 0 | 0.0058 | 0 | 0 |
| Ascomycota | Sordariomycetes | Hypocreales | Ophiocordycipitaceae | Tolypocladium_sp. | 0.000 | 0.000 | 0.000 | 0.000 | 0.000 | 0.0035 | 0 | 0 | 0 | 0 | 0 |
| Ascomycota | Sordariomycetes | Hypocreales | Clavicipitaceae | Torrubiella_hemipterigena | 0.000 | 0.000 | 0.000 | 0.000 | 0.000 | 0.000 | 0.000 | 0.000 | 0.000 | 0.000 | 0.000 |
| Ascomycota | Agaricomycetes | Polyporales | Polysporaceae | Trametes_hirsuta | 0.012 | 0.000 | 0.000 | 0.006 | 0.010 | 0.0074 | 0.0033 | 0 | 0 | 0 | 0 |
| Basidiomycota | Agaricomycetes | Polyporales | Polysporaceae | Trametes_pubescens | 0.005 | 0.000 | 0.011 | 0.077 | 0.050 | 0.0325 | 0.0066 | 0 | 0 | 0 | 0 |
| Basidiomycota | Agaricomycetes | Polyporales | Polysporaceae | Trametes_sp. | 0.000 | 0.000 | 0.000 | 0.000 | 0.000 | 0 | 0.0016 | 0 | 0 | 0 | 0 |
| Basidiomycota | Agaricomycetes | Polyporales | Polysporaceae | Trametes_versicolor | 0.000 | 0.000 | 0.000 | 0.000 | 0.000 | 0.000 | 0.000 | 0.000 | 0.000 | 0.000 | 0.000 |
| Basidiomycota | Sordariomycetes | Hypocreales | Hypocreaceae | Trichoderma_asperellum | 0.000 | 0.000 | 0.000 | 0.000 | 0.000 | 0 | 0 | 0.0055 | 0 | 0.0055 | 0 |
| Basidiomycota | Sordariomycetes | Hypocreales | Hypocreaceae | Trichoderma_atroviride | 0.000 | 0.000 | 0.000 | 0.000 | 0.000 | 0 | 0 | 0.0166 | 0.029 | 0.0055 | 0.04 |
| Basidiomycota | Sordariomycetes | Hypocreales | Hypocreaceae | Trichoderma_brevicompactum | 0.000 | 0.000 | 0.000 | 0.000 | 0.000 | 0.0071 | 0 | 0 | 0.0116 | 0.0276 | 0 |
| Basidiomycota | Sordariomycetes | Hypocreales | Hypocreaceae | Trichoderma_citrinoviride | 0.000 | 0.000 | 0.000 | 0.000 | 0.000 | 0 | 0 | 0.0111 | 0 | 0.0055 | 0.02 |
| Basidiomycota | Sordariomycetes | Hypocreales | Hypocreaceae | Trichoderma_erinaceum | 0.000 | 0.000 | 0.000 | 0.000 | 0.000 | 0.0035 | 0 | 0.0444 | 0.0232 | 0.0276 | 0 |
| Basidiomycota | Sordariomycetes | Hypocreales | Hypocreaceae | Trichoderma_gamsii | 0.000 | 0.000 | 0.000 | 0.000 | 0.000 | 0 | 0 | 0.0055 | 0.0058 | 0 | 0 |
| Basidiomycota | Sordariomycetes | Hypocreales | Hypocreaceae | Trichoderma_hamatum | 0.000 | 0.000 | 0.000 | 0.000 | 0.000 | 0 | 0.0016 | 0 | 0 | 0 | 0 |
| Basidiomycota | Sordariomycetes | Hypocreales | Hypocreaceae | Trichoderma_harzianum | 0.000 | 0.000 | 0.000 | 0.000 | 0.000 | 0 | 0 | 0.0055 | 0 | 0.0276 | 0.02 |
| Basidiomycota | Sordariomycetes | Hypocreales | Hypocreaceae | Trichoderma_koningii | 0.000 | 0.000 | 0.000 | 0.000 | 0.000 | 0 | 0 | 0 | 0 | 0.011 | 0 |
| Basidiomycota | Sordariomycetes | Hypocreales | Hypocreaceae | Trichoderma_koningiopsis | 0.000 | 0.000 | 0.000 | 0.000 | 0.000 | 0 | 0 | 0 | 0.0058 | 0 | 0 |
| Basidiomycota | Sordariomycetes | Hypocreales | Hypocreaceae | Trichoderma_lixii | 0.000 | 0.000 | 0.000 | 0.000 | 0.000 | 0.0035 | 0 | 0 | 0.0058 | 0.0166 | 0.02 |
| Basidiomycota | Sordariomycetes | Hypocreales | Hypocreaceae | Trichoderma_longibrachiatum | 0.000 | 0.000 | 0.000 | 0.000 | 0.000 | 0.0035 | 0 | 0 | 0 | 0.0055 | 0 |
| Basidiomycota | Sordariomycetes | Hypocreales | Hypocreaceae | Trichoderma_pleuroticola | 0.000 | 0.000 | 0.000 | 0.000 | 0.000 | 0 | 0 | 0 | 0.0058 | 0 | 0 |
| Basidiomycota | Sordariomycetes | Hypocreales | Hypocreaceae | Trichoderma_reesei | 0.001 | 0.000 | 0.006 | 0.000 | 0.010 | 0.1099 | 0.0197 | 0.1442 | 0.267 | 0.1877 | 0.3802 |
| Basidiomycota | Sordariomycetes | Hypocreales | Hypocreaceae | Trichoderma_sp. | 0.003 | 0.000 | 0.000 | 0.000 | 0.000 | 0.0213 | 0.0016 | 0.0222 | 0.0174 | 0.0607 | 0.02 |
| Basidiomycota | Sordariomycetes | Hypocreales | Hypocreaceae | Trichoderma_spirale | 0.000 | 0.000 | 0.000 | 0.000 | 0.000 | 0.0035 | 0 | 0.0111 | 0 | 0.0773 | 0 |
| Ascomycota | Sordariomycetes | Hypocreales | Hypocreaceae | Trichoderma_tomentosum | 0.000 | 0.000 | 0.000 | 0.000 | 0.000 | 0.0035 | 0 | 0.0055 | 0 | 0.011 | 0 |

**Table S8 continued.**

| | | | | | | | | | | | | | | | |
|---|---|---|---|---|---|---|---|---|---|---|---|---|---|---|---|
| Basidiomycota | Sordariomycetes | Hypocreales | Hypocreaceae | Trichoderma_virens | 0.001 | 0.000 | 0.056 | 0.000 | 0.000 | 0.0707 | 0.0016 | 0.0444 | 0.0232 | 0.5962 | 0.06 |
| Ascomycota | Sordariomycetes | Hypocreales | Hypocreaceae | Trichoderma_viride | 0.000 | 0.000 | 0.000 | 0.000 | 0.000 | 0 | 0 | 0 | 0 | 0 | 0.02 |
| Basidiomycota | Agaricomycetes | Agaricales | Tricholomataceae | Tricholoma_elegans | 0.000 | 0.000 | 0.000 | 0.000 | 0.000 | 0.011 | 0.000 | 0.000 | 0.000 | 0.000 | 0.000 |
| Basidiomycota | Agaricomycetes | Agaricales | Tricholomataceae | Tricholoma_lascivum | 0.001 | 0.000 | 0.000 | 0.000 | 0.000 | 0.004 | 0.000 | 0.000 | 0.000 | 0.000 | 0.000 |
| Ascomycota | Agaricomycetes | Agaricales | Tricholomataceae | Tricholoma_matsutake | 0.000 | 0.000 | 0.000 | 0.000 | 0.010 | 0 | 0.0016 | 0 | 0 | 0 | 0 |
| Basidiomycota | Agaricomycetes | Agaricales | Tricholomataceae | Tricholoma_scalpturatum | 0.001 | 0.000 | 0.000 | 0.000 | 0.000 | 0.000 | 0.000 | 0.000 | 0.000 | 0.000 | 0.000 |
| Basidiomycota | Agaricomycetes | Agaricales | Tricholomataceae | Tricholoma_sp. | 0.000 | 0.000 | 0.000 | 0.000 | 0.000 | 0.0037 | 0.0016 | 0 | 0 | 0 | 0 |
| Ascomycota | Saccharomycetes | Saccharomycetales | Trichomonascaceae | Trichomonascus_apis | 0.005 | 0.000 | 0.000 | 0.000 | 0.000 | 0.000 | 0.000 | 0.000 | 0.000 | 0.000 | 0.000 |
| Ascomycota | Eurotiomycetes | Onygenales | Arthrodermataceae | Trichophyton_sp. | 0.000 | 0.000 | 0.000 | 0.001 | 0.000 | 0.000 | 0.000 | 0.000 | 0.000 | 0.000 | 0.000 |
| Ascomycota | Eurotiomycetes | Onygenales | Arthrodermataceae | Trichophyton_terrestre | 0.000 | 0.000 | 0.000 | 0.022 | 0.015 | 0.000 | 0.000 | 0.000 | 0.000 | 0.000 | 0.000 |
| Ascomycota | Tritirachiomycetes | Tritirachiales | Tritirachiaceae | Tritirachium_sp. | 0.000 | 0.000 | 0.000 | 0.000 | 0.000 | 0 | 0 | 0 | 0 | 0.0055 | 0 |
| Basidiomycota | Sordariomycetes | Xylariales | Sporocadaceae | Truncatella_angustata | 0.001 | 0.000 | 0.000 | 0.000 | 0.000 | 0.000 | 0.000 | 0.000 | 0.000 | 0.000 | 0.000 |
| Basidiomycota | Sordariomycetes | Xylariales | Sporocadaceae | Truncatella_cf._angustata | 0.000 | 0.000 | 0.000 | 0.000 | 0.000 | 0 | 0 | 0.0055 | 0 | 0 | 0 |
| Basidiomycota | Sordariomycetes | Diaporthales | Melanconiellaceae | Tubakia_dryina | 0.000 | 0.000 | 0.000 | 0.000 | 0.000 | 0 | 0 | 0 | 0.0058 | 0 | 0 |
| Basidiomycota | Agaricomycetes | Boletales | Boletaceae | Tylopilus_porphyrosporus | 0.000 | 0.000 | 0.000 | 0.024 | 0.020 | 0.000 | 0.000 | 0.000 | 0.000 | 0.000 | 0.000 |
| Basidiomycota | Lecanoromycetes | Umbilicariales | Umbilicariaceae | Umbilicaria_muehlenbergii | 0.000 | 0.025 | 0.000 | 0.000 | 0.000 | 0.0035 | 0.000 | 0.000 | 0.000 | 0.000 | 0.000 |
| Ascomycota | Unclassified | Unclassified | Unclassified | Unclassified Ascomycota | 0.004 | 0.025 | 0.017 | 0.001 | 0.000 | 0.015 | 0.000 | 0.000 | 0.000 | 0.000 | 0.000 |
| Ascomycota | Unclassified | Unclassified | Unclassified | Uncultured _fungus | 0.000 | 0.000 | 0.000 | 0.000 | 0.000 | 0.0212 | 0.1608 | 0.3882 | 1.7298 | 0.3478 | 0.3601 |
| Ascomycota | Agaricomycetes | Agaricales | Agaricaceae | Uncultured_Agaricaceae | 0.005 | 0.000 | 0.006 | 0.000 | 0.000 | 0.0037 | 0.0033 | 0 | 0 | 0 | 0 |
| Ascomycota | Agaricomycetes | Agaricales | Unclassified | Uncultured_Agaricales | 1.208 | 1.312 | 1.200 | 0.003 | 0.140 | 2.0553 | 0.8041 | 0.0333 | 0.1045 | 0.011 | 0 |
| Ascomycota | Agaricomycetes | Agaricales | Unclassified | Uncultured_Agaricomycetes | 0.000 | 0.000 | 0.000 | 0.001 | 0.000 | 0.000 | 0.000 | 0.000 | 0.000 | 0.000 | 0.000 |
| Ascomycota | Agaricomycetes | Agaricales | Cortinariaceae | Uncultured_Alnicola | 0.000 | 0.000 | 0.000 | 0.000 | 0.000 | 0.004 | 0.000 | 0.000 | 0.000 | 0.000 | 0.000 |
| Ascomycota | Agaricomycetes | Agaricales | Amanitaceae | Uncultured_Amanita | 0.001 | 0.000 | 0.000 | 0.000 | 0.000 | 0.0035 | 0 | 0 | 0 | 0 | 0 |
| Ascomycota | Unclassified | Unclassified | Unclassified | Uncultured_AMF_fungus | 0.001 | 0.000 | 0.000 | 0.000 | 0.000 | 0.000 | 0.000 | 0.000 | 0.000 | 0.000 | 0.000 |
| Ascomycota | Agaricomycetes | Atheliales | Atheliaceae | Uncultured_Amphinema | 0.003 | 0.000 | 0.006 | 0.000 | 0.000 | 0 | 0.0016 | 0 | 0 | 0 | 0 |
| Ascomycota | Unclassified | Unclassified | Unclassified | Uncultured_Ascomycota | 0.007 | 0.025 | 0.451 | 0.013 | 0.010 | 8.6879 | 0.2412 | 28.904 | 18.279 | 8.6222 | 19.788 |
| Ascomycota | Dothideomycetes | Dothideales | Dothioraceae | Uncultured_Aureobasidium | 0.001 | 0.000 | 0.000 | 0.000 | 0.000 | 0.000 | 0.000 | 0.000 | 0.000 | 0.000 | 0.000 |
| Ascomycota | Unclassified | Unclassified | Unclassified | Uncultured_basal_lineage_fungus | 0.000 | 0.000 | 0.000 | 0.004 | 0.015 | 0.000 | 0.000 | 0.000 | 0.000 | 0.000 | 0.000 |
| Ascomycota | Unclassified | Unclassified | Unclassified | Uncultured_basidiomycete | 0.000 | 0.000 | 0.000 | 0.004 | 0.000 | 0.0072 | 0.0016 | 0 | 0 | 0 | 0 |
| Ascomycota | Unclassified | Unclassified | Unclassified | Uncultured_Basidiomycota | 0.000 | 0.000 | 0.000 | 0.000 | 0.000 | 0.000 | 0.000 | 0.000 | 0.000 | 0.000 | 0.000 |
| Ascomycota | Sordariomycetes | Hypocreales | Bionectriaceae | Uncultured_Bionectria | 0.000 | 0.000 | 0.000 | 0.000 | 0.000 | 0 | 0 | 0.0055 | 0.0058 | 0 | 0 |
| Ascomycota | Agaricomycetes | Boletales | Unclassified | Uncultured_Boletaceae | 0.000 | 0.000 | 0.000 | 0.000 | 0.000 | 0.000 | 0.000 | 0.000 | 0.000 | 0.000 | 0.000 |

**Table S8 continued.**

| | | | | | | | | | | | | | | | |
|---|---|---|---|---|---|---|---|---|---|---|---|---|---|---|---|
| Ascomycota | Dothideomycetes | Unclassified | Gloniaceae | Uncultured_Cenococcum | 0.000 | 0.000 | 0.000 | 0.001 | 0.000 | 0.000 | 0.000 | 0.000 | 0.000 | 0.000 | 0.000 |
| Ascomycota | Agaricomycetes | Cantharellales | Ceratobasidiaceae | Uncultured_Ceratobasidiaceae | 0.001 | 0.000 | 0.000 | 0.000 | 0.000 | 0 | 0.0033 | 0 | 0 | 0 | 0 |
| Basidiomycota | Dothideomycetes | Cladosporiales | Cladosporiaceae | Uncultured_Cladosporium | 0.000 | 0.000 | 0.000 | 0.002 | 0.015 | 0.000 | 0.000 | 0.000 | 0.000 | 0.000 | 0.000 |
| Basidiomycota | Sordariomycetes | Glomerellales | Plectosphaerellacea | Uncultured_Colletotrichum | 0.000 | 0.000 | 0.000 | 0.000 | 0.000 | 0 | 0 | 0.0055 | 0 | 0 | 0 |
| Basidiomycota | Unclassified | Unclassified | Unclassified | Uncultured_compost_fungus | 0.008 | 0.025 | 0.090 | 0.000 | 0.000 | 0 | 0.0016 | 0 | 0 | 0 | 0 |
| Basidiomycota | Agaricomycetes | Unclassified | Unclassified | Uncultured_Corticiales | 0.000 | 0.000 | 0.000 | 0.017 | 0.000 | 0.000 | 0.000 | 0.000 | 0.000 | 0.000 | 0.000 |
| Basidiomycota | Agaricomycetes | Agaricales | Cortinariaceae | Uncultured_Cortinarius | 0.016 | 0.025 | 0.011 | 0.001 | 0.000 | 0.0433 | 0.0181 | 0 | 0.0058 | 0 | 0 |
| Basidiomycota | Unclassified | Unclassified | Unclassified | Uncultured_Cucurbitaria | 0.000 | 0.000 | 0.000 | 0.000 | 0.000 | 0.0675 | 0 | 0 | 0 | 0 | 0 |
| Ascomycota | Unclassified | Unclassified | Unclassified | Uncultured_ectomycorrhizal_fungus | 0.038 | 0.151 | 0.062 | 0.231 | 0.150 | 0.1045 | 0.0542 | 0.0055 | 0.0232 | 0.0055 | 0.02 |
| Ascomycota | Unclassified | Unclassified | Unclassified | Uncultured_Elaphomyces | 0.001 | 0.000 | 0.000 | 0.000 | 0.000 | 0.000 | 0.000 | 0.000 | 0.000 | 0.000 | 0.000 |
| Ascomycota | Unclassified | Unclassified | Unclassified | Uncultured_endophytic_fungus | 0.075 | 0.025 | 0.056 | 0.023 | 0.085 | 0.0213 | 0.0164 | 0.0943 | 0.3309 | 0.0442 | 0.1401 |
| Basidiomycota | Unclassified | Unclassified | Unclassified | Uncultured_Entoloma | 0.003 | 0.000 | 0.000 | 0.000 | 0.000 | 0.0035 | 0.0033 | 0 | 0 | 0 | 0 |
| Ascomycota | Sordariomycetes | Hypocreales | Clavicipitaceae | Uncultured_Epichloe | 0.000 | 0.000 | 0.000 | 0.000 | 0.000 | 0.0035 | 0 | 0 | 0.0116 | 0.0055 | 0.02 |
| Ascomycota | Unclassified | Unclassified | Unclassified | Uncultured_fungus | 19.135 | 8.960 | 5.848 | 45.605 | 39.800 | 5.6863 | 3.1459 | 29.292 | 28.444 | 43.9 | 55.102 |
| Ascomycota | Pezizomycetes | Pezizales | Pyronemataceae | Uncultured_Geopora | 0.000 | 0.025 | 0.000 | 0.000 | 0.000 | 0.000 | 0.000 | 0.000 | 0.000 | 0.000 | 0.000 |
| Basidiomycota | Glomeromycetes | Glomerales | Glomeraceae | Uncultured_glomeraceous_AM_fungus | 0.003 | 0.000 | 0.000 | 0.000 | 0.000 | 0.000 | 0.000 | 0.000 | 0.000 | 0.000 | 0.000 |
| Ascomycota | Agaricomycetes | Agaricales | Cortinariaceae | Uncultured_Hebeloma | 0.007 | 0.025 | 0.011 | 0.000 | 0.000 | 0.0071 | 0.0082 | 0 | 0 | 0 | 0 |
| Unclassified | Pezizomycetes | Pezizales | Helvellaceae | Uncultured_Helvella | 0.000 | 0.000 | 0.000 | 0.000 | 0.000 | 0.000 | 0.000 | 0.000 | 0.000 | 0.000 | 0.000 |
| Basidiomycota | Agaricomycetes | Cantharellales | Hydnaceae | Uncultured_Hydnum | 0.001 | 0.000 | 0.000 | 0.000 | 0.000 | 0.000 | 0.000 | 0.000 | 0.000 | 0.000 | 0.000 |
| Basidiomycota | Agaricomycetes | Agaricales | Hymenogastraceae | Uncultured_Hymenogaster | 0.003 | 0.000 | 0.000 | 0.000 | 0.000 | 0 | 0.0066 | 0 | 0 | 0 | 0 |
| Basidiomycota | Sordariomycetes | Hypocreales | Unclassified | Uncultured_Hypocreales | 0.000 | 0.000 | 0.000 | 0.000 | 0.000 | 0.0035 | 0 | 0 | 0.0058 | 0 | 0 |
| Basidiomycota | Agaricomycetes | Agaricales | Cortinariaceae | Uncultured_Inocybe | 0.000 | 0.000 | 0.000 | 0.000 | 0.000 | 0.004 | 0.000 | 0.000 | 0.000 | 0.000 | 0.000 |
| Unclassified | Agaricomycetes | Agaricales | Tricholomataceae | Uncultured_Laccaria | 0.003 | 0.025 | 0.011 | 0.000 | 0.000 | 0.0107 | 0.0049 | 0 | 0 | 0 | 0 |
| Basidiomycota | Agaricomycetes | Russulales | Russulaceae | Uncultured_Lactarius | 0.000 | 0.000 | 0.000 | 0.003 | 0.000 | 0.000 | 0.000 | 0.000 | 0.000 | 0.000 | 0.000 |
| Ascomycota | Unclassified | Unclassified | Unclassified | Uncultured_leaf_litter_fungus | 0.001 | 0.025 | 0.011 | 0.000 | 0.000 | 0.011 | 0.000 | 0.000 | 0.000 | 0.000 | 0.000 |
| Ascomycota | Malasseziomycetes | Malasseziales | Unclassified | Uncultured_Malasseziales | 0.000 | 0.000 | 0.000 | 0.000 | 0.000 | 0.000 | 0.000 | 0.000 | 0.000 | 0.000 | 0.000 |
| Unclassified | Unclassified | Unclassified | Unclassified | Uncultured_marine_fungus | 0.011 | 0.025 | 0.011 | 0.780 | 0.661 | 0 | 0 | 0.0111 | 0.0116 | 0 | 0 |
| Basidiomycota | Unclassified | Unclassified | Unclassified | Uncultured_mycorrhizal_fungus | 0.087 | 0.076 | 0.068 | 0.077 | 0.055 | 0.1047 | 0.0509 | 0.0055 | 0.0116 | 0 | 0 |
| Basidiomycota | Neocallimastigomycetes | Neocaliimastigales | Unclassified | Uncultured_Neocallimastigales | 0.022 | 0.000 | 0.000 | 0.001 | 0.000 | 0.000 | 0.000 | 0.000 | 0.000 | 0.000 | 0.000 |
| Ascomycota | Pezizomycetes | Pezizales | Pezizaceae | Uncultured_Peziza | 0.038 | 0.000 | 0.000 | 0.000 | 0.000 | 0.000 | 0.000 | 0.000 | 0.000 | 0.000 | 0.000 |
| Basidiomycota | Pezizomycetes | Pezizales | Pezizaceae | Uncultured_Pezizales | 0.009 | 0.000 | 0.000 | 0.000 | 0.000 | 0.000 | 0.000 | 0.000 | 0.000 | 0.000 | 0.000 |
| Ascomycota | Pezizomycetes | Pezizales | Pyronemataceae | Uncultured_Pyronemataceae | 0.001 | 0.000 | 0.000 | 0.000 | 0.000 | 0.000 | 0.000 | 0.000 | 0.000 | 0.000 | 0.000 |

**Table S8 continued.**

| | | | | | | | | | | | | | | | |
|---|---|---|---|---|---|---|---|---|---|---|---|---|---|---|---|
| Basidiomycota | Unclassified | Unclassified | Unclassified | Uncultured_root-associated_fungus | 0.003 | 0.025 | 0.000 | 0.003 | 0.000 | 0.0106 | 0.0033 | 0.0055 | 0.0174 | 0 | 0.04 |
| Ascomycota | Agaricomycetes | Russulales | Russulaceae | Uncultured_Russula | 0.000 | 0.000 | 0.000 | 0.129 | 0.095 | 0.000 | 0.000 | 0.000 | 0.000 | 0.000 | 0.000 |
| Ascomycota | Agaricomycetes | Russulales | Russulaceae | Uncultured_Russulaceae | 0.000 | 0.000 | 0.000 | 0.002 | 0.000 | 0.000 | 0.000 | 0.000 | 0.000 | 0.000 | 0.000 |
| Unclassified | Agaricomycetes | Boletales | Sclerodermataceae | Uncultured_Scleroderma | 0.000 | 0.000 | 0.000 | 0.002 | 0.000 | 0.000 | 0.000 | 0.000 | 0.000 | 0.000 | 0.000 |
| Basidiomycota | Agaricomycetes | Sebacinales | Sebacinaceae | Uncultured_Sebacina | 0.000 | 0.000 | 0.000 | 0.004 | 0.005 | 0.000 | 0.000 | 0.000 | 0.000 | 0.000 | 0.000 |
| Basidiomycota | Agaricomycetes | Sebacinales | Sebacinaceae | Uncultured_Sebacinaceae | 0.000 | 0.000 | 0.000 | 0.000 | 0.005 | 0.000 | 0.000 | 0.000 | 0.000 | 0.000 | 0.000 |
| Unclassified | Unclassified | Unclassified | Unclassified | Uncultured_soil_fungus | 0.276 | 0.202 | 0.293 | 1.522 | 1.427 | 0.0789 | 0.0394 | 0.1109 | 0.1393 | 0.0938 | 0.1 |
| Unclassified | Agaricomycetes | Boletales | Suillaceae | Uncultured_Suillus | 0.000 | 0.000 | 0.000 | 0.010 | 0.020 | 0.000 | 0.000 | 0.000 | 0.000 | 0.000 | 0.000 |
| Unclassified | Agaricomycetes | Agaricales | Lyophyllaceae | Uncultured_Termitomyces | 1.498 | 1.312 | 1.758 | 0.004 | 0.165 | 2.6906 | 1.216 | 0.0444 | 0.1858 | 0 | 0 |
| Unclassified | Agaricomycetes | Cantharellales | Ceratobasidiaceae | Uncultured_Thanatephorus | 0.001 | 0.000 | 0.000 | 0.001 | 0.000 | 0.000 | 0.000 | 0.000 | 0.000 | 0.000 | 0.000 |
| Unclassified | Agaricomycetes | Thelephorales | Thelephoraceae | Uncultured_Thelephoraceae | 0.013 | 0.000 | 0.000 | 0.006 | 0.005 | 0.000 | 0.000 | 0.000 | 0.000 | 0.000 | 0.000 |
| Ascomycota | Agaricomycetes | Thelephorales | Thelephoraceae | Uncultured_Tomentella | 0.008 | 0.000 | 0.000 | 0.013 | 0.020 | 0.007 | 0.000 | 0.000 | 0.000 | 0.000 | 0.000 |
| Unclassified | Agaricomycetes | Agaricales | Tricholomataceae | Uncultured_Tricholomataceae | 0.001 | 0.000 | 0.000 | 0.000 | 0.000 | 0.000 | 0.000 | 0.000 | 0.000 | 0.000 | 0.000 |
| Ascomycota | Pezizomycetes | Pezizales | Pyronemataceae | Uncultured_Trichophaea | 0.000 | 0.025 | 0.000 | 0.000 | 0.000 | 0.000 | 0.000 | 0.000 | 0.000 | 0.000 | 0.000 |
| Mucoromycota | Pezizomycetes | Pezizales | Tuberaceae | Uncultured_Tuber | 0.000 | 0.000 | 0.000 | 0.006 | 0.000 | 0.000 | 0.000 | 0.000 | 0.000 | 0.000 | 0.000 |
| Basidiomycota | Unidentified | Unidentified | Unidentified | Unidentified | 0.663 | 2.398 | 1.279 | 49.503 | 40.020 | 0.3084 | 0.0771 | 0.2107 | 0.3367 | 0.3202 | 0.7403 |
| Basidiomycota | Sordariomycetes | Hypocreales | Clavicipitaceae | Ustilaginoidea_virens | 0.000 | 0.000 | 0.006 | 0.000 | 0.000 | 0.0072 | 0 | 0.0277 | 0.0116 | 0.0055 | 0.02 |
| Basidiomycota | Sordariomycetes | Glomerellales | Plectosphaerellacea | Verticillium_chlamydosporium | 0.000 | 0.000 | 0.000 | 0.000 | 0.000 | 0.0035 | 0 | 0 | 0 | 0 | 0.02 |
| Basidiomycota | Sordariomycetes | Glomerellales | Pectosphaerellacea | Verticillium_dahliae | 0.001 | 0.000 | 0.000 | 0.000 | 0.000 | 0.000 | 0.000 | 0.000 | 0.000 | 0.000 | 0.000 |
| Ascomycota | Unidentified | Unidentified | Unidentified | Vigna_angularis_var._angularis | 0.000 | 0.000 | 0.000 | 0.000 | 0.000 | 0 | 0 | 0.0055 | 0 | 0 | 0 |
| Basidiomycota | Wallemiomycetes | Wallemiales | Wallemiaceae | Wallemia_ichthyophaga | 0.004 | 0.000 | 0.000 | 0.000 | 0.000 | 0.000 | 0.000 | 0.000 | 0.000 | 0.000 | 0.000 |
| Basidiomycota | Sordariomycetes | Xylariales | Xylariaceae | Whalleya_microplaca | 0.000 | 0.000 | 0.000 | 0.000 | 0.000 | 0 | 0 | 0.0055 | 0.0058 | 0 | 0 |
| Basidiomycota | Unclassified | Unclassified | Unclassified | Woollsia | 0.000 | 0.000 | 0.000 | 0.000 | 0.000 | 0 | 0 | 0.0055 | 0.0058 | 0 | 0 |
| Basidiomycota | Sordariomycetes | Xylariales | Xylariaceae | Xylaria_angulosa | 0.001 | 0.000 | 0.000 | 0.000 | 0.000 | 0 | 0.0049 | 0.1497 | 0.1161 | 0.9439 | 0 |
| Unclassified | Sordariomycetes | Xylariales | Xylariaceae | Xylaria_arbuscula | 0.000 | 0.000 | 0.000 | 0.000 | 0.000 | 0.0035 | 0 | 0 | 0.0116 | 0 | 0.02 |
| Basidiomycota | Sordariomycetes | Xylariales | Xylariaceae | Xylaria_brunneovinosa | 0.000 | 0.000 | 0.017 | 0.000 | 0.000 | 0 | 0.023 | 0.3827 | 0.1741 | 0.2429 | 0.2201 |
| Unclassified | Sordariomycetes | Xylariales | Xylariaceae | Xylaria_carpophila | 0.000 | 0.000 | 0.000 | 0.000 | 0.000 | 0 | 0 | 0 | 0 | 0.0055 | 0 |
| Unclassified | Sordariomycetes | Xylariales | Xylariaceae | Xylaria_cf._heliscus | 0.000 | 0.000 | 0.000 | 0.000 | 0.000 | 0 | 0.0016 | 0.0222 | 0.1161 | 0.011 | 0.06 |
| Chytridiomycota | Sordariomycetes | Xylariales | Xylariaceae | Xylaria_enteroleuca | 0.000 | 0.000 | 0.000 | 0.000 | 0.000 | 0.0035 | 0 | 0 | 0.0058 | 0 | 0 |
| Ascomycota | Sordariomycetes | Xylariales | Xylariaceae | Xylaria_feejeensis | 0.000 | 0.000 | 0.011 | 0.000 | 0.000 | 0 | 0 | 0.0055 | 0.029 | 0 | 0 |
| Ascomycota | Sordariomycetes | Xylariales | Xylariaceae | Xylaria_hypoxylon | 0.000 | 0.000 | 0.000 | 0.000 | 0.000 | 0 | 0.0016 | 0 | 0 | 0 | 0 |
| Ascomycota | Sordariomycetes | Xylariales | Xylariaceae | Xylaria_polymorpha | 0.000 | 0.000 | 0.000 | 0.000 | 0.000 | 0 | 0.0033 | 0.0055 | 0.029 | 0.0221 | 0.04 |

**Table S8 continued.**

| | | | | | | | | | | | | | | | |
|---|---|---|---|---|---|---|---|---|---|---|---|---|---|---|---|
| Unclassified | Sordariomycetes | Xylariales | Xylariaceae | Xylaria_primorskensis | 0.000 | 0.000 | 0.000 | 0.000 | 0.000 | 0 | 0 | 0.0388 | 0.029 | 0.0386 | 0.06 |
| Basidiomycota | Sordariomycetes | Xylariales | Xylariaceae | Xylaria_psidii | 0.000 | 0.000 | 0.000 | 0.000 | 0.000 | 0 | 0 | 0 | 0.0058 | 0.0055 | 0.02 |
| Basidiomycota | Sordariomycetes | Xylariales | Xylariaceae | Xylaria_sp. | 0.000 | 0.000 | 0.090 | 0.000 | 0.000 | 0.0037 | 0.0394 | 1.209 | 0.505 | 0.3422 | 0.2201 |
| Basidiomycota | Sordariomycetes | Xylariales | Xylariaceae | Xylariaceae_sp. | 0.003 | 0.000 | 0.197 | 0.002 | 0.005 | 0.1136 | 0.43 | 7.5255 | 18.262 | 23.029 | 5.9224 |
| Basidiomycota | Sordariomycetes | Xylariales | Xylariaceae | Xylariales_sp. | 0.000 | 0.000 | 0.011 | 0.000 | 0.000 | 0 | 0.0033 | 0.1165 | 0.1567 | 0.2705 | 0.06 |
| Basidiomycota | Sordariomycetes | Xylariales | Xylariaceae | Xylariomycetidae_sp. | 0.000 | 0.000 | 0.000 | 0.000 | 0.000 | 0 | 0 | 0 | 0.0116 | 0.0055 | 0 |

**Table S8 continued.**

### Table S9 :Relative abundances of the bacterial communities across degrading fungus comb obtained from Nanopore sequencing.

| Phylum | Class | Order | Family | Genus | 0 hrs | 24 hrs | 48 hrs | 72 hrs | 96 hrs | 120 hrs |
|---|---|---|---|---|---|---|---|---|---|---|
| Acidobacteria | Holophagae | Acanthopleuribacterales | Acanthopleuribacteraceae | Acanthopleuribacter | 0.000 | 0.000 | 0.016 | 0.000 | 0.000 | 0.000 |
| Acidobacteria | Acidobacteria | Acidobacteriales | Acidobacteriaceae | Acidobacterium | 0.000 | 0.000 | 0.091 | 0.000 | 0.000 | 0.000 |
| Acidobacteria | Acidobacteria | Acidobacteriales | Acidobacteriaceae | AMD_cluster | 0.000 | 0.000 | 0.034 | 0.000 | 0.000 | 0.000 |
| Acidobacteria | Acidobacteria | Acidobacteriales | Acidobacteriaceae | Candidatus_Koribacter | 0.000 | 0.000 | 0.069 | 0.000 | 0.000 | 0.000 |
| Acidobacteria | Acidobacteria | Acidobacteriales | Acidobacteriaceae | Telmatobacter | 0.000 | 0.000 | 0.048 | 0.000 | 0.000 | 0.000 |
| Acidobacteria | Acidobacteria | Acidobacteriales | Acidobacteriaceae | Terrimonas | 0.000 | 0.000 | 0.002 | 0.000 | 0.000 | 0.000 |
| Acidobacteria | Acidobacteria | Acidobacteriales | Acidobacteriaceae | Uncultured_11 | 0.016 | 0.012 | 0.951 | 0.000 | 0.000 | 0.000 |
| Acidobacteria | Acidobacteria | Acidobacteriales | Acidobacteriaceae | Uncultured_24 | 0.065 | 0.025 | 0.135 | 0.000 | 0.005 | 0.006 |
| Acidobacteria | Acidobacteria | Acidobacteriales | Acidobacteriaceae | Uncultured_28 | 0.008 | 0.000 | 0.129 | 0.006 | 0.000 | 0.000 |
| Acidobacteria | Acidobacteria | Acidobacteriales | Acidobacteriaceae | Uncultured_33 | 0.000 | 0.000 | 0.028 | 0.000 | 0.000 | 0.000 |
| Acidobacteria | Acidobacteria | Acidobacteriales | Acidobacteriaceae | Uncultured_b | 0.000 | 0.000 | 0.018 | 0.000 | 0.000 | 0.000 |
| Acidobacteria | Acidobacteriia | Acidobacteriales | Acidobacteriaceae | Candidatus_Chloroacidobacterium | 0.024 | 0.000 | 0.377 | 0.000 | 0.000 | 0.000 |
| Acidobacteria | Acidobacteriia | Acidobacteriales | Acidobacteriaceae | Candidatus_Solibacter | 0.016 | 0.000 | 0.405 | 0.006 | 0.000 | 0.006 |
| Acidobacteria | Acidobacteriia | Acidobacteriales | Acidobacteriaceae | Edaphobacter | 0.000 | 0.000 | 0.171 | 0.000 | 0.000 | 0.000 |
| Acidobacteria | Acidobacteriia | Acidobacteriales | Acidobacteriaceae | marine_benthic_group_2 | 0.000 | 0.000 | 0.030 | 0.000 | 0.000 | 0.000 |
| Acidobacteria | Acidobacteriia | Acidobacteriales | Acidobacteriaceae | marine_group | 0.033 | 0.000 | 0.167 | 0.000 | 0.000 | 0.000 |
| Acidobacteria | Acidobacteriia | Acidobacteriales | Acidobacteriaceae | Mixed_gut_cluster | 0.000 | 0.000 | 0.143 | 0.000 | 0.000 | 0.000 |
| Acidobacteria | Acidobacteriia | Acidobacteriales | Acidobacteriaceae | Uncultured_2 | 0.789 | 0.265 | 0.665 | 0.102 | 0.015 | 0.006 |
| Acidobacteria | Acidobacteriia | Acidobacteriales | Acidobacteriaceae | Uncultured_25 | 0.033 | 0.000 | 0.121 | 0.012 | 0.000 | 0.000 |
| Acidobacteria | Acidobacteriia | Acidobacteriales | Acidobacteriaceae | Uncultured_27 | 0.033 | 0.012 | 0.276 | 0.000 | 0.000 | 0.000 |
| Acidobacteria | Blastocatellia | Blastocatellales | Blastocatellaceae | Aridibacter | 0.016 | 0.000 | 0.000 | 0.000 | 0.000 | 0.000 |
| Acidobacteria | Blastocatellia | Blastocatellales | Blastocatellaceae | Armored_scale_insect_endosymbionts | 0.000 | 0.000 | 0.091 | 0.000 | 0.000 | 0.000 |
| Acidobacteria | Holophagae | Holophagales | Holophagaceae | Holophaga | 0.000 | 0.000 | 0.010 | 0.000 | 0.000 | 0.000 |
| Acidobacteria | Holophagae | Holophagales | Holophagaceae | Uncultured_30 | 0.024 | 0.006 | 0.109 | 0.000 | 0.000 | 0.000 |
| Acidobacteria | Holophagae | Holophagales | Holophagaceae | Uncultured_61 | 0.000 | 0.000 | 0.006 | 0.000 | 0.000 | 0.000 |
| Actinobacteria | Actinobacteria | Acidimicrobiales | Acidimicrobiaceae_1 | marine_group_AEGEAN-169 | 0.000 | 0.000 | 0.018 | 0.000 | 0.000 | 0.000 |
| Actinobacteria | Actinobacteria | Acidimicrobiales | Acidimicrobiaceae_1 | Uncultured_41 | 0.000 | 0.000 | 0.010 | 0.000 | 0.000 | 0.000 |
| Actinobacteria | Actinobacteria | Acidimicrobiales | Acidimicrobiaceae_2 | Acidimicrobium | 0.000 | 0.000 | 0.004 | 0.000 | 0.000 | 0.000 |
| Actinobacteria | Actinobacteria | Acidimicrobiales | Acidimicrobiaceae_2 | Ferrithrix | 0.000 | 0.000 | 0.004 | 0.000 | 0.000 | 0.000 |
| Actinobacteria | Actinobacteria | Actinomycetales | Acidothermaceae | Acidothermus | 0.000 | 0.000 | 0.018 | 0.000 | 0.000 | 0.000 |
| Actinobacteria | Actinobacteria | Actinomycetales | Actinomycetaceae | Actinomyces | 0.000 | 0.000 | 0.000 | 0.006 | 0.000 | 0.000 |
| Actinobacteria | Actinobacteria | Actinomycetales | Actinomycetaceae | Actinomyces_4 | 0.000 | 0.000 | 0.014 | 0.000 | 0.000 | 0.000 |

**Table S9 continued.**

| | | | | | | | | | | |
|---|---|---|---|---|---|---|---|---|---|---|
| Actinobacteria | Actinobacteria | Actinomycetales | Actinomycetaceae | Mobiluncus | 0.000 | 0.000 | 0.006 | 0.000 | 0.000 | 0.000 |
| Actinobacteria | Actinomycetia | Actinomycetales | Actinomycetaceae | Flaviflexus | 0.000 | 0.000 | 0.000 | 0.006 | 0.000 | 0.000 |
| Actinobacteria | Actinobacteria | Actinomycetales | Actinomycetaceae_2 | Actinobaculum | 0.000 | 0.000 | 0.016 | 0.000 | 0.000 | 0.000 |
| Actinobacteria | Actinobacteria | Actinomycetales | Actinomycetaceae_2 | Actinomyces_1 | 0.000 | 0.000 | 0.004 | 0.000 | 0.000 | 0.000 |
| Actinobacteria | Actinobacteria | Actinomycetales | Actinomycetaceae_2 | Actinomyces_2 | 0.000 | 0.000 | 0.044 | 0.000 | 0.000 | 0.000 |
| Actinobacteria | Actinobacteria | Actinomycetales | Actinomycetaceae_2 | Actinomyces_3 | 0.008 | 0.000 | 0.060 | 0.000 | 0.000 | 0.000 |
| Actinobacteria | Actinobacteria | Actinomycetales | Actinomycetaceae_2 | Arcanobacterium | 0.000 | 0.000 | 0.020 | 0.000 | 0.000 | 0.000 |
| Actinobacteria | Actinobacteria | Actinomycetales | Actinomycetaceae_2 | Varibaculum | 0.000 | 0.000 | 0.008 | 0.000 | 0.000 | 0.000 |
| Actinobacteria | Actinobacteria | Actinopolysporales | Actinopolysporaceae | Actinospica | 0.000 | 0.000 | 0.006 | 0.000 | 0.000 | 0.000 |
| Actinobacteria | Actinobacteria | Actinomycetales_2 | Actinosynnemataceae | Actinokineospora_1 | 0.000 | 0.000 | 0.010 | 0.000 | 0.000 | 0.000 |
| Actinobacteria | Actinobacteria | Actinomycetales_2 | Actinosynnemataceae | Leclercia | 0.049 | 0.271 | 0.034 | 0.126 | 0.020 | 0.139 |
| Actinobacteria | Actinobacteria | Pseudonocardiales_6 | Actinosynnemataceae | Actinosynnema | 0.000 | 0.000 | 0.008 | 0.000 | 0.000 | 0.006 |
| Actinobacteria | Actinobacteria | Pseudonocardiales_6 | Actinosynnemataceae | Lechevalieria | 0.000 | 0.000 | 0.004 | 0.000 | 0.000 | 0.000 |
| Actinobacteria | Actinobacteria | Pseudonocardiales_6 | Actinosynnemataceae | Lechevalieria_2 | 0.000 | 0.000 | 0.004 | 0.000 | 0.000 | 0.000 |
| Actinobacteria | Actinobacteria | Pseudonocardiales_6 | Actinosynnemataceae | Lechevalieria_3 | 0.000 | 0.000 | 0.004 | 0.000 | 0.000 | 0.000 |
| Actinobacteria | Actinobacteria | Pseudonocardiales_6 | Actinosynnemataceae | Salegentibacter | 0.000 | 0.000 | 0.010 | 0.000 | 0.000 | 0.000 |
| Actinobacteria | Coriobacteriia | Coriobacteriales | Atopobiaceae | Aurantimonas_1 | 0.000 | 0.000 | 0.004 | 0.000 | 0.000 | 0.000 |
| Actinobacteria | Actinobacteria | Actinomycetales | Beutenbergiaceae | Salana | 0.000 | 0.000 | 0.004 | 0.000 | 0.000 | 0.000 |
| Actinobacteria | Actinobacteria | Micrococcales | Beutenbergiaceae | Mitochondria | 0.033 | 0.012 | 0.145 | 0.042 | 0.010 | 0.006 |
| Actinobacteria | Actinobacteria | Micrococcales | Beutenbergiaceae | Serratia | 0.008 | 0.043 | 0.046 | 0.054 | 0.488 | 0.054 |
| Actinobacteria | Actinobacteria | Bifidobacteriales | Bifidobacteriaceae | Bifidibacterium | 0.000 | 0.000 | 0.103 | 0.000 | 0.000 | 0.000 |
| Actinobacteria | Actinobacteria | Bifidobacteriales | Bifidobacteriaceae | Gardnerella | 0.000 | 0.000 | 0.010 | 0.000 | 0.000 | 0.000 |
| Actinobacteria | Actinobacteria | Bifidobacteriales | Bifidobacteriaceae | Methylobacterium | 0.016 | 0.018 | 0.117 | 0.012 | 0.030 | 0.018 |
| Actinobacteria | Actinobacteria | Bifidobacteriales | Bifidobacteriaceae | Scardovia | 0.000 | 0.000 | 0.012 | 0.000 | 0.000 | 0.000 |
| Actinobacteria | Actinobacteria | Actinomycetales | Bogoriellaceae | Georgenia | 0.000 | 0.000 | 0.010 | 0.000 | 0.000 | 0.000 |
| Actinobacteria | Actinobacteria | Micrococcales | Brevibacteriaceae | Brevibacterium | 0.000 | 0.000 | 0.044 | 0.000 | 0.000 | 0.000 |
| Actinobacteria | Actinobacteria | Acidimicrobiales | Candidatus_Microthrix | Candidatus_Microthrix | 0.016 | 0.000 | 0.006 | 0.000 | 0.000 | 0.000 |
| Actinobacteria | Actinobacteria | Catenulisporales | Catenulisporaceae | Catenulispora | 0.000 | 0.000 | 0.004 | 0.000 | 0.000 | 0.000 |
| Actinobacteria | Actinobacteria | Actinomycetales | Cellulomonadaceae | Actinotalea | 0.000 | 0.000 | 0.010 | 0.000 | 0.000 | 0.000 |
| Actinobacteria | Actinobacteria | Actinomycetales | Cellulomonadaceae | Cellulomonas_1 | 0.000 | 0.006 | 0.032 | 0.000 | 0.000 | 0.000 |
| Actinobacteria | Actinobacteria | Actinomycetales | Cellulomonadaceae | Ohtaekwangia | 0.098 | 0.006 | 0.002 | 0.006 | 0.000 | 0.000 |
| Actinobacteria | Actinobacteria | Micrococcales_2 | Cellulomonadaceae | Cellulomonas | 0.016 | 0.006 | 0.000 | 0.000 | 0.005 | 0.012 |
| Actinobacteria | Actinobacteria | Micrococcales_3 | Cellulomonadaceae | Cellulomonas_2 | 0.000 | 0.000 | 0.010 | 0.000 | 0.000 | 0.000 |
| Actinobacteria | Actinobacteria | Micrococcales_3 | Cellulomonadaceae | Cellulomonas_3 | 0.000 | 0.000 | 0.004 | 0.000 | 0.000 | 0.000 |
| Actinobacteria | Thermoleophilia | Solirubrobacterales | Conexibacteraceae | Conexibacter | 0.000 | 0.000 | 0.004 | 0.000 | 0.000 | 0.000 |
| Actinobacteria | Actinobacteria | Coriobacteriales | Coriobacteriaceae | Collinsella | 0.000 | 0.000 | 0.119 | 0.000 | 0.000 | 0.000 |
| Actinobacteria | Actinobacteria | Coriobacteriales | Coriobacteriaceae | Coriobacterium | 0.000 | 0.000 | 0.004 | 0.000 | 0.000 | 0.000 |
| Actinobacteria | Actinobacteria | Coriobacteriales | Coriobacteriaceae | Olsenella | 0.000 | 0.000 | 0.004 | 0.000 | 0.000 | 0.000 |
| Actinobacteria | Actinobacteria | Coriobacteriales | Coriobacteriaceae | OM27_clade | 0.008 | 0.000 | 0.012 | 0.000 | 0.000 | 0.000 |
| Actinobacteria | Actinobacteria | Coriobacteriales | Coriobacteriaceae | Uncultured_12 | 0.130 | 0.043 | 0.508 | 0.030 | 0.000 | 0.006 |

**Table S9 continued.**

| | | | | | | | | | | |
|---|---|---|---|---|---|---|---|---|---|---|
| Actinobacteria | Actinobacteria | Coriobacteriales | Coriobacteriaceae | Uncultured_a | 0.024 | 0.006 | 0.046 | 0.000 | 0.000 | 0.000 |
| Actinobacteria | Actinobacteria | Corynebacteriales | Corynebacteriaceae | Corynebacterium | 0.008 | 0.000 | 0.000 | 0.000 | 0.000 | 0.000 |
| Actinobacteria | Actinobacteria | Corynebacteriales | Corynebacteriaceae | Corynebacterium_1 | 0.000 | 0.000 | 0.345 | 0.006 | 0.000 | 0.000 |
| Actinobacteria | Actinobacteria | Corynebacteriales | Corynebacteriaceae | Corynebacterium_2 | 0.000 | 0.000 | 0.004 | 0.000 | 0.000 | 0.000 |
| Actinobacteria | Actinobacteria | Corynebacteriales | Corynebacteriaceae | Corynebacterium_4 | 0.000 | 0.000 | 0.008 | 0.000 | 0.000 | 0.000 |
| Actinobacteria | Actinobacteria | Corynebacteriales | Corynebacteriaceae | Corynebacterium_5 | 0.000 | 0.000 | 0.012 | 0.000 | 0.000 | 0.000 |
| Actinobacteria | Actinobacteria | Corynebacteriales | Corynebacteriaceae | Corynebacterium_6 | 0.000 | 0.000 | 0.006 | 0.000 | 0.000 | 0.000 |
| Actinobacteria | Actinobacteria | Corynebacteriales | Corynebacteriaceae | Corynebacterium_8 | 0.000 | 0.000 | 0.022 | 0.000 | 0.000 | 0.000 |
| Actinobacteria | Actinobacteria | Actinomycetales | Cryptosporangiaceae | Cryptosporangium | 0.000 | 0.006 | 0.006 | 0.000 | 0.000 | 0.000 |
| Actinobacteria | Actinobacteria | Actinomycetales | Dermabacteraceae | Brachybacterium | 0.000 | 0.006 | 0.038 | 0.012 | 0.005 | 0.000 |
| Actinobacteria | Actinobacteria | Micrococcales_1 | Dermabacteraceae | Dermabacter | 0.000 | 0.000 | 0.008 | 0.000 | 0.005 | 0.000 |
| Actinobacteria | Actinobacteria | Actinomycetales | Dermacoccaceae | Dermacoccus | 0.000 | 0.000 | 0.008 | 0.000 | 0.000 | 0.000 |
| Actinobacteria | Actinobacteria | Micrococcales_4 | Dermacoccaceae | Demetria | 0.000 | 0.000 | 0.004 | 0.000 | 0.000 | 0.000 |
| Actinobacteria | Actinobacteria | Micrococcales | Dermatophilaceae | Dermatophilus | 0.008 | 0.000 | 0.028 | 0.000 | 0.000 | 0.000 |
| Actinobacteria | Actinobacteria | Corynebacteriales | Dietziaceae | Dietzia | 0.000 | 0.000 | 0.016 | 0.000 | 0.000 | 0.000 |
| Actinobacteria | Coriobacteriia | Eggerthellales | Eggerthellaceae | Eggerthella | 0.000 | 0.000 | 0.008 | 0.000 | 0.000 | 0.000 |
| Actinobacteria | Coriobacteriia | Eggerthellales | Eggerthellaceae | Enterorhabdus | 0.000 | 0.000 | 0.058 | 0.000 | 0.000 | 0.000 |
| Actinobacteria | Coriobacteriia | Eggerthellales | Eggerthellaceae | Gordonibacter_sp_1 | 0.000 | 0.000 | 0.010 | 0.000 | 0.000 | 0.000 |
| Actinobacteria | Coriobacteriia | Eggerthellales | Eggerthellaceae | RC9_gut_group | 0.057 | 0.012 | 0.792 | 0.024 | 0.000 | 0.000 |
| Actinobacteria | Nitriliruptoria | Euzebyales | Euzebyaceae | Euzebya | 0.016 | 0.000 | 0.000 | 0.000 | 0.000 | 0.000 |
| Actinobacteria | Actinobacteria | Actinomycetales_3 | Fodinicola | Fodinicola | 0.000 | 0.000 | 0.006 | 0.000 | 0.000 | 0.000 |
| Actinobacteria | Actinobacteria | Actinomycetales | Frankiaceae | Frankia | 0.000 | 0.000 | 0.024 | 0.000 | 0.000 | 0.000 |
| Actinobacteria | Actinobacteria | Actinomycetales | Geodermatophilaceae | Actinotelluria_sp_a | 0.000 | 0.000 | 0.004 | 0.000 | 0.000 | 0.000 |
| Actinobacteria | Actinobacteria | Actinomycetales | Geodermatophilaceae | Blastococcus_2 | 0.000 | 0.000 | 0.004 | 0.000 | 0.000 | 0.000 |
| Actinobacteria | Actinobacteria | Actinomycetales | Geodermatophilaceae | Blastococcus_3 | 0.000 | 0.000 | 0.004 | 0.000 | 0.000 | 0.000 |
| Actinobacteria | Actinobacteria | Actinomycetales | Geodermatophilaceae | Blastococcus_5 | 0.008 | 0.000 | 0.004 | 0.000 | 0.000 | 0.000 |
| Actinobacteria | Actinobacteria | Actinomycetales | Geodermatophilaceae | Modestobacter | 0.000 | 0.000 | 0.008 | 0.000 | 0.000 | 0.000 |
| Actinobacteria | Actinobacteria | Actinomycetales | Geodermatophilaceae | Mogibacterium | 0.000 | 0.000 | 0.038 | 0.000 | 0.000 | 0.000 |
| Actinobacteria | Actinobacteria | Frankiales | Geodermatophilaceae | Geodermatophilus | 0.000 | 0.000 | 0.010 | 0.000 | 0.000 | 0.006 |
| Actinobacteria | Actinobacteria | Glycomycetales | Glycomycetaceae | Glycomyces | 0.000 | 0.000 | 0.016 | 0.000 | 0.000 | 0.000 |
| Actinobacteria | Actinobacteria | Corynebacteriales | Gordoniaceae | Gordonia | 0.000 | 0.000 | 0.073 | 0.000 | 0.000 | 0.000 |
| Actinobacteria | Actinobacteria | Micrococcales | Intrasporangiaceae | Ktedonobacter | 0.000 | 0.000 | 0.435 | 0.000 | 0.000 | 0.000 |
| Actinobacteria | Actinobacteria | Micrococcales | Intrasporangiaceae | Phyllobacterium | 0.008 | 0.000 | 0.024 | 0.000 | 0.000 | 0.000 |
| Actinobacteria | Actinobacteria | Micrococcales | Intrasporangiaceae | Physiococcus | 0.008 | 0.000 | 0.014 | 0.000 | 0.000 | 0.000 |
| Actinobacteria | Actinobacteria | Micrococcales | Intrasporangiaceae | Serratia_1 | 0.000 | 0.086 | 0.069 | 0.066 | 0.344 | 0.091 |
| Actinobacteria | Actinobacteria | Micrococcales | Intrasporangiaceae | Serratia_2 | 0.000 | 0.000 | 0.056 | 0.000 | 0.005 | 0.000 |
| Actinobacteria | Actinomycetia | Micrococcales | Intrasporangiaceae | Intrasporangium | 0.008 | 0.000 | 0.000 | 0.000 | 0.000 | 0.000 |
| Actinobacteria | Actinobacteria | Actinomycetales | Intrasporangiaceae_1 | Humihabitans | 0.000 | 0.000 | 0.004 | 0.000 | 0.000 | 0.000 |
| Actinobacteria | Actinobacteria | Actinomycetales | Intrasporangiaceae_1 | Ornithinimicrobium | 0.000 | 0.000 | 0.012 | 0.000 | 0.000 | 0.000 |

**Table S9 continued.**

| | | | | | | | | | | |
|---|---|---|---|---|---|---|---|---|---|---|
| Actinobacteria | Actinobacteria | Actinomycetales | Intrasporangiaceae_1 | Oryzihumus | 0.000 | 0.000 | 0.010 | 0.000 | 0.000 | 0.000 |
| Actinobacteria | Actinobacteria | Actinomycetales | Intrasporangiaceae_1 | Phytobacter | 0.000 | 0.062 | 0.018 | 0.096 | 0.209 | 0.036 |
| Actinobacteria | Actinobacteria | Actinomycetales | Intrasporangiaceae_1 | Terrabacter | 0.000 | 0.000 | 0.000 | 0.000 | 0.000 | 0.006 |
| Actinobacteria | Actinobacteria | Actinomycetales | Intrasporangiaceae_1 | Terracoccus_sp_a | 0.000 | 0.000 | 0.006 | 0.000 | 0.000 | 0.000 |
| Actinobacteria | Actinobacteria | Actinomycetales | Intrasporangiaceae_1 | Terribacillus | 0.000 | 0.000 | 0.006 | 0.000 | 0.000 | 0.000 |
| Actinobacteria | Actinobacteria | Actinomycetales | Intrasporangiaceae_1 | Terriglobus | 0.000 | 0.000 | 0.016 | 0.000 | 0.000 | 0.000 |
| Actinobacteria | Actinobacteria | Micrococcales_4 | Intrasporangiaceae_1 | Arsenicicoccus | 0.000 | 0.000 | 0.006 | 0.000 | 0.000 | 0.000 |
| Actinobacteria | Actinobacteria | Micrococcales_4 | Intrasporangiaceae_1 | Larkinella | 0.000 | 0.000 | 0.004 | 0.000 | 0.000 | 0.000 |
| Actinobacteria | Actinobacteria | Micrococcales_4 | Intrasporangiaceae_1 | Ornithinicoccus | 0.000 | 0.000 | 0.006 | 0.000 | 0.000 | 0.000 |
| Actinobacteria | Actinobacteria | Micrococcales_4 | Intrasporangiaceae_1 | Oscillibacter | 0.016 | 0.012 | 0.316 | 0.006 | 0.000 | 0.000 |
| Actinobacteria | Actinobacteria | Actinomycetales | Jonesiaceae | Jonesia | 0.000 | 0.000 | 0.004 | 0.000 | 0.000 | 0.000 |
| Actinobacteria | Actinobacteria | Actinomycetales | Kineosporiaceae | Kinetoplastibacterium | 0.000 | 0.000 | 0.004 | 0.012 | 0.005 | 0.006 |
| Actinobacteria | Actinobacteria | Kineosporiales | Kineosporiaceae | Kineosporia | 0.000 | 0.006 | 0.012 | 0.000 | 0.000 | 0.000 |
| Actinobacteria | Actinobacteria | Kineosporiales | Kineosporiaceae | Rahnella | 0.000 | 0.012 | 0.040 | 0.000 | 0.005 | 0.024 |
| Actinobacteria | Actinobacteria | Kineosporiales | Kineosporiaceae | Thauera | 0.000 | 0.000 | 0.036 | 0.000 | 0.005 | 0.000 |
| Actinobacteria | Actinobacteria | Actinomycetales | Microbacteriaceae | Cryobacterium_1 | 0.000 | 0.000 | 0.016 | 0.000 | 0.000 | 0.000 |
| Actinobacteria | Actinobacteria | Actinomycetales | Microbacteriaceae | Labrenzia | 0.000 | 0.000 | 0.016 | 0.000 | 0.000 | 0.000 |
| Actinobacteria | Actinobacteria | Actinomycetales | Microbacteriaceae | Leuconostoc | 0.000 | 0.000 | 0.050 | 0.000 | 0.000 | 0.000 |
| Actinobacteria | Actinobacteria | Actinomycetales | Microbacteriaceae | Mycetocola | 0.000 | 0.000 | 0.010 | 0.000 | 0.000 | 0.000 |
| Actinobacteria | Actinobacteria | Actinomycetales | Microbacteriaceae | Mycobacterium | 0.000 | 0.000 | 0.234 | 0.000 | 0.000 | 0.000 |
| Actinobacteria | Actinobacteria | Actinomycetales | Microbacteriaceae | Okibacterium | 0.000 | 0.000 | 0.004 | 0.006 | 0.000 | 0.000 |
| Actinobacteria | Actinobacteria | Actinomycetales | Microbacteriaceae | Rathayibacter | 0.000 | 0.000 | 0.010 | 0.006 | 0.000 | 0.000 |
| Actinobacteria | Actinobacteria | Actinomycetales | Microbacteriaceae | Reinekea | 0.000 | 0.000 | 0.008 | 0.000 | 0.000 | 0.000 |
| Actinobacteria | Actinobacteria | Actinomycetales | Microbacteriaceae | Salimicrobium | 0.000 | 0.000 | 0.008 | 0.000 | 0.000 | 0.000 |
| Actinobacteria | Actinobacteria | Actinomycetales | Microbacteriaceae | Subtercola | 0.000 | 0.000 | 0.004 | 0.000 | 0.000 | 0.000 |
| Actinobacteria | Actinobacteria | Actinomycetales | Microbacteriaceae | Uncultured_21 | 0.024 | 0.012 | 0.093 | 0.000 | 0.005 | 0.000 |
| Actinobacteria | Actinobacteria | Actinomycetales | Microbacteriaceae | Zimmermanella_3 | 0.000 | 0.000 | 0.004 | 0.000 | 0.000 | 0.000 |
| Actinobacteria | Actinobacteria | Micrococcales | Microbacteriaceae | Agromyces_2 | 0.008 | 0.000 | 0.036 | 0.000 | 0.000 | 0.000 |
| Actinobacteria | Actinobacteria | Micrococcales | Microbacteriaceae | Alloiococcus | 0.000 | 0.000 | 0.006 | 0.000 | 0.000 | 0.000 |
| Actinobacteria | Actinobacteria | Micrococcales | Microbacteriaceae | Leifsonia_1 | 0.000 | 0.000 | 0.012 | 0.000 | 0.000 | 0.000 |
| Actinobacteria | Actinobacteria | Micrococcales | Microbacteriaceae | Leifsonia_3 | 0.000 | 0.000 | 0.006 | 0.000 | 0.000 | 0.000 |
| Actinobacteria | Actinobacteria | Micrococcales | Microbacteriaceae | Leifsonia_7 | 0.000 | 0.000 | 0.008 | 0.000 | 0.000 | 0.000 |
| Actinobacteria | Actinomycetia | Micrococcales | Microbacteriaceae | Compostimonas | 0.008 | 0.000 | 0.000 | 0.000 | 0.000 | 0.000 |
| Actinobacteria | Actinomycetia | Micrococcales | Microbacteriaceae | Gryllotalpicola | 0.008 | 0.000 | 0.000 | 0.000 | 0.000 | 0.000 |
| Actinobacteria | Actinobacteria | Micrococcales_3 | Microbacteriaceae | Agrobacterium | 0.016 | 0.000 | 0.002 | 0.000 | 0.000 | 0.018 |
| Actinobacteria | Actinobacteria | Micrococcales_3 | Microbacteriaceae | Agrococcus | 0.000 | 0.000 | 0.018 | 0.000 | 0.000 | 0.000 |
| Actinobacteria | Actinobacteria | Micrococcales_3 | Microbacteriaceae | Candidatus_Aquiluna | 0.000 | 0.000 | 0.004 | 0.000 | 0.000 | 0.000 |
| Actinobacteria | Actinobacteria | Micrococcales_3 | Microbacteriaceae | Candidatus_Flavinula_a | 0.000 | 0.000 | 0.004 | 0.000 | 0.000 | 0.000 |
| Actinobacteria | Actinobacteria | Micrococcales_3 | Microbacteriaceae | Clavibacter | 0.000 | 0.000 | 0.010 | 0.006 | 0.000 | 0.006 |

**Table S9 continued.**

| | | | | | | | | | | |
|---|---|---|---|---|---|---|---|---|---|---|
| Actinobacteria | Actinobacteria | Micrococcales_3 | Microbacteriaceae | Curtobacterium | 0.008 | 0.000 | 0.030 | 0.012 | 0.010 | 0.000 |
| Actinobacteria | Actinobacteria | Micrococcales_3 | Microbacteriaceae | Frigoribacterium_1 | 0.000 | 0.000 | 0.046 | 0.000 | 0.000 | 0.000 |
| Actinobacteria | Actinobacteria | Micrococcales_3 | Microbacteriaceae | Frondihabitans | 0.000 | 0.000 | 0.004 | 0.000 | 0.000 | 0.000 |
| Actinobacteria | Actinobacteria | Micrococcales_3 | Microbacteriaceae | Leifsonia_5 | 0.000 | 0.000 | 0.006 | 0.000 | 0.000 | 0.000 |
| Actinobacteria | Actinobacteria | Micrococcales_3 | Microbacteriaceae | Salinibacterium_3 | 0.000 | 0.000 | 0.004 | 0.000 | 0.000 | 0.000 |
| Actinobacteria | Actinobacteria | Micrococcales_3 | Microbacteriaceae | Schumannella | 0.000 | 0.000 | 0.004 | 0.000 | 0.000 | 0.000 |
| Actinobacteria | Actinobacteria | Micrococcales_3 | Microbacteriaceae | Sebaldella | 0.000 | 0.000 | 0.004 | 0.000 | 0.000 | 0.000 |
| Actinobacteria | Actinobacteria | Micrococcales_3 | Microbacteriaceae | Zimmermanella_2 | 0.000 | 0.000 | 0.004 | 0.000 | 0.000 | 0.000 |
| Actinobacteria | Actinobacteria | Micrococcales_3 | Microbacteriaceae | Zimmermannella_1 | 0.000 | 0.000 | 0.004 | 0.000 | 0.000 | 0.000 |
| Actinobacteria | Actinobacteria | Micrococcales_3 | Microbacteriaceae | Zoogloea | 0.016 | 0.000 | 0.038 | 0.006 | 0.010 | 0.012 |
| Actinobacteria | Actinobacteria | Micrococcales_4 | Microbacteriaceae | Micrococcus | 0.008 | 0.000 | 0.002 | 0.000 | 0.005 | 0.000 |
| Actinobacteria | Actinobacteria | Actinomycetales | Micrococcaceae | Rs-D38_termite_group | 0.163 | 0.080 | 0.030 | 0.102 | 0.005 | 0.000 |
| Actinobacteria | Actinobacteria | Actinomycetales | Micrococcaceae | Sinomonas | 0.016 | 0.000 | 0.012 | 0.000 | 0.000 | 0.000 |
| Actinobacteria | Actinobacteria | Actinomycetales | Micrococcaceae | Sinorhizobium-Ensifer | 0.024 | 0.006 | 0.058 | 0.000 | 0.000 | 0.012 |
| Actinobacteria | Actinobacteria | Micrococcales | Micrococcaceae | Kocuria_1 | 0.000 | 0.000 | 0.018 | 0.000 | 0.000 | 0.000 |
| Actinobacteria | Actinobacteria | Micrococcales | Micrococcaceae | Kocuria_2 | 0.000 | 0.000 | 0.040 | 0.006 | 0.010 | 0.012 |
| Actinobacteria | Actinobacteria | Micrococcales | Micrococcaceae | Kofleria | 0.008 | 0.000 | 0.000 | 0.000 | 0.000 | 0.000 |
| Actinobacteria | Actinobacteria | Micrococcales | Micrococcaceae | Nevskia | 0.033 | 0.000 | 0.012 | 0.018 | 0.000 | 0.000 |
| Actinobacteria | Actinobacteria | Micrococcales_1 | Micrococcaceae | Arhtrobacter_12 | 0.000 | 0.000 | 0.010 | 0.000 | 0.000 | 0.000 |
| Actinobacteria | Actinobacteria | Micrococcales_1 | Micrococcaceae | Arthrobacter | 0.065 | 0.012 | 0.002 | 0.000 | 0.005 | 0.000 |
| Actinobacteria | Actinobacteria | Micrococcales_1 | Micrococcaceae | Arthrobacter_1 | 0.008 | 0.000 | 0.022 | 0.000 | 0.000 | 0.000 |
| Actinobacteria | Actinobacteria | Micrococcales_1 | Micrococcaceae | Arthrobacter_10 | 0.008 | 0.000 | 0.024 | 0.000 | 0.000 | 0.000 |
| Actinobacteria | Actinobacteria | Micrococcales_1 | Micrococcaceae | Arthrobacter_11 | 0.000 | 0.000 | 0.006 | 0.000 | 0.000 | 0.000 |
| Actinobacteria | Actinobacteria | Micrococcales_1 | Micrococcaceae | Arthrobacter_13 | 0.000 | 0.000 | 0.006 | 0.000 | 0.000 | 0.000 |
| Actinobacteria | Actinobacteria | Micrococcales_1 | Micrococcaceae | Arthrobacter_14 | 0.008 | 0.000 | 0.034 | 0.000 | 0.000 | 0.000 |
| Actinobacteria | Actinobacteria | Micrococcales_1 | Micrococcaceae | Arthrobacter_17 | 0.049 | 0.006 | 0.091 | 0.000 | 0.000 | 0.000 |
| Actinobacteria | Actinobacteria | Micrococcales_1 | Micrococcaceae | Arthrobacter_20 | 0.000 | 0.000 | 0.008 | 0.000 | 0.000 | 0.000 |
| Actinobacteria | Actinobacteria | Micrococcales_1 | Micrococcaceae | Arthrobacter_21 | 0.000 | 0.000 | 0.105 | 0.000 | 0.000 | 0.000 |
| Actinobacteria | Actinobacteria | Micrococcales_1 | Micrococcaceae | Arthrobacter_5 | 0.008 | 0.000 | 0.008 | 0.000 | 0.000 | 0.000 |
| Actinobacteria | Actinobacteria | Micrococcales_1 | Micrococcaceae | Arthrobacter_6 | 0.000 | 0.012 | 0.006 | 0.000 | 0.000 | 0.006 |
| Actinobacteria | Actinobacteria | Micrococcales_1 | Micrococcaceae | Arthrobacter_8 | 0.000 | 0.000 | 0.008 | 0.000 | 0.000 | 0.000 |
| Actinobacteria | Actinobacteria | Micrococcales_1 | Micrococcaceae | Arthrobacter_9 | 0.000 | 0.000 | 0.004 | 0.000 | 0.000 | 0.000 |
| Actinobacteria | Actinobacteria | Micrococcales_1 | Micrococcaceae | Citricoccus | 0.000 | 0.000 | 0.006 | 0.000 | 0.000 | 0.000 |
| Actinobacteria | Actinobacteria | Micrococcales_1 | Micrococcaceae | Micrococcus_1 | 0.000 | 0.000 | 0.044 | 0.000 | 0.005 | 0.000 |
| Actinobacteria | Actinobacteria | Micrococcales_1 | Micrococcaceae | Micrococcus_3 | 0.000 | 0.000 | 0.004 | 0.000 | 0.000 | 0.000 |
| Actinobacteria | Actinobacteria | Micrococcales_1 | Micrococcaceae | Microcoleus_3 | 0.000 | 0.000 | 0.006 | 0.000 | 0.000 | 0.000 |
| Actinobacteria | Actinobacteria | Micrococcales_1 | Micrococcaceae | Renibacterium | 0.000 | 0.000 | 0.004 | 0.000 | 0.000 | 0.000 |
| Actinobacteria | Actinobacteria | Micrococcales_1 | Micrococcaceae | Yaniella | 0.000 | 0.000 | 0.006 | 0.000 | 0.000 | 0.000 |
| Actinobacteria | Actinobacteria | Actinomycetales | Micromonosporaceae | Polymorphospora | 0.000 | 0.000 | 0.006 | 0.000 | 0.000 | 0.000 |
| Actinobacteria | Actinobacteria | Actinomycetales | Micromonosporaceae | Polynucleobacter | 0.000 | 0.000 | 0.026 | 0.000 | 0.005 | 0.000 |

**Table S9 continued.**

| | | | | | | | | | | |
|---|---|---|---|---|---|---|---|---|---|---|
| Actinobacteria | Actinobacteria | Actinomycetales | Micromonosporaceae | Spiroplasma_1 | 0.000 | 0.000 | 0.016 | 0.000 | 0.000 | 0.000 |
| Actinobacteria | Actinobacteria | Actinomycetales_4 | Micromonosporaceae | Actinocatenispora | 0.000 | 0.000 | 0.004 | 0.000 | 0.000 | 0.000 |
| Actinobacteria | Actinobacteria | Actinomycetales_4 | Micromonosporaceae | Actinoplanes_1 | 0.000 | 0.000 | 0.026 | 0.000 | 0.005 | 0.000 |
| Actinobacteria | Actinobacteria | Actinomycetales_4 | Micromonosporaceae | Actinoplanes_2 | 0.000 | 0.000 | 0.004 | 0.000 | 0.000 | 0.000 |
| Actinobacteria | Actinobacteria | Actinomycetales_4 | Micromonosporaceae | Actinoplanes_3 | 0.000 | 0.000 | 0.004 | 0.000 | 0.000 | 0.000 |
| Actinobacteria | Actinobacteria | Actinomycetales_4 | Micromonosporaceae | Actinoplanes_4 | 0.000 | 0.000 | 0.012 | 0.000 | 0.000 | 0.000 |
| Actinobacteria | Actinobacteria | Actinomycetales_4 | Micromonosporaceae | Asanoa | 0.000 | 0.000 | 0.006 | 0.000 | 0.000 | 0.000 |
| Actinobacteria | Actinobacteria | Actinomycetales_4 | Micromonosporaceae | Catelliglobosispora | 0.016 | 0.000 | 0.008 | 0.000 | 0.000 | 0.000 |
| Actinobacteria | Actinobacteria | Actinomycetales_4 | Micromonosporaceae | Dactylosporangium | 0.000 | 0.000 | 0.016 | 0.000 | 0.000 | 0.000 |
| Actinobacteria | Actinobacteria | Actinomycetales_4 | Micromonosporaceae | Hamadaea | 0.000 | 0.000 | 0.004 | 0.000 | 0.000 | 0.000 |
| Actinobacteria | Actinobacteria | Actinomycetales_4 | Micromonosporaceae | Kribbella | 0.000 | 0.000 | 0.014 | 0.000 | 0.000 | 0.000 |
| Actinobacteria | Actinobacteria | Actinomycetales_4 | Micromonosporaceae | Luedemannella | 0.000 | 0.000 | 0.004 | 0.000 | 0.000 | 0.000 |
| Actinobacteria | Actinobacteria | Actinomycetales_4 | Micromonosporaceae | Micromonosporaceae | 0.000 | 0.000 | 0.004 | 0.000 | 0.000 | 0.000 |
| Actinobacteria | Actinobacteria | Actinomycetales_4 | Micromonosporaceae | Microvirga | 0.008 | 0.006 | 0.006 | 0.000 | 0.000 | 0.000 |
| Actinobacteria | Actinobacteria | Actinomycetales_4 | Micromonosporaceae | Planosporangium | 0.000 | 0.000 | 0.004 | 0.000 | 0.000 | 0.000 |
| Actinobacteria | Actinobacteria | Actinomycetales_4 | Micromonosporaceae | Staphylococcus_1 | 0.008 | 0.006 | 0.202 | 0.000 | 0.005 | 0.012 |
| Actinobacteria | Actinobacteria | Actinomycetales_4 | Micromonosporaceae | Uncultured_10 | 0.358 | 0.136 | 0.324 | 0.066 | 0.000 | 0.006 |
| Actinobacteria | Actinobacteria | Actinomycetales_4 | Micromonosporaceae | Verrucosispora | 0.008 | 0.000 | 0.006 | 0.000 | 0.005 | 0.006 |
| Actinobacteria | Actinobacteria | Actinomycetales_4 | Micromonosporaceae | Vibrionimonas | 0.000 | 0.006 | 0.000 | 0.000 | 0.000 | 0.000 |
| Actinobacteria | Actinobacteria | Actinomycetales_4 | Micromonosporaceae | Vogesella | 0.000 | 0.000 | 0.010 | 0.000 | 0.000 | 0.000 |
| Actinobacteria | Actinobacteria | Micromonosporales | Micromonosporaceae | Actinoplanes | 0.000 | 0.000 | 0.000 | 0.000 | 0.005 | 0.000 |
| Actinobacteria | Actinobacteria | Micromonosporales | Micromonosporaceae | Actinoplanes_5 | 0.000 | 0.000 | 0.026 | 0.000 | 0.000 | 0.000 |
| Actinobacteria | Actinobacteria | Micromonosporales | Micromonosporaceae | Actinoplanes_6 | 0.000 | 0.000 | 0.016 | 0.000 | 0.000 | 0.000 |
| Actinobacteria | Actinobacteria | Micromonosporales | Micromonosporaceae | Actinoplanes_7 | 0.000 | 0.000 | 0.004 | 0.000 | 0.000 | 0.000 |
| Actinobacteria | Actinobacteria | Micromonosporales | Micromonosporaceae | Longispora | 0.000 | 0.000 | 0.004 | 0.000 | 0.000 | 0.000 |
| Actinobacteria | Actinobacteria | Micromonosporales | Micromonosporaceae | Micromonospora_1 | 0.000 | 0.000 | 0.034 | 0.000 | 0.005 | 0.000 |
| Actinobacteria | Actinobacteria | Micromonosporales | Micromonosporaceae | Micromonospora_4 | 0.000 | 0.000 | 0.004 | 0.000 | 0.000 | 0.000 |
| Actinobacteria | Actinobacteria | Micromonosporales | Micromonosporaceae | Micromonospora_9 | 0.000 | 0.000 | 0.008 | 0.000 | 0.000 | 0.000 |
| Actinobacteria | Actinobacteria | Micromonosporales | Micromonosporaceae | Pilimelia | 0.000 | 0.000 | 0.006 | 0.000 | 0.000 | 0.000 |
| Actinobacteria | Actinobacteria | Micromonosporales | Micromonosporaceae | Pseudoxanthobacter | 0.008 | 0.000 | 0.004 | 0.000 | 0.000 | 0.000 |
| Actinobacteria | Actinobacteria | Micromonosporales | Micromonosporaceae | Salinispora | 0.000 | 0.000 | 0.006 | 0.000 | 0.000 | 0.000 |
| Actinobacteria | Actinobacteria | Micromonosporales | Micromonosporaceae | Stackebrandtia | 0.000 | 0.000 | 0.004 | 0.000 | 0.000 | 0.000 |
| Actinobacteria | Actinomycetia | Micromonosporales | Micromonosporaceae | Phytomonospora | 0.000 | 0.000 | 0.000 | 0.000 | 0.005 | 0.000 |
| Actinobacteria | Actinomycetia | Micromonosporales | Micromonosporaceae | Plantactinospora | 0.000 | 0.000 | 0.000 | 0.000 | 0.010 | 0.000 |
| Actinobacteria | Actinomycetia | Micromonosporales | Micromonosporaceae | Xiangella | 0.000 | 0.000 | 0.000 | 0.000 | 0.000 | 0.006 |
| Actinobacteria | Actinobacteria | Micrococcales | Mircrobacteriaceae | Pseudoclavibacter_1 | 0.000 | 0.000 | 0.004 | 0.000 | 0.000 | 0.000 |
| Actinobacteria | Actinobacteria | Micromonosporales | Mircromonasporaceae | Catenuloplanes | 0.000 | 0.000 | 0.010 | 0.000 | 0.000 | 0.000 |
| Actinobacteria | Actinobacteria | Actinomycetales_2 | Mycobacteriaceae | Mycoplana | 0.000 | 0.000 | 0.004 | 0.000 | 0.005 | 0.000 |
| Actinobacteria | Actinobacteria | Actinomycetales | Nakamurellaceae | Saxeibacter | 0.000 | 0.000 | 0.008 | 0.000 | 0.000 | 0.000 |
| Actinobacteria | Actinobacteria | Nitriliruptorales | Nitriliruptoraceae | Nitriliruptor | 0.000 | 0.000 | 0.008 | 0.000 | 0.000 | 0.000 |

**Table S9 continued.**

| | | | | | | | | | | |
|---|---|---|---|---|---|---|---|---|---|---|
| Actinobacteria | Actinobacteria | Actinomycetales_2 | Nocardiaceae | SM1A02 | 0.000 | 0.000 | 0.085 | 0.000 | 0.000 | 0.000 |
| Actinobacteria | Actinobacteria | Corynebacteriales | Nocardiaceae | Nocardioides | 0.146 | 0.018 | 0.157 | 0.012 | 0.000 | 0.000 |
| Actinobacteria | Actinobacteria | Corynebacteriales | Nocardiaceae | Rhodococcus_1 | 0.000 | 0.000 | 0.042 | 0.000 | 0.000 | 0.006 |
| Actinobacteria | Actinobacteria | Corynebacteriales | Nocardiaceae | Rhodococcus_2 | 0.000 | 0.000 | 0.073 | 0.000 | 0.000 | 0.000 |
| Actinobacteria | Actinobacteria | Corynebacteriales | Nocardiaceae | Rhodococcus_3 | 0.000 | 0.000 | 0.010 | 0.000 | 0.000 | 0.000 |
| Actinobacteria | Actinobacteria | Corynebacteriales | Nocardiaceae | Rhodocyclus | 0.000 | 0.000 | 0.006 | 0.000 | 0.000 | 0.000 |
| Actinobacteria | Actinobacteria | Corynebacteriales | Nocardiaceae | Rhodoferax | 0.000 | 0.006 | 0.208 | 0.000 | 0.010 | 0.000 |
| Actinobacteria | Actinobacteria | Corynebacteriales | Nocardiaceae | Solimonas | 0.008 | 0.000 | 0.004 | 0.006 | 0.000 | 0.000 |
| Actinobacteria | Actinobacteria | Actinomycetales_2 | Nocardiaceae_1 | Williamsia | 0.000 | 0.000 | 0.016 | 0.000 | 0.000 | 0.000 |
| Actinobacteria | Actinobacteria | Actinomycetales_2 | Nocardiaceae_1 | Wolbachia | 9.835 | 0.924 | 0.034 | 0.701 | 0.035 | 0.030 |
| Actinobacteria | Actinobacteria | Actinomycetales | Nocardioidaceae | Actinopolymorpha | 0.000 | 0.000 | 0.006 | 0.000 | 0.000 | 0.000 |
| Actinobacteria | Actinobacteria | Actinomycetales | Nocardioidaceae | Aeromicrobium | 0.008 | 0.000 | 0.024 | 0.000 | 0.000 | 0.000 |
| Actinobacteria | Actinobacteria | Propionibacteriales | Nocardioidaceae | Actinopolyspora | 0.000 | 0.000 | 0.008 | 0.000 | 0.000 | 0.000 |
| Actinobacteria | Actinobacteria | Propionibacteriales | Nocardioidaceae | Aeromonas | 0.000 | 0.006 | 0.000 | 0.000 | 0.000 | 0.000 |
| Actinobacteria | Actinobacteria | Propionibacteriales | Nocardioidaceae | Friedmanniella | 0.000 | 0.006 | 0.024 | 0.000 | 0.000 | 0.000 |
| Actinobacteria | Actinobacteria | Propionibacteriales | Nocardioidaceae | Marvinbryantia | 0.016 | 0.000 | 0.081 | 0.000 | 0.000 | 0.000 |
| Actinobacteria | Actinobacteria | Propionibacteriales | Nocardioidaceae | Propionicimonas_2 | 0.000 | 0.000 | 0.004 | 0.000 | 0.000 | 0.000 |
| Actinobacteria | Actinobacteria | Streptosporangiales | Nocardiopsaceae | Haloactinospora | 0.000 | 0.000 | 0.004 | 0.000 | 0.000 | 0.000 |
| Actinobacteria | Actinobacteria | Streptosporangiales | Nocardiopsaceae | Nocardiopsis_1 | 0.000 | 0.000 | 0.034 | 0.000 | 0.000 | 0.000 |
| Actinobacteria | Actinobacteria | Streptosporangiales | Nocardiopsaceae | Nocardiopsis_2 | 0.000 | 0.000 | 0.004 | 0.000 | 0.000 | 0.000 |
| Actinobacteria | Actinobacteria | Streptosporangiales | Nocardiopsaceae | Streptomonospora | 0.000 | 0.000 | 0.006 | 0.000 | 0.000 | 0.000 |
| Actinobacteria | Actinobacteria | Streptosporangiales | Nocardiopsaceae | Thermobifida | 0.000 | 0.000 | 0.006 | 0.000 | 0.000 | 0.000 |
| Actinobacteria | Thermoleophilia | Solirubrobacterales | Parviterribacteraceae | Parvularcula | 0.065 | 0.006 | 0.016 | 0.030 | 0.000 | 0.000 |
| Actinobacteria | Actinobacteria | Actinomycetales | Promicromonospiraceae | Cellulosimicrobium | 0.024 | 0.000 | 0.022 | 0.000 | 0.000 | 0.012 |
| Actinobacteria | Actinobacteria | Actinomycetales | Promicromonosporaceae | Yokenella | 0.008 | 0.018 | 0.030 | 0.072 | 0.060 | 0.036 |
| Actinobacteria | Actinobacteria | Micrococcales | Promicromonosporaceae | Isoptericola_1 | 0.000 | 0.000 | 0.004 | 0.000 | 0.000 | 0.000 |
| Actinobacteria | Actinobacteria | Micrococcales | Promicromonosporaceae | Isoptericola_3 | 0.000 | 0.000 | 0.004 | 0.000 | 0.000 | 0.000 |
| Actinobacteria | Actinobacteria | Micrococcales | Promicromonosporaceae | Isosphaera | 0.000 | 0.000 | 0.060 | 0.000 | 0.000 | 0.000 |
| Actinobacteria | Actinobacteria | Actinomycetales | Propionibacteriaceae | Aestuariimicrobium | 0.008 | 0.000 | 0.024 | 0.012 | 0.000 | 0.000 |
| Actinobacteria | Actinobacteria | Actinomycetales | Propionibacteriaceae | Microlunatus | 0.000 | 0.000 | 0.018 | 0.000 | 0.000 | 0.000 |
| Actinobacteria | Actinobacteria | Actinomycetales | Propionibacteriaceae | Micromonospora | 0.000 | 0.000 | 0.000 | 0.000 | 0.010 | 0.036 |
| Actinobacteria | Actinobacteria | Actinomycetales | Propionibacteriaceae | Propionispora | 0.000 | 0.000 | 0.004 | 0.000 | 0.000 | 0.000 |
| Actinobacteria | Actinobacteria | Actinomycetales | Propionibacteriaceae | Tetrasphaera | 0.000 | 0.000 | 0.026 | 0.000 | 0.000 | 0.000 |
| Actinobacteria | Actinobacteria | Actinomycetales_3 | Propionibacteriaceae | Jiangella | 0.000 | 0.000 | 0.010 | 0.000 | 0.000 | 0.000 |
| Actinobacteria | Actinobacteria | Actinomycetales_3 | Propionibacteriaceae | Johnsonella | 0.000 | 0.000 | 0.032 | 0.000 | 0.000 | 0.000 |
| Actinobacteria | Actinobacteria | Propionibacteriales | Propionibacteriaceae | Micropruina | 0.000 | 0.000 | 0.004 | 0.000 | 0.000 | 0.000 |
| Actinobacteria | Actinobacteria | Propionibacteriales | Propionibacteriaceae | Propionibacterium_1 | 0.000 | 0.000 | 0.097 | 0.000 | 0.000 | 0.000 |
| Actinobacteria | Actinobacteria | Propionibacteriales | Propionibacteriaceae | Propionicicella | 0.000 | 0.000 | 0.006 | 0.000 | 0.000 | 0.000 |
| Actinobacteria | Actinobacteria | Propionibacteriales | Propionibacteriaceae | Propioniferax | 0.000 | 0.000 | 0.004 | 0.000 | 0.000 | 0.000 |
| Actinobacteria | Actinobacteria | Propionibacteriales | Propionibacteriaceae | Propionimicrobium_sp_a | 0.000 | 0.000 | 0.004 | 0.000 | 0.000 | 0.000 |

**Table S9 continued.**

| | | | | | | | | | | |
|---|---|---|---|---|---|---|---|---|---|---|
| Actinobacteria | Actinobacteria | Actinomycetales | Pseudocardiaceae | Allokutzneria | 0.008 | 0.000 | 0.000 | 0.000 | 0.000 | 0.000 |
| Actinobacteria | Actinobacteria | Actinomycetales | Pseudonocardiaceae | Crossiella | 0.000 | 0.000 | 0.006 | 0.000 | 0.000 | 0.000 |
| Actinobacteria | Actinobacteria | Actinomycetales_2 | Pseudonocardiaceae | Actinoalloteichus | 0.000 | 0.000 | 0.008 | 0.000 | 0.000 | 0.000 |
| Actinobacteria | Actinobacteria | Pseudonocardiales | Pseudonocardiaceae | Actinomycetospora | 0.008 | 0.000 | 0.000 | 0.000 | 0.000 | 0.000 |
| Actinobacteria | Actinobacteria | Pseudonocardiales | Pseudonocardiaceae | Actinophytocola | 0.000 | 0.000 | 0.002 | 0.000 | 0.000 | 0.000 |
| Actinobacteria | Actinobacteria | Pseudonocardiales | Pseudonocardiaceae | Lentzea | 0.024 | 0.000 | 0.000 | 0.000 | 0.000 | 0.000 |
| Actinobacteria | Actinobacteria | Pseudonocardiales | Pseudonocardiaceae | Lentzea_1 | 0.000 | 0.000 | 0.010 | 0.000 | 0.000 | 0.000 |
| Actinobacteria | Actinobacteria | Pseudonocardiales | Pseudonocardiaceae | Saccharomonospora_1 | 0.000 | 0.000 | 0.010 | 0.000 | 0.000 | 0.000 |
| Actinobacteria | Actinobacteria | Pseudonocardiales | Pseudonocardiaceae | Saccharothrix | 0.008 | 0.000 | 0.018 | 0.006 | 0.000 | 0.000 |
| Actinobacteria | Actinobacteria | Actinomycetales_2 | Pseudonocardiaceae_1 | Sciscionella | 0.000 | 0.000 | 0.004 | 0.000 | 0.000 | 0.000 |
| Actinobacteria | Actinobacteria | Actinomycetales_2 | Pseudonocardiaceae_1 | Sedimentibacter | 0.000 | 0.006 | 0.022 | 0.000 | 0.000 | 0.000 |
| Actinobacteria | Actinobacteria | Actinomycetales_2 | Pseudonocardiaceae_1 | Thermocrispum | 0.000 | 0.000 | 0.004 | 0.000 | 0.000 | 0.000 |
| Actinobacteria | Actinobacteria | Actinomycetales_2 | Pseudonocardiaceae_1 | Thermodesulfovibrio | 0.000 | 0.000 | 0.054 | 0.000 | 0.000 | 0.000 |
| Actinobacteria | Actinobacteria | Pseudonocardiales_1 | Pseudonocardiaceae_1 | Prauserella | 0.000 | 0.000 | 0.004 | 0.000 | 0.000 | 0.000 |
| Actinobacteria | Actinobacteria | Actinomycetales_2 | Pseudonocardiaceae_2 | Kiloniella | 0.016 | 0.000 | 0.000 | 0.000 | 0.000 | 0.000 |
| Actinobacteria | Actinobacteria | Actinomycetales_2 | Pseudonocardiaceae_2 | Pseudorhodoferax | 0.008 | 0.031 | 0.002 | 0.000 | 0.000 | 0.012 |
| Actinobacteria | Actinobacteria | Actinomycetales_2 | Pseudonocardiaceae_3 | Streptomyces | 0.114 | 0.006 | 0.026 | 0.246 | 1.161 | 0.902 |
| Actinobacteria | Actinobacteria | Actinomycetales_2 | Pseudonocardiaceae_4 | Kutzneria_2 | 0.000 | 0.000 | 0.004 | 0.000 | 0.000 | 0.000 |
| Actinobacteria | Actinobacteria | Actinomycetales_2 | Pseudonocardiaceae_4 | Kytococcus | 0.000 | 0.000 | 0.004 | 0.000 | 0.000 | 0.000 |
| Actinobacteria | Actinobacteria | Micrococcales_2 | Rarobacteraceae | Rarobacter | 0.000 | 0.000 | 0.006 | 0.000 | 0.000 | 0.000 |
| Actinobacteria | Actinobacteria | Rubrobacterales | Rubrobacteriaceae | Rubrobacter | 0.008 | 0.000 | 0.050 | 0.000 | 0.005 | 0.000 |
| Actinobacteria | Actinobacteria | Rubrobacterales | Rubrobacteriaceae | Rufibacter | 0.000 | 0.000 | 0.024 | 0.000 | 0.000 | 0.000 |
| Actinobacteria | Actinobacteria | Actinomycetales_2 | Segniliparaceae | Selenomonas_1 | 0.000 | 0.000 | 0.018 | 0.000 | 0.000 | 0.000 |
| Actinobacteria | Actinobacteria | Corynebacteriales | Segniliparaceae | Segniliparus | 0.000 | 0.000 | 0.006 | 0.000 | 0.000 | 0.000 |
| Actinobacteria | Actinobacteria | Solirubrobacterales | Solirubrobacteriaceae | Solobacterium | 0.008 | 0.000 | 0.026 | 0.000 | 0.000 | 0.000 |
| Actinobacteria | Actinobacteria | Frankiales_2 | Sporichthyaceae | hgcI_clade | 0.000 | 0.000 | 0.036 | 0.000 | 0.000 | 0.000 |
| Actinobacteria | Actinobacteria | Actinomycetales_1 | Streptomycetaceae | Klebsiella | 0.163 | 1.084 | 0.621 | 2.330 | 1.505 | 2.058 |
| Actinobacteria | Actinobacteria | Actinomycetales_1 | Streptomycetaceae | Streptacidiphilus | 0.000 | 0.000 | 0.012 | 0.000 | 0.000 | 0.000 |
| Actinobacteria | Actinobacteria | Actinomycetales_1 | Streptomycetaceae | Streptomyces_10 | 0.000 | 0.000 | 0.012 | 0.000 | 0.000 | 0.006 |
| Actinobacteria | Actinobacteria | Actinomycetales_1 | Streptomycetaceae | Streptomyces_11 | 0.000 | 0.000 | 0.006 | 0.000 | 0.000 | 0.000 |
| Actinobacteria | Actinobacteria | Actinomycetales_1 | Streptomycetaceae | Streptomyces_2 | 0.000 | 0.000 | 0.008 | 0.000 | 0.000 | 0.000 |
| Actinobacteria | Actinobacteria | Actinomycetales_1 | Streptomycetaceae | Streptomyces_3 | 0.016 | 0.000 | 0.095 | 0.006 | 0.045 | 0.030 |
| Actinobacteria | Actinobacteria | Actinomycetales_1 | Streptomycetaceae | Streptomyces_4 | 0.000 | 0.000 | 0.101 | 0.012 | 0.030 | 0.006 |
| Actinobacteria | Actinobacteria | Actinomycetales_1 | Streptomycetaceae | Streptomyces_5 | 0.000 | 0.000 | 0.054 | 0.006 | 0.010 | 0.024 |
| Actinobacteria | Actinobacteria | Actinomycetales_1 | Streptomycetaceae | Streptomyces_6 | 0.000 | 0.000 | 0.008 | 0.000 | 0.000 | 0.000 |
| Actinobacteria | Actinobacteria | Actinomycetales_1 | Streptomycetaceae | Streptomyces_7 | 0.000 | 0.000 | 0.010 | 0.000 | 0.000 | 0.000 |
| Actinobacteria | Actinobacteria | Actinomycetales_1 | Streptomycetaceae | Streptomyces_8 | 0.000 | 0.000 | 0.012 | 0.000 | 0.000 | 0.000 |
| Actinobacteria | Actinobacteria | Actinomycetales_1 | Streptomycetaceae | Streptomyces_9 | 0.000 | 0.000 | 0.010 | 0.000 | 0.005 | 0.012 |
| Actinobacteria | Actinobacteria | Actinomycetales_1 | Streptomycetaceae | Subcluster_Ia | 0.033 | 0.012 | 0.042 | 0.000 | 0.000 | 0.000 |
| Actinobacteria | Actinobacteria | Actinomycetales_1 | Streptomycetaceae | Ureibacillus | 0.000 | 0.000 | 0.010 | 0.000 | 0.000 | 0.000 |

**Table S9 continued.**

| | | | | | | | | | | |
|---|---|---|---|---|---|---|---|---|---|---|
| Actinobacteria | Actinobacteria | Actinomycetales_1 | Streptomycetaceae | vadinBC27_wastewater-sludge_group | 0.016 | 0.000 | 0.042 | 0.006 | 0.000 | 0.000 |
| Actinobacteria | Actinobacteria | Actinomycetales | Streptosporangiaceae | Thermosinus | 0.000 | 0.000 | 0.026 | 0.000 | 0.000 | 0.000 |
| Actinobacteria | Actinobacteria | Actinomycetales_4 | Streptosporangiaceae | Acrocarpaspora | 0.000 | 0.000 | 0.008 | 0.000 | 0.000 | 0.000 |
| Actinobacteria | Actinobacteria | Actinomycetales_4 | Streptosporangiaceae | Microbispora | 0.000 | 0.000 | 0.024 | 0.000 | 0.010 | 0.006 |
| Actinobacteria | Actinobacteria | Actinomycetales_4 | Streptosporangiaceae | Microcella | 0.000 | 0.000 | 0.004 | 0.000 | 0.000 | 0.000 |
| Actinobacteria | Actinobacteria | Actinomycetales_4 | Streptosporangiaceae | Planotetraspora | 0.000 | 0.000 | 0.008 | 0.000 | 0.000 | 0.000 |
| Actinobacteria | Actinobacteria | Actinomycetales_4 | Streptosporangiaceae | Pleomorphomonas | 0.000 | 0.000 | 0.010 | 0.000 | 0.000 | 0.000 |
| Actinobacteria | Actinobacteria | Actinomycetales_4 | Streptosporangiaceae | Streptosporangium | 0.000 | 0.000 | 0.024 | 0.000 | 0.000 | 0.006 |
| Actinobacteria | Actinobacteria | Actinomycetales_4 | Streptosporangiaceae | Subcluster_Ib | 0.024 | 0.031 | 0.038 | 0.000 | 0.000 | 0.000 |
| Actinobacteria | Actinobacteria | Streptosporangiales | Streptosporangiaceae | Microtetraspora | 0.000 | 0.000 | 0.006 | 0.000 | 0.000 | 0.000 |
| Actinobacteria | Actinobacteria | Streptosporangiales | Streptosporangiaceae | Nonomuraea_1 | 0.000 | 0.000 | 0.004 | 0.000 | 0.000 | 0.000 |
| Actinobacteria | Actinobacteria | Streptosporangiales | Streptosporangiaceae | Nonomuraea_11 | 0.000 | 0.000 | 0.008 | 0.000 | 0.000 | 0.000 |
| Actinobacteria | Actinobacteria | Streptosporangiales | Streptosporangiaceae | Nonomuraea_12 | 0.000 | 0.000 | 0.006 | 0.000 | 0.000 | 0.000 |
| Actinobacteria | Actinobacteria | Streptosporangiales | Streptosporangiaceae | Nonomuraea_15 | 0.000 | 0.000 | 0.008 | 0.000 | 0.000 | 0.000 |
| Actinobacteria | Actinobacteria | Streptosporangiales | Streptosporangiaceae | Nonomuraea_16 | 0.000 | 0.000 | 0.004 | 0.000 | 0.000 | 0.000 |
| Actinobacteria | Actinobacteria | Streptosporangiales | Streptosporangiaceae | Nonomuraea_2 | 0.000 | 0.000 | 0.004 | 0.000 | 0.000 | 0.000 |
| Actinobacteria | Actinobacteria | Streptosporangiales | Streptosporangiaceae | Nonomuraea_4 | 0.000 | 0.000 | 0.004 | 0.000 | 0.000 | 0.000 |
| Actinobacteria | Actinobacteria | Streptosporangiales | Streptosporangiaceae | Nonomuraea_5 | 0.000 | 0.000 | 0.004 | 0.000 | 0.000 | 0.000 |
| Actinobacteria | Actinobacteria | Streptosporangiales | Streptosporangiaceae | Nonomuraea_7 | 0.000 | 0.000 | 0.004 | 0.000 | 0.020 | 0.006 |
| Actinobacteria | Actinobacteria | Streptosporangiales | Streptosporangiaceae | Nonomuraea_9 | 0.000 | 0.000 | 0.004 | 0.000 | 0.000 | 0.000 |
| Actinobacteria | Actinobacteria | Streptosporangiales | Streptosporangiaceae | Nordella | 0.008 | 0.000 | 0.000 | 0.000 | 0.000 | 0.000 |
| Actinobacteria | Actinobacteria | Streptosporangiales | Streptosporangiaceae | Planobispora | 0.000 | 0.000 | 0.004 | 0.000 | 0.000 | 0.000 |
| Actinobacteria | Actinobacteria | Streptosporangiales | Streptosporangiaceae | Planomonospora_2 | 0.000 | 0.000 | 0.006 | 0.000 | 0.000 | 0.000 |
| Actinobacteria | Actinobacteria | Streptosporangiales | Streptosporangiaceae | Sphaerisporangium_2 | 0.000 | 0.000 | 0.006 | 0.000 | 0.000 | 0.000 |
| Actinobacteria | Actinobacteria | Streptosporangiales | Streptosporangiaceae | Sphaerisporangium_3 | 0.000 | 0.000 | 0.004 | 0.000 | 0.000 | 0.000 |
| Actinobacteria | Actinobacteria | Streptosporangiales | Streptosporangiaceae | Sphaerotilus | 0.000 | 0.000 | 0.012 | 0.000 | 0.000 | 0.000 |
| Actinobacteria | Actinobacteria | Actinomycetales | Termite_Cluster | Termite_cluster_3 | 0.496 | 0.283 | 0.058 | 0.174 | 0.030 | 0.042 |
| Actinobacteria | Actinobacteria | Actinomycetales | Termite_Cluster | Termite_cluster_I | 0.122 | 0.000 | 0.032 | 0.042 | 0.005 | 0.006 |
| Actinobacteria | Actinobacteria | Actinomycetales | Termite_cluster_1 | Candidatus_Ancillula_trichonymphae | 0.000 | 0.000 | 0.006 | 0.000 | 0.000 | 0.000 |
| Actinobacteria | Actinobacteria | Actinomycetales | Termite_cluster_1 | Subcluster_a | 0.000 | 0.000 | 0.020 | 0.000 | 0.000 | 0.000 |
| Actinobacteria | Actinobacteria | Actinomycetales | Termite_cluster_1 | Subcluster_b | 0.000 | 0.000 | 0.006 | 0.000 | 0.000 | 0.000 |
| Actinobacteria | Actinobacteria | Streptosporangiales | Thermomonasporaceae | Thermosediminibacter | 0.008 | 0.000 | 0.000 | 0.000 | 0.000 | 0.000 |
| Actinobacteria | Actinobacteria | Streptosporangiales | Thermomonosporaceae | Actinomadura | 0.008 | 0.000 | 0.000 | 0.000 | 0.000 | 0.000 |
| Actinobacteria | Actinobacteria | Actinomycetales_4 | Thermomonosporaceae_2 | Actinoallomurus | 0.000 | 0.000 | 0.008 | 0.000 | 0.000 | 0.000 |
| Actinobacteria | Actinobacteria | Actinomycetales_4 | Thermomonosporaceae_2 | Actinocorallia | 0.000 | 0.000 | 0.014 | 0.000 | 0.000 | 0.000 |
| Actinobacteria | Actinobacteria | Actinomycetales_4 | Thermomonosporaceae_2 | Actinomadura_1 | 0.000 | 0.000 | 0.044 | 0.000 | 0.000 | 0.000 |
| Actinobacteria | Actinobacteria | Actinomycetales_4 | Thermomonosporaceae_2 | Actinomadura_10 | 0.000 | 0.000 | 0.004 | 0.000 | 0.000 | 0.000 |
| Actinobacteria | Actinobacteria | Actinomycetales_4 | Thermomonosporaceae_2 | Actinomadura_11 | 0.000 | 0.000 | 0.004 | 0.000 | 0.000 | 0.000 |
| Actinobacteria | Actinobacteria | Actinomycetales_4 | Thermomonosporaceae_2 | Actinomadura_2 | 0.000 | 0.000 | 0.004 | 0.000 | 0.000 | 0.000 |
| Actinobacteria | Actinobacteria | Actinomycetales_4 | Thermomonosporaceae_2 | Actinomadura_5 | 0.000 | 0.000 | 0.014 | 0.000 | 0.000 | 0.000 |

**Table S9 continued.**

| | | | | | | | | | | |
|---|---|---|---|---|---|---|---|---|---|---|
| Actinobacteria | Actinobacteria | Actinomycetales_4 | Thermomonosporaceae_2 | Actinomadura_7 | 0.000 | 0.000 | 0.012 | 0.000 | 0.000 | 0.000 |
| Actinobacteria | Actinobacteria | Actinomycetales_4 | Thermomonosporaceae_2 | Actinomadura_8 | 0.000 | 0.000 | 0.008 | 0.000 | 0.000 | 0.000 |
| Actinobacteria | Actinobacteria | Corynebacteriales | Tsukumurellaceae | Turicibacter | 0.000 | 0.000 | 0.048 | 0.006 | 0.005 | 0.000 |
| Actinobacteria | Actinobacteria | Micrococcales | Unclassified | Tistrella | 0.000 | 0.006 | 0.004 | 0.000 | 0.000 | 0.000 |
| Actinobacteria | Actinobacteria | Micrococcales_1 | Unclassified | Demequina | 0.016 | 0.000 | 0.000 | 0.000 | 0.000 | 0.000 |
| Aquificae | Aquificae | Aquificales | Aquificaceae | Aquifex | 0.000 | 0.000 | 0.006 | 0.000 | 0.000 | 0.000 |
| Aquificae | Aquificae | Aquificales | Aquificaceae | Hydrogenivirga | 0.000 | 0.000 | 0.008 | 0.000 | 0.000 | 0.000 |
| Aquificae | Aquificae | Aquificales | Aquificaceae | Hydrogenobacter | 0.000 | 0.000 | 0.032 | 0.000 | 0.000 | 0.000 |
| Aquificae | Aquificae | Aquificales | Aquificaceae | Hydrogenobaculum | 0.000 | 0.000 | 0.024 | 0.000 | 0.000 | 0.000 |
| Aquificae | Aquificae | Aquificales | Aquificaceae | Thermocrinis_1 | 0.000 | 0.000 | 0.054 | 0.000 | 0.000 | 0.000 |
| Aquificae | Aquificae | Aquificales | Desulfurobacteriaceae | Balnearium | 0.000 | 0.000 | 0.004 | 0.000 | 0.000 | 0.000 |
| Aquificae | Aquificae | Aquificales | Hydrogenothermaceae | Hydrogenothermus | 0.000 | 0.000 | 0.006 | 0.000 | 0.000 | 0.000 |
| Aquificae | Aquificae | Aquificales | Hydrogenothermaceae | Persephonella_1 | 0.000 | 0.000 | 0.012 | 0.000 | 0.000 | 0.000 |
| Aquificae | Aquificae | Aquificales | Hydrogenothermaceae | Sulfurihydrogenibium | 0.000 | 0.000 | 0.071 | 0.000 | 0.000 | 0.000 |
| Bacteroidetes | Flavobacteria | Flavobacteriales | Flavobacteriaceae_1 | Winogradskyella | 0.000 | 0.000 | 0.028 | 0.000 | 0.000 | 0.000 |
| Bacteroidetes | Bacteroidia | Bacteroidales | Bacteroidaceae | Bacteroides | 1.936 | 0.518 | 1.356 | 0.168 | 0.060 | 0.109 |
| Bacteroidetes | Bacteroidia | Bacteroidales | Bacteroidaceae | Meganema | 0.000 | 0.000 | 0.010 | 0.000 | 0.000 | 0.000 |
| Bacteroidetes | Flavobacteriia | Flavobacteriales | Blattabacteriaceae | Blattabacterium | 0.000 | 0.000 | 0.036 | 0.000 | 0.000 | 0.000 |
| Bacteroidetes | Flavobacteriia | Flavobacteriales | Blattabacteriaceae | Candidatus_Sulcia | 0.000 | 0.000 | 0.030 | 0.000 | 0.000 | 0.000 |
| Bacteroidetes | Chitinophagia | Chitinophagales | Chitinophagaceae | Flavisolibacter | 0.033 | 0.006 | 0.026 | 0.006 | 0.000 | 0.000 |
| Bacteroidetes | Chitinophagia | Chitinophagales | Chitinophagaceae | Flavitalea | 0.016 | 0.006 | 0.002 | 0.000 | 0.000 | 0.000 |
| Bacteroidetes | Chitinophagia | Chitinophagales | Chitinophagaceae | Hydrocarboniphaga | 0.000 | 0.000 | 0.000 | 0.006 | 0.000 | 0.000 |
| Bacteroidetes | Chitinophagia | Chitinophagales | Chitinophagaceae | Niastella | 0.016 | 0.000 | 0.000 | 0.006 | 0.000 | 0.000 |
| Bacteroidetes | Chitinophagia | Chitinophagales | Chitinophagaceae | Pandoraea | 0.122 | 0.000 | 0.020 | 0.186 | 0.224 | 0.042 |
| Bacteroidetes | Chitinophagia | Chitinophagales | Chitinophagaceae | Parasporobacterium | 0.000 | 0.006 | 0.000 | 0.000 | 0.000 | 0.000 |
| Bacteroidetes | Chitinophagia | Chitinophagales | Chitinophagaceae | Tannerella | 0.594 | 0.253 | 0.101 | 0.066 | 0.025 | 0.030 |
| Bacteroidetes | Chitinophagia | Chitinophagales | Chitinophagaceae | Terrimonas_2 | 0.000 | 0.000 | 0.004 | 0.000 | 0.000 | 0.000 |
| Bacteroidetes | Chitinophagia | Chitinophagales | Chitinophagaceae | Tessaracoccus | 0.000 | 0.000 | 0.018 | 0.000 | 0.000 | 0.000 |
| Bacteroidetes | Chitinophagia | Chitinophagales | Chitinophagaceae | Tetragenococcus | 0.000 | 0.000 | 0.012 | 0.000 | 0.000 | 0.000 |
| Bacteroidetes | Chitinophagia | Chitinophagales | Chitinophagaceae | Virgibacillus | 0.000 | 0.000 | 0.002 | 0.000 | 0.000 | 0.000 |
| Bacteroidetes | Sphingobacteriia | Sphingobacteriales_1 | Chitinophagaceae | Niastella_2 | 0.000 | 0.000 | 0.004 | 0.000 | 0.000 | 0.000 |
| Bacteroidetes | Sphingobacteria | Sphingobacteriales_2 | Chitinophagaceae | Filimonas | 0.000 | 0.000 | 0.010 | 0.000 | 0.000 | 0.000 |
| Bacteroidetes | Sphingobacteria | Sphingobacteriales_2 | Chitinophagaceae | Segetibacter | 0.000 | 0.000 | 0.010 | 0.000 | 0.000 | 0.000 |
| Bacteroidetes | Sphingobacteriia | Sphingobacteriales_2 | Chitinophagaceae | Chitinophaga | 0.016 | 0.000 | 0.040 | 0.018 | 0.000 | 0.000 |
| Bacteroidetes | Sphingobacteriia | Sphingobacteriales_2 | Chitinophagaceae | Nisaea | 0.000 | 0.000 | 0.004 | 0.000 | 0.000 | 0.000 |
| Bacteroidetes | Sphingobacteriia | Sphingobacteriales_2 | Chitinophagaceae | Uncultured_13 | 0.041 | 0.000 | 0.079 | 0.000 | 0.000 | 0.000 |
| Bacteroidetes | Sphingobacteria | Sphingobacteriales_4 | Chitinophagaceae | Balneola | 0.000 | 0.000 | 0.006 | 0.000 | 0.000 | 0.000 |
| Bacteroidetes | Bacteroidia | Bacteroidales | Cluster_V | Subdoligranulum | 0.000 | 0.000 | 0.167 | 0.000 | 0.000 | 0.000 |
| Bacteroidetes | Bacteroidia | Bacteroidales | Cluster_V | SubgroupII | 0.000 | 0.000 | 0.212 | 0.000 | 0.000 | 0.000 |
| Bacteroidetes | Flavobacteriia | Flavobacteriales | Cryomorphaceae | Oxalicibacterium | 0.000 | 0.000 | 0.004 | 0.006 | 0.000 | 0.000 |

**Table S9 continued.**

| | | | | | | | | | | |
|---|---|---|---|---|---|---|---|---|---|---|
| Bacteroidetes | Flavobacteria | Flavobacteriales | Cryomorphaceae_1 | NS10_marine_group | 0.000 | 0.000 | 0.004 | 0.000 | 0.000 | 0.000 |
| Bacteroidetes | Flavobacteria | Flavobacteriales | Cryomorphaceae_2 | Brumimimicrobium | 0.000 | 0.000 | 0.006 | 0.000 | 0.000 | 0.000 |
| Bacteroidetes | Flavobacteria | Flavobacteriales | Cryomorphaceae_2 | Lishizhenia | 0.000 | 0.000 | 0.004 | 0.000 | 0.000 | 0.000 |
| Bacteroidetes | Flavobacteria | Flavobacteriales | Cryomorphaceae_2 | NS7_marine_group | 0.000 | 0.000 | 0.004 | 0.000 | 0.000 | 0.000 |
| Bacteroidetes | Flavobacteriia | Flavobacteriales | Cryomorphaceae_2 | Crocinitomix | 0.000 | 0.000 | 0.006 | 0.000 | 0.000 | 0.000 |
| Bacteroidetes | Flavobacteriia | Flavobacteriales | Cryomorphaceae_2 | Fluviicola | 0.016 | 0.000 | 0.030 | 0.000 | 0.000 | 0.000 |
| Bacteroidetes | Sphingobacteria | Sphingobacteriales_3 | Cyclobacteriaceae | Algoriphagus_1 | 0.000 | 0.000 | 0.004 | 0.000 | 0.000 | 0.000 |
| Bacteroidetes | Sphingobacteria | Sphingobacteriales_3 | Cyclobacteriaceae | Algoriphagus_3 | 0.000 | 0.000 | 0.004 | 0.000 | 0.000 | 0.000 |
| Bacteroidetes | Sphingobacteria | Sphingobacteriales_3 | Cyclobacteriaceae | Algoriphagus_4 | 0.000 | 0.000 | 0.018 | 0.000 | 0.000 | 0.000 |
| Bacteroidetes | Sphingobacteria | Sphingobacteriales_3 | Cyclobacteriaceae | Algoriphagus_6 | 0.000 | 0.000 | 0.008 | 0.000 | 0.000 | 0.000 |
| Bacteroidetes | Sphingobacteria | Sphingobacteriales_3 | Cyclobacteriaceae | Algoriphagus_7 | 0.000 | 0.000 | 0.004 | 0.000 | 0.000 | 0.000 |
| Bacteroidetes | Sphingobacteria | Sphingobacteriales_3 | Cyclobacteriaceae | Aquiflexum | 0.000 | 0.000 | 0.006 | 0.000 | 0.000 | 0.000 |
| Bacteroidetes | Sphingobacteria | Sphingobacteriales_3 | Cyclobacteriaceae | Belliella | 0.000 | 0.000 | 0.006 | 0.000 | 0.000 | 0.000 |
| Bacteroidetes | Sphingobacteria | Sphingobacteriales_3 | Cyclobacteriaceae | Cyclobacterium | 0.000 | 0.000 | 0.010 | 0.000 | 0.000 | 0.000 |
| Bacteroidetes | Sphingobacteria | Sphingobacteriales_3 | Cyclobacteriaceae | Echinicola | 0.000 | 0.000 | 0.006 | 0.000 | 0.000 | 0.000 |
| Bacteroidetes | Cytophagia | Cytophagales | Cytophagaceae | Cytophaga | 0.000 | 0.000 | 0.006 | 0.000 | 0.000 | 0.000 |
| Bacteroidetes | Cytophagia | Cytophagales | Cytophagaceae | Lawsonia | 0.000 | 0.000 | 0.030 | 0.000 | 0.000 | 0.000 |
| Bacteroidetes | Sphingobacteria | Sphingobacteriales_3 | Cytophagaceae_1 | Arcicella_1 | 0.000 | 0.000 | 0.004 | 0.000 | 0.000 | 0.000 |
| Bacteroidetes | Sphingobacteria | Sphingobacteriales_3 | Cytophagaceae_1 | Arcicella_2 | 0.000 | 0.000 | 0.004 | 0.000 | 0.000 | 0.000 |
| Bacteroidetes | Sphingobacteria | Sphingobacteriales_3 | Cytophagaceae_1 | Emticicia | 0.000 | 0.000 | 0.010 | 0.000 | 0.000 | 0.000 |
| Bacteroidetes | Sphingobacteria | Sphingobacteriales_3 | Cytophagaceae_1 | Flectobacillus | 0.000 | 0.000 | 0.004 | 0.000 | 0.000 | 0.000 |
| Bacteroidetes | Sphingobacteria | Sphingobacteriales_3 | Cytophagaceae_1 | Flexibacter_1 | 0.277 | 0.006 | 0.046 | 0.006 | 0.000 | 0.006 |
| Bacteroidetes | Sphingobacteria | Sphingobacteriales_3 | Cytophagaceae_1 | Leadbetterella | 0.000 | 0.000 | 0.010 | 0.000 | 0.000 | 0.000 |
| Bacteroidetes | Sphingobacteria | Sphingobacteriales_3 | Cytophagaceae_1 | Runella | 0.000 | 0.000 | 0.010 | 0.000 | 0.000 | 0.000 |
| Bacteroidetes | Sphingobacteriia | Sphingobacteriales_3 | Cytophagaceae_1 | Dyadobacter | 0.024 | 0.031 | 0.030 | 0.000 | 0.000 | 0.054 |
| Bacteroidetes | Sphingobacteria | Sphingobacteriales_3 | Cytophagaceae_2 | Sporocytophaga | 0.000 | 0.000 | 0.004 | 0.000 | 0.000 | 0.000 |
| Bacteroidetes | Sphingobacteria | Sphingobacteriales_3 | Cytophagaceae_2 | Sporomusa | 0.033 | 0.006 | 0.022 | 0.000 | 0.000 | 0.000 |
| Bacteroidetes | Sphingobacteria | Sphingobacteriales_3 | Cytophagaceae_3 | Adhaeribacter | 0.049 | 0.012 | 0.012 | 0.000 | 0.000 | 0.000 |
| Bacteroidetes | Sphingobacteria | Sphingobacteriales_3 | Flammeovirgaceae_1 | Flexithrix | 0.000 | 0.000 | 0.008 | 0.000 | 0.000 | 0.000 |
| Bacteroidetes | Sphingobacteria | Sphingobacteriales_3 | Flammeovirgaceae_1 | Fulvivirga | 0.000 | 0.000 | 0.008 | 0.000 | 0.000 | 0.000 |
| Bacteroidetes | Sphingobacteria | Sphingobacteriales_3 | Flammeovirgaceae_1 | Persicobacter | 0.000 | 0.000 | 0.006 | 0.000 | 0.000 | 0.000 |
| Bacteroidetes | Sphingobacteria | Sphingobacteriales_3 | Flammeovirgaceae_1 | Reichenbachiella | 0.008 | 0.000 | 0.012 | 0.000 | 0.000 | 0.000 |
| Bacteroidetes | Sphingobacteria | Sphingobacteriales_3 | Flammeovirgaceae_2 | Candidatus_Amoebophilus | 0.000 | 0.000 | 0.006 | 0.000 | 0.000 | 0.000 |
| Bacteroidetes | Sphingobacteria | Sphingobacteriales_3 | Flammeovirgaceae_2 | Fabibacter | 0.000 | 0.000 | 0.004 | 0.000 | 0.000 | 0.000 |
| Bacteroidetes | Sphingobacteria | Sphingobacteriales_3 | Flammeovirgaceae_2 | Flammeovirga | 0.000 | 0.000 | 0.012 | 0.000 | 0.000 | 0.000 |
| Bacteroidetes | Sphingobacteria | Sphingobacteriales_3 | Flammeovirgaceae_2 | Marinoscillum | 0.000 | 0.000 | 0.008 | 0.000 | 0.000 | 0.000 |
| Bacteroidetes | Sphingobacteria | Sphingobacteriales_3 | Flammeovirgaceae_2 | Microscilla | 0.000 | 0.000 | 0.004 | 0.000 | 0.000 | 0.000 |
| Bacteroidetes | Sphingobacteria | Sphingobacteriales_3 | Flammeovirgaceae_2 | Perexilibacter | 0.000 | 0.000 | 0.006 | 0.000 | 0.000 | 0.000 |
| Bacteroidetes | Sphingobacteria | Sphingobacteriales_3 | Flammeovirgaceae_2 | Rapidithrix | 0.000 | 0.000 | 0.004 | 0.000 | 0.000 | 0.000 |
| Bacteroidetes | Sphingobacteriia | Sphingobacteriales_3 | Flammeovirgaceae_2 | Candidatus_Cardinium | 0.000 | 0.000 | 0.008 | 0.000 | 0.000 | 0.000 |

**Table S9 continued.**

| | | | | | | | | | | |
|---|---|---|---|---|---|---|---|---|---|---|
| Bacteroidetes | Flavobacteriia | Flavobacteriales | Flavobacteriaceae | Aquimarina | 0.000 | 0.000 | 0.012 | 0.000 | 0.000 | 0.000 |
| Bacteroidetes | Flavobacteriia | Flavobacteriales | Flavobacteriaceae | Arenibacter | 0.000 | 0.000 | 0.012 | 0.000 | 0.000 | 0.000 |
| Bacteroidetes | Flavobacteriia | Flavobacteriales | Flavobacteriaceae | Capnocytophaga | 0.000 | 0.000 | 0.038 | 0.000 | 0.000 | 0.000 |
| Bacteroidetes | Flavobacteriia | Flavobacteriales | Flavobacteriaceae | Costertonia | 0.000 | 0.000 | 0.004 | 0.000 | 0.000 | 0.000 |
| Bacteroidetes | Flavobacteriia | Flavobacteriales | Flavobacteriaceae | Croceibacter | 0.000 | 0.000 | 0.004 | 0.000 | 0.000 | 0.000 |
| Bacteroidetes | Flavobacteriia | Flavobacteriales | Flavobacteriaceae | Elizabethkingia | 0.000 | 0.000 | 0.006 | 0.000 | 0.000 | 0.000 |
| Bacteroidetes | Flavobacteriia | Flavobacteriales | Flavobacteriaceae | Eudoraea | 0.000 | 0.000 | 0.004 | 0.000 | 0.000 | 0.000 |
| Bacteroidetes | Flavobacteriia | Flavobacteriales | Flavobacteriaceae | Flavobacterium | 0.033 | 0.012 | 0.004 | 0.000 | 0.005 | 0.000 |
| Bacteroidetes | Flavobacteriia | Flavobacteriales | Flavobacteriaceae | Flavobacterium_2 | 0.000 | 0.000 | 0.036 | 0.000 | 0.000 | 0.000 |
| Bacteroidetes | Flavobacteriia | Flavobacteriales | Flavobacteriaceae | Kordiimonas | 0.000 | 0.000 | 0.004 | 0.000 | 0.000 | 0.000 |
| Bacteroidetes | Flavobacteriia | Flavobacteriales | Flavobacteriaceae | Luteibacter | 0.106 | 0.080 | 0.091 | 3.762 | 5.413 | 3.880 |
| Bacteroidetes | Flavobacteriia | Flavobacteriales | Flavobacteriaceae | Mesoplasma_3 | 0.000 | 0.000 | 0.006 | 0.000 | 0.000 | 0.000 |
| Bacteroidetes | Flavobacteriia | Flavobacteriales | Flavobacteriaceae | Moorella | 0.000 | 0.000 | 0.010 | 0.000 | 0.000 | 0.000 |
| Bacteroidetes | Flavobacteriia | Flavobacteriales | Flavobacteriaceae | Nonomuraea | 0.000 | 0.000 | 0.000 | 0.000 | 0.045 | 0.024 |
| Bacteroidetes | Flavobacteriia | Flavobacteriales | Flavobacteriaceae | Quadrisphaera | 0.000 | 0.000 | 0.010 | 0.000 | 0.000 | 0.000 |
| Bacteroidetes | Flavobacteriia | Flavobacteriales | Flavobacteriaceae | Tenacibaculum_3 | 0.000 | 0.000 | 0.010 | 0.000 | 0.000 | 0.000 |
| Bacteroidetes | Flavobacteriia | Flavobacteriales | Flavobacteriaceae | Tepidanaerobacter | 0.000 | 0.000 | 0.026 | 0.000 | 0.000 | 0.000 |
| Bacteroidetes | Flavobacteriia | Flavobacteriales | Flavobacteriaceae | Tepidibacter | 0.000 | 0.000 | 0.006 | 0.000 | 0.000 | 0.000 |
| Bacteroidetes | Flavobacteria | Flavobacteriales | Flavobacteriaceae_1 | Actibacter | 0.000 | 0.000 | 0.006 | 0.000 | 0.000 | 0.000 |
| Bacteroidetes | Flavobacteria | Flavobacteriales | Flavobacteriaceae_1 | Aequorivita | 0.000 | 0.000 | 0.010 | 0.000 | 0.000 | 0.000 |
| Bacteroidetes | Flavobacteria | Flavobacteriales | Flavobacteriaceae_1 | Algibacter_1 | 0.000 | 0.000 | 0.004 | 0.000 | 0.000 | 0.000 |
| Bacteroidetes | Flavobacteria | Flavobacteriales | Flavobacteriaceae_1 | Algibacter_2 | 0.000 | 0.000 | 0.004 | 0.000 | 0.000 | 0.000 |
| Bacteroidetes | Flavobacteria | Flavobacteriales | Flavobacteriaceae_1 | Bizionia_1 | 0.000 | 0.000 | 0.008 | 0.000 | 0.000 | 0.000 |
| Bacteroidetes | Flavobacteria | Flavobacteriales | Flavobacteriaceae_1 | Bizionia_2 | 0.000 | 0.000 | 0.010 | 0.000 | 0.000 | 0.000 |
| Bacteroidetes | Flavobacteria | Flavobacteriales | Flavobacteriaceae_1 | Cellulophaga_1 | 0.000 | 0.000 | 0.008 | 0.000 | 0.000 | 0.000 |
| Bacteroidetes | Flavobacteria | Flavobacteriales | Flavobacteriaceae_1 | Cellulophaga_2 | 0.000 | 0.000 | 0.006 | 0.000 | 0.000 | 0.000 |
| Bacteroidetes | Flavobacteria | Flavobacteriales | Flavobacteriaceae_1 | Croceitalea_1 | 0.000 | 0.000 | 0.006 | 0.000 | 0.000 | 0.000 |
| Bacteroidetes | Flavobacteria | Flavobacteriales | Flavobacteriaceae_1 | Flavobacterium_1 | 0.000 | 0.000 | 0.329 | 0.006 | 0.000 | 0.000 |
| Bacteroidetes | Flavobacteria | Flavobacteriales | Flavobacteriaceae_1 | Formosa | 0.000 | 0.000 | 0.010 | 0.000 | 0.000 | 0.000 |
| Bacteroidetes | Flavobacteria | Flavobacteriales | Flavobacteriaceae_1 | Gillisia | 0.000 | 0.000 | 0.018 | 0.000 | 0.000 | 0.000 |
| Bacteroidetes | Flavobacteria | Flavobacteriales | Flavobacteriaceae_1 | Gilvibacter | 0.000 | 0.000 | 0.004 | 0.000 | 0.000 | 0.000 |
| Bacteroidetes | Flavobacteria | Flavobacteriales | Flavobacteriaceae_1 | Gramella | 0.000 | 0.000 | 0.008 | 0.000 | 0.000 | 0.000 |
| Bacteroidetes | Flavobacteria | Flavobacteriales | Flavobacteriaceae_1 | Joostella | 0.000 | 0.000 | 0.004 | 0.000 | 0.000 | 0.000 |
| Bacteroidetes | Flavobacteria | Flavobacteriales | Flavobacteriaceae_1 | Krokonibacter_sp_a | 0.000 | 0.000 | 0.004 | 0.000 | 0.000 | 0.000 |
| Bacteroidetes | Flavobacteria | Flavobacteriales | Flavobacteriaceae_1 | Lacinutrix_1 | 0.000 | 0.000 | 0.006 | 0.000 | 0.000 | 0.000 |
| Bacteroidetes | Flavobacteria | Flavobacteriales | Flavobacteriaceae_1 | Leeuwenhoekiella | 0.000 | 0.000 | 0.010 | 0.000 | 0.000 | 0.000 |
| Bacteroidetes | Flavobacteria | Flavobacteriales | Flavobacteriaceae_1 | Lutibacter | 0.000 | 0.000 | 0.006 | 0.000 | 0.000 | 0.000 |
| Bacteroidetes | Flavobacteria | Flavobacteriales | Flavobacteriaceae_1 | Maribacter | 0.000 | 0.000 | 0.022 | 0.000 | 0.000 | 0.000 |
| Bacteroidetes | Flavobacteria | Flavobacteriales | Flavobacteriaceae_1 | Marixanthomonas | 0.000 | 0.000 | 0.004 | 0.000 | 0.000 | 0.000 |
| Bacteroidetes | Flavobacteria | Flavobacteriales | Flavobacteriaceae_1 | Mesoflavibacter | 0.000 | 0.000 | 0.004 | 0.000 | 0.000 | 0.000 |

**Table S9 continued.**

| | | | | | | | | | | |
|---|---|---|---|---|---|---|---|---|---|---|
| Bacteroidetes | Flavobacteria | Flavobacteriales | Flavobacteriaceae_1 | NS2b_marine_group | 0.000 | 0.000 | 0.004 | 0.000 | 0.000 | 0.000 |
| Bacteroidetes | Flavobacteria | Flavobacteriales | Flavobacteriaceae_1 | NS3a_marine_group | 0.000 | 0.000 | 0.004 | 0.000 | 0.000 | 0.000 |
| Bacteroidetes | Flavobacteria | Flavobacteriales | Flavobacteriaceae_1 | NS4_marine_group | 0.000 | 0.000 | 0.008 | 0.000 | 0.000 | 0.000 |
| Bacteroidetes | Flavobacteria | Flavobacteriales | Flavobacteriaceae_1 | NS5_marine_group | 0.000 | 0.000 | 0.018 | 0.000 | 0.000 | 0.000 |
| Bacteroidetes | Flavobacteria | Flavobacteriales | Flavobacteriaceae_1 | Persicivirga | 0.000 | 0.000 | 0.004 | 0.000 | 0.000 | 0.000 |
| Bacteroidetes | Flavobacteria | Flavobacteriales | Flavobacteriaceae_1 | Polaribacter | 0.000 | 0.000 | 0.030 | 0.000 | 0.000 | 0.000 |
| Bacteroidetes | Flavobacteria | Flavobacteriales | Flavobacteriaceae_1 | Psychroflexus | 0.000 | 0.000 | 0.014 | 0.000 | 0.000 | 0.000 |
| Bacteroidetes | Flavobacteria | Flavobacteriales | Flavobacteriaceae_1 | Psychroserpens_2 | 0.000 | 0.000 | 0.006 | 0.000 | 0.000 | 0.000 |
| Bacteroidetes | Flavobacteria | Flavobacteriales | Flavobacteriaceae_1 | Salinimicrobium | 0.000 | 0.000 | 0.016 | 0.000 | 0.000 | 0.000 |
| Bacteroidetes | Flavobacteria | Flavobacteriales | Flavobacteriaceae_1 | Sediminibacter | 0.000 | 0.000 | 0.008 | 0.000 | 0.000 | 0.000 |
| Bacteroidetes | Flavobacteria | Flavobacteriales | Flavobacteriaceae_1 | Sediminicola | 0.000 | 0.000 | 0.004 | 0.000 | 0.000 | 0.000 |
| Bacteroidetes | Flavobacteria | Flavobacteriales | Flavobacteriaceae_1 | Stenothermobacter | 0.000 | 0.000 | 0.004 | 0.000 | 0.000 | 0.000 |
| Bacteroidetes | Flavobacteria | Flavobacteriales | Flavobacteriaceae_1 | Subsaxibacter | 0.000 | 0.000 | 0.006 | 0.000 | 0.000 | 0.000 |
| Bacteroidetes | Flavobacteria | Flavobacteriales | Flavobacteriaceae_1 | Sufflavibacter | 0.000 | 0.000 | 0.010 | 0.000 | 0.000 | 0.000 |
| Bacteroidetes | Flavobacteria | Flavobacteriales | Flavobacteriaceae_1 | Tamlana_2 | 0.000 | 0.000 | 0.004 | 0.000 | 0.000 | 0.000 |
| Bacteroidetes | Flavobacteria | Flavobacteriales | Flavobacteriaceae_1 | Ulvibacter | 0.000 | 0.000 | 0.010 | 0.000 | 0.000 | 0.000 |
| Bacteroidetes | Flavobacteria | Flavobacteriales | Flavobacteriaceae_1 | Vitellibacter | 0.000 | 0.000 | 0.006 | 0.000 | 0.000 | 0.000 |
| Bacteroidetes | Flavobacteria | Flavobacteriales | Flavobacteriaceae_1 | Yeosuana | 0.000 | 0.000 | 0.004 | 0.000 | 0.000 | 0.000 |
| Bacteroidetes | Flavobacteria | Flavobacteriales | Flavobacteriaceae_1 | Zeaxanthinibacter | 0.000 | 0.000 | 0.004 | 0.000 | 0.000 | 0.000 |
| Bacteroidetes | Flavobacteria | Flavobacteriales | Flavobacteriaceae_1 | Zunongwangia | 0.000 | 0.000 | 0.004 | 0.000 | 0.000 | 0.000 |
| Bacteroidetes | Fusobacteriia | Flavobacteriales | Flavobacteriaceae_1 | Gaetbulibacter_2 | 0.000 | 0.000 | 0.008 | 0.000 | 0.000 | 0.000 |
| Bacteroidetes | Flavobacteria | Flavobacteriales | Flavobacteriaceae_2 | Bergeyella | 0.000 | 0.000 | 0.006 | 0.000 | 0.000 | 0.000 |
| Bacteroidetes | Flavobacteria | Flavobacteriales | Flavobacteriaceae_2 | Chryseobacterium_12 | 0.000 | 0.000 | 0.014 | 0.000 | 0.000 | 0.000 |
| Bacteroidetes | Flavobacteria | Flavobacteriales | Flavobacteriaceae_2 | Chryseobacterium_2 | 0.000 | 0.000 | 0.028 | 0.000 | 0.000 | 0.000 |
| Bacteroidetes | Flavobacteria | Flavobacteriales | Flavobacteriaceae_2 | Chryseobacterium_3 | 0.000 | 0.000 | 0.004 | 0.000 | 0.000 | 0.000 |
| Bacteroidetes | Flavobacteria | Flavobacteriales | Flavobacteriaceae_2 | Chryseobacterium_6 | 0.000 | 0.000 | 0.008 | 0.000 | 0.000 | 0.000 |
| Bacteroidetes | Flavobacteria | Flavobacteriales | Flavobacteriaceae_2 | Chryseobacterium_7 | 0.000 | 0.000 | 0.004 | 0.000 | 0.000 | 0.000 |
| Bacteroidetes | Flavobacteria | Flavobacteriales | Flavobacteriaceae_2 | Chryseobacterium_9 | 0.000 | 0.000 | 0.004 | 0.000 | 0.000 | 0.000 |
| Bacteroidetes | Flavobacteria | Flavobacteriales | Flavobacteriaceae_2 | Cloacibacterium | 0.000 | 0.000 | 0.026 | 0.000 | 0.000 | 0.000 |
| Bacteroidetes | Flavobacteria | Flavobacteriales | Flavobacteriaceae_2 | Planobacterium | 0.000 | 0.000 | 0.004 | 0.000 | 0.000 | 0.000 |
| Bacteroidetes | Flavobacteria | Flavobacteriales | Flavobacteriaceae_2 | Riemerella | 0.000 | 0.000 | 0.006 | 0.000 | 0.000 | 0.000 |
| Bacteroidetes | Bacteroidia | Bacteroidales | Gut_Group_A | Nubsella | 0.008 | 0.012 | 0.008 | 0.000 | 0.000 | 0.000 |
| Bacteroidetes | Bacteroidia | Bacteroidales | Gut_Group_A | Rs-H88_termite_group | 0.000 | 0.000 | 0.006 | 0.000 | 0.000 | 0.000 |
| Bacteroidetes | Cytophagia | Cytophagales | Hymenobacteraceae | Hyphomicrobium | 0.041 | 0.000 | 0.000 | 0.000 | 0.000 | 0.000 |
| Bacteroidetes | Cytophagia | Cytophagales | Hymenobacteraceae | Pontibacter_1 | 0.000 | 0.000 | 0.008 | 0.000 | 0.000 | 0.000 |
| Bacteroidetes | Cytophagia | Cytophagales | Hymenobacteraceae | Pontibacter_2 | 0.000 | 0.000 | 0.004 | 0.000 | 0.000 | 0.000 |
| Bacteroidetes | Cytophagia | Cytophagales | Hymenobacteraceae | Porphyrobacter | 0.008 | 0.000 | 0.000 | 0.000 | 0.000 | 0.000 |
| Bacteroidetes | Cytophagia | Cytophagales | Hymenobacteraceae | Ruminococcus | 0.000 | 0.000 | 0.002 | 0.000 | 0.000 | 0.000 |
| Bacteroidetes | Bacteroidia | Bacteroidales | Marinilabiaceae | Alkaliflexus | 0.008 | 0.000 | 0.018 | 0.000 | 0.000 | 0.000 |
| Bacteroidetes | Bacteroidia | Bacteroidales | Marinilabiaceae | Anaerophaga | 0.000 | 0.000 | 0.004 | 0.000 | 0.000 | 0.000 |

**Table S9 continued.**

| | | | | | | | | | | |
|---|---|---|---|---|---|---|---|---|---|---|
| Bacteroidetes | Bacteroidia | Marinilabiliales | Marinilabiliaceae | marine_benthic_group | 0.008 | 0.000 | 0.032 | 0.000 | 0.000 | 0.000 |
| Bacteroidetes | Bacteroidia | Marinilabiliales | Marinilabiliaceae | Naxibacter | 0.024 | 0.000 | 0.060 | 0.006 | 0.010 | 0.018 |
| Bacteroidetes | Bacteroidia | Bacteroidales | Odoribacteraceae | Culturomica | 0.016 | 0.000 | 0.000 | 0.000 | 0.000 | 0.000 |
| Bacteroidetes | Cytophagia | Cytophagales | Persicobacteraceae | Fulvitalea | 0.008 | 0.000 | 0.000 | 0.000 | 0.000 | 0.000 |
| Bacteroidetes | Bacteroidia | Bacteroidales | Porphyromonadaceae | Cockroach_cluster | 0.073 | 0.006 | 0.075 | 0.018 | 0.000 | 0.000 |
| Bacteroidetes | Bacteroidia | Bacteroidales | Porphyromonadaceae | Microbacterium | 0.008 | 0.006 | 0.226 | 0.000 | 0.010 | 0.000 |
| Bacteroidetes | Bacteroidia | Bacteroidales | Porphyromonadaceae_1 | Dysgonomonas | 0.708 | 0.111 | 0.171 | 0.060 | 0.005 | 0.024 |
| Bacteroidetes | Bacteroidia | Bacteroidales | Porphyromonadaceae_1 | Tardiphaga | 0.008 | 0.000 | 0.000 | 0.000 | 0.000 | 0.000 |
| Bacteroidetes | Bacteroidia | Bacteroidales | Porphyromonadaceae_2 | Cubitermes_Cluster_A | 0.000 | 0.000 | 0.004 | 0.000 | 0.000 | 0.000 |
| Bacteroidetes | Bacteroidia | Bacteroidales | Porphyromonadaceae_2 | Panacagrimonas | 0.081 | 0.000 | 0.000 | 0.000 | 0.000 | 0.000 |
| Bacteroidetes | Bacteroidia | Bacteroidales | Porphyromonadaceae_2 | Termite_cluster_II | 0.854 | 0.240 | 0.042 | 0.246 | 0.025 | 0.018 |
| Bacteroidetes | Bacteroidia | Bacteroidales | Porphyromonadaceae_3 | Cluster_IV | 0.049 | 0.006 | 0.077 | 0.006 | 0.000 | 0.000 |
| Bacteroidetes | Bacteroidia | Bacteroidales | Porphyromonadaceae_3 | Paraburkholderia | 0.049 | 0.006 | 0.022 | 0.329 | 0.229 | 0.315 |
| Bacteroidetes | Bacteroidia | Bacteroidales | Porphyromonadaceae_7 | Barnesiella | 0.008 | 0.000 | 0.123 | 0.000 | 0.000 | 0.000 |
| Bacteroidetes | Bacteroidia | Bacteroidales | Porphyromonadaceae_Cluster_V | Candidatus_Armantifilum | 0.089 | 0.018 | 0.085 | 0.000 | 0.000 | 0.000 |
| Bacteroidetes | Bacteroidia | Bacteroidales | Porphyromonadaceae_Cluster_V | Candidatus_Azobacteroides | 0.000 | 0.000 | 0.030 | 0.000 | 0.000 | 0.000 |
| Bacteroidetes | Bacteroidia | Bacteroidales | Porphyromonadaceae_Cluster_V | Candidatus_Symbiothrix | 0.130 | 0.018 | 0.081 | 0.000 | 0.000 | 0.000 |
| Bacteroidetes | Bacteroidia | Bacteroidales | Porphyromonadaceae_Gut_group | Butyricimonas | 0.000 | 0.000 | 0.028 | 0.000 | 0.000 | 0.000 |
| Bacteroidetes | Bacteroidia | Bacteroidales | Porphyromonadaceae_Gut_group | Environmental_cluster | 0.000 | 0.000 | 0.004 | 0.000 | 0.000 | 0.000 |
| Bacteroidetes | Bacteroidia | Bacteroidales | Porphyromonadaceae_Gut_group | ML635J_40_aquatic_group | 0.000 | 0.000 | 0.006 | 0.000 | 0.000 | 0.000 |
| Bacteroidetes | Bacteroidia | Bacteroidales | Prevotellaceae | Prevotella_2 | 0.000 | 0.006 | 0.119 | 0.000 | 0.000 | 0.000 |
| Bacteroidetes | Bacteroidia | Bacteroidales | Prevotellaceae | Prolixibacter | 0.000 | 0.000 | 0.018 | 0.000 | 0.000 | 0.000 |
| Bacteroidetes | Bacteroidia | Marinilabiliales | Prolixibacteraceae | Mesonia | 0.000 | 0.000 | 0.010 | 0.000 | 0.000 | 0.000 |
| Bacteroidetes | Bacteroidia | Marinilabiliales | Prolixibacteraceae | Promicromonospora | 0.000 | 0.006 | 0.016 | 0.000 | 0.005 | 0.006 |
| Bacteroidetes | Sphingobacteria | Sphingobacteriales_4 | Rhodothermaceae | Rhodothermus | 0.000 | 0.000 | 0.004 | 0.000 | 0.000 | 0.000 |
| Bacteroidetes | Sphingobacteria | Sphingobacteriales_4 | Rhodothermaceae | Salinibacter | 0.000 | 0.000 | 0.010 | 0.000 | 0.000 | 0.000 |
| Bacteroidetes | Sphingobacteria | Sphingobacteriales_4 | Rhodothermaceae | Salisaeta | 0.000 | 0.000 | 0.004 | 0.000 | 0.000 | 0.000 |
| Bacteroidetes | Bacteroidia | Bacteroidales | Rikenellaceae | Alistipes | 0.016 | 0.000 | 0.000 | 0.000 | 0.000 | 0.000 |
| Bacteroidetes | Bacteroidia | Bacteroidales | Rikenellaceae | Alistipes_I | 0.073 | 0.012 | 0.173 | 0.018 | 0.005 | 0.000 |
| Bacteroidetes | Bacteroidia | Bacteroidales | Rikenellaceae | Alistipes_II | 4.856 | 1.386 | 0.121 | 0.545 | 0.120 | 0.157 |
| Bacteroidetes | Bacteroidia | Bacteroidales | Rikenellaceae | Alistipes_III | 0.635 | 0.055 | 0.028 | 0.018 | 0.005 | 0.006 |
| Bacteroidetes | Bacteroidia | Bacteroidales | Rikenellaceae | BCf9-17_termite_group | 1.001 | 0.166 | 0.040 | 0.072 | 0.005 | 0.006 |
| Bacteroidetes | Bacteroidia | Bacteroidales | Rikenellaceae | dgA-11_gut_group | 0.000 | 0.000 | 0.026 | 0.000 | 0.000 | 0.000 |
| Bacteroidetes | Bacteroidia | Bacteroidales | Rikenellaceae | gut_cluster_c | 0.081 | 0.025 | 0.016 | 0.036 | 0.000 | 0.006 |
| Bacteroidetes | Bacteroidia | Bacteroidales | Rikenellaceae | Gut_digester_cluster | 0.000 | 0.000 | 0.028 | 0.000 | 0.000 | 0.000 |
| Bacteroidetes | Bacteroidia | Bacteroidales | Rikenellaceae | M2PT2-76_termite_group | 0.016 | 0.000 | 0.004 | 0.000 | 0.000 | 0.000 |
| Bacteroidetes | Bacteroidia | Bacteroidales | Rikenellaceae | Paenibacillus | 0.138 | 0.049 | 0.046 | 1.276 | 3.265 | 7.529 |
| Bacteroidetes | Bacteroidia | Bacteroidales | Rikenellaceae | Reyranella | 0.008 | 0.000 | 0.000 | 0.000 | 0.000 | 0.000 |
| Bacteroidetes | Bacteroidia | Bacteroidales | Rikenellaceae | Rikenella_3 | 0.000 | 0.000 | 0.004 | 0.000 | 0.000 | 0.000 |
| Bacteroidetes | Bacteroidia | Bacteroidales | Rikenellaceae | Robiginitalea | 0.000 | 0.000 | 0.006 | 0.000 | 0.000 | 0.000 |

**Table S9 continued.**

| | | | | | | | | | | |
|---|---|---|---|---|---|---|---|---|---|---|
| Bacteroidetes | Bacteroidia | Bacteroidales | Rikenellaceae | Rs-M59_termite_group | 0.000 | 0.000 | 0.018 | 0.000 | 0.000 | 0.000 |
| Bacteroidetes | Bacteroidia | Bacteroidales | Rikenellaceae | Tissierella_3 | 0.000 | 0.000 | 0.028 | 0.000 | 0.000 | 0.000 |
| Bacteroidetes | Bacteroidia | Bacteroidales | Rikenellaceae | U29-B03 | 0.000 | 0.000 | 0.004 | 0.000 | 0.000 | 0.000 |
| Bacteroidetes | Bacteroidia | Bacteroidales | Rs-E47_termite_group | Gut_cluster | 0.024 | 0.339 | 0.069 | 1.036 | 0.379 | 1.035 |
| Bacteroidetes | Bacteroidia | Bacteroidales | Rs-E47_termite_group | Hirschia | 0.024 | 0.000 | 0.006 | 0.018 | 0.000 | 0.000 |
| Bacteroidetes | Bacteroidia | Bacteroidales | S24_7 | Cluster_I | 0.008 | 0.006 | 1.528 | 0.000 | 0.000 | 0.006 |
| Bacteroidetes | Bacteroidia | Bacteroidales | S24_7 | Cluster_II | 0.016 | 0.006 | 0.097 | 0.006 | 0.005 | 0.000 |
| Bacteroidetes | Sphingobacteria | Sphingobacteriales_2 | Saprospiraceae | Aureispira | 0.000 | 0.000 | 0.004 | 0.000 | 0.000 | 0.000 |
| Bacteroidetes | Sphingobacteria | Sphingobacteriales_2 | Saprospiraceae | Haliscomenobacter | 0.000 | 0.000 | 0.004 | 0.000 | 0.000 | 0.000 |
| Bacteroidetes | Sphingobacteria | Sphingobacteriales | Sphingobacteriaceae | Olivibacter_1 | 0.000 | 0.000 | 0.004 | 0.000 | 0.000 | 0.000 |
| Bacteroidetes | Sphingobacteria | Sphingobacteriales | Sphingobacteriaceae | Olivibacter_2 | 0.000 | 0.000 | 0.008 | 0.000 | 0.000 | 0.000 |
| Bacteroidetes | Sphingobacteriia | Sphingobacteriales | Sphingobacteriaceae | Paraprevotella | 0.000 | 0.000 | 0.032 | 0.000 | 0.000 | 0.000 |
| Bacteroidetes | Sphingobacteriia | Sphingobacteriales | Sphingobacteriaceae | Sorangium | 0.000 | 0.000 | 0.002 | 0.000 | 0.000 | 0.000 |
| Bacteroidetes | Sphingobacteriia | Sphingobacteriales | Sphingobacteriaceae | Sphingobacterium_2 | 0.000 | 0.000 | 0.022 | 0.000 | 0.000 | 0.000 |
| Bacteroidetes | Sphingobacteriia | Sphingobacteriales | Sphingobacteriaceae | Sphingobacterium_4 | 0.000 | 0.000 | 0.008 | 0.000 | 0.000 | 0.000 |
| Bacteroidetes | Sphingobacteriia | Sphingobacteriales | Sphingobacteriaceae | Sphingobium | 0.033 | 0.099 | 0.020 | 0.006 | 0.000 | 0.151 |
| Bacteroidetes | Sphingobacteriia | Sphingobacteriales | Sphingobacteriaceae | Sphingobium_1 | 0.024 | 0.018 | 0.115 | 0.012 | 0.000 | 0.012 |
| Bacteroidetes | Sphingobacteria | Sphingobacteriales_1 | Sphingobacteriaceae | Mucilaginibacter_2 | 0.000 | 0.000 | 0.006 | 0.000 | 0.000 | 0.000 |
| Bacteroidetes | Sphingobacteria | Sphingobacteriales_1 | Sphingobacteriaceae | Sphingobacterium_3 | 0.000 | 0.000 | 0.030 | 0.000 | 0.000 | 0.000 |
| Bacteroidetes | Sphingobacteriia | Sphingobacteriales_1 | Sphingobacteriaceae | Mucilaginibacter_1 | 0.000 | 0.012 | 0.036 | 0.006 | 0.000 | 0.000 |
| Bacteroidetes | Sphingobacteriia | Sphingobacteriales_1 | Sphingobacteriaceae | Mucispirillum | 0.399 | 0.129 | 0.032 | 0.174 | 0.005 | 0.006 |
| Bacteroidetes | Sphingobacteriia | Sphingobacteriales_1 | Sphingobacteriaceae | Pedobacter_1 | 0.000 | 0.000 | 0.016 | 0.000 | 0.000 | 0.000 |
| Bacteroidetes | Sphingobacteriia | Sphingobacteriales_1 | Sphingobacteriaceae | Pedobacter_2 | 0.008 | 0.000 | 0.050 | 0.000 | 0.000 | 0.000 |
| Bacteroidetes | Sphingobacteriia | Sphingobacteriales_1 | Sphingobacteriaceae | Pedobacter_3 | 0.000 | 0.000 | 0.006 | 0.000 | 0.000 | 0.000 |
| Bacteroidetes | Sphingobacteriia | Sphingobacteriales_1 | Sphingobacteriaceae | Pedobacter_4 | 0.000 | 0.000 | 0.006 | 0.000 | 0.000 | 0.000 |
| Bacteroidetes | Sphingobacteriia | Sphingobacteriales_1 | Sphingobacteriaceae | Pedobacter_5 | 0.000 | 0.000 | 0.008 | 0.000 | 0.000 | 0.000 |
| Bacteroidetes | Sphingobacteriia | Sphingobacteriales_1 | Sphingobacteriaceae | Pedobacter_6 | 0.000 | 0.000 | 0.012 | 0.000 | 0.000 | 0.000 |
| Bacteroidetes | Sphingobacteriia | Sphingobacteriales_1 | Sphingobacteriaceae | Pedomicrobium | 0.016 | 0.000 | 0.024 | 0.000 | 0.000 | 0.006 |
| Bacteroidetes | Cytophagia | Cytophagales | Unclassified | Chryseolinea | 0.008 | 0.006 | 0.000 | 0.000 | 0.000 | 0.000 |
| Balneolaeota | Balneolia | Balneolales | Balneolaceae | Aliihoeflea | 0.000 | 0.006 | 0.000 | 0.000 | 0.000 | 0.000 |
| Caldiserica | Caldisericia | Caldisericales | Caldisericaceae | Caldisericum | 0.000 | 0.000 | 0.026 | 0.000 | 0.000 | 0.000 |
| Candidate_division_TG3 | TG3_Subphylum_1 | TG3_Subphylum_1 | Termite_Cluster | Termite_cluster_III | 0.098 | 0.037 | 0.012 | 0.012 | 0.000 | 0.006 |
| Candidate_division_TG3 | TG3_Subphylum_1 | TG3_Subphylum_1 | Termite_Cluster | Termite_cockroach_cluster | 2.481 | 0.838 | 0.206 | 0.198 | 0.075 | 0.036 |
| Candidate_division_TG3 | TG3_Subphylum_2 | TG3_Subphylum_2 | Termite_Cockroach_Cluster | Natranaerovirga | 0.000 | 0.000 | 0.000 | 0.000 | 0.005 | 0.000 |
| Candidate_phylum_TG3 | Subphylum_2 | incertae_sedis | Chitinivibrionaceae | Chitinivibrio | 0.000 | 0.000 | 0.006 | 0.000 | 0.000 | 0.000 |
| Candidate_phylum_TG3 | Subphylum_1 | incertae_sedis | Termite_cluster_III | Subcluster_IIIb | 0.000 | 0.000 | 0.018 | 0.000 | 0.000 | 0.000 |
| Candidate_phylum_TG3 | Subphylum_1 | incertae_sedis | Termite_cluster_III | Subcluster_IVb | 0.000 | 0.000 | 0.006 | 0.000 | 0.000 | 0.000 |
| Candidatus_Melainabacteria | Unclassified | Vampirovibrionales | Unclassified | Variovorax_1 | 0.008 | 0.006 | 0.149 | 0.006 | 0.000 | 0.006 |
| Chlamydiae | Chlamydiae | Chlamydiales | Chlamydiaceae | Chlamydia | 0.000 | 0.000 | 0.008 | 0.000 | 0.000 | 0.000 |
| Chlamydiae | Chlamydiae | Chlamydiales | Parachlamydiaceae | Candidatus_Protochlamydia | 0.000 | 0.000 | 0.010 | 0.000 | 0.000 | 0.000 |

**Table S9 continued.**

| | | | | | | | | | | |
|---|---|---|---|---|---|---|---|---|---|---|
| Chlamydiae | Chlamydiia | Parachlamydiales | Parachlamydiaceae | Paracoccus | 0.033 | 0.018 | 0.002 | 0.012 | 0.000 | 0.012 |
| Chlamydiae | Chlamydiae | Chlamydiales | Simkaniaceae | Candidatus_Fritschea | 0.000 | 0.000 | 0.006 | 0.000 | 0.000 | 0.000 |
| Chlamydiae | Chlamydiia | Chlamydiales | Simkaniaceae | Candidatus_Rhabdochlamydia | 0.000 | 0.000 | 0.018 | 0.000 | 0.000 | 0.000 |
| Chlamydiae | Chlamydiae | Chlamydiales | Waddliaceae | Waddlia | 0.000 | 0.000 | 0.008 | 0.000 | 0.000 | 0.000 |
| Chlorobi | Chlorobia | Chlorobiales | Chlorobiaceae | Chlorobaculum | 0.000 | 0.000 | 0.040 | 0.000 | 0.000 | 0.000 |
| Chlorobi | Chlorobia | Chlorobiales | Chlorobiaceae | Chlorobium_1 | 0.000 | 0.000 | 0.034 | 0.000 | 0.000 | 0.000 |
| Chlorobi | Chlorobia | Chlorobiales | Chlorobiaceae | Chlorobium_2 | 0.000 | 0.000 | 0.018 | 0.000 | 0.000 | 0.000 |
| Chlorobi | Chlorobia | Chlorobiales | Chlorobiaceae | Chloroherpeton | 0.000 | 0.000 | 0.006 | 0.000 | 0.000 | 0.000 |
| Chlorobi | Chlorobia | Chlorobiales | Chlorobiaceae | Prosthecochloris_1 | 0.000 | 0.000 | 0.004 | 0.000 | 0.000 | 0.000 |
| Chlorobi | Chlorobia | Chlorobiales | Chlorobiaceae | Prosthecochloris_2 | 0.000 | 0.000 | 0.004 | 0.000 | 0.000 | 0.000 |
| Chlorobi | Chlorobia | Chlorobiales | Chlorobiaceae | Prosthecochloris_3 | 0.000 | 0.000 | 0.004 | 0.000 | 0.000 | 0.000 |
| Chlorobi | Chlorobia | Chlorobiales | Chlorobiaceae | Prosthecochloris_4 | 0.000 | 0.000 | 0.004 | 0.000 | 0.000 | 0.000 |
| Chlorobi | Chlorobia | Chlorobiales | Chlorobiaceae | Prosthecochloris_5 | 0.000 | 0.000 | 0.004 | 0.000 | 0.000 | 0.000 |
| Chloroflexi | Anaerolineae | Anaerolineales | Anaerolineaceae | Anaerolinea | 0.000 | 0.000 | 0.075 | 0.000 | 0.000 | 0.000 |
| Chloroflexi | Anaerolineae | Anaerolineales | Anaerolineaceae | Bellilinea | 0.000 | 0.000 | 0.014 | 0.000 | 0.000 | 0.000 |
| Chloroflexi | Anaerolineae | Anaerolineales | Anaerolineaceae | Leptolinea | 0.000 | 0.000 | 0.123 | 0.000 | 0.000 | 0.000 |
| Chloroflexi | Anaerolineae | Anaerolineales | Anaerolineaceae | Levilinea | 0.000 | 0.000 | 0.018 | 0.000 | 0.000 | 0.000 |
| Chloroflexi | Anaerolineae | Anaerolineales | Anaerolineaceae | Longilinea | 0.000 | 0.000 | 0.246 | 0.000 | 0.000 | 0.000 |
| Chloroflexi | Caldilineae | Caldilineales | Caldilineaceae | Caldilinea | 0.016 | 0.000 | 0.161 | 0.000 | 0.000 | 0.000 |
| Chloroflexi | Chloroflexi | Chloroflexales | Candidatus_Chlorothrix | Candidatus_Chlorothrix | 0.000 | 0.000 | 0.052 | 0.000 | 0.000 | 0.000 |
| Chloroflexi | Chloroflexi | Chloroflexales | Chloroflexaceae | Chloroflexus | 0.000 | 0.000 | 0.016 | 0.000 | 0.000 | 0.000 |
| Chloroflexi | Chloroflexi | Chloroflexales | Chloroflexaceae | Chloronema | 0.000 | 0.000 | 0.004 | 0.000 | 0.000 | 0.000 |
| Chloroflexi | Dehalococcoidetes | Dehalococcoidetes | Dehalococcoidetes | Dehalococcoides | 0.000 | 0.000 | 0.020 | 0.000 | 0.000 | 0.000 |
| Chloroflexi | Dehalococcoidetes | Dehalococcoidetes | Dehalococcoidetes | Dehalogenimonas | 0.000 | 0.000 | 0.008 | 0.000 | 0.000 | 0.000 |
| Chloroflexi | Chloroflexi | Herpetosiphonales | Herpetosiphonaceae | Hespellia | 0.000 | 0.000 | 0.006 | 0.000 | 0.000 | 0.000 |
| Chloroflexi | Ktedonobacteria | Ktedonobacterales | Ktedonobacteraceae | Kurthia | 0.000 | 0.000 | 0.032 | 0.024 | 0.000 | 0.000 |
| Chloroflexi | Thermomicrobia | Sphaerobacterales | Sphaerobacteraceae | Sphaerobacter | 0.000 | 0.000 | 0.004 | 0.000 | 0.000 | 0.000 |
| Chloroflexi | Thermomicrobia | Sphaerobacteridae | Sphaerobacterales | Sphingobacterium | 0.000 | 0.012 | 0.012 | 0.006 | 0.000 | 0.000 |
| Chloroflexi | Thermomicrobia | Thermomicrobiales | Thermomicrobiaceae | Thermomicrobia | 0.000 | 0.000 | 0.010 | 0.000 | 0.000 | 0.000 |
| Chrysiogenetes | Chrysiogenetes | Chrysiogenales | Chrysiogenaceae | Desulfurispirillum | 0.000 | 0.000 | 0.004 | 0.000 | 0.000 | 0.000 |
| Cyanobacteria | Unclassified | Synechococcales | Acaryochloridaceae | Acaryochloris | 0.000 | 0.000 | 0.012 | 0.000 | 0.000 | 0.000 |
| Cyanobacteria | Cyanobacteria | Cyanobacteriales | Brasilonema | Brasilonema | 0.000 | 0.000 | 0.012 | 0.000 | 0.000 | 0.000 |
| Cyanobacteria | Cyanobacteria | Cyanobacteriales | Chloroplast | Chloroplast | 0.049 | 0.012 | 1.911 | 0.006 | 0.000 | 0.006 |
| Cyanobacteria | Cyanobacteria | Cyanobacteriales | Chroogloeocystis | Chroogloeocystis | 0.000 | 0.000 | 0.006 | 0.000 | 0.000 | 0.000 |
| Cyanobacteria | Cyanobacteria | Cyanobacteriales | Mastigocladopsis | Mastigocladopsis | 0.000 | 0.000 | 0.006 | 0.000 | 0.000 | 0.000 |
| Cyanobacteria | Cyanobacteria | Cyanobacteriales | Merismopedia | Merismopedia | 0.000 | 0.000 | 0.004 | 0.000 | 0.000 | 0.000 |
| Cyanobacteria | Cyanobacteria | Cyanobacteriales | ML635J-21 | Cluster_TG2 | 0.073 | 0.055 | 0.018 | 0.054 | 0.005 | 0.006 |
| Cyanobacteria | Cyanobacteria | Cyanobacteriales | SubsectionI | Chroococcus | 0.000 | 0.000 | 0.004 | 0.000 | 0.000 | 0.000 |
| Cyanobacteria | Cyanobacteria | Cyanobacteriales | SubsectionI | Cyanobacterium | 0.000 | 0.000 | 0.028 | 0.000 | 0.000 | 0.000 |
| Cyanobacteria | Cyanobacteria | Cyanobacteriales | SubsectionI | Cyanothece | 0.000 | 0.000 | 0.018 | 0.000 | 0.000 | 0.000 |

**Table S9 continued.**

| | | | | | | | | | | |
|---|---|---|---|---|---|---|---|---|---|---|
| Cyanobacteria | Cyanobacteria | Cyanobacteriales | SubsectionI | Gleocapsa | 0.000 | 0.000 | 0.010 | 0.000 | 0.000 | 0.000 |
| Cyanobacteria | Cyanobacteria | Cyanobacteriales | SubsectionI | Gloeobacter | 0.000 | 0.000 | 0.004 | 0.000 | 0.000 | 0.000 |
| Cyanobacteria | Cyanobacteria | Cyanobacteriales | SubsectionI | Gloeothece | 0.000 | 0.000 | 0.004 | 0.000 | 0.000 | 0.000 |
| Cyanobacteria | Cyanobacteria | Cyanobacteriales | SubsectionI | Microcystis | 0.000 | 0.000 | 0.050 | 0.000 | 0.000 | 0.000 |
| Cyanobacteria | Cyanobacteria | Cyanobacteriales | SubsectionI | Prochlorococcus | 0.000 | 0.000 | 0.127 | 0.000 | 0.000 | 0.000 |
| Cyanobacteria | Cyanobacteria | Cyanobacteriales | SubsectionI | Synechococcus | 0.000 | 0.000 | 0.026 | 0.000 | 0.000 | 0.000 |
| Cyanobacteria | Cyanobacteria | Cyanobacteriales | SubsectionI | Synechococcus_1 | 0.000 | 0.006 | 0.109 | 0.000 | 0.000 | 0.000 |
| Cyanobacteria | Cyanobacteria | Cyanobacteriales | SubsectionI | Synechococcus_2 | 0.000 | 0.000 | 0.006 | 0.000 | 0.000 | 0.000 |
| Cyanobacteria | Cyanobacteria | Cyanobacteriales | SubsectionI | Synechococcus_3 | 0.000 | 0.000 | 0.066 | 0.000 | 0.000 | 0.000 |
| Cyanobacteria | Cyanobacteria | Cyanobacteriales | SubsectionI | Synechococcus_4 | 0.000 | 0.000 | 0.020 | 0.000 | 0.000 | 0.000 |
| Cyanobacteria | Cyanobacteria | Cyanobacteriales | SubsectionI | Thermosynechococcus | 0.000 | 0.000 | 0.012 | 0.000 | 0.000 | 0.000 |
| Cyanobacteria | Cyanobacteria | Cyanobacteriales | SubsectionII | SubgroupI | 0.000 | 0.000 | 0.030 | 0.000 | 0.000 | 0.000 |
| Cyanobacteria | Cyanobacteria | Cyanobacteriales | SubsectionII | Succiniclasticum | 0.000 | 0.000 | 0.026 | 0.000 | 0.000 | 0.000 |
| Cyanobacteria | Cyanobacteria | Cyanobacteriales | SubsectionIII | Arthronema | 0.000 | 0.000 | 0.006 | 0.000 | 0.000 | 0.000 |
| Cyanobacteria | Cyanobacteria | Cyanobacteriales | SubsectionIII | Arthrospira | 0.000 | 0.000 | 0.014 | 0.000 | 0.000 | 0.000 |
| Cyanobacteria | Cyanobacteria | Cyanobacteriales | SubsectionIII | Chamaesiphon | 0.000 | 0.000 | 0.022 | 0.000 | 0.000 | 0.000 |
| Cyanobacteria | Cyanobacteria | Cyanobacteriales | SubsectionIII | Dactylococcopsis | 0.000 | 0.000 | 0.004 | 0.000 | 0.000 | 0.000 |
| Cyanobacteria | Cyanobacteria | Cyanobacteriales | SubsectionIII | Euhalothece | 0.000 | 0.000 | 0.016 | 0.000 | 0.000 | 0.000 |
| Cyanobacteria | Cyanobacteria | Cyanobacteriales | SubsectionIII | Halomicronema | 0.000 | 0.000 | 0.006 | 0.000 | 0.000 | 0.000 |
| Cyanobacteria | Cyanobacteria | Cyanobacteriales | SubsectionIII | Leptolyngbya_1 | 0.000 | 0.000 | 0.030 | 0.000 | 0.000 | 0.000 |
| Cyanobacteria | Cyanobacteria | Cyanobacteriales | SubsectionIII | Leptolyngbya_2 | 0.000 | 0.000 | 0.042 | 0.000 | 0.000 | 0.000 |
| Cyanobacteria | Cyanobacteria | Cyanobacteriales | SubsectionIII | Leptolyngbya_3 | 0.000 | 0.000 | 0.121 | 0.000 | 0.000 | 0.000 |
| Cyanobacteria | Cyanobacteria | Cyanobacteriales | SubsectionIII | Leptospira | 0.000 | 0.000 | 0.048 | 0.000 | 0.000 | 0.000 |
| Cyanobacteria | Cyanobacteria | Cyanobacteriales | SubsectionIII | Limnothrix | 0.000 | 0.000 | 0.016 | 0.000 | 0.000 | 0.000 |
| Cyanobacteria | Cyanobacteria | Cyanobacteriales | SubsectionIII | Lyngbya | 0.000 | 0.000 | 0.012 | 0.000 | 0.000 | 0.000 |
| Cyanobacteria | Cyanobacteria | Cyanobacteriales | SubsectionIII | Lyngbya_1 | 0.000 | 0.000 | 0.014 | 0.000 | 0.000 | 0.000 |
| Cyanobacteria | Cyanobacteria | Cyanobacteriales | SubsectionIII | Lyngbya_2 | 0.000 | 0.000 | 0.014 | 0.000 | 0.000 | 0.000 |
| Cyanobacteria | Cyanobacteria | Cyanobacteriales | SubsectionIII | Microcoleus_1 | 0.000 | 0.000 | 0.014 | 0.000 | 0.000 | 0.000 |
| Cyanobacteria | Cyanobacteria | Cyanobacteriales | SubsectionIII | Microcoleus_2 | 0.000 | 0.000 | 0.014 | 0.000 | 0.000 | 0.000 |
| Cyanobacteria | Cyanobacteria | Cyanobacteriales | SubsectionIII | Oscillatoria | 0.000 | 0.000 | 0.040 | 0.000 | 0.000 | 0.000 |
| Cyanobacteria | Cyanobacteria | Cyanobacteriales | SubsectionIII | Phormidium_1 | 0.000 | 0.000 | 0.018 | 0.000 | 0.000 | 0.000 |
| Cyanobacteria | Cyanobacteria | Cyanobacteriales | SubsectionIII | Phormidium_2 | 0.000 | 0.000 | 0.034 | 0.000 | 0.000 | 0.000 |
| Cyanobacteria | Cyanobacteria | Cyanobacteriales | SubsectionIII | Phormidium_3 | 0.000 | 0.000 | 0.010 | 0.000 | 0.000 | 0.000 |
| Cyanobacteria | Cyanobacteria | Cyanobacteriales | SubsectionIII | Phormidium_5 | 0.000 | 0.000 | 0.004 | 0.000 | 0.000 | 0.000 |
| Cyanobacteria | Cyanobacteria | Cyanobacteriales | SubsectionIII | Planktothrix | 0.000 | 0.000 | 0.022 | 0.000 | 0.000 | 0.000 |
| Cyanobacteria | Cyanobacteria | Cyanobacteriales | SubsectionIII | Prochlorothrix | 0.000 | 0.000 | 0.010 | 0.000 | 0.000 | 0.000 |
| Cyanobacteria | Cyanobacteria | Cyanobacteriales | SubsectionIII | Pseudanabaena | 0.000 | 0.000 | 0.004 | 0.000 | 0.000 | 0.000 |
| Cyanobacteria | Cyanobacteria | Cyanobacteriales | SubsectionIII | Pseudoanabaena | 0.000 | 0.000 | 0.004 | 0.000 | 0.000 | 0.000 |
| Cyanobacteria | Cyanobacteria | Cyanobacteriales | SubsectionIII | Rubidibacter | 0.000 | 0.000 | 0.004 | 0.000 | 0.000 | 0.000 |
| Cyanobacteria | Cyanobacteria | Cyanobacteriales | SubsectionIII | Spirulina | 0.000 | 0.000 | 0.022 | 0.000 | 0.000 | 0.000 |

**Table S9 continued.**

| | | | | | | | | | | |
|---|---|---|---|---|---|---|---|---|---|---|
| Cyanobacteria | Cyanobacteria | Cyanobacteriales | SubsectionIII | Trichodesmium | 0.000 | 0.000 | 0.004 | 0.000 | 0.000 | 0.000 |
| Cyanobacteria | Cyanobacteria | Cyanobacteriales | SubsectionIV | Anabaena_1 | 0.000 | 0.000 | 0.008 | 0.000 | 0.000 | 0.000 |
| Cyanobacteria | Cyanobacteria | Cyanobacteriales | SubsectionIV | Anabaena_10 | 0.000 | 0.000 | 0.006 | 0.000 | 0.000 | 0.000 |
| Cyanobacteria | Cyanobacteria | Cyanobacteriales | SubsectionIV | Anabaena_11 | 0.000 | 0.000 | 0.008 | 0.000 | 0.000 | 0.000 |
| Cyanobacteria | Cyanobacteria | Cyanobacteriales | SubsectionIV | Anabaena_3 | 0.000 | 0.000 | 0.014 | 0.000 | 0.000 | 0.000 |
| Cyanobacteria | Cyanobacteria | Cyanobacteriales | SubsectionIV | Anabaena_6 | 0.000 | 0.000 | 0.008 | 0.000 | 0.000 | 0.000 |
| Cyanobacteria | Cyanobacteria | Cyanobacteriales | SubsectionIV | Anabaena_9 | 0.000 | 0.000 | 0.008 | 0.000 | 0.000 | 0.000 |
| Cyanobacteria | Cyanobacteria | Cyanobacteriales | SubsectionIV | Anabaenopsis_1 | 0.000 | 0.000 | 0.010 | 0.000 | 0.000 | 0.000 |
| Cyanobacteria | Cyanobacteria | Cyanobacteriales | SubsectionIV | Anabaenopsis_2 | 0.000 | 0.000 | 0.004 | 0.000 | 0.000 | 0.000 |
| Cyanobacteria | Cyanobacteria | Cyanobacteriales | SubsectionIV | Aphanizomenon_3 | 0.000 | 0.000 | 0.004 | 0.000 | 0.000 | 0.000 |
| Cyanobacteria | Cyanobacteria | Cyanobacteriales | SubsectionIV | Aphanizomenon_4 | 0.000 | 0.000 | 0.010 | 0.000 | 0.000 | 0.000 |
| Cyanobacteria | Cyanobacteria | Cyanobacteriales | SubsectionIV | Calothrix | 0.000 | 0.000 | 0.004 | 0.000 | 0.000 | 0.000 |
| Cyanobacteria | Cyanobacteria | Cyanobacteriales | SubsectionIV | Cylindrospermosis | 0.000 | 0.000 | 0.008 | 0.000 | 0.000 | 0.000 |
| Cyanobacteria | Cyanobacteria | Cyanobacteriales | SubsectionIV | Gloeotrichia | 0.000 | 0.000 | 0.004 | 0.000 | 0.000 | 0.000 |
| Cyanobacteria | Cyanobacteria | Cyanobacteriales | SubsectionIV | Nodularia_1 | 0.000 | 0.000 | 0.032 | 0.000 | 0.000 | 0.000 |
| Cyanobacteria | Cyanobacteria | Cyanobacteriales | SubsectionIV | Nostoc_1 | 0.000 | 0.000 | 0.010 | 0.000 | 0.000 | 0.000 |
| Cyanobacteria | Cyanobacteria | Cyanobacteriales | SubsectionIV | Nostoc_2 | 0.000 | 0.000 | 0.004 | 0.000 | 0.000 | 0.000 |
| Cyanobacteria | Cyanobacteria | Cyanobacteriales | SubsectionIV | Nostoc_3 | 0.000 | 0.000 | 0.004 | 0.000 | 0.000 | 0.000 |
| Cyanobacteria | Cyanobacteria | Cyanobacteriales | SubsectionIV | Nostoc_5 | 0.000 | 0.000 | 0.008 | 0.000 | 0.000 | 0.000 |
| Cyanobacteria | Cyanobacteria | Cyanobacteriales | SubsectionIV | Nostoc_6 | 0.000 | 0.000 | 0.087 | 0.000 | 0.000 | 0.000 |
| Cyanobacteria | Cyanobacteria | Cyanobacteriales | SubsectionIV | Nostoc_7 | 0.000 | 0.000 | 0.004 | 0.000 | 0.000 | 0.000 |
| Cyanobacteria | Cyanobacteria | Cyanobacteriales | SubsectionIV | Noviherbaspirillum | 0.000 | 0.000 | 0.000 | 0.006 | 0.000 | 0.000 |
| Cyanobacteria | Cyanobacteria | Cyanobacteriales | SubsectionIV | Trichormus | 0.000 | 0.000 | 0.004 | 0.000 | 0.000 | 0.000 |
| Cyanobacteria | Cyanobacteria | Cyanobacteriales | SubsectionV | Chlorogloeopsis | 0.000 | 0.000 | 0.006 | 0.000 | 0.000 | 0.000 |
| Cyanobacteria | Cyanobacteria | Cyanobacteriales | SubsectionV | Fischerella | 0.000 | 0.000 | 0.030 | 0.000 | 0.000 | 0.000 |
| Cyanobacteria | Cyanobacteria | Cyanobacteriales | SubsectionV | Nostochopsis | 0.000 | 0.000 | 0.004 | 0.000 | 0.000 | 0.000 |
| Cyanobacteria | Cyanobacteria | Cyanobacteriales | SubsectionV | Stigonema | 0.000 | 0.000 | 0.006 | 0.000 | 0.000 | 0.000 |
| Cyanobacteria | Unclassified | Synechcoccales | Trichocoleusaceae | Tropicimonas | 0.000 | 0.000 | 0.010 | 0.000 | 0.000 | 0.000 |
| Deferribacteres | Deferribacteres | Deferribacterales | Deferribacteraceae | Calditerrivibrio | 0.000 | 0.000 | 0.014 | 0.000 | 0.000 | 0.000 |
| Deferribacteres | Deferribacteres | Deferribacterales | Deferribacteraceae | Deferribacter | 0.000 | 0.000 | 0.012 | 0.000 | 0.000 | 0.000 |
| Deferribacteres | Deferribacteres | Deferribacterales | Deferribacteraceae | Denitrovibrio | 0.000 | 0.000 | 0.004 | 0.000 | 0.000 | 0.000 |
| Deferribacteres | Deferribacteres | Deferribacterales | Deferribacteraceae | Geovibrio | 0.000 | 0.000 | 0.004 | 0.000 | 0.000 | 0.000 |
| Deferribacteres | Deferribacteres | Deferribacterales | Deferribacteraceae | Muricauda | 0.000 | 0.000 | 0.018 | 0.000 | 0.000 | 0.000 |
| Deinococcus-Thermus | Deinococci | Deinococcales | Deinococcaceae | Deinococcus | 0.000 | 0.000 | 0.206 | 0.000 | 0.000 | 0.000 |
| Deinococcus-Thermus | Deinococci | Thermales | Thermaceae | Melissococcus | 0.000 | 0.000 | 0.004 | 0.000 | 0.000 | 0.000 |
| Deinococcus-Thermus | Deinococcus-Thermus | Thermales | Thermaceae | Oceanithermus | 0.000 | 0.000 | 0.006 | 0.000 | 0.000 | 0.000 |
| Deinococcus-Thermus | Deinococcus-Thermus | Thermales | Thermaceae | Thermus | 0.000 | 0.000 | 0.095 | 0.000 | 0.000 | 0.000 |
| Deinococcus-Thermus | Deinococcus-Thermus | Thermales | Thermaceae | Thioalkalivibrio_1 | 0.000 | 0.000 | 0.016 | 0.000 | 0.000 | 0.000 |
| Deinococcus-Thermus | Deinococcus-Thermus | Thermales | Thermaceae | Vulcanithermus | 0.000 | 0.000 | 0.004 | 0.000 | 0.000 | 0.000 |
| Deinococcus-Thermus | Deinococcus-Thermus | Deinococcales | Trueperaceae | Truepera | 0.000 | 0.000 | 0.052 | 0.000 | 0.000 | 0.000 |

**Table S9 continued.**

| | | | | | | | | | | |
|---|---|---|---|---|---|---|---|---|---|---|
| Dictyoglomi | Dictyoglomia | Dictyoglomales | Dictyoglomaceae | Dictyoglomus | 0.000 | 0.000 | 0.012 | 0.000 | 0.000 | 0.000 |
| Elusimicrobia | Elusimicrobia | Elusimicrobiales | Elusimicrobiaceae | Listeria | 0.000 | 0.000 | 0.014 | 0.006 | 0.000 | 0.000 |
| Elusimicrobia | Elusimicrobia | Elusimicrobiales | Elusimicrobiaceae | Undibacterium | 0.041 | 0.043 | 0.038 | 0.006 | 0.005 | 0.006 |
| Elusimicrobia | Elusimicrobia | Endomicrobiales | Endomicrobiaceae | Environmental_cluster_II | 0.000 | 0.000 | 0.006 | 0.000 | 0.000 | 0.000 |
| Elusimicrobia | Elusimicrobia | Endomicrobiales | Endomicrobiaceae | Endomicrobium | 0.000 | 0.000 | 0.195 | 0.000 | 0.000 | 0.000 |
| Elusimicrobia | Elusimicrobia | Endomicrobiales | Endomicrobiaceae | Environmental_cluster_I | 0.000 | 0.000 | 0.010 | 0.000 | 0.000 | 0.000 |
| Fibrobacteres | Subphylum_2 | Insect_cluster | Cockroach_cluster_II | Subcluster_IIa | 0.000 | 0.000 | 0.012 | 0.000 | 0.000 | 0.000 |
| Fibrobacteres | Subphylum_2 | Insect_cluster | Cockroach_cluster_II | Subcluster_IIb | 0.000 | 0.000 | 0.008 | 0.000 | 0.000 | 0.000 |
| Fibrobacteres | Fibrobacteres_sp1 | Fibrobacterales | Fibrobacteraceae | Fibrobacter | 0.008 | 0.000 | 0.062 | 0.000 | 0.000 | 0.000 |
| Fibrobacteres | Subphylum_2 | Insect_cluster | Termite_cluster_I | Subcluster_IIIa | 0.000 | 0.000 | 0.048 | 0.000 | 0.000 | 0.006 |
| Fibrobacteres | Subphylum_2 | Insect_cluster | Termite_cluster_I | Subcluster_IVa | 0.138 | 0.185 | 0.006 | 0.012 | 0.005 | 0.000 |
| Firmicutes | Mollicutes | Acholeplasmatales | Acholeplasmataceae | Acholeplasma_2 | 0.000 | 0.000 | 0.016 | 0.000 | 0.000 | 0.000 |
| Firmicutes | Negativicutes | Acidaminococcales | Acidaminococcaceae | Sulfuricurvum | 0.000 | 0.000 | 0.022 | 0.000 | 0.000 | 0.000 |
| Firmicutes | Bacilli | Lactobacillales | Aerococcaceae | Abiotrophia | 0.000 | 0.000 | 0.004 | 0.000 | 0.000 | 0.000 |
| Firmicutes | Bacilli | Lactobacillales | Aerococcaceae | Eremococcus | 0.000 | 0.000 | 0.008 | 0.000 | 0.000 | 0.000 |
| Firmicutes | Bacilli | Lactobacillales | Aerococcaceae | Flacklamia | 0.000 | 0.000 | 0.014 | 0.000 | 0.000 | 0.000 |
| Firmicutes | Bacilli | Lactobacillales | Aerococcaceae | Ignavigranum | 0.000 | 0.000 | 0.008 | 0.000 | 0.000 | 0.000 |
| Firmicutes | Bacilli | Bacillales | Alicyclobacillaceae | Alicyclobacillus | 0.000 | 0.000 | 0.066 | 0.000 | 0.000 | 0.000 |
| Firmicutes | Mollicutes | Anaeroplasmatales | Anaeroplasmataceae | Anaeroplasma | 0.008 | 0.006 | 0.022 | 0.000 | 0.000 | 0.006 |
| Firmicutes | Bacilli | Bacillales | Bacillaceae | Amorphus | 0.000 | 0.006 | 0.000 | 0.000 | 0.000 | 0.000 |
| Firmicutes | Bacilli | Bacillales | Bacillaceae | Bacillus_1 | 0.000 | 0.000 | 0.036 | 0.030 | 0.005 | 0.000 |
| Firmicutes | Bacilli | Bacillales | Bacillaceae | Bacillus_10 | 0.008 | 0.000 | 0.022 | 0.006 | 0.005 | 0.000 |
| Firmicutes | Bacilli | Bacillales | Bacillaceae | Bacillus_11 | 0.008 | 0.068 | 0.073 | 1.887 | 0.474 | 0.000 |
| Firmicutes | Bacilli | Bacillales | Bacillaceae | Bacillus_12 | 0.000 | 0.000 | 0.016 | 0.000 | 0.000 | 0.000 |
| Firmicutes | Bacilli | Bacillales | Bacillaceae | Bacillus_14 | 0.155 | 0.111 | 0.189 | 0.323 | 0.040 | 0.024 |
| Firmicutes | Bacilli | Bacillales | Bacillaceae | Bacillus_15 | 0.000 | 0.000 | 0.006 | 0.000 | 0.000 | 0.000 |
| Firmicutes | Bacilli | Bacillales | Bacillaceae | Bacillus_16 | 0.008 | 0.000 | 0.014 | 0.000 | 0.000 | 0.000 |
| Firmicutes | Bacilli | Bacillales | Bacillaceae | Bacillus_2 | 0.000 | 0.006 | 0.038 | 0.000 | 0.000 | 0.000 |
| Firmicutes | Bacilli | Bacillales | Bacillaceae | Bacillus_3 | 0.000 | 0.000 | 0.032 | 0.000 | 0.010 | 0.000 |
| Firmicutes | Bacilli | Bacillales | Bacillaceae | Bacillus_4 | 0.024 | 0.043 | 0.079 | 0.054 | 0.005 | 0.061 |
| Firmicutes | Bacilli | Bacillales | Bacillaceae | Bacillus_5 | 0.024 | 0.006 | 0.071 | 0.012 | 0.005 | 0.000 |
| Firmicutes | Bacilli | Bacillales | Bacillaceae | Bacillus_6 | 0.000 | 0.006 | 0.060 | 0.012 | 0.005 | 0.000 |
| Firmicutes | Bacilli | Bacillales | Bacillaceae | Bacillus_7 | 0.008 | 0.487 | 0.032 | 0.024 | 0.010 | 0.018 |
| Firmicutes | Bacilli | Bacillales | Bacillaceae | Bacillus_9 | 0.000 | 0.000 | 0.030 | 0.006 | 0.000 | 0.006 |
| Firmicutes | Bacilli | Bacillales | Bacillaceae | Caldalkalibacillus | 0.000 | 0.000 | 0.004 | 0.000 | 0.000 | 0.000 |
| Firmicutes | Bacilli | Bacillales | Bacillaceae | Domibacillus | 0.008 | 0.000 | 0.000 | 0.000 | 0.000 | 0.000 |
| Firmicutes | Bacilli | Bacillales | Bacillaceae | Fictibacillus | 0.000 | 0.006 | 0.002 | 0.000 | 0.000 | 0.000 |
| Firmicutes | Bacilli | Bacillales | Bacillaceae | Gracilibacillus | 0.000 | 0.000 | 0.016 | 0.000 | 0.000 | 0.000 |
| Firmicutes | Bacilli | Bacillales | Bacillaceae | Halocella | 0.000 | 0.000 | 0.026 | 0.000 | 0.000 | 0.000 |
| Firmicutes | Bacilli | Bacillales | Bacillaceae | Halochromatium | 0.000 | 0.000 | 0.006 | 0.000 | 0.000 | 0.000 |

**Table S9 continued.**

| | | | | | | | | | | |
|---|---|---|---|---|---|---|---|---|---|---|
| Firmicutes | Bacilli | Bacillales | Bacillaceae | Natronoflexus | 0.008 | 0.000 | 0.000 | 0.000 | 0.000 | 0.000 |
| Firmicutes | Bacilli | Bacillales | Bacillaceae | Pontibacter | 0.049 | 0.000 | 0.000 | 0.000 | 0.000 | 0.000 |
| Firmicutes | Bacilli | Bacillales | Bacillaceae | Psychromonas | 0.000 | 0.000 | 0.030 | 0.000 | 0.000 | 0.000 |
| Firmicutes | Bacilli | Bacillales | Bacillaceae | Salmonella | 0.822 | 6.235 | 0.238 | 1.546 | 0.698 | 1.089 |
| Firmicutes | Bacilli | Bacillales | Bacillaceae_1 | Anoxybacillus | 0.000 | 0.006 | 0.040 | 0.012 | 0.000 | 0.000 |
| Firmicutes | Bacilli | Bacillales | Bacillaceae_1 | Geobacillus_1 | 0.000 | 0.000 | 0.038 | 0.006 | 0.000 | 0.000 |
| Firmicutes | Bacilli | Bacillales | Bacillaceae_1 | Geobacillus_2 | 0.000 | 0.000 | 0.010 | 0.000 | 0.000 | 0.006 |
| Firmicutes | Bacilli | Bacillales | Bacillaceae_3 | Bacillus | 0.342 | 1.047 | 0.240 | 7.728 | 1.979 | 0.145 |
| Firmicutes | Bacilli | Bacillales | Bacillaceae_3 | Paucisalibacillus | 0.000 | 0.000 | 0.008 | 0.000 | 0.000 | 0.000 |
| Firmicutes | Bacilli | Bacillales | Bacillaceae_3 | Pectobacterium | 0.016 | 0.012 | 0.038 | 0.006 | 0.010 | 0.012 |
| Firmicutes | Bacilli | Bacillales | Bacillaceae_5 | Alkalibacillus | 0.000 | 0.000 | 0.010 | 0.000 | 0.000 | 0.000 |
| Firmicutes | Bacilli | Bacillales | Bacillaceae_5 | Amphibacillus_2 | 0.000 | 0.000 | 0.006 | 0.000 | 0.000 | 0.000 |
| Firmicutes | Bacilli | Bacillales | Bacillaceae_5 | Cerasibacillus | 0.000 | 0.000 | 0.004 | 0.000 | 0.000 | 0.000 |
| Firmicutes | Bacilli | Bacillales | Bacillaceae_5 | Halobacillus_1 | 0.000 | 0.000 | 0.010 | 0.000 | 0.000 | 0.000 |
| Firmicutes | Bacilli | Bacillales | Bacillaceae_5 | Halobacillus_7 | 0.000 | 0.000 | 0.004 | 0.000 | 0.000 | 0.000 |
| Firmicutes | Bacilli | Bacillales | Bacillaceae_5 | Halolactibacillus | 0.000 | 0.000 | 0.004 | 0.000 | 0.000 | 0.000 |
| Firmicutes | Bacilli | Bacillales | Bacillaceae_5 | Lentibacillus | 0.000 | 0.000 | 0.018 | 0.000 | 0.000 | 0.000 |
| Firmicutes | Bacilli | Bacillales | Bacillaceae_5 | Ochrobactrum | 0.016 | 0.000 | 0.002 | 0.012 | 0.010 | 0.067 |
| Firmicutes | Bacilli | Bacillales | Bacillaceae_5 | Ornithinibacillus_1 | 0.000 | 0.000 | 0.004 | 0.000 | 0.000 | 0.000 |
| Firmicutes | Bacilli | Bacillales | Bacillaceae_5 | Piscibacillus | 0.000 | 0.000 | 0.004 | 0.000 | 0.000 | 0.000 |
| Firmicutes | Bacilli | Bacillales | Bacillaceae_5 | Salinibacillus | 0.000 | 0.000 | 0.006 | 0.000 | 0.000 | 0.000 |
| Firmicutes | Bacilli | Bacillales | Bacillaceae_5 | Sediminibacillus | 0.000 | 0.000 | 0.004 | 0.000 | 0.000 | 0.000 |
| Firmicutes | Bacilli | Bacillales | Bacillaceae_5 | Virgibacillus_3 | 0.000 | 0.000 | 0.008 | 0.000 | 0.000 | 0.006 |
| Firmicutes | Bacilli | Bacillales | Bacillaceae_5 | Virgibacillus_5 | 0.000 | 0.000 | 0.012 | 0.000 | 0.000 | 0.000 |
| Firmicutes | Bacilli | Bacillales | Bacillaceae_5 | Virgibacillus_6 | 0.000 | 0.000 | 0.004 | 0.000 | 0.000 | 0.000 |
| Firmicutes | Bacilli | Bacillales | Bacillaceae_6 | Marinococcus | 0.000 | 0.000 | 0.004 | 0.006 | 0.000 | 0.000 |
| Firmicutes | Bacilli | Bacillales | Bacillaceae_6 | Salsuginibacillus | 0.000 | 0.000 | 0.004 | 0.000 | 0.000 | 0.000 |
| Firmicutes | Clostridia_2 | Candidatus_Desulforudis | Candidatus_Desulforudis | Candidatus_Desulforudis | 0.000 | 0.000 | 0.004 | 0.000 | 0.000 | 0.000 |
| Firmicutes | Bacilli | Lactobacillales | Carnobacteriaceae | Alkalibacterium_1 | 0.000 | 0.000 | 0.008 | 0.000 | 0.000 | 0.000 |
| Firmicutes | Bacilli | Lactobacillales | Carnobacteriaceae | Alkalibacterium_2 | 0.000 | 0.000 | 0.004 | 0.000 | 0.000 | 0.000 |
| Firmicutes | Bacilli | Lactobacillales | Carnobacteriaceae | Atopococcus | 0.000 | 0.000 | 0.006 | 0.000 | 0.000 | 0.000 |
| Firmicutes | Bacilli | Lactobacillales | Carnobacteriaceae | Aurantimonas_2 | 0.000 | 0.000 | 0.004 | 0.000 | 0.000 | 0.000 |
| Firmicutes | Bacilli | Lactobacillales | Carnobacteriaceae | Cardiobacterium | 0.000 | 0.000 | 0.006 | 0.000 | 0.000 | 0.000 |
| Firmicutes | Bacilli | Lactobacillales | Carnobacteriaceae | Carnobacterium_1 | 0.000 | 0.000 | 0.010 | 0.000 | 0.000 | 0.000 |
| Firmicutes | Bacilli | Lactobacillales | Carnobacteriaceae | Carnobacterium_3 | 0.000 | 0.000 | 0.012 | 0.000 | 0.000 | 0.000 |
| Firmicutes | Bacilli | Lactobacillales | Carnobacteriaceae | Desemzia | 0.000 | 0.000 | 0.008 | 0.000 | 0.000 | 0.000 |
| Firmicutes | Bacilli | Lactobacillales | Carnobacteriaceae | Dolosigranulum | 0.000 | 0.000 | 0.004 | 0.000 | 0.000 | 0.000 |
| Firmicutes | Bacilli | Lactobacillales | Carnobacteriaceae | Granulicatella | 0.016 | 0.000 | 0.008 | 0.000 | 0.000 | 0.000 |
| Firmicutes | Bacilli | Lactobacillales | Carnobacteriaceae | Mariniphaga | 0.008 | 0.006 | 0.000 | 0.000 | 0.000 | 0.000 |
| Firmicutes | Bacilli | Lactobacillales | Carnobacteriaceae | Trichocoleus | 0.000 | 0.000 | 0.004 | 0.000 | 0.000 | 0.000 |

**Table S9 continued.**

| | | | | | | | | | | |
|---|---|---|---|---|---|---|---|---|---|---|
| Firmicutes | Clostridia_1 | Clostridiales | Christensenellaceae | Christensenella | 0.016 | 0.006 | 0.000 | 0.000 | 0.000 | 0.000 |
| Firmicutes | Clostridia | Clostridiales | Clostridiaceae | Alkaliphilus_2 | 0.000 | 0.000 | 0.006 | 0.000 | 0.000 | 0.000 |
| Firmicutes | Clostridia | Clostridiales | Clostridiaceae | Lactovum | 1.180 | 0.246 | 0.026 | 0.036 | 0.015 | 0.024 |
| Firmicutes | Clostridia | Clostridiales | Clostridiaceae | Morganella | 0.000 | 0.000 | 0.014 | 0.000 | 0.000 | 0.000 |
| Firmicutes | Clostridia | Clostridiales | Clostridiaceae | Oceanisphaera | 0.000 | 0.000 | 0.004 | 0.000 | 0.000 | 0.000 |
| Firmicutes | Clostridia | Clostridiales | Clostridiaceae | Thermodesulfatator | 0.000 | 0.000 | 0.004 | 0.000 | 0.000 | 0.000 |
| Firmicutes | Clostridia_1 | Clostridiales | Clostridiaceae | Clostridium | 0.130 | 0.018 | 0.022 | 0.024 | 0.000 | 0.000 |
| Firmicutes | Clostridia_1 | Clostridiales | Clostridiaceae_1 | Arthropod_cluster | 0.000 | 0.006 | 0.006 | 0.000 | 0.000 | 0.000 |
| Firmicutes | Clostridia_1 | Clostridiales | Clostridiaceae_1 | Asteroleplasma | 0.057 | 0.216 | 0.016 | 0.677 | 2.078 | 5.968 |
| Firmicutes | Clostridia_1 | Clostridiales | Clostridiaceae_1 | Caloramator_1 | 0.000 | 0.000 | 0.004 | 0.000 | 0.000 | 0.000 |
| Firmicutes | Clostridia_1 | Clostridiales | Clostridiaceae_1 | Caloramator_2 | 0.000 | 0.000 | 0.004 | 0.000 | 0.000 | 0.000 |
| Firmicutes | Clostridia_1 | Clostridiales | Clostridiaceae_1 | Candidatus_Savagella | 0.000 | 0.000 | 0.006 | 0.000 | 0.000 | 0.000 |
| Firmicutes | Clostridia_1 | Clostridiales | Clostridiaceae_1 | Clostridium_1 | 0.008 | 0.000 | 0.385 | 0.000 | 0.000 | 0.000 |
| Firmicutes | Clostridia_1 | Clostridiales | Clostridiaceae_1 | Clostridium_10 | 0.000 | 0.000 | 0.012 | 0.000 | 0.000 | 0.000 |
| Firmicutes | Clostridia_1 | Clostridiales | Clostridiaceae_1 | Clostridium_11 | 0.008 | 0.000 | 0.016 | 0.000 | 0.000 | 0.000 |
| Firmicutes | Clostridia_1 | Clostridiales | Clostridiaceae_1 | Clostridium_2 | 0.000 | 0.000 | 0.008 | 0.000 | 0.000 | 0.000 |
| Firmicutes | Clostridia_1 | Clostridiales | Clostridiaceae_1 | Clostridium_3 | 0.033 | 0.012 | 0.012 | 0.000 | 0.000 | 0.000 |
| Firmicutes | Clostridia_1 | Clostridiales | Clostridiaceae_1 | Clostridium_5 | 0.000 | 0.000 | 0.010 | 0.000 | 0.000 | 0.000 |
| Firmicutes | Clostridia_1 | Clostridiales | Clostridiaceae_1 | Clostridium_7 | 0.000 | 0.000 | 0.020 | 0.000 | 0.000 | 0.000 |
| Firmicutes | Clostridia_1 | Clostridiales | Clostridiaceae_1 | Clostridium_8 | 0.000 | 0.000 | 0.093 | 0.000 | 0.000 | 0.000 |
| Firmicutes | Clostridia_1 | Clostridiales | Clostridiaceae_1 | Clostridium_9 | 0.008 | 0.000 | 0.022 | 0.000 | 0.000 | 0.000 |
| Firmicutes | Clostridia_1 | Clostridiales | Clostridiaceae_2 | Alkaliphilus_1 | 0.000 | 0.000 | 0.014 | 0.000 | 0.000 | 0.000 |
| Firmicutes | Clostridia_1 | Clostridiales | Clostridiaceae_2 | Tindallia | 0.000 | 0.000 | 0.004 | 0.000 | 0.000 | 0.000 |
| Firmicutes | Clostridia_1 | Clostridiales | Clostridiaceae_4 | Caminicella | 0.000 | 0.000 | 0.006 | 0.000 | 0.000 | 0.000 |
| Firmicutes | Clostridia_1 | Clostridiales | Clostridiaceae_5 | Caloranaerobacter | 0.000 | 0.000 | 0.004 | 0.000 | 0.000 | 0.000 |
| Firmicutes | Clostridia_1 | Clostridiales | Clostridiaceae_5 | Clostridiisalibacter | 0.000 | 0.000 | 0.006 | 0.000 | 0.000 | 0.000 |
| Firmicutes | Erysipelotrichia | Enterobacteriales | Enterobacteriaceae | Breznakia | 0.073 | 0.012 | 0.006 | 0.012 | 0.000 | 0.030 |
| Firmicutes | Bacilli | Lactobacillales | Enterococcaceae | Enterococcus | 0.179 | 0.043 | 0.137 | 0.006 | 0.000 | 0.000 |
| Firmicutes | Bacilli | Lactobacillales | Enterococcaceae | Enterococcus_2 | 0.049 | 0.006 | 0.008 | 0.000 | 0.000 | 0.000 |
| Firmicutes | Bacilli | Lactobacillales | Enterococcaceae | Enterococcus_3 | 0.000 | 0.000 | 0.024 | 0.000 | 0.000 | 0.000 |
| Firmicutes | Bacilli | Lactobacillales | Enterococcaceae | Tetrathiobacter | 0.106 | 0.006 | 0.012 | 0.108 | 0.075 | 0.061 |
| Firmicutes | Bacilli | Lactobacillales | Enterococcaceae_2 | Enterococcus_1 | 0.024 | 0.000 | 0.022 | 0.000 | 0.000 | 0.000 |
| Firmicutes | Mollicutes | Entomoplasmatales | Entomoplasmataceae | Mesorhizobium | 0.000 | 0.006 | 0.004 | 0.000 | 0.000 | 0.000 |
| Firmicutes | Erysipelotrichi | Erysipelotrichales | Erysipelotrichaceae | Atopobium | 0.000 | 0.000 | 0.040 | 0.000 | 0.000 | 0.000 |
| Firmicutes | Erysipelotrichi | Erysipelotrichales | Erysipelotrichaceae | Shigella | 0.016 | 0.006 | 0.002 | 0.000 | 0.000 | 0.000 |
| Firmicutes | Erysipelotrichia | Erysipelotrichales | Erysipelotrichaceae | Allobaculum | 0.000 | 0.000 | 0.101 | 0.000 | 0.000 | 0.000 |
| Firmicutes | Erysipelotrichia | Erysipelotrichales | Erysipelotrichaceae | Catenibacterium | 0.000 | 0.000 | 0.042 | 0.000 | 0.000 | 0.000 |
| Firmicutes | Erysipelotrichia | Erysipelotrichales | Erysipelotrichaceae | Coprobacillus | 0.000 | 0.000 | 0.004 | 0.000 | 0.000 | 0.000 |
| Firmicutes | Erysipelotrichia | Erysipelotrichales | Erysipelotrichaceae | Erysipelothrix | 0.016 | 0.000 | 0.022 | 0.012 | 0.000 | 0.000 |
| Firmicutes | Erysipelotrichia | Erysipelotrichales | Erysipelotrichaceae | Uliginosibacterium | 0.000 | 0.000 | 0.006 | 0.000 | 0.000 | 0.000 |

**Table S9 continued.**

| | | | | | | | | | | |
|---|---|---|---|---|---|---|---|---|---|---|
| Firmicutes | Clostridia_1 | Clostridiales | Eubacteriaceae_1 | Acetobacterium | 0.000 | 0.000 | 0.024 | 0.000 | 0.000 | 0.000 |
| Firmicutes | Clostridia_1 | Clostridiales | Eubacteriaceae_1 | Anaerofustis | 0.000 | 0.000 | 0.020 | 0.000 | 0.000 | 0.000 |
| Firmicutes | Clostridia_1 | Clostridiales | Eubacteriaceae_1 | Eubacterium_1 | 0.000 | 0.006 | 0.030 | 0.000 | 0.000 | 0.000 |
| Firmicutes | Clostridia_1 | Clostridiales | Eubacteriaceae_1 | Pseudoramibacter | 0.000 | 0.000 | 0.006 | 0.000 | 0.000 | 0.000 |
| Firmicutes | Clostridia | Thermoanaerobacterales | Family_III._Incertae_Sedis | Caldicellulosiruptor | 0.000 | 0.000 | 0.028 | 0.000 | 0.000 | 0.000 |
| Firmicutes | Clostridia | Thermoanaerobacterales | Family_III_Incertae_Sedis | Caldanaerobius | 0.000 | 0.000 | 0.006 | 0.000 | 0.000 | 0.000 |
| Firmicutes | Bacilli | Bacillales | Family_X_Incertae_Sedis | Thermicanus | 0.000 | 0.000 | 0.008 | 0.000 | 0.000 | 0.000 |
| Firmicutes | Bacilli | Bacillales | Family_X_Incertae_Sedis | Thermoactinomyces | 0.000 | 0.000 | 0.004 | 0.000 | 0.000 | 0.000 |
| Firmicutes | Bacilli | Bacillales | Family_XI_Incertae_Sedis | Gemella | 0.000 | 0.000 | 0.024 | 0.006 | 0.000 | 0.000 |
| Firmicutes | Clostridia | Clostridiales | Family_XI_Incertae_Sedis | Tepidiphilus | 0.008 | 0.006 | 0.014 | 0.006 | 0.000 | 0.000 |
| Firmicutes | Clostridia_0 | Clostridiales | Family_XI_Incertae_Sedis | Anaerococcus | 0.000 | 0.006 | 0.000 | 0.006 | 0.000 | 0.000 |
| Firmicutes | Clostridia_1 | Clostridiales | Family_XI_Incertae_Sedis | Aerococcus | 0.000 | 0.000 | 0.022 | 0.000 | 0.000 | 0.000 |
| Firmicutes | Clostridia_1 | Clostridiales | Family_XI_Incertae_Sedis | Anaerococcus_1 | 0.000 | 0.000 | 0.101 | 0.000 | 0.000 | 0.000 |
| Firmicutes | Clostridia_1 | Clostridiales | Family_XI_Incertae_Sedis | Finegoldia_2 | 0.000 | 0.000 | 0.012 | 0.000 | 0.000 | 0.000 |
| Firmicutes | Clostridia_1 | Clostridiales | Family_XI_Incertae_Sedis | Parvimonas | 0.000 | 0.000 | 0.014 | 0.000 | 0.000 | 0.000 |
| Firmicutes | Clostridia_1 | Clostridiales | Family_XI_Incertae_Sedis | Peredibacter | 0.000 | 0.000 | 0.012 | 0.000 | 0.000 | 0.000 |
| Firmicutes | Clostridia_1 | Clostridiales | Family_XI_Incertae_Sedis | Soehngenia | 0.000 | 0.000 | 0.006 | 0.000 | 0.000 | 0.000 |
| Firmicutes | Clostridia_1 | Clostridiales | Family_XI_Incertae_Sedis | Sporobacter | 0.000 | 0.006 | 0.004 | 0.000 | 0.000 | 0.000 |
| Firmicutes | Clostridia_1 | Clostridiales | Family_XI_Incertae_Sedis | Tepidimicrobium | 0.000 | 0.000 | 0.010 | 0.000 | 0.000 | 0.000 |
| Firmicutes | Clostridia_1 | Clostridiales | Family_XI_Incertae_Sedis | Tissierella_1 | 0.000 | 0.000 | 0.006 | 0.000 | 0.000 | 0.000 |
| Firmicutes | Clostridia_1 | Clostridiales | Family_XI_Incertae_Sedis | Tissierella_2 | 0.000 | 0.000 | 0.006 | 0.000 | 0.000 | 0.000 |
| Firmicutes | Bacilli | Bacillales | Family_XII_Incertae_Sedis | Exiguobacterium | 0.000 | 0.000 | 0.026 | 0.006 | 0.005 | 0.000 |
| Firmicutes | Clostridia | Clostridiales | Family_XII_Incertae_Sedis | Acidaminobacter | 0.000 | 0.000 | 0.006 | 0.000 | 0.000 | 0.000 |
| Firmicutes | Clostridia | Clostridiales | Family_XII_Incertae_Sedis | Fusibacter | 0.000 | 0.000 | 0.020 | 0.000 | 0.000 | 0.000 |
| Firmicutes | Clostridia_1 | Clostridiales | Family_XII_Incertae_Sedis | Guggenheimella | 0.008 | 0.000 | 0.006 | 0.000 | 0.000 | 0.000 |
| Firmicutes | Clostridia_1 | Clostridiales | Family_XIII_Incertae_Sedis | Anaerovorax_1 | 0.000 | 0.000 | 0.024 | 0.012 | 0.000 | 0.000 |
| Firmicutes | Clostridia_1 | Clostridiales | Family_XIII_Incertae_Sedis | Anaerovorax_2 | 0.000 | 0.000 | 0.020 | 0.000 | 0.000 | 0.000 |
| Firmicutes | Clostridia_1 | Clostridiales | Family_XIII_Incertae_Sedis | Eubacterium_2 | 0.000 | 0.000 | 0.064 | 0.000 | 0.000 | 0.000 |
| Firmicutes | Clostridia_1 | Clostridiales | Family_XIII_Incertae_Sedis | Eubacterium_3 | 0.033 | 0.000 | 0.034 | 0.000 | 0.000 | 0.000 |
| Firmicutes | Clostridia_1 | Clostridiales | Family_XIII_Incertae_Sedis | Termite_group_aaa | 0.065 | 0.031 | 0.016 | 0.012 | 0.000 | 0.000 |
| Firmicutes | Clostridia_1 | Clostridiales | Family_XIV_Incertae_Sedis | Anaerobranca | 0.000 | 0.000 | 0.008 | 0.000 | 0.000 | 0.000 |
| Firmicutes | Clostridia | Clostridiales | Family_XVI_Incertae_Sedis | Carboxydocella | 0.000 | 0.000 | 0.004 | 0.000 | 0.000 | 0.000 |
| Firmicutes | Clostridia | Clostridiales | Family_XVII_Incertae_Sedis | Thermaerobacter | 0.000 | 0.000 | 0.006 | 0.000 | 0.000 | 0.000 |
| Firmicutes | Clostridia | Clostridiales | Family_XVII_Incertae_Sedis | Thermanaerovibrio | 0.000 | 0.000 | 0.004 | 0.000 | 0.000 | 0.000 |
| Firmicutes | Clostridia_2 | Clostridiales_2 | Family_XVII_Incertae_Sedis | Sulfobacillus | 0.000 | 0.000 | 0.026 | 0.000 | 0.000 | 0.000 |
| Firmicutes | Clostridia_2 | Clostridiales_2 | Family_XVIII_Incertae_Sedis | Symbiobacterium | 0.000 | 0.000 | 0.040 | 0.000 | 0.000 | 0.000 |
| Firmicutes | Tissierellia | Tissierellales | Gottschalkiaceae | Anderseniella | 0.000 | 0.000 | 0.004 | 0.000 | 0.000 | 0.000 |
| Firmicutes | Tissierellia | Tissierellales | Gottschalkiaceae | Gottschalkia | 0.000 | 0.000 | 0.004 | 0.000 | 0.000 | 0.000 |
| Firmicutes | Clostridia_1 | Clostridiales | Gracilibacteraceae | Lysinibacillus | 0.008 | 0.000 | 0.036 | 0.210 | 0.010 | 0.121 |
| Firmicutes | Clostridia | Clostridiales | Hungateiclostridiaceae | Ruminococcus_1 | 0.008 | 0.000 | 0.278 | 0.000 | 0.000 | 0.000 |

**Table S9 continued.**

| | | | | | | | | | | |
|---|---|---|---|---|---|---|---|---|---|---|
| Firmicutes | Clostridia | Clostridiales | Hungateiclostridiaceae | Saccharopolyspora | 0.000 | 0.000 | 0.038 | 0.000 | 0.000 | 0.000 |
| Firmicutes | Clostridia | Clostridiales | Lachnospiraceae | Catonella | 0.000 | 0.000 | 0.004 | 0.000 | 0.000 | 0.000 |
| Firmicutes | Clostridia | Clostridiales | Lachnospiraceae | Cuneatibacter | 0.024 | 0.006 | 0.000 | 0.000 | 0.000 | 0.000 |
| Firmicutes | Clostridia | CLostridiales | Lachnospiraceae | Pseudochrobactrum | 0.000 | 0.012 | 0.018 | 0.000 | 0.000 | 0.012 |
| Firmicutes | Clostridia | Clostridiales | Lachnospiraceae | Roseburia_4 | 0.000 | 0.000 | 0.044 | 0.000 | 0.000 | 0.000 |
| Firmicutes | Clostridia | Clostridiales | Lachnospiraceae | Uncultured_47 | 0.000 | 0.000 | 0.004 | 0.000 | 0.000 | 0.000 |
| Firmicutes | Clostridia | Clostridiales | Lachnospiraceae | Uncultured_62 | 0.033 | 0.037 | 0.008 | 0.012 | 0.000 | 0.000 |
| Firmicutes | Clostridia | Clostridiales | Lachnospiraceae | Uncultured_63 | 0.000 | 0.000 | 0.006 | 0.000 | 0.000 | 0.000 |
| Firmicutes | Clostridia | Clostridiales | Lachnospiraceae | Uncultured_68 | 0.000 | 0.000 | 0.012 | 0.000 | 0.000 | 0.000 |
| Firmicutes | Clostridia | Clostridiales | Lachnospiraceae | Uncultured_7 | 0.130 | 0.018 | 0.637 | 0.018 | 0.010 | 0.000 |
| Firmicutes | Clostridia_0 | Clostridiales | Lachnospiraceae | Parasporobacterium-Sporobacterium | 0.041 | 0.018 | 0.008 | 0.006 | 0.000 | 0.000 |
| Firmicutes | Clostridia_1 | Clostridiales | Lachnospiraceae | Acetitomaculum | 0.000 | 0.000 | 0.046 | 0.000 | 0.000 | 0.000 |
| Firmicutes | Clostridia_1 | Clostridiales | Lachnospiraceae | Anaerostipes | 0.000 | 0.000 | 0.006 | 0.000 | 0.000 | 0.000 |
| Firmicutes | Clostridia_1 | Clostridiales | Lachnospiraceae | Blautia | 0.024 | 0.012 | 1.028 | 0.006 | 0.000 | 0.000 |
| Firmicutes | Clostridia_1 | Clostridiales | Lachnospiraceae | Butyrivibrio | 0.008 | 0.000 | 0.000 | 0.000 | 0.000 | 0.000 |
| Firmicutes | Clostridia_1 | Clostridiales | Lachnospiraceae | Butyrivibrio_1 | 0.000 | 0.000 | 0.020 | 0.000 | 0.000 | 0.000 |
| Firmicutes | Clostridia_1 | Clostridiales | Lachnospiraceae | Butyrivibrio_2 | 0.016 | 0.000 | 0.069 | 0.000 | 0.000 | 0.000 |
| Firmicutes | Clostridia_1 | Clostridiales | Lachnospiraceae | Butyrivibrio_3 | 0.008 | 0.006 | 0.077 | 0.000 | 0.000 | 0.000 |
| Firmicutes | Clostridia_1 | Clostridiales | Lachnospiraceae | Butyrivibrio_4 | 0.000 | 0.000 | 0.004 | 0.000 | 0.000 | 0.000 |
| Firmicutes | Clostridia_1 | Clostridiales | Lachnospiraceae | Candidatus_Arthromitus | 2.717 | 0.776 | 0.236 | 0.473 | 0.000 | 0.000 |
| Firmicutes | Clostridia_1 | Clostridiales | Lachnospiraceae | Catabacter | 0.041 | 0.006 | 0.022 | 0.006 | 0.000 | 0.000 |
| Firmicutes | Clostridia_1 | Clostridiales | Lachnospiraceae | Clostridium_piliforme_cluster | 0.016 | 0.006 | 0.026 | 0.000 | 0.000 | 0.000 |
| Firmicutes | Clostridia_1 | Clostridiales | Lachnospiraceae | Coprococcus_1 | 0.000 | 0.000 | 0.123 | 0.000 | 0.000 | 0.000 |
| Firmicutes | Clostridia_1 | Clostridiales | Lachnospiraceae | Coprococcus_2 | 0.008 | 0.000 | 0.036 | 0.000 | 0.000 | 0.000 |
| Firmicutes | Clostridia_1 | Clostridiales | Lachnospiraceae | Dorea | 0.008 | 0.000 | 0.123 | 0.000 | 0.000 | 0.000 |
| Firmicutes | Clostridia_1 | Clostridiales | Lachnospiraceae | Epulopiscium | 0.000 | 0.000 | 0.020 | 0.000 | 0.000 | 0.000 |
| Firmicutes | Clostridia_1 | Clostridiales | Lachnospiraceae | Gut_cluster_10 | 0.000 | 0.006 | 0.187 | 0.000 | 0.000 | 0.000 |
| Firmicutes | Clostridia_1 | Clostridiales | Lachnospiraceae | Gut_cluster_13 | 0.879 | 0.505 | 0.808 | 0.078 | 0.030 | 0.042 |
| Firmicutes | Clostridia_1 | Clostridiales | Lachnospiraceae | Gut_cluster_14 | 1.163 | 0.351 | 0.111 | 0.156 | 0.015 | 0.012 |
| Firmicutes | Clostridia_1 | Clostridiales | Lachnospiraceae | Gut_cluster_15 | 0.789 | 0.080 | 0.040 | 0.114 | 0.005 | 0.000 |
| Firmicutes | Clostridia_1 | Clostridiales | Lachnospiraceae | Gut_cluster_2 | 0.179 | 0.018 | 0.147 | 0.030 | 0.005 | 0.000 |
| Firmicutes | Clostridia_1 | Clostridiales | Lachnospiraceae | Gut_cluster_7 | 0.618 | 0.222 | 0.401 | 0.060 | 0.030 | 0.030 |
| Firmicutes | Clostridia_1 | Clostridiales | Lachnospiraceae | Howardella | 0.000 | 0.000 | 0.008 | 0.000 | 0.000 | 0.000 |
| Firmicutes | Clostridia_1 | Clostridiales | Lachnospiraceae | Humibacter | 0.000 | 0.000 | 0.004 | 0.000 | 0.000 | 0.000 |
| Firmicutes | Clostridia_1 | Clostridiales | Lachnospiraceae | Incertae_Sedis | 0.000 | 0.000 | 0.008 | 0.000 | 0.000 | 0.000 |
| Firmicutes | Clostridia_1 | Clostridiales | Lachnospiraceae | Incertae_Sedis_1 | 0.187 | 0.080 | 0.480 | 0.024 | 0.005 | 0.012 |
| Firmicutes | Clostridia_1 | Clostridiales | Lachnospiraceae | Incertae_Sedis_10 | 0.049 | 0.000 | 0.087 | 0.000 | 0.000 | 0.000 |
| Firmicutes | Clostridia_1 | Clostridiales | Lachnospiraceae | Incertae_Sedis_11 | 0.106 | 0.012 | 0.198 | 0.006 | 0.000 | 0.000 |
| Firmicutes | Clostridia_1 | Clostridiales | Lachnospiraceae | Incertae_Sedis_12 | 0.008 | 0.000 | 0.197 | 0.000 | 0.000 | 0.000 |
| Firmicutes | Clostridia_1 | Clostridiales | Lachnospiraceae | Incertae_Sedis_13 | 0.000 | 0.000 | 0.050 | 0.000 | 0.000 | 0.000 |

**Table S9 continued.**

| | | | | | | | | | | |
|---|---|---|---|---|---|---|---|---|---|---|
| Firmicutes | Clostridia_1 | Clostridiales | Lachnospiraceae | Incertae_Sedis_14 | 0.016 | 0.006 | 0.220 | 0.006 | 0.000 | 0.000 |
| Firmicutes | Clostridia_1 | Clostridiales | Lachnospiraceae | Incertae_Sedis_15 | 0.000 | 0.006 | 0.044 | 0.000 | 0.000 | 0.000 |
| Firmicutes | Clostridia_1 | Clostridiales | Lachnospiraceae | Incertae_Sedis_16 | 0.041 | 0.006 | 0.145 | 0.000 | 0.000 | 0.000 |
| Firmicutes | Clostridia_1 | Clostridiales | Lachnospiraceae | Incertae_Sedis_17 | 0.008 | 0.000 | 0.111 | 0.000 | 0.000 | 0.000 |
| Firmicutes | Clostridia_1 | Clostridiales | Lachnospiraceae | Incertae_Sedis_18 | 0.000 | 0.000 | 0.008 | 0.000 | 0.000 | 0.000 |
| Firmicutes | Clostridia_1 | Clostridiales | Lachnospiraceae | Incertae_Sedis_2 | 0.008 | 0.006 | 0.087 | 0.006 | 0.000 | 0.000 |
| Firmicutes | Clostridia_1 | Clostridiales | Lachnospiraceae | Incertae_Sedis_20 | 0.000 | 0.000 | 0.046 | 0.000 | 0.000 | 0.000 |
| Firmicutes | Clostridia_1 | Clostridiales | Lachnospiraceae | Incertae_Sedis_21 | 0.008 | 0.000 | 0.197 | 0.000 | 0.000 | 0.000 |
| Firmicutes | Clostridia_1 | Clostridiales | Lachnospiraceae | Incertae_Sedis_22 | 0.000 | 0.000 | 0.004 | 0.000 | 0.000 | 0.000 |
| Firmicutes | Clostridia_1 | Clostridiales | Lachnospiraceae | Incertae_Sedis_23 | 0.008 | 0.000 | 0.107 | 0.000 | 0.000 | 0.000 |
| Firmicutes | Clostridia_1 | Clostridiales | Lachnospiraceae | Incertae_Sedis_25 | 0.000 | 0.000 | 0.020 | 0.000 | 0.000 | 0.000 |
| Firmicutes | Clostridia_1 | Clostridiales | Lachnospiraceae | Incertae_Sedis_26 | 0.000 | 0.000 | 0.020 | 0.000 | 0.000 | 0.000 |
| Firmicutes | Clostridia_1 | Clostridiales | Lachnospiraceae | Incertae_Sedis_27 | 0.000 | 0.006 | 0.123 | 0.000 | 0.000 | 0.000 |
| Firmicutes | Clostridia_1 | Clostridiales | Lachnospiraceae | Incertae_Sedis_28 | 0.000 | 0.000 | 0.062 | 0.000 | 0.000 | 0.000 |
| Firmicutes | Clostridia_1 | Clostridiales | Lachnospiraceae | Incertae_Sedis_3 | 0.008 | 0.000 | 0.123 | 0.030 | 0.010 | 0.024 |
| Firmicutes | Clostridia_1 | Clostridiales | Lachnospiraceae | Incertae_Sedis_31 | 0.000 | 0.006 | 0.018 | 0.006 | 0.000 | 0.000 |
| Firmicutes | Clostridia_1 | Clostridiales | Lachnospiraceae | Incertae_Sedis_32 | 0.008 | 0.000 | 0.036 | 0.000 | 0.000 | 0.000 |
| Firmicutes | Clostridia_1 | Clostridiales | Lachnospiraceae | Incertae_Sedis_33 | 0.000 | 0.000 | 0.022 | 0.000 | 0.000 | 0.000 |
| Firmicutes | Clostridia_1 | Clostridiales | Lachnospiraceae | Incertae_Sedis_34 | 0.041 | 0.006 | 0.036 | 0.006 | 0.000 | 0.000 |
| Firmicutes | Clostridia_1 | Clostridiales | Lachnospiraceae | Incertae_Sedis_35 | 0.000 | 0.000 | 0.008 | 0.000 | 0.000 | 0.000 |
| Firmicutes | Clostridia_1 | Clostridiales | Lachnospiraceae | Incertae_Sedis_4 | 0.008 | 0.000 | 0.016 | 0.000 | 0.000 | 0.000 |
| Firmicutes | Clostridia_1 | Clostridiales | Lachnospiraceae | Incertae_Sedis_6 | 0.016 | 0.006 | 0.232 | 0.006 | 0.000 | 0.000 |
| Firmicutes | Clostridia_1 | Clostridiales | Lachnospiraceae | Incertae_Sedis_9 | 0.033 | 0.006 | 0.113 | 0.000 | 0.000 | 0.000 |
| Firmicutes | Clostridia_1 | Clostridiales | Lachnospiraceae | Inhella | 0.000 | 0.000 | 0.006 | 0.000 | 0.000 | 0.000 |
| Firmicutes | Clostridia_1 | Clostridiales | Lachnospiraceae | Lachnobacterium | 0.000 | 0.000 | 0.004 | 0.000 | 0.000 | 0.000 |
| Firmicutes | Clostridia_1 | Clostridiales | Lachnospiraceae | Moryella | 0.000 | 0.006 | 0.028 | 0.000 | 0.000 | 0.000 |
| Firmicutes | Clostridia_1 | Clostridiales | Lachnospiraceae | Orienta | 0.000 | 0.000 | 0.004 | 0.000 | 0.000 | 0.000 |
| Firmicutes | Clostridia_1 | Clostridiales | Lachnospiraceae | Parvibaculum | 0.000 | 0.000 | 0.008 | 0.000 | 0.000 | 0.000 |
| Firmicutes | Clostridia_1 | Clostridiales | Lachnospiraceae | Robinsoniella_Insects | 0.000 | 0.000 | 0.010 | 0.000 | 0.000 | 0.000 |
| Firmicutes | Clostridia_1 | Clostridiales | Lachnospiraceae | Roseburia_2 | 0.000 | 0.000 | 0.022 | 0.000 | 0.000 | 0.000 |
| Firmicutes | Clostridia_1 | Clostridiales | Lachnospiraceae | Roseburia_3 | 0.000 | 0.000 | 0.006 | 0.000 | 0.000 | 0.000 |
| Firmicutes | Clostridia_1 | Clostridiales | Lachnospiraceae | Roseiflexus | 0.008 | 0.000 | 0.195 | 0.000 | 0.000 | 0.000 |
| Firmicutes | Clostridia_1 | Clostridiales | Lachnospiraceae | Syntrophomonas_2 | 0.000 | 0.000 | 0.006 | 0.000 | 0.000 | 0.000 |
| Firmicutes | Clostridia_1 | Clostridiales | Lachnospiraceae | Uncultured_15 | 0.000 | 0.000 | 0.101 | 0.000 | 0.000 | 0.000 |
| Firmicutes | Clostridia_1 | Clostridiales | Lachnospiraceae | Uncultured_23 | 0.016 | 0.000 | 0.292 | 0.000 | 0.000 | 0.000 |
| Firmicutes | Clostridia_1 | Clostridiales | Lachnospiraceae | Uncultured_32 | 0.000 | 0.000 | 0.034 | 0.000 | 0.000 | 0.000 |
| Firmicutes | Clostridia_1 | Clostridiales | Lachnospiraceae | Uncultured_38 | 0.000 | 0.000 | 0.004 | 0.000 | 0.000 | 0.000 |
| Firmicutes | Clostridia_1 | Clostridiales | Lachnospiraceae | Uncultured_4 | 0.073 | 0.012 | 0.570 | 0.006 | 0.005 | 0.006 |
| Firmicutes | Clostridia_1 | Clostridiales | Lachnospiraceae | Uncultured_40 | 0.000 | 0.000 | 0.004 | 0.000 | 0.000 | 0.000 |
| Firmicutes | Clostridia_1 | Clostridiales | Lachnospiraceae | Uncultured_43 | 0.000 | 0.000 | 0.079 | 0.000 | 0.000 | 0.000 |

**Table S9 continued.**

| | | | | | | | | | | |
|---|---|---|---|---|---|---|---|---|---|---|
| Firmicutes | Clostridia_1 | Clostridiales | Lachnospiraceae | Uncultured_44 | 0.000 | 0.006 | 0.081 | 0.006 | 0.005 | 0.000 |
| Firmicutes | Clostridia_1 | Clostridiales | Lachnospiraceae | Uncultured_45 | 0.000 | 0.000 | 0.008 | 0.000 | 0.000 | 0.000 |
| Firmicutes | Clostridia_1 | Clostridiales | Lachnospiraceae | Uncultured_48 | 0.000 | 0.000 | 0.010 | 0.000 | 0.000 | 0.000 |
| Firmicutes | Clostridia_1 | Clostridiales | Lachnospiraceae | Uncultured_49 | 0.000 | 0.000 | 0.022 | 0.000 | 0.000 | 0.000 |
| Firmicutes | Clostridia_1 | Clostridiales | Lachnospiraceae | Uncultured_5 | 0.317 | 0.099 | 0.572 | 0.036 | 0.010 | 0.000 |
| Firmicutes | Clostridia_1 | Clostridiales | Lachnospiraceae | Uncultured_50 | 0.000 | 0.000 | 0.040 | 0.000 | 0.000 | 0.000 |
| Firmicutes | Clostridia_1 | Clostridiales | Lachnospiraceae | Uncultured_53 | 0.000 | 0.000 | 0.006 | 0.000 | 0.000 | 0.000 |
| Firmicutes | Clostridia_1 | Clostridiales | Lachnospiraceae | Uncultured_54 | 0.374 | 0.062 | 0.008 | 0.024 | 0.000 | 0.000 |
| Firmicutes | Clostridia_1 | Clostridiales | Lachnospiraceae | Uncultured_55 | 0.016 | 0.000 | 0.014 | 0.000 | 0.000 | 0.000 |
| Firmicutes | Clostridia_1 | Clostridiales | Lachnospiraceae | Uncultured_56 | 0.000 | 0.000 | 0.004 | 0.000 | 0.000 | 0.000 |
| Firmicutes | Clostridia_1 | Clostridiales | Lachnospiraceae | Uncultured_57 | 0.008 | 0.000 | 0.008 | 0.000 | 0.000 | 0.000 |
| Firmicutes | Clostridia_1 | Clostridiales | Lachnospiraceae | Uncultured_58 | 0.000 | 0.000 | 0.008 | 0.000 | 0.000 | 0.000 |
| Firmicutes | Clostridia_1 | Clostridiales | Lachnospiraceae | Uncultured_6 | 0.195 | 0.043 | 0.383 | 0.024 | 0.000 | 0.000 |
| Firmicutes | Clostridia_1 | Clostridiales | Lachnospiraceae | Uncultured_60 | 0.008 | 0.000 | 0.024 | 0.000 | 0.000 | 0.000 |
| Firmicutes | Clostridia_1 | Clostridiales | Lachnospiraceae | Uncultured_64 | 0.008 | 0.006 | 0.028 | 0.000 | 0.000 | 0.000 |
| Firmicutes | Clostridia_1 | Clostridiales | Lachnospiraceae | Uncultured_66 | 0.000 | 0.000 | 0.004 | 0.000 | 0.000 | 0.000 |
| Firmicutes | Clostridia_1 | Clostridiales | Lachnospiraceae | Uncultured_8 | 0.049 | 0.018 | 0.246 | 0.018 | 0.000 | 0.000 |
| Firmicutes | Clostridia_2 | Clostridiales | Lachnospiraceae | Butyrivibrio-Pseudobutyrivibrio | 0.000 | 0.000 | 0.040 | 0.000 | 0.000 | 0.000 |
| Firmicutes | Clostridia_2 | Clostridiales | Lachnospiraceae | Lachnospira | 0.000 | 0.000 | 0.050 | 0.000 | 0.000 | 0.000 |
| Firmicutes | Clostridia_2 | Clostridiales | Lachnospiraceae | Lacibacter | 0.000 | 0.000 | 0.004 | 0.000 | 0.000 | 0.000 |
| Firmicutes | Clostridia_3 | Clostridiales | Lachnospiraceae | Anaerocolumna | 0.008 | 0.000 | 0.000 | 0.000 | 0.000 | 0.000 |
| Firmicutes | Bacilli | Lactobacillales | Lactobacillaceae | Lactobacillus_1 | 0.008 | 0.000 | 0.490 | 0.000 | 0.000 | 0.000 |
| Firmicutes | Bacilli | Lactobacillales | Lactobacillaceae | Lactobacillus_2 | 0.000 | 0.000 | 0.046 | 0.000 | 0.000 | 0.000 |
| Firmicutes | Bacilli | Lactobacillales | Lactobacillaceae | Lactobacillus_3 | 0.000 | 0.000 | 0.054 | 0.000 | 0.000 | 0.000 |
| Firmicutes | Bacilli | Lactobacillales | Lactobacillaceae | Lactobacillus_4 | 0.000 | 0.000 | 0.062 | 0.000 | 0.000 | 0.000 |
| Firmicutes | Bacilli | Lactobacillales | Lactobacillaceae | Lactobacillus_5 | 0.000 | 0.000 | 0.016 | 0.000 | 0.000 | 0.000 |
| Firmicutes | Bacilli | Lactobacillales | Lactobacillaceae | Lactobacillus_6 | 0.000 | 0.000 | 0.004 | 0.000 | 0.000 | 0.000 |
| Firmicutes | Bacilli | Lactobacillales | Lactobacillaceae | Lactobacillus_8 | 0.000 | 0.000 | 0.121 | 0.000 | 0.000 | 0.000 |
| Firmicutes | Bacilli | Lactobacillales | Lactobacillaceae | Lactobacillus_9 | 0.000 | 0.000 | 0.026 | 0.000 | 0.000 | 0.000 |
| Firmicutes | Bacilli | Lactobacillales | Lactobacillaceae | Lactobacillus_sp_a | 0.000 | 0.000 | 0.004 | 0.000 | 0.000 | 0.000 |
| Firmicutes | Bacilli | Lactobacillales | Lactobacillaceae | Lactococcus | 0.024 | 0.000 | 0.000 | 0.000 | 0.000 | 0.000 |
| Firmicutes | Bacilli | Lactobacillales | Lactobacillaceae | Parapedobacter | 0.000 | 0.000 | 0.006 | 0.000 | 0.000 | 0.000 |
| Firmicutes | Bacilli | Lactobacillales | Lactobacillaceae | Pediococcus | 0.000 | 0.000 | 0.022 | 0.000 | 0.000 | 0.000 |
| Firmicutes | Bacilli | Lactobacillales | Lactobacillaceae | Pedobacter | 0.016 | 0.074 | 0.000 | 0.000 | 0.000 | 0.000 |
| Firmicutes | Bacilli | Lactobacillales | Leuconostocaceae | hoa5-07d05_gut_group | 0.000 | 0.000 | 0.016 | 0.000 | 0.000 | 0.000 |
| Firmicutes | Bacilli | Lactobacillales | Leuconostocaceae | Lewinella | 0.000 | 0.000 | 0.026 | 0.000 | 0.000 | 0.006 |
| Firmicutes | Bacilli | Lactobacillales | Leuconostocaceae | Oenococcus | 0.000 | 0.000 | 0.006 | 0.000 | 0.000 | 0.000 |
| Firmicutes | Bacilli | Lactobacillales | Leuconostocaceae | Oerskovia | 0.000 | 0.000 | 0.010 | 0.000 | 0.000 | 0.000 |
| Firmicutes | Bacilli | Lactobacillales | Leuconostocaceae | Weissella_1 | 0.000 | 0.000 | 0.036 | 0.000 | 0.000 | 0.000 |
| Firmicutes | Bacilli | Lactobacillales | Leuconostocaceae | Weissella_2 | 0.000 | 0.000 | 0.004 | 0.000 | 0.000 | 0.000 |

**Table S9 continued.**

| | | | | | | | | | | |
|---|---|---|---|---|---|---|---|---|---|---|
| Firmicutes | Bacilli | Bacillales | Listeriaceae | Brochothrix | 0.000 | 0.000 | 0.006 | 0.000 | 0.000 | 0.000 |
| Firmicutes | Clostridia_3 | Natranaerobiales | Natranaerobiaceae | Natranaerobius | 0.000 | 0.000 | 0.006 | 0.000 | 0.000 | 0.000 |
| Firmicutes | Bacilli | Bacillales | Paenibacillaceae | Cohnella | 0.000 | 0.000 | 0.022 | 0.024 | 0.184 | 0.375 |
| Firmicutes | Bacilli | Bacillales | Paenibacillaceae | Fontibacillus | 0.000 | 0.000 | 0.000 | 0.000 | 0.025 | 0.048 |
| Firmicutes | Bacilli | Bacillales | Paenibacillaceae | Gorillibacterium | 0.024 | 0.018 | 0.012 | 0.000 | 0.000 | 0.000 |
| Firmicutes | Bacilli | Bacillales | Paenibacillaceae_2 | Brevibacillus | 0.000 | 0.000 | 0.058 | 0.012 | 0.015 | 0.036 |
| Firmicutes | Bacilli | Bacillales | Paenibacillaceae_2 | Paenibacillus_1 | 0.016 | 0.000 | 0.032 | 0.036 | 0.145 | 0.502 |
| Firmicutes | Bacilli | Bacillales | Paenibacillaceae_2 | Paenibacillus_10 | 0.000 | 0.000 | 0.004 | 0.006 | 0.000 | 0.006 |
| Firmicutes | Bacilli | Bacillales | Paenibacillaceae_2 | Paenibacillus_12 | 0.024 | 0.000 | 0.046 | 0.312 | 0.075 | 0.260 |
| Firmicutes | Bacilli | Bacillales | Paenibacillaceae_2 | Paenibacillus_13 | 0.008 | 0.000 | 0.079 | 0.048 | 0.045 | 0.188 |
| Firmicutes | Bacilli | Bacillales | Paenibacillaceae_2 | Paenibacillus_14 | 0.000 | 0.000 | 0.024 | 0.006 | 0.010 | 0.000 |
| Firmicutes | Bacilli | Bacillales | Paenibacillaceae_2 | Paenibacillus_15 | 0.000 | 0.000 | 0.004 | 0.000 | 0.000 | 0.006 |
| Firmicutes | Bacilli | Bacillales | Paenibacillaceae_2 | Paenibacillus_16 | 0.000 | 0.000 | 0.012 | 0.000 | 0.000 | 0.012 |
| Firmicutes | Bacilli | Bacillales | Paenibacillaceae_2 | Paenibacillus_17 | 0.008 | 0.000 | 0.079 | 0.060 | 0.075 | 0.242 |
| Firmicutes | Bacilli | Bacillales | Paenibacillaceae_2 | Paenibacillus_18 | 0.000 | 0.000 | 0.012 | 0.000 | 0.030 | 0.042 |
| Firmicutes | Bacilli | Bacillales | Paenibacillaceae_2 | Paenibacillus_19 | 0.000 | 0.006 | 0.014 | 0.000 | 0.000 | 0.006 |
| Firmicutes | Bacilli | Bacillales | Paenibacillaceae_2 | Paenibacillus_2 | 0.000 | 0.000 | 0.008 | 0.000 | 0.005 | 0.012 |
| Firmicutes | Bacilli | Bacillales | Paenibacillaceae_2 | Paenibacillus_20 | 0.000 | 0.000 | 0.004 | 0.000 | 0.000 | 0.000 |
| Firmicutes | Bacilli | Bacillales | Paenibacillaceae_2 | Paenibacillus_4 | 0.000 | 0.000 | 0.006 | 0.000 | 0.000 | 0.000 |
| Firmicutes | Bacilli | Bacillales | Paenibacillaceae_2 | Paenibacillus_6 | 0.000 | 0.000 | 0.004 | 0.000 | 0.005 | 0.012 |
| Firmicutes | Bacilli | Bacillales | Paenibacillaceae_2 | Paenibacillus_7 | 0.000 | 0.000 | 0.008 | 0.000 | 0.000 | 0.000 |
| Firmicutes | Bacilli | Bacillales | Paenibacillaceae_2 | Paenibacillus_8 | 0.000 | 0.000 | 0.010 | 0.006 | 0.000 | 0.036 |
| Firmicutes | Bacilli | Bacillales | Paenibacillaceae_2 | Paenibacillus_9 | 0.000 | 0.000 | 0.004 | 0.000 | 0.000 | 0.000 |
| Firmicutes | Bacilli | Bacillales | Paenibacillaceae_2 | Paenochrobactrum | 0.000 | 0.000 | 0.002 | 0.000 | 0.000 | 0.000 |
| Firmicutes | Bacilli | Bacillales_2 | Pasteuriaceae | Pasteuria | 0.000 | 0.000 | 0.004 | 0.000 | 0.000 | 0.000 |
| Firmicutes | Bacilli | Bacillales_2 | Pasteuriaceae | Patulibacter | 0.065 | 0.006 | 0.016 | 0.012 | 0.000 | 0.006 |
| Firmicutes | Clostridia | Clostridiales | Peptococcaceae | Desulfotomaculum_7 | 0.000 | 0.000 | 0.024 | 0.000 | 0.000 | 0.000 |
| Firmicutes | Clostridia_2 | Clostridiales | Peptococcaceae | Desulfitibacter | 0.000 | 0.000 | 0.004 | 0.000 | 0.000 | 0.000 |
| Firmicutes | Clostridia_3 | Clostridiales | Peptococcaceae | Desulfitobacterium | 0.000 | 0.006 | 0.024 | 0.000 | 0.000 | 0.000 |
| Firmicutes | Clostridia_2 | Clostridiales_1 | Peptococcaceae | Desulfosporosinus | 0.000 | 0.000 | 0.020 | 0.000 | 0.000 | 0.000 |
| Firmicutes | Clostridia_2 | Clostridiales_1 | Peptococcaceae_1 | Peptococcus | 0.000 | 0.000 | 0.016 | 0.000 | 0.000 | 0.000 |
| Firmicutes | Clostridia_2 | Clostridiales_1 | Peptococcaceae_1 | Thermoanaerobacter | 0.000 | 0.000 | 0.016 | 0.000 | 0.000 | 0.000 |
| Firmicutes | Clostridia_2 | Clostridiales_1 | Peptococcaceae_2 | Dehalobacter | 0.000 | 0.000 | 0.004 | 0.000 | 0.000 | 0.000 |
| Firmicutes | Clostridia_2 | Clostridiales_1 | Peptococcaceae_3 | Cryptanaerobacter | 0.000 | 0.000 | 0.004 | 0.000 | 0.000 | 0.000 |
| Firmicutes | Clostridia_2 | Clostridiales_1 | Peptococcaceae_3 | Desulfotomaculum_1 | 0.000 | 0.000 | 0.028 | 0.000 | 0.000 | 0.000 |
| Firmicutes | Clostridia_2 | Clostridiales_1 | Peptococcaceae_3 | Desulfotomaculum_4 | 0.000 | 0.000 | 0.008 | 0.000 | 0.000 | 0.000 |
| Firmicutes | Clostridia_2 | Clostridiales_1 | Peptococcaceae_3 | Desulfotomaculum_5 | 0.000 | 0.000 | 0.006 | 0.000 | 0.000 | 0.000 |
| Firmicutes | Clostridia_2 | Clostridiales_1 | Peptococcaceae_3 | Desulfotomaculum_6 | 0.000 | 0.000 | 0.004 | 0.000 | 0.000 | 0.000 |
| Firmicutes | Clostridia_2 | Clostridiales_1 | Peptococcaceae_3 | Desulfurispora | 0.000 | 0.000 | 0.006 | 0.000 | 0.000 | 0.000 |
| Firmicutes | Clostridia_2 | Clostridiales_1 | Peptococcaceae_3 | Staphylococcus | 0.000 | 0.012 | 0.000 | 0.012 | 0.010 | 0.000 |

**Table S9 continued.**

| | | | | | | | | | | |
|---|---|---|---|---|---|---|---|---|---|---|
| Firmicutes | Tissierellia | Tissierellales | Peptoniphilaceae | Helicobacter | 0.000 | 0.000 | 0.123 | 0.000 | 0.000 | 0.000 |
| Firmicutes | Clostridia_1 | Clostridiales | Peptostreptococcaceae | Frigovirgula_patagoniensis | 0.000 | 0.000 | 0.004 | 0.000 | 0.000 | 0.000 |
| Firmicutes | Clostridia_1 | Clostridiales | Peptostreptococcaceae | Peptostreptococcus | 0.000 | 0.000 | 0.016 | 0.000 | 0.000 | 0.000 |
| Firmicutes | Clostridia_1 | Clostridiales | Peptostreptococcaceae | Sporichthya | 0.000 | 0.000 | 0.008 | 0.000 | 0.000 | 0.000 |
| Firmicutes | Clostridia_2 | Clostridiales | Peptostreptococcaceae | Filifactor | 0.000 | 0.000 | 0.006 | 0.000 | 0.000 | 0.000 |
| Firmicutes | Clostridia_3 | Clostridiales | Peptostreptococcaceae | Peptoniphilus | 0.000 | 0.000 | 0.058 | 0.000 | 0.000 | 0.000 |
| Firmicutes | Bacilli | Bacillales | Planococcaceae | Sporosarcina_4 | 0.000 | 0.000 | 0.010 | 0.000 | 0.000 | 0.000 |
| Firmicutes | Bacilli | Bacillales | Planococcaceae | Sporosarcina_5 | 0.000 | 0.000 | 0.006 | 0.000 | 0.000 | 0.000 |
| Firmicutes | Bacilli | Bacillales | Planococcaceae | Vagococcus | 0.008 | 0.000 | 0.014 | 0.000 | 0.005 | 0.000 |
| Firmicutes | Bacilli | Bacillales | Planococcaceae | Viridibacillus | 0.000 | 0.000 | 0.002 | 0.018 | 0.010 | 0.000 |
| Firmicutes | Bacilli | Bacillales | Planococcaceae_1 | Jeotgalibacillus | 0.000 | 0.000 | 0.004 | 0.006 | 0.000 | 0.000 |
| Firmicutes | Bacilli | Bacillales | Planococcaceae_1 | Marinibacillus | 0.000 | 0.000 | 0.004 | 0.000 | 0.000 | 0.000 |
| Firmicutes | Bacilli | Bacillales | Planococcaceae_2 | Caryophanon | 0.000 | 0.000 | 0.006 | 0.000 | 0.000 | 0.000 |
| Firmicutes | Bacilli | Bacillales | Planococcaceae_2 | Incertae_Sedis_30 | 0.008 | 0.000 | 0.020 | 0.000 | 0.000 | 0.000 |
| Firmicutes | Bacilli | Bacillales | Planococcaceae_2 | Incertae_Sedis_7 | 0.000 | 0.000 | 0.066 | 0.000 | 0.000 | 0.000 |
| Firmicutes | Bacilli | Bacillales | Planococcaceae_2 | Lysobacter | 0.114 | 0.012 | 0.000 | 0.012 | 0.000 | 0.000 |
| Firmicutes | Bacilli | Bacillales | Planococcaceae_2 | Planococcus_1 | 0.000 | 0.018 | 0.060 | 0.012 | 0.005 | 0.000 |
| Firmicutes | Bacilli | Bacillales | Planococcaceae_2 | Planococcus_2 | 0.000 | 0.000 | 0.006 | 0.000 | 0.000 | 0.006 |
| Firmicutes | Bacilli | Bacillales | Planococcaceae_2 | Sporosarcina_1 | 0.000 | 0.000 | 0.008 | 0.000 | 0.000 | 0.000 |
| Firmicutes | Bacilli | Bacillales | Planococcaceae_2 | Sporosarcina_2 | 0.000 | 0.000 | 0.004 | 0.000 | 0.000 | 0.000 |
| Firmicutes | Bacilli | Bacillales | Planococcaceae_3 | Planomicrobium | 0.000 | 0.000 | 0.000 | 0.006 | 0.000 | 0.000 |
| Firmicutes | Clostridia | Clostridiales | Ruminococcaceae | Acetivibrio | 0.000 | 0.000 | 0.004 | 0.000 | 0.000 | 0.000 |
| Firmicutes | Clostridia | Clostridiales | Ruminococcaceae | Helcococcus | 0.000 | 0.000 | 0.018 | 0.000 | 0.000 | 0.000 |
| Firmicutes | Clostridia | CLostridiales | Ruminococcaceae | Pseudolabrys | 0.016 | 0.000 | 0.030 | 0.006 | 0.000 | 0.000 |
| Firmicutes | Clostridia | Clostridiales | Ruminococcaceae | Ruminococcus_2 | 0.000 | 0.000 | 0.064 | 0.000 | 0.000 | 0.000 |
| Firmicutes | Clostridia | Clostridiales | Ruminococcaceae | Saccharibacillus | 0.000 | 0.000 | 0.004 | 0.000 | 0.000 | 0.000 |
| Firmicutes | Clostridia_1 | Clostridiales | Ruminococcaceae | Acetanaerobacterium | 0.000 | 0.000 | 0.006 | 0.000 | 0.000 | 0.000 |
| Firmicutes | Clostridia_1 | Clostridiales | Ruminococcaceae | Anaerofilum | 0.000 | 0.006 | 0.010 | 0.000 | 0.000 | 0.000 |
| Firmicutes | Clostridia_1 | Clostridiales | Ruminococcaceae | Anaerotruncus | 0.089 | 0.080 | 0.175 | 0.018 | 0.000 | 0.000 |
| Firmicutes | Clostridia_1 | Clostridiales | Ruminococcaceae | Faecalibacterium | 0.008 | 0.000 | 0.381 | 0.000 | 0.000 | 0.000 |
| Firmicutes | Clostridia_1 | Clostridiales | Ruminococcaceae | Fastidiosipila | 0.008 | 0.006 | 0.048 | 0.000 | 0.000 | 0.000 |
| Firmicutes | Clostridia_1 | Clostridiales | Ruminococcaceae | Gut_cluster_11 | 0.016 | 0.012 | 0.343 | 0.000 | 0.000 | 0.000 |
| Firmicutes | Clostridia_1 | Clostridiales | Ruminococcaceae | Gut_cluster_5 | 0.268 | 0.154 | 0.288 | 0.054 | 0.010 | 0.000 |
| Firmicutes | Clostridia_1 | Clostridiales | Ruminococcaceae | Gut_cluster_6 | 0.277 | 0.105 | 0.691 | 0.018 | 0.005 | 0.000 |
| Firmicutes | Clostridia_1 | Clostridiales | Ruminococcaceae | Gut_cluster_8 | 0.813 | 0.437 | 0.318 | 0.168 | 0.025 | 0.006 |
| Firmicutes | Clostridia_1 | Clostridiales | Ruminococcaceae | Gut_cluster_9 | 0.098 | 0.018 | 0.183 | 0.024 | 0.000 | 0.000 |
| Firmicutes | Clostridia_1 | Clostridiales | Ruminococcaceae | Hydrogenophaga | 0.000 | 0.006 | 0.000 | 0.000 | 0.000 | 0.000 |
| Firmicutes | Clostridia_1 | Clostridiales | Ruminococcaceae | Incertae_Sedis_19 | 0.016 | 0.000 | 0.165 | 0.000 | 0.000 | 0.000 |
| Firmicutes | Clostridia_1 | Clostridiales | Ruminococcaceae | Incertae_Sedis_5 | 0.033 | 0.006 | 0.320 | 0.006 | 0.000 | 0.000 |
| Firmicutes | Clostridia_1 | Clostridiales | Ruminococcaceae | Incertae_Sedis_8 | 0.203 | 0.006 | 0.177 | 0.000 | 0.000 | 0.000 |

**Table S9 continued.**

| | | | | | | | | | | |
|---|---|---|---|---|---|---|---|---|---|---|
| Firmicutes | Clostridia_1 | Clostridiales | Ruminococcaceae | Oscillochloris | 0.000 | 0.000 | 0.006 | 0.000 | 0.000 | 0.000 |
| Firmicutes | Clostridia_1 | Clostridiales | Ruminococcaceae | Ottowia | 0.000 | 0.000 | 0.050 | 0.000 | 0.000 | 0.000 |
| Firmicutes | Clostridia_1 | Clostridiales | Ruminococcaceae | Parabacteroides | 0.016 | 0.000 | 0.139 | 0.006 | 0.005 | 0.000 |
| Firmicutes | Clostridia_1 | Clostridiales | Ruminococcaceae | Phormidium | 0.000 | 0.000 | 0.030 | 0.000 | 0.000 | 0.000 |
| Firmicutes | Clostridia_1 | Clostridiales | Ruminococcaceae | Termite_cluster_2 | 0.561 | 0.277 | 0.099 | 0.072 | 0.000 | 0.006 |
| Firmicutes | Clostridia_1 | Clostridiales | Ruminococcaceae | Uncultured_14 | 0.000 | 0.000 | 0.079 | 0.000 | 0.000 | 0.000 |
| Firmicutes | Clostridia_1 | Clostridiales | Ruminococcaceae | Uncultured_20 | 0.057 | 0.012 | 0.443 | 0.006 | 0.000 | 0.000 |
| Firmicutes | Clostridia_1 | Clostridiales | Ruminococcaceae | Uncultured_22 | 0.065 | 0.000 | 0.230 | 0.006 | 0.000 | 0.000 |
| Firmicutes | Clostridia_1 | Clostridiales | Ruminococcaceae | Uncultured_26 | 0.041 | 0.006 | 0.625 | 0.000 | 0.000 | 0.006 |
| Firmicutes | Clostridia_1 | Clostridiales | Ruminococcaceae | Uncultured_29 | 0.073 | 0.006 | 0.175 | 0.030 | 0.000 | 0.000 |
| Firmicutes | Clostridia_1 | Clostridiales | Ruminococcaceae | Uncultured_3 | 0.171 | 0.018 | 0.963 | 0.036 | 0.015 | 0.030 |
| Firmicutes | Clostridia_1 | Clostridiales | Ruminococcaceae | Uncultured_35 | 0.016 | 0.000 | 0.060 | 0.000 | 0.005 | 0.000 |
| Firmicutes | Clostridia_1 | Clostridiales | Ruminococcaceae | Uncultured_37 | 0.000 | 0.000 | 0.008 | 0.000 | 0.000 | 0.000 |
| Firmicutes | Clostridia_1 | Clostridiales | Ruminococcaceae | Uncultured_39 | 0.000 | 0.000 | 0.006 | 0.000 | 0.000 | 0.000 |
| Firmicutes | Negativicutes | Selenomonadales | Selenomonadaceae | Mixed_environment_cluster | 0.008 | 0.000 | 0.111 | 0.000 | 0.000 | 0.000 |
| Firmicutes | Negativicutes | Selenomonadales | Selenomonadaceae | Serinicoccus | 0.000 | 0.000 | 0.004 | 0.000 | 0.000 | 0.000 |
| Firmicutes | Mollicutes | Entomoplasmatales | Spiroplasmataceae | Spirosoma | 0.000 | 0.000 | 0.040 | 0.000 | 0.000 | 0.000 |
| Firmicutes | Mollicutes | Entomoplasmatales | Spiroplasmataceae | Spongiimicrobium | 0.000 | 0.000 | 0.004 | 0.000 | 0.000 | 0.000 |
| Firmicutes | Bacilli | Bacillales | Sporolactobacillaceae | Sporolactobacillus | 0.000 | 0.000 | 0.012 | 0.000 | 0.000 | 0.000 |
| Firmicutes | Negativicutes | Selenomonadales | Sporomusaceae | Amycolatopsis | 0.000 | 0.000 | 0.079 | 0.000 | 0.000 | 0.000 |
| Firmicutes | Negativicutes | Selenomonadales | Sporomusaceae | Anaerospora | 0.000 | 0.000 | 0.004 | 0.000 | 0.000 | 0.000 |
| Firmicutes | Negativicutes | Selenomonadales | Sporomusaceae | Propionivibrio | 0.033 | 0.031 | 0.032 | 0.012 | 0.000 | 0.012 |
| Firmicutes | Bacilli | Bacillales | Staphylococcaceae | Macrococcus_1 | 0.000 | 0.000 | 0.006 | 0.000 | 0.000 | 0.000 |
| Firmicutes | Bacilli | Bacillales | Staphylococcaceae | Salinicoccus | 0.000 | 0.000 | 0.022 | 0.000 | 0.000 | 0.000 |
| Firmicutes | Bacilli | Bacillales | Staphylococcaceae | Staphylococcus_2 | 0.000 | 0.000 | 0.020 | 0.000 | 0.010 | 0.000 |
| Firmicutes | Bacilli | Bacillales | Staphylococcaceae | Stappia_2 | 0.000 | 0.000 | 0.006 | 0.000 | 0.000 | 0.000 |
| Firmicutes | Bacilli | Lactobacillales | Streptococcaceae | Lactococcus_1 | 0.065 | 1.201 | 0.060 | 0.012 | 0.005 | 0.000 |
| Firmicutes | Bacilli | Lactobacillales | Streptococcaceae | Lactococcus_2 | 0.000 | 0.000 | 0.008 | 0.000 | 0.000 | 0.000 |
| Firmicutes | Bacilli | Lactobacillales | Streptococcaceae | Lactococcus_3 | 0.000 | 0.000 | 0.020 | 0.000 | 0.000 | 0.000 |
| Firmicutes | Bacilli | Lactobacillales | Streptococcaceae | Lactonifactor | 0.008 | 0.000 | 0.000 | 0.000 | 0.000 | 0.000 |
| Firmicutes | Bacilli | Lactobacillales | Streptococcaceae | Streptococcus | 0.228 | 0.037 | 0.393 | 0.054 | 0.005 | 0.000 |
| Firmicutes | Bacilli | Lactobacillales | Streptococcaceae | Streptomyces_1 | 0.008 | 0.006 | 0.564 | 0.036 | 0.229 | 0.188 |
| Firmicutes | Clostridia | Clostridiales | Syntrophomonadaceae | Syntrophomonas_3 | 0.000 | 0.000 | 0.006 | 0.000 | 0.000 | 0.000 |
| Firmicutes | Clostridia_2 | Clostridiales | Syntrophomonadaceae | Pelotomaculum_1 | 0.000 | 0.000 | 0.004 | 0.000 | 0.000 | 0.000 |
| Firmicutes | Clostridia_2 | Clostridiales | Syntrophomonadaceae | Pelotomaculum_2 | 0.000 | 0.000 | 0.004 | 0.000 | 0.000 | 0.000 |
| Firmicutes | Clostridia_2 | Clostridiales | Syntrophomonadaceae | Pelotomaculum_3 | 0.000 | 0.000 | 0.010 | 0.000 | 0.000 | 0.000 |
| Firmicutes | Clostridia_2 | Clostridiales | Syntrophomonadaceae | Syntrophomonas_1 | 0.000 | 0.000 | 0.024 | 0.000 | 0.000 | 0.000 |
| Firmicutes | Clostridia_2 | Clostridiales | Syntrophomonadaceae | Syntrophomonas_4 | 0.000 | 0.000 | 0.004 | 0.000 | 0.000 | 0.000 |
| Firmicutes | Clostridia_2 | Clostridiales | Syntrophomonadaceae | Syntrophus | 0.024 | 0.000 | 0.018 | 0.000 | 0.000 | 0.000 |
| Firmicutes | Clostridia_2 | Clostridiales | Syntrophomonadaceae | Thermovirga | 0.008 | 0.000 | 0.062 | 0.000 | 0.000 | 0.000 |

**Table S9 continued.**

| | | | | | | | | | | |
|---|---|---|---|---|---|---|---|---|---|---|
| Firmicutes | Clostridia_3 | Clostridiales | Syntrophomonadaceae | Dethiobacter | 0.000 | 0.000 | 0.006 | 0.000 | 0.000 | 0.000 |
| Firmicutes | Bacilli | Bacillales | Thermoactinomycetaceae | Marinobacter | 0.008 | 0.000 | 0.107 | 0.000 | 0.000 | 0.000 |
| Firmicutes | Bacilli | Bacillales | Thermoactinomycetaceae | Thermoanaerobacter_1 | 0.000 | 0.000 | 0.044 | 0.000 | 0.000 | 0.000 |
| Firmicutes | Bacilli | Bacillales_2 | Thermoactinomycetaceae | Desmospora | 0.000 | 0.000 | 0.004 | 0.000 | 0.000 | 0.000 |
| Firmicutes | Bacilli | Bacillales_2 | Thermoactinomycetaceae | Planifilum | 0.000 | 0.000 | 0.006 | 0.000 | 0.000 | 0.000 |
| Firmicutes | Bacilli | Bacillales_2 | Thermoactinomycetaceae | Shimazuella | 0.000 | 0.000 | 0.006 | 0.000 | 0.000 | 0.006 |
| Firmicutes | Bacilli | Bacillales_2 | Thermoactinomycetaceae | Shimwellia | 0.008 | 0.025 | 0.016 | 0.030 | 0.040 | 0.006 |
| Firmicutes | Closridia | Thermoanaerobacterales | Thermoanaerobacteraceae | Gallibacterium | 0.000 | 0.000 | 0.016 | 0.000 | 0.000 | 0.000 |
| Firmicutes | Clostridia | Thermoanaerobacterales | Thermoanaerobacteraceae | Thermobacillus | 0.000 | 0.000 | 0.008 | 0.000 | 0.000 | 0.000 |
| Firmicutes | Clostridia_3 | Thermoanaerobacterales_3 | Thermoanaerobacteraceae | Caldanaerobacter_1 | 0.000 | 0.000 | 0.018 | 0.000 | 0.000 | 0.000 |
| Firmicutes | Clostridia_3 | Thermoanaerobacterales_3 | Thermoanaerobacteraceae | Caldanaerobacter_2 | 0.000 | 0.000 | 0.012 | 0.000 | 0.000 | 0.000 |
| Firmicutes | Clostridia_3 | Thermoanaerobacterales_3 | Thermoanaerobacteraceae | Caldanaerobacter_5 | 0.000 | 0.000 | 0.004 | 0.000 | 0.000 | 0.000 |
| Firmicutes | Clostridia_2 | Thermoanerobacterales | Thermoanaerobacteraceae | Ammonifex | 0.000 | 0.000 | 0.006 | 0.000 | 0.000 | 0.000 |
| Firmicutes | Clostridia_2 | Thermoanerobacterales | Thermoanaerobacteraceae | Carboxydothermus | 0.000 | 0.000 | 0.006 | 0.000 | 0.000 | 0.000 |
| Firmicutes | Clostridia_2 | Thermoanerobacterales | Thermoanaerobacteraceae | Gelria | 0.000 | 0.000 | 0.026 | 0.000 | 0.000 | 0.000 |
| Firmicutes | Clostridia_2 | Thermoanerobacterales | Thermoanaerobacteraceae | Thermacetogenium | 0.000 | 0.000 | 0.008 | 0.000 | 0.000 | 0.000 |
| Firmicutes | Clostridia | Thermoanaerobacterales | Thermodesulfobiaceae | Coprothermobacter | 0.000 | 0.006 | 0.121 | 0.000 | 0.000 | 0.000 |
| Firmicutes | Clostridia_4 | Thermoanaerobacterales_4 | Thermodesulfobiaceae | Thermodesulfobium | 0.000 | 0.000 | 0.004 | 0.000 | 0.000 | 0.000 |
| Firmicutes | Tissierellia | Tissierellales | Tissierellaceae | Trabulsiella | 2.440 | 12.809 | 0.482 | 5.523 | 1.196 | 3.238 |
| Firmicutes | Clostridia | Clostridiales | Unclassified | Isoptericola | 0.000 | 0.000 | 0.000 | 0.006 | 0.000 | 0.012 |
| Firmicutes | Clostridia_4 | Clostridiales | Unclassified | Fenollaria | 0.024 | 0.000 | 0.000 | 0.000 | 0.000 | 0.000 |
| Firmicutes | Clostridia_2 | Clostridiales_1 | Veillonellaceae | Acetonema_a | 0.000 | 0.000 | 0.020 | 0.000 | 0.000 | 0.000 |
| Firmicutes | Clostridia_2 | Clostridiales_1 | Veillonellaceae | Acidaminococcus | 0.000 | 0.000 | 0.008 | 0.000 | 0.000 | 0.000 |
| Firmicutes | Clostridia_2 | Clostridiales_1 | Veillonellaceae | Allisonella | 0.000 | 0.000 | 0.006 | 0.000 | 0.000 | 0.000 |
| Firmicutes | Clostridia_2 | Clostridiales_1 | Veillonellaceae | Anaeroarcus-Anaeromusa | 0.000 | 0.000 | 0.014 | 0.000 | 0.000 | 0.000 |
| Firmicutes | Clostridia_2 | Clostridiales_1 | Veillonellaceae | Anaerovibrio | 0.000 | 0.000 | 0.008 | 0.000 | 0.000 | 0.000 |
| Firmicutes | Clostridia_2 | Clostridiales_1 | Veillonellaceae | Dendrosporobacter | 0.016 | 0.000 | 0.004 | 0.000 | 0.005 | 0.000 |
| Firmicutes | Clostridia_2 | Clostridiales_1 | Veillonellaceae | Dialister_1 | 0.000 | 0.000 | 0.054 | 0.000 | 0.000 | 0.000 |
| Firmicutes | Clostridia_2 | Clostridiales_1 | Veillonellaceae | Megamonas | 0.000 | 0.000 | 0.030 | 0.000 | 0.000 | 0.000 |
| Firmicutes | Clostridia_2 | Clostridiales_1 | Veillonellaceae | Pectinatus | 0.000 | 0.000 | 0.004 | 0.000 | 0.000 | 0.000 |
| Firmicutes | Clostridia_2 | Clostridiales_1 | Veillonellaceae | Phascolarctobacterium | 0.000 | 0.000 | 0.060 | 0.000 | 0.000 | 0.000 |
| Firmicutes | Clostridia_2 | Clostridiales_1 | Veillonellaceae | Phaselicystis | 0.008 | 0.000 | 0.008 | 0.006 | 0.000 | 0.000 |
| Firmicutes | Clostridia_2 | Clostridiales_1 | Veillonellaceae | Propionispira-Zymophilus | 0.000 | 0.000 | 0.008 | 0.000 | 0.000 | 0.000 |
| Firmicutes | Clostridia_2 | Clostridiales_1 | Veillonellaceae | Quinella | 0.000 | 0.000 | 0.006 | 0.000 | 0.000 | 0.000 |
| Firmicutes | Clostridia_2 | Clostridiales_1 | Veillonellaceae | Schwartzia | 0.000 | 0.000 | 0.004 | 0.000 | 0.000 | 0.000 |
| Firmicutes | Clostridia_2 | Clostridiales_1 | Veillonellaceae | Selenomonas_2 | 0.000 | 0.000 | 0.006 | 0.000 | 0.000 | 0.000 |
| Firmicutes | Clostridia_2 | Clostridiales_1 | Veillonellaceae | Selenomonas_5 | 0.000 | 0.000 | 0.008 | 0.000 | 0.000 | 0.000 |
| Firmicutes | Clostridia_2 | Clostridiales_1 | Veillonellaceae | Selenomonas_7 | 0.000 | 0.000 | 0.010 | 0.000 | 0.000 | 0.000 |
| Firmicutes | Clostridia_2 | Clostridiales_1 | Veillonellaceae | Selenomonas_sp_a | 0.000 | 0.000 | 0.004 | 0.000 | 0.000 | 0.000 |
| Firmicutes | Clostridia_2 | Clostridiales_1 | Veillonellaceae | Sulfurimonas | 0.000 | 0.000 | 0.038 | 0.000 | 0.000 | 0.000 |

**Table S9 continued.**

| | | | | | | | | | | |
|---|---|---|---|---|---|---|---|---|---|---|
| Firmicutes | Clostridia_2 | Clostridiales_1 | Veillonellaceae | Uncultured_9 | 0.016 | 0.006 | 0.623 | 0.000 | 0.000 | 0.006 |
| Firmicutes | Clostridia_2 | Clostridiales_1 | Veillonellaceae | Verrucomicrobium | 0.000 | 0.000 | 0.008 | 0.000 | 0.000 | 0.000 |
| Firmicutes | Negativicutes | Veillonellales | Veillonellaceae | Meiothermus | 0.000 | 0.000 | 0.040 | 0.000 | 0.000 | 0.000 |
| Firmicutes | Negativicutes | Veillonellales | Veillonellaceae | Neisseria_1 | 0.008 | 0.000 | 0.026 | 0.000 | 0.000 | 0.000 |
| Firmicutes | Negativicutes | Veillonellales | Veillonellaceae | Neisseria_2 | 0.000 | 0.000 | 0.012 | 0.000 | 0.000 | 0.000 |
| Firmicutes | Negativicutes | Veillonellales | Veillonellaceae | Neisseria_3 | 0.000 | 0.000 | 0.006 | 0.000 | 0.000 | 0.000 |
| Firmicutes | Negativicutes | Veillonellales | Veillonellaceae | Neisseria_4 | 0.000 | 0.000 | 0.008 | 0.000 | 0.000 | 0.000 |
| Firmicutes_1 | Clostridia | Halanaerobiales | Halanaerobiaceae | Halanaerobium | 0.000 | 0.000 | 0.062 | 0.000 | 0.000 | 0.000 |
| Firmicutes_1 | Clostridia | Halanaerobiales | Halobacteroidaceae | Halanaerobacter_1 | 0.000 | 0.000 | 0.004 | 0.000 | 0.000 | 0.000 |
| Firmicutes_1 | Clostridia | Halanaerobiales | Halobacteroidaceae | Halanaerobacter_2 | 0.000 | 0.000 | 0.006 | 0.000 | 0.000 | 0.000 |
| Firmicutes_1 | Clostridia | Halanaerobiales | Halobacteroidaceae | Halobacteroides_2 | 0.000 | 0.000 | 0.010 | 0.000 | 0.000 | 0.000 |
| Firmicutes_1 | Clostridia | Halanaerobiales | Halobacteroidaceae | Orenia | 0.000 | 0.000 | 0.016 | 0.000 | 0.000 | 0.000 |
| Fusobacteria | Fusobacteria | Fusobacteriales | Fusobacteriaceae | Ilyobacter | 0.000 | 0.000 | 0.006 | 0.000 | 0.000 | 0.000 |
| Fusobacteria | Fusobacteria | Fusobacteriales | Fusobacteriaceae | Propionigenium | 0.000 | 0.000 | 0.012 | 0.000 | 0.000 | 0.000 |
| Fusobacteria | Fusobacteria | Fusobacteriales | Fusobacteriaceae | Psychrilyobacter | 0.000 | 0.000 | 0.014 | 0.000 | 0.000 | 0.000 |
| Fusobacteria | Fusobacteriia | Fusobacteriales | Fusobacteriaceae | Cetobacterium | 0.000 | 0.000 | 0.087 | 0.000 | 0.000 | 0.000 |
| Fusobacteria | Fusobacteriia | Fusobacteriales | Fusobacteriaceae | Fusobacterium | 0.008 | 0.000 | 0.000 | 0.006 | 0.000 | 0.000 |
| Fusobacteria | Fusobacteriia | Fusobacteriales | Fusobacteriaceae | Fusobacterium_1 | 0.000 | 0.000 | 0.115 | 0.000 | 0.000 | 0.000 |
| Fusobacteria | Fusobacteriia | Fusobacteriales | Fusobacteriaceae | Fusobacterium_2 | 0.000 | 0.000 | 0.006 | 0.000 | 0.000 | 0.000 |
| Fusobacteria | Fusobacteriia | Fusobacteriales | Fusobacteriaceae | Iamia | 0.016 | 0.000 | 0.020 | 0.006 | 0.000 | 0.000 |
| Fusobacteria | Fusobacteria | Fusobacteriales | Leptotrichiaceae | Sediminibacterium | 0.041 | 0.000 | 0.030 | 0.000 | 0.000 | 0.000 |
| Fusobacteria | Fusobacteria | Fusobacteriales | Leptotrichiaceae | Sneathia | 0.000 | 0.000 | 0.004 | 0.000 | 0.000 | 0.000 |
| Fusobacteria | Fusobacteria | Fusobacteriales | Leptotrichiaceae | Streptobacillus | 0.000 | 0.000 | 0.006 | 0.000 | 0.000 | 0.000 |
| Fusobacteria | Fusobacteriia | Fusobacteriales | Leptotrichiaceae | Leucobacter | 0.000 | 0.006 | 0.038 | 0.012 | 0.000 | 0.000 |
| Fusobacteria | Fusobacteriia | Fusobacteriales | Leptotrichiaceae | Oceanobacillus | 0.000 | 0.000 | 0.036 | 0.000 | 0.000 | 0.006 |
| Gemmatimonadetes | Gemmatimonadetes | Gemmatimonadales | Gemmatimonadaceae | Gemmatimonas | 0.089 | 0.000 | 0.183 | 0.000 | 0.000 | 0.000 |
| Lentisphaerae | Lentisphaeria | Lentisphaerales | Lentisphaeraceae | Lentisphaera | 0.000 | 0.000 | 0.016 | 0.000 | 0.000 | 0.000 |
| Lentisphaerae | Lentisphaeria | Victivallales | Victivallaceae | Victivallis | 0.000 | 0.000 | 0.075 | 0.000 | 0.000 | 0.000 |
| Nitrospinae | Nitrospinia | Nitrospinales | Nitrospinaceae | Nitrospira | 0.089 | 0.012 | 0.270 | 0.006 | 0.000 | 0.000 |
| Nitrospirae | Nitrospira | Nitrospirales | Nitrospiraceae | Candidatus_Magnetobacterium | 0.000 | 0.000 | 0.008 | 0.000 | 0.000 | 0.000 |
| Nitrospirae | Nitrospira | Nitrospirales | Nitrospiraceae | Leptospirillum | 0.000 | 0.000 | 0.119 | 0.000 | 0.000 | 0.000 |
| Nitrospirae | Nitrospira | Nitrospirales | Nitrospiraceae | Nocardia | 0.000 | 0.000 | 0.214 | 0.000 | 0.000 | 0.000 |
| Nitrospirae | Nitrospira | Nitrospirales | Nitrospiraceae | Thermomonas_1 | 0.033 | 0.000 | 0.010 | 0.000 | 0.000 | 0.000 |
| Planctomycetes | Planctomycetacia | Brocadiales | Brocadiaceae | Candidatus_Anammoxoglobus | 0.000 | 0.000 | 0.004 | 0.000 | 0.000 | 0.000 |
| Planctomycetes | Planctomycetacia | Brocadiales | Brocadiaceae | Candidatus_Brocadia_anammoxidans | 0.000 | 0.000 | 0.012 | 0.000 | 0.000 | 0.000 |
| Planctomycetes | Planctomycetacia | Brocadiales | Brocadiaceae | Candidatus_Brocadia_fulgida | 0.000 | 0.000 | 0.012 | 0.000 | 0.000 | 0.000 |
| Planctomycetes | Planctomycetacia | Brocadiales | Brocadiaceae | Candidatus_Jettenia | 0.000 | 0.000 | 0.010 | 0.000 | 0.000 | 0.000 |
| Planctomycetes | Planctomycetacia | Brocadiales | Brocadiaceae | Candidatus_Kuenenia | 0.000 | 0.000 | 0.004 | 0.000 | 0.000 | 0.000 |
| Planctomycetes | Planctomycetacia | Brocadiales | Brocadiaceae | Candidatus_Scalindua | 0.000 | 0.000 | 0.064 | 0.000 | 0.000 | 0.000 |
| Planctomycetes | Phycisphaerae | Phycisphaerales | Phycisphaeraceae | Albidovulum | 0.000 | 0.000 | 0.002 | 0.000 | 0.000 | 0.000 |

**Table S9 continued.**

| | | | | | | | | | | |
|---|---|---|---|---|---|---|---|---|---|---|
| Planctomycetes | Phycisphaerae | Phycisphaerales | Phycisphaeraceae | CL500_3 | 0.000 | 0.000 | 0.032 | 0.000 | 0.000 | 0.000 |
| Planctomycetes | Phycisphaerae | Phycisphaerales | Phycisphaeraceae | FS140-16B-02_marine_group | 0.000 | 0.000 | 0.010 | 0.000 | 0.000 | 0.000 |
| Planctomycetes | Phycisphaerae | Phycisphaerales | Phycisphaeraceae | I_8 | 0.000 | 0.000 | 0.006 | 0.000 | 0.000 | 0.000 |
| Planctomycetes | Phycisphaerae | Phycisphaerales | Phycisphaeraceae | Phycisphaera | 0.000 | 0.000 | 0.067 | 0.000 | 0.000 | 0.000 |
| Planctomycetes | Phycisphaerae | Phycisphaerales | Phycisphaeraceae | Sneathiella | 0.000 | 0.000 | 0.006 | 0.000 | 0.000 | 0.000 |
| Planctomycetes | Phycisphaerae | Phycisphaerales | Phycisphaeraceae | Urania-1B-19_marine_sediment_group | 0.000 | 0.000 | 0.054 | 0.000 | 0.000 | 0.000 |
| Planctomycetes | Phycisphaerae | Phycisphaerales | Phycisphaeraceae | Z195MB87 | 0.000 | 0.000 | 0.006 | 0.000 | 0.000 | 0.000 |
| Planctomycetes | Planctomycetacia | Planctomycetales | Planctomycetaceae | Blastopirellula | 0.000 | 0.000 | 0.157 | 0.000 | 0.000 | 0.000 |
| Planctomycetes | Planctomycetacia | Planctomycetales | Planctomycetaceae | Gemmata | 0.000 | 0.006 | 0.179 | 0.000 | 0.000 | 0.000 |
| Planctomycetes | Planctomycetacia | Planctomycetales | Planctomycetaceae | Hypersaline_mat_cluster | 0.000 | 0.000 | 0.004 | 0.000 | 0.000 | 0.000 |
| Planctomycetes | Planctomycetacia | Planctomycetales | Planctomycetaceae | Janibacter | 0.000 | 0.000 | 0.018 | 0.000 | 0.000 | 0.000 |
| Planctomycetes | Planctomycetacia | Planctomycetales | Planctomycetaceae | Pir1_lineage | 0.000 | 0.000 | 0.006 | 0.000 | 0.000 | 0.000 |
| Planctomycetes | Planctomycetacia | Planctomycetales | Planctomycetaceae | Pir2_lineage | 0.000 | 0.000 | 0.008 | 0.000 | 0.000 | 0.000 |
| Planctomycetes | Planctomycetacia | Planctomycetales | Planctomycetaceae | Pir3_lineage | 0.000 | 0.000 | 0.006 | 0.000 | 0.000 | 0.000 |
| Planctomycetes | Planctomycetacia | Planctomycetales | Planctomycetaceae | Pir4_lineage | 0.000 | 0.000 | 0.343 | 0.000 | 0.000 | 0.000 |
| Planctomycetes | Planctomycetacia | Planctomycetales | Planctomycetaceae | Pirellula | 0.000 | 0.000 | 0.262 | 0.000 | 0.000 | 0.000 |
| Planctomycetes | Planctomycetacia | Planctomycetales | Planctomycetaceae | Planctomyces_1 | 0.000 | 0.000 | 0.502 | 0.000 | 0.000 | 0.000 |
| Planctomycetes | Planctomycetacia | Planctomycetales | Planctomycetaceae | Planctomyces_2 | 0.000 | 0.000 | 0.042 | 0.000 | 0.000 | 0.000 |
| Planctomycetes | Planctomycetacia | Planctomycetales | Planctomycetaceae | Planococcus | 0.000 | 0.000 | 0.000 | 0.006 | 0.000 | 0.000 |
| Planctomycetes | Planctomycetacia | Planctomycetales | Planctomycetaceae | Rhodopseudomonas | 0.008 | 0.006 | 0.020 | 0.000 | 0.005 | 0.006 |
| Planctomycetes | Planctomycetacia | Planctomycetales | Planctomycetaceae | Schlesneria | 0.000 | 0.000 | 0.028 | 0.000 | 0.000 | 0.000 |
| Planctomycetes | Planctomycetacia | Planctomycetales | Planctomycetaceae | Sinorhizobium | 0.016 | 0.018 | 0.002 | 0.000 | 0.000 | 0.024 |
| Planctomycetes | Planctomycetacia | Planctomycetales | Planctomycetaceae | Uncultured_18 | 0.016 | 0.000 | 0.611 | 0.006 | 0.000 | 0.000 |
| Planctomycetes | Planctomycetacia | Planctomycetales | Planctomycetaceae | Uncultured_34 | 0.000 | 0.006 | 0.034 | 0.000 | 0.000 | 0.000 |
| Planctomycetes | Planctomycetacia | Planctomycetales | Planctomycetaceae | Uncultured_c | 0.000 | 0.000 | 0.014 | 0.000 | 0.000 | 0.000 |
| Planctomycetes | Planctomycetacia | Planctomycetales | Planctomycetaceae | Uncultured_gut_Group_A | 0.000 | 0.000 | 0.006 | 0.000 | 0.000 | 0.000 |
| Planctomycetes | Planctomycetacia | Planctomycetales | Planctomycetaceae | Zavarzinella | 0.000 | 0.000 | 0.042 | 0.000 | 0.000 | 0.000 |
| Planctomycetes | Planctomycetacia | Planctomycetales | Planctomycetaceae | Zobellia | 0.000 | 0.000 | 0.006 | 0.000 | 0.000 | 0.000 |
| Proteobacteria | Alphaproteobacteria | Rhodospirillales | Acetobacteraceae | Acidicaldus | 0.000 | 0.000 | 0.014 | 0.000 | 0.000 | 0.000 |
| Proteobacteria | Alphaproteobacteria | Rhodospirillales | Acetobacteraceae | Gluconobacter | 0.000 | 0.006 | 0.012 | 0.000 | 0.000 | 0.000 |
| Proteobacteria | Alphaproteobacteria | Rhodospirillales | Acetobacteraceae | Granulibacter | 0.000 | 0.000 | 0.002 | 0.000 | 0.000 | 0.000 |
| Proteobacteria | Alphaproteobacteria | Rhodospirillales | Acetobacteraceae | Kordia | 0.000 | 0.000 | 0.004 | 0.000 | 0.000 | 0.000 |
| Proteobacteria | Alphaproteobacteria | Rhodospirillales | Acetobacteraceae | Neorickettsia | 0.000 | 0.000 | 0.012 | 0.000 | 0.000 | 0.000 |
| Proteobacteria | Alphaproteobacteria | Rhodospirillales | Acetobacteraceae | Parafilimonas | 0.000 | 0.000 | 0.000 | 0.006 | 0.000 | 0.000 |
| Proteobacteria | Alphaproteobacteria | Rhodospirillales | Acetobacteraceae | Rhodovulum | 0.000 | 0.000 | 0.004 | 0.000 | 0.000 | 0.000 |
| Proteobacteria | Alphaproteobacteria | Rhodospirillales | Acetobacteraceae | Roseococcus | 0.000 | 0.000 | 0.004 | 0.000 | 0.000 | 0.000 |
| Proteobacteria | Alphaproteobacteria | Rhodospirillales | Acetobacteraceae | Roseomonas_1 | 0.000 | 0.000 | 0.026 | 0.000 | 0.005 | 0.000 |
| Proteobacteria | Alphaproteobacteria | Rhodospirillales | Acetobacteraceae | Roseomonas_2 | 0.000 | 0.000 | 0.010 | 0.000 | 0.000 | 0.006 |
| Proteobacteria | Alphaproteobacteria | Rhodospirillales | Acetobacteraceae | Roseomonas_4 | 0.000 | 0.000 | 0.008 | 0.000 | 0.000 | 0.000 |
| Proteobacteria | Alphaproteobacteria | Rhodospirillales | Acetobacteraceae | Rothia | 0.008 | 0.000 | 0.024 | 0.000 | 0.000 | 0.000 |

**Table S9 continued.**

| | | | | | | | | | | |
|---|---|---|---|---|---|---|---|---|---|---|
| Proteobacteria | Alphaproteobacteria | Rhodospirillales_1 | Acetobacteraceae | Acetobacter | 0.000 | 0.000 | 0.042 | 0.006 | 0.000 | 0.000 |
| Proteobacteria | Alphaproteobacteria | Rhodospirillales_1 | Acetobacteraceae | Acidiphilium | 0.000 | 0.000 | 0.042 | 0.000 | 0.000 | 0.000 |
| Proteobacteria | Alphaproteobacteria | Rhodospirillales_1 | Acetobacteraceae | Acidisoma | 0.000 | 0.006 | 0.006 | 0.000 | 0.000 | 0.000 |
| Proteobacteria | Alphaproteobacteria | Rhodospirillales_1 | Acetobacteraceae | Acidocella | 0.000 | 0.000 | 0.012 | 0.000 | 0.000 | 0.000 |
| Proteobacteria | Alphaproteobacteria | Rhodospirillales_1 | Acetobacteraceae | Asaia_1 | 0.000 | 0.000 | 0.006 | 0.000 | 0.000 | 0.000 |
| Proteobacteria | Alphaproteobacteria | Rhodospirillales_1 | Acetobacteraceae | Craurococcus | 0.000 | 0.000 | 0.006 | 0.000 | 0.000 | 0.000 |
| Proteobacteria | Alphaproteobacteria | Rhodospirillales_1 | Acetobacteraceae | Gluconacetobacter | 0.000 | 0.000 | 0.022 | 0.000 | 0.005 | 0.000 |
| Proteobacteria | Alphaproteobacteria | Rhodospirillales_1 | Acetobacteraceae | Rhodopila | 0.000 | 0.000 | 0.004 | 0.000 | 0.000 | 0.000 |
| Proteobacteria | Alphaproteobacteria | Rhodospirillales_1 | Acetobacteraceae | Rhodoplanes | 0.000 | 0.000 | 0.018 | 0.000 | 0.000 | 0.000 |
| Proteobacteria | Alphaproteobacteria | Rhodospirillales_1 | Acetobacteraceae | Rhodovarius | 0.000 | 0.000 | 0.004 | 0.000 | 0.000 | 0.000 |
| Proteobacteria | Alphaproteobacteria | Rhodospirillales_1 | Acetobacteraceae | Rhodovibrio | 0.000 | 0.000 | 0.004 | 0.000 | 0.000 | 0.000 |
| Proteobacteria | Alphaproteobacteria | Rhodospirillales_1 | Acetobacteraceae | Saccharibacter | 0.000 | 0.000 | 0.018 | 0.000 | 0.000 | 0.000 |
| Proteobacteria | Alphaproteobacteria | Rhodospirillales_1 | Acetobacteraceae | Saccharofermentans | 0.000 | 0.000 | 0.000 | 0.006 | 0.000 | 0.000 |
| Proteobacteria | Gammaproteobacteria_2 | Acidithiobacillales | Acidithiobacillaceae | Acidithiobacillus | 0.000 | 0.000 | 0.052 | 0.000 | 0.000 | 0.000 |
| Proteobacteria | Gammaproteobacteria_1 | Aeromonadales | Aeromonadaceae | Aeromonas_1 | 0.049 | 0.012 | 0.365 | 0.018 | 0.020 | 0.018 |
| Proteobacteria | Gammaproteobacteria_1 | Aeromonadales | Aeromonadaceae | Aeromonas_2 | 0.000 | 0.000 | 0.006 | 0.000 | 0.000 | 0.000 |
| Proteobacteria | Gammaproteobacteria_1 | Aeromonadales | Aeromonadaceae | Oceanimonas | 0.000 | 0.000 | 0.012 | 0.000 | 0.000 | 0.000 |
| Proteobacteria | Gammaproteobacteria_1 | Aeromonadales | Aeromonadaceae | Tolumonas | 0.000 | 0.000 | 0.010 | 0.000 | 0.005 | 0.000 |
| Proteobacteria | Betaproteobacteria | Burkholderiales | Alacaligenaceae | Derxia | 0.041 | 0.000 | 0.018 | 0.006 | 0.000 | 0.018 |
| Proteobacteria | Betaproteobacteria | Burkholderiales | Alcaligenaceae | Advenella | 0.033 | 0.012 | 0.004 | 0.030 | 0.070 | 0.061 |
| Proteobacteria | Betaproteobacteria | Burkholderiales | Alcaligenaceae | Amphritea | 0.000 | 0.000 | 0.010 | 0.000 | 0.000 | 0.000 |
| Proteobacteria | Betaproteobacteria | Burkholderiales | Alcaligenaceae | Ampullimonas | 0.016 | 0.000 | 0.000 | 0.000 | 0.005 | 0.000 |
| Proteobacteria | Betaproteobacteria | Burkholderiales | Alcaligenaceae | Basilea | 0.008 | 0.000 | 0.000 | 0.000 | 0.000 | 0.000 |
| Proteobacteria | Betaproteobacteria | Burkholderiales | Alcaligenaceae | Brackiella | 0.000 | 0.000 | 0.000 | 0.006 | 0.000 | 0.000 |
| Proteobacteria | Betaproteobacteria | Burkholderiales | Alcaligenaceae | Candidimonas | 0.041 | 0.006 | 0.000 | 0.024 | 0.065 | 0.030 |
| Proteobacteria | Betaproteobacteria | Burkholderiales | Alcaligenaceae | Eoetvoesia | 0.008 | 0.000 | 0.000 | 0.006 | 0.030 | 0.000 |
| Proteobacteria | Betaproteobacteria | Burkholderiales | Alcaligenaceae | Kerstersia | 0.016 | 0.012 | 0.004 | 0.048 | 0.105 | 0.054 |
| Proteobacteria | Betaproteobacteria | Burkholderiales | Alcaligenaceae | Paenalcaligenes | 0.000 | 0.000 | 0.000 | 0.000 | 0.005 | 0.006 |
| Proteobacteria | Betaproteobacteria | Burkholderiales | Alcaligenaceae | Paralcaligenes | 0.024 | 0.000 | 0.000 | 0.000 | 0.000 | 0.000 |
| Proteobacteria | Betaproteobacteria | Burkholderiales | Alcaligenaceae | Parapusillimonas | 0.016 | 0.000 | 0.000 | 0.012 | 0.005 | 0.006 |
| Proteobacteria | Betaproteobacteria | Burkholderiales | Alcaligenaceae | Pelobacter | 0.008 | 0.000 | 0.012 | 0.000 | 0.000 | 0.000 |
| Proteobacteria | Betaproteobacteria | Burkholderiales | Alcaligenaceae_0 | Achromobacter | 0.057 | 0.025 | 0.010 | 0.066 | 0.169 | 0.436 |
| Proteobacteria | Betaproteobacteria | Burkholderiales | Alcaligenaceae_1 | Achromobacter_1 | 0.016 | 0.000 | 0.004 | 0.006 | 0.015 | 0.000 |
| Proteobacteria | Betaproteobacteria | Burkholderiales | Alcaligenaceae_1 | Achromobacter_2 | 0.260 | 0.074 | 0.048 | 0.276 | 0.324 | 0.272 |
| Proteobacteria | Betaproteobacteria | Burkholderiales | Alcaligenaceae_1 | Achromobacter_3 | 0.049 | 0.012 | 0.014 | 0.042 | 0.030 | 0.018 |
| Proteobacteria | Betaproteobacteria | Burkholderiales | Alcaligenaceae_1 | Alcaligenes | 0.016 | 0.000 | 0.018 | 0.000 | 0.010 | 0.012 |
| Proteobacteria | Betaproteobacteria | Burkholderiales | Alcaligenaceae_1 | Bordetella_2 | 0.138 | 0.031 | 0.014 | 0.096 | 0.189 | 0.248 |
| Proteobacteria | Betaproteobacteria | Burkholderiales | Alcaligenaceae_1 | Castellaniella | 0.106 | 0.025 | 0.014 | 0.060 | 0.090 | 0.097 |
| Proteobacteria | Betaproteobacteria | Burkholderiales | Alcaligenaceae_1 | GKS98_freshwater_group | 0.081 | 0.006 | 0.010 | 0.084 | 0.080 | 0.048 |
| Proteobacteria | Betaproteobacteria | Burkholderiales | Alcaligenaceae_1 | Glaciecola | 0.000 | 0.000 | 0.046 | 0.000 | 0.000 | 0.000 |

**Table S9 continued.**

| | | | | | | | | | | |
|---|---|---|---|---|---|---|---|---|---|---|
| Proteobacteria | Betaproteobacteria | Burkholderiales | Alcaligenaceae_1 | Oligella | 0.000 | 0.000 | 0.004 | 0.000 | 0.000 | 0.000 |
| Proteobacteria | Betaproteobacteria | Burkholderiales | Alcaligenaceae_1 | Parasutterella | 0.000 | 0.000 | 0.046 | 0.000 | 0.000 | 0.000 |
| Proteobacteria | Betaproteobacteria | Burkholderiales | Alcaligenaceae_1 | Pusillimonas_1 | 0.000 | 0.000 | 0.008 | 0.000 | 0.000 | 0.000 |
| Proteobacteria | Betaproteobacteria | Burkholderiales | Alcaligenaceae_1 | Pusillimonas_4 | 0.000 | 0.000 | 0.006 | 0.012 | 0.010 | 0.006 |
| Proteobacteria | Betaproteobacteria | Burkholderiales | Alcaligenaceae_1 | Sutterella | 0.016 | 0.000 | 0.064 | 0.000 | 0.005 | 0.000 |
| Proteobacteria | Betaproteobacteria | Burkholderiales | Alcaligenaceae_1 | Taylorella | 0.000 | 0.000 | 0.008 | 0.000 | 0.000 | 0.006 |
| Proteobacteria | Betaproteobacteria | Burkholderiales | Alcaligenaceae_1 | Thalassospira | 0.708 | 0.228 | 0.113 | 0.174 | 0.020 | 0.030 |
| Proteobacteria | Betaproteobacteria | Burkholderiales | Alcanigenaceae | Bordetella | 9.257 | 2.347 | 1.024 | 10.400 | 14.818 | 16.578 |
| Proteobacteria | Gammaproteobacteria_1 | Oceanospirillales_2 | Alcanivoracaceae | Kangiella | 0.000 | 0.000 | 0.012 | 0.000 | 0.000 | 0.000 |
| Proteobacteria | Gammaproteobacteria | Alteromonadales | Alteromonadaceae | Marinobacterium | 0.000 | 0.000 | 0.000 | 0.000 | 0.005 | 0.000 |
| Proteobacteria | Gammaproteobacteria_1 | Alteromonadales | Alteromonadaceae | BD1-7_clade | 0.000 | 0.000 | 0.048 | 0.000 | 0.000 | 0.000 |
| Proteobacteria | Gammaproteobacteria_1 | Alteromonadales | Alteromonadaceae | BD2_7 | 0.000 | 0.000 | 0.004 | 0.000 | 0.000 | 0.000 |
| Proteobacteria | Gammaproteobacteria_1 | Alteromonadales | Alteromonadaceae | Candidatus_Endobugula | 0.000 | 0.000 | 0.018 | 0.000 | 0.000 | 0.000 |
| Proteobacteria | Gammaproteobacteria_1 | Alteromonadales | Alteromonadaceae | Marinimicrobium | 0.000 | 0.000 | 0.010 | 0.000 | 0.000 | 0.000 |
| Proteobacteria | Gammaproteobacteria_1 | Alteromonadales | Alteromonadaceae | Microbulbifer | 0.000 | 0.000 | 0.030 | 0.000 | 0.000 | 0.000 |
| Proteobacteria | Gammaproteobacteria_1 | Alteromonadales | Alteromonadaceae | Opitutus | 0.024 | 0.000 | 0.135 | 0.000 | 0.000 | 0.006 |
| Proteobacteria | Gammaproteobacteria_1 | Alteromonadales | Alteromonadaceae | SAR92_clade | 0.000 | 0.000 | 0.022 | 0.000 | 0.000 | 0.000 |
| Proteobacteria | Gammaproteobacteria_1 | Alteromonadales | Alteromonadaceae | Teredinibacter | 0.000 | 0.000 | 0.030 | 0.000 | 0.000 | 0.000 |
| Proteobacteria | Gammaproteobacteria_1 | Alteromonadales | Alteromonadaceae | Termite_cluster_1 | 0.000 | 0.000 | 0.036 | 0.000 | 0.000 | 0.000 |
| Proteobacteria | Gammaproteobacteria_1 | Alteromonadales_1 | Alteromonadaceae_1 | Aestuariibacter | 0.000 | 0.000 | 0.008 | 0.000 | 0.000 | 0.000 |
| Proteobacteria | Gammaproteobacteria_1 | Alteromonadales_1 | Alteromonadaceae_1 | Alteromonas_1 | 0.000 | 0.000 | 0.032 | 0.000 | 0.000 | 0.000 |
| Proteobacteria | Gammaproteobacteria_1 | Alteromonadales_1 | Alteromonadaceae_1 | Alteromonas_2 | 0.000 | 0.000 | 0.012 | 0.000 | 0.000 | 0.000 |
| Proteobacteria | Gammaproteobacteria_1 | Alteromonadales_1 | Alteromonadaceae_1 | Salinimonas | 0.000 | 0.000 | 0.004 | 0.000 | 0.000 | 0.000 |
| Proteobacteria | Gammaproteobacteria_1 | Alteromonadales_1 | Alteromonadaceae_2 | Aliagarivorans | 0.000 | 0.000 | 0.004 | 0.000 | 0.000 | 0.000 |
| Proteobacteria | Alphaproteobacteria | Rickettsiales | Anaplasmataceae | Anaplasma | 0.000 | 0.000 | 0.008 | 0.000 | 0.000 | 0.000 |
| Proteobacteria | Alphaproteobacteria | Rickettsiales | Anaplasmataceae | Candidatus_Xenohaliotis | 0.000 | 0.000 | 0.004 | 0.000 | 0.000 | 0.000 |
| Proteobacteria | Alphaproteobacteria | Rickettsiales | Anaplasmataceae | Ehrlichia | 0.000 | 0.000 | 0.014 | 0.000 | 0.010 | 0.000 |
| Proteobacteria | Alphaproteobacteria | Rickettsiales | Anaplasmataceae | Nesiotobacter | 0.000 | 0.000 | 0.004 | 0.000 | 0.000 | 0.000 |
| Proteobacteria | Alphaproteobacteria | Rickettsiales | Anaplasmataceae | Xanthomonas | 0.016 | 0.012 | 0.020 | 0.006 | 0.005 | 0.091 |
| Proteobacteria | Deltaproteobacteria | Myxococcales | Archangiaceae | Aneurinibacillus | 0.000 | 0.000 | 0.012 | 0.000 | 0.000 | 0.000 |
| Proteobacteria | Deltaproteobacteria | Myxococcales | Archangiaceae | Cystobacter | 0.008 | 0.000 | 0.000 | 0.000 | 0.000 | 0.000 |
| Proteobacteria | Alphaproteobacteria | Hyphomicrobiales | Aurantimonadaceae | Martelella | 0.000 | 0.006 | 0.000 | 0.006 | 0.000 | 0.000 |
| Proteobacteria | Alphaproteobacteria | Rhizobiales | Aurantimonadaceae | Fructobacillus | 0.000 | 0.000 | 0.010 | 0.000 | 0.000 | 0.000 |
| Proteobacteria | Alphaproteobacteria | Rhizobiales_1 | Aurantimonadaceae | Aurantimonas_3 | 0.000 | 0.000 | 0.014 | 0.000 | 0.000 | 0.000 |
| Proteobacteria | Alphaproteobacteria | Rhizobiales_1 | Aurantimonadaceae | Aureimonas | 0.000 | 0.000 | 0.000 | 0.000 | 0.000 | 0.006 |
| Proteobacteria | Alphaproteobacteria | Rhizobiales_1 | Aurantimonadaceae | Azoarcus | 0.000 | 0.000 | 0.000 | 0.000 | 0.005 | 0.000 |
| Proteobacteria | Alphaproteobacteria | Rhizobiales_1 | Aurantimonadaceae | Azoarcus_2 | 0.033 | 0.000 | 0.024 | 0.000 | 0.000 | 0.000 |
| Proteobacteria | Alphaproteobacteria | Rhizobiales_1 | Aurantimonadaceae | Fulvimarina | 0.000 | 0.000 | 0.004 | 0.000 | 0.000 | 0.000 |
| Proteobacteria | Betaproteobacteria | Rhodocyclales | Azonexaceae | Azospira | 0.000 | 0.000 | 0.028 | 0.000 | 0.000 | 0.000 |
| Proteobacteria | Betaproteobacteria | Rhodocyclales | Azonexaceae | Azospirillum | 0.008 | 0.000 | 0.000 | 0.000 | 0.000 | 0.000 |

**Table S9 continued.**

| | | | | | | | | | | |
|---|---|---|---|---|---|---|---|---|---|---|
| Proteobacteria | Betaproteobacteria | Rhodocyclales | Azonexaceae | Azospirillum_1 | 0.000 | 0.000 | 0.040 | 0.000 | 0.000 | 0.000 |
| Proteobacteria | Betaproteobacteria | Rhodocyclales | Azonexaceae | Azospirillum_2 | 0.000 | 0.000 | 0.004 | 0.000 | 0.000 | 0.000 |
| Proteobacteria | Deltaproteobacteria | Bdellovibrionales | Bacteriovoraceae | Bacteriovorax_1 | 0.000 | 0.000 | 0.016 | 0.000 | 0.000 | 0.000 |
| Proteobacteria | Deltaproteobacteria | Bdellovibrionales | Bacteriovoraceae | Bacteriovorax_2 | 0.000 | 0.000 | 0.008 | 0.000 | 0.000 | 0.000 |
| Proteobacteria | Deltaproteobacteria | Bdellovibrionales | Bacteriovoraceae | Perlucidibaca | 0.049 | 0.006 | 0.022 | 0.000 | 0.000 | 0.000 |
| Proteobacteria | Gammaproteobacteria | Oceanospirillales | Balneatrichaceae | Balneatrix | 0.000 | 0.000 | 0.012 | 0.000 | 0.000 | 0.000 |
| Proteobacteria | Alphaproteobacteria | Rhizobiales_1 | Bartonellaceae | Bartonella | 0.000 | 0.006 | 0.048 | 0.006 | 0.000 | 0.006 |
| Proteobacteria | Deltaproteobacteria | Bdellovibrionales | Bdellovibrionaceae | Bdellovibrio | 0.000 | 0.006 | 0.040 | 0.006 | 0.000 | 0.000 |
| Proteobacteria | Deltaproteobacteria | Bdellovibrionales | Bdellovibrionaceae | OM60_NOR5__clade | 0.000 | 0.000 | 0.097 | 0.000 | 0.000 | 0.000 |
| Proteobacteria | Alphaproteobacteria | Hyphomicrobiales | Beijerinckiaceae | Methyloferula | 0.008 | 0.000 | 0.000 | 0.000 | 0.000 | 0.000 |
| Proteobacteria | Alphaproteobacteria | Rhizobiales | Beijerinckiaceae | Beijerinckia | 0.016 | 0.000 | 0.008 | 0.000 | 0.000 | 0.006 |
| Proteobacteria | Alphaproteobacteria | Rhizobiales | Beijerinckiaceae | Methylovorus | 0.000 | 0.000 | 0.002 | 0.000 | 0.000 | 0.000 |
| Proteobacteria | Alphaproteobacteria | Rhizobiales | Beijernckiaceae | Methylocella | 0.000 | 0.000 | 0.004 | 0.000 | 0.000 | 0.000 |
| Proteobacteria | Alphaproteobacteria | Rhizobiales | Bradyrhizobiaceae | Afipia_1 | 0.000 | 0.000 | 0.008 | 0.000 | 0.000 | 0.000 |
| Proteobacteria | Alphaproteobacteria | Rhizobiales | Bradyrhizobiaceae | Bradyrhizobium | 0.179 | 0.049 | 0.006 | 0.030 | 0.090 | 0.085 |
| Proteobacteria | Alphaproteobacteria | Rhizobiales | Bradyrhizobiaceae | Tatumella | 0.382 | 6.118 | 0.058 | 5.625 | 3.140 | 1.543 |
| Proteobacteria | Alphaproteobacteria | Rhizobiales_0 | Bradyrhizobiaceae | Bradyrhizobium_1 | 0.000 | 0.000 | 0.004 | 0.000 | 0.005 | 0.000 |
| Proteobacteria | Alphaproteobacteria | Rhizobiales_1 | Bradyrhizobiaceae | Bradyrhizobium_10 | 0.081 | 0.012 | 0.004 | 0.012 | 0.020 | 0.006 |
| Proteobacteria | Alphaproteobacteria | Rhizobiales_2 | Bradyrhizobiaceae | Afipia | 0.098 | 0.012 | 0.000 | 0.018 | 0.055 | 0.054 |
| Proteobacteria | Alphaproteobacteria | Rhizobiales_2 | Bradyrhizobiaceae | Balneimonas | 0.016 | 0.000 | 0.012 | 0.000 | 0.000 | 0.000 |
| Proteobacteria | Alphaproteobacteria | Rhizobiales_2 | Bradyrhizobiaceae | Blastobacter | 0.000 | 0.000 | 0.004 | 0.000 | 0.000 | 0.000 |
| Proteobacteria | Alphaproteobacteria | Rhizobiales_2 | Bradyrhizobiaceae | Bosea | 0.024 | 0.006 | 0.022 | 0.006 | 0.000 | 0.006 |
| Proteobacteria | Alphaproteobacteria | Rhizobiales_2 | Bradyrhizobiaceae | Bradyrhizobium_12 | 0.065 | 0.000 | 0.020 | 0.012 | 0.015 | 0.006 |
| Proteobacteria | Alphaproteobacteria | Rhizobiales_2 | Bradyrhizobiaceae | Bradyrhizobium_13 | 0.008 | 0.018 | 0.006 | 0.000 | 0.010 | 0.012 |
| Proteobacteria | Alphaproteobacteria | Rhizobiales_2 | Bradyrhizobiaceae | Bradyrhizobium_6 | 0.049 | 0.006 | 0.004 | 0.006 | 0.015 | 0.018 |
| Proteobacteria | Alphaproteobacteria | Rhizobiales_2 | Bradyrhizobiaceae | Nitrosococcus | 0.016 | 0.006 | 0.024 | 0.000 | 0.000 | 0.000 |
| Proteobacteria | Alphaproteobacteria | Rhizobiales_2 | Bradyrhizobiaceae | Rhodoblastus | 0.000 | 0.000 | 0.004 | 0.000 | 0.000 | 0.000 |
| Proteobacteria | Alphaproteobacteria | Rhizobiales_0 | Brucellaceae | Ochrobactrum_1 | 0.000 | 0.000 | 0.020 | 0.000 | 0.000 | 0.000 |
| Proteobacteria | Alphaproteobacteria | Rhizobiales_1 | Brucellaceae | Brucella | 0.000 | 0.000 | 0.008 | 0.000 | 0.000 | 0.000 |
| Proteobacteria | Alphaproteobacteria | Rhizobiales_1 | Brucellaceae | Ochrobactrum_2 | 0.016 | 0.000 | 0.006 | 0.006 | 0.000 | 0.000 |
| Proteobacteria | Alphaproteobacteria | Rhizobiales_1 | Brucellaceae | Ochrobactrum_3 | 0.000 | 0.000 | 0.022 | 0.000 | 0.000 | 0.018 |
| Proteobacteria | Alphaproteobacteria | Rhizobiales_1 | Brucellaceae | Ochrobactrum_4 | 0.024 | 0.000 | 0.006 | 0.000 | 0.000 | 0.006 |
| Proteobacteria | Alphaproteobacteria | Rhizobiales_1 | Brucellaceae | Ochrobactrum_6 | 0.000 | 0.000 | 0.004 | 0.000 | 0.000 | 0.000 |
| Proteobacteria | Alphaproteobacteria | Rhizobiales_1 | Brucellaceae | Odoribacter | 0.049 | 0.012 | 0.040 | 0.006 | 0.000 | 0.000 |
| Proteobacteria | Alphaproteobacteria | Rhizobiales_1 | Brucellaceae | Pseudocitrobacter | 0.016 | 0.006 | 0.022 | 0.090 | 0.015 | 0.284 |
| Proteobacteria | Betaproteobacteria | Burkholderiales | Burkholderiacea | Burkholderia | 4.019 | 0.333 | 1.435 | 7.087 | 15.546 | 9.835 |
| Proteobacteria | Betaproteobacteria | Burkholderiales | Burkholderiaceae | Burkholderia_1 | 0.781 | 0.099 | 0.220 | 1.893 | 2.951 | 2.506 |
| Proteobacteria | Betaproteobacteria | Burkholderiales | Burkholderiaceae | Caballeronia | 0.000 | 0.000 | 0.000 | 0.000 | 0.005 | 0.000 |
| Proteobacteria | Betaproteobacteria | Burkholderiales | Burkholderiaceae | Chitinimonas | 0.000 | 0.000 | 0.006 | 0.000 | 0.000 | 0.000 |
| Proteobacteria | Betaproteobacteria | Burkholderiales | Burkholderiaceae | Pontibacillus | 0.000 | 0.000 | 0.008 | 0.000 | 0.000 | 0.000 |

**Table S9 continued.**

| | | | | | | | | | | |
|---|---|---|---|---|---|---|---|---|---|---|
| Proteobacteria | Betaproteobacteria | Burkholderiales | Burkholderiaceae_1 | Burkholderia_10 | 0.008 | 0.000 | 0.006 | 0.012 | 0.085 | 0.030 |
| Proteobacteria | Betaproteobacteria | Burkholderiales | Burkholderiaceae_1 | Burkholderia_11 | 0.000 | 0.000 | 0.004 | 0.000 | 0.000 | 0.000 |
| Proteobacteria | Betaproteobacteria | Burkholderiales | Burkholderiaceae_1 | Burkholderia_15 | 0.024 | 0.000 | 0.004 | 0.006 | 0.020 | 0.012 |
| Proteobacteria | Betaproteobacteria | Burkholderiales | Burkholderiaceae_1 | Burkholderia_2 | 0.000 | 0.000 | 0.004 | 0.000 | 0.000 | 0.000 |
| Proteobacteria | Betaproteobacteria | Burkholderiales | Burkholderiaceae_1 | Burkholderia_9 | 0.000 | 0.000 | 0.004 | 0.006 | 0.080 | 0.024 |
| Proteobacteria | Betaproteobacteria | Burkholderiales | Burkholderiaceae_1 | Candidatus_Tremblaya | 0.000 | 0.000 | 0.006 | 0.000 | 0.000 | 0.000 |
| Proteobacteria | Betaproteobacteria | Burkholderiales | Burkholderiaceae_1 | Pannonibacter | 0.033 | 0.000 | 0.004 | 0.000 | 0.005 | 0.000 |
| Proteobacteria | Betaproteobacteria | Burkholderiales | Burkholderiaceae_2 | Cupriavidus | 0.130 | 0.049 | 0.067 | 0.168 | 0.110 | 0.121 |
| Proteobacteria | Betaproteobacteria | Burkholderiales | Burkholderiaceae_2 | Raoultella | 0.073 | 0.173 | 0.058 | 0.210 | 0.274 | 0.200 |
| Proteobacteria | Betaproteobacteria | Burkholderiales | Burkholderiales_genera_incertae_sedis | Rhizobacter | 0.000 | 0.006 | 0.000 | 0.000 | 0.000 | 0.000 |
| Proteobacteria | Alphaproteobacteria | Rickettsiales | Caedibacter | Caedibacter | 0.000 | 0.000 | 0.008 | 0.006 | 0.000 | 0.000 |
| Proteobacteria | Epsilonproteobacteria | Campylobacterales | Campylobacteraceae | Arcobacter | 0.317 | 0.062 | 0.073 | 0.072 | 0.000 | 0.018 |
| Proteobacteria | Epsilonproteobacteria | Campylobacterales | Campylobacteraceae | Campylobacter | 0.000 | 0.000 | 0.109 | 0.000 | 0.000 | 0.000 |
| Proteobacteria | Epsilonproteobacteria | Campylobacterales | Campylobacteraceae | Sulfurovum | 0.008 | 0.000 | 0.069 | 0.000 | 0.000 | 0.000 |
| Proteobacteria | Alphaproteobacteria | Rhodospirillales_2 | Candidatus_Alysiosphaera | Candidatus_Alysiosphaera | 0.000 | 0.006 | 0.014 | 0.000 | 0.000 | 0.000 |
| Proteobacteria | Alphaproteobacteria | Rickettsiales | Candidatus_Captivus | Candidatus_Captivus | 0.000 | 0.006 | 0.040 | 0.000 | 0.000 | 0.000 |
| Proteobacteria | Alphaproteobacteria | Rickettsiales | Candidatus_Hepatincola | Candidatus_Hepatincola | 0.008 | 0.000 | 0.040 | 0.000 | 0.000 | 0.000 |
| Proteobacteria | Alphaproteobacteria | Rhizobiales_3 | Candidatus_Liberibacter | Candidatus_Liberibacter | 0.000 | 0.000 | 0.004 | 0.000 | 0.000 | 0.000 |
| Proteobacteria | Alphaproteobacteria | Rickettsiales | Candidatus_Midichloria | Candidatus_Midichloria | 0.000 | 0.000 | 0.006 | 0.000 | 0.000 | 0.000 |
| Proteobacteria | Alphaproteobacteria | Rickettsiales | Candidatus_Odyssella | Candidatus_Odyssella | 0.000 | 0.000 | 0.004 | 0.000 | 0.000 | 0.000 |
| Proteobacteria | Alphaproteobacteria | Caulobacterales | Caulobacteraceae | Asticcacaulis | 0.008 | 0.006 | 0.010 | 0.012 | 0.000 | 0.000 |
| Proteobacteria | Alphaproteobacteria | Caulobacterales | Caulobacteraceae | Atopostipes | 0.016 | 0.000 | 0.008 | 0.000 | 0.000 | 0.006 |
| Proteobacteria | Alphaproteobacteria | Caulobacterales | Caulobacteraceae | Brevundimonas | 0.024 | 0.043 | 0.073 | 0.006 | 0.000 | 0.006 |
| Proteobacteria | Alphaproteobacteria | Caulobacterales | Caulobacteraceae | Caulobacter | 0.065 | 0.012 | 0.014 | 0.000 | 0.000 | 0.012 |
| Proteobacteria | Alphaproteobacteria | Caulobacterales | Caulobacteraceae | Phocea | 0.000 | 0.000 | 0.000 | 0.000 | 0.000 | 0.006 |
| Proteobacteria | Alphaproteobacteria | Rhizobiales | Chelatococcaceae | Chelatococcus_2 | 0.000 | 0.000 | 0.006 | 0.000 | 0.000 | 0.000 |
| Proteobacteria | Gammaproteobacteria | Chromatiales | Chromatiaceae | Thiorhodovibrio | 0.000 | 0.000 | 0.004 | 0.000 | 0.000 | 0.000 |
| Proteobacteria | Gammaproteobacteria_0 | Chromatiales | Chromatiaceae | Rheinheimera_3 | 0.000 | 0.000 | 0.008 | 0.000 | 0.000 | 0.000 |
| Proteobacteria | Gammaproteobacteria_1 | Chromatiales | Chromatiaceae | Alisitpes_IV | 2.872 | 0.826 | 0.111 | 0.371 | 0.075 | 0.145 |
| Proteobacteria | Gammaproteobacteria_1 | Chromatiales | Chromatiaceae | Alkalimonas | 0.000 | 0.000 | 0.006 | 0.000 | 0.000 | 0.000 |
| Proteobacteria | Gammaproteobacteria_1 | Chromatiales | Chromatiaceae | Rheinheimera_4 | 0.000 | 0.000 | 0.004 | 0.000 | 0.000 | 0.000 |
| Proteobacteria | Gammaproteobacteria_1 | Chromatiales | Chromatiaceae | Rhizobium | 0.081 | 0.080 | 0.010 | 0.036 | 0.045 | 0.163 |
| Proteobacteria | Gammaproteobacteria_1 | Chromatiales | Chromatiaceae | Rhizobium-Agrobacterium | 0.041 | 0.166 | 0.234 | 0.024 | 0.035 | 0.151 |
| Proteobacteria | Gammaproteobacteria_2 | Chromatiales | Chromatiaceae | Allochromatium | 0.000 | 0.000 | 0.014 | 0.000 | 0.000 | 0.000 |
| Proteobacteria | Gammaproteobacteria_2 | Chromatiales | Chromatiaceae | Chromatium | 0.000 | 0.000 | 0.004 | 0.000 | 0.000 | 0.000 |
| Proteobacteria | Gammaproteobacteria_2 | Chromatiales | Chromatiaceae | Lamprocystis | 0.000 | 0.000 | 0.006 | 0.000 | 0.000 | 0.000 |
| Proteobacteria | Gammaproteobacteria_2 | Chromatiales | Chromatiaceae | Marichromatium | 0.000 | 0.000 | 0.004 | 0.000 | 0.000 | 0.000 |
| Proteobacteria | Gammaproteobacteria_2 | Chromatiales | Chromatiaceae | Nitrosomonas | 0.008 | 0.000 | 0.036 | 0.000 | 0.000 | 0.000 |
| Proteobacteria | Gammaproteobacteria_2 | Chromatiales | Chromatiaceae | Thiocapsa | 0.000 | 0.000 | 0.010 | 0.000 | 0.000 | 0.000 |
| Proteobacteria | Gammaproteobacteria_2 | Chromatiales | Chromatiaceae | Thiocystis_1 | 0.000 | 0.000 | 0.006 | 0.000 | 0.000 | 0.000 |

**Table S9 continued.**

| | | | | | | | | | | |
|---|---|---|---|---|---|---|---|---|---|---|
| Proteobacteria | Gammaproteobacteria_2 | Chromatiales | Chromatiaceae | Thiorhodococcus_1 | 0.000 | 0.000 | 0.004 | 0.000 | 0.000 | 0.000 |
| Proteobacteria | Gammaproteobacteria_2 | Chromatiales | Chromatiaceae | Thiorhodococcus_2 | 0.000 | 0.000 | 0.006 | 0.000 | 0.000 | 0.000 |
| Proteobacteria | Betaproteobacteria | Neisseriales | Chromobacteriaceae | Chitinilyticum | 0.000 | 0.000 | 0.004 | 0.000 | 0.000 | 0.000 |
| Proteobacteria | Betaproteobacteria | Neisseriales | Chromobacteriaceae | Chromobacterium | 0.000 | 0.000 | 0.006 | 0.000 | 0.000 | 0.000 |
| Proteobacteria | Betaproteobacteria | Neisseriales | Chromobacteriaceae | Chryseobacterium | 0.000 | 0.000 | 0.002 | 0.000 | 0.000 | 0.006 |
| Proteobacteria | Betaproteobacteria | Neisseriales | Chromobacteriaceae | Chryseobacterium_1 | 0.000 | 0.000 | 0.048 | 0.000 | 0.000 | 0.000 |
| Proteobacteria | Betaproteobacteria | Neisseriales | Chromobacteriaceae | Chryseobacterium_10 | 0.000 | 0.000 | 0.004 | 0.000 | 0.000 | 0.000 |
| Proteobacteria | Betaproteobacteria | Neisseriales | Chromobacteriaceae | Chryseobacterium_11 | 0.000 | 0.000 | 0.036 | 0.000 | 0.000 | 0.000 |
| Proteobacteria | Betaproteobacteria | Neisseriales | Chromobacteriaceae | Chryseobacterium_8 | 0.000 | 0.000 | 0.020 | 0.000 | 0.000 | 0.000 |
| Proteobacteria | Deltaproteobacteria | Rs-K70_termite_group | Cluster_1 | Cohaesibacter | 0.000 | 0.000 | 0.006 | 0.000 | 0.000 | 0.000 |
| Proteobacteria | Gammaproteobacteria_1 | Alteromonadales_1 | Colwelliaceaea | Colwellia_1 | 0.000 | 0.006 | 0.054 | 0.000 | 0.005 | 0.006 |
| Proteobacteria | Gammaproteobacteria_1 | Alteromonadales_1 | Colwelliaceaea | Colwellia_2 | 0.000 | 0.000 | 0.012 | 0.000 | 0.000 | 0.000 |
| Proteobacteria | Gammaproteobacteria_1 | Alteromonadales_1 | Colwelliaceaea | Thalassomonas | 0.000 | 0.000 | 0.016 | 0.000 | 0.000 | 0.000 |
| Proteobacteria | Betaproteobacteria | Burkholderiales | Comamonadaceae | Acidovorax | 0.033 | 0.012 | 0.000 | 0.006 | 0.000 | 0.012 |
| Proteobacteria | Betaproteobacteria | Burkholderiales | Comamonadaceae | Acidovorax-Hylemonella | 0.000 | 0.006 | 0.042 | 0.006 | 0.000 | 0.000 |
| Proteobacteria | Betaproteobacteria | Burkholderiales | Comamonadaceae | Acidovorax-Verminephrobacter | 0.041 | 0.006 | 0.153 | 0.000 | 0.005 | 0.000 |
| Proteobacteria | Betaproteobacteria | Burkholderiales | Comamonadaceae | Alicycliphilus | 0.000 | 0.000 | 0.026 | 0.000 | 0.000 | 0.000 |
| Proteobacteria | Betaproteobacteria | Burkholderiales | Comamonadaceae | Aquabacterium | 0.065 | 0.018 | 0.111 | 0.000 | 0.005 | 0.012 |
| Proteobacteria | Betaproteobacteria | Burkholderiales | Comamonadaceae | Aquamonas | 0.016 | 0.000 | 0.022 | 0.006 | 0.000 | 0.000 |
| Proteobacteria | Betaproteobacteria | Burkholderiales | Comamonadaceae | Azonexus_2 | 0.000 | 0.000 | 0.004 | 0.000 | 0.005 | 0.000 |
| Proteobacteria | Betaproteobacteria | Burkholderiales | Comamonadaceae | Brachymonas_1 | 0.008 | 0.000 | 0.040 | 0.000 | 0.000 | 0.000 |
| Proteobacteria | Betaproteobacteria | Burkholderiales | Comamonadaceae | Brachymonas_2 | 0.008 | 0.000 | 0.010 | 0.000 | 0.000 | 0.000 |
| Proteobacteria | Betaproteobacteria | Burkholderiales | Comamonadaceae | Caenimonas | 0.008 | 0.000 | 0.020 | 0.000 | 0.000 | 0.000 |
| Proteobacteria | Betaproteobacteria | Burkholderiales | Comamonadaceae | Comamonas | 0.008 | 0.000 | 0.000 | 0.000 | 0.000 | 0.000 |
| Proteobacteria | Betaproteobacteria | Burkholderiales | Comamonadaceae | Comamonas_1 | 0.008 | 0.000 | 0.036 | 0.000 | 0.000 | 0.000 |
| Proteobacteria | Betaproteobacteria | Burkholderiales | Comamonadaceae | Comamonas_2 | 0.000 | 0.000 | 0.008 | 0.000 | 0.000 | 0.000 |
| Proteobacteria | Betaproteobacteria | Burkholderiales | Comamonadaceae | Comamonas_3 | 0.008 | 0.000 | 0.016 | 0.000 | 0.000 | 0.000 |
| Proteobacteria | Betaproteobacteria | Burkholderiales | Comamonadaceae | Comamonas_4 | 0.000 | 0.000 | 0.014 | 0.000 | 0.000 | 0.000 |
| Proteobacteria | Betaproteobacteria | Burkholderiales | Comamonadaceae | Comamonas_5 | 0.008 | 0.006 | 0.042 | 0.000 | 0.005 | 0.000 |
| Proteobacteria | Betaproteobacteria | Burkholderiales | Comamonadaceae | Comamonas_6 | 0.016 | 0.006 | 0.022 | 0.000 | 0.000 | 0.000 |
| Proteobacteria | Betaproteobacteria | Burkholderiales | Comamonadaceae | Comamonas_7 | 0.000 | 0.000 | 0.006 | 0.000 | 0.000 | 0.000 |
| Proteobacteria | Betaproteobacteria | Burkholderiales | Comamonadaceae | Comamonas_8 | 0.008 | 0.000 | 0.034 | 0.000 | 0.005 | 0.000 |
| Proteobacteria | Betaproteobacteria | Burkholderiales | Comamonadaceae | Comamonas_R | 0.000 | 0.000 | 0.004 | 0.000 | 0.000 | 0.000 |
| Proteobacteria | Betaproteobacteria | Burkholderiales | Comamonadaceae | Curvibacter | 0.008 | 0.000 | 0.014 | 0.000 | 0.000 | 0.000 |
| Proteobacteria | Betaproteobacteria | Burkholderiales | Comamonadaceae | Delftia | 0.000 | 0.000 | 0.040 | 0.006 | 0.000 | 0.000 |
| Proteobacteria | Betaproteobacteria | Burkholderiales | Comamonadaceae | Diaphorobacter | 0.000 | 0.000 | 0.024 | 0.000 | 0.000 | 0.006 |
| Proteobacteria | Betaproteobacteria | Burkholderiales | Comamonadaceae | Geothrix | 0.000 | 0.000 | 0.022 | 0.000 | 0.000 | 0.000 |
| Proteobacteria | Betaproteobacteria | Burkholderiales | Comamonadaceae | Giesbergeria | 0.008 | 0.000 | 0.004 | 0.000 | 0.000 | 0.000 |
| Proteobacteria | Betaproteobacteria | Burkholderiales | Comamonadaceae | Hydrogenophaga_1 | 0.008 | 0.000 | 0.054 | 0.000 | 0.000 | 0.000 |
| Proteobacteria | Betaproteobacteria | Burkholderiales | Comamonadaceae | Hydrogenophaga_2 | 0.000 | 0.000 | 0.012 | 0.000 | 0.000 | 0.000 |

**Table S9 continued.**

| | | | | | | | | | | |
|---|---|---|---|---|---|---|---|---|---|---|
| Proteobacteria | Betaproteobacteria | Burkholderiales | Comamonadaceae | Hydrogenophaga_3 | 0.000 | 0.000 | 0.012 | 0.000 | 0.000 | 0.000 |
| Proteobacteria | Betaproteobacteria | Burkholderiales | Comamonadaceae | Hydrogenophaga_4 | 0.000 | 0.000 | 0.004 | 0.000 | 0.000 | 0.000 |
| Proteobacteria | Betaproteobacteria | Burkholderiales | Comamonadaceae | Hydrogenophaga_5 | 0.000 | 0.006 | 0.020 | 0.000 | 0.000 | 0.000 |
| Proteobacteria | Betaproteobacteria | Burkholderiales | Comamonadaceae | Hydrogenophilus | 0.000 | 0.000 | 0.006 | 0.000 | 0.000 | 0.000 |
| Proteobacteria | Betaproteobacteria | Burkholderiales | Comamonadaceae | Hymenobacter | 0.024 | 0.012 | 0.077 | 0.000 | 0.000 | 0.000 |
| Proteobacteria | Betaproteobacteria | Burkholderiales | Comamonadaceae | Ideonella_1 | 0.000 | 0.000 | 0.004 | 0.000 | 0.000 | 0.000 |
| Proteobacteria | Betaproteobacteria | Burkholderiales | Comamonadaceae | Ideonella_2 | 0.008 | 0.000 | 0.012 | 0.000 | 0.000 | 0.000 |
| Proteobacteria | Betaproteobacteria | Burkholderiales | Comamonadaceae | Ignatzschineria | 0.000 | 0.000 | 0.004 | 0.000 | 0.000 | 0.000 |
| Proteobacteria | Betaproteobacteria | Burkholderiales | Comamonadaceae | Inquilinus | 0.008 | 0.000 | 0.006 | 0.000 | 0.000 | 0.000 |
| Proteobacteria | Betaproteobacteria | Burkholderiales | Comamonadaceae | Kitasatospora | 0.008 | 0.000 | 0.050 | 0.000 | 0.010 | 0.000 |
| Proteobacteria | Betaproteobacteria | Burkholderiales | Comamonadaceae | Lapillicoccus | 0.000 | 0.000 | 0.006 | 0.000 | 0.000 | 0.000 |
| Proteobacteria | Betaproteobacteria | Burkholderiales | Comamonadaceae | Leptothrix_1 | 0.000 | 0.000 | 0.004 | 0.000 | 0.000 | 0.000 |
| Proteobacteria | Betaproteobacteria | Burkholderiales | Comamonadaceae | Leptothrix_2 | 0.008 | 0.000 | 0.050 | 0.000 | 0.000 | 0.006 |
| Proteobacteria | Betaproteobacteria | Burkholderiales | Comamonadaceae | Leptotrichia | 0.008 | 0.000 | 0.103 | 0.000 | 0.000 | 0.000 |
| Proteobacteria | Betaproteobacteria | Burkholderiales | Comamonadaceae | Macromonas | 0.000 | 0.000 | 0.004 | 0.000 | 0.000 | 0.000 |
| Proteobacteria | Betaproteobacteria | Burkholderiales | Comamonadaceae | Malikia | 0.000 | 0.000 | 0.006 | 0.000 | 0.000 | 0.000 |
| Proteobacteria | Betaproteobacteria | Burkholderiales | Comamonadaceae | Methylibium_1 | 0.130 | 0.012 | 0.091 | 0.006 | 0.005 | 0.000 |
| Proteobacteria | Betaproteobacteria | Burkholderiales | Comamonadaceae | Methylibium_2 | 0.000 | 0.000 | 0.006 | 0.000 | 0.000 | 0.006 |
| Proteobacteria | Betaproteobacteria | Burkholderiales | Comamonadaceae | Mitsuokella | 0.000 | 0.000 | 0.010 | 0.000 | 0.000 | 0.000 |
| Proteobacteria | Betaproteobacteria | Burkholderiales | Comamonadaceae | Owenweeksia | 0.000 | 0.000 | 0.022 | 0.000 | 0.000 | 0.000 |
| Proteobacteria | Betaproteobacteria | Burkholderiales | Comamonadaceae | Pelospora | 0.000 | 0.000 | 0.006 | 0.006 | 0.000 | 0.000 |
| Proteobacteria | Betaproteobacteria | Burkholderiales | Comamonadaceae | Polaromonas | 0.000 | 0.000 | 0.066 | 0.000 | 0.000 | 0.000 |
| Proteobacteria | Betaproteobacteria | Burkholderiales | Comamonadaceae | Pseudacidovorax | 0.000 | 0.000 | 0.004 | 0.000 | 0.000 | 0.000 |
| Proteobacteria | Betaproteobacteria | Burkholderiales | Comamonadaceae | Pseudospirillum | 0.024 | 0.000 | 0.048 | 0.000 | 0.000 | 0.000 |
| Proteobacteria | Betaproteobacteria | Burkholderiales | Comamonadaceae | Raoultibacter | 0.033 | 0.000 | 0.000 | 0.000 | 0.000 | 0.000 |
| Proteobacteria | Betaproteobacteria | Burkholderiales | Comamonadaceae | Rhodomicrobium | 0.000 | 0.000 | 0.004 | 0.000 | 0.000 | 0.000 |
| Proteobacteria | Betaproteobacteria | Burkholderiales | Comamonadaceae | Rubrivivax | 0.008 | 0.000 | 0.010 | 0.000 | 0.005 | 0.000 |
| Proteobacteria | Betaproteobacteria | Burkholderiales | Comamonadaceae | Rudaibacter | 0.000 | 0.000 | 0.002 | 0.000 | 0.000 | 0.000 |
| Proteobacteria | Betaproteobacteria | Burkholderiales | Comamonadaceae | Schlegelella | 0.000 | 0.012 | 0.008 | 0.000 | 0.000 | 0.000 |
| Proteobacteria | Betaproteobacteria | Burkholderiales | Comamonadaceae | Simplicispira_1 | 0.000 | 0.000 | 0.006 | 0.000 | 0.000 | 0.000 |
| Proteobacteria | Betaproteobacteria | Burkholderiales | Comamonadaceae | Simplicispira_3 | 0.000 | 0.000 | 0.006 | 0.000 | 0.000 | 0.000 |
| Proteobacteria | Betaproteobacteria | Burkholderiales | Comamonadaceae | Singulisphaera | 0.000 | 0.000 | 0.073 | 0.000 | 0.000 | 0.000 |
| Proteobacteria | Betaproteobacteria | Burkholderiales | Comamonadaceae | Sphingobacterium_1 | 0.000 | 0.000 | 0.022 | 0.000 | 0.000 | 0.000 |
| Proteobacteria | Betaproteobacteria | Burkholderiales | Comamonadaceae | Tepidicella | 0.000 | 0.000 | 0.006 | 0.000 | 0.000 | 0.000 |
| Proteobacteria | Betaproteobacteria | Burkholderiales | Comamonadaceae | Tepidimonas | 0.000 | 0.000 | 0.018 | 0.000 | 0.005 | 0.000 |
| Proteobacteria | Betaproteobacteria | Burkholderiales | Comamonadaceae | Uncultured_16 | 0.000 | 0.000 | 0.067 | 0.000 | 0.000 | 0.000 |
| Proteobacteria | Betaproteobacteria | Burkholderiales | Comamonadaceae | Uncultured_17 | 0.024 | 0.000 | 0.165 | 0.000 | 0.000 | 0.000 |
| Proteobacteria | Betaproteobacteria | Burkholderiales | Comamonadaceae | Uncultured_19 | 0.000 | 0.006 | 0.101 | 0.000 | 0.000 | 0.000 |
| Proteobacteria | Betaproteobacteria | Burkholderiales | Comamonadaceae | Uncultured_31 | 0.610 | 0.043 | 2.257 | 0.030 | 0.005 | 0.006 |
| Proteobacteria | Betaproteobacteria | Burkholderiales | Comamonadaceae | Uncultured_d | 0.000 | 0.000 | 0.004 | 0.000 | 0.000 | 0.000 |

**Table S9 continued.**

| | | | | | | | | | | |
|---|---|---|---|---|---|---|---|---|---|---|
| Proteobacteria | Betaproteobacteria | Burkholderiales | Comamonadaceae | Variovorax_2 | 0.024 | 0.000 | 0.040 | 0.000 | 0.005 | 0.000 |
| Proteobacteria | Betaproteobacteria | Burkholderiales | Comamonadaceae | Variovorax_a | 0.000 | 0.000 | 0.004 | 0.000 | 0.000 | 0.000 |
| Proteobacteria | Betaproteobacteria | Burkholderiales | Comamonadaceae | Variovorax_b | 0.000 | 0.000 | 0.004 | 0.000 | 0.000 | 0.000 |
| Proteobacteria | Betaproteobacteria | Burkholderiales | Comamonadaceae | Veillonella | 0.000 | 0.000 | 0.048 | 0.006 | 0.000 | 0.000 |
| Proteobacteria | Betaproteobacteria | Burkholderiales | Comamonadaceae | Xylanimonas | 0.008 | 0.000 | 0.008 | 0.000 | 0.000 | 0.000 |
| Proteobacteria | Betaproteobacteria | Burkholderiales | Comamonadaceae | Xylophilus | 0.000 | 0.000 | 0.004 | 0.000 | 0.000 | 0.000 |
| Proteobacteria | Gammaproteobacteria_2 | Legionellales | Coxiellaceae | Aquicella | 0.008 | 0.000 | 0.054 | 0.000 | 0.000 | 0.000 |
| Proteobacteria | Gammaproteobacteria_2 | Legionellales | Coxiellaceae | Coxiella | 0.008 | 0.000 | 0.026 | 0.000 | 0.000 | 0.006 |
| Proteobacteria | Gammaproteobacteria_2 | Legionellales | Coxiellaceae | Rikenella_1 | 0.000 | 0.000 | 0.014 | 0.000 | 0.000 | 0.000 |
| Proteobacteria | Deltaproteobacteria | Myxococcales | Cystobacteraceae | Anaeromyxobacter | 0.016 | 0.000 | 0.016 | 0.000 | 0.000 | 0.000 |
| Proteobacteria | Deltaproteobacteria | Myxococcales | Cystobacteraceae | Hyalangium | 0.000 | 0.000 | 0.004 | 0.000 | 0.000 | 0.000 |
| Proteobacteria | Deltaproteobacteria | Desulfovibrionales | Desufovibrionaceae | Desulfovibrio | 0.041 | 0.000 | 0.000 | 0.000 | 0.005 | 0.000 |
| Proteobacteria | Deltaproteobacteria | Desulfovibrionales | Desufovibrionaceae | Desulfovibrio_5 | 0.000 | 0.000 | 0.113 | 0.000 | 0.000 | 0.000 |
| Proteobacteria | Deltaproteobacteria | Desulfarculales | Desulfarculaceae | Desulfarculus | 0.008 | 0.000 | 0.012 | 0.000 | 0.000 | 0.000 |
| Proteobacteria | Deltaproteobacteria | Desulfobacterales | Desulfobacteraceae | Desulfatiferula | 0.000 | 0.000 | 0.016 | 0.000 | 0.000 | 0.000 |
| Proteobacteria | Deltaproteobacteria | Desulfobacterales | Desulfobacteraceae | Desulfobacter | 0.000 | 0.000 | 0.020 | 0.000 | 0.000 | 0.000 |
| Proteobacteria | Deltaproteobacteria | Desulfobacterales | Desulfobacteraceae | Desulfobacterium | 0.000 | 0.000 | 0.010 | 0.000 | 0.000 | 0.000 |
| Proteobacteria | Deltaproteobacteria | Desulfobacterales | Desulfobacteraceae | Desulfobacterium_1 | 0.000 | 0.000 | 0.012 | 0.000 | 0.000 | 0.000 |
| Proteobacteria | Deltaproteobacteria | Desulfobacterales | Desulfobacteraceae | Desulfobacterium_2 | 0.000 | 0.000 | 0.008 | 0.000 | 0.000 | 0.000 |
| Proteobacteria | Deltaproteobacteria | Desulfobacterales | Desulfobacteraceae | Desulfobacula | 0.000 | 0.000 | 0.010 | 0.000 | 0.000 | 0.000 |
| Proteobacteria | Deltaproteobacteria | Desulfobacterales | Desulfobacteraceae | Desulfobotulus | 0.000 | 0.000 | 0.004 | 0.000 | 0.000 | 0.000 |
| Proteobacteria | Deltaproteobacteria | Desulfobacterales | Desulfobacteraceae | Desulfocella | 0.000 | 0.000 | 0.004 | 0.000 | 0.000 | 0.000 |
| Proteobacteria | Deltaproteobacteria | Desulfobacterales | Desulfobacteraceae | Desulfococcus_1 | 0.000 | 0.000 | 0.006 | 0.000 | 0.000 | 0.000 |
| Proteobacteria | Deltaproteobacteria | Desulfobacterales | Desulfobacteraceae | Desulfococcus_2 | 0.000 | 0.000 | 0.004 | 0.000 | 0.000 | 0.000 |
| Proteobacteria | Deltaproteobacteria | Desulfobacterales | Desulfobacteraceae | Desulfofaba_1 | 0.000 | 0.000 | 0.004 | 0.000 | 0.000 | 0.000 |
| Proteobacteria | Deltaproteobacteria | Desulfobacterales | Desulfobacteraceae | Desulfofaba_2 | 0.000 | 0.000 | 0.004 | 0.000 | 0.000 | 0.000 |
| Proteobacteria | Deltaproteobacteria | Desulfobacterales | Desulfobacteraceae | Desulfofrigus | 0.000 | 0.000 | 0.010 | 0.000 | 0.000 | 0.000 |
| Proteobacteria | Deltaproteobacteria | Desulfobacterales | Desulfobacteraceae | Desulfonema | 0.000 | 0.000 | 0.008 | 0.000 | 0.000 | 0.000 |
| Proteobacteria | Deltaproteobacteria | Desulfobacterales | Desulfobacteraceae | Desulfosarcina | 0.000 | 0.000 | 0.004 | 0.000 | 0.000 | 0.000 |
| Proteobacteria | Deltaproteobacteria | Desulfobacterales | Desulfobacteraceae | Desulfotignum | 0.000 | 0.000 | 0.008 | 0.000 | 0.000 | 0.000 |
| Proteobacteria | Deltaproteobacteria | Desulfobacterales | Desulfobulbaceae | Desulfobulbus | 0.000 | 0.000 | 0.042 | 0.000 | 0.000 | 0.000 |
| Proteobacteria | Deltaproteobacteria | Desulfobacterales | Desulfobulbaceae | Desulfocapsa_1 | 0.000 | 0.000 | 0.016 | 0.000 | 0.000 | 0.000 |
| Proteobacteria | Deltaproteobacteria | Desulfobacterales | Desulfobulbaceae | Desulfocapsa_2 | 0.000 | 0.000 | 0.006 | 0.000 | 0.000 | 0.000 |
| Proteobacteria | Deltaproteobacteria | Desulfobacterales | Desulfobulbaceae | Desulfofustis | 0.000 | 0.000 | 0.004 | 0.000 | 0.000 | 0.000 |
| Proteobacteria | Deltaproteobacteria | Desulfobacterales | Desulfobulbaceae | Desulfopila | 0.000 | 0.000 | 0.008 | 0.000 | 0.000 | 0.000 |
| Proteobacteria | Deltaproteobacteria | Desulfobacterales | Desulfobulbaceae | Desulforhopalus | 0.000 | 0.000 | 0.014 | 0.000 | 0.000 | 0.000 |
| Proteobacteria | Deltaproteobacteria | Desulfobacterales | Desulfobulbaceae | Desulfotalea | 0.000 | 0.000 | 0.012 | 0.000 | 0.000 | 0.000 |
| Proteobacteria | Deltaproteobacteria | Desulfobacterales | Desulfobulbaceae | Desulfurivibrio | 0.000 | 0.000 | 0.008 | 0.000 | 0.000 | 0.000 |
| Proteobacteria | Deltaproteobacteria | Desulfobacterales | Desulfobulbaceae | Dissulfuribacter | 0.000 | 0.000 | 0.006 | 0.000 | 0.000 | 0.000 |
| Proteobacteria | Deltaproteobacteria | Desulfovibrionales | Desulfohalobiaceae_1 | Desulfonatronospira | 0.000 | 0.000 | 0.004 | 0.000 | 0.000 | 0.000 |

**Table S9 continued.**

| | | | | | | | | | | |
|---|---|---|---|---|---|---|---|---|---|---|
| Proteobacteria | Deltaproteobacteria | Desulfovibrionales | Desulfohalobiaceae_1 | Desulfonatronovibrio | 0.000 | 0.000 | 0.004 | 0.000 | 0.000 | 0.000 |
| Proteobacteria | Deltaproteobacteria | Desulfovibrionales | Desulfohalobiaceae_2 | Desulfohalobium | 0.000 | 0.000 | 0.004 | 0.000 | 0.000 | 0.000 |
| Proteobacteria | Deltaproteobacteria | Desulfovibrionales | Desulfohalobiaceae_2 | Desulfonauticus | 0.000 | 0.000 | 0.004 | 0.000 | 0.000 | 0.000 |
| Proteobacteria | Deltaproteobacteria | Desulfovibrionales | Desulfohalobiaceae_2 | Desulfothermus | 0.000 | 0.000 | 0.004 | 0.000 | 0.000 | 0.000 |
| Proteobacteria | Deltaproteobacteria | Desulfovibrionales | Desulfohalobiaceae_2 | Desulfovermiculus | 0.000 | 0.000 | 0.008 | 0.000 | 0.000 | 0.000 |
| Proteobacteria | Deltaproteobacteria | Desulfovibrionales | Desulfomicrobiaceae | Desulfomicrobium | 0.008 | 0.000 | 0.018 | 0.012 | 0.000 | 0.000 |
| Proteobacteria | Deltaproteobacteria | Desulfovibrionales | Desulfonatronaceae | Desulfonatronum | 0.000 | 0.000 | 0.006 | 0.000 | 0.000 | 0.000 |
| Proteobacteria | Deltaproteobacteria | Desulfovibrionales | Desulfovibrionaceae | Bilophila | 0.000 | 0.000 | 0.010 | 0.000 | 0.000 | 0.000 |
| Proteobacteria | Deltaproteobacteria | Desulfovibrionales | Desulfovibrionaceae | Desulfovibrio_2 | 0.000 | 0.000 | 0.008 | 0.000 | 0.000 | 0.000 |
| Proteobacteria | Deltaproteobacteria | Desulfovibrionales | Desulfovibrionaceae | Desulfovibrio_4 | 0.000 | 0.000 | 0.024 | 0.000 | 0.000 | 0.000 |
| Proteobacteria | Deltaproteobacteria | Desulfovibrionales | Desulfovibrionaceae | Desulfurobacterium_1 | 0.000 | 0.000 | 0.016 | 0.000 | 0.000 | 0.000 |
| Proteobacteria | Deltaproteobacteria | Desulfovibrionales | Desulfovibrionaceae | Gut_cluster_3 | 1.594 | 0.388 | 0.405 | 0.108 | 0.015 | 0.042 |
| Proteobacteria | Deltaproteobacteria | Desulfovibrionales | Desulfovibrionaceae | Gut_cluster_4 | 0.041 | 0.037 | 0.262 | 0.006 | 0.000 | 0.000 |
| Proteobacteria | Deltaproteobacteria | Desulfovibrionales | Desulfovibrionaceae | Mammalian_cluster | 0.000 | 0.000 | 0.228 | 0.000 | 0.000 | 0.000 |
| Proteobacteria | Deltaproteobacteria | Desulfovibrionales | Desulfovibrionaceae | Termite_cockroach_cluster_1 | 0.179 | 0.092 | 0.087 | 0.030 | 0.005 | 0.006 |
| Proteobacteria | Deltaproteobacteria | Desulfurellales | Desulfurellaceae | Desulfurella | 0.000 | 0.000 | 0.004 | 0.000 | 0.000 | 0.000 |
| Proteobacteria | Deltaproteobacteria | Desulfuromonadales | Desulfuromonadaceae | Desulfuromonas_1 | 0.000 | 0.000 | 0.004 | 0.000 | 0.000 | 0.000 |
| Proteobacteria | Deltaproteobacteria | Desulfuromonadales | Desulfuromonadaceae | Desulfuromonas_3 | 0.008 | 0.000 | 0.020 | 0.000 | 0.000 | 0.000 |
| Proteobacteria | Deltaproteobacteria | Desulfuromonadales | Desulfuromonadaceae | Pelomonas | 0.114 | 0.018 | 0.060 | 0.012 | 0.000 | 0.000 |
| Proteobacteria | Deltaproteobacteria | Desulfuromonadales | Desulfuromonadaceae_1 | Desulfuromusa_1 | 0.000 | 0.000 | 0.006 | 0.000 | 0.000 | 0.000 |
| Proteobacteria | Gammaproteobacteria | Chromatiales | Ectothiorhodospiraceae | Ectothiorhodospira | 0.000 | 0.000 | 0.012 | 0.000 | 0.000 | 0.000 |
| Proteobacteria | Gammaproteobacteria | Chromatiales | Ectothiorhodospiraceae | Tianweitania | 0.008 | 0.000 | 0.000 | 0.000 | 0.000 | 0.000 |
| Proteobacteria | Gammaproteobacteria_2 | Chromatiales | Ectothiorhodospiraceae | Thioalkalispira | 0.008 | 0.000 | 0.006 | 0.000 | 0.000 | 0.000 |
| Proteobacteria | Gammaproteobacteria_2 | Chromatiales | Ectothiorhodospiraceae_1 | Thioalkalivibrio_2 | 0.000 | 0.000 | 0.004 | 0.000 | 0.000 | 0.000 |
| Proteobacteria | Gammaproteobacteria_2 | Chromatiales | Ectothiorhodospiraceae_1 | Thiobacillus_1 | 0.000 | 0.000 | 0.044 | 0.006 | 0.005 | 0.000 |
| Proteobacteria | Gammaproteobacteria_2 | Chromatiales | Ectothiorhodospiraceae_2 | Aquasalimonas | 0.000 | 0.000 | 0.004 | 0.000 | 0.000 | 0.000 |
| Proteobacteria | Gammaproteobacteria_2 | Chromatiales | Ectothiorhodospiraceae_3 | Thiohalospira | 0.000 | 0.000 | 0.004 | 0.000 | 0.000 | 0.000 |
| Proteobacteria | Gammaproteobacteria | Enterobacterales | Enterobacteriaceae | Cedecea | 0.033 | 0.025 | 0.000 | 0.000 | 0.010 | 0.006 |
| Proteobacteria | Gammaproteobacteria | Enterobacterales | Enterobacteriaceae | Cronobacter | 0.089 | 0.314 | 0.155 | 0.198 | 0.274 | 0.206 |
| Proteobacteria | Gammaproteobacteria | Enterobacterales | Enterobacteriaceae | Mangrovibacter | 0.000 | 0.012 | 0.002 | 0.006 | 0.000 | 0.012 |
| Proteobacteria | Gammaproteobacteria | Enterobacterales | Enterobacteriaceae | Pluralibacter | 0.000 | 0.006 | 0.002 | 0.024 | 0.020 | 0.006 |
| Proteobacteria | Gammaproteobacteria | Enterobacterales | Enterobacteriaceae | Pseudoclavibacter | 0.000 | 0.000 | 0.000 | 0.006 | 0.000 | 0.000 |
| Proteobacteria | Gammaproteobacteria | Enterobacterales | Enterobacteriaceae | Shinella_genera_incertae_sedis | 0.000 | 0.000 | 0.030 | 0.000 | 0.000 | 0.006 |
| Proteobacteria | Gammaproteobacteria | Enterobacterales | Enterobacteriaceae | Siccibacter | 0.000 | 0.006 | 0.004 | 0.000 | 0.000 | 0.000 |
| Proteobacteria | Gammaproteobacteria | Enterobacteriales | Enterobacteriaceae | Krasilnikovia | 0.000 | 0.000 | 0.004 | 0.000 | 0.000 | 0.000 |
| Proteobacteria | Gammaproteobacteria | Enterobacteriales | Enterobacteriaceae | Lelliottia | 0.000 | 0.006 | 0.036 | 0.012 | 0.025 | 0.012 |
| Proteobacteria | Gammaproteobacteria | Enterobacteriales | Enterobacteriaceae | Shimia | 0.000 | 0.000 | 0.008 | 0.000 | 0.000 | 0.000 |
| Proteobacteria | Gammaproteobacteria_1 | Enterobacteriales | Enterobacteriaceae | Brenneria | 0.000 | 0.006 | 0.024 | 0.006 | 0.000 | 0.000 |
| Proteobacteria | Gammaproteobacteria_1 | Enterobacteriales | Enterobacteriaceae | Brenneria-Erwinia-Pectobacterium | 0.366 | 5.379 | 0.159 | 3.876 | 5.049 | 5.484 |
| Proteobacteria | Gammaproteobacteria_1 | Enterobacteriales | Enterobacteriaceae | Budvicia | 0.000 | 0.000 | 0.006 | 0.000 | 0.000 | 0.000 |

**Table S9 continued.**

| | | | | | | | | | | |
|---|---|---|---|---|---|---|---|---|---|---|
| Proteobacteria | Gammaproteobacteria_1 | Enterobacteriales | Enterobacteriaceae | Buttiauxella | 0.114 | 1.479 | 0.171 | 0.767 | 0.927 | 0.369 |
| Proteobacteria | Gammaproteobacteria_1 | Enterobacteriales | Enterobacteriaceae | Candidatus_Rohrkolberia | 0.000 | 0.006 | 0.010 | 0.000 | 0.010 | 0.000 |
| Proteobacteria | Gammaproteobacteria_1 | Enterobacteriales | Enterobacteriaceae | Citrobacter | 0.106 | 0.499 | 0.145 | 0.389 | 0.344 | 0.254 |
| Proteobacteria | Gammaproteobacteria_1 | Enterobacteriales | Enterobacteriaceae | Citrobacter_1 | 0.399 | 1.670 | 0.071 | 0.725 | 0.424 | 0.527 |
| Proteobacteria | Gammaproteobacteria_1 | Enterobacteriales | Enterobacteriaceae | Citrobacter_2 | 0.309 | 1.793 | 0.073 | 0.875 | 0.558 | 1.211 |
| Proteobacteria | Gammaproteobacteria_1 | Enterobacteriales | Enterobacteriaceae | Dickeya | 0.000 | 0.006 | 0.022 | 0.006 | 0.000 | 0.000 |
| Proteobacteria | Gammaproteobacteria_1 | Enterobacteriales | Enterobacteriaceae | Edwardsiella | 0.000 | 0.000 | 0.010 | 0.000 | 0.005 | 0.006 |
| Proteobacteria | Gammaproteobacteria_1 | Enterobacteriales | Enterobacteriaceae | Enterobacter | 0.260 | 5.003 | 0.977 | 2.079 | 2.014 | 1.931 |
| Proteobacteria | Gammaproteobacteria_1 | Enterobacteriales | Enterobacteriaceae | Enterobacter_1 | 1.277 | 10.702 | 0.191 | 7.788 | 9.964 | 7.886 |
| Proteobacteria | Gammaproteobacteria_1 | Enterobacteriales | Enterobacteriaceae | Enterobacter_2 | 0.008 | 0.117 | 0.004 | 0.012 | 0.010 | 0.054 |
| Proteobacteria | Gammaproteobacteria_1 | Enterobacteriales | Enterobacteriaceae | Enterobacter_3 | 0.000 | 0.006 | 0.010 | 0.018 | 0.010 | 0.012 |
| Proteobacteria | Gammaproteobacteria_1 | Enterobacteriales | Enterobacteriaceae | Enterobacter_4 | 0.431 | 1.627 | 0.048 | 2.270 | 1.246 | 1.241 |
| Proteobacteria | Gammaproteobacteria_1 | Enterobacteriales | Enterobacteriaceae | Escherichia | 0.016 | 0.099 | 0.077 | 0.186 | 0.115 | 0.073 |
| Proteobacteria | Gammaproteobacteria_1 | Enterobacteriales | Enterobacteriaceae | Escherichia_blattae_cluster | 0.008 | 0.043 | 0.016 | 0.012 | 0.035 | 0.030 |
| Proteobacteria | Gammaproteobacteria_1 | Enterobacteriales | Enterobacteriaceae | Escherichia-Shigella | 0.106 | 0.092 | 0.262 | 0.084 | 0.075 | 0.103 |
| Proteobacteria | Gammaproteobacteria_1 | Enterobacteriales | Enterobacteriaceae | Gut_cluster_1 | 0.065 | 0.018 | 0.615 | 0.018 | 0.005 | 0.000 |
| Proteobacteria | Gammaproteobacteria_1 | Enterobacteriales | Enterobacteriaceae | Hafnia | 0.000 | 0.000 | 0.020 | 0.000 | 0.005 | 0.006 |
| Proteobacteria | Gammaproteobacteria_1 | Enterobacteriales | Enterobacteriaceae | Klebsiella_1 | 0.293 | 2.132 | 0.145 | 2.804 | 2.188 | 2.463 |
| Proteobacteria | Gammaproteobacteria_1 | Enterobacteriales | Enterobacteriaceae | Klebsiella_2 | 0.163 | 0.357 | 0.046 | 0.449 | 0.538 | 0.424 |
| Proteobacteria | Gammaproteobacteria_1 | Enterobacteriales | Enterobacteriaceae | Kluyvera | 0.155 | 0.573 | 0.226 | 0.383 | 0.424 | 0.333 |
| Proteobacteria | Gammaproteobacteria_1 | Enterobacteriales | Enterobacteriaceae | Leminorella | 0.000 | 0.031 | 0.014 | 0.048 | 0.040 | 0.061 |
| Proteobacteria | Gammaproteobacteria_1 | Enterobacteriales | Enterobacteriaceae | Papillibacter | 0.024 | 0.031 | 0.040 | 0.006 | 0.000 | 0.000 |
| Proteobacteria | Gammaproteobacteria_1 | Enterobacteriales | Enterobacteriaceae | Pibocella | 0.000 | 0.000 | 0.010 | 0.000 | 0.000 | 0.000 |
| Proteobacteria | Gammaproteobacteria_1 | Enterobacteriales | Enterobacteriaceae | Pragia | 0.000 | 0.000 | 0.004 | 0.000 | 0.000 | 0.000 |
| Proteobacteria | Gammaproteobacteria_1 | Enterobacteriales | Enterobacteriaceae | Prevotella_1 | 0.049 | 0.006 | 1.427 | 0.006 | 0.000 | 0.000 |
| Proteobacteria | Gammaproteobacteria_1 | Enterobacteriales | Enterobacteriaceae | Ramlibacter | 0.065 | 0.000 | 0.024 | 0.006 | 0.000 | 0.006 |
| Proteobacteria | Gammaproteobacteria_1 | Enterobacteriales | Enterobacteriaceae | Sandaracinus | 0.016 | 0.000 | 0.000 | 0.006 | 0.000 | 0.000 |
| Proteobacteria | Gammaproteobacteria_1 | Enterobacteriales | Enterobacteriaceae | Shewanella | 0.000 | 0.043 | 0.216 | 0.060 | 0.045 | 0.030 |
| Proteobacteria | Gammaproteobacteria_1 | Enterobacteriales | Enterobacteriaceae | Sodalis_1 | 0.008 | 0.006 | 0.022 | 0.018 | 0.005 | 0.000 |
| Proteobacteria | Gammaproteobacteria_1 | Enterobacteriales | Enterobacteriaceae | Thorsellia | 0.000 | 0.000 | 0.004 | 0.000 | 0.000 | 0.000 |
| Proteobacteria | Gammaproteobacteria_1 | Enterobacteriales | Enterobacteriaceae | Treponema_caldaria_cluster | 0.024 | 0.000 | 0.016 | 0.000 | 0.000 | 0.000 |
| Proteobacteria | Gammaproteobacteria_2 | Enterobacteriales | Enterobacteriaceae | Knoellia | 0.000 | 0.000 | 0.008 | 0.000 | 0.000 | 0.000 |
| Proteobacteria | Gammaproteobacteria_1 | Enterobacteriales | Enterobacteriaceae_1 | Photorhabdus | 0.000 | 0.000 | 0.034 | 0.012 | 0.000 | 0.000 |
| Proteobacteria | Gammaproteobacteria_1 | Enterobacteriales | Enterobacteriaceae_1 | Proteus | 0.008 | 0.000 | 0.048 | 0.006 | 0.010 | 0.000 |
| Proteobacteria | Gammaproteobacteria_1 | Enterobacteriales | Enterobacteriaceae_1 | Providencia | 0.000 | 0.006 | 0.034 | 0.000 | 0.000 | 0.000 |
| Proteobacteria | Gammaproteobacteria_1 | Enterobacteriales | Enterobacteriaceae_1 | Pseudaminobacter | 0.000 | 0.006 | 0.000 | 0.000 | 0.000 | 0.000 |
| Proteobacteria | Gammaproteobacteria_1 | Enterobacteriales | Enterobacteriaceae_1 | Xenorhabdus | 0.000 | 0.006 | 0.002 | 0.036 | 0.010 | 0.006 |
| Proteobacteria | Gammaproteobacteria_1 | Enterobacteriales | Enterobacteriaceae_1 | Xenorhabdus_1 | 0.000 | 0.000 | 0.006 | 0.000 | 0.000 | 0.000 |
| Proteobacteria | Gammaproteobacteria_1 | Enterobacteriales | Enterobacteriaceae_1 | Xenorhabdus_2 | 0.000 | 0.000 | 0.008 | 0.000 | 0.000 | 0.000 |
| Proteobacteria | Gammaproteobacteria_1 | Enterobacteriales | Enterobacteriaceae_1 | Xenorhabdus_3 | 0.000 | 0.000 | 0.008 | 0.000 | 0.000 | 0.000 |

**Table S9 continued.**

| | | | | | | | | | | |
|---|---|---|---|---|---|---|---|---|---|---|
| Proteobacteria | Gammaproteobacteria_1 | Enterobacteriales | Enterobacteriaceae_1 | Xenorhabdus_4 | 0.000 | 0.000 | 0.006 | 0.000 | 0.000 | 0.000 |
| Proteobacteria | Gammaproteobacteria_1 | Enterobacteriales | Enterobacteriaceae_1 | Xenorhabdus_5 | 0.000 | 0.000 | 0.030 | 0.000 | 0.000 | 0.000 |
| Proteobacteria | Gammaproteobacteria_1 | Enterobacteriales | Enterobacteriaceae_1 | Xenorhabdus_6 | 0.000 | 0.000 | 0.004 | 0.000 | 0.000 | 0.000 |
| Proteobacteria | Gammaproteobacteria_1 | Enterobacteriales | Enterobacteriaceae_1 | Yersinia | 0.057 | 0.105 | 0.048 | 0.084 | 0.065 | 0.085 |
| Proteobacteria | Gammaproteobacteria_1 | Enterobacteriales | Enterobacteriaceae_2 | Candidatus_Hamiltonella | 0.000 | 0.000 | 0.004 | 0.000 | 0.000 | 0.000 |
| Proteobacteria | Gammaproteobacteria_1 | Enterobacteriales | Enterobacteriaceae_2 | Candidatus_Regiella | 0.000 | 0.000 | 0.006 | 0.000 | 0.000 | 0.000 |
| Proteobacteria | Gammaproteobacteria_1 | Enterobacteriales | Enterobacteriaceae_2 | endosymbionts | 0.000 | 0.000 | 0.020 | 0.000 | 0.000 | 0.000 |
| Proteobacteria | Gammaproteobacteria_1 | Enterobacteriales | Enterobacteriaceae_3 | Buchnera_2 | 0.000 | 0.006 | 0.012 | 0.000 | 0.000 | 0.000 |
| Proteobacteria | Gammaproteobacteria_1 | Enterobacteriales | Enterobacteriaceae_3 | Candidatus_Baumannia | 0.000 | 0.000 | 0.036 | 0.000 | 0.000 | 0.000 |
| Proteobacteria | Gammaproteobacteria_1 | Enterobacteriales | Enterobacteriaceae_3 | Candidatus_Blochmannia | 0.000 | 0.000 | 0.018 | 0.000 | 0.000 | 0.000 |
| Proteobacteria | Gammaproteobacteria_1 | Enterobacteriales | Enterobacteriaceae_3 | Candidatus_Carsonella | 0.000 | 0.000 | 0.042 | 0.000 | 0.000 | 0.000 |
| Proteobacteria | Gammaproteobacteria_1 | Enterobacteriales | Enterobacteriaceae_3 | Candidatus_Curculioniphilus | 0.000 | 0.000 | 0.004 | 0.000 | 0.000 | 0.000 |
| Proteobacteria | Gammaproteobacteria_1 | Enterobacteriales | Enterobacteriaceae_3 | Candidatus_Ishikawaella | 0.000 | 0.000 | 0.016 | 0.006 | 0.000 | 0.006 |
| Proteobacteria | Gammaproteobacteria_1 | Enterobacteriales | Enterobacteriaceae_3 | Candidatus_Riesia | 0.000 | 0.000 | 0.006 | 0.000 | 0.000 | 0.000 |
| Proteobacteria | Gammaproteobacteria_1 | Enterobacteriales | Enterobacteriaceae_3 | endosymbionts_1 | 0.000 | 0.000 | 0.010 | 0.000 | 0.000 | 0.000 |
| Proteobacteria | Gammaproteobacteria_1 | Enterobacteriales | Enterobacteriaceae_3 | endosymbionts_3 | 0.000 | 0.000 | 0.006 | 0.000 | 0.000 | 0.000 |
| Proteobacteria | Gammaproteobacteria | Enterobacterales | Erwiniaceae | Izhakiella | 0.000 | 0.006 | 0.000 | 0.006 | 0.005 | 0.000 |
| Proteobacteria | Gammaproteobacteria | Enterobacteriales | Erwiniaceae | Erwinia | 0.033 | 0.407 | 0.212 | 0.401 | 0.364 | 0.345 |
| Proteobacteria | Alphaproteobacteria | Sphingomonadales | Erythrobacteraceae | Altererythrobacter | 0.033 | 0.006 | 0.032 | 0.000 | 0.000 | 0.000 |
| Proteobacteria | Alphaproteobacteria | Sphingomonadales | Erythrobacteraceae | Erythrobacter | 0.073 | 0.006 | 0.056 | 0.006 | 0.000 | 0.000 |
| Proteobacteria | Alphaproteobacteria | Sphingomonadales | Erythrobacteraceae | Porphyrobacter_3 | 0.000 | 0.000 | 0.004 | 0.000 | 0.000 | 0.000 |
| Proteobacteria | Alphaproteobacteria | Sphingomonadales | Erythrobacteraceae | Porphyromonas | 0.000 | 0.000 | 0.073 | 0.000 | 0.000 | 0.000 |
| Proteobacteria | Gammaproteobacteria_1 | Alteromonadales | Ferrimonadaceae | Ferrimonas | 0.000 | 0.000 | 0.026 | 0.000 | 0.000 | 0.000 |
| Proteobacteria | Gammaproteobacteria_2 | Thiotrichales | Francisellaceae | Francisella | 0.000 | 0.000 | 0.008 | 0.000 | 0.000 | 0.000 |
| Proteobacteria | Betaproteobacteria | Nitrosomonadales | Gallionellaceae | Candidatus_Nitrotoga | 0.000 | 0.000 | 0.008 | 0.000 | 0.000 | 0.000 |
| Proteobacteria | Betaproteobacteria | Nitrosomonadales | Gallionellaceae | Gallionella | 0.008 | 0.000 | 0.006 | 0.000 | 0.000 | 0.000 |
| Proteobacteria | Alphaproteobacteria | Rhodospirillales | Geminicoccaceae | Geminicoccus | 0.008 | 0.000 | 0.000 | 0.000 | 0.000 | 0.000 |
| Proteobacteria | Deltaproteobacteria | Desulfuromonadales | Geobacteraceae | Geoalkalibacter | 0.000 | 0.000 | 0.004 | 0.000 | 0.000 | 0.000 |
| Proteobacteria | Deltaproteobacteria | Desulfuromonadales | Geobacteraceae | Geobacter | 0.016 | 0.000 | 0.000 | 0.000 | 0.000 | 0.000 |
| Proteobacteria | Deltaproteobacteria | Desulfuromonadales | Geobacteraceae | Geobacter_1 | 0.000 | 0.000 | 0.020 | 0.000 | 0.000 | 0.000 |
| Proteobacteria | Deltaproteobacteria | Desulfuromonadales | Geobacteraceae | Geobacter_2 | 0.000 | 0.000 | 0.006 | 0.000 | 0.000 | 0.000 |
| Proteobacteria | Deltaproteobacteria | Desulfuromonadales | Geobacteraceae | Geobacter_5 | 0.000 | 0.000 | 0.008 | 0.000 | 0.000 | 0.000 |
| Proteobacteria | Deltaproteobacteria | Desulfuromonadales | Geobacteraceae | Geobacter_6 | 0.008 | 0.000 | 0.008 | 0.000 | 0.000 | 0.000 |
| Proteobacteria | Deltaproteobacteria | Desulfuromonadales | Geobacteraceae | Geopsychrobacter | 0.000 | 0.000 | 0.004 | 0.000 | 0.000 | 0.000 |
| Proteobacteria | Deltaproteobacteria | Desulfuromonadales | Geobacteraceae | Geothermobacter | 0.000 | 0.000 | 0.004 | 0.000 | 0.000 | 0.000 |
| Proteobacteria | Gammaproteobacteria_5 | Chromatiales | Granulosicoccaceae | Granulosicoccus | 0.000 | 0.000 | 0.008 | 0.000 | 0.000 | 0.000 |
| Proteobacteria | Gammaproteobacteria | Enterobacteriales | Hafniaceae | Enterobacillus | 0.000 | 0.006 | 0.000 | 0.006 | 0.000 | 0.000 |
| Proteobacteria | Gammaproteobacteria_1 | Oceanospirillales | Hahellaceae | Hahella | 0.000 | 0.000 | 0.010 | 0.000 | 0.000 | 0.000 |
| Proteobacteria | Gammaproteobacteria_1 | Oceanospirillales_4 | Hahellaceae | Endozoicomonas | 0.000 | 0.000 | 0.034 | 0.006 | 0.000 | 0.000 |
| Proteobacteria | Gammaproteobacteria_1 | Oceanospirillales_4 | Hahellaceae | Zooshikella | 0.000 | 0.000 | 0.004 | 0.000 | 0.000 | 0.000 |

**Table S9 continued.**

| | | | | | | | | | | |
|---|---|---|---|---|---|---|---|---|---|---|
| Proteobacteria | Deltaproteobacteria | Myxococcales | Haliangiaceae | Haliangium | 0.212 | 0.006 | 0.046 | 0.006 | 0.000 | 0.000 |
| Proteobacteria | Deltaproteobacteria | Myxococcales | Haliangiaceae | Haloferula | 0.000 | 0.000 | 0.060 | 0.000 | 0.000 | 0.000 |
| Proteobacteria | Gammaproteobacteria | Oceanospirillales | Halomonadaceae | Carnimonas | 0.000 | 0.000 | 0.002 | 0.000 | 0.000 | 0.000 |
| Proteobacteria | Gammaproteobacteria | Oceanospirillales | Halomonadaceae | Halomonas | 0.016 | 0.000 | 0.002 | 0.006 | 0.000 | 0.000 |
| Proteobacteria | Gammaproteobacteria_1 | Oceanospirillales_1 | Halomonadaceae | Candidatus_Portiera | 0.000 | 0.000 | 0.010 | 0.000 | 0.000 | 0.000 |
| Proteobacteria | Gammaproteobacteria_1 | Oceanospirillales_1 | Halomonadaceae | Chromohalobacter | 0.000 | 0.000 | 0.016 | 0.000 | 0.000 | 0.000 |
| Proteobacteria | Gammaproteobacteria_1 | Oceanospirillales_1 | Halomonadaceae | Cobetia | 0.000 | 0.000 | 0.004 | 0.000 | 0.000 | 0.000 |
| Proteobacteria | Gammaproteobacteria_1 | Oceanospirillales_1 | Halomonadaceae | Halomonas_2 | 0.000 | 0.000 | 0.004 | 0.000 | 0.000 | 0.000 |
| Proteobacteria | Gammaproteobacteria_1 | Oceanospirillales_1 | Halomonadaceae | Halomonas_3 | 0.000 | 0.000 | 0.008 | 0.000 | 0.000 | 0.000 |
| Proteobacteria | Gammaproteobacteria_1 | Oceanospirillales_1 | Halomonadaceae | Halomonas_4 | 0.000 | 0.000 | 0.012 | 0.000 | 0.000 | 0.000 |
| Proteobacteria | Gammaproteobacteria_1 | Oceanospirillales_1 | Halomonadaceae | Halomonas_5 | 0.000 | 0.000 | 0.004 | 0.000 | 0.000 | 0.000 |
| Proteobacteria | Gammaproteobacteria_1 | Oceanospirillales_1 | Halomonadaceae | Halomonas_6 | 0.000 | 0.000 | 0.004 | 0.000 | 0.000 | 0.000 |
| Proteobacteria | Gammaproteobacteria_1 | Oceanospirillales_1 | Halomonadaceae | Halomonas_7 | 0.000 | 0.000 | 0.004 | 0.000 | 0.000 | 0.000 |
| Proteobacteria | Gammaproteobacteria_1 | Oceanospirillales_1 | Halomonadaceae | Kushneria | 0.000 | 0.000 | 0.012 | 0.000 | 0.000 | 0.000 |
| Proteobacteria | Gammaproteobacteria_1 | Oceanospirillales_4 | Halomonadaceae | Halovibrio_2 | 0.000 | 0.000 | 0.008 | 0.000 | 0.000 | 0.000 |
| Proteobacteria | Gammaproteobacteria | Chromatiales | Halothiobacillaceae | Halothiobacillus_1 | 0.000 | 0.000 | 0.004 | 0.000 | 0.000 | 0.000 |
| Proteobacteria | Gammaproteobacteria | Chromatiales | Halothiobacillaceae | Halothiobacillus_2 | 0.000 | 0.000 | 0.004 | 0.000 | 0.000 | 0.000 |
| Proteobacteria | Gammaproteobacteria | Chromatiales | Halothiobacillaceae | Thiofaba | 0.000 | 0.000 | 0.004 | 0.000 | 0.000 | 0.000 |
| Proteobacteria | Gammaproteobacteria | Chromatiales | Halothiobacillaceae | Thiovirga | 0.000 | 0.000 | 0.006 | 0.000 | 0.000 | 0.000 |
| Proteobacteria | Epsilonproteobacteria | Campylobacterales | Helicobacteraceae | Heliobacterium | 0.000 | 0.000 | 0.006 | 0.000 | 0.000 | 0.000 |
| Proteobacteria | Epsilonproteobacteria | Campylobacterales | Helicobacteraceae | Rubritepida | 0.000 | 0.000 | 0.006 | 0.000 | 0.000 | 0.000 |
| Proteobacteria | Epsilonproteobacteria | Campylobacterales | Helicobacteraceae | Sulfurospirillum | 2.164 | 0.493 | 0.042 | 0.252 | 0.055 | 0.024 |
| Proteobacteria | Alphaproteobacteria | Rickettsiales | Holosporaceae | Holospora | 0.000 | 0.000 | 0.006 | 0.000 | 0.000 | 0.000 |
| Proteobacteria | Betaproteobacteria | Hydrogenophilales | Hydrogenophilaceae | Petrobacter | 0.000 | 0.000 | 0.004 | 0.000 | 0.000 | 0.000 |
| Proteobacteria | Betaproteobacteria | Hydrogenophilales | Hydrogenophilaceae | Termite_cluster | 2.400 | 0.850 | 0.117 | 1.162 | 0.085 | 0.048 |
| Proteobacteria | Betaproteobacteria | Hydrogenophilales | Hydrogenophilaceae | Thiobacillus | 0.000 | 0.000 | 0.012 | 0.000 | 0.000 | 0.000 |
| Proteobacteria | Betaproteobacteria | Hydrogenophilales | Hydrogenophilaceae | Thiobacter | 0.008 | 0.000 | 0.000 | 0.000 | 0.000 | 0.000 |
| Proteobacteria | Betaproteobacteria | Hydrogenophilales | Hydrogenophilaceae | Thiorhodospira | 0.000 | 0.000 | 0.008 | 0.000 | 0.000 | 0.000 |
| Proteobacteria | Hydrogenophilalia | Hydrogenophilales | Hydrogenophilaceae | Hydrotalea | 0.024 | 0.000 | 0.000 | 0.006 | 0.000 | 0.000 |
| Proteobacteria | Alphaproteobacteria | Rhizobiales | Hyphomicrobiaceae | Blastochloris | 0.033 | 0.000 | 0.006 | 0.000 | 0.000 | 0.000 |
| Proteobacteria | Alphaproteobacteria | Rhizobiales | Hyphomicrobiaceae | Methylosinus | 0.000 | 0.000 | 0.010 | 0.000 | 0.000 | 0.000 |
| Proteobacteria | Alphaproteobacteria | Rhizobiales | Hyphomicrobiaceae | Methyloversatilis | 0.000 | 0.000 | 0.004 | 0.000 | 0.000 | 0.000 |
| Proteobacteria | Alphaproteobacteria | Rhizobiales_0 | Hyphomicrobiaceae | Devosia | 0.008 | 0.031 | 0.000 | 0.006 | 0.000 | 0.000 |
| Proteobacteria | Alphaproteobacteria | Rhizobiales_1 | Hyphomicrobiaceae | Devosia-Prosthecomicrobium | 0.041 | 0.012 | 0.069 | 0.000 | 0.005 | 0.000 |
| Proteobacteria | Alphaproteobacteria | Rhizobiales_1 | Hyphomicrobiaceae | Maritalea | 0.000 | 0.000 | 0.006 | 0.000 | 0.000 | 0.000 |
| Proteobacteria | Alphaproteobacteria | Rhizobiales_2 | Hyphomicrobiaceae | Proteiniphilum | 0.000 | 0.000 | 0.046 | 0.000 | 0.000 | 0.000 |
| Proteobacteria | Alphaproteobacteria | Rhizobiales_2 | Hyphomicrobiaceae | Rhodopirellula | 0.000 | 0.000 | 0.187 | 0.000 | 0.000 | 0.000 |
| Proteobacteria | Alphaproteobacteria | Rhizobiales_2 | Hyphomicrobiaceae | Rhodospirillum | 0.000 | 0.000 | 0.006 | 0.000 | 0.000 | 0.000 |
| Proteobacteria | Alphaproteobacteria | Rhizobiales_3 | Hyphomicrobiaceae | Filomicrobium_1 | 0.000 | 0.000 | 0.006 | 0.000 | 0.000 | 0.000 |
| Proteobacteria | Alphaproteobacteria | Rhizobiales_3 | Hyphomicrobiaceae | Hyphomicrobium_1 | 0.008 | 0.000 | 0.022 | 0.000 | 0.000 | 0.000 |

**Table S9 continued.**

| | | | | | | | | | | |
|---|---|---|---|---|---|---|---|---|---|---|
| Proteobacteria | Alphaproteobacteria | Rhizobiales_3 | Hyphomicrobiaceae | Hyphomicrobium_2 | 0.008 | 0.000 | 0.024 | 0.000 | 0.005 | 0.000 |
| Proteobacteria | Alphaproteobacteria | Rhizobiales_3 | Hyphomicrobiaceae | Hyphomonas | 0.000 | 0.006 | 0.022 | 0.000 | 0.000 | 0.000 |
| Proteobacteria | Alphaproteobacteria | Rhizobiales_3 | Hyphomicrobiaceae | Pelagibius | 0.000 | 0.006 | 0.012 | 0.000 | 0.000 | 0.000 |
| Proteobacteria | Alphaproteobacteria | Rhizobiales_2 | Hyphomicrobiaceae_1 | Ancalomicrobium | 0.000 | 0.000 | 0.012 | 0.000 | 0.000 | 0.000 |
| Proteobacteria | Alphaproteobacteria | Rhizobiales_2 | Hyphomicrobiaceae_1 | Azorhizobium_1 | 0.000 | 0.000 | 0.004 | 0.000 | 0.000 | 0.000 |
| Proteobacteria | Alphaproteobacteria | Rhizobiales_2 | Hyphomicrobiaceae_1 | Azorhizobium_2 | 0.000 | 0.000 | 0.004 | 0.000 | 0.000 | 0.000 |
| Proteobacteria | Alphaproteobacteria | Rhizobiales_2 | Hyphomicrobiaceae_1 | Xanthobacter_1 | 0.000 | 0.000 | 0.006 | 0.000 | 0.000 | 0.000 |
| Proteobacteria | Alphaproteobacteria | Rhizobiales_2 | Hyphomicrobiaceae_1 | Xanthobacter_2 | 0.000 | 0.000 | 0.008 | 0.000 | 0.000 | 0.000 |
| Proteobacteria | Alphaproteobacteria | Caulobacterales | Hyphomonadaceae | Hellea | 0.000 | 0.000 | 0.004 | 0.000 | 0.000 | 0.000 |
| Proteobacteria | Alphaproteobacteria | Caulobacterales | Hyphomonadaceae | Maricaulis | 0.000 | 0.000 | 0.008 | 0.000 | 0.000 | 0.000 |
| Proteobacteria | Alphaproteobacteria | Caulobacterales | Hyphomonadaceae | Oceanicaulis | 0.000 | 0.000 | 0.006 | 0.000 | 0.000 | 0.000 |
| Proteobacteria | Alphaproteobacteria | Caulobacterales | Hyphomonadaceae | Robiginitomaculum | 0.000 | 0.000 | 0.004 | 0.000 | 0.000 | 0.000 |
| Proteobacteria | Gammaproteobacteria_1 | Alteromonadales_1 | Idiomarinaceae | Idiomarina | 0.000 | 0.000 | 0.016 | 0.000 | 0.000 | 0.000 |
| Proteobacteria | Gammaproteobacteria_1 | Alteromonadales_1 | Idiomarinaceae | Pseudidiomarina | 0.000 | 0.000 | 0.022 | 0.000 | 0.000 | 0.000 |
| Proteobacteria | Betaproteobacteria | Burkholderiales | incertae_sedis | Thiomonas | 0.000 | 0.000 | 0.018 | 0.000 | 0.000 | 0.000 |
| Proteobacteria | Alphaproteobacteria | Kiloniellales | Kiloniellaceae | Kineococcus | 0.000 | 0.018 | 0.020 | 0.000 | 0.000 | 0.000 |
| Proteobacteria | Alphaproteobacteria | Kordiimonadales | Kordiimonadaceae | Kosakonia | 0.277 | 0.986 | 1.935 | 2.816 | 2.108 | 1.658 |
| Proteobacteria | Gammaproteobacteria_2 | Legionellales | Legionellaceae | Legionella_1 | 0.000 | 0.000 | 0.073 | 0.000 | 0.000 | 0.000 |
| Proteobacteria | Gammaproteobacteria_2 | Legionellales | Legionellaceae | Legionella_4 | 0.000 | 0.000 | 0.006 | 0.000 | 0.000 | 0.000 |
| Proteobacteria | Gammaproteobacteria_2 | Legionellales | Legionellaceae | Legionella_6 | 0.000 | 0.000 | 0.006 | 0.000 | 0.000 | 0.000 |
| Proteobacteria | Gammaproteobacteria_3 | Legionellales | Legionellaceae | Legionella_2 | 0.000 | 0.000 | 0.010 | 0.000 | 0.000 | 0.000 |
| Proteobacteria | Gammaproteobacteria_3 | Legionellales | Legionellaceae | Legionella_7 | 0.000 | 0.000 | 0.018 | 0.000 | 0.000 | 0.000 |
| Proteobacteria | Gammaproteobacteria_3 | Legionellales | Legionellaceae | Legionella_9 | 0.000 | 0.000 | 0.004 | 0.000 | 0.000 | 0.000 |
| Proteobacteria | Gammaproteobacteria_4 | Legionellales | Legionellaceae | Legionella_5 | 0.000 | 0.000 | 0.010 | 0.000 | 0.000 | 0.000 |
| Proteobacteria | Gammaproteobacteria_5 | Legionellales | Legionellaceae | Leifsonia | 0.008 | 0.000 | 0.000 | 0.000 | 0.000 | 0.006 |
| Proteobacteria | Gammaproteobacteria_1 | Oceanospirillales_3 | Litoricolaceae | Litoricola | 0.000 | 0.000 | 0.006 | 0.000 | 0.000 | 0.000 |
| Proteobacteria | Alphaproteobacteria | Rhizobiales | Methylobacteriaceae | Megasphaera | 0.000 | 0.000 | 0.028 | 0.000 | 0.000 | 0.000 |
| Proteobacteria | Gammaproteobacteria | Methylococcales | Methylococcaceae | Methylocaldum | 0.000 | 0.000 | 0.010 | 0.000 | 0.000 | 0.006 |
| Proteobacteria | Gammaproteobacteria_4 | Methylococcales | Methylococcaceae | Methylomicrobium_1 | 0.000 | 0.000 | 0.010 | 0.000 | 0.000 | 0.000 |
| Proteobacteria | Gammaproteobacteria_4 | Methylococcales | Methylococcaceae | Methylomicrobium_2 | 0.000 | 0.000 | 0.004 | 0.000 | 0.000 | 0.000 |
| Proteobacteria | Gammaproteobacteria_4 | Methylococcales | Methylococcaceae | Methylomonas | 0.000 | 0.000 | 0.014 | 0.000 | 0.000 | 0.000 |
| Proteobacteria | Gammaproteobacteria_2 | Methylococcales | Methylococcaceae_1 | Methylococcus | 0.000 | 0.000 | 0.006 | 0.000 | 0.000 | 0.000 |
| Proteobacteria | Gammaproteobacteria_4 | Methylococcales | Methylococcaceae_1 | Methylosarcina | 0.000 | 0.000 | 0.004 | 0.000 | 0.000 | 0.000 |
| Proteobacteria | Gammaproteobacteria_2 | Methylococcales | Methylococcaceae_2 | Methylothermus | 0.000 | 0.000 | 0.004 | 0.000 | 0.000 | 0.000 |
| Proteobacteria | Gammaproteobacteria_4 | Methylococcales | Methylococcaceae_3 | Methylosoma | 0.000 | 0.000 | 0.004 | 0.000 | 0.000 | 0.000 |
| Proteobacteria | Alphaproteobacteria | Rhizobiales | Methylocystaceae | Harryflintia | 0.000 | 0.006 | 0.000 | 0.000 | 0.000 | 0.000 |
| Proteobacteria | Alphaproteobacteria | Rhizobiales | Methylocystaceae | Plesiomonas | 0.000 | 0.000 | 0.087 | 0.000 | 0.010 | 0.006 |
| Proteobacteria | Alphaproteobacteria | Rhizobiales_2 | Methylocystaceae_1 | Methylocystis_3 | 0.008 | 0.000 | 0.008 | 0.000 | 0.000 | 0.000 |
| Proteobacteria | Alphaproteobacteria | Rhizobiales_2 | Methylocystaceae_1 | Methylonatrum | 0.000 | 0.000 | 0.004 | 0.000 | 0.000 | 0.000 |
| Proteobacteria | Alphaproteobacteria | Rhizobiales_2 | Methylocystaceae_1 | Methylotenera | 0.008 | 0.000 | 0.000 | 0.000 | 0.000 | 0.000 |

**Table S9 continued.**

| | | | | | | | | | | |
|---|---|---|---|---|---|---|---|---|---|---|
| Proteobacteria | Gammaproteobacteria_2 | Methylonatrum | Methylonatrum | Methylophilus | 0.073 | 0.031 | 0.018 | 0.006 | 0.000 | 0.000 |
| Proteobacteria | Betaproteobacteria | Methylophilales | Methylophilaceae | LD28_freshwater_group | 0.000 | 0.000 | 0.004 | 0.000 | 0.000 | 0.000 |
| Proteobacteria | Betaproteobacteria | Methylophilales | Methylophilaceae | Methylobacillus | 0.008 | 0.000 | 0.008 | 0.000 | 0.000 | 0.000 |
| Proteobacteria | Betaproteobacteria | Methylophilales | Methylophilaceae | Methylopila | 0.000 | 0.006 | 0.006 | 0.000 | 0.000 | 0.000 |
| Proteobacteria | Betaproteobacteria | Methylophilales | Methylophilaceae | OM43_clade | 0.016 | 0.000 | 0.018 | 0.000 | 0.000 | 0.000 |
| Proteobacteria | Gammaproteobacteria | Pseudomonadales | Moraxellaceae | Cavicella | 0.016 | 0.000 | 0.000 | 0.000 | 0.000 | 0.000 |
| Proteobacteria | Gammaproteobacteria | Pseudomonadales | Moraxellaceae | Paraperlucidibaca | 0.008 | 0.000 | 0.000 | 0.000 | 0.000 | 0.000 |
| Proteobacteria | Gammaproteobacteria__1 | Pseudomonadales | Moraxellaceae | Acinetobacter | 0.944 | 1.183 | 3.529 | 0.982 | 0.055 | 0.127 |
| Proteobacteria | Gammaproteobacteria_1 | Pseudomonadales | Moraxellaceae | Alkanindiges | 0.000 | 0.000 | 0.022 | 0.012 | 0.000 | 0.000 |
| Proteobacteria | Gammaproteobacteria_1 | Pseudomonadales | Moraxellaceae | Moraxella_1 | 0.000 | 0.000 | 0.008 | 0.000 | 0.000 | 0.000 |
| Proteobacteria | Gammaproteobacteria_1 | Pseudomonadales | Moraxellaceae | Moraxella_2 | 0.000 | 0.000 | 0.040 | 0.000 | 0.000 | 0.000 |
| Proteobacteria | Gammaproteobacteria_1 | Pseudomonadales | Moraxellaceae | Petrimonas | 0.000 | 0.000 | 0.008 | 0.000 | 0.000 | 0.000 |
| Proteobacteria | Gammaproteobacteria | Enterobacterales | Morganellaceae | Arsenophonus | 0.000 | 0.000 | 0.018 | 0.000 | 0.000 | 0.000 |
| Proteobacteria | Gammaproteobacteria | Enterobacterales | Morganellaceae | Mucilaginibacter | 0.000 | 0.018 | 0.000 | 0.000 | 0.000 | 0.000 |
| Proteobacteria | Gammaproteobacteria_1 | Alteromonadales | Moritellaceae | Paramoritella | 0.000 | 0.000 | 0.006 | 0.000 | 0.000 | 0.000 |
| Proteobacteria | Gammaproteobacteria_1 | Alteromonadales_2 | Moritellaceae | Moritella | 0.000 | 0.006 | 0.016 | 0.000 | 0.005 | 0.006 |
| Proteobacteria | Deltaproteobacteria | Myxococcales | Myxococcaceae | Agreia | 0.000 | 0.000 | 0.004 | 0.000 | 0.000 | 0.000 |
| Proteobacteria | Deltaproteobacteria | Myxococcales | Myxococcaceae | Corallococcus | 0.008 | 0.000 | 0.000 | 0.000 | 0.000 | 0.000 |
| Proteobacteria | Deltaproteobacteria | Myxococcales | Myxococcaceae | Nakamurella | 0.000 | 0.000 | 0.004 | 0.000 | 0.000 | 0.000 |
| Proteobacteria | Deltaproteobacteria | Myxococcales | Nannocystaceae | Pseudoalteromonas | 0.008 | 0.000 | 0.113 | 0.000 | 0.005 | 0.006 |
| Proteobacteria | Epsilonproteobacteria | Nautiliales | Nautiliaceae | Caminibacter_2 | 0.000 | 0.000 | 0.004 | 0.000 | 0.000 | 0.000 |
| Proteobacteria | Epsilonproteobacteria | Nautiliales | Nautiliaceae | Nautilia | 0.000 | 0.000 | 0.004 | 0.000 | 0.000 | 0.000 |
| Proteobacteria | Epsilonproteobacteria | Nautiliales | Nautiliaceae | Thioreductor | 0.000 | 0.000 | 0.004 | 0.000 | 0.000 | 0.000 |
| Proteobacteria | Betaproteobacteria | Neisseriales | Neisseriaceae | Leeia | 0.000 | 0.000 | 0.012 | 0.000 | 0.000 | 0.000 |
| Proteobacteria | Betaproteobacteria | Neisseriales | Neisseriaceae | Silicimonas | 0.000 | 0.000 | 0.000 | 0.000 | 0.000 | 0.006 |
| Proteobacteria | Betaproteobacteria | Neisseriales | Neisseriaceae | Simplicispira_2 | 0.000 | 0.000 | 0.054 | 0.000 | 0.000 | 0.000 |
| Proteobacteria | Betaproteobacteria | Neisseriales | Neisseriaceae | Uruburuella | 0.008 | 0.000 | 0.000 | 0.000 | 0.000 | 0.000 |
| Proteobacteria | Betaproteobacteria | Neisseriales | Neisseriaceae_1 | Alysiella | 0.000 | 0.000 | 0.008 | 0.000 | 0.000 | 0.000 |
| Proteobacteria | Betaproteobacteria | Neisseriales | Neisseriaceae_1 | Aquaspirillum | 0.000 | 0.000 | 0.004 | 0.000 | 0.005 | 0.000 |
| Proteobacteria | Betaproteobacteria | Neisseriales | Neisseriaceae_1 | Conchiformibius | 0.000 | 0.000 | 0.014 | 0.000 | 0.000 | 0.000 |
| Proteobacteria | Betaproteobacteria | Neisseriales | Neisseriaceae_1 | Laribacter | 0.000 | 0.000 | 0.004 | 0.000 | 0.005 | 0.000 |
| Proteobacteria | Betaproteobacteria | Neisseriales | Neisseriaceae_1 | Microvirgula | 0.000 | 0.000 | 0.006 | 0.000 | 0.000 | 0.000 |
| Proteobacteria | Betaproteobacteria | Neisseriales | Neisseriaceae_1 | Neochlamydia | 0.008 | 0.000 | 0.014 | 0.000 | 0.000 | 0.000 |
| Proteobacteria | Betaproteobacteria | Neisseriales | Neisseriaceae_1 | Paludimonas | 0.000 | 0.000 | 0.006 | 0.000 | 0.005 | 0.000 |
| Proteobacteria | Betaproteobacteria | Neisseriales | Neisseriaceae_1 | Stenoxybacter | 0.000 | 0.000 | 0.004 | 0.000 | 0.000 | 0.000 |
| Proteobacteria | Betaproteobacteria | Neisseriales | Neisseriaceae_1 | Uncultured_42 | 0.000 | 0.000 | 0.010 | 0.000 | 0.000 | 0.000 |
| Proteobacteria | Betaproteobacteria | Neisseriales | Neisseriaceae_1 | Vitreoscilla | 0.000 | 0.000 | 0.004 | 0.000 | 0.000 | 0.000 |
| Proteobacteria | Betaproteobacteria | Neisseriales | Neisseriaceae_1 | Vulgatibacter | 0.008 | 0.000 | 0.000 | 0.000 | 0.000 | 0.000 |
| Proteobacteria | Betaproteobacteria | Neisseriales | Neisseriaceae_2 | Andreprevotia | 0.000 | 0.000 | 0.004 | 0.000 | 0.000 | 0.000 |
| Proteobacteria | Betaproteobacteria | Neisseriales | Neisseriaceae_2 | Deefgea | 0.000 | 0.000 | 0.010 | 0.000 | 0.000 | 0.000 |

**Table S9 continued.**

| | | | | | | | | | | |
|---|---|---|---|---|---|---|---|---|---|---|
| Proteobacteria | Betaproteobacteria | Nitrosomonadales | Nitrosomonadaceae | Nitrospina | 0.000 | 0.000 | 0.175 | 0.000 | 0.000 | 0.000 |
| Proteobacteria | Deltaproteobacteria | Myxococcales | nocystaceae | nocystis | 0.000 | 0.000 | 0.006 | 0.000 | 0.000 | 0.000 |
| Proteobacteria | Gammaproteobacteria | Oceanospirillales | Oceanospirillaceae | Marinomonas | 0.000 | 0.000 | 0.050 | 0.000 | 0.000 | 0.000 |
| Proteobacteria | Gammaproteobacteria | Oceanospirillales | Oceanospirillaceae | Nitrobacter | 0.016 | 0.006 | 0.018 | 0.000 | 0.010 | 0.000 |
| Proteobacteria | Gammaproteobacteria_1 | Oceanospirillales | Oceanospirillaceae | Pseudovibrio | 0.000 | 0.000 | 0.012 | 0.000 | 0.000 | 0.000 |
| Proteobacteria | Gammaproteobacteria_1 | Oceanospirillales | Oceanospirillaceae_1 | Marinobacterium_1 | 0.000 | 0.000 | 0.016 | 0.000 | 0.000 | 0.000 |
| Proteobacteria | Gammaproteobacteria_1 | Oceanospirillales | Oceanospirillaceae_1 | Marinobacterium_2 | 0.000 | 0.000 | 0.026 | 0.000 | 0.000 | 0.000 |
| Proteobacteria | Gammaproteobacteria_1 | Oceanospirillales | Oceanospirillaceae_1 | Marinobacterium_3 | 0.000 | 0.000 | 0.004 | 0.000 | 0.000 | 0.000 |
| Proteobacteria | Gammaproteobacteria_1 | Oceanospirillales | Oceanospirillaceae_1 | Marinobacterium_4 | 0.000 | 0.000 | 0.006 | 0.000 | 0.000 | 0.000 |
| Proteobacteria | Gammaproteobacteria_1 | Oceanospirillales | Oceanospirillaceae_1 | Marinospirillum | 0.000 | 0.000 | 0.012 | 0.000 | 0.000 | 0.000 |
| Proteobacteria | Gammaproteobacteria_1 | Oceanospirillales | Oceanospirillaceae_1 | Neptuniibacter | 0.000 | 0.000 | 0.008 | 0.006 | 0.000 | 0.000 |
| Proteobacteria | Gammaproteobacteria_1 | Oceanospirillales | Oceanospirillaceae_1 | Neptunomonas | 0.000 | 0.006 | 0.018 | 0.000 | 0.000 | 0.000 |
| Proteobacteria | Gammaproteobacteria_1 | Oceanospirillales | Oceanospirillaceae_2 | Oceaniserpentilla | 0.000 | 0.000 | 0.008 | 0.000 | 0.000 | 0.000 |
| Proteobacteria | Gammaproteobacteria_1 | Oceanospirillales | Oceanospirillaceae_2 | Oceanobacter | 0.000 | 0.000 | 0.008 | 0.000 | 0.000 | 0.000 |
| Proteobacteria | Gammaproteobacteria_1 | Oceanospirillales | Oceanospirillaceae_2 | Oceanospirillum | 0.000 | 0.006 | 0.012 | 0.000 | 0.000 | 0.000 |
| Proteobacteria | Gammaproteobacteria_1 | Oceanospirillales | Oceanospirillaceae_2 | Oleispira | 0.000 | 0.000 | 0.008 | 0.000 | 0.000 | 0.000 |
| Proteobacteria | Gammaproteobacteria_1 | Oceanospirillales_3 | Oleiphilaceae | Oleiphilus | 0.000 | 0.000 | 0.008 | 0.000 | 0.000 | 0.000 |
| Proteobacteria | Betaproteobacteria | Burkholderiales | Oxalobacteraceae | Aquaspirillum_sp_a | 0.000 | 0.000 | 0.004 | 0.000 | 0.000 | 0.000 |
| Proteobacteria | Betaproteobacteria | Burkholderiales | Oxalobacteraceae | Collimonas | 0.008 | 0.000 | 0.012 | 0.000 | 0.005 | 0.000 |
| Proteobacteria | Betaproteobacteria | Burkholderiales | Oxalobacteraceae | Duganella | 0.000 | 0.000 | 0.004 | 0.000 | 0.005 | 0.000 |
| Proteobacteria | Betaproteobacteria | Burkholderiales | Oxalobacteraceae | Duganella_sp_a | 0.000 | 0.006 | 0.004 | 0.000 | 0.000 | 0.000 |
| Proteobacteria | Betaproteobacteria | Burkholderiales | Oxalobacteraceae | Herbaspirillum_1 | 0.000 | 0.000 | 0.016 | 0.000 | 0.000 | 0.000 |
| Proteobacteria | Betaproteobacteria | Burkholderiales | Oxalobacteraceae | Herbaspirillum_2 | 0.000 | 0.000 | 0.006 | 0.000 | 0.000 | 0.000 |
| Proteobacteria | Betaproteobacteria | Burkholderiales | Oxalobacteraceae | Herbaspirillum_3 | 0.000 | 0.000 | 0.018 | 0.006 | 0.005 | 0.000 |
| Proteobacteria | Betaproteobacteria | Burkholderiales | Oxalobacteraceae | Herminiimonas_1 | 0.008 | 0.000 | 0.018 | 0.000 | 0.000 | 0.000 |
| Proteobacteria | Betaproteobacteria | Burkholderiales | Oxalobacteraceae | Herpetosiphon | 0.000 | 0.006 | 0.008 | 0.000 | 0.000 | 0.000 |
| Proteobacteria | Betaproteobacteria | Burkholderiales | Oxalobacteraceae | Janthinobacterium_1 | 0.008 | 0.000 | 0.054 | 0.006 | 0.000 | 0.000 |
| Proteobacteria | Betaproteobacteria | Burkholderiales | Oxalobacteraceae | Janthinobacterium_2 | 0.000 | 0.000 | 0.014 | 0.000 | 0.000 | 0.000 |
| Proteobacteria | Betaproteobacteria | Burkholderiales | Oxalobacteraceae | Jeotgalicoccus | 0.000 | 0.000 | 0.008 | 0.000 | 0.000 | 0.000 |
| Proteobacteria | Betaproteobacteria | Burkholderiales | Oxalobacteraceae | Massilia_1 | 0.008 | 0.000 | 0.004 | 0.000 | 0.000 | 0.000 |
| Proteobacteria | Betaproteobacteria | Burkholderiales | Oxalobacteraceae | Massilia_10 | 0.000 | 0.000 | 0.012 | 0.000 | 0.000 | 0.000 |
| Proteobacteria | Betaproteobacteria | Burkholderiales | Oxalobacteraceae | Massilia_5 | 0.008 | 0.000 | 0.004 | 0.000 | 0.000 | 0.000 |
| Proteobacteria | Betaproteobacteria | Burkholderiales | Oxalobacteraceae | Massilia_6 | 0.000 | 0.000 | 0.004 | 0.000 | 0.000 | 0.000 |
| Proteobacteria | Betaproteobacteria | Burkholderiales | Oxalobacteraceae | Massilia_8 | 0.000 | 0.000 | 0.004 | 0.000 | 0.000 | 0.000 |
| Proteobacteria | Betaproteobacteria | Burkholderiales | Oxalobacteraceae | Massilia_9 | 0.008 | 0.006 | 0.018 | 0.006 | 0.000 | 0.000 |
| Proteobacteria | Betaproteobacteria | Burkholderiales | Oxalobacteraceae | Massilia_sp_a | 0.000 | 0.000 | 0.004 | 0.000 | 0.000 | 0.000 |
| Proteobacteria | Betaproteobacteria | Burkholderiales | Oxalobacteraceae | Novispirillum | 0.024 | 0.012 | 0.004 | 0.006 | 0.000 | 0.000 |
| Proteobacteria | Betaproteobacteria | Burkholderiales | Oxalobacteraceae | Oxalobacter | 0.024 | 0.012 | 0.018 | 0.000 | 0.000 | 0.000 |
| Proteobacteria | Betaproteobacteria | Burkholderiales | Oxalobacteraceae | Telluria | 0.000 | 0.000 | 0.004 | 0.000 | 0.000 | 0.000 |
| Proteobacteria | Gammaproteobacteria_1 | Pasteurellales | Pasteurellaceae | Actinobacillus_2 | 0.000 | 0.000 | 0.004 | 0.000 | 0.000 | 0.000 |

**Table S9 continued.**

| | | | | | | | | | | |
|---|---|---|---|---|---|---|---|---|---|---|
| Proteobacteria | Gammaproteobacteria_1 | Pasteurellales | Pasteurellaceae | Actinobacillus_11 | 0.000 | 0.000 | 0.008 | 0.000 | 0.000 | 0.000 |
| Proteobacteria | Gammaproteobacteria_1 | Pasteurellales | Pasteurellaceae | Actinobacillus_4 | 0.000 | 0.000 | 0.006 | 0.000 | 0.000 | 0.000 |
| Proteobacteria | Gammaproteobacteria_1 | Pasteurellales | Pasteurellaceae | Actinobacillus_5 | 0.000 | 0.000 | 0.008 | 0.000 | 0.000 | 0.000 |
| Proteobacteria | Gammaproteobacteria_1 | Pasteurellales | Pasteurellaceae | Actinobacillus_6 | 0.000 | 0.000 | 0.022 | 0.000 | 0.000 | 0.000 |
| Proteobacteria | Gammaproteobacteria_1 | Pasteurellales | Pasteurellaceae | Actinobacillus_7 | 0.000 | 0.000 | 0.010 | 0.000 | 0.000 | 0.000 |
| Proteobacteria | Gammaproteobacteria_1 | Pasteurellales | Pasteurellaceae | Actinobacillus_8 | 0.000 | 0.000 | 0.006 | 0.000 | 0.000 | 0.000 |
| Proteobacteria | Gammaproteobacteria_1 | Pasteurellales | Pasteurellaceae | Actinobacillus_9 | 0.000 | 0.000 | 0.006 | 0.000 | 0.000 | 0.000 |
| Proteobacteria | Gammaproteobacteria_1 | Pasteurellales | Pasteurellaceae | Aggregatibacter_2 | 0.000 | 0.000 | 0.004 | 0.000 | 0.000 | 0.000 |
| Proteobacteria | Gammaproteobacteria_1 | Pasteurellales | Pasteurellaceae | Avibacterium_2 | 0.000 | 0.000 | 0.012 | 0.000 | 0.000 | 0.000 |
| Proteobacteria | Gammaproteobacteria_1 | Pasteurellales | Pasteurellaceae | Haemophilus_1 | 0.000 | 0.000 | 0.032 | 0.000 | 0.000 | 0.000 |
| Proteobacteria | Gammaproteobacteria_1 | Pasteurellales | Pasteurellaceae | Haemophilus_2 | 0.000 | 0.006 | 0.004 | 0.000 | 0.000 | 0.000 |
| Proteobacteria | Gammaproteobacteria_1 | Pasteurellales | Pasteurellaceae | Haemophilus_4 | 0.000 | 0.000 | 0.004 | 0.000 | 0.000 | 0.000 |
| Proteobacteria | Gammaproteobacteria_1 | Pasteurellales | Pasteurellaceae | Haemophilus_5 | 0.000 | 0.000 | 0.014 | 0.000 | 0.000 | 0.000 |
| Proteobacteria | Gammaproteobacteria_1 | Pasteurellales | Pasteurellaceae | Haemophilus_6 | 0.000 | 0.000 | 0.004 | 0.000 | 0.000 | 0.000 |
| Proteobacteria | Gammaproteobacteria_1 | Pasteurellales | Pasteurellaceae | Haemophilus_7 | 0.016 | 0.000 | 0.018 | 0.000 | 0.000 | 0.000 |
| Proteobacteria | Gammaproteobacteria_1 | Pasteurellales | Pasteurellaceae | Histophilus | 0.000 | 0.000 | 0.008 | 0.000 | 0.000 | 0.000 |
| Proteobacteria | Gammaproteobacteria_1 | Pasteurellales | Pasteurellaceae | Mannheimia | 0.000 | 0.000 | 0.022 | 0.000 | 0.000 | 0.000 |
| Proteobacteria | Gammaproteobacteria_1 | Pasteurellales | Pasteurellaceae | Pasteurella_1 | 0.000 | 0.000 | 0.004 | 0.000 | 0.000 | 0.000 |
| Proteobacteria | Gammaproteobacteria_1 | Pasteurellales | Pasteurellaceae | Pasteurella_11 | 0.000 | 0.000 | 0.004 | 0.000 | 0.000 | 0.000 |
| Proteobacteria | Gammaproteobacteria_1 | Pasteurellales | Pasteurellaceae | Pasteurella_4 | 0.000 | 0.000 | 0.008 | 0.000 | 0.000 | 0.000 |
| Proteobacteria | Gammaproteobacteria_1 | Pasteurellales | Pasteurellaceae | Pasteurella_5 | 0.000 | 0.000 | 0.010 | 0.000 | 0.000 | 0.000 |
| Proteobacteria | Gammaproteobacteria_1 | Pasteurellales | Pasteurellaceae | Pasteurella_6 | 0.000 | 0.000 | 0.008 | 0.000 | 0.000 | 0.000 |
| Proteobacteria | Gammaproteobacteria_1 | Pasteurellales | Pasteurellaceae | Pasteurella_7 | 0.000 | 0.000 | 0.008 | 0.000 | 0.000 | 0.000 |
| Proteobacteria | Gammaproteobacteria_1 | Pasteurellales | Pasteurellaceae | Pasteurella_8 | 0.000 | 0.000 | 0.004 | 0.000 | 0.000 | 0.000 |
| Proteobacteria | Gammaproteobacteria_1 | Pasteurellales | Pasteurellaceae | Phocoenobacter | 0.000 | 0.000 | 0.004 | 0.000 | 0.000 | 0.000 |
| Proteobacteria | Gammaproteobacteria_1 | Pasteurellales | Pasteurellaceae | Volucribacter_1 | 0.000 | 0.000 | 0.006 | 0.000 | 0.000 | 0.000 |
| Proteobacteria | Gammaproteobacteria_1 | Pasteurellales | Pasteurellaceae | Volucribacter_2 | 0.000 | 0.000 | 0.006 | 0.000 | 0.000 | 0.000 |
| Proteobacteria | Deltaproteobacteria | Myxococcales | Phaselicystidaceae | Phenylobacterium | 0.057 | 0.018 | 0.028 | 0.006 | 0.000 | 0.000 |
| Proteobacteria | Alphaproteobacteria | Hyphomicrobiales | Phyllobacteriaceae | Lentilitoribacter | 0.008 | 0.000 | 0.000 | 0.000 | 0.000 | 0.000 |
| Proteobacteria | Alphaproteobacteria | Hyphomicrobiales | Phyllobacteriaceae | Thermovum | 0.008 | 0.000 | 0.000 | 0.000 | 0.000 | 0.000 |
| Proteobacteria | Alphaproteobacteria | Rhizobiales | Phyllobacteriaceae | Holdemania | 0.000 | 0.000 | 0.004 | 0.000 | 0.000 | 0.000 |
| Proteobacteria | Alphaproteobacteria | Rhizobiales | Phyllobacteriaceae | Nitratiruptor | 0.000 | 0.000 | 0.004 | 0.000 | 0.000 | 0.000 |
| Proteobacteria | Alphaproteobacteria | Rhizobiales | Phyllobacteriaceae | Pseudenhygromyxa | 0.008 | 0.000 | 0.000 | 0.000 | 0.000 | 0.000 |
| Proteobacteria | Alphaproteobacteria | Rhizobiales_1 | Phyllobacteriaceae | Aminobacter_1 | 0.000 | 0.006 | 0.012 | 0.000 | 0.005 | 0.006 |
| Proteobacteria | Alphaproteobacteria | Rhizobiales_1 | Phyllobacteriaceae | Aminobacter_2 | 0.000 | 0.000 | 0.006 | 0.000 | 0.000 | 0.000 |
| Proteobacteria | Alphaproteobacteria | Rhizobiales_1 | Phyllobacteriaceae | Defluvibacter | 0.000 | 0.000 | 0.004 | 0.000 | 0.000 | 0.000 |
| Proteobacteria | Alphaproteobacteria | Rhizobiales_1 | Phyllobacteriaceae | Mesorhizobium_10 | 0.000 | 0.000 | 0.004 | 0.000 | 0.000 | 0.000 |
| Proteobacteria | Alphaproteobacteria | Rhizobiales_1 | Phyllobacteriaceae | Mesorhizobium_11 | 0.000 | 0.006 | 0.004 | 0.000 | 0.000 | 0.000 |
| Proteobacteria | Alphaproteobacteria | Rhizobiales_1 | Phyllobacteriaceae | Mesorhizobium_2 | 0.000 | 0.000 | 0.006 | 0.000 | 0.000 | 0.000 |
| Proteobacteria | Alphaproteobacteria | Rhizobiales_1 | Phyllobacteriaceae | Mesorhizobium_8 | 0.000 | 0.000 | 0.004 | 0.000 | 0.000 | 0.000 |

**Table S9 continued.**

| | | | | | | | | | | |
|---|---|---|---|---|---|---|---|---|---|---|
| Proteobacteria | Alphaproteobacteria | Rhizobiales_2 | Phyllobacteriaceae | Mesorhizobium_12 | 0.000 | 0.000 | 0.006 | 0.000 | 0.000 | 0.000 |
| Proteobacteria | Gammaproteobacteria | Thiotrichale | Piscirickettsiaceae | Piscirickettsiaceae | 0.000 | 0.000 | 0.004 | 0.000 | 0.000 | 0.000 |
| Proteobacteria | Gammaproteobacteria | Thiotrichales | Piscirickettsiaceae | Cycloclasticus | 0.000 | 0.000 | 0.010 | 0.000 | 0.000 | 0.000 |
| Proteobacteria | Gammaproteobacteria | Thiotrichales | Piscirickettsiaceae | endosymbionts_2 | 0.000 | 0.000 | 0.008 | 0.000 | 0.000 | 0.000 |
| Proteobacteria | Gammaproteobacteria | Thiotrichales | Piscirickettsiaceae | Mariprofundus | 0.000 | 0.000 | 0.008 | 0.000 | 0.000 | 0.000 |
| Proteobacteria | Gammaproteobacteria | Thiotrichales | Piscirickettsiaceae | Methylophaga | 0.000 | 0.000 | 0.022 | 0.000 | 0.000 | 0.000 |
| Proteobacteria | Gammaproteobacteria | Thiotrichales | Piscirickettsiaceae | Thioalkalimicrobium | 0.000 | 0.000 | 0.004 | 0.000 | 0.000 | 0.000 |
| Proteobacteria | Gammaproteobacteria | Thiotrichales | Piscirickettsiaceae | Thiomicrospira_1 | 0.000 | 0.000 | 0.024 | 0.000 | 0.000 | 0.000 |
| Proteobacteria | Deltaproteobacteria | Myxococcales | Polyangiaceae | Byssovorax | 0.008 | 0.000 | 0.004 | 0.000 | 0.000 | 0.000 |
| Proteobacteria | Deltaproteobacteria | Myxococcales | Polyangiaceae | Chondromyces | 0.008 | 0.000 | 0.000 | 0.000 | 0.000 | 0.000 |
| Proteobacteria | Deltaproteobacteria | Myxococcales | Polyangiaceae | Jahnella | 0.008 | 0.000 | 0.000 | 0.012 | 0.000 | 0.000 |
| Proteobacteria | Deltaproteobacteria | Myxococcales | Polyangiaceae | Ralstonia | 0.081 | 0.018 | 0.044 | 0.006 | 0.025 | 0.012 |
| Proteobacteria | Deltaproteobacteria | Myxococcales | Polyangiaceae | Sorangium_1 | 0.008 | 0.000 | 0.004 | 0.000 | 0.000 | 0.000 |
| Proteobacteria | Deltaproteobacteria | Myxococcales | Polyangiaceae | Sorangium_2 | 0.008 | 0.006 | 0.012 | 0.000 | 0.000 | 0.000 |
| Proteobacteria | Deltaproteobacteria | Myxococcales | Polyangiaceae | SP3-e08 | 0.008 | 0.000 | 0.012 | 0.000 | 0.000 | 0.000 |
| Proteobacteria | Deltaproteobacteria | Myxococcales | Polyangiaceae | Sphaerisporangium | 0.000 | 0.000 | 0.000 | 0.000 | 0.005 | 0.000 |
| Proteobacteria | Gammaproteobacteria | Cellvibrionales | Porticoccaceae | Povalibacter | 0.008 | 0.006 | 0.006 | 0.012 | 0.000 | 0.000 |
| Proteobacteria | Betaproteobacteria | Procabacteriales | Procabacteriaceae | Procabacter | 0.008 | 0.000 | 0.004 | 0.000 | 0.000 | 0.000 |
| Proteobacteria | Gammaproteobacteria | Alteromonadales | Pseudoalteromonadaceae | Pseudoaminobacter | 0.000 | 0.000 | 0.010 | 0.000 | 0.000 | 0.000 |
| Proteobacteria | Gammaproteobacteria_1 | Alteromonadales_1 | Pseudoalteromonadaceae | Algicola | 0.000 | 0.000 | 0.012 | 0.000 | 0.000 | 0.000 |
| Proteobacteria | Gammaproteobacteria | Pseudomonadales | Pseudomonadaceae | Azomonas | 0.000 | 0.006 | 0.000 | 0.000 | 0.000 | 0.000 |
| Proteobacteria | Gammaproteobacteria | Pseudomonadales | Pseudomonadaceae | Azotobacter | 0.000 | 0.000 | 0.002 | 0.000 | 0.005 | 0.006 |
| Proteobacteria | Gammaproteobacteria | Pseudomonadales | Pseudomonadaceae | Oblitimonas | 0.008 | 0.000 | 0.000 | 0.000 | 0.000 | 0.000 |
| Proteobacteria | Gammaproteobacteria | Pseudomonadales | Pseudomonadaceae | Permianibacter | 0.000 | 0.000 | 0.000 | 0.006 | 0.000 | 0.000 |
| Proteobacteria | Gammaproteobacteria_0 | Pseudomonadales | Pseudomonadaceae | Pseudomonas_4 | 0.163 | 0.062 | 0.044 | 0.006 | 0.005 | 0.006 |
| Proteobacteria | Gammaproteobacteria_1 | Pseudomonadales | Pseudomonadaceae | Azotobacter-Azomonas | 0.033 | 0.012 | 0.036 | 0.000 | 0.005 | 0.000 |
| Proteobacteria | Gammaproteobacteria_1 | Pseudomonadales | Pseudomonadaceae | Cellvibrio | 0.024 | 0.000 | 0.022 | 0.012 | 0.000 | 0.000 |
| Proteobacteria | Gammaproteobacteria_1 | Pseudomonadales | Pseudomonadaceae | Insect_cluster_I | 0.000 | 0.000 | 0.020 | 0.000 | 0.000 | 0.000 |
| Proteobacteria | Gammaproteobacteria_1 | Pseudomonadales | Pseudomonadaceae | Insect_cluster_II | 0.000 | 0.000 | 0.006 | 0.000 | 0.000 | 0.000 |
| Proteobacteria | Gammaproteobacteria_1 | Pseudomonadales | Pseudomonadaceae | Pseudomonas_2 | 0.268 | 0.253 | 0.314 | 0.258 | 0.279 | 0.369 |
| Proteobacteria | Gammaproteobacteria_1 | Pseudomonadales | Pseudomonadaceae | Pseudomonas_3 | 0.000 | 0.148 | 0.024 | 0.000 | 0.000 | 0.000 |
| Proteobacteria | Gammaproteobacteria_1 | Pseudomonadales | Pseudomonadaceae | Pseudomonas_5 | 0.008 | 0.000 | 0.012 | 0.000 | 0.000 | 0.000 |
| Proteobacteria | Gammaproteobacteria_1 | Pseudomonadales | Pseudomonadaceae | Pseudomonas_6 | 0.008 | 0.062 | 0.016 | 0.018 | 0.020 | 0.012 |
| Proteobacteria | Gammaproteobacteria_1 | Pseudomonadales | Pseudomonadaceae | Pseudomonas_7 | 0.000 | 0.000 | 0.014 | 0.006 | 0.005 | 0.000 |
| Proteobacteria | Gammaproteobacteria_1 | Pseudomonadales | Pseudomonadaceae | Pseudonocardia | 0.024 | 0.000 | 0.083 | 0.000 | 0.000 | 0.000 |
| Proteobacteria | Gammaproteobacteria_1 | Pseudomonadales | Pseudomonadaceae | Serpens | 0.000 | 0.000 | 0.006 | 0.000 | 0.000 | 0.000 |
| Proteobacteria | Gammaproteobacteria_3 | Pseudomonadales | Pseudomonadaceae | Pseudomonas_1 | 1.342 | 4.307 | 0.857 | 2.810 | 4.291 | 3.341 |
| Proteobacteria | Gammaproteobacteria | Alteromonadales | Psychromonadaceae | Pusillimonas | 0.024 | 0.000 | 0.004 | 0.024 | 0.020 | 0.036 |
| Proteobacteria | Alphaproteobacteria | Rhizobiales | Rhizobiaceae | Agromyces | 0.000 | 0.000 | 0.000 | 0.006 | 0.000 | 0.000 |
| Proteobacteria | Alphaproteobacteria | Rhizobiales | Rhizobiaceae | Ciceribacter | 0.008 | 0.000 | 0.000 | 0.000 | 0.000 | 0.000 |

**Table S9 continued.**

| | | | | | | | | | | |
|---|---|---|---|---|---|---|---|---|---|---|
| Proteobacteria | Alphaproteobacteria | Rhizobiales | Rhizobiaceae | Ensifer | 0.008 | 0.000 | 0.000 | 0.000 | 0.000 | 0.006 |
| Proteobacteria | Alphaproteobacteria | Rhizobiales | Rhizobiaceae | Shuttleworthia | 0.000 | 0.000 | 0.016 | 0.000 | 0.000 | 0.000 |
| Proteobacteria | Alphaproteobacteria | Rhizobiales_1 | Rhizobiaceae | Rhodanobacter_1 | 0.000 | 0.000 | 0.044 | 0.000 | 0.005 | 0.000 |
| Proteobacteria | Alphaproteobacteria | Rhizobiales_1 | Rhizobiaceae | Rhodanobacter_2 | 0.000 | 0.000 | 0.010 | 0.000 | 0.005 | 0.000 |
| Proteobacteria | Alphaproteobacteria | Rhizobiales_1 | Rhizobiaceae | Skermanella | 0.000 | 0.000 | 0.014 | 0.000 | 0.000 | 0.000 |
| Proteobacteria | Gammaproteobacteria | Xanthomonadales | Rhodanobacteraceae | Aquimonas | 0.000 | 0.000 | 0.004 | 0.000 | 0.000 | 0.000 |
| Proteobacteria | Gammaproteobacteria | Xanthomonadales | Rhodanobacteraceae | Frateuria | 0.000 | 0.000 | 0.006 | 0.078 | 0.075 | 0.054 |
| Proteobacteria | Gammaproteobacteria | Xanthomonadales | Rhodanobacteraceae | Fulvimonas | 0.000 | 0.000 | 0.000 | 0.018 | 0.040 | 0.036 |
| Proteobacteria | Gammaproteobacteria | Xanthomonadales | Rhodanobacteraceae | Metallibacterium | 0.000 | 0.000 | 0.000 | 0.000 | 0.005 | 0.000 |
| Proteobacteria | Gammaproteobacteria | Xanthomonadales | Rhodanobacteraceae | Tahibacter | 0.008 | 0.000 | 0.000 | 0.030 | 0.000 | 0.006 |
| Proteobacteria | Alphaproteobacteria | Rhodobacterales | Rhodobacteraceae | Acuticoccus | 0.008 | 0.000 | 0.000 | 0.000 | 0.000 | 0.000 |
| Proteobacteria | Alphaproteobacteria | Rhodobacterales | Rhodobacteraceae | Ahrensia | 0.000 | 0.000 | 0.010 | 0.000 | 0.000 | 0.000 |
| Proteobacteria | Alphaproteobacteria | Rhodobacterales | Rhodobacteraceae | Albimonas | 0.000 | 0.000 | 0.006 | 0.000 | 0.000 | 0.000 |
| Proteobacteria | Alphaproteobacteria | Rhodobacterales | Rhodobacteraceae | Gemmobacter | 0.000 | 0.000 | 0.004 | 0.000 | 0.000 | 0.000 |
| Proteobacteria | Alphaproteobacteria | Rhodobacterales | Rhodobacteraceae | Kibdelosporangium | 0.000 | 0.000 | 0.002 | 0.000 | 0.000 | 0.000 |
| Proteobacteria | Alphaproteobacteria | Rhodobacterales | Rhodobacteraceae | Labrys | 0.008 | 0.006 | 0.010 | 0.000 | 0.000 | 0.024 |
| Proteobacteria | Alphaproteobacteria | Rhodobacterales | Rhodobacteraceae | Lutispora | 0.000 | 0.006 | 0.008 | 0.000 | 0.000 | 0.000 |
| Proteobacteria | Alphaproteobacteria | Rhodobacterales | Rhodobacteraceae | Nesterenkonia | 0.000 | 0.000 | 0.030 | 0.000 | 0.000 | 0.000 |
| Proteobacteria | Alphaproteobacteria | Rhodobacterales | Rhodobacteraceae | Polymorphum | 0.016 | 0.000 | 0.000 | 0.000 | 0.000 | 0.000 |
| Proteobacteria | Alphaproteobacteria | Rhodobacterales | Rhodobacteraceae | Pseudoxanthomonas | 0.098 | 0.179 | 0.040 | 0.006 | 0.005 | 0.145 |
| Proteobacteria | Alphaproteobacteria | Rhodobacterales | Rhodobacteraceae | Rhodobacter_2 | 0.000 | 0.000 | 0.028 | 0.000 | 0.000 | 0.000 |
| Proteobacteria | Alphaproteobacteria | Rhodobacterales | Rhodobacteraceae | Rhodobacter_3 | 0.000 | 0.000 | 0.018 | 0.000 | 0.000 | 0.000 |
| Proteobacteria | Alphaproteobacteria | Rhodobacterales | Rhodobacteraceae | Rhodobacter_4 | 0.000 | 0.000 | 0.006 | 0.000 | 0.000 | 0.000 |
| Proteobacteria | Alphaproteobacteria | Rhodobacterales | Rhodobacteraceae | Rhodobacter_6 | 0.000 | 0.000 | 0.004 | 0.000 | 0.000 | 0.000 |
| Proteobacteria | Alphaproteobacteria | Rhodobacterales | Rhodobacteraceae | Rhodobium | 0.000 | 0.000 | 0.002 | 0.000 | 0.000 | 0.000 |
| Proteobacteria | Alphaproteobacteria | Rhodobacterales | Rhodobacteraceae | Rhodobium_2 | 0.016 | 0.000 | 0.024 | 0.006 | 0.005 | 0.000 |
| Proteobacteria | Alphaproteobacteria | Rhodobacterales | Rhodobacteraceae | Rickettsiella | 0.000 | 0.000 | 0.010 | 0.000 | 0.005 | 0.000 |
| Proteobacteria | Alphaproteobacteria | Rhodobacterales | Rhodobacteraceae | Roseobacter_clade | 0.008 | 0.000 | 0.326 | 0.000 | 0.000 | 0.000 |
| Proteobacteria | Alphaproteobacteria | Rhodobacterales | Rhodobacteraceae | Stenotrophomonas | 0.106 | 0.062 | 0.125 | 0.006 | 0.025 | 0.157 |
| Proteobacteria | Alphaproteobacteria | Rhodobacterales | Rhodobacteraceae | Tsukamurella | 0.000 | 0.000 | 0.004 | 0.000 | 0.000 | 0.000 |
| Proteobacteria | Alphaproteobacteria | Rhodobacterales | Rhodobacteraceae_1 | Alcanivorax | 0.000 | 0.000 | 0.064 | 0.000 | 0.000 | 0.000 |
| Proteobacteria | Alphaproteobacteria | Rhodobacterales | Rhodobacteraceae_1 | Amaricoccus | 0.000 | 0.000 | 0.014 | 0.000 | 0.000 | 0.000 |
| Proteobacteria | Alphaproteobacteria | Rhodobacterales | Rhodobacteraceae_1 | Catellibacterium | 0.000 | 0.000 | 0.004 | 0.000 | 0.000 | 0.000 |
| Proteobacteria | Alphaproteobacteria | Rhodobacterales | Rhodobacteraceae_1 | Dinoroseobacter | 0.000 | 0.000 | 0.006 | 0.000 | 0.000 | 0.000 |
| Proteobacteria | Alphaproteobacteria | Rhodobacterales | Rhodobacteraceae_1 | Diverse_a | 0.000 | 0.000 | 0.008 | 0.000 | 0.000 | 0.000 |
| Proteobacteria | Alphaproteobacteria | Rhodobacterales | Rhodobacteraceae_1 | Donghicola | 0.000 | 0.000 | 0.004 | 0.000 | 0.000 | 0.000 |
| Proteobacteria | Alphaproteobacteria | Rhodobacterales | Rhodobacteraceae_1 | Loktanella_1 | 0.000 | 0.000 | 0.004 | 0.000 | 0.000 | 0.000 |
| Proteobacteria | Alphaproteobacteria | Rhodobacterales | Rhodobacteraceae_1 | Methylarcula | 0.000 | 0.000 | 0.004 | 0.000 | 0.000 | 0.000 |
| Proteobacteria | Alphaproteobacteria | Rhodobacterales | Rhodobacteraceae_1 | Nereida | 0.000 | 0.000 | 0.006 | 0.000 | 0.000 | 0.000 |
| Proteobacteria | Alphaproteobacteria | Rhodobacterales | Rhodobacteraceae_1 | Palleronia | 0.000 | 0.000 | 0.004 | 0.000 | 0.000 | 0.000 |

**Table S9 continued.**

| | | | | | | | | | | |
|---|---|---|---|---|---|---|---|---|---|---|
| Proteobacteria | Alphaproteobacteria | Rhodobacterales | Rhodobacteraceae_1 | Paludibacter | 0.008 | 0.000 | 0.073 | 0.000 | 0.000 | 0.000 |
| Proteobacteria | Alphaproteobacteria | Rhodobacterales | Rhodobacteraceae_1 | Paracoccus_1 | 0.008 | 0.000 | 0.069 | 0.000 | 0.000 | 0.000 |
| Proteobacteria | Alphaproteobacteria | Rhodobacterales | Rhodobacteraceae_1 | Paracoccus_2 | 0.000 | 0.000 | 0.010 | 0.000 | 0.000 | 0.000 |
| Proteobacteria | Alphaproteobacteria | Rhodobacterales | Rhodobacteraceae_1 | Paracoccus_4 | 0.000 | 0.000 | 0.008 | 0.000 | 0.000 | 0.000 |
| Proteobacteria | Alphaproteobacteria | Rhodobacterales | Rhodobacteraceae_1 | Paracoccus_5 | 0.000 | 0.000 | 0.016 | 0.000 | 0.000 | 0.000 |
| Proteobacteria | Alphaproteobacteria | Rhodobacterales | Rhodobacteraceae_1 | Paracoccus_6 | 0.000 | 0.000 | 0.006 | 0.000 | 0.000 | 0.000 |
| Proteobacteria | Alphaproteobacteria | Rhodobacterales | Rhodobacteraceae_1 | Pelagibaca | 0.000 | 0.000 | 0.004 | 0.000 | 0.000 | 0.000 |
| Proteobacteria | Alphaproteobacteria | Rhodobacterales | Rhodobacteraceae_1 | Pseudorhodobacter_2 | 0.000 | 0.000 | 0.004 | 0.000 | 0.000 | 0.000 |
| Proteobacteria | Alphaproteobacteria | Rhodobacterales | Rhodobacteraceae_1 | Rhodobacter_5 | 0.000 | 0.000 | 0.004 | 0.000 | 0.000 | 0.000 |
| Proteobacteria | Alphaproteobacteria | Rhodobacterales | Rhodobacteraceae_1 | Rhodovulum_2 | 0.000 | 0.000 | 0.032 | 0.000 | 0.000 | 0.000 |
| Proteobacteria | Alphaproteobacteria | Rhodobacterales | Rhodobacteraceae_1 | Roseisalinus | 0.000 | 0.000 | 0.004 | 0.000 | 0.000 | 0.000 |
| Proteobacteria | Alphaproteobacteria | Rhodobacterales | Rhodobacteraceae_1 | Roseobacter_1 | 0.000 | 0.000 | 0.012 | 0.000 | 0.000 | 0.000 |
| Proteobacteria | Alphaproteobacteria | Rhodobacterales | Rhodobacteraceae_1 | Roseobacter_2 | 0.000 | 0.000 | 0.004 | 0.000 | 0.000 | 0.000 |
| Proteobacteria | Alphaproteobacteria | Rhodobacterales | Rhodobacteraceae_1 | Roseomonas | 0.000 | 0.031 | 0.000 | 0.000 | 0.000 | 0.000 |
| Proteobacteria | Alphaproteobacteria | Rhodobacterales | Rhodobacteraceae_1 | Roseovarius | 0.000 | 0.000 | 0.004 | 0.000 | 0.000 | 0.000 |
| Proteobacteria | Alphaproteobacteria | Rhodobacterales | Rhodobacteraceae_1 | Roseovarius_3 | 0.000 | 0.000 | 0.008 | 0.000 | 0.000 | 0.000 |
| Proteobacteria | Alphaproteobacteria | Rhodobacterales | Rhodobacteraceae_1 | Rubrimonas | 0.000 | 0.000 | 0.006 | 0.000 | 0.000 | 0.000 |
| Proteobacteria | Alphaproteobacteria | Rhodobacterales | Rhodobacteraceae_1 | Sediminimonas | 0.000 | 0.000 | 0.004 | 0.000 | 0.000 | 0.000 |
| Proteobacteria | Alphaproteobacteria | Rhodobacterales | Rhodobacteraceae_1 | Sulfitobacter_6 | 0.000 | 0.000 | 0.004 | 0.000 | 0.000 | 0.000 |
| Proteobacteria | Alphaproteobacteria | Rhodobacterales | Rhodobacteraceae_1 | Tateyamaria | 0.000 | 0.000 | 0.006 | 0.000 | 0.000 | 0.000 |
| Proteobacteria | Alphaproteobacteria | Rhodobacterales | Rhodobacteraceae_1 | Thalassobacter | 0.000 | 0.000 | 0.008 | 0.000 | 0.000 | 0.000 |
| Proteobacteria | Alphaproteobacteria | Rhodobacterales | Rhodobacteraceae_1 | Thalassobius | 0.000 | 0.000 | 0.022 | 0.000 | 0.000 | 0.000 |
| Proteobacteria | Alphaproteobacteria | Rhodobacterales | Rhodobacteraceae_1 | Thalassococcus | 0.000 | 0.000 | 0.006 | 0.000 | 0.000 | 0.000 |
| Proteobacteria | Alphaproteobacteria | Rhodobacterales | Rhodobacteraceae_1 | Thioclava | 0.000 | 0.000 | 0.004 | 0.000 | 0.000 | 0.000 |
| Proteobacteria | Alphaproteobacteria | Rhodobacterales | Rhodobacteraceae_1 | Yangia | 0.000 | 0.000 | 0.004 | 0.000 | 0.000 | 0.000 |
| Proteobacteria | Alphaproteobacteria | Rhodobacterales | Rhodobacteraceae_2 | Paracoccus_3 | 0.000 | 0.000 | 0.006 | 0.000 | 0.000 | 0.000 |
| Proteobacteria | Alphaproteobacteria | Rhodobacterales | Rhodobacteraceae_3 | Pantoea | 0.285 | 5.613 | 1.139 | 2.875 | 3.649 | 4.176 |
| Proteobacteria | Alphaproteobacteria | Rhodobacterales | Rhodobacteraceae_3 | Rhodothalassium | 0.000 | 0.000 | 0.004 | 0.000 | 0.000 | 0.000 |
| Proteobacteria | Alphaproteobacteria | Rhizobiales | Rhodobiaceae | Afifella | 0.000 | 0.000 | 0.004 | 0.000 | 0.000 | 0.000 |
| Proteobacteria | Alphaproteobacteria | Rhizobiales | Rhodobiaceae | Parviterribacter | 0.041 | 0.000 | 0.000 | 0.000 | 0.000 | 0.000 |
| Proteobacteria | Alphaproteobacteria | Rhizobiales | Rhodobiaceae | Rhodococcus | 0.000 | 0.000 | 0.000 | 0.000 | 0.005 | 0.006 |
| Proteobacteria | Betaproteobacteria | Rhodocyclales | Rhodocyclaceae | Dechloromonas | 0.000 | 0.000 | 0.008 | 0.000 | 0.000 | 0.000 |
| Proteobacteria | Betaproteobacteria | Rhodocyclales | Rhodocyclaceae | Dechloromonas_1 | 0.016 | 0.006 | 0.030 | 0.000 | 0.000 | 0.000 |
| Proteobacteria | Betaproteobacteria | Rhodocyclales | Rhodocyclaceae | Rhodoligotrophos | 0.008 | 0.000 | 0.000 | 0.000 | 0.000 | 0.000 |
| Proteobacteria | Betaproteobacteria | Rhodocyclales | Rhodocyclaceae_1 | Sterolibacterium | 0.000 | 0.000 | 0.004 | 0.000 | 0.000 | 0.000 |
| Proteobacteria | Betaproteobacteria | Rhodocyclales | Rhodocyclaceae_2 | Azoarcus_1 | 0.000 | 0.000 | 0.010 | 0.000 | 0.000 | 0.000 |
| Proteobacteria | Betaproteobacteria | Rhodocyclales | Rhodocyclaceae_2 | Azohydromonas_1 | 0.000 | 0.000 | 0.004 | 0.000 | 0.000 | 0.000 |
| Proteobacteria | Betaproteobacteria | Rhodocyclales | Rhodocyclaceae_2 | Azovibrio | 0.000 | 0.000 | 0.006 | 0.000 | 0.000 | 0.000 |
| Proteobacteria | Betaproteobacteria | Rhodocyclales | Rhodocyclaceae_3 | Azohydromonas_2 | 0.000 | 0.000 | 0.008 | 0.000 | 0.000 | 0.000 |
| Proteobacteria | Betaproteobacteria | Rhodocyclales | Rhodocyclaceae_3 | Ideonella | 0.016 | 0.000 | 0.000 | 0.000 | 0.000 | 0.000 |

**Table S9 continued.**

| | | | | | | | | | | |
|---|---|---|---|---|---|---|---|---|---|---|
| Proteobacteria | Betaproteobacteria | Rhodocyclales | Rhodocyclaceae_3 | Prosthecobacter | 0.000 | 0.000 | 0.066 | 0.000 | 0.000 | 0.000 |
| Proteobacteria | Alphaproteobacteria | Rhodospirillales | Rhodospirilaceae | Pelistega | 0.008 | 0.000 | 0.000 | 0.012 | 0.015 | 0.018 |
| Proteobacteria | Alphaproteobacteria | Rhodospirillales | Rhodospirillaceae | Aetherobacter | 0.008 | 0.000 | 0.000 | 0.000 | 0.000 | 0.000 |
| Proteobacteria | Alphaproteobacteria | Rhodospirillales | Rhodospirillaceae | Aggregatibacter_1 | 0.000 | 0.000 | 0.006 | 0.000 | 0.000 | 0.000 |
| Proteobacteria | Alphaproteobacteria | Rhodospirillales | Rhodospirillaceae | Defluviicoccus | 0.008 | 0.000 | 0.040 | 0.000 | 0.000 | 0.000 |
| Proteobacteria | Alphaproteobacteria | Rhodospirillales | Rhodospirillaceae | Desertibacter | 0.008 | 0.000 | 0.000 | 0.000 | 0.000 | 0.000 |
| Proteobacteria | Alphaproteobacteria | Rhodospirillales | Rhodospirillaceae | Dongia | 0.008 | 0.000 | 0.000 | 0.006 | 0.000 | 0.000 |
| Proteobacteria | Alphaproteobacteria | Rhodospirillales | Rhodospirillaceae | Haemophilus | 0.024 | 0.000 | 0.000 | 0.006 | 0.000 | 0.000 |
| Proteobacteria | Alphaproteobacteria | Rhodospirillales | Rhodospirillaceae | Lacibacterium | 0.000 | 0.006 | 0.000 | 0.000 | 0.000 | 0.000 |
| Proteobacteria | Alphaproteobacteria | Rhodospirillales | Rhodospirillaceae | Nitratifractor | 0.000 | 0.000 | 0.006 | 0.000 | 0.000 | 0.000 |
| Proteobacteria | Alphaproteobacteria | Rhodospirillales | Rhodospirillaceae | Novosphingobium | 0.065 | 0.080 | 0.024 | 0.012 | 0.010 | 0.018 |
| Proteobacteria | Alphaproteobacteria | Rhodospirillales | Rhodospirillaceae | Rickettsia | 0.008 | 0.000 | 0.024 | 0.000 | 0.010 | 0.000 |
| Proteobacteria | Alphaproteobacteria | Rhodospirillales | Rhodospirillaceae | Tatlockia | 0.000 | 0.000 | 0.004 | 0.000 | 0.000 | 0.000 |
| Proteobacteria | Alphaproteobacteria | Rhodospirillales | Rhodospirillaceae | Treponema | 0.024 | 0.006 | 0.000 | 0.000 | 0.000 | 0.000 |
| Proteobacteria | Alphaproteobacteria | Rhodospirillales_1 | Rhodospirillaceae | Magnetospirillum_1 | 0.000 | 0.000 | 0.004 | 0.000 | 0.000 | 0.000 |
| Proteobacteria | Alphaproteobacteria | Rhodospirillales_1 | Rhodospirillaceae | Magnetospirillum_2 | 0.000 | 0.000 | 0.008 | 0.000 | 0.000 | 0.000 |
| Proteobacteria | Alphaproteobacteria | Rhodospirillales_1 | Rhodospirillaceae | Phaeospirillum | 0.000 | 0.000 | 0.004 | 0.000 | 0.000 | 0.000 |
| Proteobacteria | Alphaproteobacteria | Rhodospirillales_1 | Rhodospirillaceae | Roseospira | 0.008 | 0.000 | 0.010 | 0.000 | 0.000 | 0.000 |
| Proteobacteria | Alphaproteobacteria | Rhodospirillales_1 | Rhodospirillaceae | Telmatospirillum | 0.000 | 0.000 | 0.006 | 0.000 | 0.000 | 0.000 |
| Proteobacteria | Alphaproteobacteria | Rhodospirillales_1 | Rhodospirillaceae | Tenacibaculum_2 | 0.000 | 0.000 | 0.056 | 0.000 | 0.000 | 0.000 |
| Proteobacteria | Alphaproteobacteria | Rhodspirillales | Rhodospirillaceae | Insect_cluster | 0.927 | 0.481 | 0.097 | 0.138 | 0.005 | 0.000 |
| Proteobacteria | Alphaproteobacteria | Rhodspirillales | Rhodospirillaceae | Limnobacter | 0.000 | 0.000 | 0.016 | 0.006 | 0.005 | 0.000 |
| Proteobacteria | Alphaproteobacteria | Rhodspirillales | Rhodospirillaceae | Magnetovibrio | 0.057 | 0.018 | 0.000 | 0.006 | 0.000 | 0.000 |
| Proteobacteria | Alphaproteobacteria | Rhodspirillales | Rhodospirillaceae | Marmoricola | 0.008 | 0.006 | 0.028 | 0.006 | 0.000 | 0.000 |
| Proteobacteria | Alphaproteobacteria | Rhodospirillales_2 | Rhodospirillaceae_1 | Thalassobaculum | 0.000 | 0.000 | 0.004 | 0.000 | 0.000 | 0.000 |
| Proteobacteria | Alphaproteobacteria | Rhodospirillales_2 | Rhodospirillaceae_2 | Slackia | 0.000 | 0.000 | 0.022 | 0.000 | 0.000 | 0.000 |
| Proteobacteria | Alphaproteobacteria | Rhodospirillales | Rhodspirillaceae | Elstera | 0.016 | 0.006 | 0.000 | 0.000 | 0.000 | 0.000 |
| Proteobacteria | Alphaproteobacteria | Rickettsiales | Rickettsiaceae | Occidentia | 0.000 | 0.000 | 0.000 | 0.000 | 0.000 | 0.006 |
| Proteobacteria | Alphaproteobacteria | Rickettsiales | Rickettsiaceae | Orientia | 0.000 | 0.000 | 0.000 | 0.000 | 0.000 | 0.006 |
| Proteobacteria | Alphaproteobacteria | Rickettsiales | Rickettsiaceae | Rikenella | 0.008 | 0.006 | 0.000 | 0.006 | 0.000 | 0.000 |
| Proteobacteria | Gammaproteobacteria | Oceanospirillales | Saccharospirillaceae | Rheinheimera | 0.024 | 0.006 | 0.002 | 0.012 | 0.000 | 0.000 |
| Proteobacteria | Gammaproteobacteria_1 | Oceanospirillales_2 | Saccharospirillaceae | Saccharospirillum | 0.000 | 0.000 | 0.006 | 0.000 | 0.000 | 0.000 |
| Proteobacteria | Gammaproteobacteria_2 | Salinisphaerales | Salinisphaeraceae | marine_group_ZD0417 | 0.000 | 0.000 | 0.006 | 0.000 | 0.000 | 0.000 |
| Proteobacteria | Gammaproteobacteria_2 | Salinisphaerales | Salinisphaeraceae | Salinisphaera | 0.000 | 0.000 | 0.006 | 0.000 | 0.000 | 0.000 |
| Proteobacteria | Deltaproteobacteria | Myxococcales | Sandaracinaceae | Sanguibacter | 0.000 | 0.000 | 0.026 | 0.000 | 0.000 | 0.000 |
| Proteobacteria | Gammaproteobacteria_4 | Sedimenticola | Sedimenticola | Sedimenticola | 0.000 | 0.000 | 0.018 | 0.006 | 0.000 | 0.000 |
| Proteobacteria | Alphaproteobacteria | Rhizobiales_1 | Shinella_genera_incertae_sedis | Silanimonas | 0.008 | 0.006 | 0.008 | 0.000 | 0.005 | 0.006 |
| Proteobacteria | Gammaproteobacteria | Nevskiales | Sinobacteraceae | Hydrogenoanaerobacterium | 0.073 | 0.068 | 0.012 | 0.000 | 0.000 | 0.000 |
| Proteobacteria | Gammaproteobacteria_2 | Xanthomonadales | Sinobacteraceae | marine_benthic_group_1 | 0.000 | 0.000 | 0.073 | 0.000 | 0.000 | 0.000 |
| Proteobacteria | Gammaproteobacteria_2 | Xanthomonadales | Sinobacteraceae | Niabella | 0.016 | 0.000 | 0.010 | 0.000 | 0.000 | 0.000 |

**Table S9 continued.**

| | | | | | | | | | | |
|---|---|---|---|---|---|---|---|---|---|---|
| Proteobacteria | Gammaproteobacteria_2 | Xanthomonadales | Sinobacteraceae | Solitalea | 0.008 | 0.000 | 0.008 | 0.000 | 0.000 | 0.000 |
| Proteobacteria | Alphaproteobacteria | Sneathiellales | Sneathiellaceae | Solirubrobacter | 0.195 | 0.006 | 0.032 | 0.012 | 0.000 | 0.000 |
| Proteobacteria | Alphaproteobacteria | Sphingomonadales | Sphingomonadaceae | Blastomonas | 0.016 | 0.000 | 0.010 | 0.000 | 0.000 | 0.000 |
| Proteobacteria | Alphaproteobacteria | Sphingomonadales | Sphingomonadaceae | Herbaspirillum | 0.008 | 0.006 | 0.000 | 0.006 | 0.000 | 0.006 |
| Proteobacteria | Alphaproteobacteria | Sphingomonadales | Sphingomonadaceae | Novosphingobium_2 | 0.041 | 0.043 | 0.101 | 0.006 | 0.005 | 0.006 |
| Proteobacteria | Alphaproteobacteria | Sphingomonadales | Sphingomonadaceae | Rhizorhabdus | 0.008 | 0.000 | 0.000 | 0.000 | 0.000 | 0.000 |
| Proteobacteria | Alphaproteobacteria | Sphingomonadales | Sphingomonadaceae | Sandarakinorhabdus | 0.000 | 0.000 | 0.008 | 0.000 | 0.000 | 0.000 |
| Proteobacteria | Alphaproteobacteria | Sphingomonadales | Sphingomonadaceae | Sarcina | 0.008 | 0.000 | 0.085 | 0.000 | 0.000 | 0.000 |
| Proteobacteria | Alphaproteobacteria | Sphingomonadales | Sphingomonadaceae | Sphingomicrobium | 0.008 | 0.000 | 0.000 | 0.000 | 0.000 | 0.000 |
| Proteobacteria | Alphaproteobacteria | Sphingomonadales | Sphingomonadaceae | Sphingomonas | 0.350 | 0.062 | 0.008 | 0.006 | 0.025 | 0.036 |
| Proteobacteria | Alphaproteobacteria | Sphingomonadales | Sphingomonadaceae | Sphingomonas_1 | 0.000 | 0.006 | 0.008 | 0.000 | 0.000 | 0.000 |
| Proteobacteria | Alphaproteobacteria | Sphingomonadales | Sphingomonadaceae | Sphingomonas_2 | 0.268 | 0.012 | 0.097 | 0.012 | 0.000 | 0.006 |
| Proteobacteria | Alphaproteobacteria | Sphingomonadales | Sphingomonadaceae | Sphingomonas_3 | 0.130 | 0.031 | 0.169 | 0.018 | 0.005 | 0.012 |
| Proteobacteria | Alphaproteobacteria | Sphingomonadales | Sphingomonadaceae | Sphingomonas_4 | 0.000 | 0.000 | 0.004 | 0.000 | 0.000 | 0.000 |
| Proteobacteria | Alphaproteobacteria | Sphingomonadales | Sphingomonadaceae | Sphingopyxis | 0.016 | 0.000 | 0.000 | 0.000 | 0.000 | 0.006 |
| Proteobacteria | Alphaproteobacteria | Sphingomonadales | Sphingomonadaceae | Sphingopyxis_1 | 0.016 | 0.006 | 0.052 | 0.000 | 0.005 | 0.000 |
| Proteobacteria | Alphaproteobacteria | Sphingomonadales | Sphingomonadaceae | Sphingopyxis_2 | 0.000 | 0.000 | 0.004 | 0.000 | 0.000 | 0.000 |
| Proteobacteria | Alphaproteobacteria | Sphingomonadales | Sphingomonadaceae | Sphingopyxis_3 | 0.000 | 0.000 | 0.006 | 0.000 | 0.000 | 0.000 |
| Proteobacteria | Alphaproteobacteria | Sphingomonadales | Sphingomonadaceae | Sphingosinicella | 0.008 | 0.000 | 0.004 | 0.000 | 0.000 | 0.000 |
| Proteobacteria | Alphaproteobacteria | Sphingomonadales | Sphingomonadaceae | Spirochaeta | 0.057 | 0.037 | 0.478 | 0.018 | 0.000 | 0.000 |
| Proteobacteria | Alphaproteobacteria | Sphingomonadales | Sphingomonadaceae | Zymomonas | 0.000 | 0.000 | 0.012 | 0.000 | 0.000 | 0.000 |
| Proteobacteria | Betaproteobacteria | Nitrosomonadales | Spirillaceae | Spirillum | 0.000 | 0.006 | 0.000 | 0.000 | 0.000 | 0.000 |
| Proteobacteria | Gammaproteobacteria_1 | Aeromonadales | Succinivibrionaceae | Anaerobiospirillum_2 | 0.000 | 0.000 | 0.014 | 0.000 | 0.000 | 0.000 |
| Proteobacteria | Gammaproteobacteria_1 | Aeromonadales | Succinivibrionaceae | Succinivibrio | 0.000 | 0.000 | 0.012 | 0.000 | 0.000 | 0.000 |
| Proteobacteria | Alphaproteobacteria | SAR11_clade | Surface_1 | Candidatus_Pelagibacter | 0.000 | 0.000 | 0.004 | 0.000 | 0.000 | 0.000 |
| Proteobacteria | Deltaproteobacteria | Syntrophobacterales | Syntrophaceae | Desulfobacca | 0.000 | 0.000 | 0.008 | 0.000 | 0.000 | 0.000 |
| Proteobacteria | Deltaproteobacteria | Syntrophobacterales | Syntrophaceae | Desulfomonile | 0.000 | 0.000 | 0.008 | 0.000 | 0.000 | 0.000 |
| Proteobacteria | Deltaproteobacteria | Syntrophobacterales | Syntrophaceae | Smithella | 0.008 | 0.000 | 0.018 | 0.000 | 0.000 | 0.000 |
| Proteobacteria | Deltaproteobacteria | Syntrophobacterales | Syntrophaceae | Taibaiella | 0.008 | 0.012 | 0.000 | 0.000 | 0.000 | 0.000 |
| Proteobacteria | Deltaproteobacteria | Syntrophobacterales | Syntrophobacteraceae | Desulforhabdus | 0.000 | 0.000 | 0.004 | 0.000 | 0.000 | 0.000 |
| Proteobacteria | Deltaproteobacteria | Syntrophobacterales | Syntrophobacteraceae | Syntrophobacter_1 | 0.000 | 0.000 | 0.020 | 0.000 | 0.000 | 0.000 |
| Proteobacteria | Deltaproteobacteria | Syntrophobacterales | Syntrophobacteraceae | Syntrophobacter_2 | 0.000 | 0.000 | 0.004 | 0.000 | 0.000 | 0.000 |
| Proteobacteria | Deltaproteobacteria | Syntrophobacterales | Syntrophorhabdaceae | Syntrophorhabdus | 0.000 | 0.000 | 0.016 | 0.000 | 0.000 | 0.000 |
| Proteobacteria | Gammaproteobacteria_3 | Chromatiales | Thioalkalispiraceae | Thiohalophilus | 0.000 | 0.000 | 0.006 | 0.000 | 0.000 | 0.000 |
| Proteobacteria | Gammaproteobacteria_2 | Thiohalomonas | Thiohalomonas | Thiohalomonas | 0.000 | 0.000 | 0.004 | 0.000 | 0.000 | 0.000 |
| Proteobacteria | Gammaproteobacteria | Chromatiales | Thioprofundaceae | Thioprofundum | 0.000 | 0.000 | 0.000 | 0.006 | 0.000 | 0.000 |
| Proteobacteria | Gammaproteobacteria | Thiotrichales | Thiotrichaceae | Tidjanibacter | 0.008 | 0.000 | 0.000 | 0.000 | 0.000 | 0.000 |
| Proteobacteria | Gammaproteobacteria_2 | Thiotrichales | Thiotrichaceae | Beggiatoa | 0.008 | 0.000 | 0.006 | 0.000 | 0.000 | 0.000 |
| Proteobacteria | Gammaproteobacteria_2 | Thiotrichales | Thiotrichaceae | Thioploca | 0.000 | 0.000 | 0.004 | 0.000 | 0.000 | 0.000 |
| Proteobacteria | Gammaproteobacteria_4 | Thiotrichales | Thiotrichaceae | Leucothrix | 0.000 | 0.000 | 0.014 | 0.000 | 0.000 | 0.000 |

**Table S9 continued.**

| | | | | | | | | | | |
|---|---|---|---|---|---|---|---|---|---|---|
| Proteobacteria | Betaproteobacteria | Betaproteobacteria_incertae_sedis | Unclassified | Chitinivorax | 0.008 | 0.000 | 0.000 | 0.000 | 0.000 | 0.000 |
| Proteobacteria | Betaproteobacteria | Burkholderiales | Unclassified | Paucimonas | 0.008 | 0.006 | 0.022 | 0.012 | 0.015 | 0.006 |
| Proteobacteria | Betaproteobacteria | Burkholderiales | Unclassified | Roseatales_1 | 0.000 | 0.000 | 0.004 | 0.000 | 0.000 | 0.000 |
| Proteobacteria | Betaproteobacteria | Burkholderiales | Unclassified | Roseatales_2 | 0.000 | 0.000 | 0.004 | 0.000 | 0.000 | 0.000 |
| Proteobacteria | Betaproteobacteria | Burkholderiales | Unclassified | Roseburia_1 | 0.000 | 0.000 | 0.095 | 0.000 | 0.000 | 0.000 |
| Proteobacteria | Epsilonproteobacteria | Campylobacterales | Unclassified | Nitratireductor | 0.000 | 0.000 | 0.012 | 0.000 | 0.000 | 0.000 |
| Proteobacteria | Deltaproteobacteria | Myxococcales | Unclassified | Enhygromyxa | 0.000 | 0.000 | 0.006 | 0.000 | 0.000 | 0.000 |
| Proteobacteria | Alphaproteobacteria | Rhizobiales_2 | Unclassified | Bauldia | 0.008 | 0.000 | 0.000 | 0.000 | 0.005 | 0.000 |
| Proteobacteria | Alphaproteobacteria | Rhodospirillales | Unclassified | Enhydrobacter | 0.000 | 0.000 | 0.018 | 0.000 | 0.000 | 0.000 |
| Proteobacteria | Alphaproteobacteria | Rhodospirillales | Unclassified | Rheinheimera_1 | 0.016 | 0.006 | 0.016 | 0.000 | 0.000 | 0.000 |
| Proteobacteria | Betaproteobacteria | Unclassified | Unclassified | Aquincola | 0.000 | 0.000 | 0.006 | 0.000 | 0.000 | 0.000 |
| Proteobacteria | Betaproteobacteria | Unclassified | Unclassified | Thiothrix | 0.000 | 0.000 | 0.028 | 0.006 | 0.005 | 0.000 |
| Proteobacteria | Epsilonproteobacteria | Unclassified | Unclassified | Nitrincola | 0.000 | 0.000 | 0.008 | 0.000 | 0.005 | 0.000 |
| Proteobacteria | Epsilonproteobacteria | Unclassified | Unclassified | Symploca | 0.000 | 0.000 | 0.014 | 0.000 | 0.000 | 0.000 |
| Proteobacteria | Gammaproteobacteria | Unclassified | Unclassified | Ilumatobacter | 0.016 | 0.000 | 0.008 | 0.006 | 0.000 | 0.000 |
| Proteobacteria | Gammaproteobacteria | Unclassified | Unclassified | Methylohalomonas | 0.000 | 0.000 | 0.004 | 0.000 | 0.000 | 0.000 |
| Proteobacteria | Gammaproteobacteria | Vibrionales | Vibrionaceae | Alishewanella | 0.016 | 0.000 | 0.010 | 0.000 | 0.000 | 0.000 |
| Proteobacteria | Gammaproteobacteria | Vibrionales | Vibrionaceae | Vibrio | 0.008 | 0.006 | 0.000 | 0.000 | 0.000 | 0.006 |
| Proteobacteria | Gammaproteobacteria_1 | Vibrionales | Vibrionaceae | Enterovibrio | 0.000 | 0.000 | 0.014 | 0.000 | 0.000 | 0.000 |
| Proteobacteria | Gammaproteobacteria_1 | Vibrionales | Vibrionaceae | Grimontia | 0.000 | 0.000 | 0.006 | 0.000 | 0.000 | 0.000 |
| Proteobacteria | Gammaproteobacteria_1 | Vibrionales | Vibrionaceae | Photobacterium | 0.000 | 0.000 | 0.071 | 0.000 | 0.000 | 0.000 |
| Proteobacteria | Gammaproteobacteria_1 | Vibrionales | Vibrionaceae | Phycicoccus | 0.008 | 0.000 | 0.000 | 0.000 | 0.005 | 0.000 |
| Proteobacteria | Gammaproteobacteria_1 | Vibrionales | Vibrionaceae | Salinivibrio | 0.000 | 0.000 | 0.010 | 0.000 | 0.000 | 0.000 |
| Proteobacteria | Deltaproteobacteria | Myxococcales | Vulgatibacteraceae | Weeksella | 0.000 | 0.000 | 0.008 | 0.000 | 0.000 | 0.000 |
| Proteobacteria | Alphaproteobacteria | Rhizobiales | Xanthobacteraceae | Xenophilus | 0.008 | 0.000 | 0.006 | 0.000 | 0.000 | 0.000 |
| Proteobacteria | Alphaproteobacteria | Rhizobiales_2 | Xanthobacteraceae | Pseudomonas | 0.724 | 1.639 | 0.312 | 1.611 | 2.876 | 1.913 |
| Proteobacteria | Alphaproteobacteria | Rhizobiales_3 | Xanthobacteraceae | Ancylobacter | 0.000 | 0.000 | 0.016 | 0.000 | 0.000 | 0.000 |
| Proteobacteria | Gammaproteobacteria | Xanthomonadales | Xanthomonadaceae | Arenimonas | 0.016 | 0.000 | 0.000 | 0.000 | 0.000 | 0.000 |
| Proteobacteria | Gammaproteobacteria | Xanthomonadales | Xanthomonadaceae | Arenimonas_1 | 0.016 | 0.000 | 0.026 | 0.000 | 0.000 | 0.000 |
| Proteobacteria | Gammaproteobacteria | Xanthomonadales | Xanthomonadaceae | Arenimonas_2 | 0.000 | 0.000 | 0.020 | 0.000 | 0.000 | 0.000 |
| Proteobacteria | Gammaproteobacteria | Xanthomonadales | Xanthomonadaceae | Aspromonas | 0.000 | 0.000 | 0.004 | 0.000 | 0.000 | 0.000 |
| Proteobacteria | Gammaproteobacteria | Xanthomonadales | Xanthomonadaceae | Dokdonella | 0.016 | 0.006 | 0.016 | 0.012 | 0.000 | 0.000 |
| Proteobacteria | Gammaproteobacteria | Xanthomonadales | Xanthomonadaceae | Dyella | 0.008 | 0.000 | 0.004 | 0.054 | 0.120 | 0.079 |
| Proteobacteria | Gammaproteobacteria | Xanthomonadales | Xanthomonadaceae | Dyella_2 | 0.000 | 0.000 | 0.008 | 0.012 | 0.025 | 0.018 |
| Proteobacteria | Gammaproteobacteria | Xanthomonadales | Xanthomonadaceae | Leifsonia_a | 0.000 | 0.000 | 0.004 | 0.000 | 0.000 | 0.000 |
| Proteobacteria | Gammaproteobacteria | Xanthomonadales | Xanthomonadaceae | Luteimonas | 0.008 | 0.031 | 0.000 | 0.000 | 0.000 | 0.000 |
| Proteobacteria | Gammaproteobacteria | Xanthomonadales | Xanthomonadaceae | Luteimonas_1 | 0.000 | 0.000 | 0.004 | 0.000 | 0.000 | 0.000 |
| Proteobacteria | Gammaproteobacteria | Xanthomonadales | Xanthomonadaceae | Luteimonas_3 | 0.008 | 0.000 | 0.010 | 0.000 | 0.000 | 0.000 |
| Proteobacteria | Gammaproteobacteria | Xanthomonadales | Xanthomonadaceae | Luteococcus | 0.000 | 0.000 | 0.008 | 0.000 | 0.000 | 0.000 |
| Proteobacteria | Gammaproteobacteria | Xanthomonadales | Xanthomonadaceae | Lysobacter_1 | 0.000 | 0.000 | 0.012 | 0.000 | 0.000 | 0.000 |

**Table S9 continued.**

| | | | | | | | | | | |
|---|---|---|---|---|---|---|---|---|---|---|
| Proteobacteria | Gammaproteobacteria | Xanthomonadales | Xanthomonadaceae | Lysobacter_2 | 0.024 | 0.000 | 0.040 | 0.006 | 0.000 | 0.000 |
| Proteobacteria | Gammaproteobacteria | Xanthomonadales | Xanthomonadaceae | Lysobacter_3 | 0.000 | 0.000 | 0.014 | 0.000 | 0.000 | 0.000 |
| Proteobacteria | Gammaproteobacteria | Xanthomonadales | Xanthomonadaceae | M2PB4-61_termite_group | 1.489 | 0.308 | 0.038 | 0.413 | 0.025 | 0.042 |
| Proteobacteria | Gammaproteobacteria | Xanthomonadales | Xanthomonadaceae | Psychrobacter | 0.008 | 0.000 | 0.193 | 0.000 | 0.000 | 0.000 |
| Proteobacteria | Gammaproteobacteria | Xanthomonadales | Xanthomonadaceae | Rhodanobacter | 0.000 | 0.000 | 0.002 | 0.000 | 0.010 | 0.000 |
| Proteobacteria | Gammaproteobacteria | Xanthomonadales | Xanthomonadaceae | Rhodobacter | 0.000 | 0.012 | 0.000 | 0.000 | 0.000 | 0.000 |
| Proteobacteria | Gammaproteobacteria | Xanthomonadales | Xanthomonadaceae | Steroidobacter | 0.130 | 0.018 | 0.018 | 0.012 | 0.000 | 0.000 |
| Proteobacteria | Gammaproteobacteria | Xanthomonadales | Xanthomonadaceae | Thermomonas_2 | 0.000 | 0.000 | 0.030 | 0.000 | 0.000 | 0.000 |
| Proteobacteria | Gammaproteobacteria | Xanthomonadales | Xanthomonadaceae | Thermomonospora | 0.000 | 0.000 | 0.008 | 0.000 | 0.000 | 0.000 |
| Proteobacteria | Gammaproteobacteria | Xanthomonadales | Xanthomonadaceae | Uncultured_f | 0.000 | 0.000 | 0.004 | 0.000 | 0.000 | 0.000 |
| Proteobacteria | Gammaproteobacteria | Xanthomonadales | Xanthomonadaceae | Wohlfahrtiimonas | 0.000 | 0.000 | 0.004 | 0.000 | 0.000 | 0.000 |
| Proteobacteria | Gammaproteobacteria | Xanthomonadales | Xanthomonadaceae | Xylella | 0.000 | 0.000 | 0.012 | 0.000 | 0.000 | 0.000 |
| Proteobacteria | Gammaproteobacteria | Enterobacterales | Yersiniaceae | Chania | 0.000 | 0.018 | 0.006 | 0.018 | 0.020 | 0.012 |
| Proteobacteria | Gammaproteobacteria | Enterobacteriales | Yersiniaceae | Ewingella | 0.000 | 0.000 | 0.000 | 0.000 | 0.000 | 0.006 |
| Proteobacteria | Gammaproteobacteria | Enterobacteriales | Yersiniaceae | Gibbsiella | 0.000 | 0.000 | 0.000 | 0.000 | 0.015 | 0.000 |
| Proteobacteria | Betaproteobacteria | Rhodocyclales | Zoogloeaceae | Thermanaeromonas | 0.000 | 0.000 | 0.008 | 0.000 | 0.000 | 0.000 |
| Proteobacteria | Betaproteobacteria | Rhodocyclales | Zoogloeaceae | Uncultured | 0.390 | 0.025 | 0.951 | 0.006 | 0.005 | 0.000 |
| Proteobacteria_1 | Deltaproteobacteria | Desulfobacterales | Nitrospinaceae | Candidatus_Entotheonella | 0.008 | 0.000 | 0.127 | 0.000 | 0.000 | 0.000 |
| Spirochaetes | Spirochaetes | Spirochaetales | Brachyspiraceae | Brachyspira | 0.000 | 0.000 | 0.016 | 0.000 | 0.000 | 0.000 |
| Spirochaetes | Spirochaetes | Candidatus_Cloacamonas | Candidatus_Cloacamonas | Candidatus_Cloacamonas | 0.000 | 0.000 | 0.083 | 0.000 | 0.000 | 0.000 |
| Spirochaetes | Spirochaetes | Spirochaetales | Leptospiraceae | Turneriella | 0.000 | 0.000 | 0.010 | 0.000 | 0.000 | 0.000 |
| Spirochaetes | Spirochaetia | Spirochaetales | Leptospiraceae | Rubellimicrobium | 0.000 | 0.000 | 0.016 | 0.000 | 0.000 | 0.000 |
| Spirochaetes | Spirochaetia | Spirochaetales | Spirochaetaceae | Spiroplasma_2 | 0.000 | 0.000 | 0.012 | 0.000 | 0.000 | 0.000 |
| Spirochaetes | Spirochaetia | Spirochaetales | Spirochaetaceae | Treponema_Ia | 4.254 | 1.737 | 0.631 | 0.665 | 0.105 | 0.133 |
| Spirochaetes | Spirochaetia | Spirochaetales | Spirochaetaceae_10 | Treponema_Ib | 0.057 | 0.000 | 0.091 | 0.006 | 0.000 | 0.000 |
| Spirochaetes | Spirochaetia | Spirochaetales | Spirochaetaceae_11 | Digestor_cluster | 0.000 | 0.000 | 0.145 | 0.000 | 0.000 | 0.000 |
| Spirochaetes | Spirochaetes | Spirochaetales | Spirochaetaceae_2 | Borrelia | 0.000 | 0.000 | 0.030 | 0.000 | 0.000 | 0.000 |
| Spirochaetes | Spirochaetes | Spirochaetales | Spirochaetaceae_5 | Macrococcus | 0.000 | 0.000 | 0.000 | 0.000 | 0.000 | 0.006 |
| Spirochaetes | Spirochaetes | Spirochaetales | Spirochaetaceae_9 | Brevinema | 0.000 | 0.000 | 0.012 | 0.000 | 0.000 | 0.000 |
| Spirochaetes | Spirochaetes | Spirochaetales | Spirochaetaceae_Treponema | Animal_cluster_1 | 0.000 | 0.000 | 0.099 | 0.000 | 0.000 | 0.000 |
| Spirochaetes | Spirochaetes | Spirochaetales | Spirochaetaceae_Treponema | Animal_cluster_2 | 0.000 | 0.000 | 0.062 | 0.006 | 0.000 | 0.000 |
| Spirochaetes | Spirochaetia | Spirochaetales | Spirochaetaceae_Treponema | Treponema_Ic | 0.155 | 0.074 | 0.276 | 0.018 | 0.005 | 0.000 |
| Spirochaetes | Spirochaetia | Spirochaetales | Spirochaetaceae_Treponema | Treponema_Id | 0.000 | 0.000 | 0.032 | 0.000 | 0.000 | 0.000 |
| Spirochaetes | Spirochaetia | Spirochaetales | Spirochaetaceae_Treponema | Treponema_Ie | 0.748 | 0.308 | 0.012 | 0.114 | 0.025 | 0.012 |
| Spirochaetes | Spirochaetia | Spirochaetales | Spirochaetaceae_Treponema | Treponema_If | 1.066 | 0.160 | 0.183 | 0.102 | 0.010 | 0.024 |
| Spirochaetes | Spirochaetia | Spirochaetales | Spirochaetaceae_Treponema | Treponema_Ig | 0.033 | 0.000 | 0.095 | 0.012 | 0.000 | 0.000 |
| Spirochaetes | Spirochaetia | Spirochaetales | Spirochaetaceae_Treponema | Treponema_Ih | 0.415 | 0.129 | 0.014 | 0.042 | 0.005 | 0.000 |
| Spirochaetes | Spirochaetia | Spirochaetales | Spirochaetaceae_Treponema | Treponema_II | 0.000 | 0.000 | 0.095 | 0.000 | 0.005 | 0.000 |
| Spirochaetes | Spirochaetia | Spirochaetales | Spirochaetaceae_Treponema | Treponema_III | 0.146 | 0.086 | 0.008 | 0.006 | 0.000 | 0.000 |
| Spirochaetes | Spirochaetia | Spirochaetales | Spirochaetaceae_Treponema | Trichococcus_1 | 0.000 | 0.000 | 0.010 | 0.000 | 0.000 | 0.000 |

**Table S9 continued.**

| | | | | | | | | | | |
|---|---|---|---|---|---|---|---|---|---|---|
| Spirochaetes | Spirochaetales | Spirochaetaceae | Treponema_2 | Magnetospira | 0.008 | 0.000 | 0.000 | 0.000 | 0.000 | 0.000 |
| Synergistetes | Synergistia | Synergistales | Synergistaceae | Aminiphilus | 0.000 | 0.000 | 0.014 | 0.000 | 0.000 | 0.000 |
| Synergistetes | Synergistia | Synergistales | Synergistaceae | Aminobacterium | 0.000 | 0.000 | 0.028 | 0.000 | 0.000 | 0.000 |
| Synergistetes | Synergistia | Synergistales | Synergistaceae | Anaerobaculum | 0.000 | 0.000 | 0.042 | 0.000 | 0.000 | 0.000 |
| Synergistetes | Synergistia | Synergistales | Synergistaceae | Candidatus_Tammella | 0.146 | 0.031 | 0.050 | 0.000 | 0.000 | 0.000 |
| Synergistetes | Synergistia | Synergistales | Synergistaceae | Cloacibacillus | 0.000 | 0.000 | 0.032 | 0.000 | 0.000 | 0.000 |
| Synergistetes | Synergistia | Synergistales | Synergistaceae | Dethiosulfovibrio | 0.000 | 0.000 | 0.008 | 0.000 | 0.000 | 0.000 |
| Synergistetes | Synergistia | Synergistales | Synergistaceae | Digester_cluster_1 | 0.000 | 0.000 | 0.024 | 0.000 | 0.000 | 0.000 |
| Synergistetes | Synergistia | Synergistales | Synergistaceae | Digester_cluster_2 | 0.000 | 0.000 | 0.052 | 0.000 | 0.000 | 0.000 |
| Synergistetes | Synergistia | Synergistales | Synergistaceae | Digester_cluster_3 | 0.008 | 0.000 | 0.258 | 0.000 | 0.000 | 0.000 |
| Synergistetes | Synergistia | Synergistales | Synergistaceae | Digester_cluster_4 | 0.000 | 0.000 | 0.012 | 0.000 | 0.000 | 0.000 |
| Synergistetes | Synergistia | Synergistales | Synergistaceae | Digester_cluster_5 | 0.000 | 0.000 | 0.141 | 0.000 | 0.000 | 0.000 |
| Synergistetes | Synergistia | Synergistales | Synergistaceae | Haematospirillum | 0.008 | 0.000 | 0.000 | 0.006 | 0.000 | 0.000 |
| Synergistetes | Synergistia | Synergistales | Synergistaceae | Kaistia | 0.000 | 0.000 | 0.014 | 0.000 | 0.000 | 0.000 |
| Synergistetes | Synergistia | Synergistales | Synergistaceae | Lactobacillus | 0.000 | 0.000 | 0.000 | 0.006 | 0.000 | 0.000 |
| Synergistetes | Synergistia | Synergistales | Synergistaceae | Oribacterium | 0.000 | 0.000 | 0.054 | 0.000 | 0.000 | 0.000 |
| Synergistetes | Synergistia | Synergistales | Synergistaceae | Pyramidobacter | 0.000 | 0.000 | 0.016 | 0.000 | 0.000 | 0.000 |
| Synergistetes | Synergistia | Synergistales | Synergistaceae | Syntrophococcus | 0.000 | 0.000 | 0.018 | 0.000 | 0.000 | 0.000 |
| Synergistetes | Synergistia | Synergistales | Synergistaceae | Termite_cockroach_cluster_2 | 0.041 | 0.062 | 0.107 | 0.006 | 0.000 | 0.000 |
| Synergistetes | Synergistia | Synergistales | Synergistaceae | Thermincola | 0.008 | 0.000 | 0.012 | 0.000 | 0.000 | 0.000 |
| Synergistetes | Synergistia | Synergistales | Synergistaceae | Thioalkalivibrio | 0.000 | 0.000 | 0.000 | 0.006 | 0.000 | 0.000 |
| Tenericutes | Mollicutes | Acholeplasmatales | Acholeplasmataceae | Acholeplasma_1 | 0.000 | 0.000 | 0.016 | 0.000 | 0.000 | 0.000 |
| Tenericutes | Mollicutes | Acholeplasmatales | Acholeplasmataceae | Acholeplasma_3 | 0.000 | 0.000 | 0.010 | 0.000 | 0.000 | 0.000 |
| Tenericutes | Mollicutes | Candidatus_Phytoplasma | Candidatus_Phytoplasma | Candidatus_Phytoplasma | 0.000 | 0.000 | 0.046 | 0.000 | 0.000 | 0.000 |
| Tenericutes | Mollicutes | Entomoplasmatales | Entomoplasmataceae | Entomoplasma | 0.000 | 0.000 | 0.010 | 0.000 | 0.000 | 0.000 |
| Tenericutes | Mollicutes | Entomoplasmatales | Entomoplasmataceae | Mesoplasma_1 | 0.000 | 0.000 | 0.006 | 0.000 | 0.000 | 0.000 |
| Tenericutes | Mollicutes | Entomoplasmatales | Entomoplasmataceae | Mesoplasma_2 | 0.000 | 0.000 | 0.004 | 0.000 | 0.000 | 0.000 |
| Tenericutes | Mollicutes | Mycoplasmatales | Mycoplasmataceae | Candidatus_Bacilloplasma | 0.000 | 0.000 | 0.008 | 0.000 | 0.000 | 0.000 |
| Tenericutes | Mollicutes | Mycoplasmatales | Mycoplasmataceae | Candidatus_Lumbricincola | 0.000 | 0.000 | 0.004 | 0.000 | 0.000 | 0.000 |
| Tenericutes | Mollicutes | Mycoplasmatales | Mycoplasmataceae | Mycoplasma_1 | 0.000 | 0.000 | 0.163 | 0.000 | 0.000 | 0.000 |
| Tenericutes | Mollicutes | Mycoplasmatales | Mycoplasmataceae | Mycoplasma_2 | 0.114 | 0.037 | 0.030 | 0.018 | 0.015 | 0.042 |
| Tenericutes | Mollicutes | Mycoplasmatales | Mycoplasmataceae | Mycoplasma_3 | 0.000 | 0.000 | 0.008 | 0.000 | 0.000 | 0.000 |
| Tenericutes | Mollicutes | Mycoplasmatales | Mycoplasmataceae | Mycoplasma_4 | 0.000 | 0.000 | 0.012 | 0.000 | 0.000 | 0.000 |
| Tenericutes | Mollicutes | Mycoplasmatales | Mycoplasmataceae | Myroides | 0.000 | 0.000 | 0.012 | 0.000 | 0.000 | 0.000 |
| Tenericutes | Mollicutes | Mycoplasmatales | Mycoplasmataceae | Ureaplasma | 0.000 | 0.000 | 0.028 | 0.000 | 0.000 | 0.000 |
| Tenericutes | Mollicutes | Entomoplasmatales | Spiroplasmataceae | Spiroplasma | 0.008 | 0.000 | 0.002 | 0.000 | 0.000 | 0.000 |
| Tenericutes | Mollicutes | Entomoplasmatales | Spiroplasmataceae | Spiroplasma_3 | 0.000 | 0.000 | 0.010 | 0.000 | 0.000 | 0.012 |
| Thermodesulfobacteria | Thermodesulfobacteria | Thermodesulfobacteriales | Thermodesulfobacteriaceae | Caldimicrobium | 0.000 | 0.000 | 0.038 | 0.000 | 0.000 | 0.000 |
| Thermodesulfobacteria | Thermodesulfobacteria | Thermodesulfobacteriales | Thermodesulfobacteriaceae | Thermodesulfobacterium | 0.000 | 0.000 | 0.006 | 0.000 | 0.000 | 0.000 |
| Thermotogae | Thermotogae | Thermotogales | Thermotogaceae | Fervidobacterium | 0.000 | 0.000 | 0.040 | 0.000 | 0.000 | 0.000 |

**Table S9 continued.**

| | | | | | | | | | | |
|---|---|---|---|---|---|---|---|---|---|---|
| Thermotogae | Thermotogae | Thermotogales | Thermotogaceae | Geotoga | 0.000 | 0.000 | 0.010 | 0.000 | 0.000 | 0.000 |
| Thermotogae | Thermotogae | Thermotogales | Thermotogaceae | Kosmotoga | 0.000 | 0.000 | 0.048 | 0.000 | 0.000 | 0.000 |
| Thermotogae | Thermotogae | Thermotogales | Thermotogaceae | Marinitoga | 0.000 | 0.000 | 0.012 | 0.000 | 0.000 | 0.000 |
| Thermotogae | Thermotogae | Thermotogales | Thermotogaceae | Petrotoga | 0.000 | 0.000 | 0.012 | 0.000 | 0.000 | 0.000 |
| Thermotogae | Thermotogae | Thermotogales | Thermotogaceae | Thermosipho | 0.000 | 0.000 | 0.020 | 0.000 | 0.000 | 0.000 |
| Thermotogae | Thermotogae | Thermotogales | Thermotogaceae | Thermotoga_1 | 0.000 | 0.000 | 0.018 | 0.000 | 0.000 | 0.000 |
| Thermotogae | Thermotogae | Thermotogales | Thermotogaceae | Thermotoga_2 | 0.000 | 0.000 | 0.006 | 0.000 | 0.000 | 0.000 |
| Unclassified | Unclassified | Haloplasmatales | Haloplasmataceae | Haloplasma | 0.000 | 0.000 | 0.006 | 0.000 | 0.000 | 0.000 |
| Unclassified | Unclassified | Haloplasmatales | Haloplasmataceae | Halorhodospira | 0.000 | 0.000 | 0.006 | 0.000 | 0.000 | 0.000 |
| unclassified | unclassified | unclassified | unclassified | Uncultured_1 | 1.456 | 0.271 | 4.023 | 0.114 | 0.040 | 0.042 |
| unclassified | unclassified | unclassified | unclassified | Nitrosospira | 0.000 | 0.000 | 0.010 | 0.000 | 0.000 | 0.000 |
| unclassified | unclassified | unclassified | unclassified | Pigmentiphaga | 0.138 | 0.025 | 0.018 | 0.090 | 0.174 | 0.121 |
| Verrucomicrobia | Opitutae | Opitutales | Opitutaceae | Alterococcus | 0.000 | 0.006 | 0.012 | 0.000 | 0.000 | 0.000 |
| Verrucomicrobia | Opitutae | Opitutales | Opitutaceae | Lampropedia | 0.000 | 0.000 | 0.004 | 0.000 | 0.000 | 0.000 |
| Verrucomicrobia | Opitutae | Opitutales | Opitutaceae | Oral_cluster | 0.000 | 0.000 | 0.022 | 0.000 | 0.000 | 0.000 |
| Verrucomicrobia | Opitutae | Puniceicoccales | Puniceicoccaceae | Cerasicoccus | 0.000 | 0.000 | 0.016 | 0.000 | 0.000 | 0.000 |
| Verrucomicrobia | Opitutae | Puniceicoccales | Puniceicoccaceae | Coraliomargarita | 0.000 | 0.000 | 0.012 | 0.000 | 0.000 | 0.000 |
| Verrucomicrobia | Opitutae | Puniceicoccales | Puniceicoccaceae | Lentimonas | 0.000 | 0.000 | 0.006 | 0.000 | 0.000 | 0.000 |
| Verrucomicrobia | Opitutae | Puniceicoccales | Puniceicoccaceae | Pelagicoccus | 0.000 | 0.000 | 0.014 | 0.000 | 0.000 | 0.000 |
| Verrucomicrobia | Opitutae | Puniceicoccales | Puniceicoccaceae | Puniceicoccus | 0.000 | 0.000 | 0.016 | 0.000 | 0.000 | 0.000 |
| Verrucomicrobia | Verrucomicrobiae | Verrucomicrobiales | Rubritaleaceae | Rubritalea | 0.000 | 0.000 | 0.028 | 0.000 | 0.000 | 0.000 |
| Verrucomicrobia | Verrucomicrobiae | Verrucomicrobiales | Verrucomicrobiaceae | Akkermansia | 0.008 | 0.000 | 0.262 | 0.006 | 0.010 | 0.000 |
| Verrucomicrobia | Verrucomicrobiae | Verrucomicrobiales | Verrucomicrobiaceae | AKYG587 | 0.000 | 0.000 | 0.026 | 0.000 | 0.000 | 0.000 |
| Verrucomicrobia | Verrucomicrobiae | Verrucomicrobiales | Verrucomicrobiaceae | Persicirhabdus | 0.000 | 0.000 | 0.018 | 0.000 | 0.000 | 0.000 |
| Verrucomicrobia | Verrucomicrobiae | Verrucomicrobiales | Verrucomicrobiaceae | Roseibacillus | 0.000 | 0.000 | 0.034 | 0.000 | 0.000 | 0.000 |
| Verrucomicrobia | Verrucomicrobiae | Verrucomicrobiales | Verrucomicrobiaceae_1 | Halomonas_1 | 0.000 | 0.000 | 0.195 | 0.006 | 0.000 | 0.006 |
| Verrucomicrobia | Verrucomicrobiae | Verrucomicrobiales | Verrucomicrobiaceae_1 | Luteolibacter_2 | 0.000 | 0.000 | 0.006 | 0.000 | 0.000 | 0.000 |
| Verrucomicrobia | Verrucomicrobiae | Verrucomicrobiales | Verrucomicrobiaceae_2 | Prosthecomicrobium | 0.000 | 0.000 | 0.016 | 0.000 | 0.000 | 0.000 |
| Verrucomicrobia | Verrucomicrobiae | Verrucomicrobiales | Verrucomicrobiaceae_2 | Vibrio-Alivibrio | 0.008 | 0.000 | 0.439 | 0.012 | 0.000 | 0.006 |
| Verrucomicrobia | Spartobacteria | Chthoniobacterales | Xiphinematobacteraceae | Candidatus_Xiphinematobacter | 0.008 | 0.000 | 0.054 | 0.000 | 0.000 | 0.000 |
| Unidentified | Unidentified | Unidentified | Unidentified | Unidentified | 0.081 | 0.006 | 0.004 | 0.012 | 0.000 | 0.000 |

**Table S9 continued.**
